# Spatial Single-cell Analysis Decodes Cortical Layer and Area Specification

**DOI:** 10.1101/2024.06.05.597673

**Authors:** Xuyu Qian, Kyle Coleman, Shunzhou Jiang, Andrea J. Kriz, Jack H. Marciano, Chunyu Luo, Chunhui Cai, Monica Devi Manam, Emre Caglayan, Aoi Otani, Urmi Ghosh, Diane D. Shao, Rebecca E. Andersen, Jennifer E. Neil, Robert Johnson, Alexandra LeFevre, Jonathan L. Hecht, Michael B. Miller, Liang Sun, Carsen Stringer, Mingyao Li, Christopher A. Walsh

## Abstract

The human cerebral cortex, pivotal for advanced cognitive functions, is composed of six distinct layers and dozens of functionally specialized areas^1,2^. The layers and areas are distinguished both molecularly, by diverse neuronal and glial cell subtypes, and structurally, through intricate spatial organization^3,4^. While single-cell transcriptomics studies have advanced molecular characterization of human cortical development, a critical gap exists due to the loss of spatial context during cell dissociation^5,6,7,8^. Here, we utilized multiplexed error-robust fluorescence in situ hybridization (MERFISH)^9^, augmented with deep-learning-based cell segmentation, to examine the molecular, cellular, and cytoarchitectural development of human fetal cortex with spatially resolved single-cell resolution. Our extensive spatial atlas, encompassing 16 million single cells, spans eight cortical areas across four time points in the second and third trimesters. We uncovered an early establishment of the six-layer structure, identifiable in the laminar distribution of excitatory neuronal subtypes by mid-gestation, long before the emergence of cytoarchitectural layers. Notably, while anterior-posterior gradients of neuronal subtypes were generally observed in most cortical areas, a striking exception was the sharp molecular border between primary (V1) and secondary visual cortices (V2) at gestational week 20. Here we discovered an abrupt binary shift in neuronal subtype specification at the earliest stages, challenging the notion that continuous morphogen gradients dictate mid-gestation cortical arealization^6,10^. Moreover, integrating single-nuclei RNA-sequencing and in situ whole transcriptomics revealed an early upregulation of synaptogenesis in V1-specific Layer 4 neurons, suggesting a role of synaptogenesis in this discrete border formation. Collectively, our findings underscore the crucial role of spatial relationships in determining the molecular specification of cortical layers and areas. This work not only provides a valuable resource for the field, but also establishes a spatially resolved single-cell analysis paradigm that paves the way for a comprehensive developmental atlas of the human brain.

## Main

Abnormal development of the cerebral cortex is linked to a wide array of neurological disorders, including autism spectrum disorder (ASD), epilepsy, intellectual disability, and various neuropsychiatric conditions^1^. The human neocortex contains a myriad of neuronal and glial cell types, a diversity that emerges from the interplay of both intrinsic and extrinsic factors during development^3,4,5,11,12^. Different areas of the cortex, such as the frontal lobe and occipital lobe, display significant variations in neuronal subtype repertoire and cytoarchitecture of the layers, providing the cellular and structural basis for area-specific circuitry^3,5^. During early embryonic stages, polarized secreted factors, called morphogens, induce neural progenitor cells to acquire preliminary areal identities, creating the initial blueprint of cortical areas, often referred to as the “protomap”^13,14^. As development progresses, thalamic afferents innervating the cortex refine area-specific neuronal subtype identities through synaptic inputs and localized signaling factors, transforming the relatively homogeneous cortical sheet, or “protocortex”, into highly distinct cortical areas^16,17^.

Achieving a comprehensive understanding of cortical development necessitates simultaneous analysis of molecular and spatial organization in intact tissues. While in-situ whole-transcriptome sequencing methods detect mRNA within a spatial grid, they cannot guarantee single-cell resolution as each capture spot is typically occupied by multiple cells^18,19^. In contrast, MERFISH provides an imaging-based solution with sub-micron spatial resolution for precise transcript detection within a pre-designed gene panel^9,20,21^. In this study, we employed MERFISH to analyze human fetal cortex samples across major cortical areas, focusing on the molecular and cellular specification of cortical layers and areas during the second and third trimesters (**Fig. 1a**). We enriched this analysis with single-nucleus RNA-sequencing (snRNAseq) and in situ whole-transcriptomics (Visium) on consecutive tissue sections for a subset of samples. The snRNAseq-augmented MERFISH analysis then enabled accurate imputation of gene expression beyond the MERFISH gene panel through environmental variational inference (ENVI)^22^, and imputed results were validated by ground truth measurement from Visium (**Fig. 1a**). Collectively, our integrated approach enables spatially resolved single-cell analysis of the developing human cortex on an unprecedented scale, revealing critical insights into the development of cortical layers and areas that were previously elusive with traditional methodologies.

**Fig. 1:**
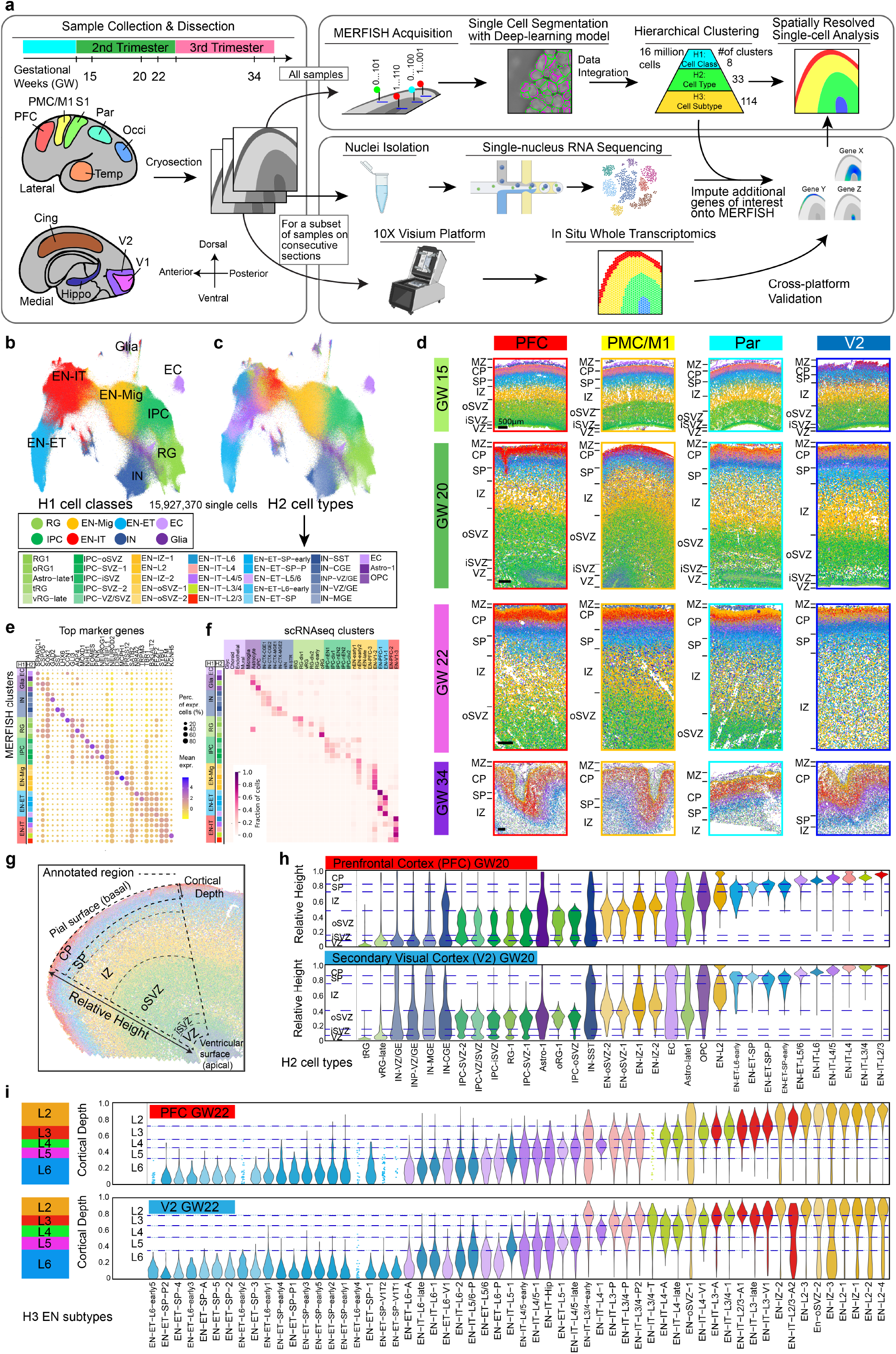
A spatially resolved single-cell atlas of human fetal cortical development. **a**, Schematics of sampling and workflow. PFC; prefrontal cortex; PMC, premotor cortex; M1, primary motor cortex; S1, primary somatosensory cortex; Par, parietal cortex; Occi, occipital cortex; Temp, temporal cortex; Cing, cingulate cortex; Hippo, hippocampus; V1, primary visual cortex; V2, secondary visual cortex. **b, c**, Uniform Manifold Approximation and Projection (UMAP) of single cells analyzed by MERFISH, colored by H1 cell classes (**b**), and by H2 cell types (**c**). RG, radial glia; IPC, intermediate progenitor cells; EN-Mig, migrating excitatory neurons; EN-IT, intratelencephalic excitatory neurons; EN-ET, extratelencephalic excitatory neurons; IN, inhibitory neurons; EC, endothelial cells; oRG, outer radial glia; tRC, truncated radial glia; vRG, ventricular radial glia; INP, inhibitory neuron progenitors; CGE, caudal ganglionic eminence; MGE, medial ganglionic eminence; OPC, oligodendrocyte precursor cells. **d**, Spatial maps of H2 cell types from major cortical areas across gestational week (GW) 15 to 34. MZ, marginal zone; CP, cortical plate; SP, subplate; IZ, intermediate zone; oSVZ, outer subventricular zone; iSVZ, inner subventricular zone; VZ, ventricular zone. Scale bars, 500 µm. **e**, Dot plot showing expression of marker genes for H2 cell types. **f**, Cell type correspondence heatmap shows the fraction of cells from MERFISH H2 clusters that associate to clusters from published mid-gestation scRNA-seq dataset^7^. **g**, Schematics illustrating the annotated fan-shaped cortical area used for relative height (RH) and cortical depth (CD) calculation. **h**, Violin plots showing the laminar distribution of H2 cell types from the apical to basal surface quantified by the RH. The width of the violin for each cell type is normalized to the maximum value. **i**, Violin plots showing CD distribution of H3 EN subtypes within the CP. Dash lines represent the borders between cortical layers (L2-L6) calculated based on the CD distribution of layer-defining clusters. Clusters with fewer than 50 cells within the annotated region are represented by dots for individual cells instead of a violin.

### A spatial atlas of human fetal cortex

We analyzed fetal human tissues from six individuals across gestational weeks (GW) 15, 20, 22, and 34, covering eight major cortical areas along the anterior-posterior (A-P) axis (**Fig. 1a, Supplementary Table 1**). We curated a panel of three hundred genes for MERFISH analysis. This panel included canonical marker genes for major cell types, alongside genes selected for their cluster-specific enrichment in a published single-cell RNA sequencing (scRNAseq) dataset of the mid-gestation human fetal cortex (**Methods, Supplementary Table 2, 3**)^7^.

The extraordinarily high cell density in the mid-gestation human fetal cortex presented a unique challenge for achieving precise single-cell resolution. To address this, we developed a custom deep-learning model based on the CellPose 2.0 framework^23,24^, which performs automated single-cell segmentation using nucleus staining images co-captured during MERFISH imaging. After iterative human-in-the-loop training on our images, the model achieved robust agreement with manual labeling (**Extended Data Fig. 1a-c**). The distribution of cell volume remained consistent across samples, experiments, and clusters (**Extended Data Fig. 1d-f**). Following barcode decoding, detected transcripts were accurately assigned to segmented single cells, and gene expression data from technical and biological replicates proved to be highly reproducible, despite some variability in total RNA abundance among different samples, which likely reflects some differences in sample condition (**Extended Data Fig. 1g, h**).

In total, we analyzed approximately 16 million single cells that met quality control criteria, and we integrated all experiments to cluster the cells based on their gene expression following a hierarchical strategy (**Fig. 1a**). At the first hierarchy, referred to as H1, eight cell classes were identified, including radial glia (RG), intermediate progenitor cells (IPC), migrating excitatory neurons (EN-Mig), intratelencephalic excitatory neurons (EN-IT), extratelencephalic excitatory neurons (EN-ET), inhibitory neurons (IN), other glia, and endothelial cells (ECs) (**Fig. 1b**). We broadly define all non-IT excitatory projection neurons, including the corticothalamic (CT) and pyramidal tract (PT) neurons as EN-ETs^25^. At the second hierarchy (H2), these eight H1 cell classes were divided into 33 cell types, which were then subdivided into 114 subtypes at the third hierarchy (H3), with 58 of these subtypes being EN subtypes (**Fig. 1b, Supplementary Table 4**). Our sampling strategy ensured robust representations of different cortical areas and gestational ages, with cell type proportions remaining consistent across samples from the same gestational age (**Extended Data Fig. 1i-k**).

Clusters across all three hierarchical levels revealed distinct spatial distribution patterns. The localization of H2 cell types delineated the apical-basal laminar structures, including the ventricular zone (VZ), inner and outer subventricular zones (iSVZ and oSVZ), intermediate zone (IZ), subplate (SP), cortical plate (CP), and marginal zone (MZ) (**Fig. 1d, Extended Data Fig 1l**). Clusters exhibited dynamic marker expression, and cell label transfer analysis showed close alignment to the fetal human cortex scRNAseq datasets (**Fig. 1e, f, Extended Data Fig. 2a, b**)^7^. For quantitative analysis of cell localization, we manually annotated each sample based on cytoarchitecture, creating a framework within a fan-shaped region that captured a geometrically uniform cortical area (**Fig. 1g**). Referencing the Allen Reference Atlas for GW15 and GW21^26^, we divided this region into major laminar structures based on the localization of H2 cell types (**Methods**). Within this framework, we assigned each cell to a structure and calculated its relative height (RH) from the apical to basal surfaces. The RH distributions for H2 cell types were consistent across experiments and cortical areas, accurately reflecting laminar structures (**Fig. 1h, Extended Data Fig. 2c**).

**Fig. 2:**
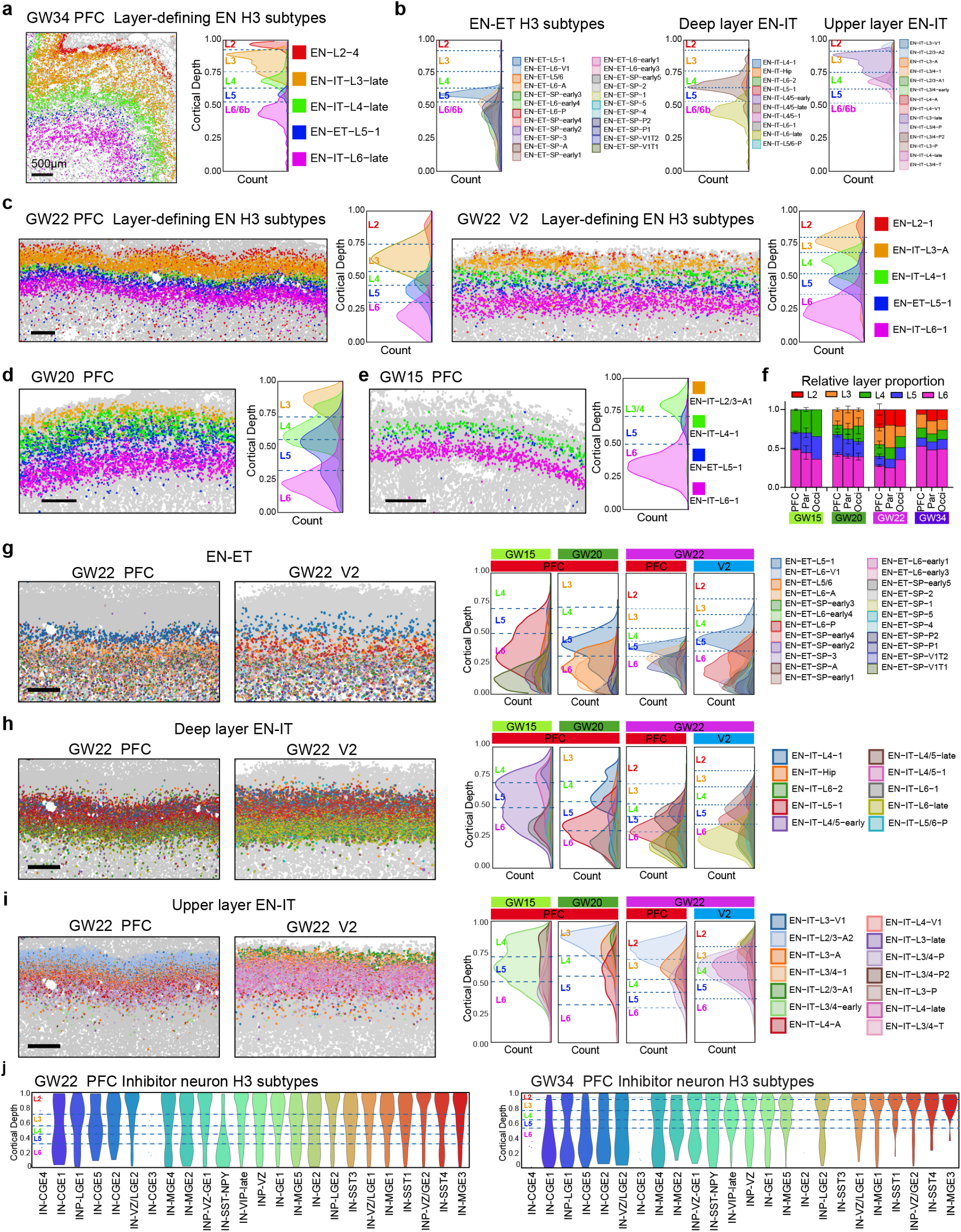
Progressive formation and specification of cortical layers. **a**, Spatial map (left) shows the cortical layers 2 to 6 are defined by the organization of selected EN subtypes in the prefrontal cortex (PFC) at GW34; and ridgeline plot (right) shows the cortical depth (CD) distribution of layer-defining EN subtypes. The height of the ridgeline represents cell density. Dash lines represent the border between layers, calculated based on the CD distribution of layer-defining EN subtypes. **b**, Ridgeline plots reveal further spatial complexity among EN-ETs, deep layer (Layers 5&6) EN-IT, and upper layer (Layers 2-4) EN-ITs in the PFC at GW34. **c**, Spatial maps and ridgeline plots show the six-layer structure can be visualized and quantitatively defined by the distribution of layer-defining EN subtypes at GW22, despite lack of cytoarchitectural differences between layers. **d, e**, Spatial maps and ridgeline plots showing the cortical layers in the PFC at GW15 and GW20. **f**, Histogram showing the relative proportion of each cortical layer at GW15 to 34 in the PFC, parietal cortex (Par), and occipital cortex (Occi). Average is taken for replicate experiments; error bars represent standard deviation when applicable. **g, h, i**, Spatial maps (left) and ridgeline plots (right) showing the distribution of EN-ETs (**g**), deep layer EN-ITs (**h**), and upper layer EN-ITs (**i**) in the PFC and V2 at GW22. While many clusters exhibited area-dependent abundance, their laminar localization is highly consistent between PFC and V2. **j**, Violin plots showing the laminar distribution of H3 inhibitory neuron (IN) subtypes within the CP of PFC at GW22 and GW34. Scale bars, 500 µm.

Surprisingly, we discovered a significant concentration of inhibitory neurons (IN) and immature IN precursors (INP) in the VZ of the dorsal forebrain during GW20-22, a phenomenon not seen at GW15 (**Fig. 1d, h, Extended Data Fig. 2c, 3a, b**). By GW20, these IN populations outnumbered both RG and IPC within the VZ across most cortical areas, except in the VZ of occipital cortex where INs were notably sparse (**Extended Data Fig. 3b, c**). The degree of VZ localization varied between IN subtypes, but VZ-enriched subtypes included those with transcriptional signatures resembling each of the three ganglionic eminence (GE) structures: lateral (LGE), caudal (CGE), and medial (MGE) (**Extended Data Fig. 3d, e**). MERFISH and Visium analyses showed high expression of IN marker genes in the VZ, including those traditionally associated with olfactory bulb interneurons such as *CALB2, PBX3, TSHZ1* (**Extended Data Fig. 3f-j**)^27^. Although our analysis did not trace the lineage of these dorsal VZ-concentrated INs, their unexpected localization raises the possibility that, contrary to being ventrally born and migrating into the cortex, they could include some dorsal-born INs, aligning with findings from several recent studies^28,29,30,31^.

**Fig. 3:**
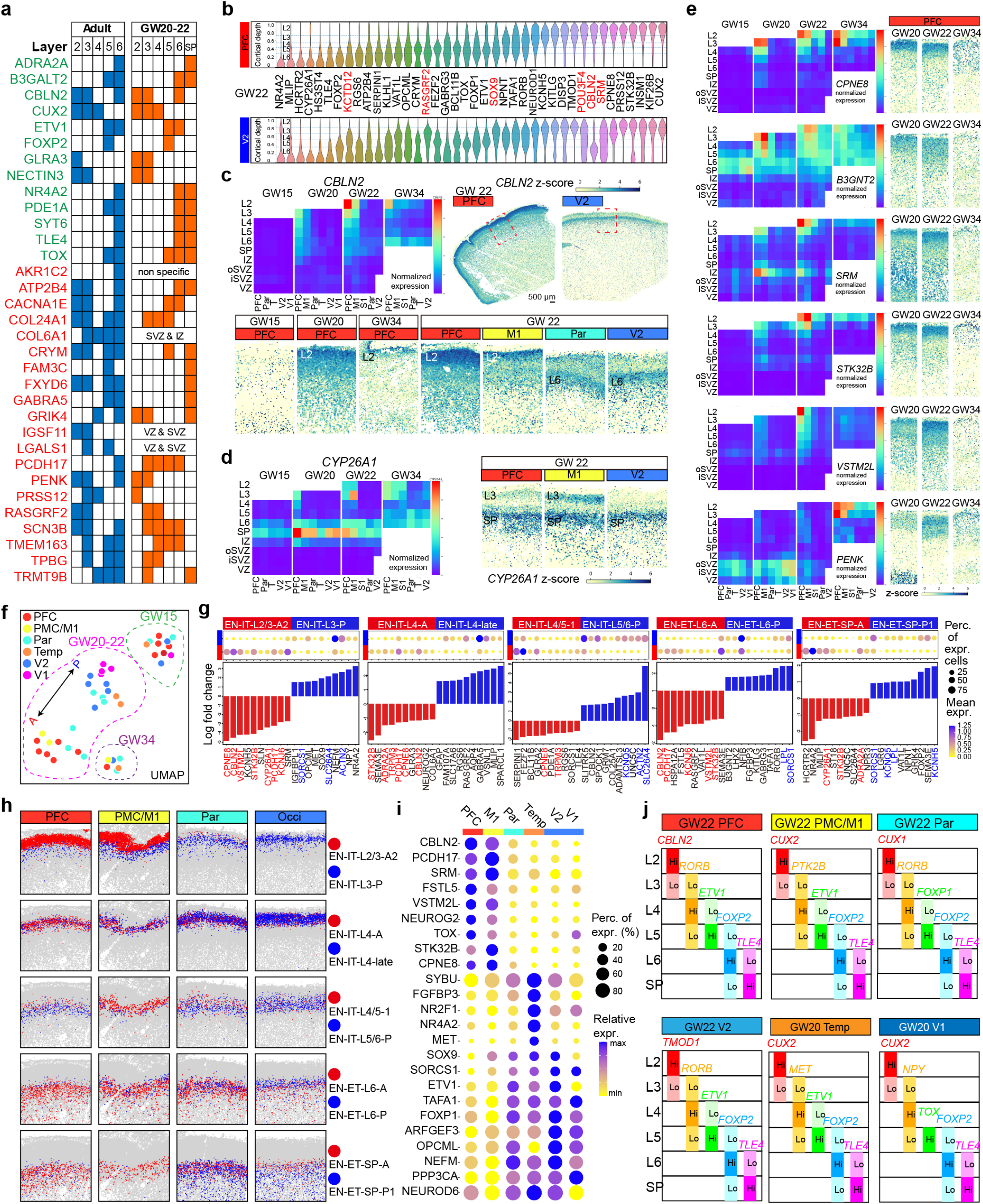
Laminar and areal dynamic gene expression underlies neuronal specification. **a**, Table compares the laminar expression patterns of validated layer marker genes between the adult human PFC^40^ and mid-gestation PFC. Genes that show conserved laminar expression pattern are colored in green, otherwise in red. **b**, Violin plots show the laminar expression patterns of layer-dependent genes within the CP of the PFC and V2 at GW22. Width of the violins represent the cumulative normalized expression in the cells located at a cortical depth (CD). Genes that are enriched in different layers between PFC and V2 are highlighted in red. **c**, Summary expression heatmap and z-score spatial maps showing the spatial-temporal expression pattern of *CBLN2*. The summary heatmap shows different cortical layers and laminar structures from top to bottom as rows, and cortical areas and gestational ages as columns, organized from anterior to posterior and from young to old. Average is taken for replicate samples from the same area and GW. T, temporal cortex. Scale bar, 500 µm. **d**, Summary expression heatmap and z-score spatial maps for *CYP26A1*. **e**, Summary expression heatmaps and z-score spatial maps of genes identified with Layers 2&3 enrichment in the PFC. **f**, UMAP plot of EN subtype composition in the CP shows an anterior-posterior spectrum for GW20-22. **g**, Dot plots and histograms of fold change for top differentially expressed genes (DEGs) between pairs of anterior- and posterior-enriched EN subtypes for each cortical layer at GW22. Genes that appear repeatedly are highlighted in red (for anterior-enriched) or in blue (for posterior-enriched). **h**, Spatial graphs showing the laminar and areal distribution of pairs of anterior- and posterior-enriched EN subtypes. **i**, Dot plot showing the expression of top areally-enriched genes at GW20 and 22 in all post-migratory EN (EN-IT and EN-ET cell classes). **j**, Schematic summary of the combinations of 5 marker genes that enable the approximation of cortical layers of different cortical areas at GW20 to 22.

### Progressive formation of cortical layers

The establishment of cortical layer structures during human fetal development has long been elusive due to the lack of cytoarchitectural features and validated molecular markers.^26,32,33,34^ In the adult cerebral cortex, a clear six-layer structure is observed, with neurons arranged into morphologically distinct layers^35^. However, such cytoarchitectural distinctions were not present between GW15 and 22, becoming only discernable at GW34 (**Extended Data Fig. 4a**), consistent with previous observations^36^. At GW34, the cytoarchitecture of cortical layers in the CP was comparable to that of the postnatal human cortex, featuring a granule layer (Layer 4) of densely packed ENs, flanked by a less dense Layer 3 and a more compact Layer 2 above, and Layers 5 and 6 below (**Extended Data Fig. 4a**).

**Fig. 4:**
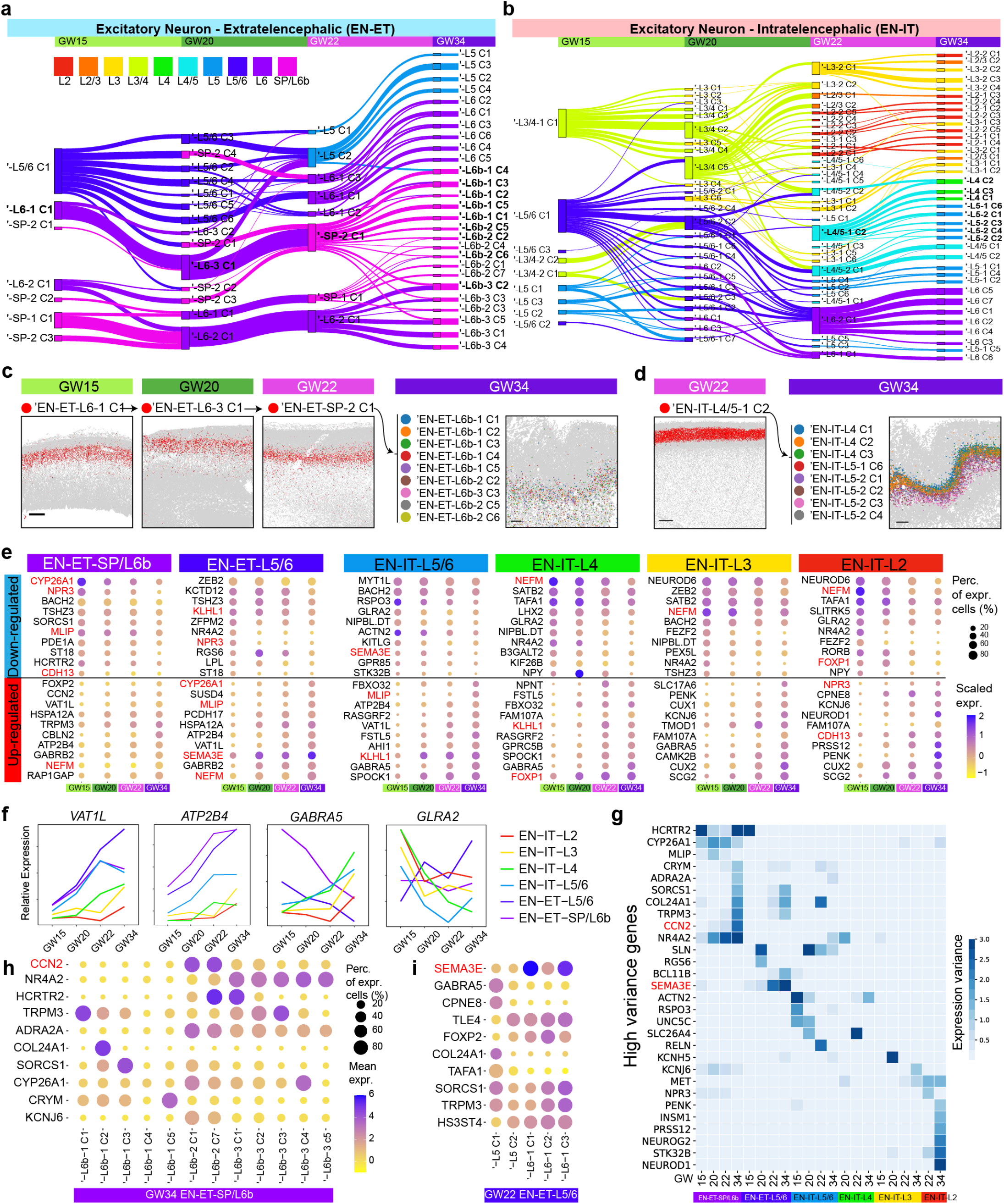
Diversification of EN subtypes over time show continued post-mitotic fate specification. **a, b**, Sankey diagram showing the correspondence between EN-ET (**a**) and EN-IT (**b**) subclusters from different gestational time points. Clustering was performed separately for each gestational age and the number of subclusters was determined single cell significant hierarchical clustering (scSHC).^42^ Nodes represent EN subtypes from this alternative clustering strategy. To differentiate these clusters from those identified through integrated analysis, an apostrophe (‘) prefixes the names of scSHC clusters. Thickness of edges represents the fraction of cells showing with correspondence, and edges with <0.1 fraction were hidden. Nodes and edges are colored based on the most enriched layer identity of the EN subtype. **c**, Spatial graphs for an inferred lineage (bolded in **a**) of Layer 6/SP EN-ETs across GW15 to GW34 show a conserved stream from GW15 to 22 before dramatic diversification at GW34. Scale bars, 500 µm. **d**, Spatial graphs showing a GW22 cluster EN-ITs spamming both Layers 4 and 5 specifying into seven GW34 clusters with refined layer specificity in either Layer 4 or 5 (bolded in **b**). Scale bars, 500 µm. **e**, Dot plots showing the top genes that are down- or up-regulated with gestational age among EN subtypes from each layer-based group. The genes that are downregulated over time in some categories but upregulated in others are highlighted in red. **f**, Curve plots show the change of relative expression over time in different groups for selected genes. **g**, Heatmap showing the genes with high expression variance among EN subtypes within the same group and gestational age. **h, i**, Dot plots showing the expression of high variance genes among EN subtypes in EN-ET-SP/L6b group at GW34 and EN-ET-L5/6 group at GW22.

Our MERFISH analysis revealed that most EN subtypes showed layer-dependent distribution within the CP and the SP, resulting in detailed layer divisions that evolved over gestational weeks. To precisely analyze the cortical layer distribution of H3 EN subtypes, we measured their relative position within the CP as cortical depth (CD) (**Fig. 1g, Methods**). For individual H3 EN subtypes, laminar distribution was remarkably stable across cortical areas and gestational ages, with gradual refinement in layer specificity over time, particularly evident when comparing GW34 to earlier time points (**Extended Data Fig. 4b**). While the degree of laminar dispersion varied between clusters, we identified a subset of narrowly dispersed subtypes exhibiting distribution patterns that aligned with one of the six morphologically defined cortical layers at GW34 (**Fig. 2a, b, Extended Data Fig. 4f**). Based on the CD distribution of these clusters, we defined the border between cortical layers quantitatively for each annotated region, enabling the assignment of individual cells to specific cortical layers according to their CD values (**Fig. 2a, Methods**). EN clusters at H2 and H3 were annotated for their predominant localization in one or two cortical layers at GW34 (**Supplementary Table 4**), a classification supported by cell type correspondence analysis with datasets from the human adult cortex (**Extended Data Fig. 4c**)^37^.

In striking contrast to conventional characterizations, the distribution of layer-defining H3 EN subtypes clearly delineated all six cortical layers at GW22, despite the absence of morphological features (**Fig. 2c**). Cortical layers evolved progressively over time, with Layers 4-6 visualized at GW15 and Layers 3-6 at GW20, consistent with the inside-out sequence of cortical layer formation and proposed corticogenesis timeline (**Fig. 2d, e, Extended Data Fig 4d-f**)^2,10,^. We quantified the relative proportions of cortical layers across various areas and gestational ages (**Fig. 2f**). Comparison between the prefrontal cortex (PFC) and the secondary visual cortex (V2) at GW22 highlighted distinct layer proportions, with the PFC exhibiting relatively larger Layers 2 and 3, while V2 displayed a larger Layer 4 (**Fig. 2c**). This alignment with the known relative layer proportions in the adult human cortex demonstrated that the mid-gestation blueprint of cortical layer thickness is predictive of developmental outcomes^3^.

Contrary to the stability of their laminar localization, the relative abundance of EN subtypes varied dynamically across areas and ages. For example, cluster EN-ET-L6-P, the most abundant subtype in Layer 6 of V2, was sparse in the PFC; while EN-IT-L3-A, the most abundant subtype in Layer 3 of PFC, was dramatically outnumbered by EN-IT-L3/4-P in V2 (**Fig. 2g-i**). At GW15, two clusters of EN-ITs (EN-IT-L4/5-early and EN-IT-L3/4-early) showed broad distribution across the CP and outnumbered other EN-IT subtypes with clearer layer specificity, but became mostly absent in GW20-34 (**Fig. 2h, i**). These two clusters also lacked unique marker gene expression (**Extended Data Fig. 2a**), indicating that the spatial organization of EN subtypes co-developed with their specified molecular identities.

The distribution of EN-ET subtypes was restricted to Layer 5 or Layer 6 with minimal crossing of the Layer 4-5 border, but the spatial complexity of EN-IT distribution surpassed the conventional six-layer definition (**Fig. 2g-i**). Most EN-IT subtypes spanned multiple layers and exhibited bell-shaped distribution curves irrespective of layer boundaries, resulting in extensive laminar intermixing (**Fig. 2h, i, Extended Data Fig 3-f**). This complexity was retained even in GW34, despite the emergence of distinct cytoarchitectures (**Fig. 2b**). The finding that different cortical layers do not contain mutually exclusive subtype repertoires mirrors observations in the adult cortex, implying that this developmental feature persists into adulthood^37^. Additionally, some EN subtypes showed enrichment within narrower regions inside a cortical layer. For example, EN-IT-L6 subtypes were more enriched in the upper portion of Layer 6, while EN-ET-SP subclusters were enriched in the lower portion of Layer 6, potentially providing the blueprint for distinct Layers 6a and 6b (**Extended Data Fig. 4b**). Between GW34 and earlier gestational ages, several EN-ET-SP clusters displayed an upward shift into the upper Layer 6, supporting the existence of a secondary translocation of SP neurons after their initial migration after neurogenesis reported in a prior study^39^.

We extended our analysis to quantify the CD distribution of IN subtypes. Cortical layer specificity was not evident during GW15-22, suggesting that layer-dependent IN organization had not been established at mid-gestation (**Fig. 2j**). Nevertheless, some layer specificity began to emerge at GW34, particularly among IN-SSTs, which primarily localized in Layers 2&3, consistent with mouse and human adult cortices^37^.

### Dynamic laminar gene expression

Because canonical marker genes identified in the adult human cortex or in mice exhibit reduced specificity for both laminar positioning and neuronal subtypes in mid-gestation fetal cortex, we sought to identify more applicable markers^7,32^. Building on our CD framework, we assessed the expression patterns of marker genes validated in the adult human cortex and observed significant divergence in their expression patterns^40^ (**Fig. 3a**). Notably, genes associated with deep layer EN-ETs, such as *B3GALT2, ETV1, NR4A2, SYT6, TLE4*, and *TOX*, showed greater conservation of expression patterns than those for EN-ITs. Seeking to identify cortical layer markers specific to the human fetal cortex, we identified genes with layer-dependent enrichment and quantified their laminar expression within the CP for each experiment (**Fig. 3b, Extended Data Fig. 5a**). Immunostaining for selected markers using validated antibodies showed concordance of mRNA localization detected by MERFISH with protein expression (**Extended Data Fig. 5b**). While many genes exhibited laminar-restricted expression, their span across multiple layers suggested the improbability of pinpointing a singular gene as a definitive marker for any specific cortical layer at mid-gestation. Instead, combinations of several genes delineate cortical layers more effectively.

**Fig. 5:**
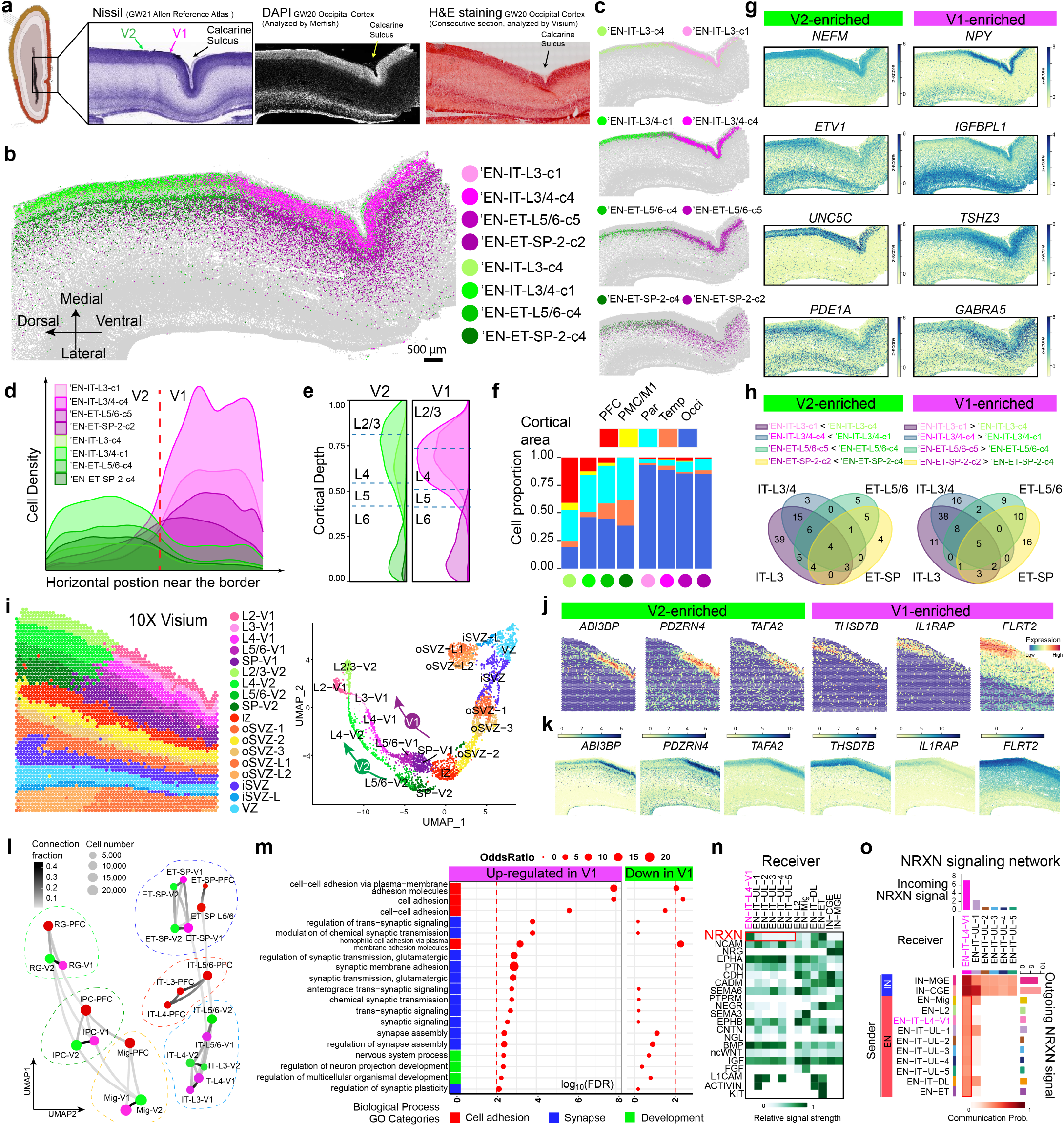
Sharp molecular border between V1 and V2 at GW20 reveals early V1 specification. **a**, The primary (V1) and secondary visual cortices (V2) do not exhibit morphological differences at GW20-21. Schematics and Nissl staining are taken from the Allen Reference Atlas for GW21 human fetal brain^48^. DAPI staining image co-captured with MERFISH; H&E staining image co-captured with Visium. **b, c** Spatial maps for selected EN subtypes show distinct border between V1 and V2 marked by sharp transition at GW20. Scale bars 500 µm. **d**, Ridgeline plots showing the horizontal cell density profile for V1- and V2-enriched EN subtypes in the CP near the border. **e**, Ridgeline plots showing the laminar distribution for V1- and V2-enriched EN subtypes. Dash lines represent the borders between cortical layers. **f**, Histograms for areal distribution show V2-enriched subtypes distribute broadly in other cortical areas, while V1-enriched subtypes are exclusive to the occipital cortex. **g**, Z-score spatial maps of V1- and V2-enriched genes. **h**, Venn diagrams for the overlap between V2-enriched (left) and V1-(right) DEGs across the four pairs of subtypes in **c**. Only strong DEGs with Log fold change >0.5 and adjusted p-value <0.0001 were considered. See **Supplementary Table 7. i**, Spatial graph (left) and UMAP (right) from Visium analysis show clear V1-V2 border across all cortical layers and two parallel UMAP trajectories for V1 and V2 that bifurcated from the SP. Capture area, 6.5 × 6.5 mm. **j**, Spatial graphs showing additional genes identified by Visium exhibiting clear transition at the V1-V2 border. **k**, Imputed expression patterns for additional V1-V2 border genes matches with Visium results. **l**, Constellation plots of cell types in different cortical areas show V1 and V2 share a developmental lineage that is distinct with PFC. Matching nodes between V1 and V2 have connection fraction > 0.2 except for EN-IT-L3 and EN-IT-L4. See **Supplementary Table 8. m**, Gene ontology (GO) analysis reveals synapse and cell adhesion-associated biological processes are upregulated in V1-specific Layer 4 neurons. **n**, Heatmap for incoming signaling pathways shows neurexin (NRXN) signaling is specifically enriched in EN-IT-L4-V1 cluster comparing to other upper layer (UL) EN-IT clusters. **o**, Network heatmap shows EN-IT-L4-V1 is unique among upper layer EN-ITs to receive NRXN signaling from both IN and EN sources, leading to significant increase in overall interaction strength. The bar graphs on the top and right side are sum of interaction strength as incoming and outgoing signals, respectively.

Remarkably, some genes displayed laminar expression patterns that varied significantly with gestational age and cortical area. For example, *CBLN2*, a synaptic organizer with a hominin-specific variant in a retinoid acid responsive enhancer^41^, was specifically expressed at relatively low levels in both EN-ITs and EN-ETs in Layer 6 across all cortical areas from GW15 to 34. Additionally, *CBLN2* displayed strong frontal enrichment in Layers 2&3 at GW22 and GW34, correlating with its role in synaptic spine development (**Fig. 3c**)^41^. Consequently, *CBLN2* could serve as a Layer 6 marker in parietal, occipital, and temporal cortices, and as a Layer 2&3 marker in the frontal cortex post-GW20. Similarly, *CYP26A1*, a retinoic acid metabolizing enzyme^13^, manifested frontal-specific expression in Layers 2&3, while remaining highly expressed in EN-ETs across Layer 6 and the SP in all areas (**Fig. 3d**). Moreover, a group of genes demonstrated pronounced frontal enrichment in Layers 2&3, but their peak expression timing varied: *CPNE8* maintained high expression throughout GW20-34; *B3GNT2* had peak expression at GW20, with *SRM, STK32B*, and *VSTM2L* following at GW22, and *PENK* peaking at GW34. This suggested the frontal specification of Layers 2&3 underwent temporally dynamic transcriptional changes (**Fig. 3e**). For visualization and comparison, we compiled the expression patterns of all genes within our MERFISH panel into a structured heatmap, serving as a compendium across cortical areas, layers, and developmental stages (**Source Data Fig. 3**).

### Areal specification of neuronal subtypes

Bulk and single-cell RNA sequencing has unveiled gene expression patterns that vary by cortical area in the fetal human cortex, often manifesting as gradients along the anterior-posterior (A-P) or medial-lateral (M-L) axes^6,7,10,33^. Our methodology enabled precise analysis of the area-dependent specification of neuronal subtypes within each cortical layer. While H2 EN cell types exhibited clear layer-dependent localization, their distributions remained largely uniform along the A-P axis (**Extended Data Fig. 1k, l**). In contrast, at the H3 subtype level, 7/25 EN-ET H3 subtypes and 15/24 EN-IT subtypes displayed gradient-like area enrichment, albeit modestly at GW15 (**Extended Data Fig. 5c**). Interestingly, the temporal (Temp) and primary visual cortices (V1) were unique in containing exclusive EN-IT subtypes absent in other cortical areas. Unsupervised clustering by EN subtype compositions showcased the distinction between major cortical areas at GW20 and 22, presenting an A-P spectrum on the UMAP projection (**Fig. 3f**). In contrast to post-migratory ENs (EN-IT and EN-ET), subtypes of migrating EN (EN-Mig), RGs, and IPCs displayed no discernible areal preference even in the Temp and V1 (**Extended Data Fig. 5c, d**).

To elucidate the genes driving cortical arealization among transcriptionally and spatially related EN subtypes, we selected 5 pairs of subtypes with opposing A-P distributions at GW20 and 22, serving as representatives of areal specification in 5 categories of neurons: Layers 2&3, Layer 4, deep layer EN-IT, deep layer EN-ET, and SP EN-ET (**Fig. 3g, h**). The opposing clusters were transcriptionally similar, leading us to hypothesize that the differentially expressed genes (DEGs) within each pair were decisive in areal differentiation. The lists of top anteriorly and posteriorly enriched DEGs were notably distinct across the five categories, indicating that arealization was not orchestrated by a uniform transcriptional program across the entire CP, but was instead regulated in a layer- and subtype-specific manner (**Fig. 3g, Supplementary Table 5**). However, several genes with strong anterior enrichment, such as *CBLN2, CPNE8, VSTML2*, and *STK32B*, frequently emerged as top DEGs in multiple categories (**Fig. 3e**). Interestingly, *CBLN2* was among the top anterior DEGs for both upper layer EN-ITs and deep layer EN-ETs, but not for deep layer EN-ITs, suggesting its role in arealization is not limited to one neuronal class (**Fig. 3g**). Upon examining all significant DEGs (adjusted p-value <0.05 and Log fold change > 0.5), we found 4 genes consistently enriched in anterior regions (*STK32B, HCRTR2, PCDH17*, and *SERPINI1*) and 8 in posterior regions (*FOXP1, KITLG, KLHL1, MET, NPY, OPCML, RORB*, and *SOX9*) across all 5 neuronal categories (**Extended Data Fig. 6a**). These common DEGs also manifested strong anterior or posterior enrichment in the post-migratory EN population as a whole, suggesting their potential as overarching markers for cortical arealization (**Fig. 3i**). Their areal enrichment patterns varied in other cell classes, limited among RG, IPC, INs, but notably consistent within EN-Mig (**Extended Data Fig. 6b**). This observation indicates that the transcriptional mechanisms underpinning arealization exhibit considerable dynamism along the differentiation trajectory, with ENs acquiring their areal identities both during and subsequent to radial migration.

Drawing from our comprehensive spatiotemporal gene expression data, we developed context-aware marker sets to facilitate the approximated visualization of mid-gestation cortical layers using conventional methods, such as multichannel in situ hybridization (**Fig. 3j**). During GW20-22, *FOXP2* and *TLE4* labeled Layer 6 and SP respectively in all cortical areas. *ETV1* marked Layer 5 in PFC, PMC, M1, V2 and Temp, while *FOXP1* and *TOX* marked Layer 5 in Par and V1, respectively. Markers for Layers 2, 3 and 4 showed increasing areal differences, including *CBLN2, CUX2, CUX1, TMOD1, RORB, PTK2B, MET* and *NPY* (**Extended Data Fig. 6c, d**). It is important to recognize that while these genes’ expressions peaked within their assigned layers, they were not confined exclusively to those layers, but we can leverage the overlapping patterns of these genes to distinguish layers.

### Neuronal subtype diversification over time

To analyze EN subtype specification over time, we devised an alternative approach, by clustering cells from each gestational age separately instead of combining them. Utilizing the single-cell significant hierarchical clustering (scSHC) pipeline^42^, we determined the number of EN subclusters that achieved transcriptional differences above a fixed statistical significance threshold between clusters (**Methods**). To differentiate these clusters from those identified through integrated analysis, an apostrophe (‘) precedes the names of scSHC clusters. This analysis involved down-sampling to 500,000 single cells for each of the four gestational time points to negate any bias arising from varying cell counts. Within EN-ET and EN-IT cell classes, the number of distinct scSHC clusters increased over gestational time, as the analysis yielded 16, 39, 40 and 62 subclusters at GW15, 20, 22, and 34, respectively (**Supplementary Table 6**).

To infer the lineage relationships between EN-IT and EN-ET subtypes from different gestational ages, we applied a deep-learning cell type classifier to predict the likely origin of clusters at the earlier time (**Methods**)^43^. The resulting correspondence flows showed high stability of layer identities across subtypes from different ages, and uncovered continued diversification particularly dramatic between GW22 and GW34, aligning with the predicted role of the third trimester in neuronal subtype specification (**Fig. 4a, b**)^44^. Deep layer EN-ET subtypes demonstrated more stability than EN-ITs during mid-gestation but also underwent more extensive diversification from GW22 to GW34, likely associated to their earlier birthdate and more advanced maturity (**Extended Data Fig. 7a**). For instance,’EN-ET-L6-2-Cluster1 at GW15 corresponded strongly with a single cluster at GW20 and GW22, before branching into nine Layer 6b EN-ET clusters by GW34 (**Fig. 4c, Extended Data Fig. 7b**). The third trimester specification also led to the refinement of layer specificity, exemplified by the ‘EN-IT-L4/5-1-Cluster2 from GW22 differentiating into five Layer 5 subtypes and three Layer 4 subtypes by GW34 (**Fig. 4d**).

Analyzing transcriptional trajectories from GW15 to GW34, we analyzed the dynamics of EN transcriptional specification by identifying genes up- or down-regulated across gestational ages within EN subtypes grouped by correspondence flows (**Fig. 4e**). Notably, among the top DEGs, those associated with the general functional maturation of neurons, like *ATP2B4* (ATPase plasma membrane Ca2+ transporting 4) and *VAT1L* (vesicle amine transport protein 1 homolog), showed synchronous regulation across all layers (**Fig. 4f**). In contrast, genes exhibiting upregulation in specific groups while being downregulated in others were often specific neurotransmitter receptors such as *GABRA5* (GABA receptor subunit alpha5) and *GLRA2* (glycine receptor alpha2), hinting at transcriptional changes stemming from differential functional specification (**Fig. 4f, Extended Data Fig. 7c**).

To pinpoint the genes driving subtype specification, we focused on those with the highest variance between EN subtypes within the same group and gestational age (**Fig. 4g**). Many high variance genes demonstrated specific laminar and areal enrichment (**Extended Data Fig. 6a, c, 7d**). For example, *CCN2* (or CTGF), emerged as one of the genes with the highest variance among EN-ETs in SP and Layer 6b specifically at GW34 (**Fig. 4h**). Given *CCN2*’s known role in differentiating between upright pyramidal neurons (PN) and other PNs in Layer 6b, its upregulation in certain EN subtypes could indicate the commencement of PN subtype specification^45^. Conversely, *SEMA3E* (semaphorin 3E), known for its involvement in specifying a subset of L5 PNs projecting to higher-order thalamic and pontine nuclei^46^, showed high variance among EN-ET-L5/6 subtypes as early as GW22 (**Fig. 4i**). This suggests the specification process for these Layer 5 neurons might occur between GW20-22, preceding Layer 6b PN specification, despite their later birthdate. These examples underscore that subtype specification follows independent, rather than synchronized, timelines.

### A precociously sharp transcriptional border between V1 and V2

To elucidate the precise differentiation of adjacent cortical areas, we focused on the primary (V1) and secondary visual cortices (V2), with the calcarine sulcus serving as a morphological landmark for their location. While V1 and V2 in the adult cortex are distinguished by a pronounced morphological border^3,47^, they appeared completely homogeneous in mid-gestation, as neurons were densely packed without cytoarchitectural distinctions (**Fig. 5a, Extended Data Fig. 8a**). Strikingly, MERFISH analysis as early as gestational week 20 identified a clear border between V1 and V2, defined by four opposing pairs of scSHC subtypes, two EN-IT pairs in Layers 3&4 and two EN-ET pairs in Layers 5&6 and SP (**Fig. 5b, c**). Their boundary featured a swift transition and was most abrupt in Layers 3&4 (**Fig. 5d**). The two subtypes within each pair (for example, ‘EN-IT-L3-c1 and -c4) were derived from the same H2 cell type and showed identical laminar distribution (**Fig. 5c, e**). In contrast, cell subtype localization remained uniform in the IZ, oSVZ, iSVZ, and VZ, indicating that V1 and V2 identities are distinctly specified in post-migratory ENs, but not in migrating ENs, IPCs, RGs (**Extended Data Fig. 8b, c**). V1-specific EN subtypes were absent in all other areas, unlike V2-enriched subtypes, which were broadly distributed with some presence even in the anterior areas (**Fig. 5f, Extended Data Fig. 8d**). This pattern showcased a unique, localized program for neuronal specification within V1, a finding consistently replicated across multiple experiments and independent samples (**Extended Data Fig. 8e and f**).

A spatial transition of gene expression underlies the neuronal subtype border (**Fig. 5g**). While most DEGs showed expression level changes within 1-2 mm of the border, *NPY* particularly stood out by displaying a pronounced binary switch, with a dramatic enrichment observed in Layers 3&4 of V1 (**Fig. 5g**). Other V1-enriched genes included *IGFBPL1* in Layers 3&4, *TSHZ3* in Layers 5&6, and *GABRA5* in SP, while V2-enriched genes included *NEFM* and *ETV1* in Layers 3&4, *UNC5C* in Layers 5&6, and *PDE1A* in SP (**Fig. 5g**). Differential gene expression analysis within each pair of V1- and V2-clusters revealed that few DEGs were universally shared among all four pairs (5 for V1, 4 for V2), but more DEGs were shared within EN-IT and EN-ET, suggesting the V1-specification of EN-IT and EN-ET may take place via separate programs (**Fig. 5h, Supplementary Table 7**). The absence of a neuronal subtype border at GW15 suggested that V1 specification occurred between GW15 to 20, likely concurrent with the formation of Layers 3&4 in the CP (**Extended Data Fig. 9a**). Nevertheless, some V1- and V2-enriched genes in deep layers at GW20, such as *TSHZ3* and *GABRA5*, showed conserved differential expression patterns in the corresponding putative areas at GW15 (**Extended Data Fig. 9b, c**).

To expand our analysis of the V1-V2 border to the whole transcriptome, we conducted Visium and snRNAseq analysis on consecutive sections analyzed by MERFISH. At 55µm-diameter spatial resolution, Visium analysis confirmed a clear border between V1 and V2 across all cortical layers and SP, whereas no border was observed in the IZ, oSVZ, iSVZ, and VZ (**Fig. 5i**). Such distinction was supported by the UMAP pattern, demonstrating two parallel trajectories for V1 and V2 that bifurcated at the SP and beyond (**Fig. 5i**). The spatial gene expression patterns measured in Visium precisely replicated MERFISH (**Extended Data Fig. 10a, b**). Comparison between matching V1 and V2 spots revealed additional genes that exhibited pronounced changes at the border (**Fig. 5j, Extended Data Fig. 10c**). Among them, *ABI3BP* and *PDZRN4* showed V1-exclusive expression pattern with a binary switch at the border of Layers 3&4, akin to that of *NPY*, while V2-enriched genes such as *FLRT2* and *IL1RAP* demonstrated more gradual transitions over a short distance (**Fig. 5j**).

To expand the gene content of the MERFISH cells, snRNAseq analysis of 91,898 cells from the same samples was integrated with the MERFISH cells to impute expression of genes outside of the MERFISH panel using environmental variational inference (ENVI) (**Extended Data Fig. 10d**)^22^. The imputed spatial expression patterns of additional V1-V2 border markers, such as *ABI3BP* and *PDZRN4*, precisely matched Visium results, showcasing the sharp Layers 3&4 border that coincided with the *NPY* expression (**Fig. 5k**). Utilizing the imputation results, we expanded our MERFISH analysis to include the top 1000 highly variable genes from snRNAseq, and constructed a constellation plot to analyze the nearest neighbor connection between cell type nodes of PFC, V1, and V2 (**Fig. 5l, Methods**). Nodes representing RG, IPC, and EN-Mig between V1 and V2 were connected strongly (connection fraction >0.2) but not with PFC, suggesting that the transcriptional distinction between V1 and V2, unlike A-P distinction, was not well-established from the progenitor stage. Notably, EN nodes of V1 and V2 were strongly connected in SP and Layers 5&6, but not in Layers 3&4 (**Fig. 5l, Supplementary Table 8**). The transcriptional proximity between V1 and V2 throughout the differentiation trajectory, despite the sharp molecular border, suggested a model wherein V1-specific EN subtypes shared a developmental lineage with V2-enriched EN subtypes but were induced to diverge due to external factors post-migration.

To uncover potential mechanisms driving V1 specification, we conducted cell label transfer analysis, revealing that the EN-IT-L4-V1 cluster from snRNAseq strongly and exclusively corresponded to V1-specific Layer 3&4 subtypes from MER-FISH, while the EN-IT-UL-2 cluster corresponded to those enriched in V2 (**Extended Data Fig. 10e**). The exclusivity of V1-specific subtype was validated by the absence of EN-IT-L4-V1 cells in the PFC sample, and DEGs between the two clusters were highly consistent with MERFISH and Visium results (**Extended Data Fig. 10f-i, Supplementary Table 9**). Gene ontology (GO) analysis of V1-enriched DEGs revealed cell adhesion and synapse assembly as the most associated biological processes, with most associated cellular components being synapse-related membrane components (**Fig. 5m, Extended Data Fig. 10j**). These GO terms were not significant for V2-enriched DEGs, suggesting that V1-specific Layers 3&4 EN-IT subtypes underwent unique aspects of synaptogenesis at this early stage of development. This observation was corroborated by ligand-receptor cell-cell communication analysis using CellChat^49^, which highlighted neurexin-neuroligin (NRXN-NLGN) signaling, fundamental to synapse formation^50^, as the most uniquely enriched signal received by EN-IT-L4-V1 (**Fig. 5n**). While other upper layer EN-IT clusters also received NRXN signal from IN sources, EN-IT-L4-V1 was unique in also receiving strong NRXN signals from EN sources (**Fig. 5o, Extended Data Fig. 10k**). However, as our data only included cells in the cerebral cortex, further investigation is required to determine whether external sources of NRXN-NLGN, such as thalamic neurons projecting to V1, contribute to the heightened synaptogenesis in V1. Recent studies on postnatal mouse V1 identifies critical roles of visual inputs during the postnatal “critical period (P21-38)” in establishing V1-specific neuronal identities, marked by *CDH13* down-regulation and *TRPC6* up-regulation in Layers 2&3^51^. Our GW20 V1 data exhibited expression patterns of *CDH13* and *TRPC6* that aligned with mouse P28 but not with earlier times (**Extended Data Fig. 10l**). Notably, lower *CDH13* and higher *TRPC6* levels observed in V1 compared to V2 at GW20 suggested V1-specific *CDH13* downregulation and *TRPC6* upregulation might accompany V1-V2 fate divergence. This implies that thalamic afferents may contribute to human V1-V2 differentiation at a much earlier developmental stage than previously predicted^4,10^.

## Discussion

In this study, we employed MERFISH to create the most extensive spatially resolved single-cell atlas of the developing human cerebral cortex to date. To facilitate broad accessibility of our data, we have built an interactive web-based browser for our spatial atlas, which will become publicly available upon publication of the study. Our study emphasizes the synergistic analysis of molecular and spatial data, yielding insights previously unattainable through droplet-based single-cell methodologies. To achieve systematic integration of spatial information with molecular profiling, we developed a novel framework that converts tissue cytoarchitecture into measurable parameters. This approach allowed us to delineate cortical layers based on the quantitative distribution of specific neuronal subtypes, and find that the CP’s six-layer structure is already in place by GW22, long before morphological differentiation is evident.

Contrary to the dynamic nature of gene expression patterns within laminae, we observed a remarkable stability in the laminar distribution of transcriptionally defined EN subtypes across cortical areas and time. Nevertheless, while all EN subtypes showed distinct laminar-dependent distribution, most EN-IT subtypes exhibited a near-normal distribution curve within the cortical plate (CP), irrespective of cortical layer boundaries. This observation aligns with recent discoveries in the adult cortex^37^, and suggests that while the six-layer model facilitates our analysis design and data interpretation, the structural complexity of neuronal lamination likely surpasses the traditional six-layer model, underscoring the advantage of direct spatial profiling over microdissection.

Our investigation into cortical arealization suggests an integrated view of the “protomap” and “protocortex” models^10^. Echoing prior studies, we observed gradients of transcriptional changes and cell subtype variation along the A-P axis, indicating a foundational role for morphogen gradients in establishing preliminary areal identities^6^. Yet, our identification of a distinct border between V1 and V2 by GW20 suggests that detailed areal specification may occur concurrently with the peak of neurogenesis, prompting a reevaluation of the cortical development timeline. The distinctness of the V1-V2 border, coupled with the exclusivity of V1-specific subtypes and the uniformity of migrating neurons and progenitors, starkly contrasts the outcome of continuous A-P morphogen gradient as seen across other areas analyzed.

The pronounced V1-V2 demarcation implicates a potential role for afferents, presumably from the dorsal lateral geniculate nucleus (dLGN) of the thalamus, which reach the SP and CP by GW12-14 in human, much earlier than in rodents^52^. These afferents initiate synapse formation and transmit spontaneous thalamic activity to the cortex, altering V1 neuronal identities via activity-induced mechanisms^10,52,53,54,55^. Damage to these afferents in macaque disrupts normal V1 cytoarchitecture formation^56,57^, whereas studies in rodents demonstrated visual disruptions lead to failure in fate specification of V1 neurons, resulting in a shift towards V2-like “default” molecular identities and blurring of V1-V2 border^51,53,58,59^. The V1-V2 boundary observed in EN-ITs of Layers 3&4 and in EN-ETs of the deep layer and SP, but not in deep layer EN-ITs, may result from thalamic axons’ initial interaction with SP neurons and subsequent targeting into Layer 4^60^. Furthermore, the similarity in key marker expression patterns observed after vision-dependent V1 specification in postnatal mice hints at a much earlier V1 specification timeline in human than rodents^53^. This development is in line with the timing of eye-opening in human fetuses around GW26, compared to P14 in mice^61^. While the calcarine sulcus allowed identification of V1 and V2 independent of gene expression, the lack of similar landmarks precluded identification of the primary somatosensory cortex and primary auditory cortex in our samples, but it is possible that further analysis with comprehensive areal coverage would uncover border formation mechanisms akin to V1 in other primary sensory areas.

Our findings reveal that the expression of individual genes varies considerably across different cortical areas and developmental stages, challenging the traditional reliance on a limited set of canonical layer markers to delineate the evolving landscape of the developing human cortex. The lack of accurate markers has impeded the advancement of brain organoid studies, which depend on comparisons with primary fetal samples for benchmarking. Our analysis indicates that, during midgestation, many markers lack the layer specificity canonically attributed to them, and the observed lack of layering in cortical organoids might partially reflect uninformative marker choice^62^. Therefore, an accurate characterization of neuronal layers in organoids should incorporate markers tailored to the matching developmental stage and cortical area. Our comprehensive molecular and structural mapping provides a foundational reference for future studies in organoid development.

## Methods

### Sample preparation and acquisition

Research performed on samples of human origin was conducted according to protocols approved by the institutional review boards (IRB) of Boston Children’s Hospital and Beth Israel Deaconess Medical Center. Samples were collected after obtaining written informed consent. De-identified human fetal tissues were acquired from two sources: (1) Samples were received after release from clinical pathology at Beth Israel Deaconess Medical Center, with a maximum post-mortem interval of 4 hours. Tissue was transported in Hibernate-E media on ice to the laboratory for research processing. Cortical tissue was then dissected into coronal pieces of 1-3 cm^2^ cross-section area and 0.5-1 cm thickness. The dissection was performed under the supervision of a neuropathologist to provide annotation of cortical areas based on anatomical location and features.

Tissue pieces were then directly frozen in liquid nitrogen and stored in -80°C. (2) Banked de-identified fresh-frozen tissues were obtained from the University of Maryland Brain and Tissue Bank through the NIH NeuroBioBank. Only samples with postnatal interval < 12 hours were used. Cortical area annotation was provided by the NeuroBioBank. Tissue was shipped overnight in dry ice and stored in -80°C. For both sources, only samples with no neurological anomalies were analyzed. Samples were screened for RNA quality by collecting 50µm-thick cryosections, isolating total RNA and measuring RNA Integrity Number (RIN) using the Agilent 4200 TapeStation System, and RNA Integrity Number (RIN) >= 6.5 were used in the study. The samples used are summarized in **Supplementary Table 1**.

### Annotation of cortical areas

The cortical areas of tissue sections used in MERFISH analysis were annotated based on the Reference Atlas for human fetal brain developed by Allen Institute for Brain Science (https://atlas.brain-map.org/)^26^. For GW20 and GW22 samples, dissection of the tissue was performed under the supervision of a neuropathologist to distinguish between the frontal lobe, parietal lobe, occipital lobe, and temporal lobe. The relative anterior-posterior, dorsal-ventral, and medial-lateral location of each piece of dissected tissue was recorded by notes and photographs. For GW15 samples obtained from NIH Neuro-BioBank, annotation for cortical lobes were provided by NeuroBioBank staff. For more specific annotation of cortical areas, the GW15 samples were compared with the GW15 Reference Atlas (https://atlas.brain-map.org/atlas?atlas=138322603), and the GW20 and GW22 samples were compared with the GW21 Reference Atlas (https://atlas.brain-map.org/atlas?atlas=3). The relative anterior-posterior locations of tissue sections were matched with the Reference Atlas and cytoarchitectural features like early sulci were used as reference landmarks. The transition between premotor (PMC) and primary motor cortices (M1) lacks clear definition in mid-gestation, and therefore some tissues analyzed were annotated as “PMC/M1”. In contrast, the calcarine sulcus was well-defined morphologically in GW15-22, enabling identification of primary (V1) and secondary visual cortices (V2). For GW34 samples from NeuroBioBank, the specific Brodmann’s Area (BA) corresponding to the postnatal human cortex was determined by NeuroBioBank, and was provided when tissues were requested. Definition of specific BA is possible only for GW34 because by this gestational age the major sulci and gyri structures have formed, which is not the case in GW15-22. The cortical area annotation for all samples is summarized in **Supplementary Table 1**.

### Immunohistochemistry and microscopy

10µm-thick cryosections of tissue were placed on Superfrost slides (Fisher) for immunohistochemistry. The sample was first fixed in 4% PFA for 15 minutes at room temperature, followed by permeabilization with 0.5% Triton-X in PBS for 1hr and blocked with blocking solution of 10% donkey serum in PBS and 0.05% Triton-X (PBST) for 30 min. Primary antibodies diluted 1:500 in blocking solution were applied to the sections overnight at 4°C. Primary antibodies used were rat anti-CTIP2 (Abcam, ab18465), mouse anti-SATB2 (Abcam, ab9244), and rabbit anti-TBR1 (Abcam, ab31940). After washing with PBST for a minimum of 5 times, secondary antibodies and DAPI (1:2000) diluted in blocking solution were applied to the sections for 1-4 hrs at room temperature or overnight at 4°C. Secondary anti-bodies were: AlexaFluor 488, 555, or 647 -conjugated donkey antibodies (Invitrogen) used at 1:500 dilution. Finally, sections were washed with PBST for a minimum of 5 times before mounting with Vectashield Vibrance Antifade Mounting Medium. Images were captured by a Zeiss LSM 980 confocal microscope. Z-stack function was used to image a 10µm thickness, and tile-stitching function was used. Sample images were prepared in ImageJ software.

### MERFISH gene panel selection and probe construction

We designed a panel of 300 genes (**Supplementary Table 2**) that contained 39 canonical markers for major cell types in the human fetal cortex, including markers for excitatory (ENs) and inhibitory neurons (INs), intermediate progenitor cells (IPCs) and neural progenitor cells (NPCs); 32 were validated cortical layer markers in adult human cerebral cortex from a previous study^40^. 30 autism spectrum disorders (ASD)-associated genes were selected from the SFARI database^63^, and one putative regulator long non-coding RNAs (LncRNA) for each SFARI gene was included (30 total). The remaining genes on the panel were obtained from the top enriched cluster markers in published single-cell RNA SMART-seq data of mid-gestation human fetal cerebral cortex^7^. Single cell clustering and marker identification based on differential expression were performed in the original study^7^. The top 20 marker genes based on fold change for excitatory neuron clusters, and the top 5-10 marker genes for other clusters were manually curated. The genes that did not overlap with previous selected categories were included in the gene panel, resulting in a total of 300 genes. All genes within the panel were included for subsequent analysis. Merscope encoding probes for the 300 genes were constructed by Vizgen using commercial pipeline. Each of the 300 genes was assigned a unique binary barcode drawn from a 22-bit, Hamming-Distance-4, Hamming-Weight-4 encoding scheme (**Supplementary Table 3**)^9^. 15 extra barcodes were included as “blank” barcodes, which were not assigned to any genes to provide a measure of the false positive rate in MERFISH as previously described^9^.

### MERFISH imaging

MERFISH analysis was performed using the Vizgen Merscope system. Sample preparation was performed according to manufacturer’s instructions (MERSCOPE Fresh and Fixed Frozen Tissue Sample Preparation User Guide, Doc. number 91600002). Briefly, fresh frozen tissues were embedded in OCT and sectioned into 10µm-thick sections using a cryostat (Leica) and adhered to a Merscope slide (Vizgen, 1050001) placed in a 6-cm petri dish. For GW15-22 samples, slides were kept inside the cryostat maintained at -15°C for 30 minutes to allow the section to dry and firmly adhere to the glass slides before fixation; for GW34 samples, slides were kept inside the cryostat for 5 minutes, and then transferred to room temperature for 15 minutes before fixation. Slides were then fixed in 4% paraformaldehyde (PFA), and permeabilized by 70% ethanol overnight, with parafilm sealing the petri dish to prevent evaporation. Slides were then treated in the MERSCOPE Photo-bleacher instrument for autofluorescence quenching for 3 hours. Slides were stored overnight or up to one week before proceeding to the next step. The encoding probe mix was added directly on top of the tissue section for hybridization at 37°C for 36 hours in a humidified incubator. Post probe hybridization, sections were fixed again using formamide and embedded in gel. After gel embedding, tissue samples were cleared using a clearing mix solution supplemented with proteinase K for 24-48 hours at 37°C until no visible tissue was evident in the gel. After clearing, sections were stained for DAPI and PolyT and fixed with formamide prior to imaging. No additional cell boundary staining was used. Reagents used for these steps were included in Merscope sample prep kit (Vizgen 10400012).

The MERFISH imaging process was done according to the Merscope Instrument Preparation Guide (Doc. Number 91500001). Briefly, an imaging kit was thawed in a 37°C water bath for 45 minutes, activated and loaded into the Merscope instrument. The flow chamber was then assembled, fluidics were primed, and flow chamber filled with liquid. A low-resolution image for the DAPI and PolyT staining was taken under 10x magnification, and a region of interest (ROI) was manually drawn in the Merscope software, followed by automated image acquisition and fluidic control in the Merscope instrument. For each section, the ROI of up to 1cm^2^ area is imaged as 2,000-2,500 tiles at 40x magnification, and images were collected in the 750-nm, 650-nm, and 560-nm channels for the readout probes and in the 488-nm and 405-nm channels for PolyT and DAPI staining, respectively. 7 z-stacks were captured over a thickness of 10µm. After imaging, image processing and transcript decoding were performed using Merscope proprietary software. The transcript matrix with spatial coordinates, and the stitched tiled DAPI images acquired were transferred for subsequent processing and single-cell segmentation.

### Cell segmentation and MERFISH data processing

Automated segmentation was performed on the DAPI channel using a custom CellPose model^23,24^. The model was initialized with the CellPose “cyto” weights, then trained for 300 epochs with a learning rate of 0.1 and weight decay of 1e-4 using 145 manually segmented images for training and 3 for testing. All images, before training and running segmentation, were filtered using a difference of Gaussians filter, consisting of a positive Gaussian having zero standard deviation (all weight at zero) and a negative Gaussian with a standard deviation of 20 pixels. When running the CellPose segmentation, for testing and in the Vizgen pipeline, the cell diameter was set to 55 pixels (average from training data), the flow threshold was set to 0.5, the cell probability threshold was set to -3, and the minimum mask size was 500 pixels.

The Vizgen MERSCOPE output consisted of 7 planes evenly spaced across 10µm. Given the wide point spread function (PSF) and the small spacing, we used the maximum projection image over the first six planes as input to the CellPose segmentation algorithm, using the CellPose parameters and filtering as specified above. Because the nucleus staining (DAPI) was used, we dilated the masks by 10 pixels to approximate the cytoplasmic area of the cells. These processing steps were added to the Vizgen processing pipeline, the modified code is available here: https://github.com/carsen-stringer/vizgen-postprocessing. The Vizgen processing pipeline uses the CellPose masks to define ROIs and then assigns the RNA transcripts to each ROI to return a cell by gene matrix. In **Extended Data Fig. 1**, we quantified the segmentation accuracy using the average precision (AP) metric, which is the number of true positives divided by the total number of true positives, false negatives and false positives^24^. The true positives were defined as segmented ROIs that matched ground-truth ROIs at or above a defined intersection-over-union (IoU) threshold. The false negatives were the ground-truth ROIs that were missed, and the false positives were the predicted ROIs which did not match any ground-truth ROIs.

### MERFISH data quality control and integrated hierarchical clustering

After performing cell segmentation, for each experiment, we filtered out all cells with a total transcript count below the tenth percentile specific to that experiment. We then normalized each cell by its total transcript count using scanpy.pp.normalize_ total(). Transcript counts were then log-transformed and Z-score normalized using scanpy.pp.log1p() and scanpy.pp.scale(), respectively. Once preprocessed, we integrated the gene expression data from all samples. Subsequently, hierarchical clustering was performed on the integrated dataset to identify groups of cells with similar gene expression profiles. Initially, all cells were grouped into 8 H1 clusters at the first hierarchy (cell class). Following this, each H1 cluster was sub-clustered into 5 preliminary H2 clusters, resulting in 40 preliminary H2 clusters (cell type). Each H2 cluster was further subdivided into 5 preliminary H3 clusters, forming a total of 200 preliminary H3 clusters (cell subtypes). The clustering results at each hierarchy were obtained using sklearn.cluster.KMeans(). Marker genes were then identified using scanpy.tl.rank_gene_groups() and scanpy.get.rank_genes_group_df(). For a given H1 cluster, genes with a log2 fold change of at least 0.25 compared to the cells in all other H1 clusters were denoted as marker genes for that cluster. Marker genes for each H2 or H3 cluster were identified using the log2 fold change between that cluster and the other four clusters belonging to the same H1 or H2 group, respectively.

Cluster annotations were conducted sequentially from H1-H3, based on marker gene list and spatial distribution of each cluster. The 8 H1 cell classes were annotated based on expression of canonical markers. Some preliminary H2 and H3 clusters were manually merged. Because endothelial cells (EC) were not of particular interest in our study and our gene panel has very limited relevant genes for ECs, all EC clusters were merged into one cluster. Similarly, astrocytes and oligodendrocyte (OPC) clusters were merged into 3 H2 clusters, and subsequently 5 H3 clusters. H3 clusters for intermediate progenitor cells (IPC) derived from each H2 IPC cluster were highly similar in spatial distribution and gene expression, and were therefore also merged. Finally, we noticed that a few clusters exhibited aberrant spatial distribution reflecting technical artifact during the imaging process. These clusters, easily recognizable with exclusive localization at the edge of certain tissue sections or surrounding bubbles within the tissue section, were the accidental result of tissue hydrogel detaching from the surface during imaging. We removed cells from these artifact clusters from all subsequent analysis.

After transcript count filtering, artifact cluster removal and supervised merging, we analyzed a total of 15,927,370 cells that were clustered into 8 H1 clusters, 33 H2 clusters, and 114 H3 clusters (**Supplementary Table 4**). H2 cell types were annotated to reflect their spatial distribution. For excitatory neuron (EN-IT, EN-ET, EN-Mig) clusters, their approximated layer enrichment at GW34 were denoted. For radial glia (RG) and IPC, their enriched laminar structures (such as ventricular zone (VZ), subventricular zone (SVZ), intermediate zone(IZ)) were denoted in cluster annotation. In addition, if a cluster is predominantly present only in GW15 samples, it was denoted with “early”, while a cluster predominantly present only in GW34 was denoted with “late”. H3 EN clusters were annotated similarly to reflect their layer enrichment as well as their areal distribution enrichment (A for anterior, P for posterior, T for temporal lobe, etc.).

### Cell label transfer analysis

Reference-based cell type annotation was conducted using SingleR^64^ (v.1.8.1) to transfer cell type labels from the reference data to the testing data with the following parameters: “genes”: “de”; “de.method”: “classic”; “fine.tune”: TRUE; “prune”: TRUE. The three separate analyses are summarized below: (1) For **Fig. 1f** and **Extended Data Fig. 2b**, the fetal human cortex scRNA-seq data^7^ was used as a reference to transfer labels to our MERFISH data. In the reference, we excluded “MGE-” (cells from dorsal cortical specimens) and “unknown” clusters and aggregated all “nIN” subclusters into one and denoted it as “IN”. (2) For **Extended Data Fig. 4c**, cells from EN clusters in the adult human cortex scRNA-seq data^37^ were extracted and served as a reference to transfer cell type labels to EN-ET and EN-IT cells in our MERFISH data. For simplicity, the 56 original excitatory neuron clusters in the adult human cortex dataset were merged into 14 groups based on their shared layer and molecular marker identities. (3) For **Extended Data Fig. 10e**, cells from the EN clusters in our snRNA-seq data were employed as a reference to transfer labels to EN cells/spots in our MERFISH and Visium data separately. The pruned labels returned from SingleR were used to annotate the target cells. Correspondences between raw and transferred cell types in the target data were visualized through heatmaps.

### Quantitative spatial analysis of MERFISH data

For most tissue sections analyzed by MERFISH, one or two fan-shaped regions were manually drawn for quantitative spatial analysis. The fan-shaped regions were selected on the basis of relative geometrical uniformity, avoiding anatomical structures such as sulci, which may complicate location quantification. Areas with tissue section tears and bubbles were also avoided. For each tissue section analyzed by MERFISH, spatial graphs of H1 cell class distribution were generated. Based on these graphs, each fan-shaped region is made of hand-traced vector lines marking the apical surface and basal surface, and connected by two straight lines using the Pen tool in Adobe Photoshop software. The basal borders were defined by the pial surface of the cortex. For GW15 and GW20, the apical borders were drawn at the ventricular surface. For GW22, the apical borders were drawn within the outer subventricular zone (oSVZ) as the VZ could not fit within the imaged area due to larger tissue size. For GW34, the apical surfaces were drawn at the approximated transition between Layer 6b and the white matter.

Within each fan-shaped region, laminar structures of the developing cerebral cortex were further defined manually with respect to the distribution and morphology of cell classes. The cortical plate (CP) was defined as the condensed layer of excitatory neurons (EN-IT and EN-ET) near the pial surface. The thin layer above the CP to the pial surface was defined as the marginal zone (MZ). The subplate (SP) was defined as the layer containing only EN-ET, below the CP, and with lower cell density than the CP. The intermediate zone (IZ) was defined as the relatively cell-sparse layer predominantly composed of EN-Mig below the SP. The oSVZ was defined as the region composed of a mixture of EN-Mig, IPC, and RG below the IZ. The VZ was defined as the condensed layer of RG showing clear ventricular morphology at the apical surface. The thin layer above the VZ composed of high density of IPC was defined as the inner subventricular zone (iSVZ). The boundaries between each neighboring laminar structures were manually traced using the Pen tool in Photoshop, establishing an annotated vector mask for each fan-shaped region.

The vector mask was then imported into the computational analysis object by overlapping the spatial coordinates of cells with the vectors. Therefore, each cell within the fan-shaped region was also assigned with its laminar structure (VZ, iSVZ, oSVZ, IZ, SP, CP, and MZ). For each annotated region, three images were exported from Photoshop: (1) a mask containing the boundary of the fan-shaped region along with the lines delineating the boundaries of the annotated structures; (2) a mask containing only the boundary of the fan-shaped region; (3) a mask containing the boundary of the fan-shaped region, where the lines marking the apical surface and basal surface were replaced by straight lines. The initial objective was to determine the coordinates of the apical and basal surfaces. Using image (3), we applied cv2.goodFeaturesToTrack() to identify the four corners of the fan shape. Using the orientation of the annotated region (up, down, left, right), corner coordinates were assigned to their respective ends of the apical and basal surfaces. We calculated the equations of the lines connecting the corners of the apical and basal surfaces, and the point at which the lines intersected (O) was determined. To identify all points comprising the apical and basal surfaces, we first set all pixels equal to 0 in the mask of image (1) that were in a 3-pixel radius of each of the corners. We then used cv2.connectedComponents() to categorize each of the four lines in the mask into different components. The component IDs for the apical and basal surfaces were then determined based on the previously calculated corner locations. To extract the cells within a specific layer, we first subtracted image (2) from image (1) to eliminate the boundaries of the fan-shaped region, including the apical and basal surfaces. With only the layer-defining boundaries remaining, we applied cv2.connectedComponents() to assign each boundary to a different component. The component IDs corresponding to each of the layer boundaries were determined using the orientation of the fan-shaped region. To calculate the mask boundary for a given layer, we used cv2.line() to draw lines between the corresponding endpoints on the top and bottom boundaries for that layer. We then created the mask by applying cv2.binary_fill_holes() to the boundary. Using the coordinates of the cells located within the annotated region, we then extracted all cells with coordinates contained within the layer-specific mask.

For each cell within the annotated fan-shaped region, its “Relative Height (RH)” is calculated to represent its laminar location from the apical to basal surfaces. The RH calculation normalizes for the tilting of the fan-shape created by the asymmetrical morphology of the human cerebral cortex. Similarly, for a cell within the CP of an annotated fan-shaped region, its “Cortical Depth (CD)” is calculated to quantify its relative laminar location within the CP. To calculate the RH for a given cell, we utilized cv2.line() to draw a line from point O to the cell’s location at point C. Using the mask containing this line, we identified intersection points with the apical and basal surfaces (designated as C_3_ and C_1_, respectively). The Euclidean distances between C_3_ and C_1_, and between C and C_1_ were then computed. The relative height of the cell was then calculated as the ratio of these distances. To calculate CD, we first extracted all cells located within the cortical plate of the fan-shaped region. Then, for each cell located within the cortical plate, we used cv2.line() to draw a line connecting O and the location of the cell, C. Using the mask containing this line, we identified the points where the line intersected with the top and bottom boundaries of the cortical plate (denoted by C_3_ and C_2_, respectively). We then calculated the Euclidean distances between C_3_ and C, and between C and C_2_. The ratio of these distances defined the CD of the cell. For a given annotated region, the RH violin plot was constructed using the H2 cell type annotations for all cells within that region. The CD violin plot was created using the H3 cell type annotations for all EN-IT, EN-ET, and EN-Mig cells within the cortical plate of that region. For any clusters with fewer than 50 cells within that region, we represented the RH or CD distribution with individual cell dots instead of a violin plot.

Cortical layers within the CP of an annotated region were defined computationally by the distribution of CD values for specific H3 EN subtypes. While the laminar location of most H3 EN subtypes were consistent across gestational ages, some clusters showed varying abundance with gestational age, preventing us from using the same set of H3 EN subtypes for the definition of cortical layers in samples from all GWs analyzed. For GW15, EN-IT-L4-1, EN-ET-L5-1, EN-IT-L6-1 were used to defined Layer 3/4, Layer 5, and Layer 6 respectively. For GW20, EN-IT-L2/3-A1, EN-IT-L4-1, EN-ET-L5-1, EN-IT-L6-1 were used to defined Layer 2/3, Layer 4, Layer 5, and Layer 6 respectively. For GW22, EN-L2-1, EN-IT-L3-A, EN-IT-L4-1, EN-ET-L5-1, EN-IT-L6-1 were used to define Layer 2, Layer 3, Layer 4, Layer 5, and Layer 6 respectively. For GW34, EN-L2-4, EN-IT-L3-late, EN-IT-L4-late, EN-ET-L5-1, EN-IT-L6-late were used to define Layer 2, Layer 3, Layer 4, Layer 5, and Layer 6 respectively. The borders between adjacent layers, for example, between Layer 5 and Layer 6, were calculated as the mean CD value between the lower 25% of the Layer 5 cells and the upper 75% of the Layer 6 cells. Therefore, the CD value for each cell within the CP was used to assign a cortical layer identity for that cell.

### Spatio-temporal expression patterns of genes

For each gestational week and cortical area, we isolated all cells within each annotated layer using their coordinates and the corresponding layer-defining masks. Cells within the CP were assigned to specific cortical layers based on their CD values. Subsequently, for each gene, we calculated the mean expression across all cells within each annotated layer for a given gestational age. We then standardize the expression for that gene to the [0, 1] range across all layers and gestational weeks. The results were visualized in the summary expression heatmaps (**Source Data Fig. 3**). To construct z-score expression spatial graphs for a given cortical area, the normalized expression values for all cells within the area were extracted and gene expression values were Z-score normalized using scanpy.pp.scale(). Gene expression heatmaps were then plotted using scanpy.pl.embedding().

### Cell number proportion and enrichment quantifications

The two panels at the top of **Extended Data Fig. 5c** illustrate (i) the proportion of cells in an H3 cell type for each cortical area (PFC, PMC/M1, Par, Temp, and Occi) and (ii) the cell number of an H3 cell type for each gestational week (GW 15, 20, 22, and 34). For the cortical area proportion plot in (i), the proportion of cells for each cortical area (CA) was calculated as the total number of cells for each CA for the H3 cell type divided by the total number of cells for that H3 cell type across all the CAs. 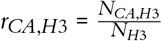 where *N*_*CA,H*3_ is the number of cells for a CA at H3 and *N*_*H*3_ is the total number of cells at H3. For the number of cells plot in (ii), the number of cells for each gestational week at each H3 subcluster is shown.

The results of spatial-temporal enrichment analysis of EN subtypes (EN-ET, EN-IT, and EN-Mig) in the bottom panel of **Extended Data Fig. 5c** and RG, IPC, and IN subtypes in **Extended Data Fig. 5d** were done through the normalization process described below. The total number of cells for each sample-region (SR) pair at H1 and H3 were counted separately for each gestational week. The number of cells of SR pairs at each H3 cell type is then divided by the number of cells of SR pairs at the corresponding H1 cell type. 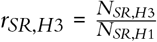 where *N*_*SR,H*1_ is the number of cells of a SR pair at H1 and *N*_*SR,H*3_ is the number of cells of a SR pair at H3. The ratio above is then divided by the maximum number of cells of SR pairs at H3. 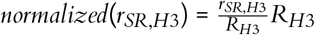 is the maximum of *r*_*SR,H*3_ among all the SR pairs at that H3 cell type.

For **Extended Data Fig. 1k**, the proportions of all H2 cell types in each SR pair was calculated. The number of cells of SR pairs at each H2 cell type is divided by the total number of cells in that SR. 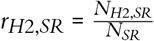 where *N*_*H*2,*SR*_ is the number of cells of an SR pair at H2 and *N*_*SR*_ is the total number of cells of an SR pair. **Extended Data Fig. 3e** and **Fig. 5c** were calculated similarly.

For UMAP of EN subtype composition (**Fig. 3f**), all EN-ET and EN-IT cells from each annotated cortical region were extracted. The relative proportion of the EN-ET and EN-IT H3 subtypes was calculated for each cortical area. We computed the UMAP projection of the matrix of relative EN H3 cell type proportions, providing a 2D representation for each cortical area based on EN subtype composition.

### Differential gene expression analysis between MERFISH clusters

The results of differentially expressed gene (DEG) analysis between clusters are visualized in **Fig. 3g** and **Extended Data Fig. 9b**. (1) In **Fig. 3g**, five pairs of anteriorly-enriched and posteriorly-enriched neuronal subtype H3 clusters were compared based on their DEGs: EN-IT-L2/3-A2 vs. EN-IT-L3-P, EN-IT-L4-A vs. EN-IT-L4-late, EN-IT-L4/5-1 vs. EN-IT-L5/6-P, EN-ET-L6-A vs. EN-ET-L6-P, and EN-ET-SP-A vs. EN-ET-SP-P1. (2) In **Extended Data Fig. 9b**, three pairs of H1 cell types (EN-IT, EN-ET, and EN-Mig) in V1 and V2 regions in the sample UMB1367-O1 were compared based on their DEG. The DEGs were detected through the “scanpy.tl.rank_genes_groups()” function with the default t-test from the Scanpy_1.7.2 in Python version 3.10. The top 10 and bottom 10 genes were selected separately based on their log fold change, and the log_10_(p-value) is used to break the ties. The top 10 genes represent the most upregulated genes in the DEG analysis, while the bottom 10 genes represent the most downregulated genes. In the bubble plots, the mean expressions for each gene across all the cells are calculated and represented by the spot color. The percentage of expressed cells is calculated by dividing the number of cells expressed for each gene by the total number of cells expressed across all the genes, represented by the spot size.

### Identification of area-enriched genes

To identify area-enriched genes in **Fig. 3i**, we analyzed EN-ET and EN-IT cells from GW20 and 22 within the annotated fan-shaped regions. Cells in PFC and PMC/M1 were grouped as anterior regions and cells in Par and V2 were grouped as posterior regions. We identified anteriorly and posteriorly enriched genes by t-test through the “scanpy.tl.rank_gene_groups” function. The top 10 genes with the highest anterior enrichment and posterior enrichment were selected separately based on their p-values. Additionally, the top 5 genes with the most enrichment in Temp exclusively were detected using the same methods.

### Clusteringby individual gestationalageandscSHCsub-clustering

To compare neuronal subtypes at different gestational ages, we employed an alternative clustering strategy in **Fig. 4 & 5**, where only cells from the same gestational age were clustered together. Using the preprocessed data, all cells belonging to samples from the same gestational week were combined. Subsequently, hierarchical clustering was conducted for each gestational week independently. Analogous to the method applied for the integrated analysis, for each time point, we first clustered the cells into 8 H1 and 40 preliminary H2 clusters using sklearn.cluster.KMeans(). H1 and H2 clusters were annotated following the same strategy as the integrated clustering described above.

Single-cell significant hierarchical clustering (scSHC) pipeline (v.0.1.0) was used to unbiasedly determine the number of significant H3 subclusters^42^. ScSHC adopts a hypothesis testing approach, recursively dividing the cells into two groups, and testing the statistical significance of each split with an adjusted threshold to control the family-wised error rate (FWER). It defines Ward linkage as test statistics and employs parametric bootstrap to estimate its null distribution. If the p-value at a given node is larger than the adjusted threshold, the two subsequent branches are merged. For each gestational age, we randomly sampled 500,000 cells and ran scSHC within each H2 cluster. To ensure model robustness, we increased the number of cells used for null distribution estimation from 1,000 in the original method to one-third of the cluster size. Furthermore, to avoid overly small subclusters, if one subcluster comprised fewer than 10% of the cluster size, we stopped further splitting. For each run of scSHC, “alpha”, the FWER, was fixed at 5e-4; “num_features”, the number of genes used as features, was set to 300 (all MERFISH panel genes); “num_PCs”, the number of top principal components retained for gene expression dimension reduction, was set to be 30. The H3 subclusters generated by scSHC are summarized in **Supplementary Table 6**.

### ScSHCsubclustercorrespondenceanalysisacrossgestational ages

To evaluate the transcriptomic correspondence of EN-ET and EN-IT scSHC subclusters across different ages, we applied XG-Boost^43^, a distributed gradient-boosted decision tree-based classification method. For each adjacent pair of gestational stages, we trained an XGBoost classifier to learn cluster labels from gene expression data within the earlier stage. Consequently, we used the classifier to classify cells from the later stage. The correspondence between the original clusters and the classifier-assigned labels in the later gestational age dataset were utilized to map clusters between ages. The general classification workflow is described below and was applied to each adjacent gestational age pair.

Let *X*_*E*_ represent the earlier stage dataset grouped into *N*_*E*_ clusters, and *X*_*L*_ denote the later stage dataset grouped into *N*_*L*_ clusters. Two gene expression matrices are normalized and log-transformed. The main steps are as follows:

1. We trained a multi-class XGBoost classifier on *X*_*E*_ using all 300 genes as features. For clusters with fewer than 15,000 cells, we upsampled by bootstrapping to 15,000 cells to make the dataset more balanced. XGBoost classifier parameters were set to the following values: “objective”: “multi:softmax”; “eval_metric”: “mlogloss”; “num_class”: *N*_*E*_; “eta”: 0.2; “max_depth”: 20; “subsample”: 0.6; “num_boost_round”: 1000.
2. For validation, we randomly sampled 80% of cells in each cluster of *X*_*E*_ to train the classifier and predicted the cluster labels for the remaining 20% of cells. For each testing cell *k*, the classifier returns a cluster assignment probability vector 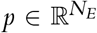, and the final cluster label is assigned as *cluster*(*k*) = *argmax*_*i*_*p*_*i*_. For EN-ET cells, the classifiers achieved over 90% accuracy (96.17%, 90.40%, 92.98%); while for EN-IT cells, the accuracy was more than 83% (93.65%, 88.96%, 83.09%).
3. For prediction, we re-trained the classifier on 100% of cells in *X*_*E*_ and applied it to *X*_*L*_ to obtain the predicted label for each cell. The overall correspondence flows between clusters across gestational ages were visualized through Sankey diagrams (**Fig. 4a, b, Extended Data Fig. 7a**).

To identify genes exhibiting up-or down-regulation across gestational ages within one layer-based EN group, we calculated the expressed cell fraction and mean expression for each gene at each time point. We then fit gene-specific linear regression models for each of these two metrics, using time points as input, where the time points were encoded as 1, 2, 3, and 4 for GW15, 20, 22, and 34 respectively. Genes were ranked according to the regression coefficient, with the top 10 positive ones indicating up-regulated genes, and the top 10 negative ones indicating down-regulated genes. The analyses were repeated for each EN group, and the expression patterns of identified genes were visualized in dot plots (**Fig. 4e**). Cluster-specific expressions of identified genes were shown in **Extended Data Fig. 7b**.

Moreover, to pinpoint genes that drive subcluster specification, we first created a pseudobulk for each cluster within the same EN group and gestational age by averaging cell expression. Subsequently, if there were more than one cluster within the same EN group and gestational age pair, we computed the pseudobulk expression variance for each gene. Next, we selected genes with expression variances greater than 1 in at least one EN group and gestational age pair and showed their expression variances using a heatmap (**Fig. 4g**). Additionally, we picked the top 10 genes with the highest expression variance within each EN group and gestational age pair and visualized their expression patterns as dot plots (**Fig. 4h, i**).

### Visium spatial transcriptomics analysis

Visium Spatial Gene Expression Analysis (10x Genomics) was performed according to manufacturer’s user guide and demonstrated protocol (Visium CytAssist Spatial Gene Expression for Fresh Frozen – Methanol Fixation, H&E Staining, Imaging & Destaining, CG000614, Rev A; Visium CytAssist Spatial Gene Expression Reagent Kits User Guide, CG000495, Rev D). Briefly, 10µm-thick cryosections adjacent to the sections used for MERFISH analysis were adhered to Superfrost slides.

Two samples were used for Visium analysis, FB080-O1, GW20 occipital cortex sample containing the V1 and V2, and FB121-F1, GW20 prefrontal cortex. FB080-O1 is coded as “A1” and FB121-F1 is coded as “D1” in the data processing and analysis. The tissues were fixed in chilled methanol at -20°C for 30 minutes and stained with Hematoxylin and Eosin. The slides were mounted with 85% glycerol in water. H&E staining was imaged at 20x magnification using a Zeiss Axioscan 7 microscope. After imaging, the slides were placed in Visium Tissue Slide Cassette with 6.5 mm gasket and destained with 0.1N HCL for 15 min at 42°C. Probe hybridization was performed overnight at 50°C on a thermal cycler using the provided slide adaptor, followed by post hybridization buff. Probe ligation mix was then added into the slide gasket and incubated for 1 hour at 37°C, followed by post-ligation wash. RNA digestion and tissue removal were performed using the CytAssist instrument (10x Genomics) before probe extension and probe elution. Pre-amplification, SPRIselect, and library construction were performed according to the user’s guide. GEX post-library construction QC was performed using the Agilent 4200 TapeStation System.

Visium libraries were sequenced on a NovaSeq6000 according to 10x Genomics’ recommended parameters to the depth of 214 to 235 million PE150 reads. Demultiplexed reads were processed with Spaceranger-count with default parameters and manual alignment. Visium data analysis was conducted separately on two slices, A1 (FB080-O1) and D1(FB121-F1). For visualization and clustering, Seurat_5.0.1 and R version 4.3.1 (2023-06-16) were utilized on the BCH compute nodes. SCTransform normalization was performed with the “spatial” assay parameter. Optimal clusters were generated, and clusters with negligible cells were removed. Marker identification was performed using FindAllMarkers analysis from the Seurat package using Wilcoxon rank sum test with the following parameters: (1) Only return positive markers; (2) Only test genes that are detected in a minimum fraction of 1% of cells in either of the two populations. This is the default. Following these results, we filtered for markers grouped by clusters with an average log2 fold change greater than 1. We generated different visualizations using top genes from this analysis. R version 4.1.2 (2021-11-01) along with Seurat_4.1.1 were used for both exploratory analysis and final visualization on local machine. Spatial Feature Plots were generated to visualize the expression of genes of interest within the tissue sections.

### Nuclei isolation and single-nucleus RNA sequencing

100µm-thick cryosections consecutive to the ones collected for MERFISH and Visium analyses were collected in a 1.5mL tube. Nuclei were isolated from the cryosection as previously described with minor modification^65^. Briefly, one cryosection per sample was resuspended in 1mL homogenization buffer with additives (10mM Tris Buffer pH 8.0, 250mM Sucrose, 25mM KCl, 5mM MgCl2, 0.1% Triton X-100, 0.1mM DTT, 1X cOmplete™, Mini, EDTA-free Protease Inhibitor Cocktail (Roche 11836170001), 25ul Protector RNAse inhibitor (Roche 3335399001, 0.2U/µl)) and transferred to a 7mL douncer and dounced 10 times with a “tight” pestle. Homogenized nuclei were spun for 10min at 900g at 4C, then washed once with Blocking Buffer (1X PBS pH 7.4, 1% BSA), spinning for 5min at 400g at 4C. All spins were done in a bucket centrifuge. Nuclei were resuspended in 300ul Blocking Buffer with Protector RNAse inhibitor (Roche 3335399001, 1U/ul) and Dapi (final concentration 1ug/ml) and passed through a 40µm filter. Dapi positive nuclei (13,500 nuclei per 10X reaction) were Fluorescence-Activated Nuclei Sorted (FANS) directly into Chromium Next GEM Single Cell 3*′* Reagent Kit v3.1 GEM master mix (Step 1.1) minus RT Enzyme C. RT Enzyme C was then added and reactions loaded into the Chromium Next GEM Chip and single-nuclei RNA-seq libraries generated according to the Chromium Next GEM Single Cell 3*′* Reagent Kit v3.1 manual with 11 PCR cycles for cDNA amplification and 11-12 PCR cycles used for final amplification of Gene Expression Libraries. 5 reactions were performed for FB080-O1 and 3 reactions were performed for FB121-F1 samples. In addition, 3 reactions were performed on FB080-O2, a consecutive tissue block posterior to FB080-O1, near the occipital pole of the brain. Single-nuclei RNA-seq libraries were sequenced on a NovaSeq6000 according to 10x Genomics’ recommended parameters to the depth of 909 PE150 million reads for FB080-O1, 541 million PE150 reads for FB080-O2, and 484 million PE150 reads for FB121-F1.

### SnRNAseq processing, clustering, and analysis

Demultiplexed reads were processed with cellranger-count with default parameters, resulting in 41,740 estimated cells for FB080-O1, 23,001 estimated cells for FB080-O2 and 26,161 estimated cells for FB121-F1. For visualization and clustering, Seurat_5.0.1 and R version 4.3.1 (2023-06-16) were employed on BCH Compute nodes, ensuring efficient clustering. Additionally, R version 4.1.2 (2021-11-01) along with Seurat_4.1.1 was utilized for generating figures on a local machine. Clustering optimization was achieved through iterative application of the FindClusters algorithm to attain optimal cluster resolution. Feature plots were constructed for genes of interest, and integration into cell-by-gene tool was verified. Contribution plots were generated to observe the contribution of cell types to the overall dataset. To find differentially expressed genes (DEGs) between V1-enriched (EN-IT-L4-V1) and V2-enriched (EN-IT-UL-2) clusters, we performed Wilcoxon rank sum test at log2FC > 0.5 and Bonferroni corrected p-value < 0.05. Only test genes that are detected in a minimum fraction of 25% of cells in either of the two populations are considered, filtering the genes that are infrequently expressed and only returning positive markers. To verify that we captured DEGs between V1-V2, we also identified DEGs between V1-V2 regions in Merscope which revealed overlap only between V1-upregulated genes (50%) or V2-upregulated genes (73%) but not vice versa (0%). For gene ontology (GO) analysis, we used the clusterProfiler package to perform gene ontology analyses in R (https://bioconductor.org/packages/release/bioc/html/motifmatchr.html). Specifically, DEGs upregulated in V1 or V2 were both tested for gene ontology enrichments using a background of all genes tested for differential expression (odds ratio > 2 and FDR < 0.01).

### ENVI-imputation from scRNAseq to MERSCOPE

We utilized environmental variational inference (ENVI) (v.0.1.0)^22^ on our MERFISH and snRNA-seq data to expand the MER-FISH gene panel. ENVI leverages a conditional variational autoencoder to integrate spatial and snRNA-seq data, which can simultaneously impute missing gene expression for genes that are not included in the spatial transcriptomics data and transfer spatial information onto the snRNA-seq data. We selected the top 1,000 highly variable genes from snRNA-seq, along with additional genes of interest for imputation: *ABI3BP, PDZRN4, FLRT2, TAFA2, NR1D1, IL1RAP, CCBE1, THSD7B, TRPC6, CHRM2, LUZP2, LRP1B, LRRTM4, HDAC9, FBXL7, DTNA, SYNDIG1, SDK1, LMO3, TRIQK, UNC13C, CNTNAP2, KCNIP4, PDZRN3, DLX6, DLX6-AS1, ADARB2, ERBB4, NRXN3, DLX2, ZNF536, PRKCA, THRB, TSHZ1, PBX3, MEIS2, CALB2, CDCA7L, SYNPR, SP8, CASZ1, FOXP4*.

We ran ENVI with the following key parameters: “k_nearest”: 100; “num_cov_genes”: 50; “num_layers”: 3; “num_neurons”: 1024; “latent_dim”: 512; “spatial_dist”: “pois”; “sc_dist”: “nb”; “cov_dist”: “OT”; “prior_dist”: “norm”; “spatial_coeff”: 1; “sc_coeff”: 1; “cov_coeff”: 1; “kl_coeff”: 0.3.

### Constellation plot analysis

The constellation plot was generated using the ENVI imputation results. We selected all RG, IPC, EN-Mig, EN-ET, and EN-IT cells from the primary (V1) and secondary visual cortices (V2) of FB080-O1c and the prefrontal cortex (PFC) of FB121-F1. Based on the H2 annotations, we combined all EN-ET cells from layers 5 and 6 into a single node (EN-ET L5/6) and did the same for layer 5 and 6 EN-IT cells (EN-IT L5/6). Other EN-ET and EN-IT cells were categorized into nodes according to their H2 annotations, while RG and IPC cells were classified by their H1 annotations. Cells were further subclassified based on the sample and annotated region in which they were located. The positions of the cell types on the constellation plot were determined using the median UMAP coordinates of all cells of each type. Node sizes were scaled according to the number of cells of each type. To establish edges between pairs of cell types, we first performed principal component analysis (PCA) on the ENVI gene expression matrix. Using the top 50 principal components, we then identified the 15 nearest neighbors for each cell by applying the Ball Tree algorithm. For any given pair of cell types, A and B, we added a connection from A to B if a cell in A had a nearest neighbor in B. We then calculated the total number of connections from A to B (*n*_*A*_*→*_*B*_), and the total number of connections from A to all types (*n*_*A*_). An edge was drawn between A and B on the constellation plot if both *n*_*A*_ *→*_*B*_/*n*_*A*_ and *n*_*B*_*→*_*A*_/*n*_*B*_ exceeded 0.02. The edge from A to B was colored according to the proportion of connections from A to B relative to the total connections from A (*n*_*A→B*_/*n*_*A*_).

### Cell-cell communication analysis

We employed the CellChat package (version 1.6.1) to decipher cell-cell communication networks using its extensive database of known human ligand-receptor interactions. To assess the communication probability, we followed the methodologies outlined in the original research paper by Jin et al^49^. This method applies the law of mass action to the average expression levels of ligands and receptors across annotated cell groups and evaluates their significance using the random permutation statistical test. Interactions with p-values below 0.05 are considered significant and those identified in fewer than 10 cells in each cell group are discarded to enhance the robustness of our findings. For this study, we analyzed all cell types available in our dataset but specifically focused on EN-IT-L4-V1, EN-IT-UL-1, EN-IT-UL-2, EN-IT-UL-3, EN-IT-UL-4, and EN-IT-UL-5 as receivers in the NRXN pathway. Visualization of the communication networks and pathways was performed using CellChat’s native plotting functions, which facilitated a comprehensive representation of the cellular interactions within our dataset.

### CELLxGENE web-based data browser

For the visualization and interactive exploration of our singlecell RNA sequencing and MERFISH data, we utilized in-house self-host CELLxGENE portal which was deployed with CELLxGENE (v1.1.0) and CELLxGENE gateway (v0.3.10)^66^. The processed Seurat object data for MERFISH data were converted into the .h5ad file format via the SeuratDisk package (v0.0.0.9020), which is compatible with the self-host CELLx-GENE portal. This setup enabled us to launch a locally hosted web-based interactive interface for real-time data analysis and exploration, facilitating UMAP visualization, analysis of cell type subpopulations, and gene expression querying.

### Statistics and reproducibility

Replicates and statistical tests are described in the figure legends and Methods. Sample sizes were not predetermined utilizing statistical methods. Tissue samples were not randomized, nor were the investigators blinded during collection as no subjective measurements were taken. Data for snRNA-seq, Visium and MERFISH were collected from all available samples and no randomization was necessary. To identify differentially expressed genes between clusters, a Wilcoxon rank sum tests were performed.

## Supporting information

Expression heatmaps of all genes analyzed

## Data availability

Raw sequencing data from Visium, processed Merfish data, and processed snRNAseq data will be deposited prior to publication. Raw data that support the findings of this study are available from the lead contact Dr. Christopher A. Walsh upon reasonable request.

## Code availability

Codes for Merscope Processing and Cellpose cell segmentation pipelines are available at: https://github.com/carsenstringer/vizgen-postprocessing. Code used for MERFISH data analysis in this manuscript is available at GitHub: https://github.com/ShunzhouJiang/Spatial-Single-cell-Analysis-Decodes-Cortical-Layer-and-Area-Specification

## Acknowledgement

We thank all members of our teams for discussion and help. We thank R. S. Hill at the Walsh Lab, R. Mathieu at the Boston Children’s Hospital (BCH) and Harvard Stem Cell Institute Flow Cytometry Research Facility, the Research Computing group at Harvard Medical School, the BCH Intellectual and Developmental Disabilities Research Center (IDDRC) Molecular Genetics Core, H. Cramer at BCH Cellular Imaging Core, Cell Function and Imaging Core at BCH for assistance. The analysis was performed with the computational resources provided by the Research Computing Group at BCH and Harvard Medical School, including High-Performance Computing Clusters Enkefalos 2 (E2), and the BioGrids scientific software made available for data analysis. We thank J. Maffei and S. Mitter at Vizgen Inc. for technical support with Merscope instrument. We thank Department of Pathology at Beth Israel Deaconess Medical Center at Harvard medical school, University of Maryland Brain and Tissue Biobank and the NIH NeuroBioBank for providing de-identified human samples with consent and approval from relevant IRB and ethics board. Illustrations in the figures were created in part with BioRender.com. X.Q. was supported by K99NS135123 from the NINDS, and Postdoctoral Fellowship from the Helen Hay Whitney Foundation and the Howard Hughes Medical Institute (HHMI). A.J.K. was supported by HHMI Jane Coffin Childs Postdoctoral Fellowship. D.D.S. was supported by K08 NS128074 from the NINDS. R.E.A. was supported T32 GM007748 from the NIGMS and Autism Speaks Post-doctoral Fellowship 13008. M.B.M. was supported by NIH Director’s New Innovator Award DP2 AG086138 and R01 AG082346. C.S. was funded by the HHMI at the Janelia Research Campus. M.L. was partly supported by R01HG013185 from the NHGRI. C.A.W. was supported by R01NS035129 and R01NS032457 from the NINDS, the Allen Discovery Center program, a Paul G. Allen Frontiers Group advised program of the Paul G. Allen Family Foundation, and John

Templeton Foundation research grant (#62587). C.A.W. is an Investigator of the Howard Hughes Medical Institute.

## Author information

These authors contributed equally: Xuyu Qian, Kyle Coleman, Shunzhou Jiang

## Authors and Affiliations

Division of Genetics and Genomics, Department of Pediatrics, Boston Children’s Hospital, Harvard Medical School, Boston, Massachusetts, USA.

Xuyu Qian, Andrea J Kriz, Jack H Marciano, Monica D. Manam, Emre Caglayan, Aoi Otani, Urmi Ghosh, Diane D. Shao, Rebecca E. Andersen, Jennifer E. Neil, Michael B. Miller & Christopher A. Walsh

Howard Hughes Medical Institute, Boston Children’s Hospital, Harvard Medical School, Boston, Massachusetts, USA. Xuyu Qian, Andrea J Kriz, Jack H Marciano, Monica D. Manam, Emre Caglayan, Aoi Otani, Urmi Ghosh, Diane D. Shao, Rebecca E. Andersen, Jennifer E. Neil & Christopher A. Walsh

Statistical Center for Single-Cell and Spatial Genomics, Department of Biostatistics, Epidemiology and Informatics, Perelman School of Medicine, University of Pennsylvania, Philadelphia, Pennsylvania, USA.

Kyle Coleman, Shunzhou Jiang, Chunyu Luo & Mingyao Li

Research Computing, Department of Information Technology, Boston Children’s Hospital, Boston, Massachusetts, USA. Chunhui Cai & Liang Sun

Department of Neurology, Boston Children’s Hospital, Boston, Massachusetts, USA.

Diane D. Shao & Christopher A. Walsh

Broad Institute of MIT and Harvard, Cambridge, Massachusetts, USA.

Rebecca E. Andersen, Diane D. Shao, Michael B. Miller & Christopher A. Walsh

University of Maryland Brain and Tissue Bank, Department of Pediatrics, University of Maryland School of Medicine, Baltimore, Maryland, USA.

Robert Johnson & Alexandra LeFevre

Department of Pathology, Beth Israel Deaconess Medical Center, Boston, Massachusetts, USA.

Jonathan L. Hecht

Division of Neuropathology, Department of Pathology, Brigham and Women’s Hospital, Harvard Medical School, Boston, Massachusetts, USA

Michael B. Miller

Department of Neurology, Brigham and Women’s Hospital, Boston, Massachusetts, USA.

Michael B. Miller

Janelia Research Campus, Howard Hughes Medical Institute, Ashburn, Virginia, USA

Carsen Stringer

Department of Pathology and Laboratory Medicine, Perelman School of Medicine, University of Pennsylvania, Philadelphia, Pennsylvania, USA.

Mingyao Li

Departments of Pediatrics and Neurology, Harvard Medical School, Boston, Massachusetts, USA.

Christopher A. Walsh

## Contributions

M.L. and C.A.W. supervised the study. X.Q., M.L. and C.A.W. conceptualized the study and wrote the manuscript. X.Q., A.J.K., J.H.M., A.O. and U.G. performed the experiments. X.Q., K.C., S.J., A.J.K., C.L., C.C., M.D.M. and E.C. analyzed the data. J.E.N., R.J., A.L., J.L.H. and M.B.M. contributed to sample collection, distribution, dissection, or management. C.C. and L.S. constructed data web browser. D.D.S. and R.E.A. gave relevant advice. C.S. developed data processing and cell segmentation pipeline.

