## Extended Data Fig. 1-10 for "Spatial Single-cell Analysis Decodes Cortical Layer and Area Specification"

**Supplementary Information**

**Supplementary Table 1:**
Meta information table for the human fetal cortex samples used in the study.

**Supplementary Table 2:**
MERFISH gene panel list with annotation for selection criterion.

**Supplementary Table 3:**
Merscope encoding probe design for the gene panel.

**Supplementary Table 4:**
Cluster annotations for integrated clustering.

**Supplementary Table 5:**
Differentially expressed genes between anteriorly and posteriorly enriched EN subtypes.

**Supplementary Table 6:**
Annotation for clustering by individual gestational age and scSHC.

**Supplementary Table 7:**
Differentially expressed genes between V1 and V2 EN subtypes and their overlaps from MERFISH analysis.

**Supplementary Table 8:**
Connection fraction values in constellation plot.

**Supplementary Table 9:**
Differentially expressed genes between EN-IT-L4-V1 and EN-IT-UL-2 clusters from snRNAseq, used for GO analysis.

**Source Data Fig. 3:**
Expression pattern heatmap for all 300 genes in the MERFISH panel.

### Extended Data Fig. 1: Single cell segmentation and quality control analyses of MERFISH data.

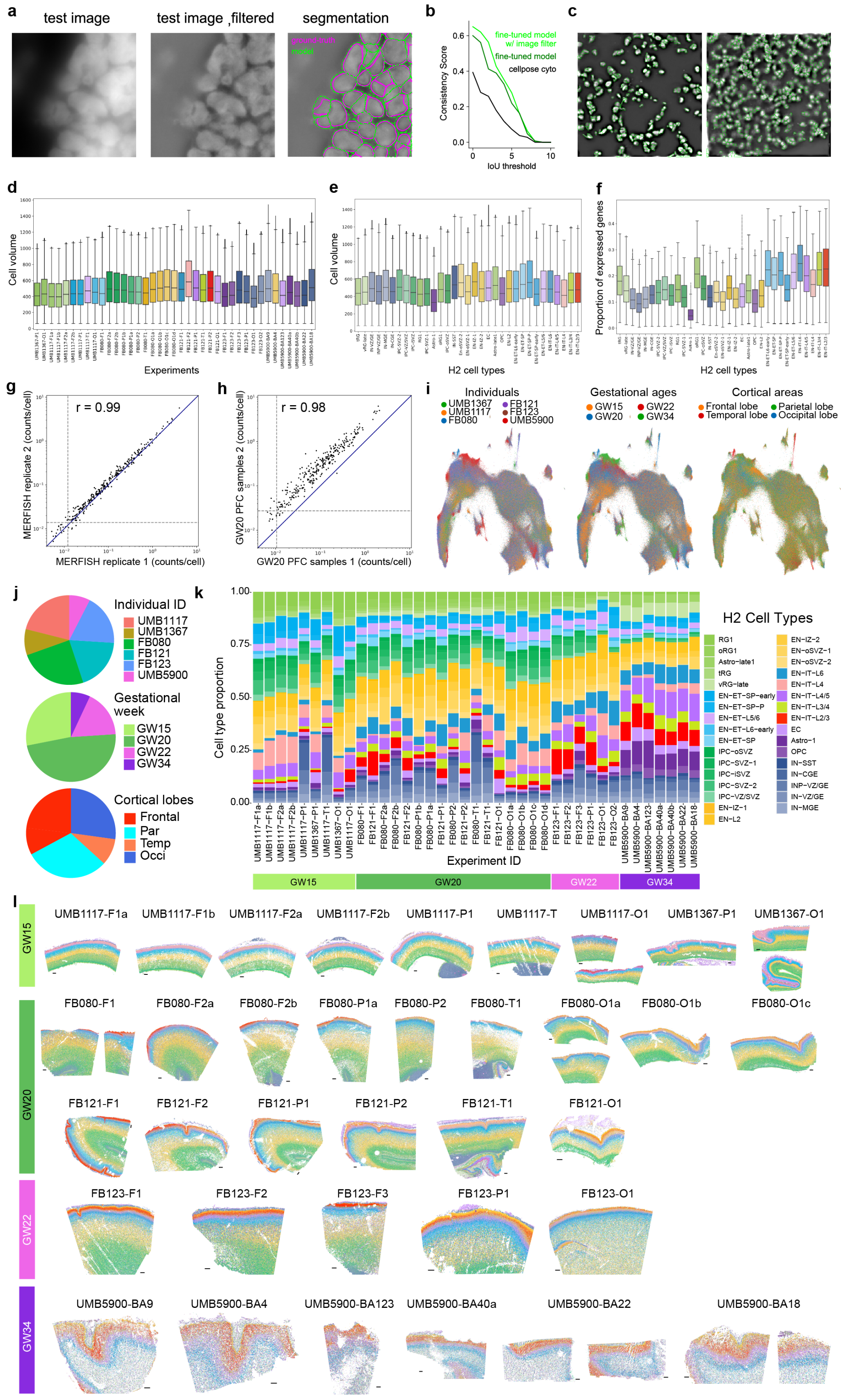

**Extended Data Fig. 1: Single cell segmentation and quality control analyses of MERFISH data. a,** Example test image (left), test image filtered with difference of Gaussians filter (mid), and filtered test image overlaid with ground-truth manual segmentation (magenta) and segmentation from the fine-tuned CellPose 2.0 model trained using filtered images (green)(right). **b,** Consistency of the segmentation algorithm comparing with human manual labeling as the ground truth. The consistency is quantified using the average precision (AP) metric, at varying intersection-over-union (IoU) thresholds. The AP is defined as the number of true positives divided by the total number of true positives, false negatives, and false positives. The fine-tuned model trained with filtered images outperformed the fine-tuned model trained on the raw images and the built-in "CellPose cyto" model. **c,** Example segmentations of fields of view from two experiments. **d, e,** Box graphs show that the distribution of volumes of individual cells after CellPose segmentation were consistent across experiments (**d**) and across cell types (**e**). **f,** Box graph showing the distribution of proportion of expressed genes among the 300 panel genes in cells from different H2 cell types. Our MERFISH panel design placed an emphasis on excitatory neurons, leading to more genes detected in EN cell types than others. **g, h,** Scatterplots show the average count per cell of individual genes is consistent between two replicate experiments from the same sample (**g**), and between two independent samples with matching gestational age and cortical area (**h**). The blue solid line indicates equality. The grey dashed lines indicate the average counts per cell of the blank barcodes, which provides an estimate of the false-positive rate. The Pearson correlation coefficient is $r = 0.99$ (**g**) and 0.98 (**h**). **i,** UMAP plots showing cells from different individuals (left), gestational ages (mid), and cortical lobes (right) integrated evenly into a combined dataset. **j,** Pie charts showing the proportion contributions of number of cells from different individuals (top), gestational ages (mid) and cortical lobes (bottom). **k,** Histogram showing the compositions of H2 cell types across all MERFISH experiments. **l,** MERFISH spatial maps colored by H2 cell types. The experiment names are shown above each graph, which includes the sample ID, and a suffix indicating the cortical lobe. F, frontal; P, parietal; O, occipital; T, temporal; BA, Brodmann's Area (only identifiable at GW34).

### Extended Data Fig. 2: Molecular characterization of H3 subtypes and spatial quantification of H2 cell types.

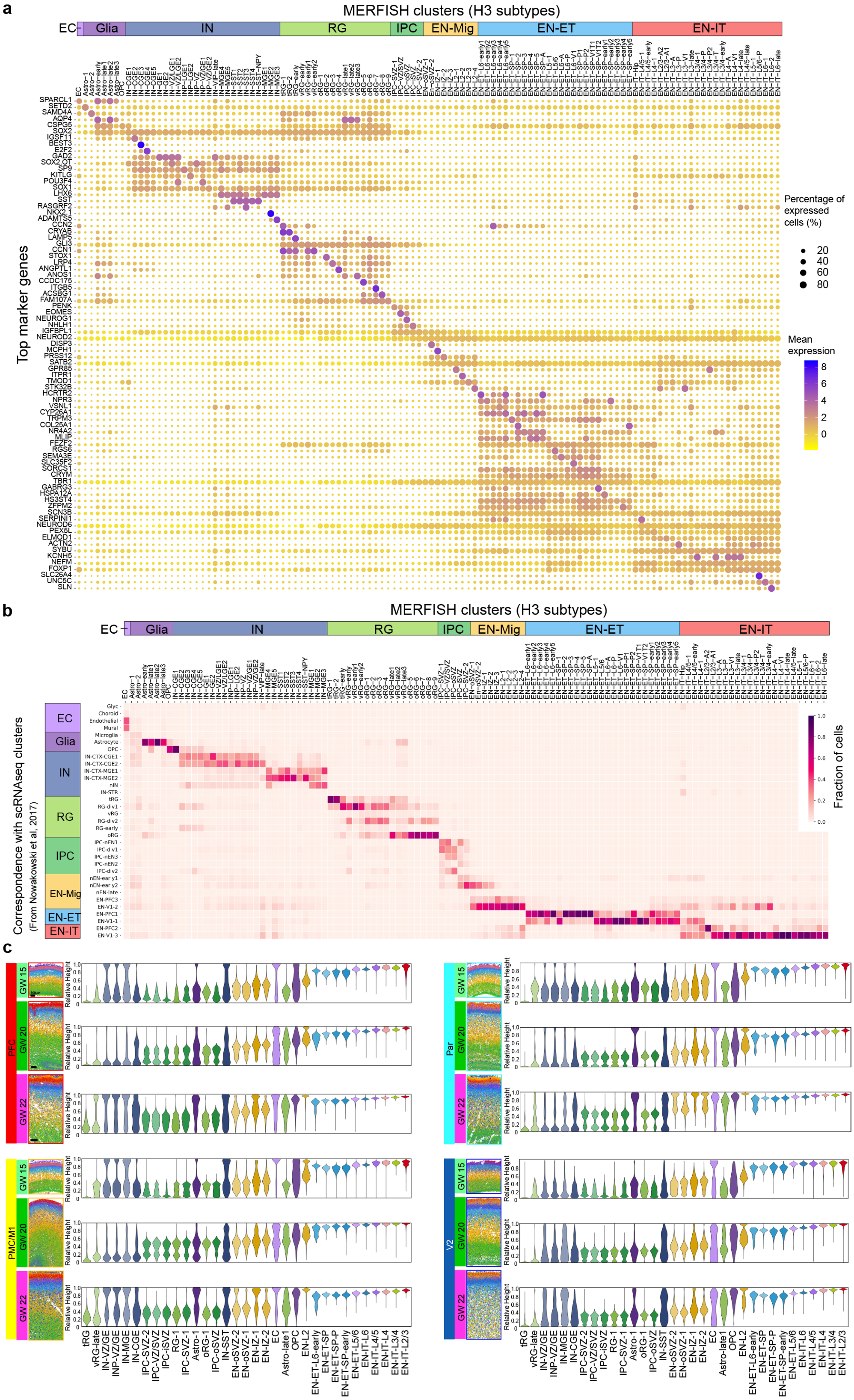

**Extended Data Fig. 2: Molecular characterization of H3 subtypes and spatial quantification of H2 cell types. a,** Dot plot showing the expression of marker genes for H3 subtypes. **b,** Cell type correspondence heatmap showing the fraction of cells from MERFISH H3 clusters associated to clusters from published mid-gestation scRNA-seq dataset from Nowakowski et al. 2017[1]. **c,** Spatial maps and violin plots showing the laminar distribution of H2 cell types from the apical to basal surface quantified by the RH across major cortical areas in the second trimester. The width of the violin for each cell type is normalized to the maximum value. Image scales are the same for different areas from the same GW. Scale bars, 500 μm.

### Extended Data Fig. 3: High concentration of INs in the dorsal VZ at mid-gestation.

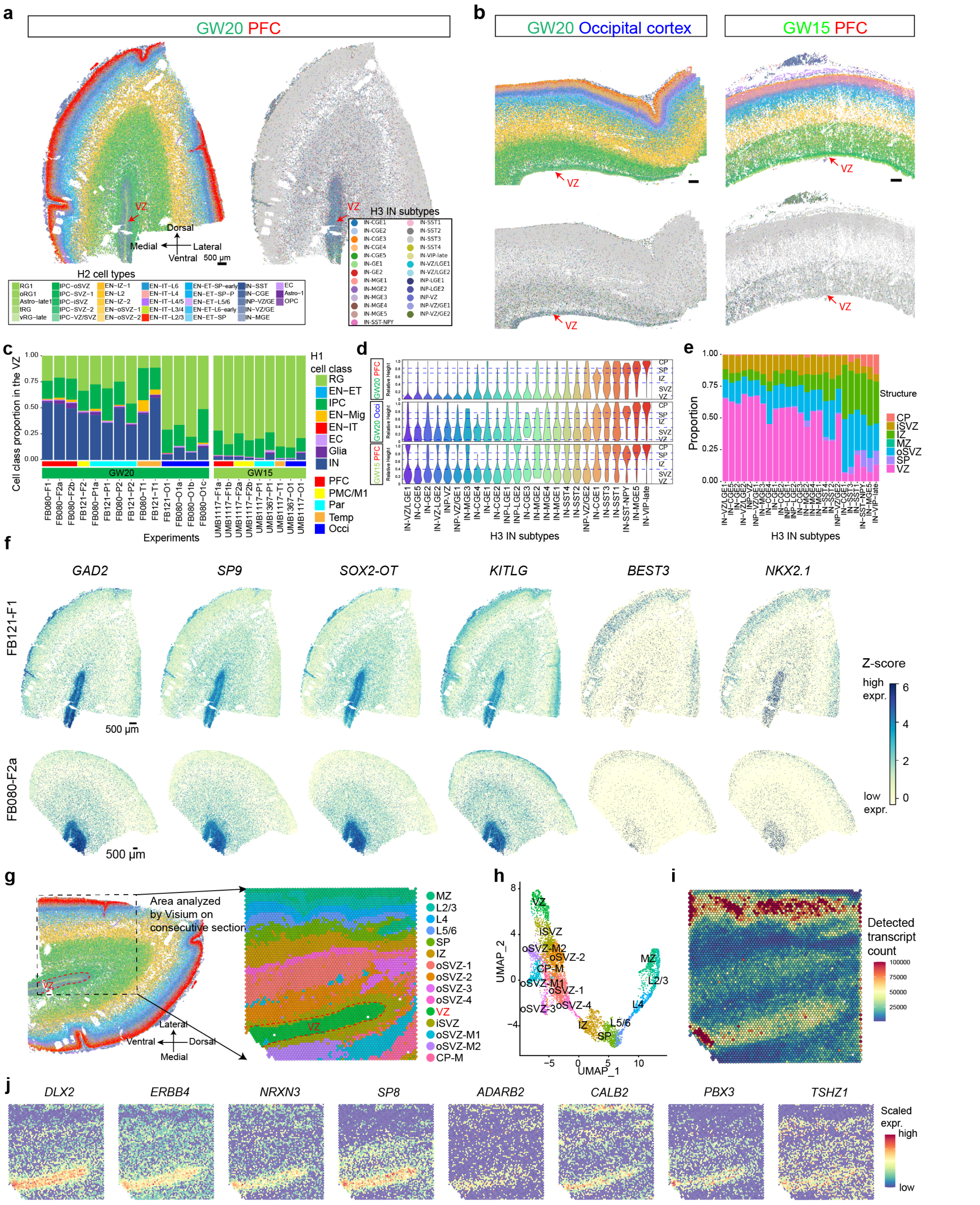

**Extended Data Fig. 3: High concentration of INs in the dorsal VZ at mid-gestation. a,** Spatial map shows inhibitory neurons (IN) are highly concentrated in the ventricular zone (VZ) of the prefrontal cortex (PFC) at GW20. Scale bars, 500 μm. **b,** INs are not enriched in the VZ of the occipital cortex at GW20, and are also absent in the VZ of all cortical areas at GW15. **c,** Histograms quantifying the cell class proportions within the VZ areas across experiments show INs outnumber RG and IPCs in all cortical areas except occipital cortex at GW20, but are largely absent at GW15. Due to the size restriction of MERFISH imaging area, the VZ was not systematically analyzed for GW22 and GW34 samples. **d, e,** Relative height (RH) violin plots and histograms for the distribution of H3 IN subtypes show the VZ localization is subtype dependent. **f,** Z-score spatial plots show IN markers are highly expressed in the VZ, but MGE marker *NKX2.1* and CGE marker *BEST3* were expressed sparsely. Scale bars, 500 μm. **g, h,** Spatial graph and UMAP of 10x Visium analysis of the PFC at GW20, performed on consecutive tissue section analyzed by MERFISH. Clusters for detection spots were annotated based on tissue structure and gene expression. "M" in the cluster annotation means medial, as opposed to lateral. Capture area: 6.5mm x 6.5mm. **i,** Spatial heatmap for detected transcript count in each spot. Because each spot has a diameter of 55 μm and contains variable number of cells, the variation in transcript count is largely due to difference in cell density in different structures. **j,** Scaled expression spatial heatmaps showing additional marker genes for various IN subtypes are highly expressed in the VZ, including olfactory bulb-like IN markers *CALB2*, *PBX3*, and *TSHZ1*.

### Extended Data Fig. 4: Additional spatial analysis of cortical layers and neuronal subtypes.

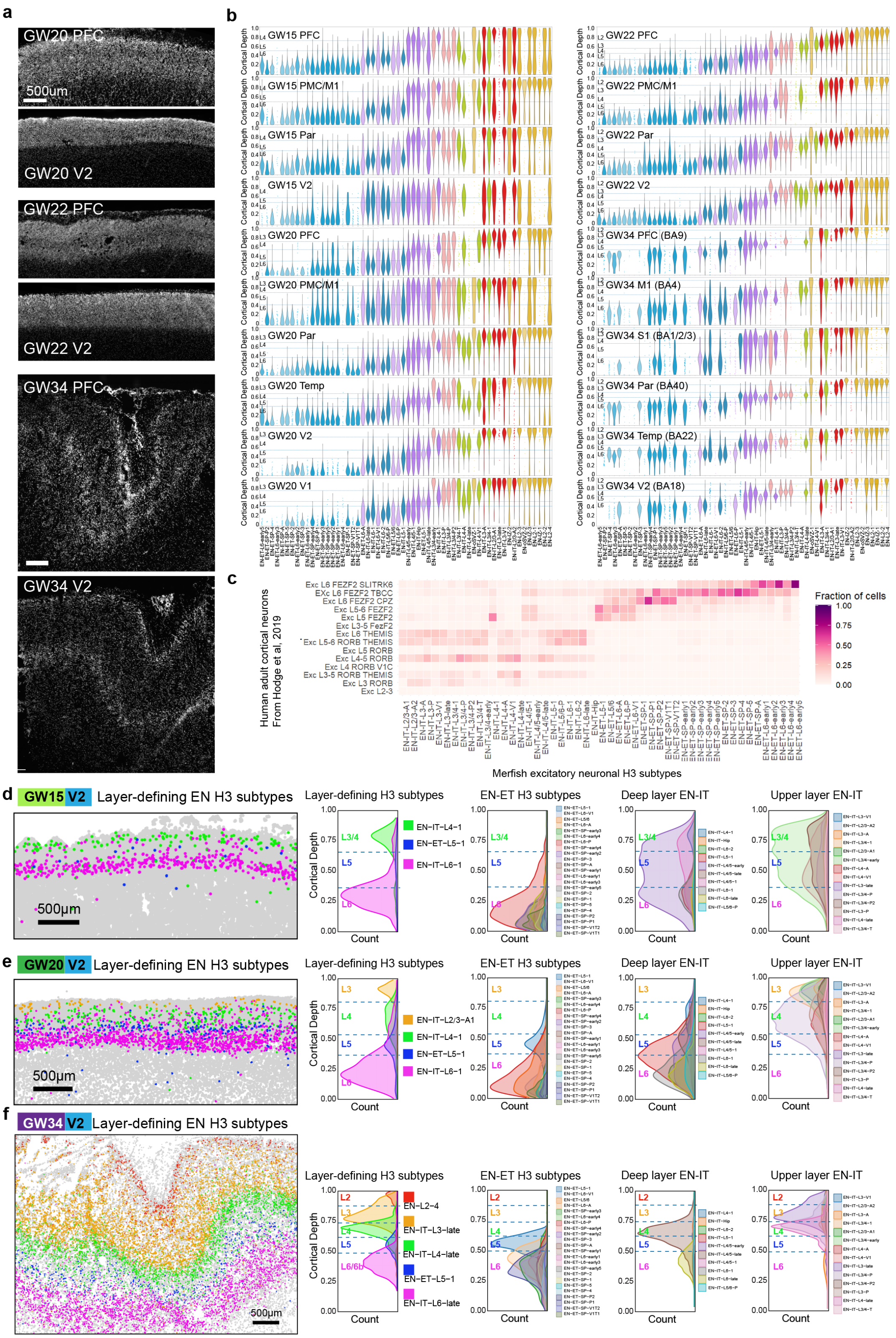

**Extended Data Fig. 4: Additional spatial analysis of cortical layers and neuronal subtypes. a,** Microscopy images of nuclei staining (DAPI) for the CP of PFC and V2 at GW20, 22 and 34, demonstrating that cytoarchitectural difference between cortical layers is not visible until GW34. These images were co-captured doing MERFISH imaging. Scale bars, 500 µm. **b,** Violin plots showing the CD distribution of H3 EN subtypes across major cortical areas at GW15 to 34. **c,** Cell type correspondence heatmap shows the fraction of cells from H3 EN subtypes associated to neuronal cell types from published adult human cortex scRNA-seq dataset from Hodge et al, 2019[2]. **d–f,** Spatial maps of layer-defining EN subtypes, and ridgeline plots showing the laminar distribution of H3 EN subtypes in the V2 at GW15 (**d**), 20 (**e**), and 34 (**f**). Scale bars, 500 µm.

### Extended Data Fig. 5: EN subtypes show distinct marker expression and area-dependent distribution.

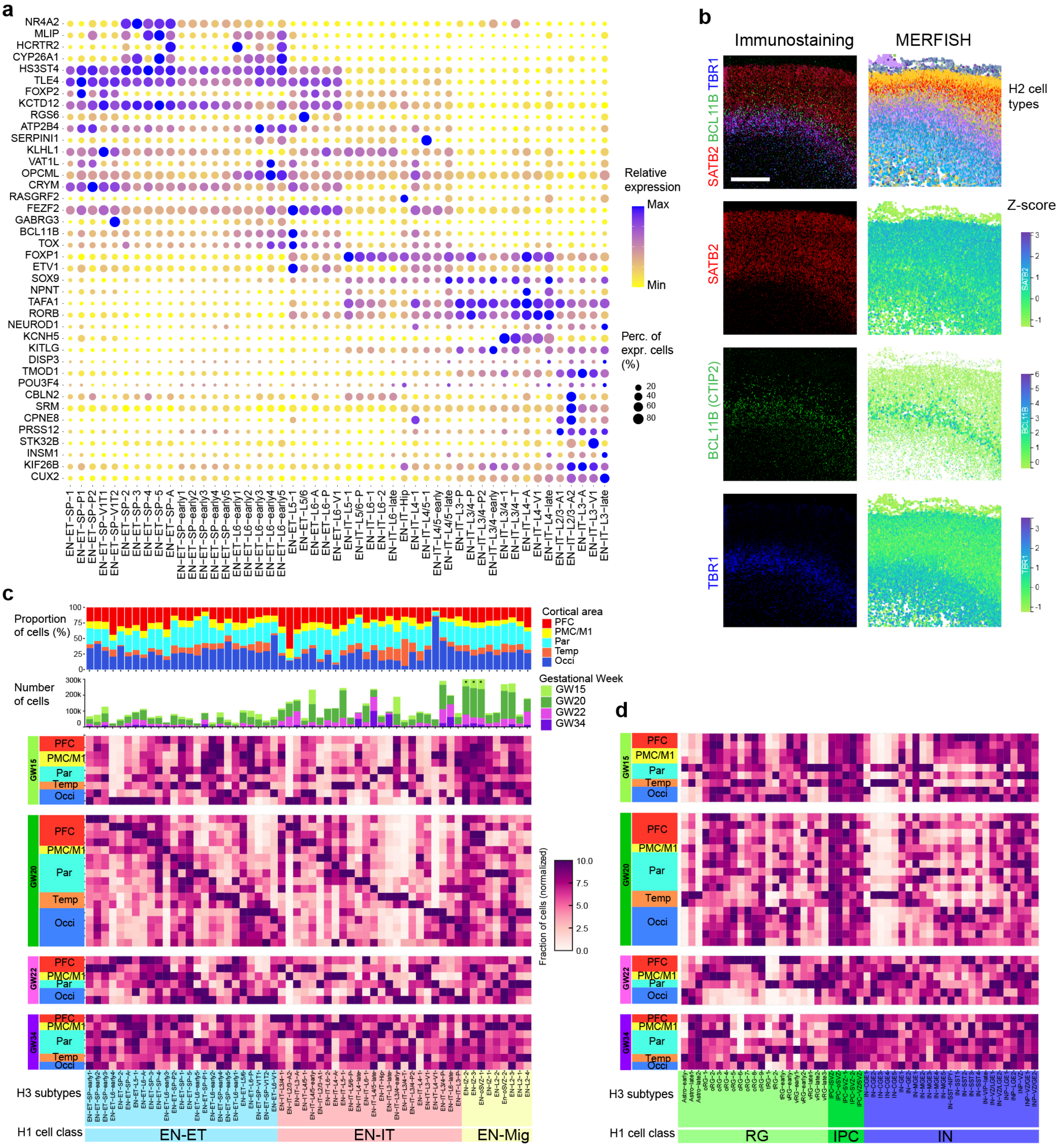

**Extended Data Fig. 5: EN subtypes show distinct marker expression and area-dependent distribution. a,** Dot plots showing the expression of genes exhibiting cortical layer-dependent distribution from **Fig. 3b** across H3 EN subtypes. **b,** Z-score spatial maps (right) from MERFISH experiments on the parietal cortex at GW22, and matching confocal microscopy images (left) of immunostaining for SATB2, CTIP2 (*BCL11B*), and TBR1. The immunostaining was performed on a tissue section adjacent to the one analyzed by MERFISH, and the same area is shown, demonstrating high spatial correspondence between mRNA detection by MERFISH and protein detection by immunohistology. Scale bar, 500 µm. **c,** Top, histogram showing the contribution of major cortical areas towards each H3 EN subtype. Mid, histogram showing the number of cells and contribution from each gestational age for H3 EN subtypes. Three columns that have values exceeding 300k cells are indicated with an asterisk. Bottom, enrichment heatmap showing the normalized fraction of cells from each H3 EN subtype that came from a given age and cortical area. Each row represents an independent MERFISH experiment, and each column represents an EN subtype under EN-IT, EN-ET, and EN-Mig. For each column the number of cells from each experiment was normalized to the total number of the cluster, and then normalized to the maximum value of the column within the same gestational week to show relative enrichment (**Methods**). **d,** Heatmap for areal enrichment of RG, IPC, and IN H3 subtypes, similar to **c**. Parts of the CGE were included in the temporal lobe samples for GW15 and 20, leading to strong enrichment of several IN-CGE subtypes.

### Extended Data Fig. 6: Areal and temporal dynamics of laminar gene expression.

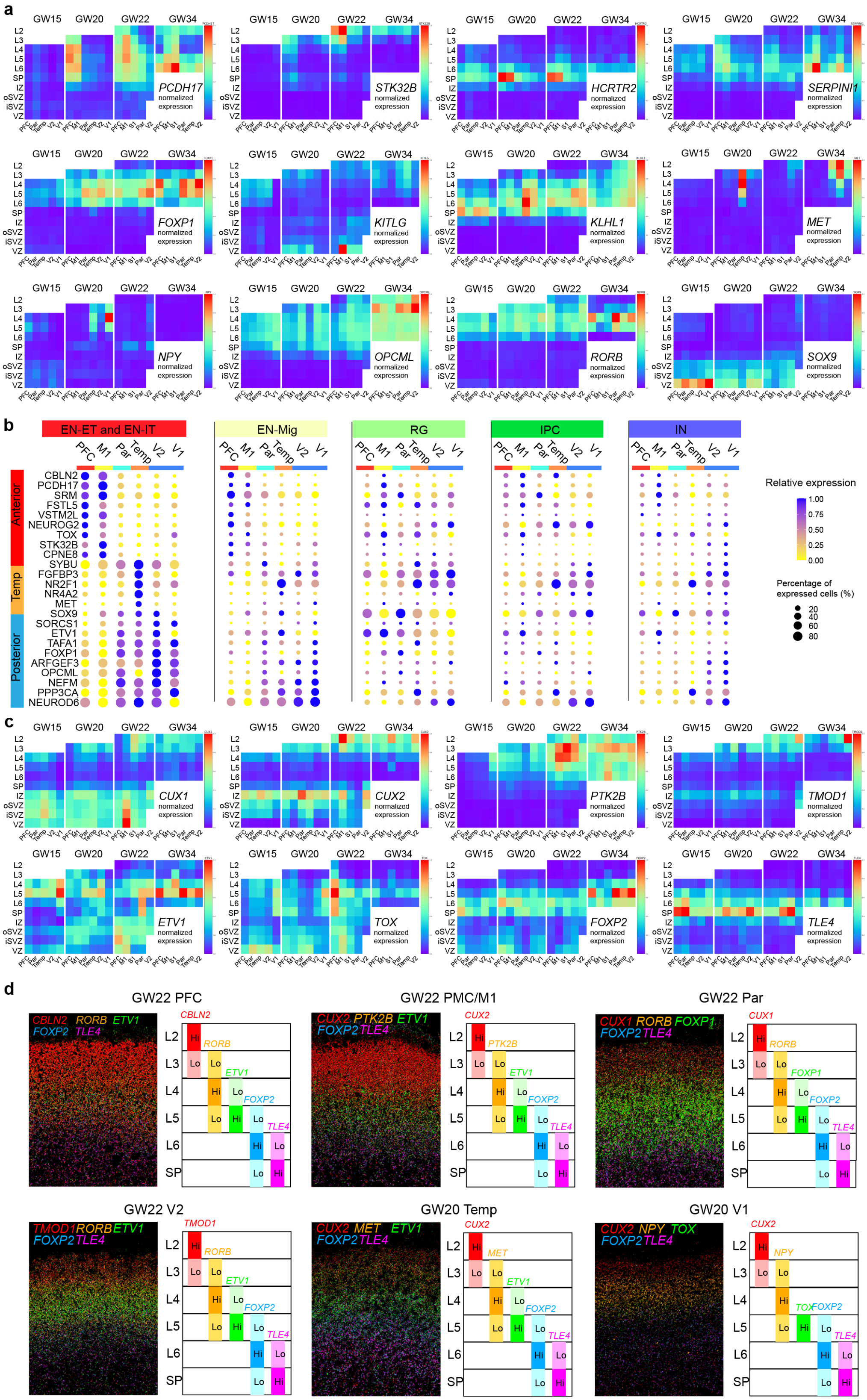

**Extended Data Fig. 6: Areal and temporal dynamics of laminar gene expression. a,** Summary expression heatmaps of genes that are anteriorly or posteriorly enriched consistently across all layer categories in **Fig. 3g. b,** Dot plots for the expression of areal markers in EN-Mig, RG, IPC and INs at GW20 show that only EN-Mig exhibited similar areal enrichment patterns as EN-IT and EN-ET, from which the markers are identified. **c,** Summary expression heatmaps showing the spatial temporal expression patterns of area-specific cortical layer markers listed in **Fig. 3j. d,** Pseudo-colored microscopy images of transcript dots for area-specific cortical layer marker combinations detected by MERFISH.

### Extended Data Fig. 7: Dynamic gene expression underlies diversification of EN subtypes.

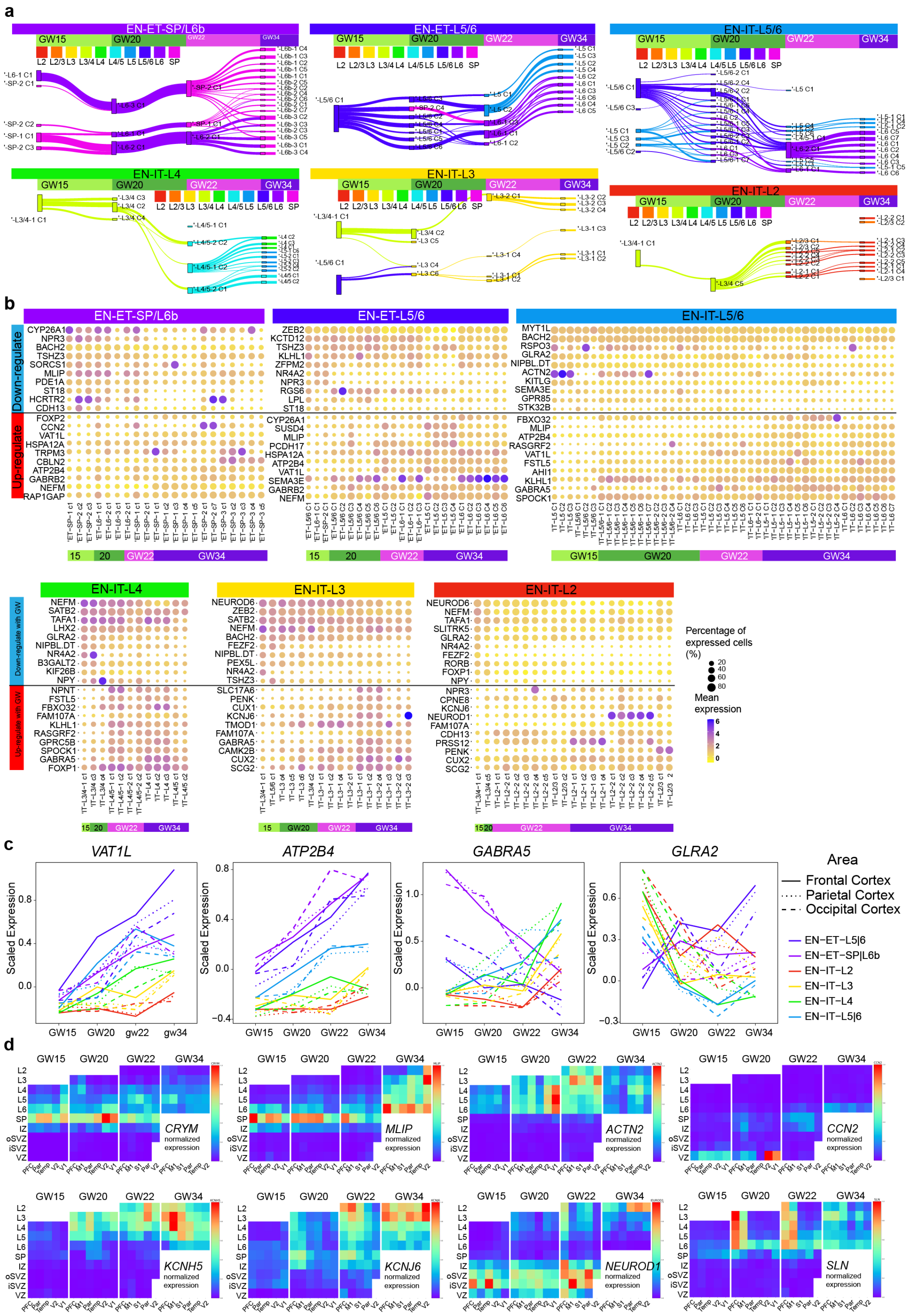

**Extended Data Fig. 7: Dynamic gene expression underlies diversification of EN subtypes. a,** Sankey diagrams showing the correspondence between scSHC EN subclusters from different gestational time points. Each layer-based group is shown separately. **b,** Dot plots showing the expression of temporally down- and up-regulated genes identified in **Fig. 4e** among scSHC EN subtypes within each group. **c,** Curve plots showing the change of scaled expression over time in different EN groups and cortical lobes for selected genes. Data from PFC, PMC and M1 are averaged for frontal cortex; V1 and V2 are averaged for occipital cortex. **d,** Summary expression heatmaps for high variance genes among EN subtypes display distinct laminar and areal enrichment.

### Extended Data Fig. 8: Spatial organization of cell subtypes at the V1-V2 border.

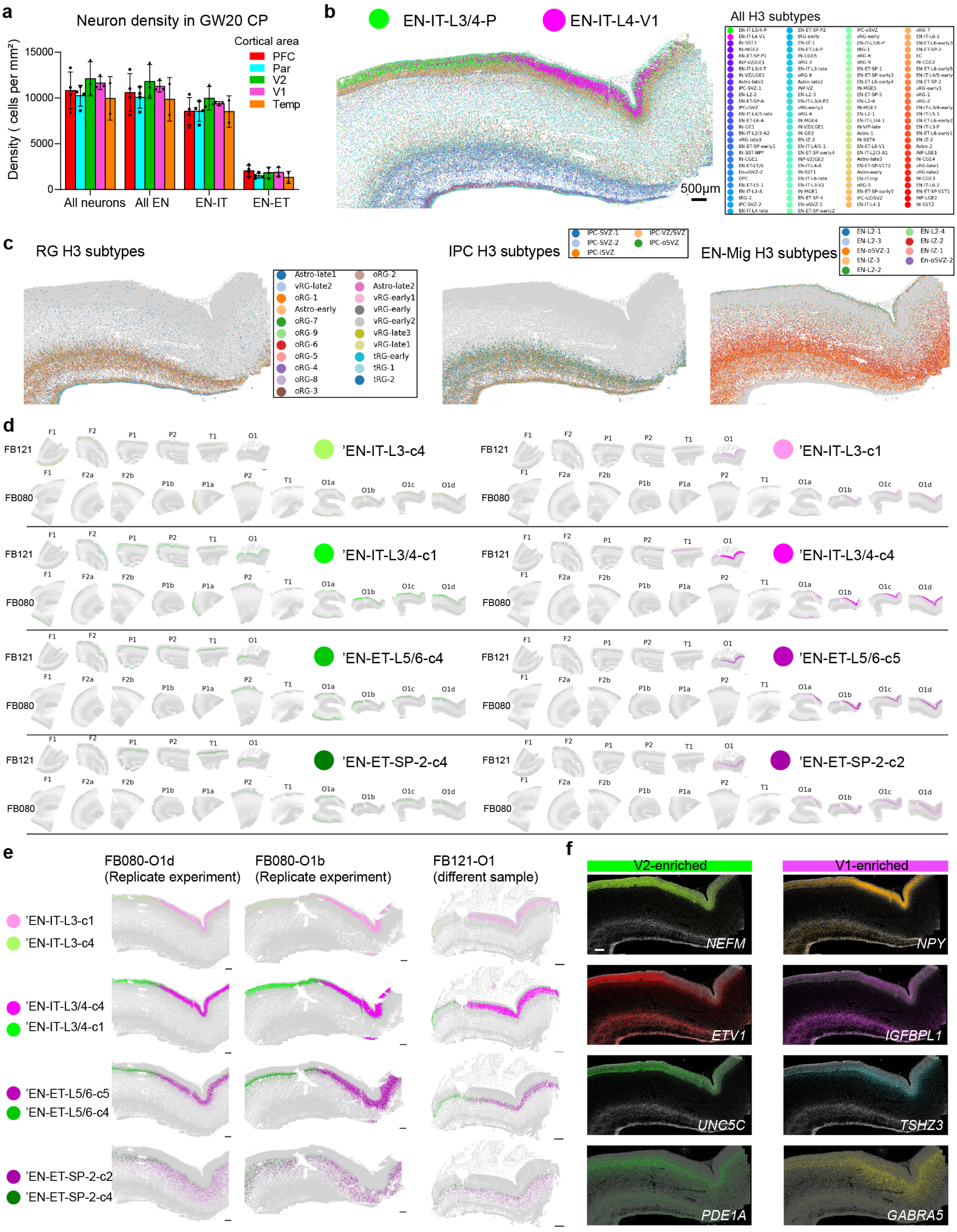

**Extended Data Fig. 8: Spatial organization of cell subtypes at the V1-V2 border. a,** Histograms of quantification of neuron density in the CP show no significant difference between major cortical areas at GW20. Error bars represent standard deviation. **b,** Spatial graph for the medial occipital cortex at GW20, colored by all H3 subtypes from integrated clustering, show the V1-V2 border is distinctly visible in the CP. Scale bar, 500 µm. **c,** Spatial graphs for EN-Mig, IPC, and RG subtypes show that a border between V1 and V2 is not seen in the IZ, SVZ, or VZ. **d,** Spatial graphs for the distribution of V1- and V2-enriched EN subtypes across all GW20 MERFISH experiments show that V1-enriched clusters are exclusively found in the V1, while V2-enriched clusters exhibit broad distributions despite overall posterior enrichment. **e,** Spatial graphs for V1- and V2- enriched EN subtypes from replicate experiments and an independent replicate sample consistently show subtype transition at the border in all cortical layers and the SP. Scale bars, 500 µm. **f,** Pseudo-colored microscopy images of transcript dots detected by MERFISH for genes that are differentially expressed at the V1-V2 border. Scale bar, 500 µm.

### Extended Data Fig. 9: V1 and V2 show distinct gene expressions at GW15, but a clear border has not formed.

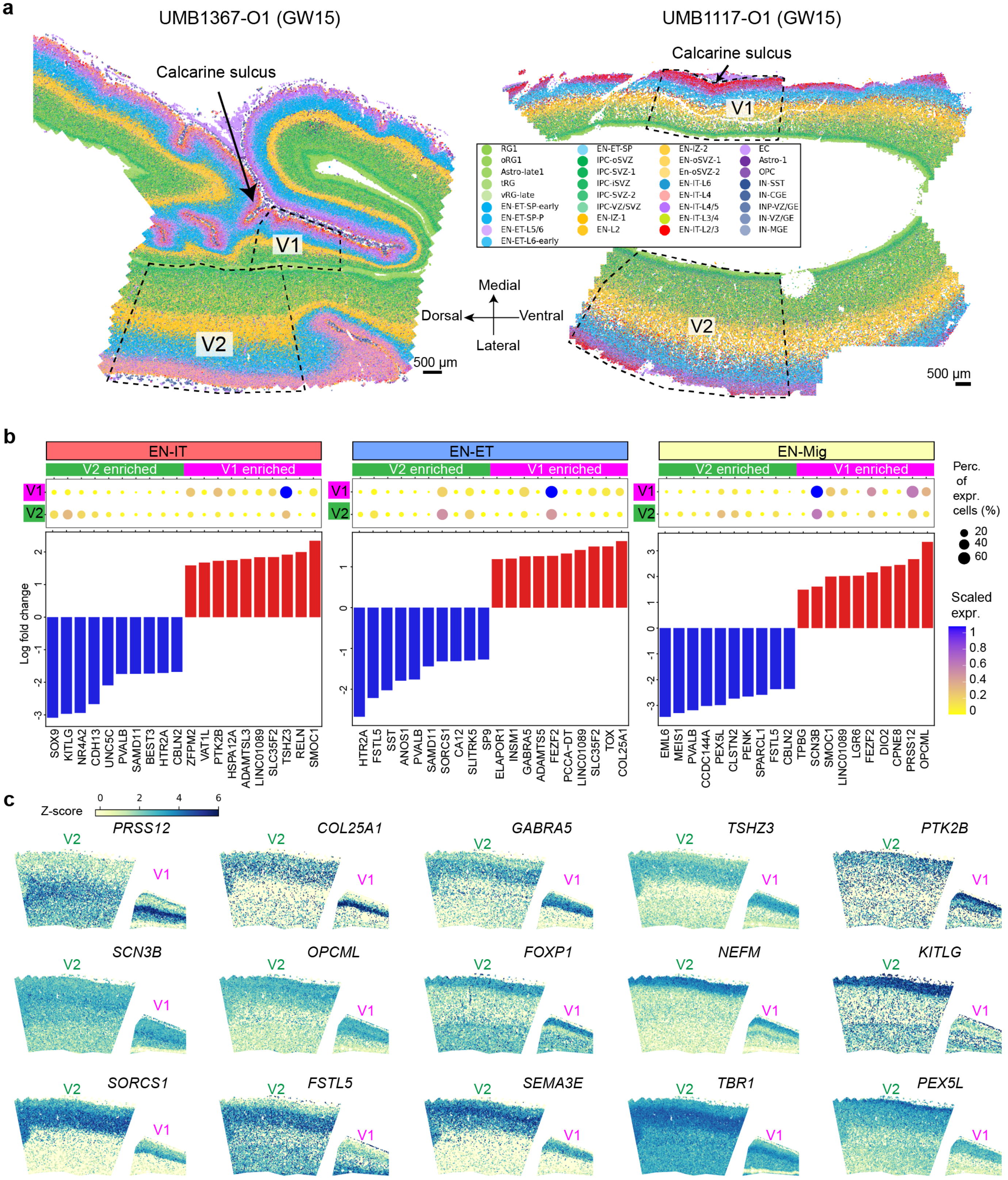

**Extended Data Fig. 9: V1 and V2 show distinct gene expressions at GW15, but a clear border has not formed. a,** Spatial graphs for the occipital cortex at GW15 from two samples. Because the regions of V2 cannot be reliably located at the medial side, the V2 at the lateral side is used for comparison, as indicated by the dash lined area. Note the ventral end of UMB1367-O1 (left) was bent to the top during the freezing process. Scale bars, 500 μm. **b,** Dot plots and histograms of fold change for top DEGs between V1 and V2 identified in EN-ITs, EN-ETs, and EN-Mig. **c,** Z-score spatial graphs for V1-V2 DEGs at GW15. The putative V1 and V2 regions are cropped from UMB1367-O1, shown as dash lined areas in **a**.

### Extended Data Fig. 10: Whole transcriptomics analyses at the V1-V2 border.

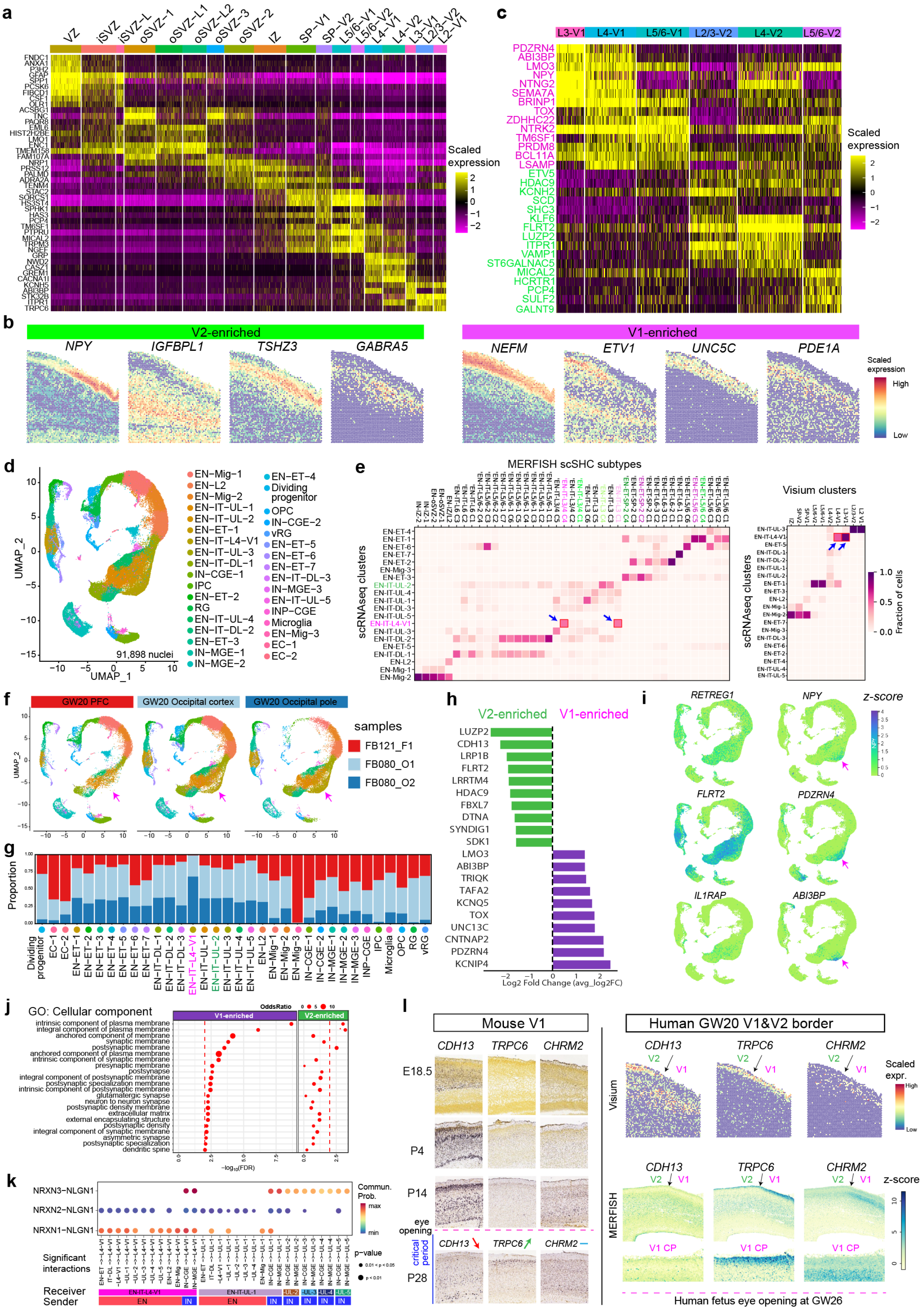

**Extended Data Fig. 10: Whole transcriptomics analyses at the V1-V2 border. a,** Heatmap showing the expression of top enriched marker genes for each Visium cluster. Columns represent individual Visium detection spots. **b,** Visium spatial graphs for V1- and V2-enriched genes show high consistency with MERFISH measurements in **Fig. 5c**. **c,** Heatmap showing the expression of top DEGs between V1 and V2 identified separately for each cortical layer. **d,** UMAP for 91,898 nuclei analyzed by snRNAseq, colored by unsupervised clustering. **e,** Heatmaps showing cell type correspondence of snRNAseq clusters with MERFISH scSHC EN subtypes (left), and with Visium spot clusters (right). EN-IT-L4-V1 cluster from snRNAseq corresponds strongly to V1-specific Layer 3&4 EN-ITs (arrows), whereas EN-IT-UL-2 corresponds to V2-enriched Layer 3&4 EN-ITs. **f,** UMAPs showing cells from the three specimens separately highlight the absence of EN-IT-L4-V1 cluster (arrow) in the PFC, validating its exclusive presence in the V1. FB121_F1 is GW20 PFC sample, FB080_O1 and FB080_O2 are both GW20 occipital cortex sample, containing both V1 and V2. FB080_O2 is closer to the occipital pole, and therefore contains a higher proportion of V1 than FB080_O1. **g,** Histogram for the cell number contribution from each sample towards each cluster, highlighting the V1-exclusive presence of EN-IT-L4-V1. **h,** Bar graph showing top DEGs between EN-IT-L4-V1 (V1-enriched) and EN-IT-UL-2 (V2 enriched) clusters. **i,** Z-score plots for the expression of V1- and V2- enriched genes demonstrate high consistency between snRNAseq, MERFISH, and Visium analysis. **j,** Gene ontology (GO) analysis reveals genes associated with synapse-related membrane components are upregulated in EN-IT-L4-V1, comparing to EN-IT-UL-2. **k,** Dot plot for specific signaling interactions between neuronal clusters highlight EN-IT-L4-V1 is unique among upper layer (UL) EN-ITs in receiving strong NRXN1-NLGN1 signaling from EN sources. Only significant interactions with p-value<0.05 are shown. **l,** Left, in situ hybridization images from Allen Developmental Mouse Brain Atlas (https://developingmouse.brain-map.org/) showing the expression of *CDH13*, *TRPC6* and *CHRM2* in the mouse V1 at different developmental time points. Right, Visium spatial heatmaps and MERFISH z-score spatial graphs for the expression of *CDH13*, *TRPC6* and *CHRM2* at the human V1-V2 border at GW20. *TRPC6* and *CHRM2* expressions on MERFISH data are imputed based on snRNAseq data.
