## Supplementary figures and images for "Spatial Single-cell Analysis Decodes Cortical Layer and Area Specification"

### FOXP1.png

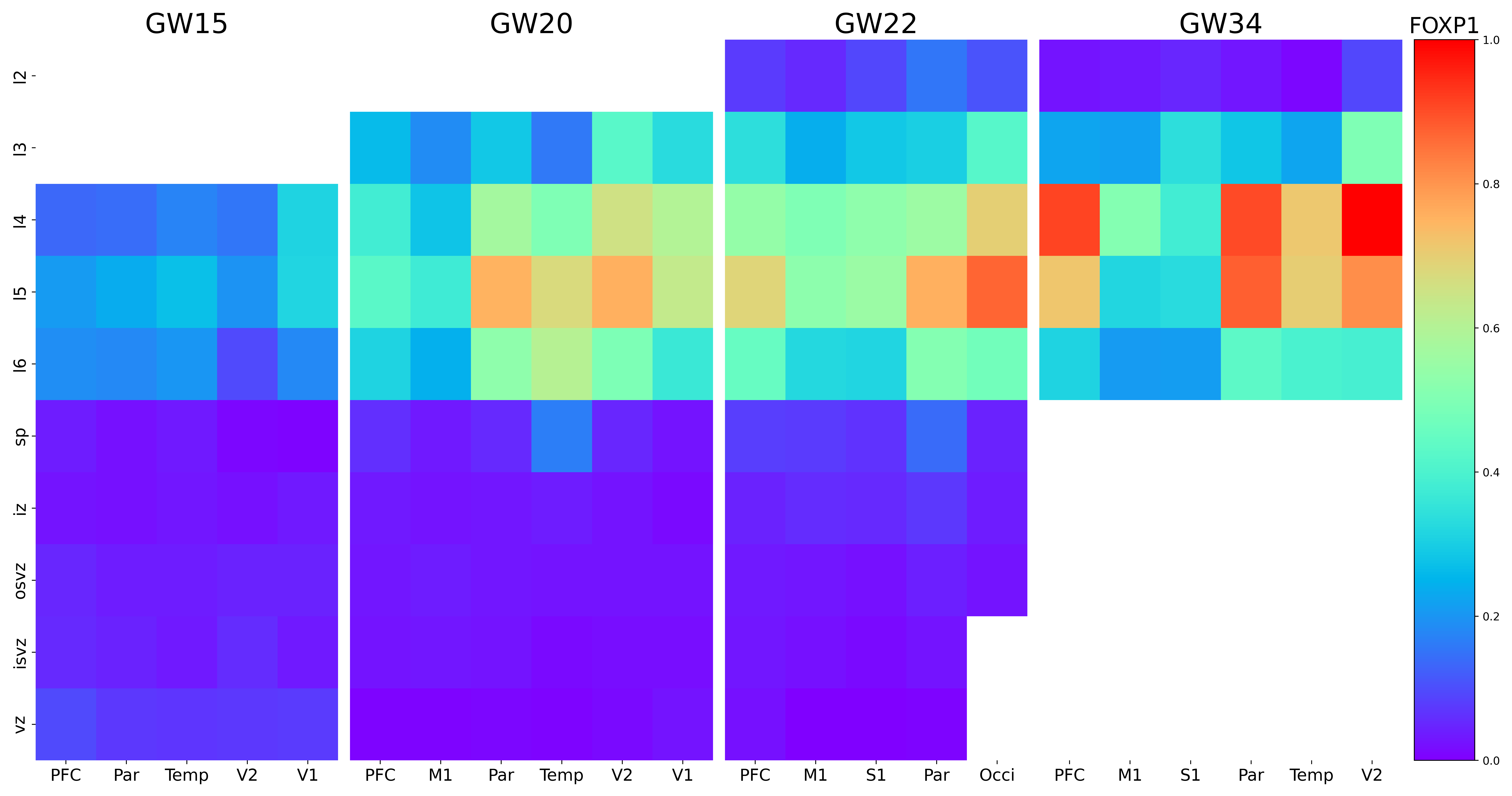

### FOXP2.png

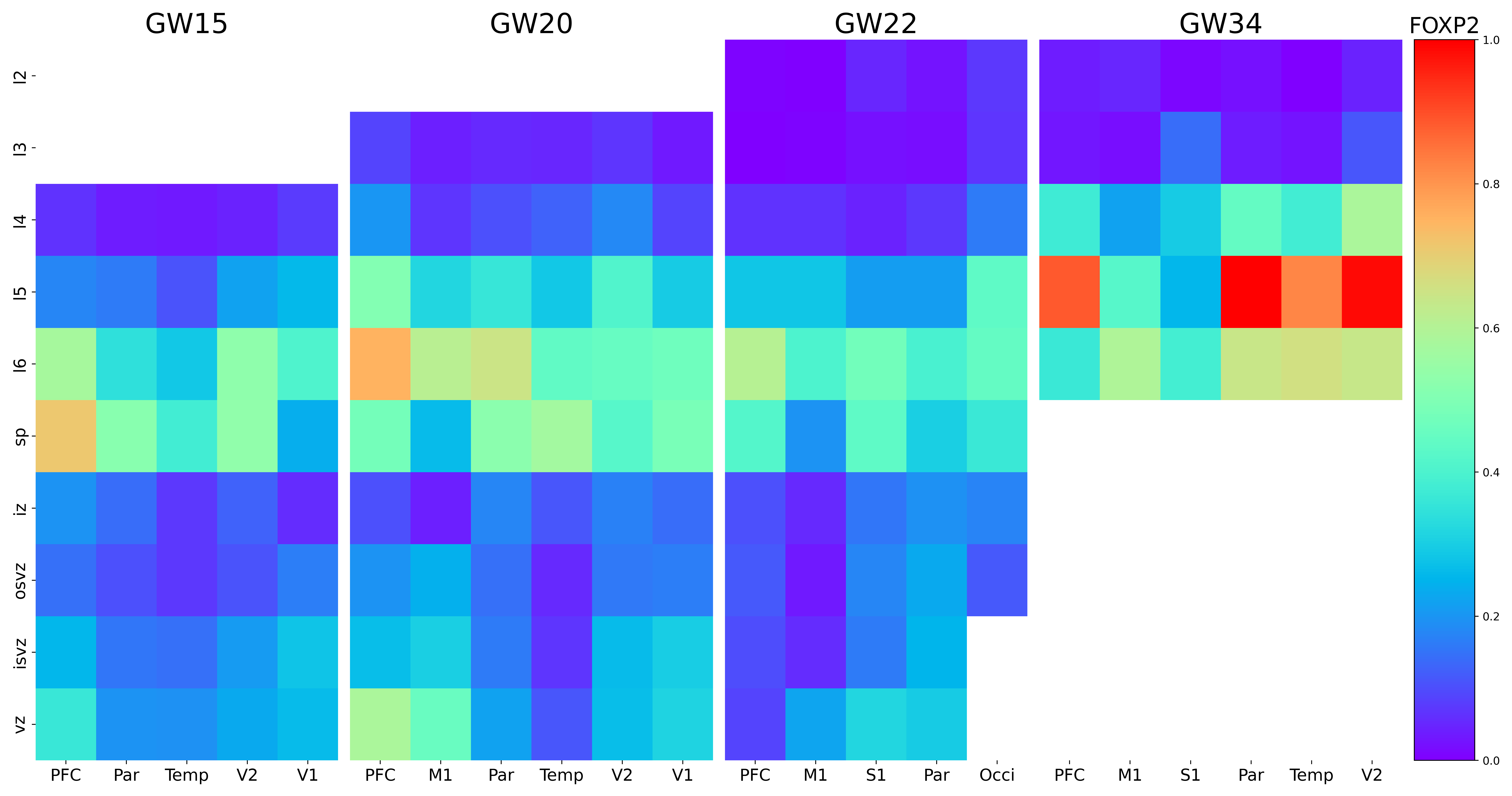

### FRG1-DT.png

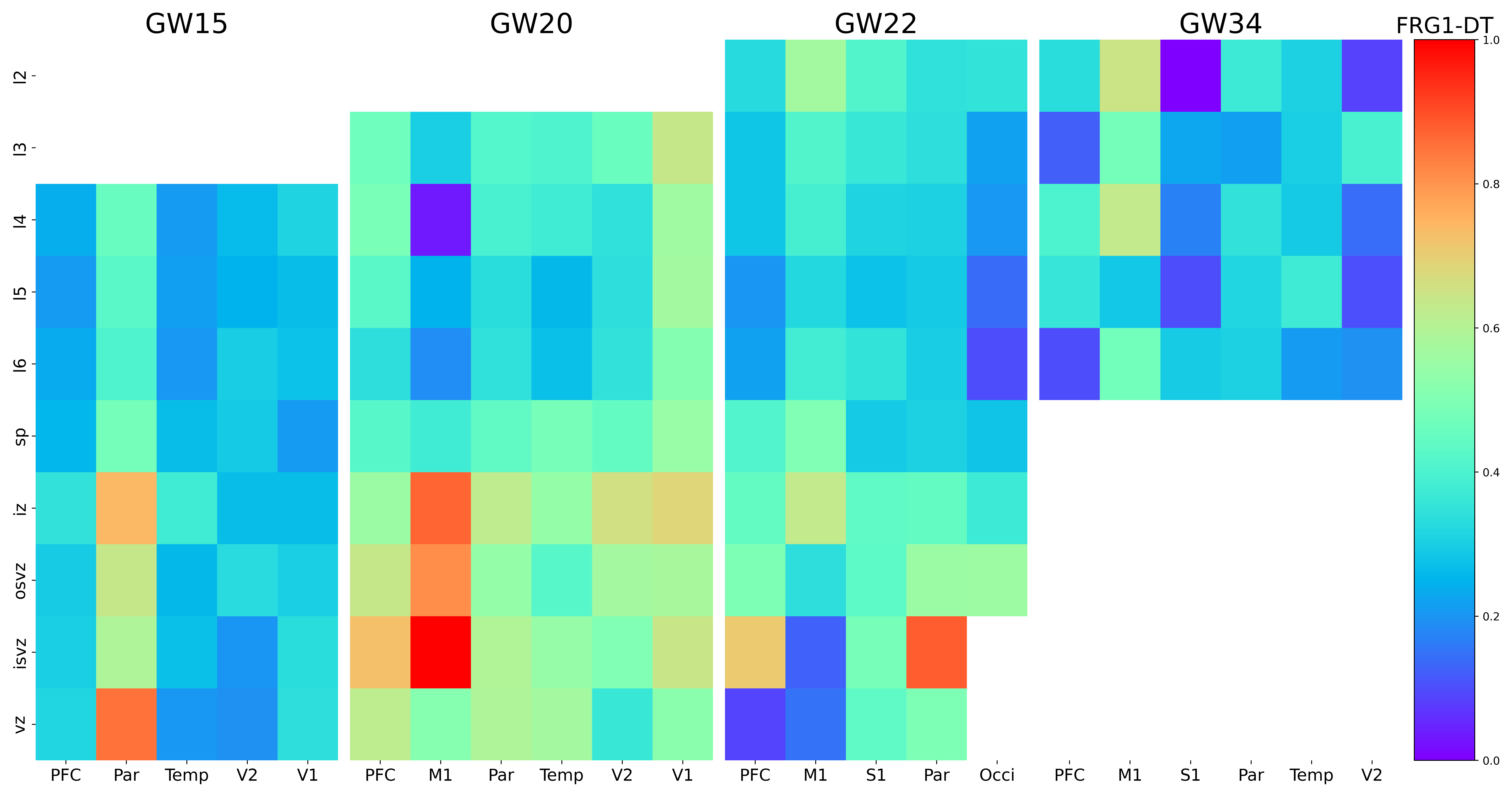

### FRG1.png

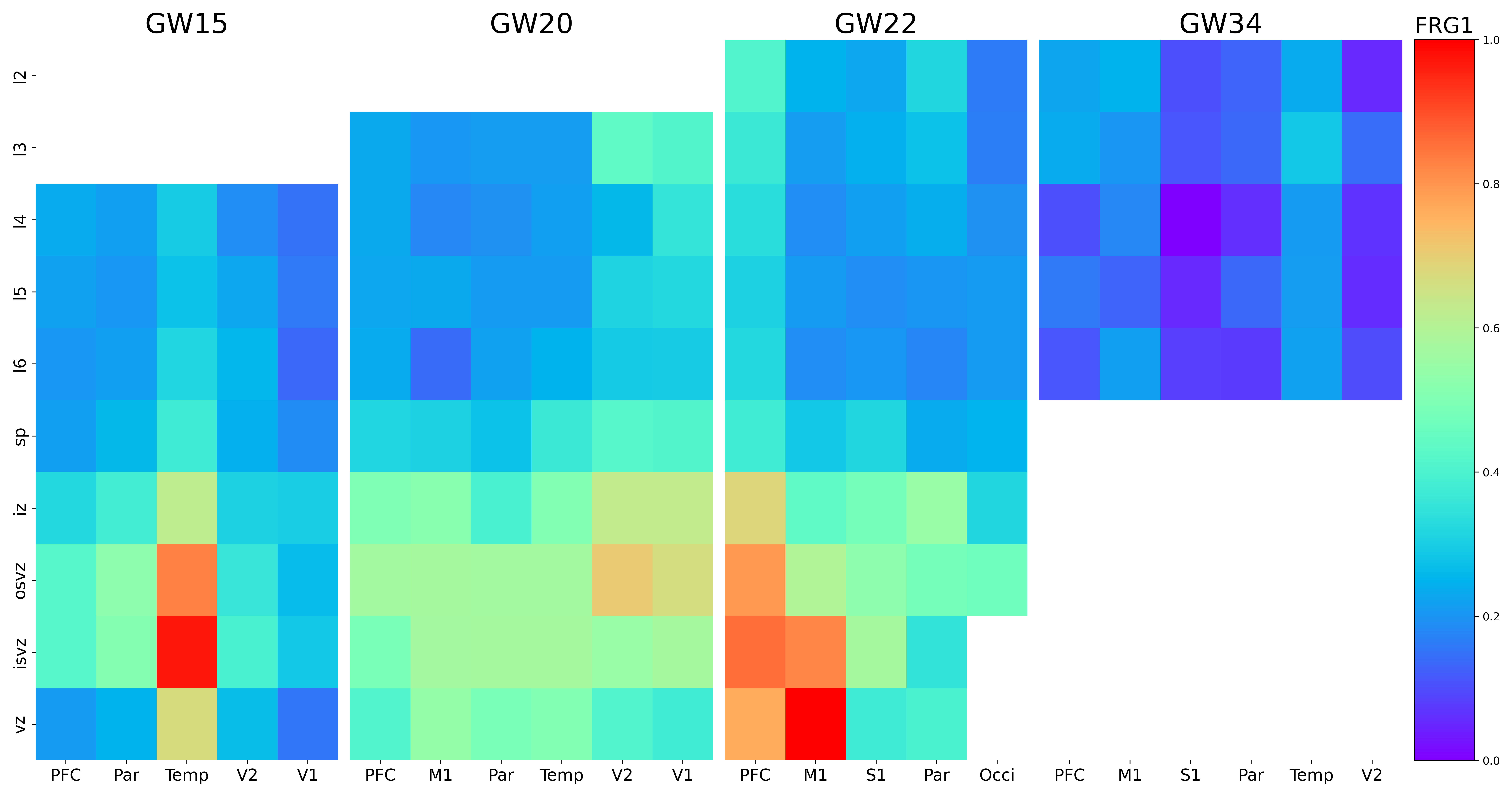

### FSTL5.png

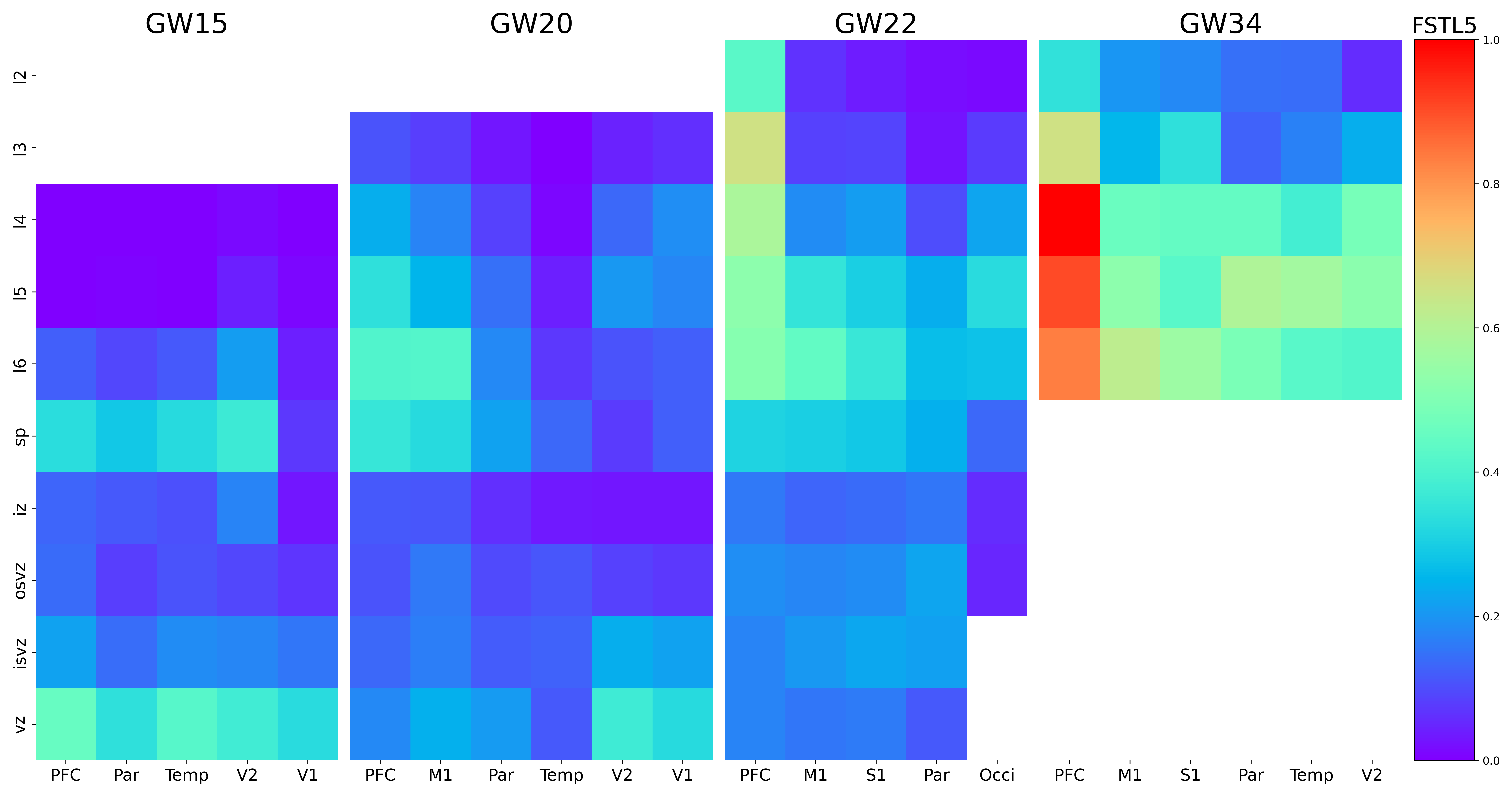

### FXYD6.png

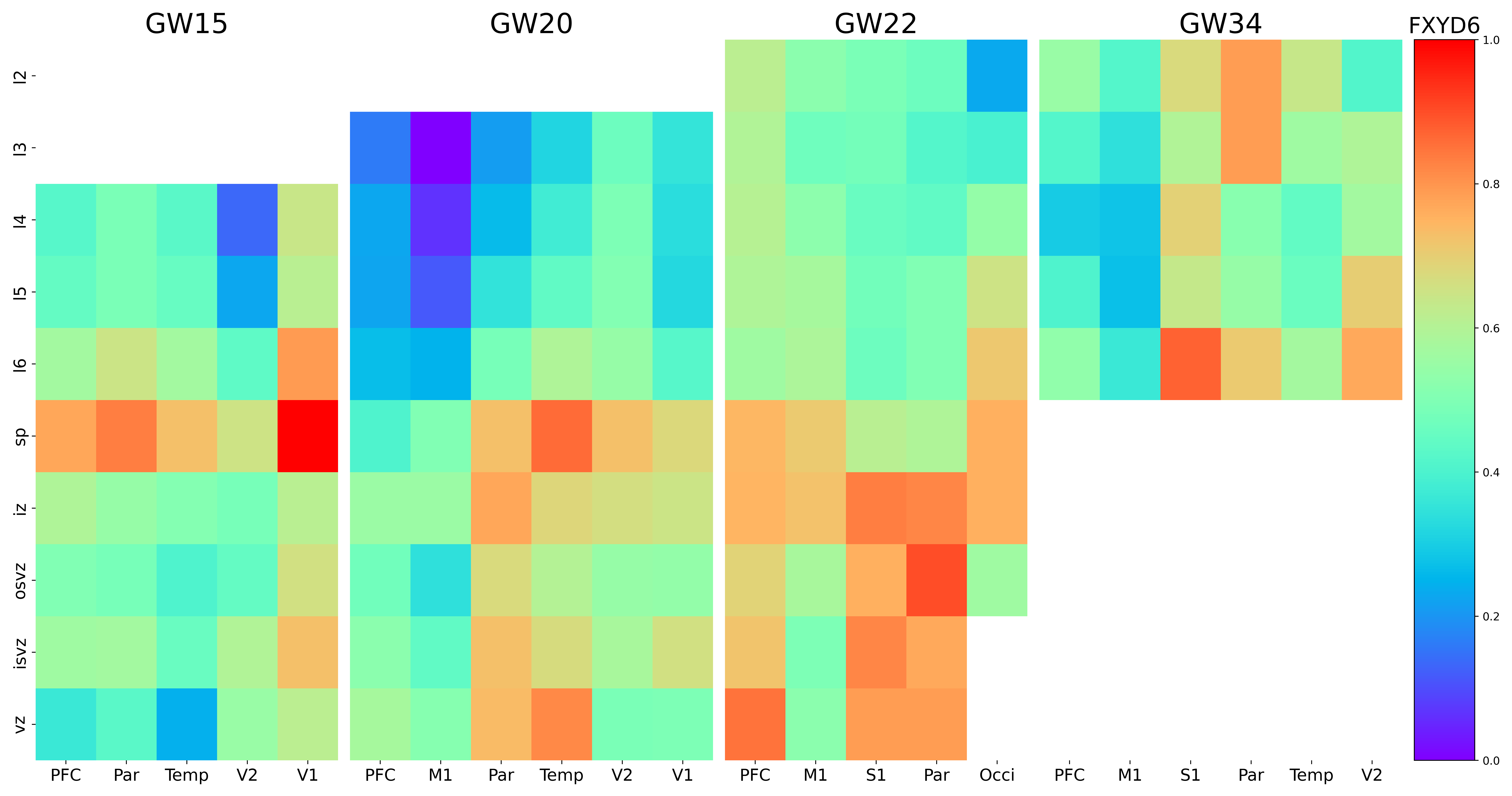

### GABRA5.png

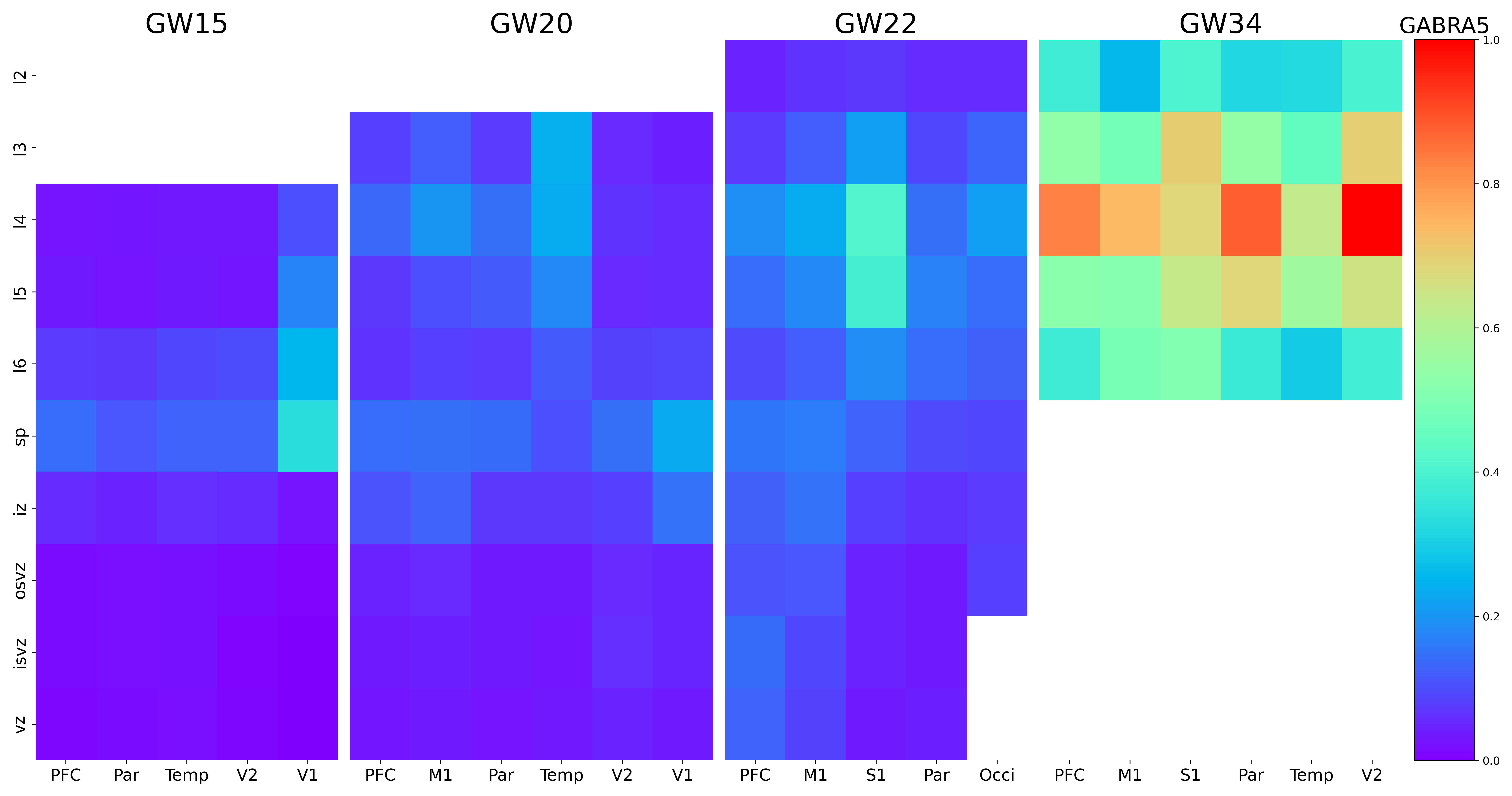

### GABRB2.png

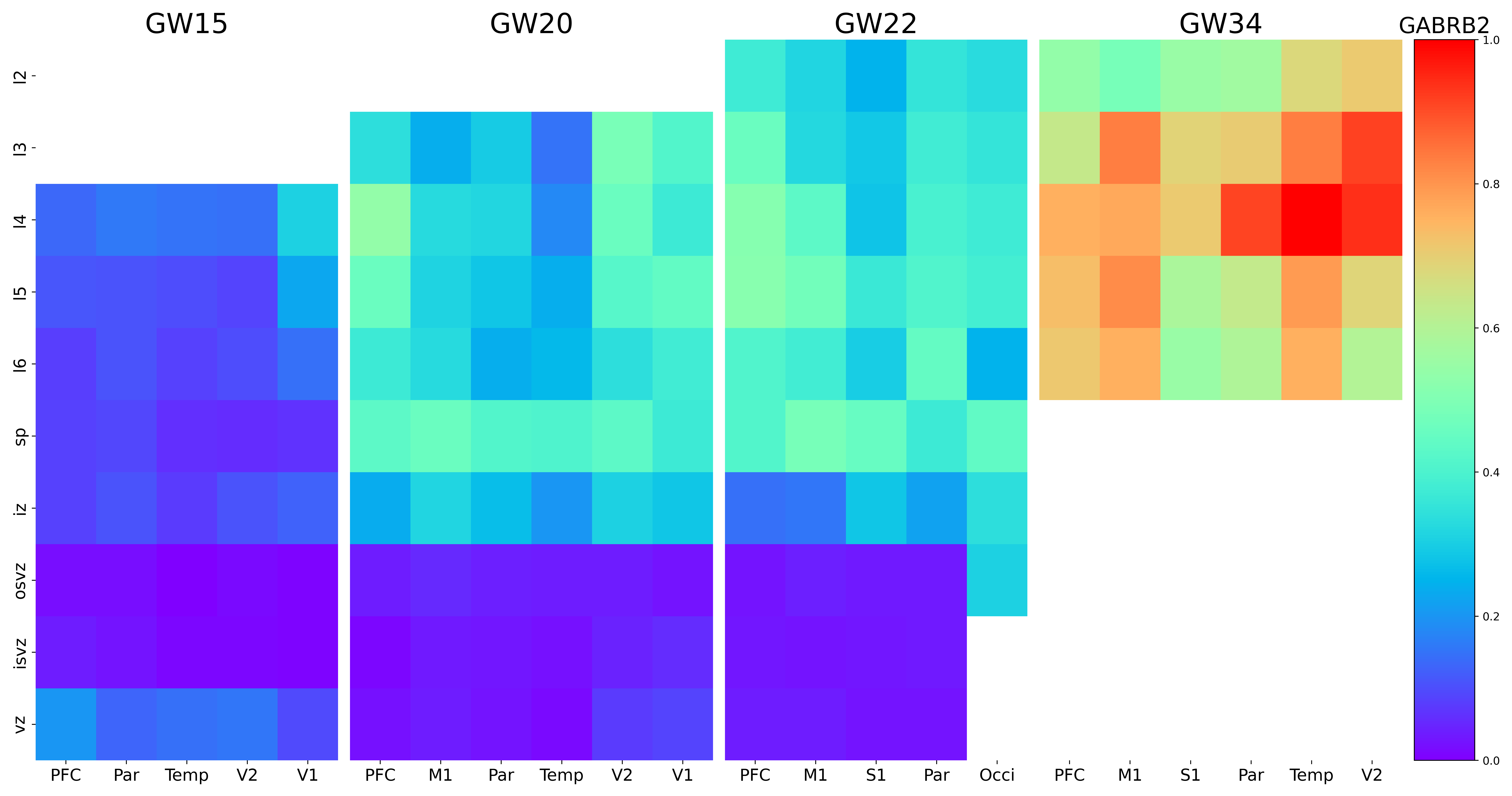

### GABRG3.png

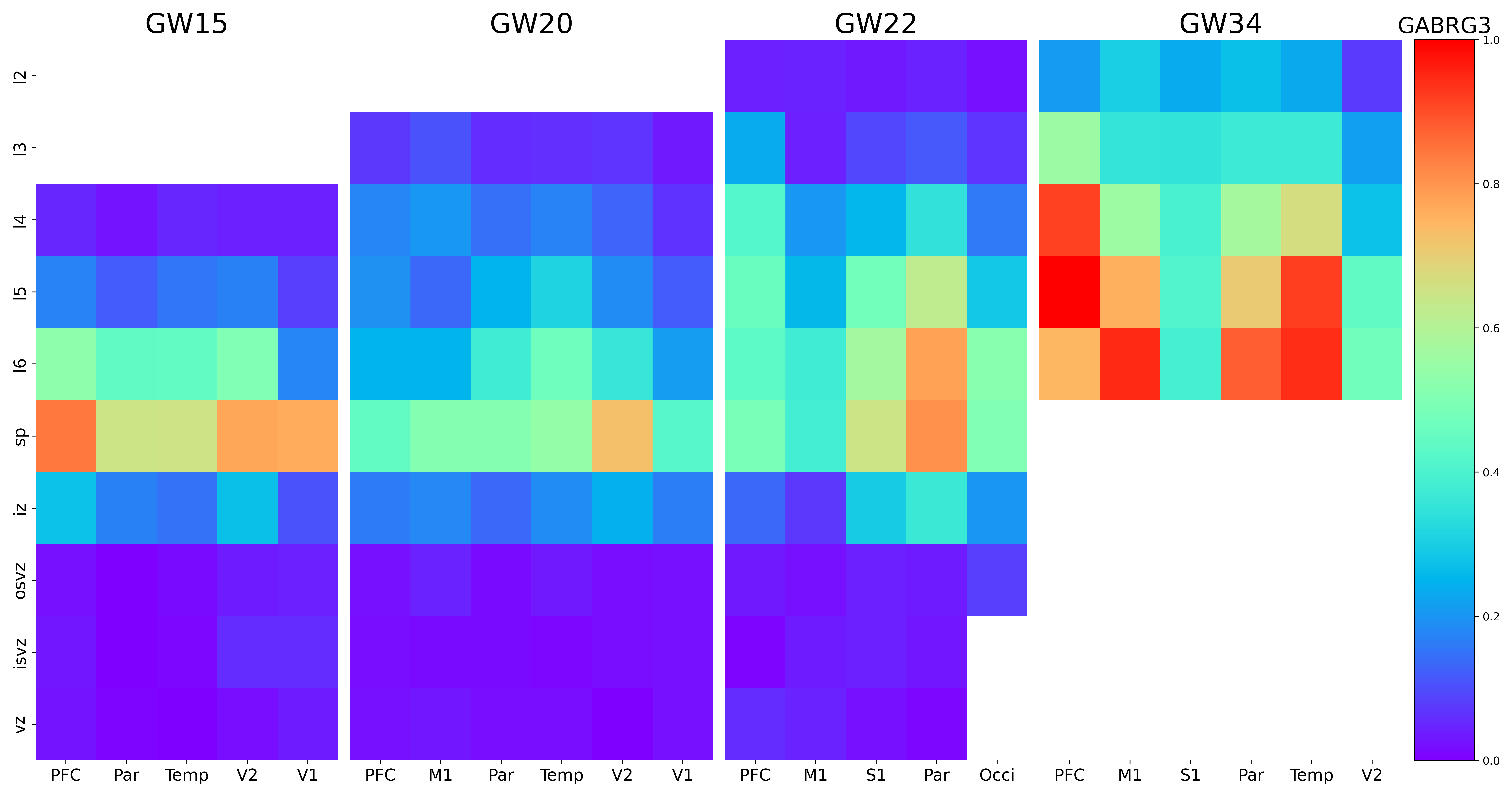

### GAD2.png

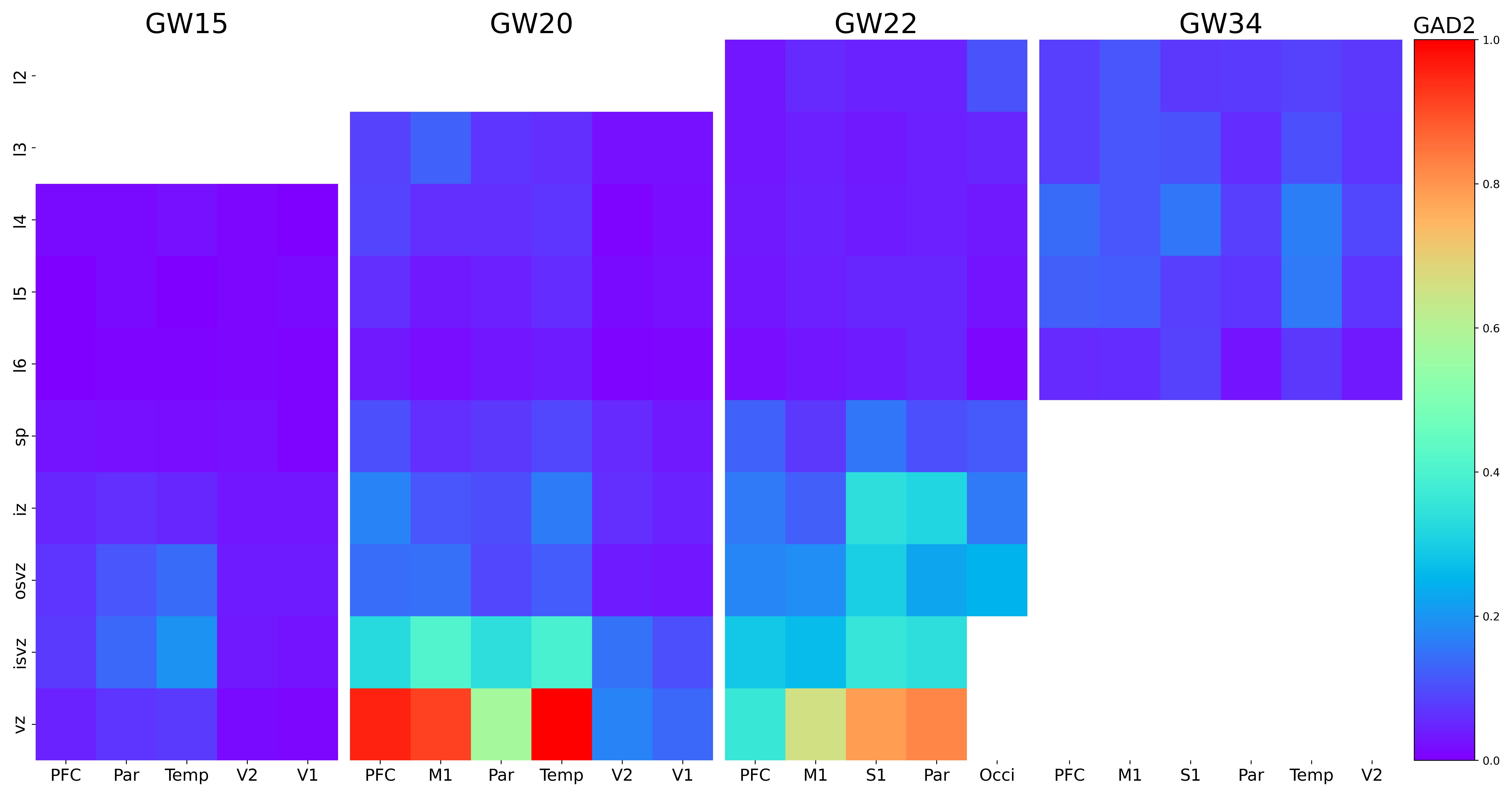

### GFAP.png

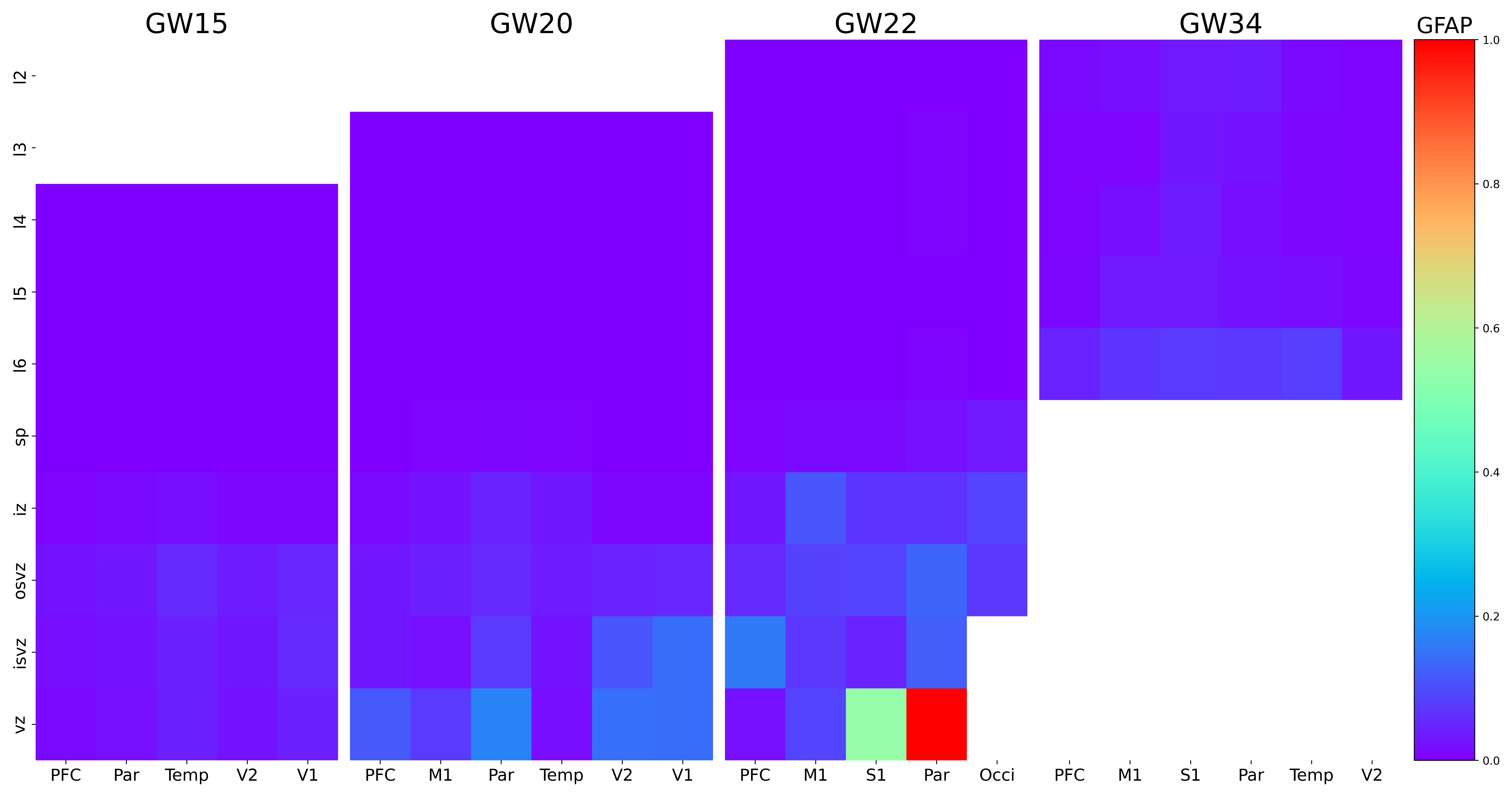

### GLB1L2.png

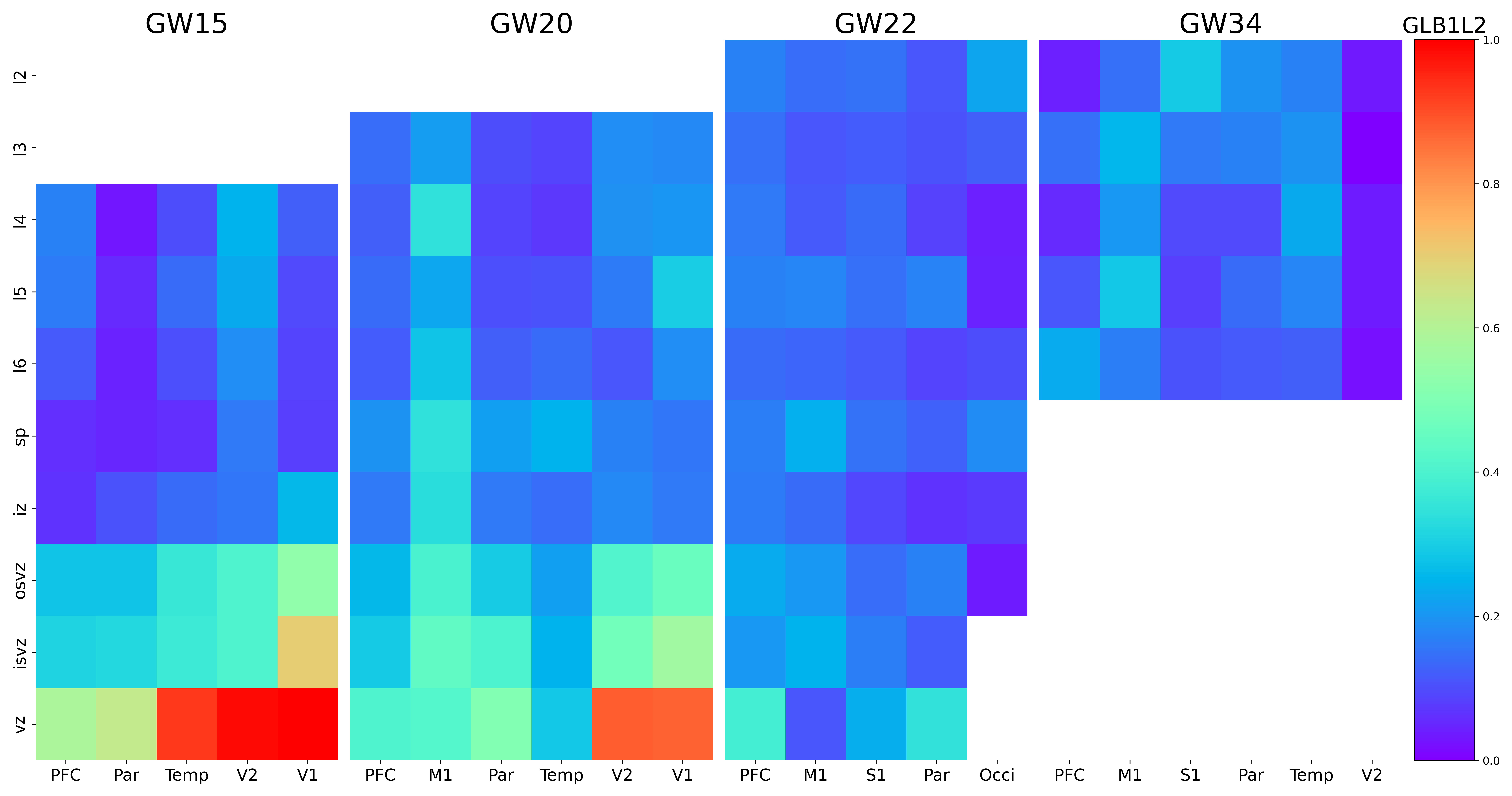

### GLI3.png

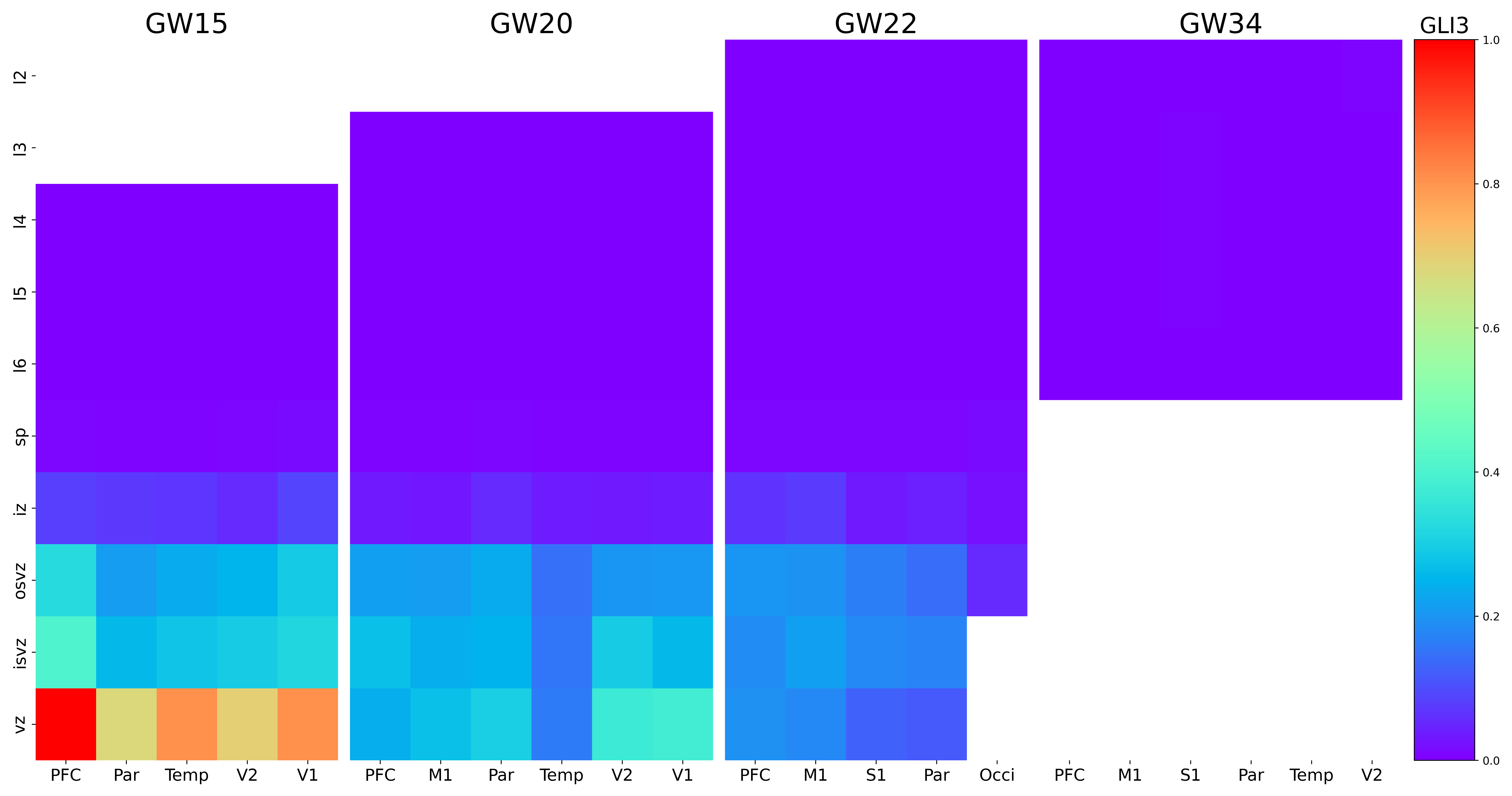

### GLRA2.png

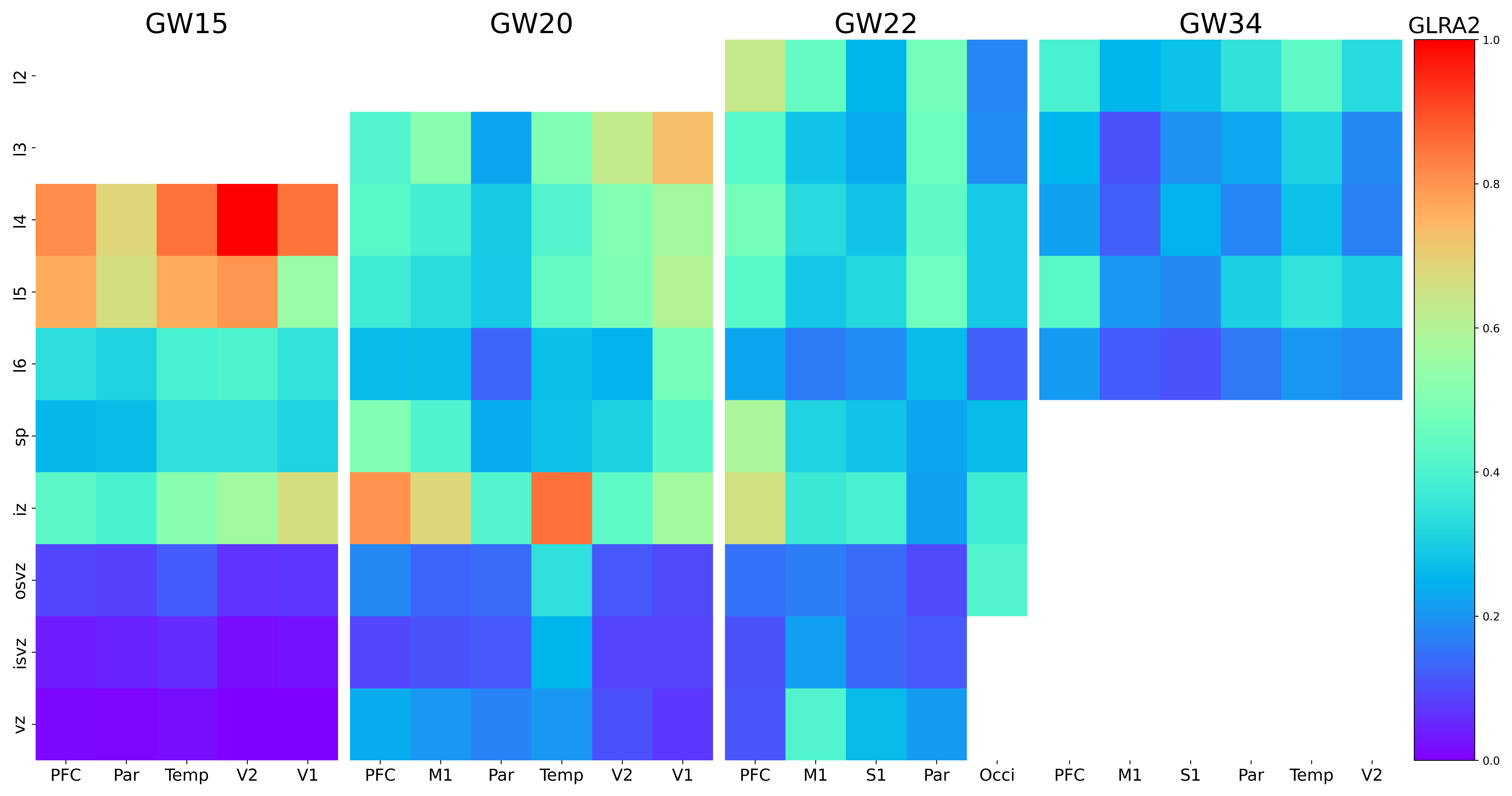

### GLRA3.png

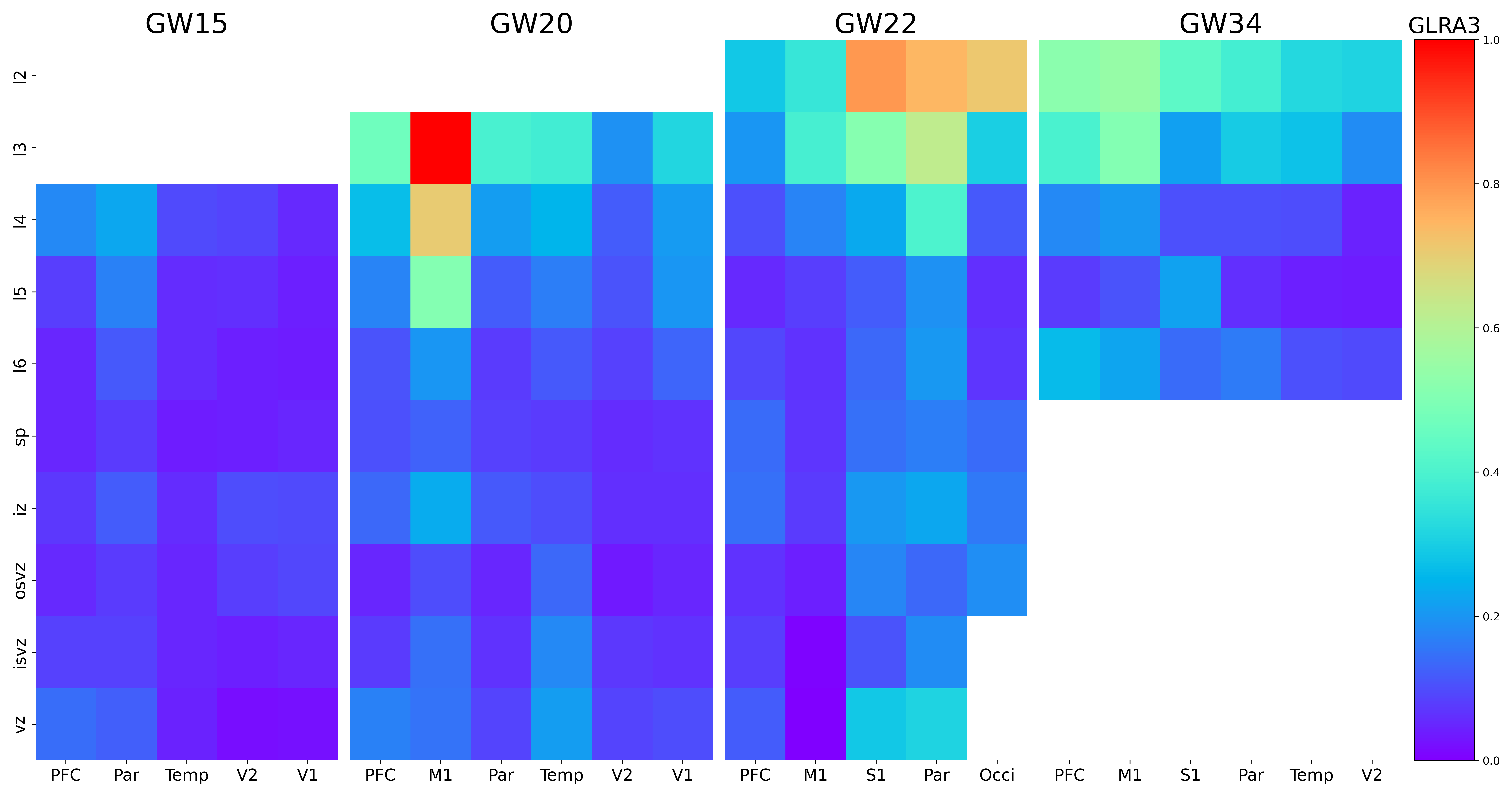

### GNAL.png

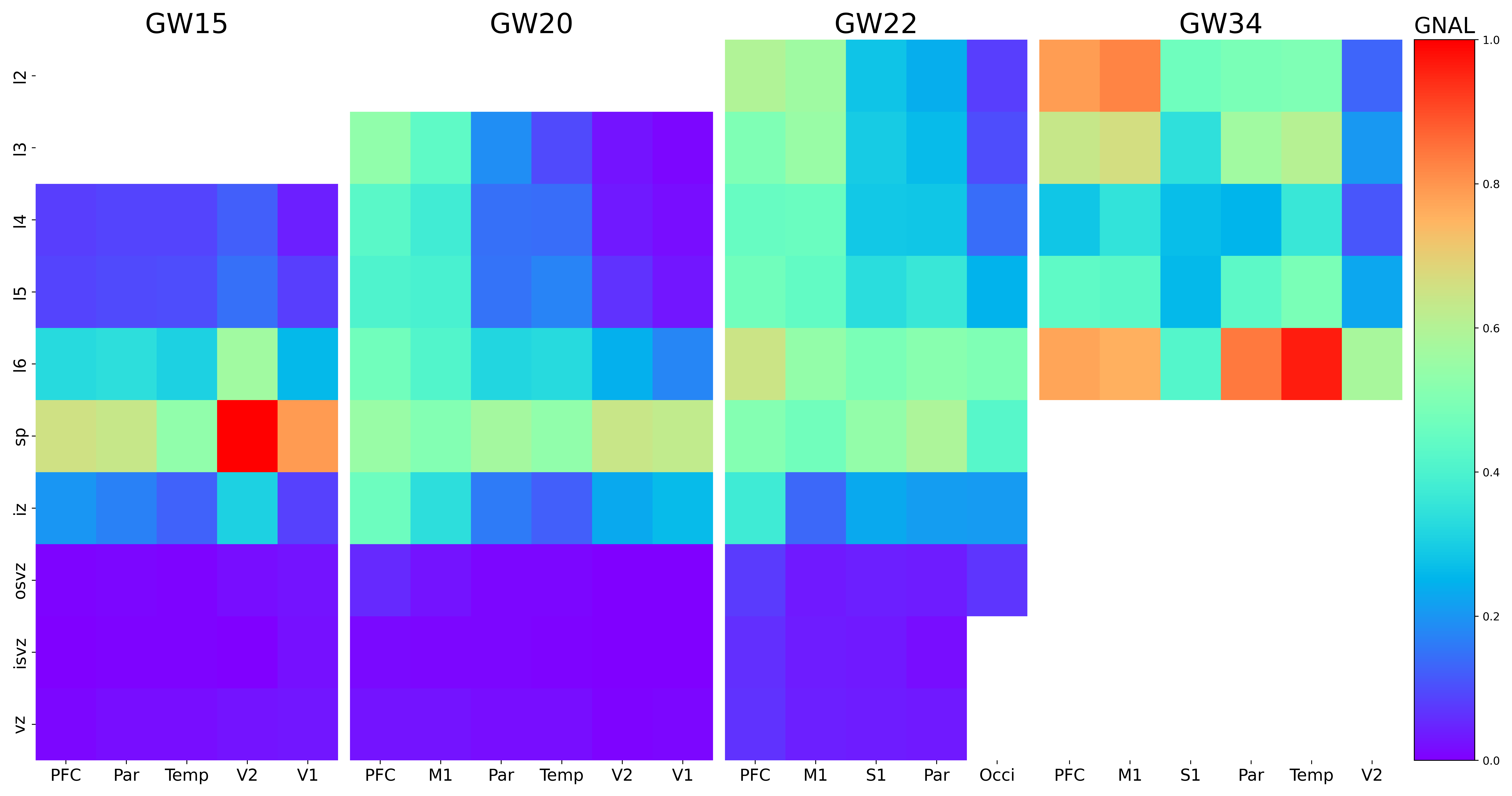

### GPR37L1.png

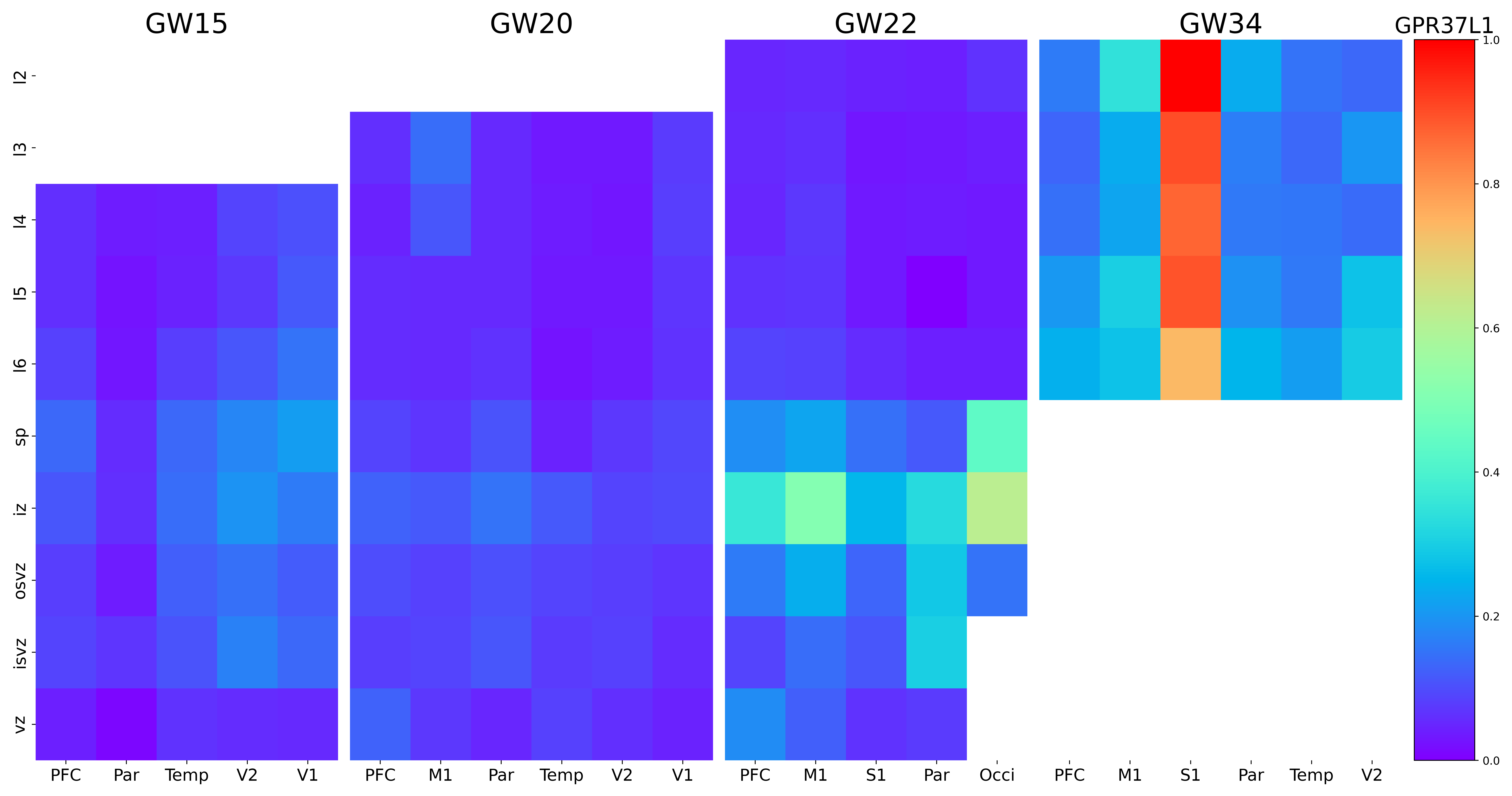

### GPR85.png

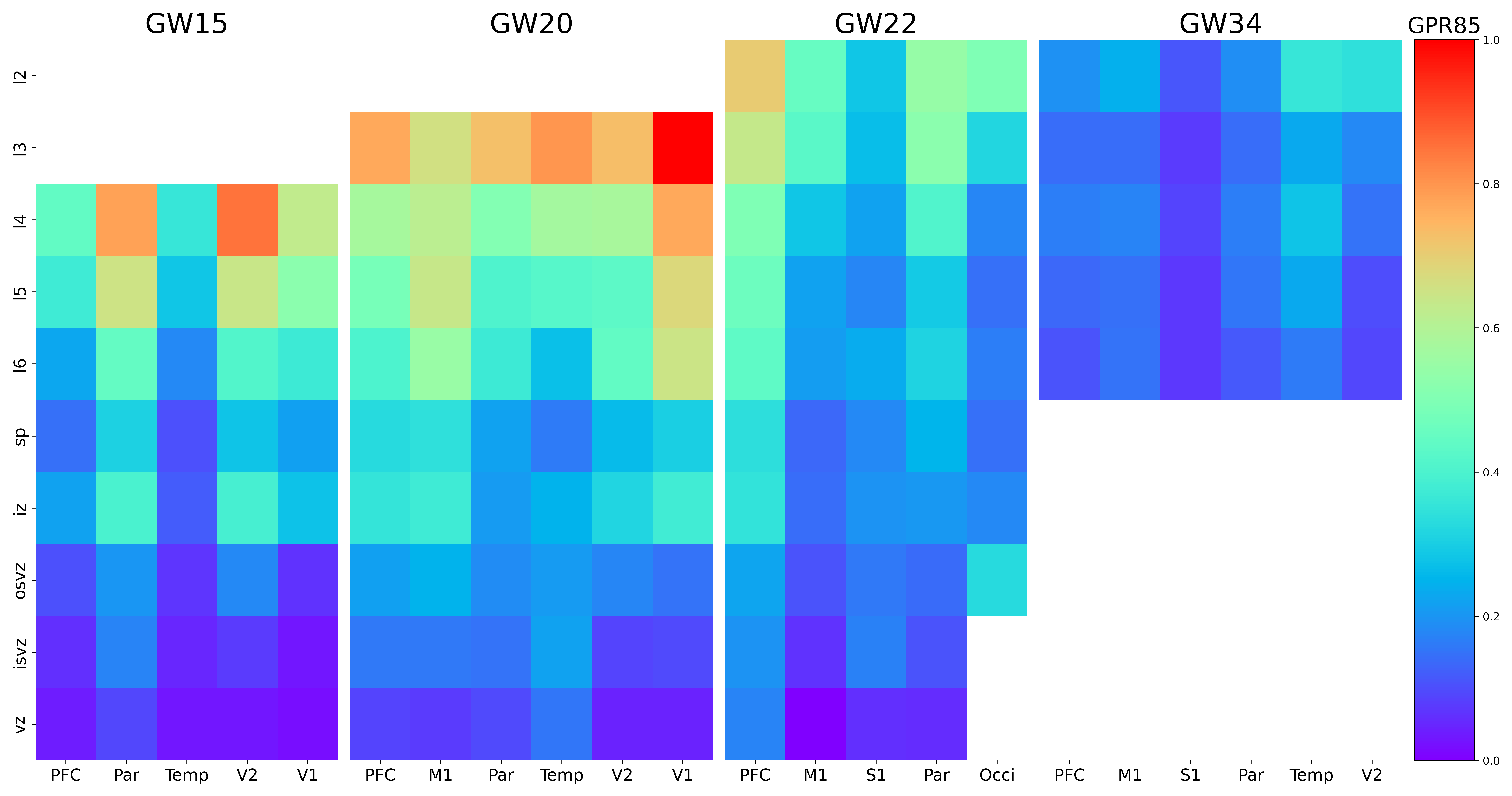

### GPRC5B.png

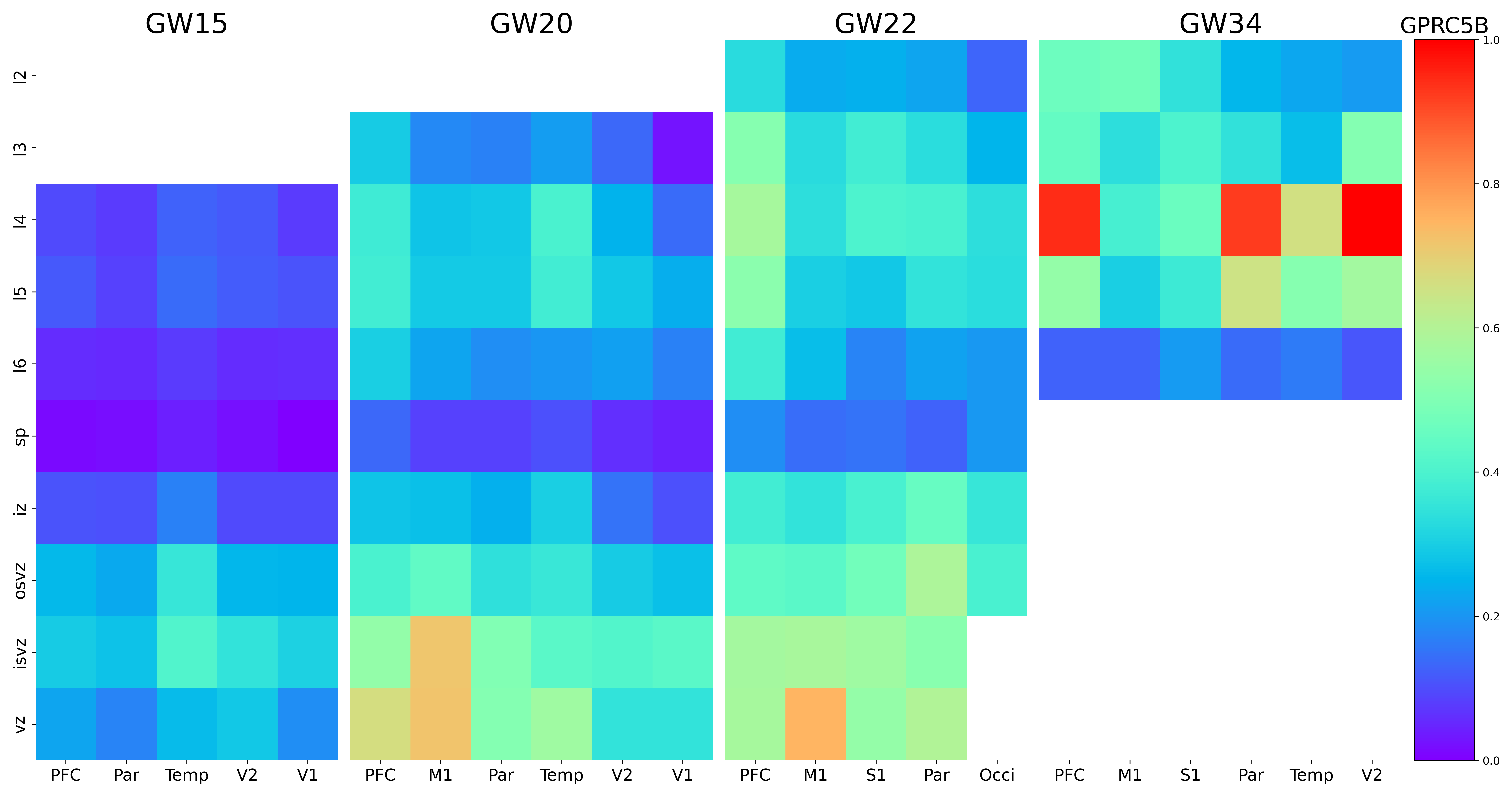

### GPRIN3.png

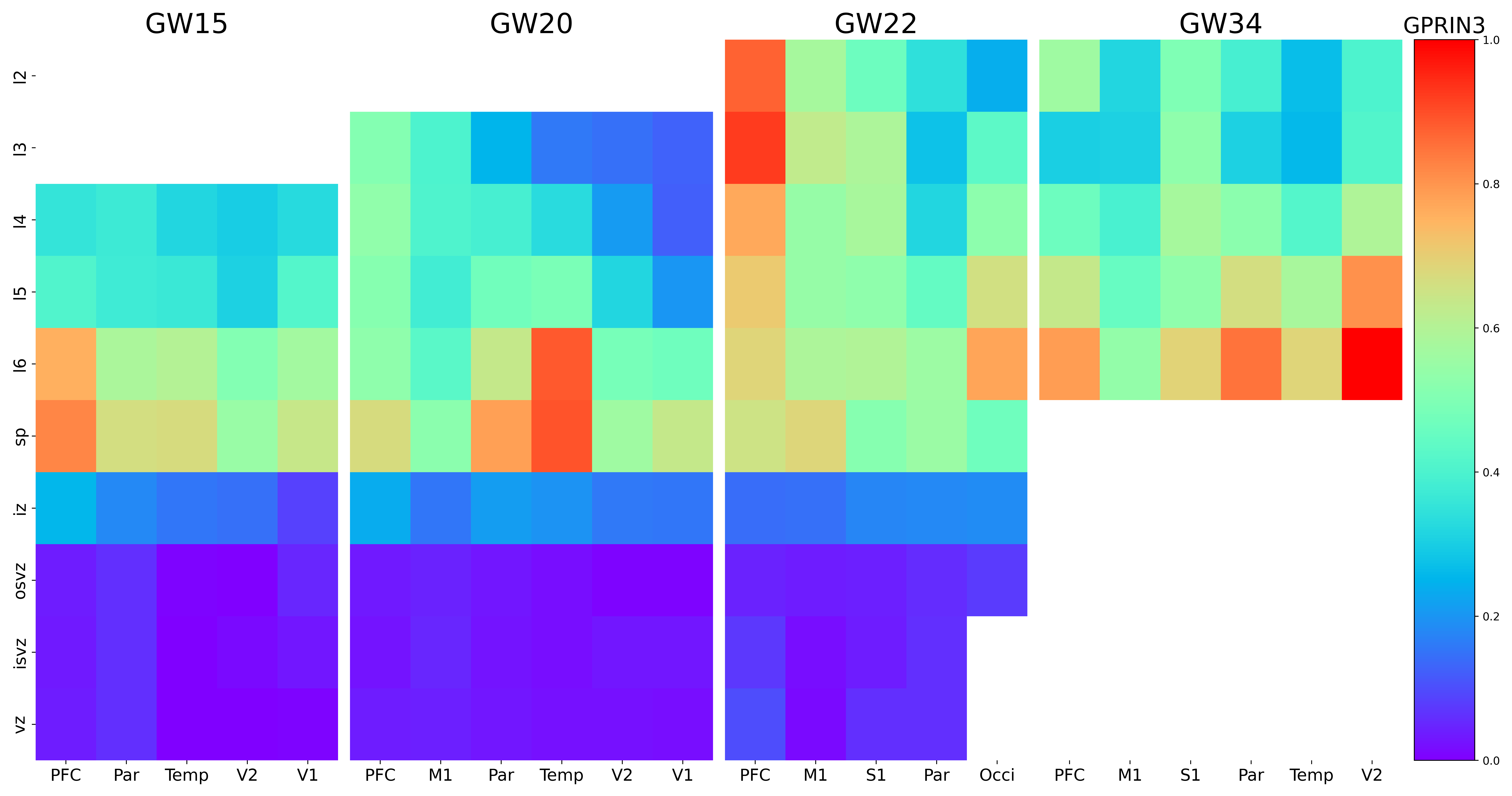

### GPX3.png

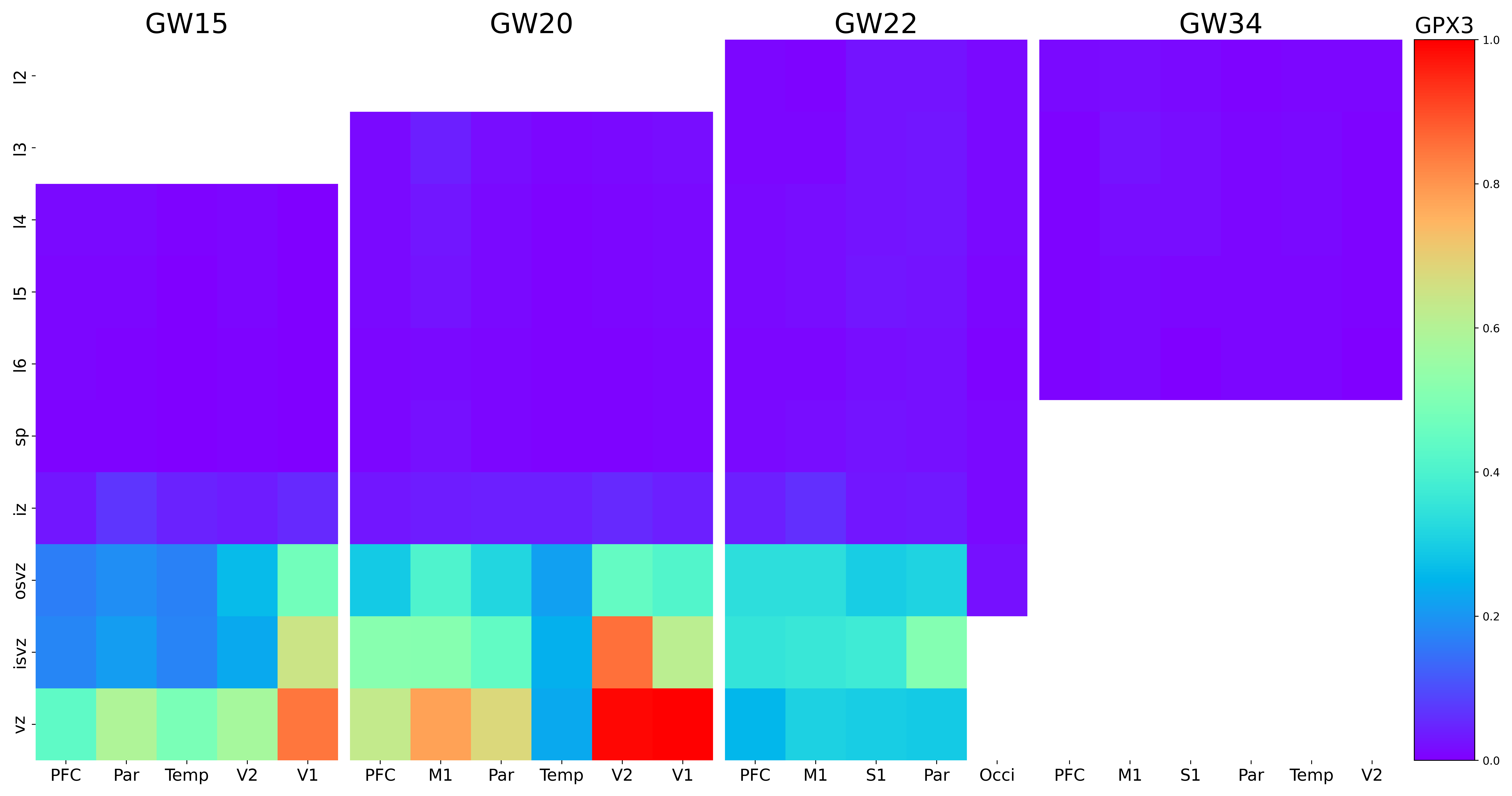

### GRAMD2B.png

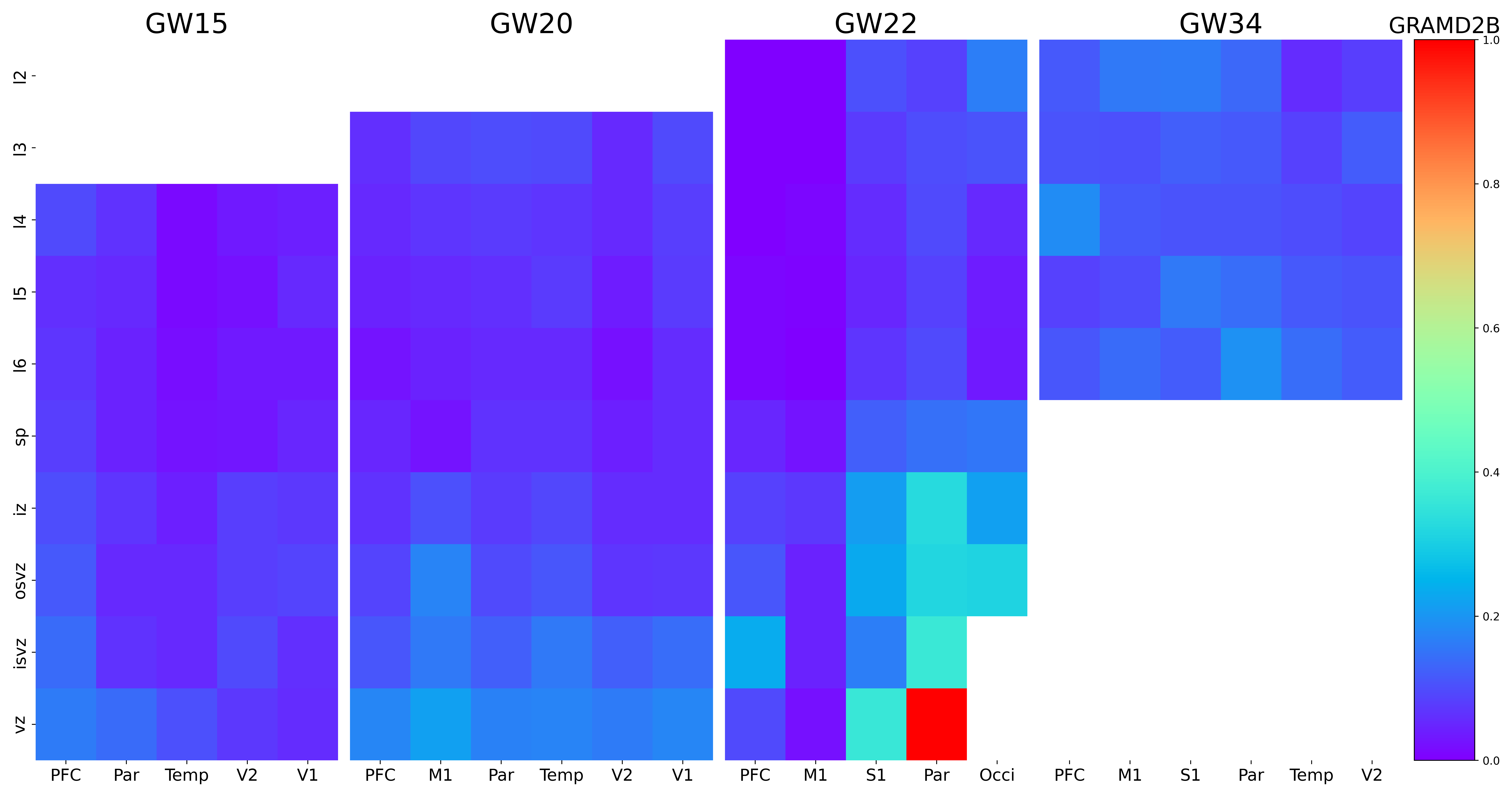

### GRIK4.png

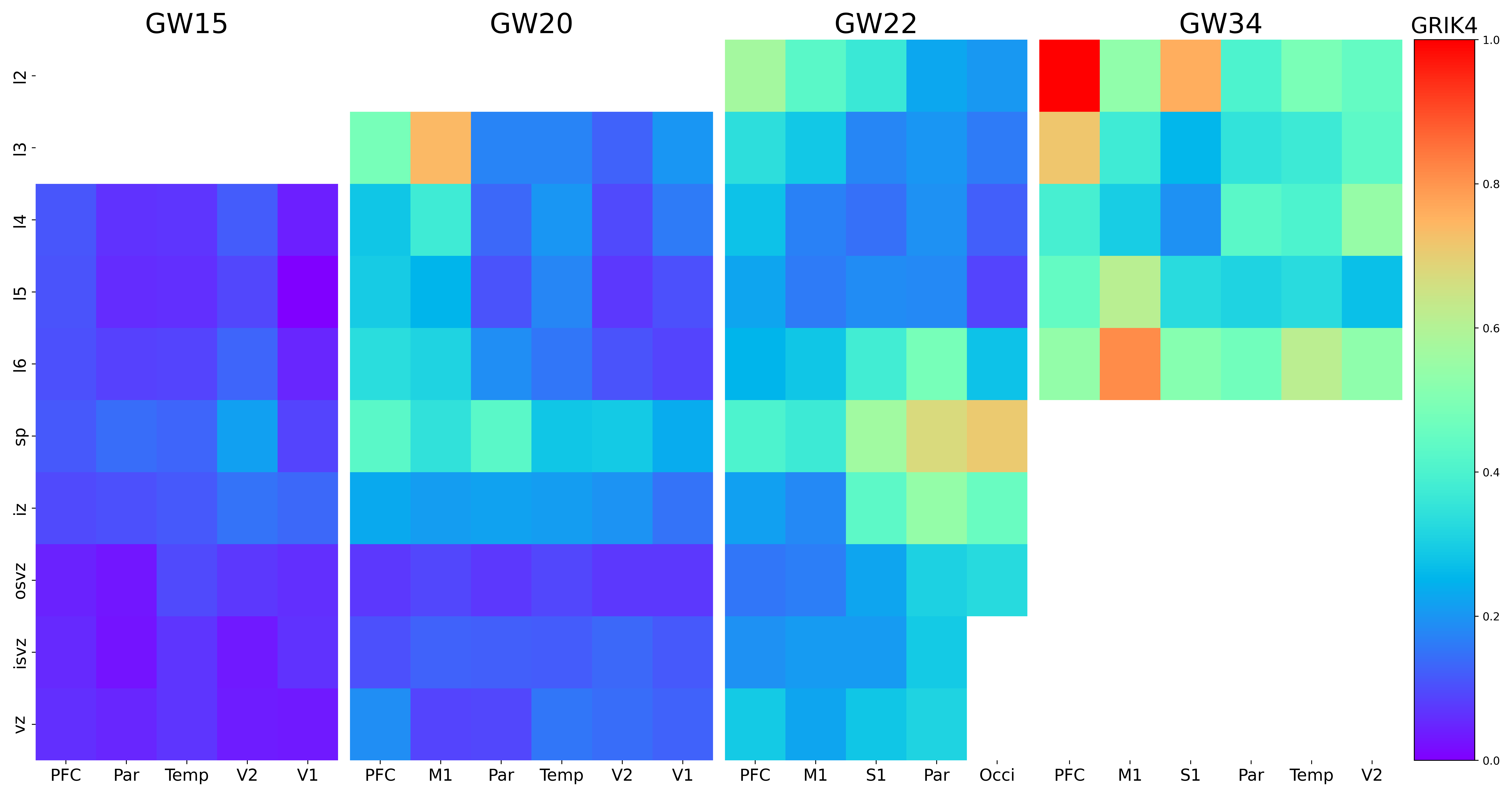

### GRM7.png

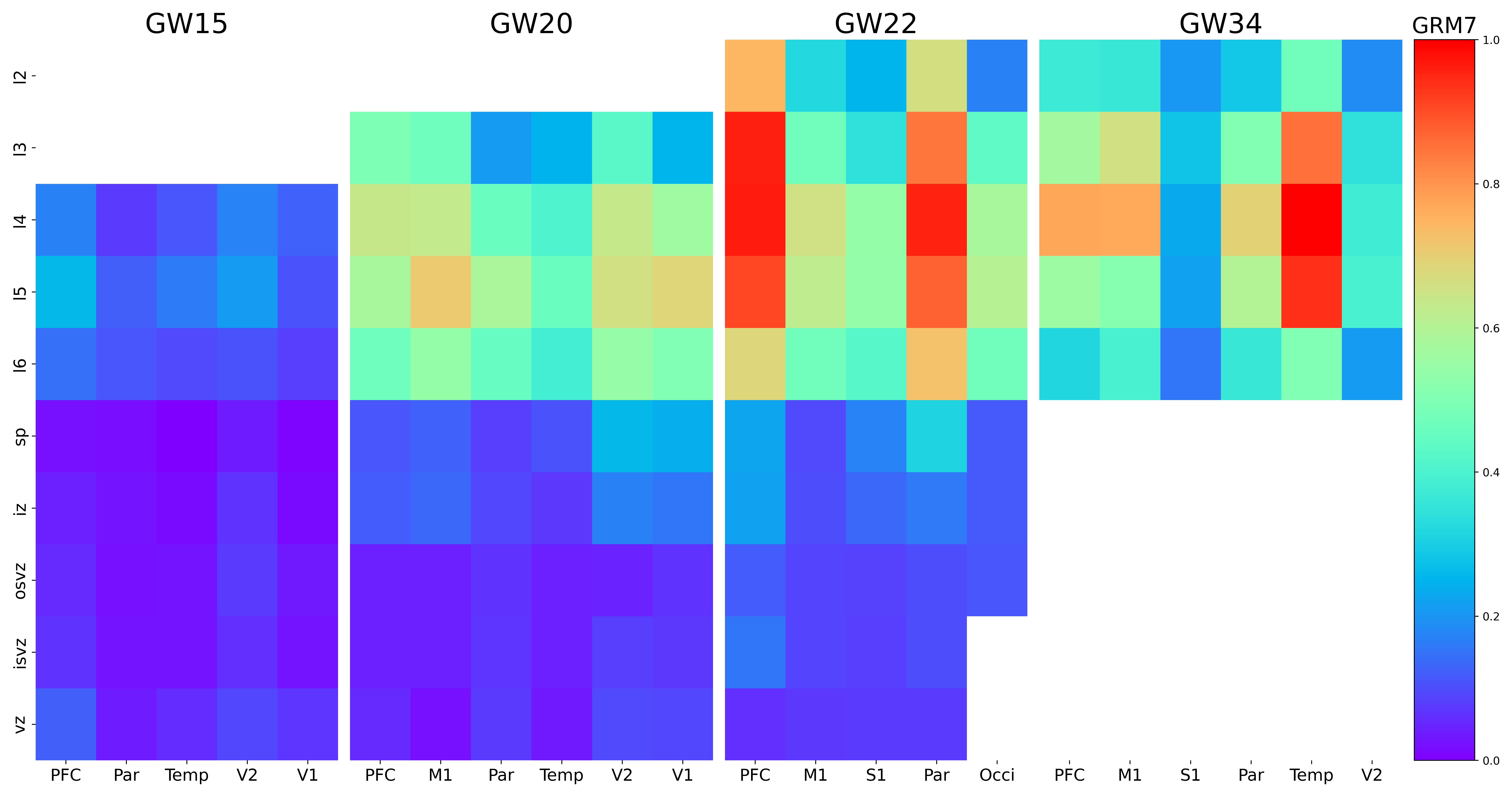

### HCRTR2.png

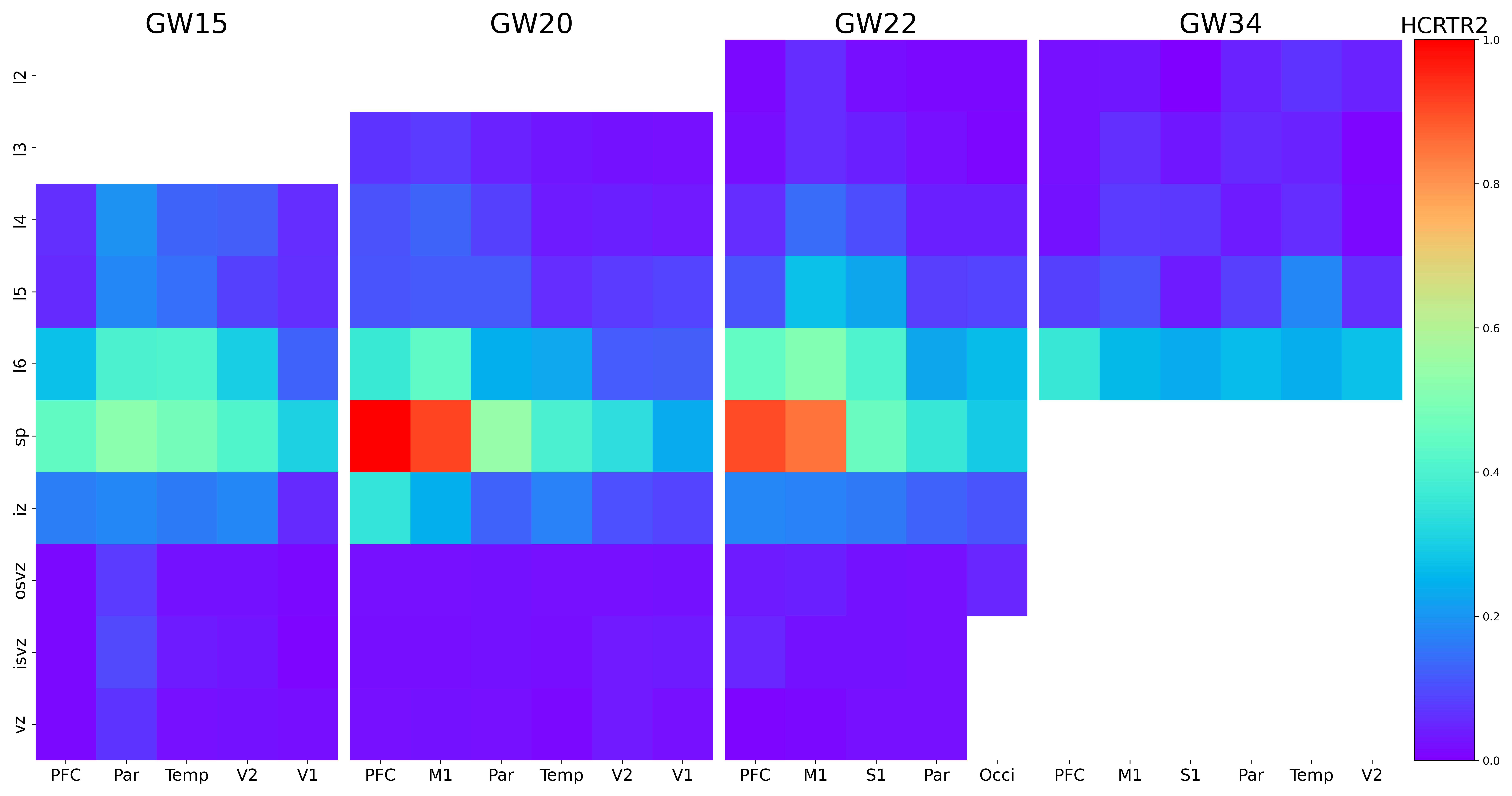

### HES1.png

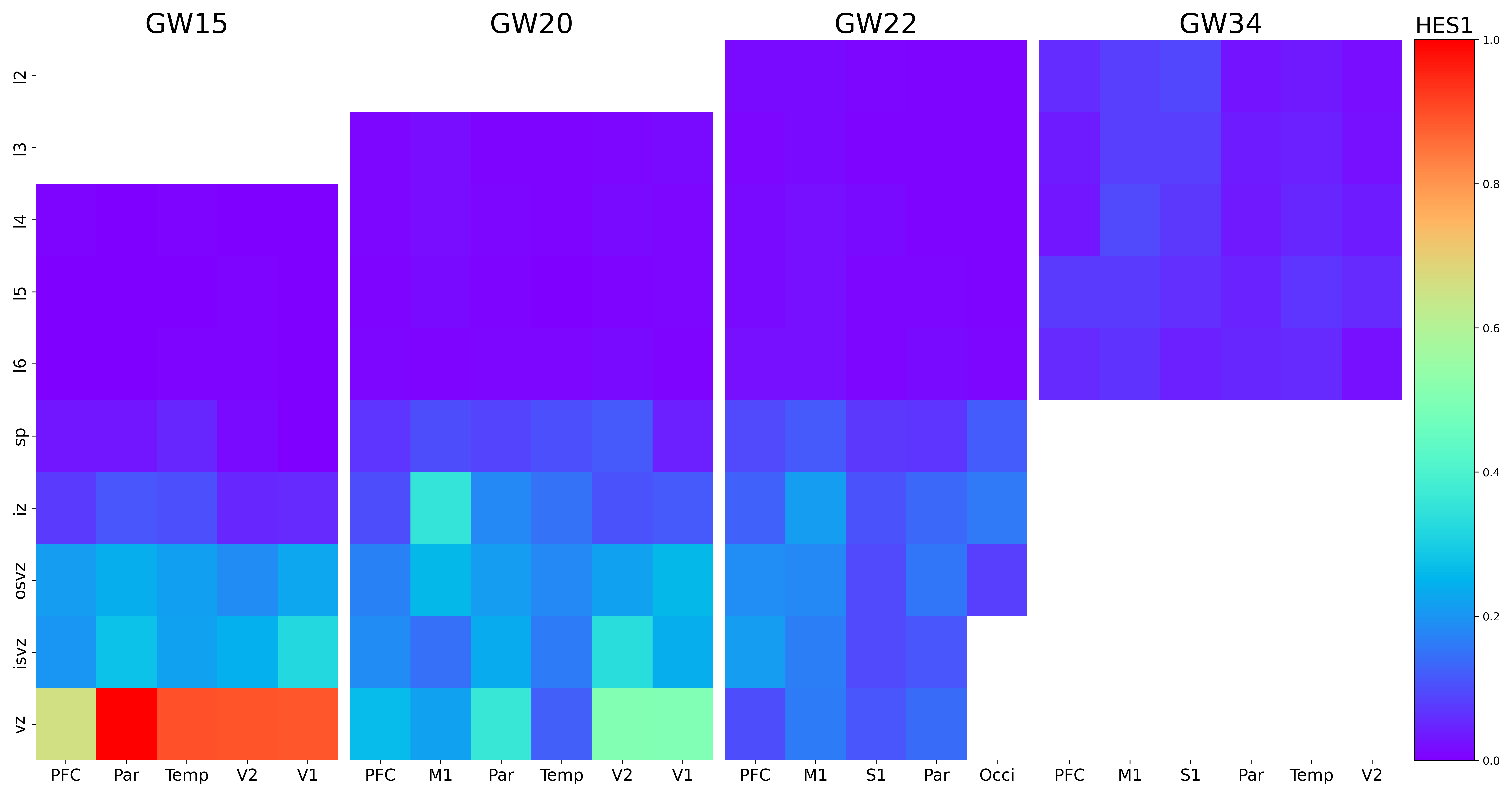

### HES5.png

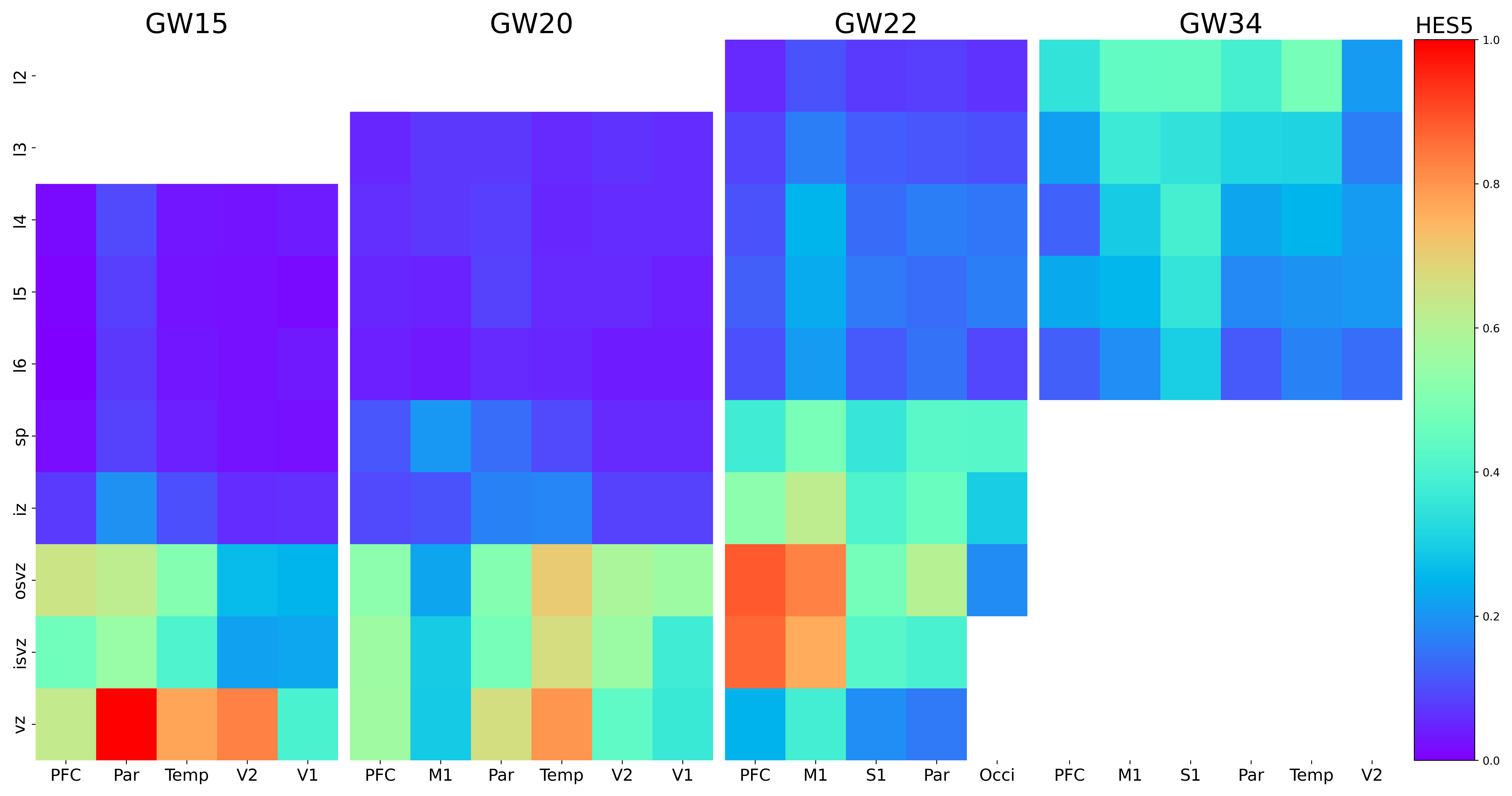

### HEY1.png

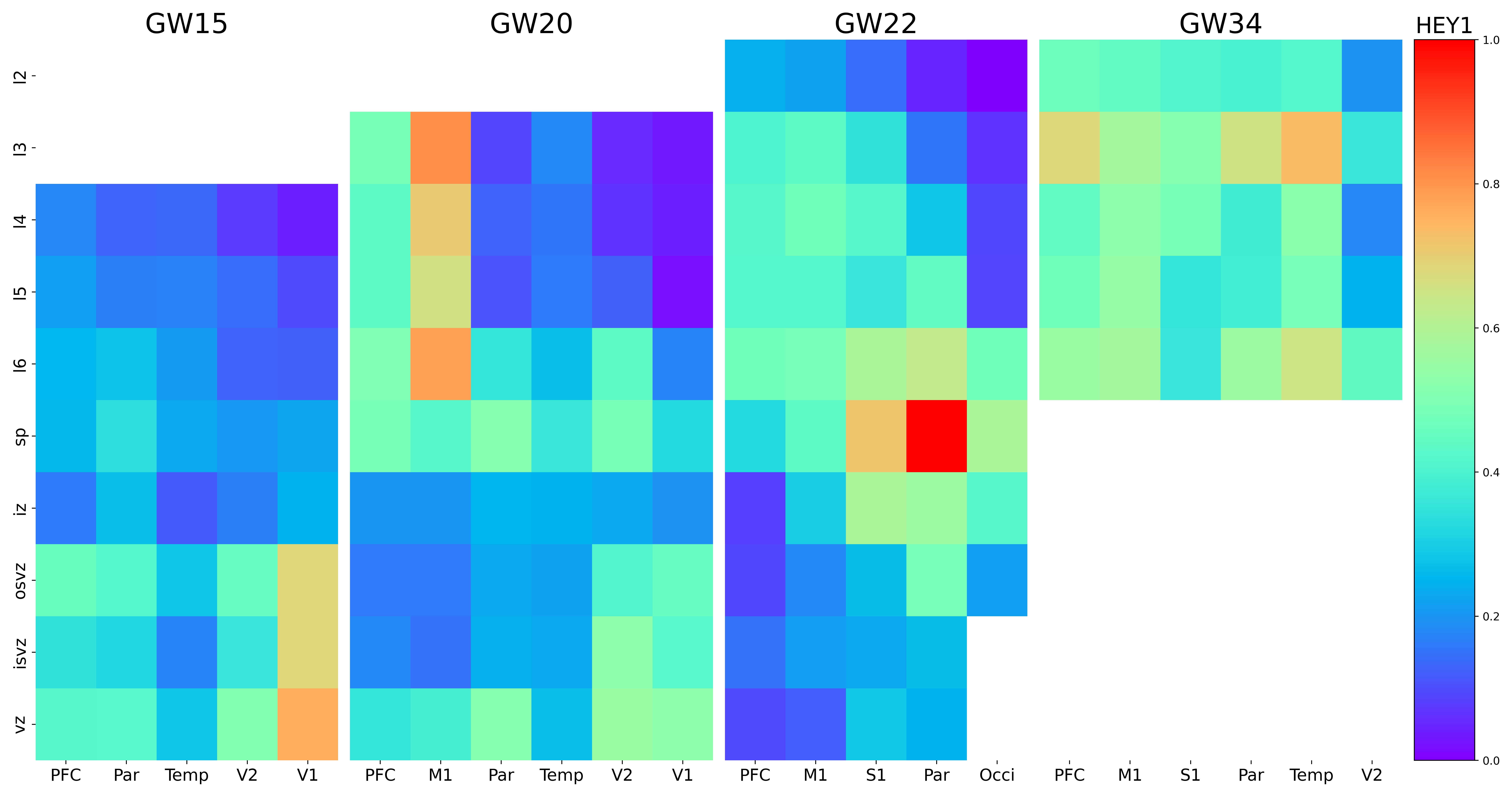

### HOPX.png

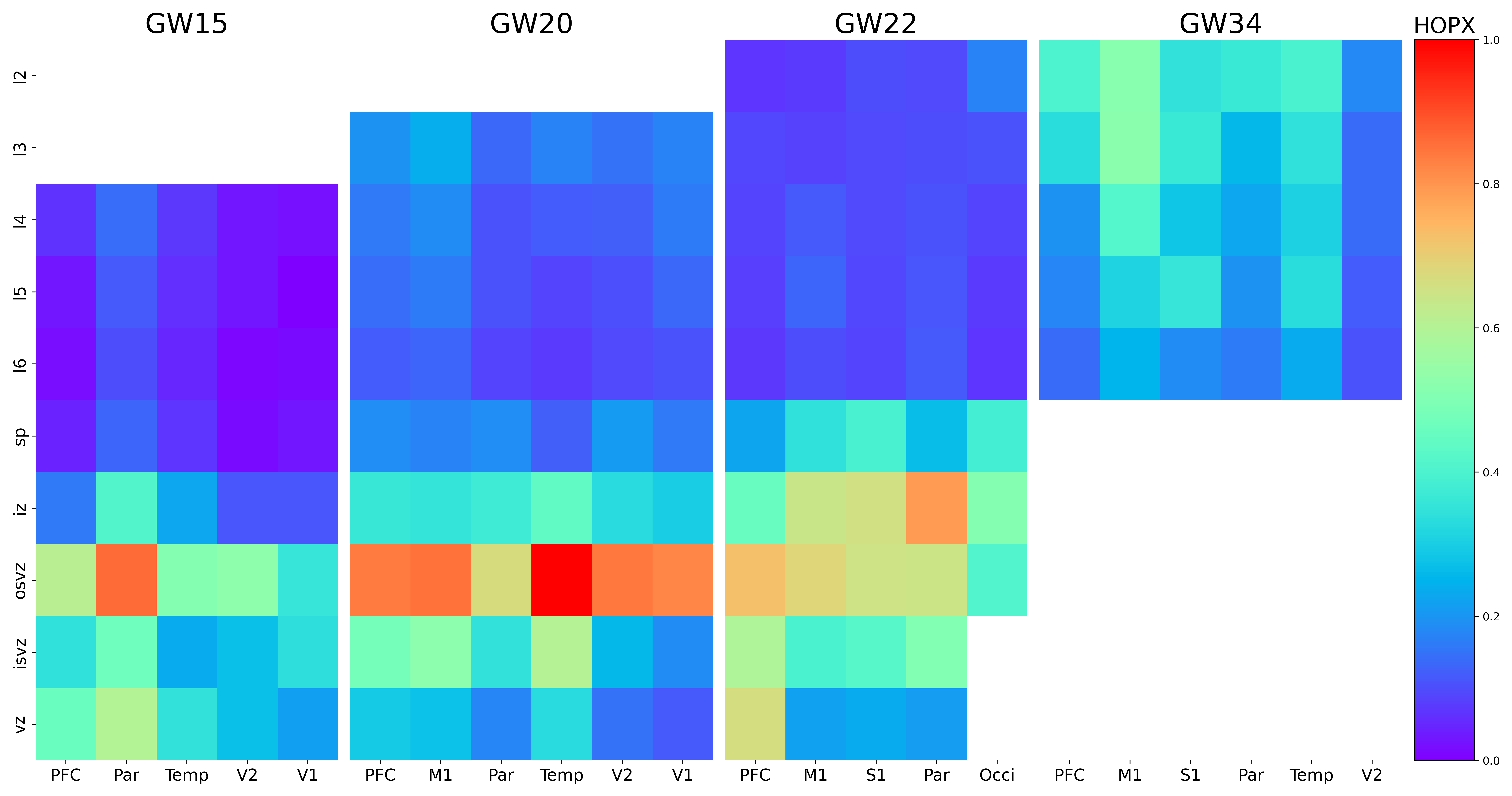

### HS3ST1.png

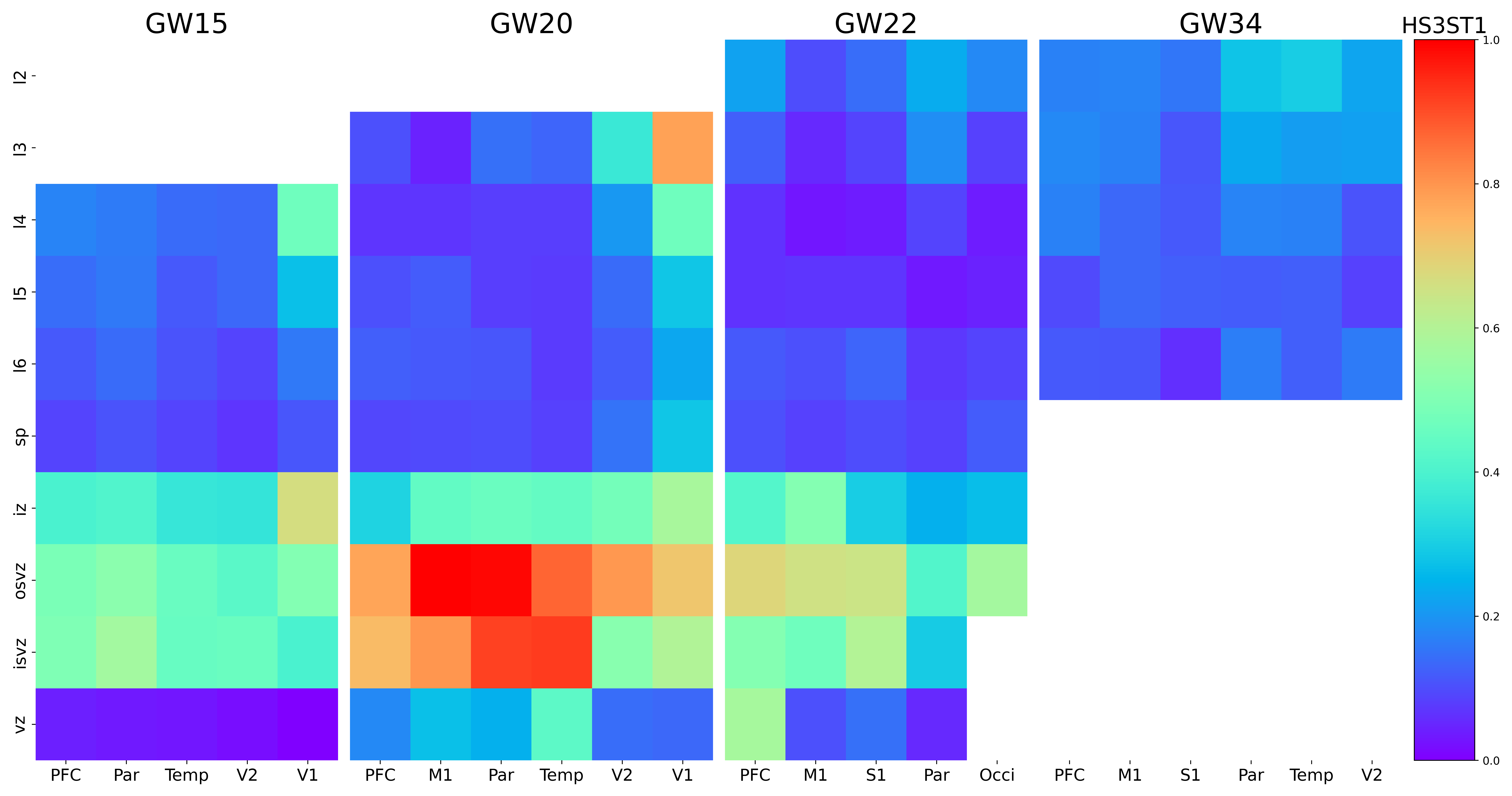

### HS3ST4.png

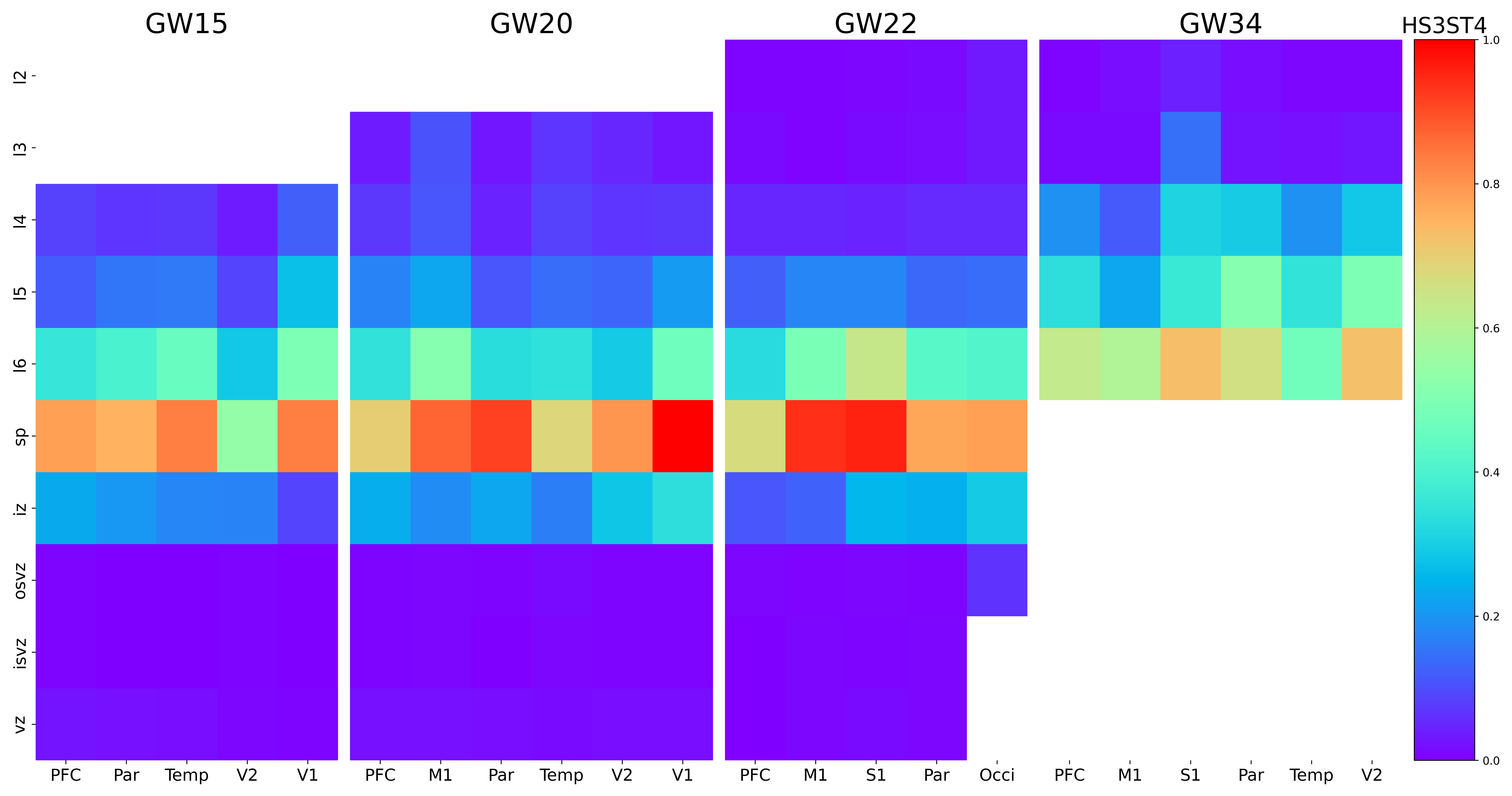

### HSPA12A.png

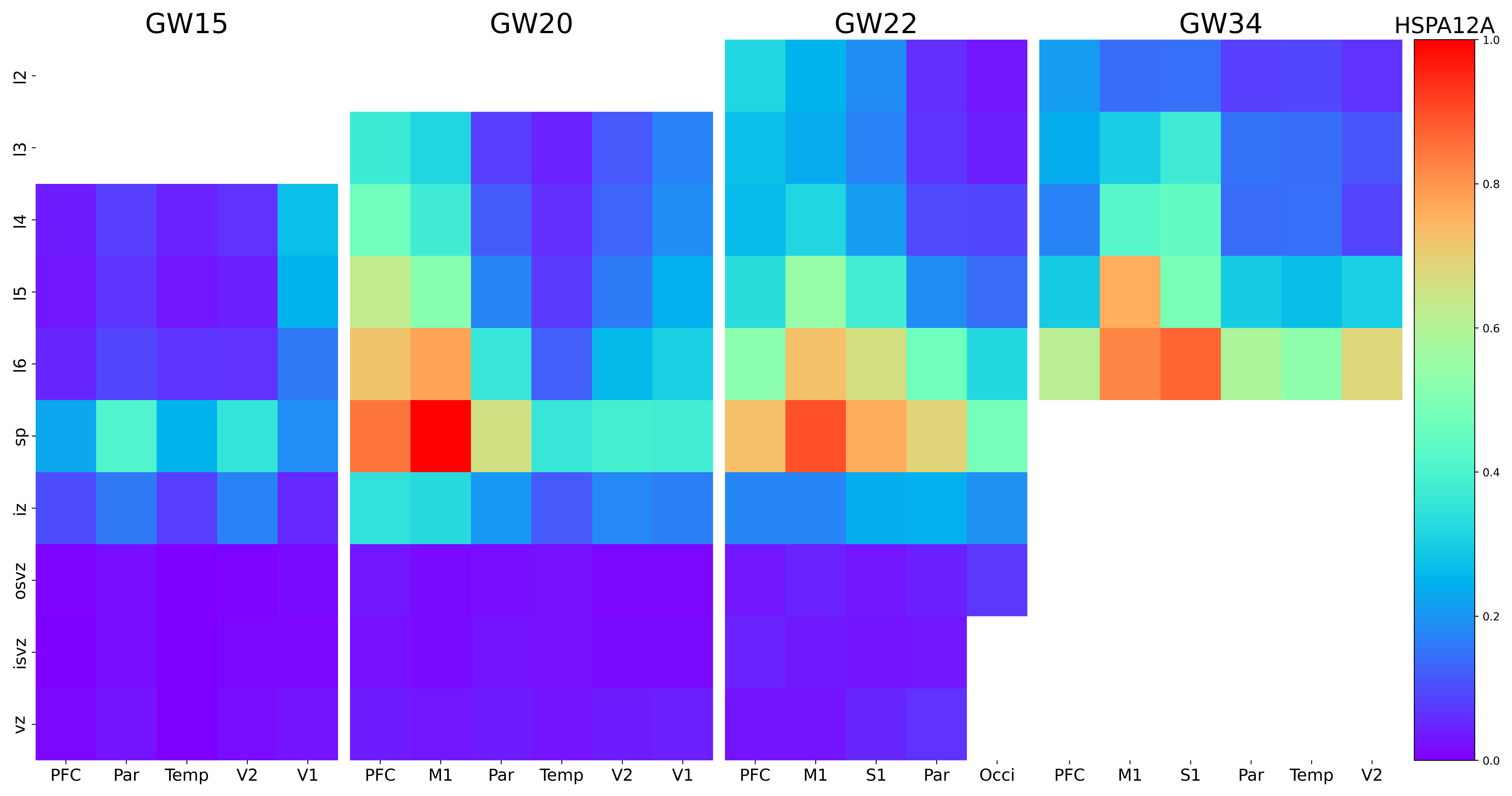

### HTR2A.png

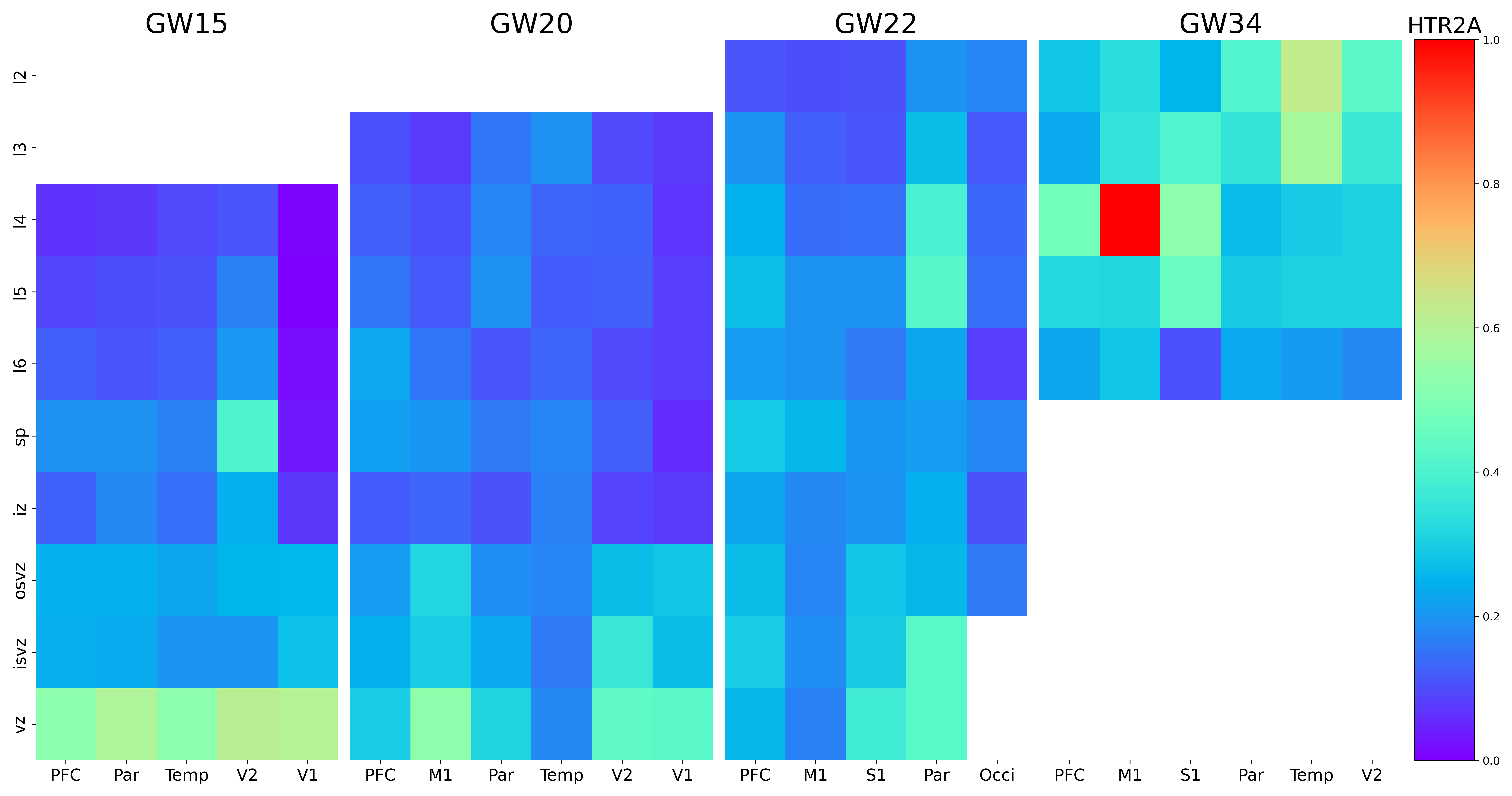

### IGFBPL1.png

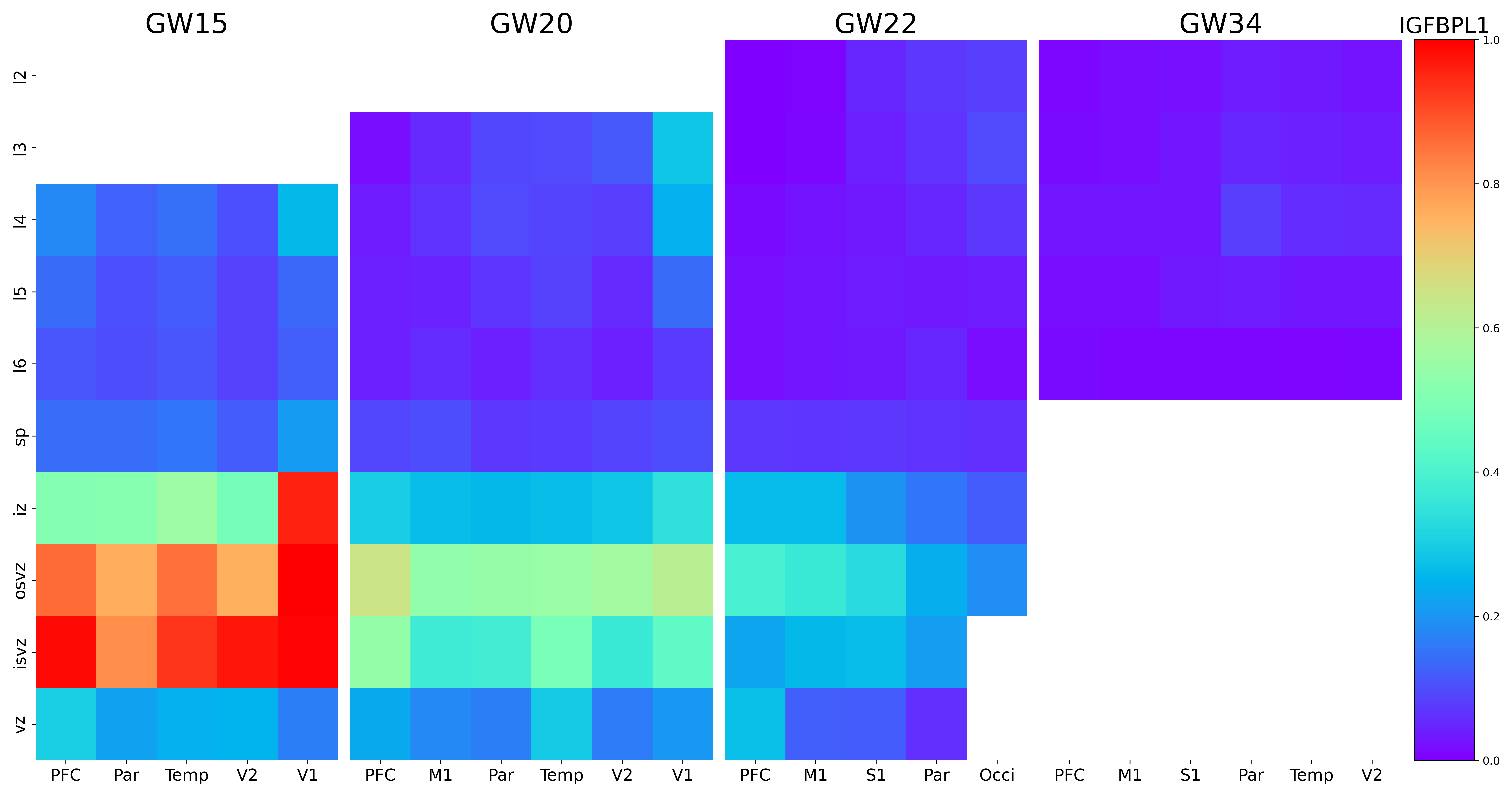

### IGSF11.png

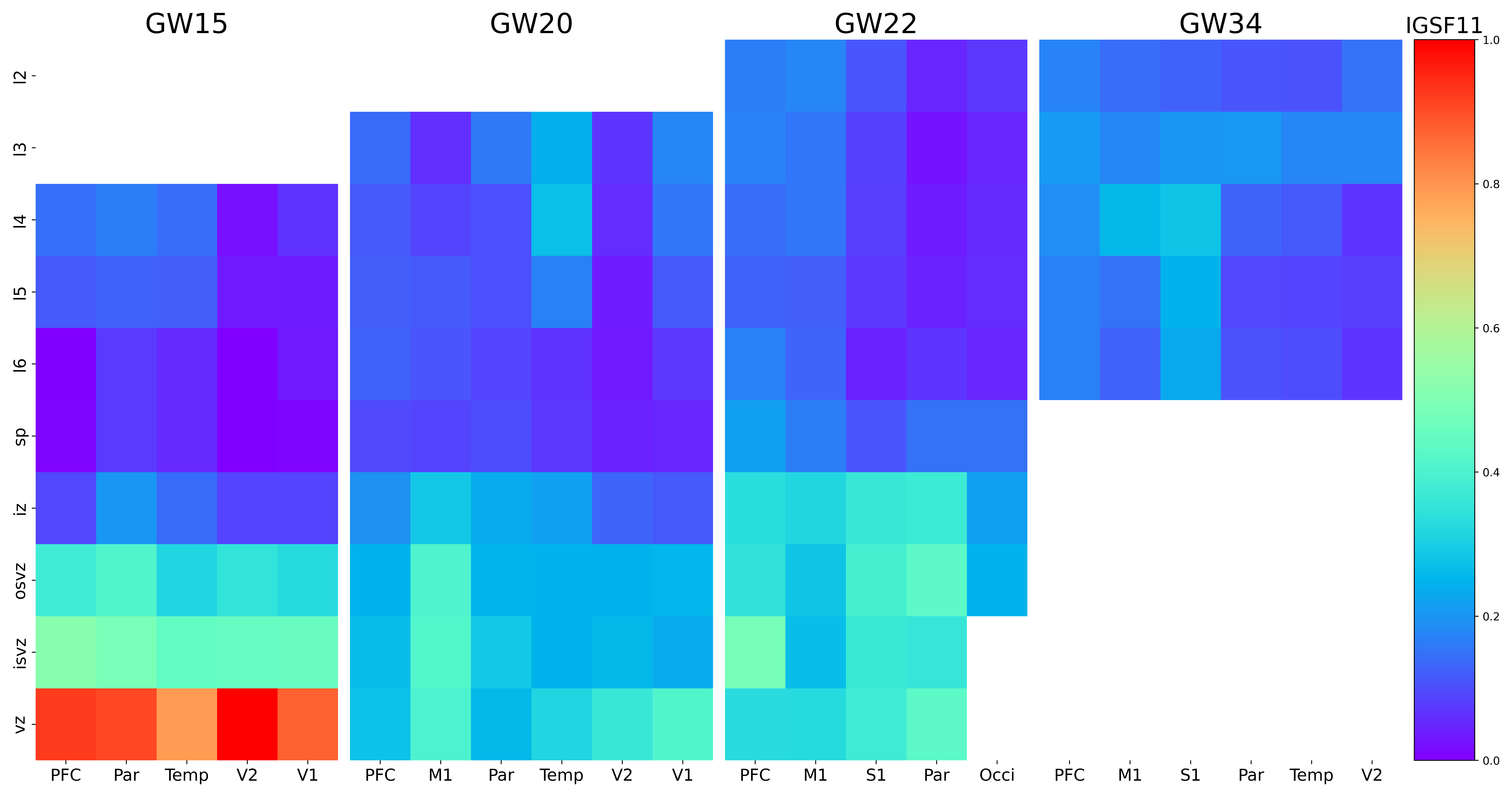

### INPP4B.png

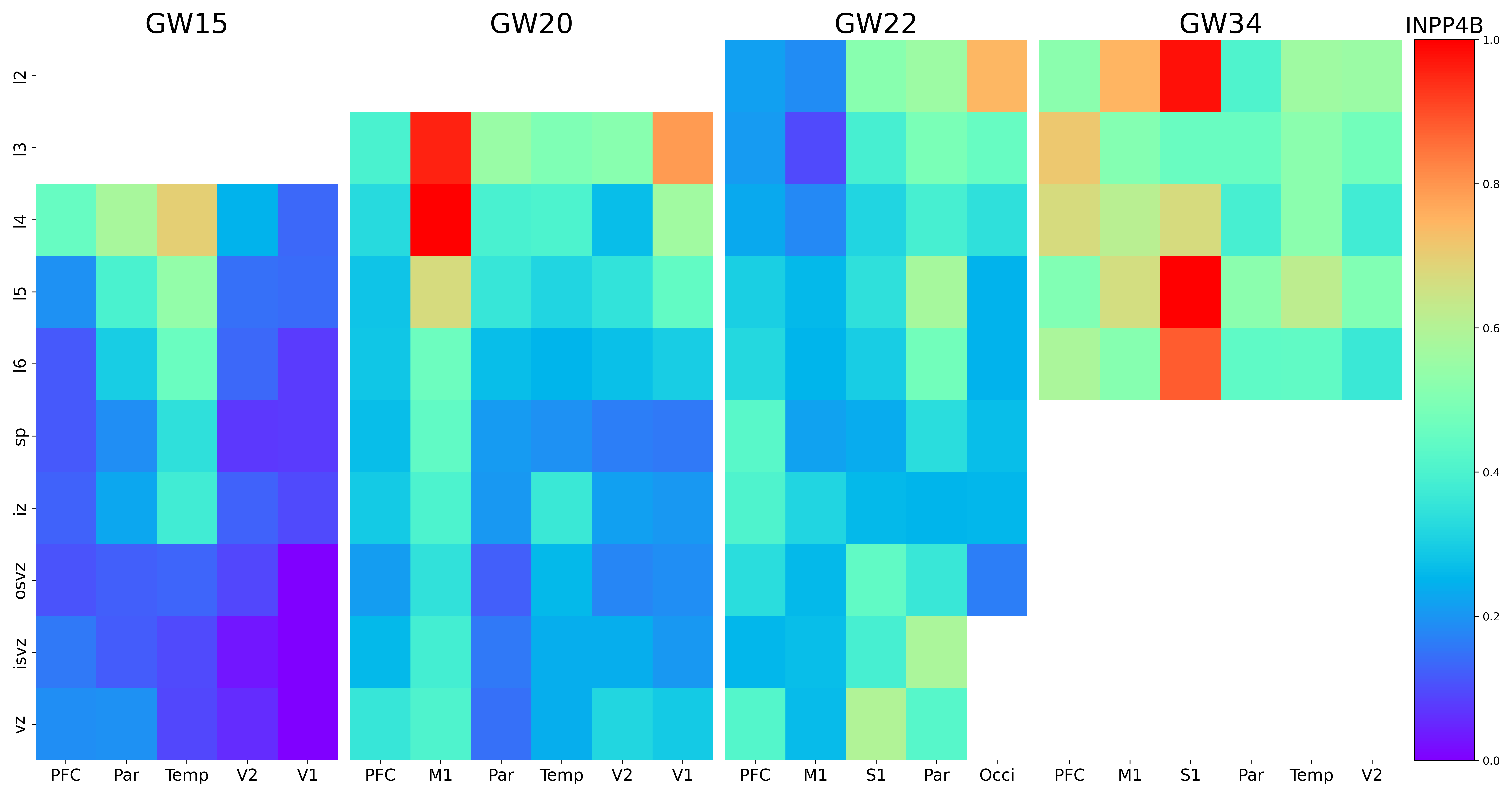

### INSM1.png

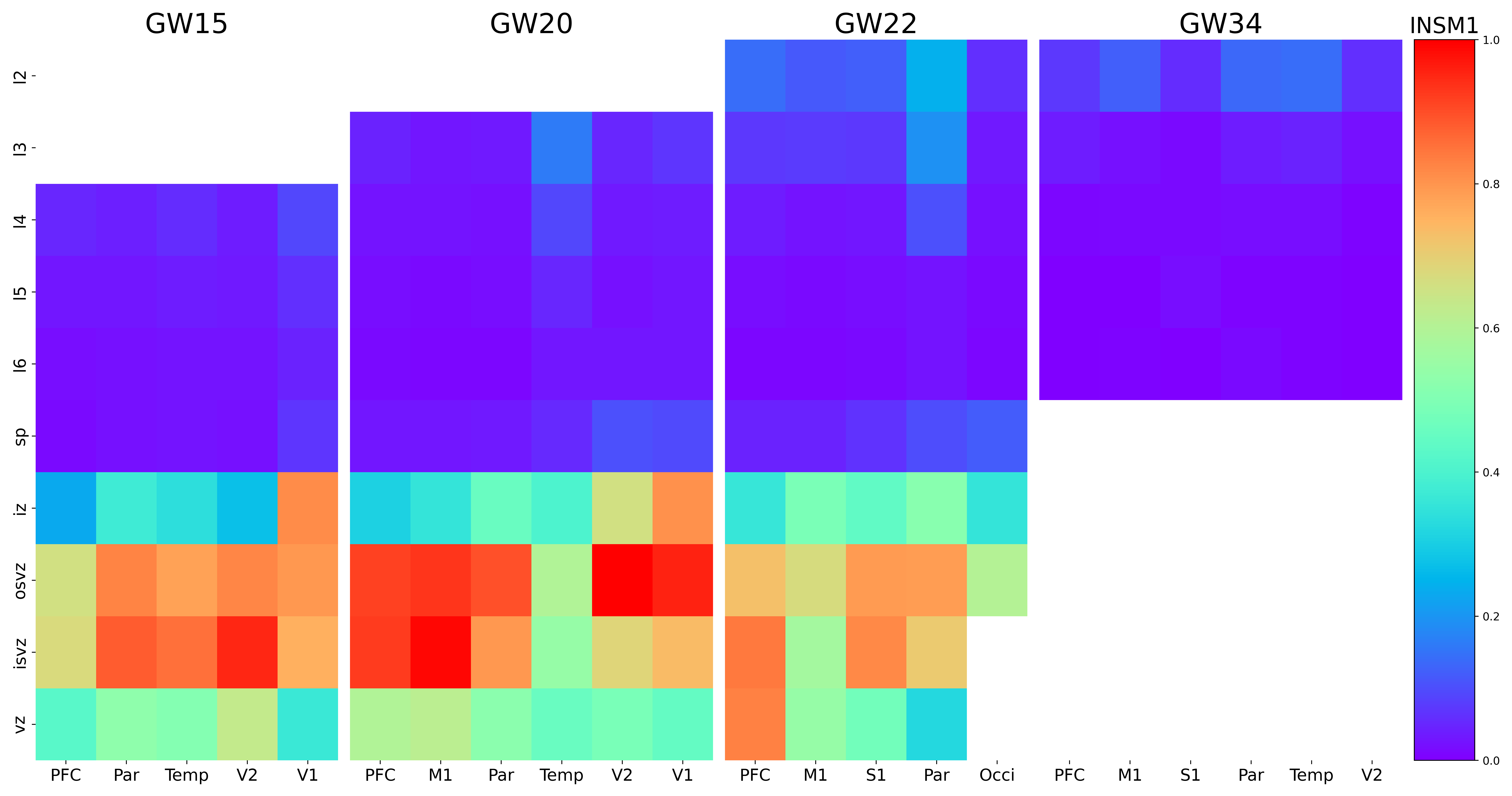

### ITGA2.png

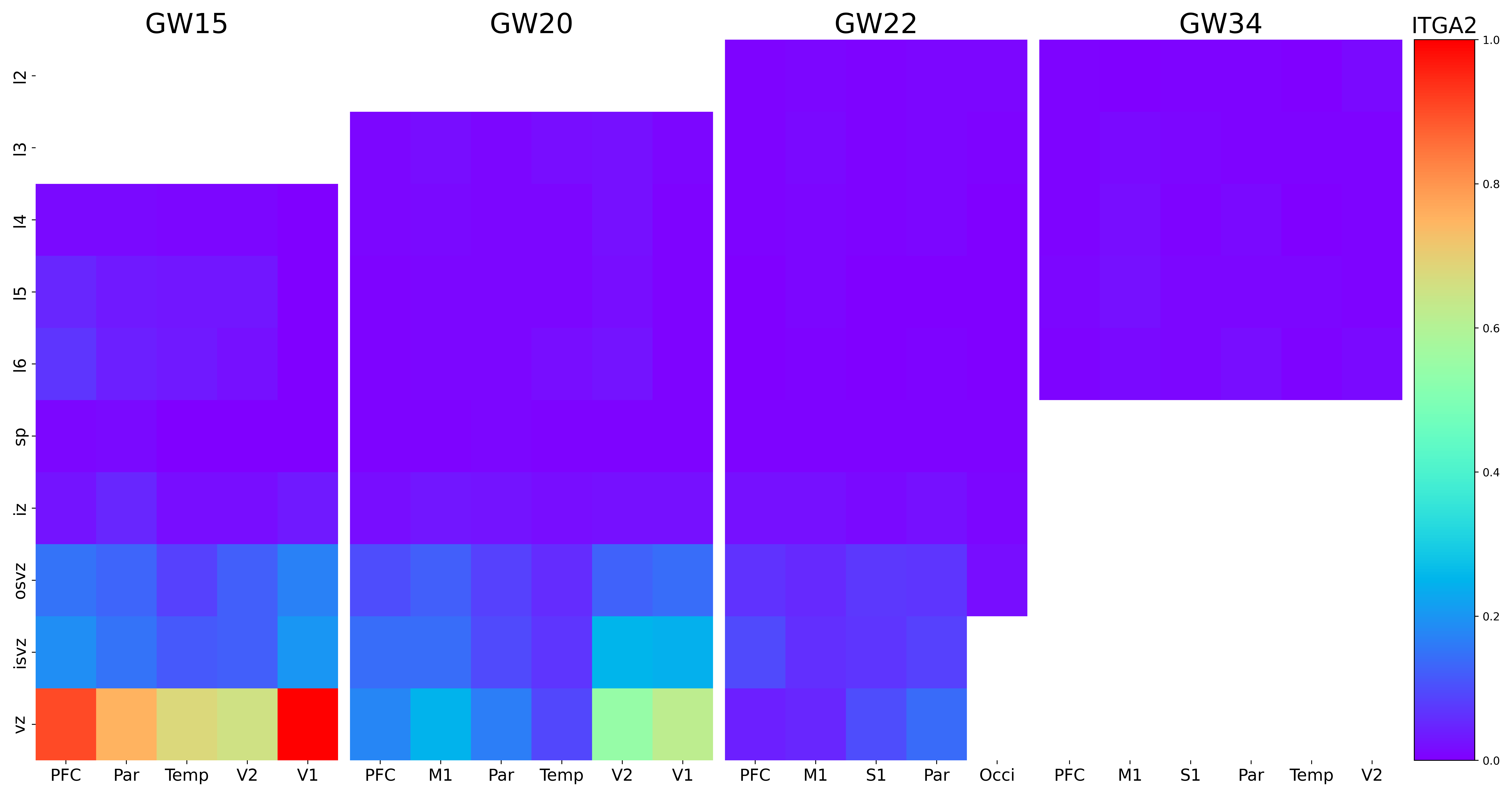

### ITGB5.png

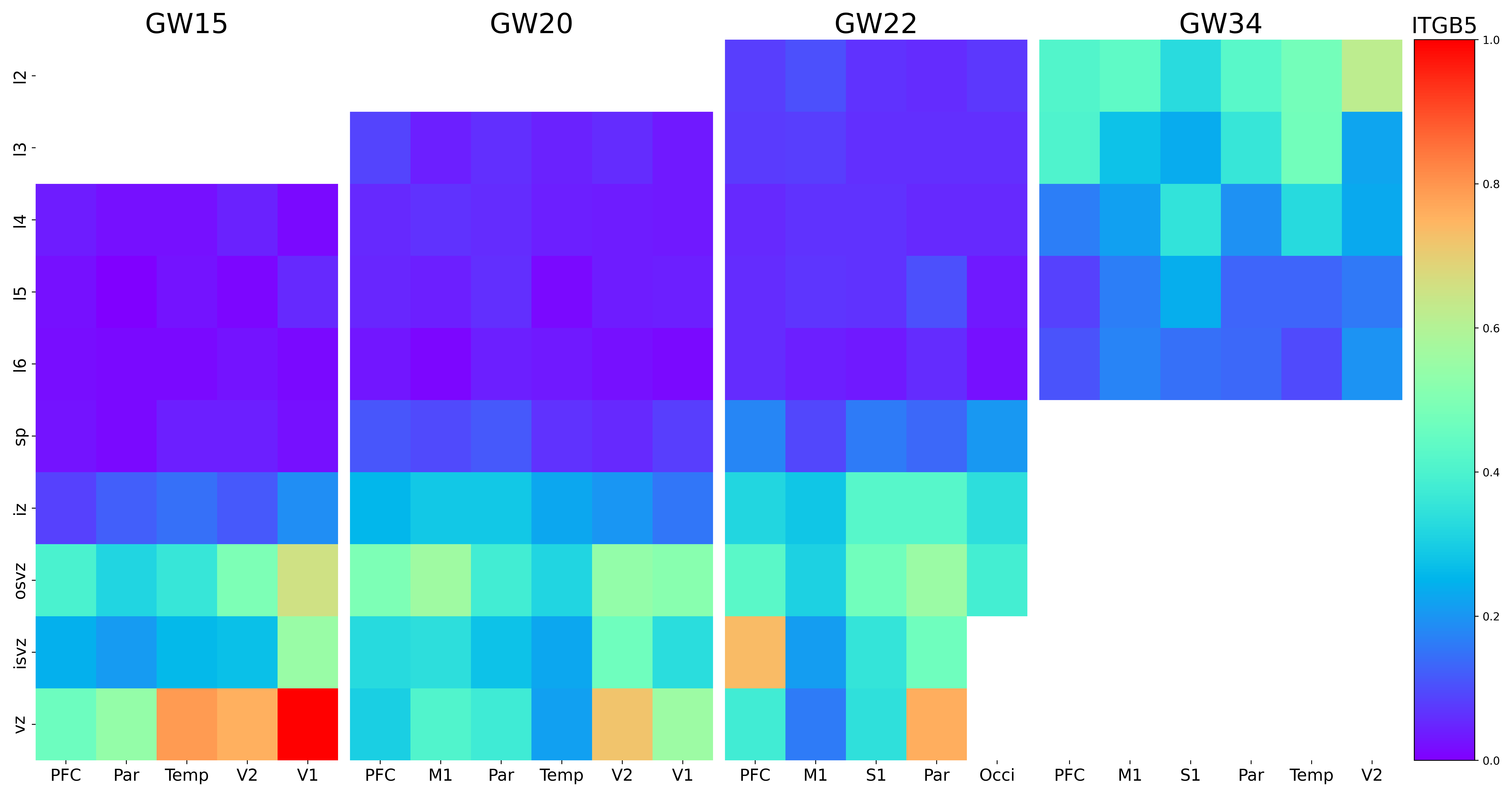

### ITPR1.png

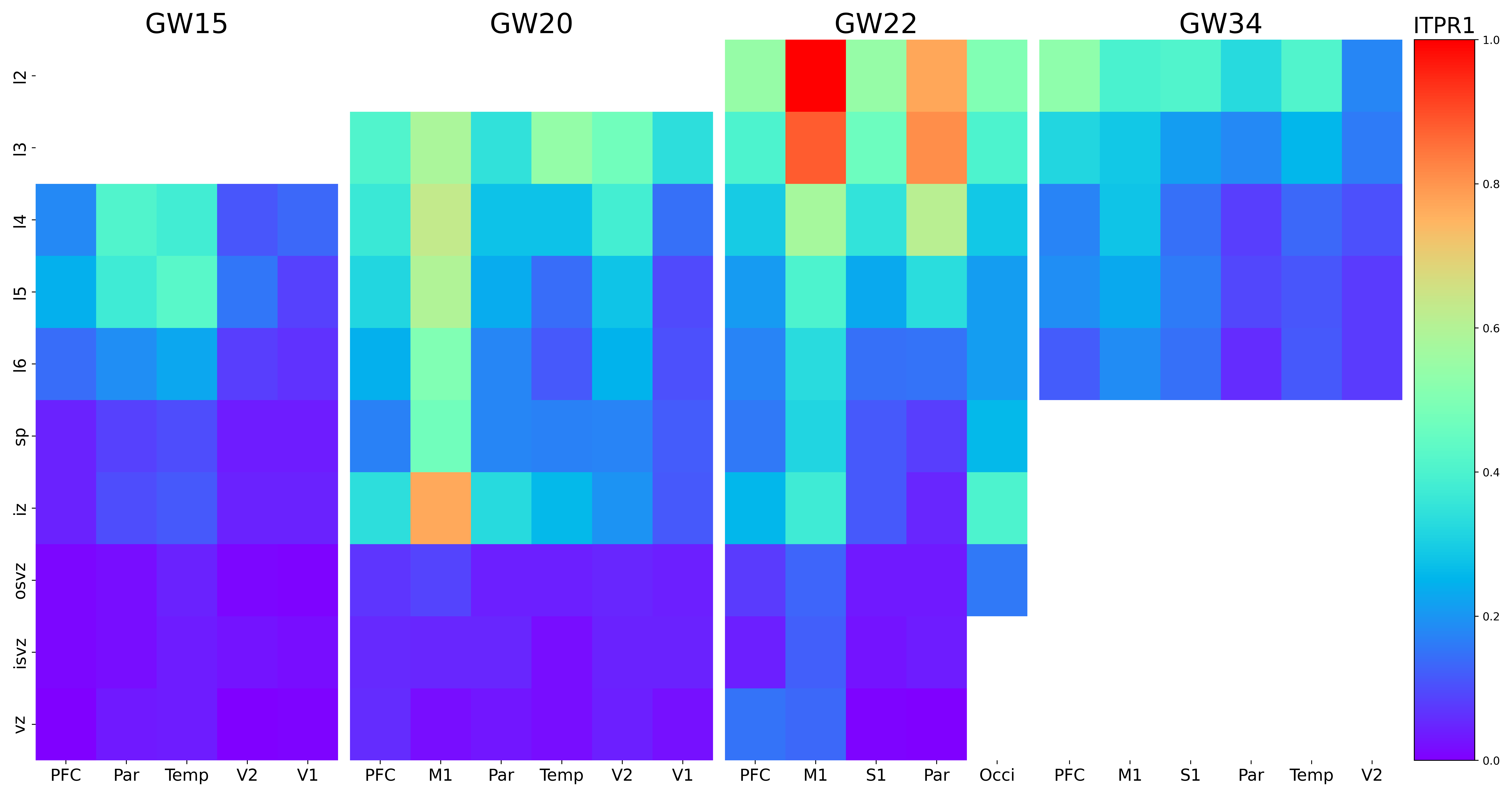

### KCNH5.png

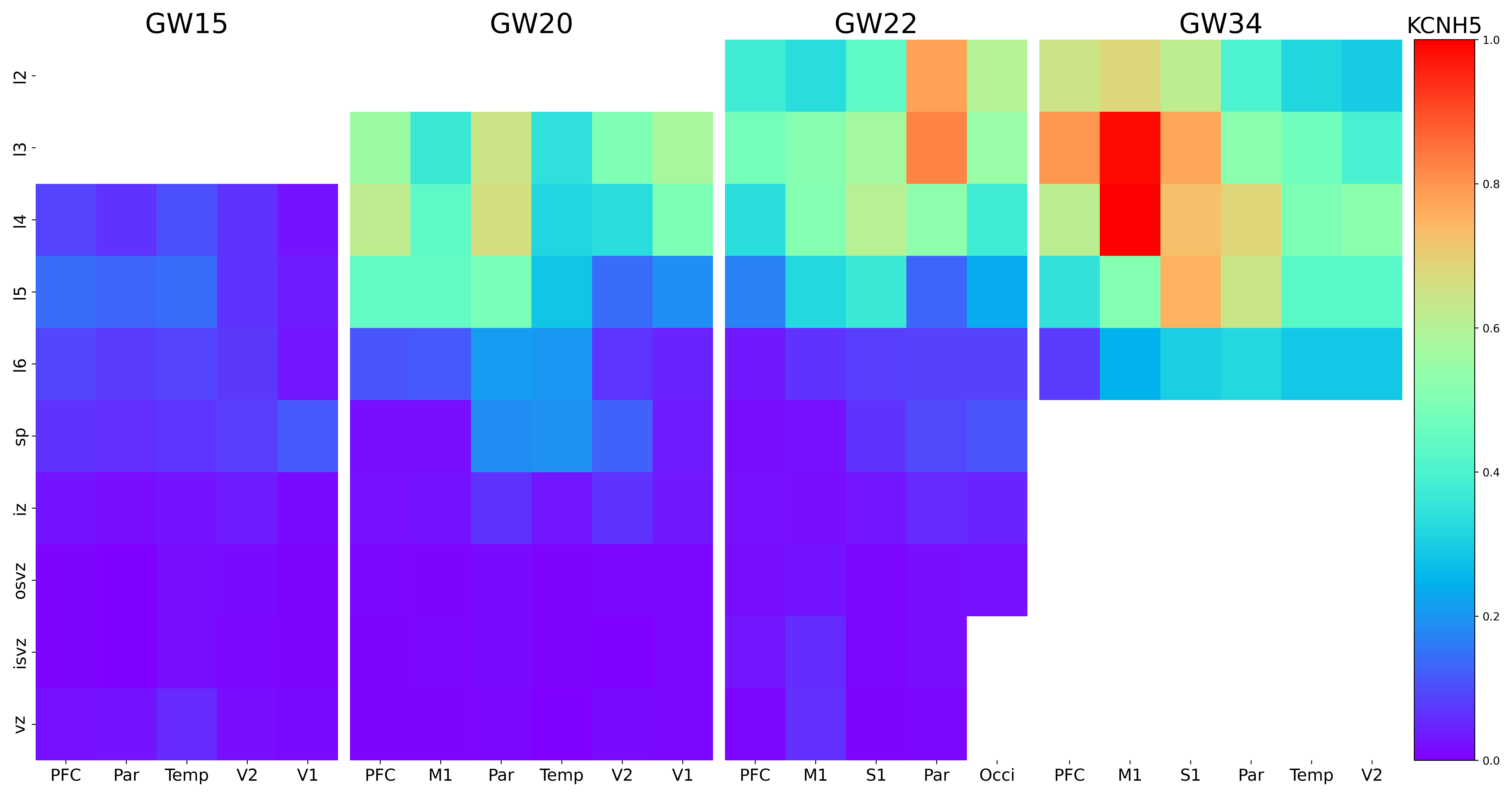

### KCNJ6.png

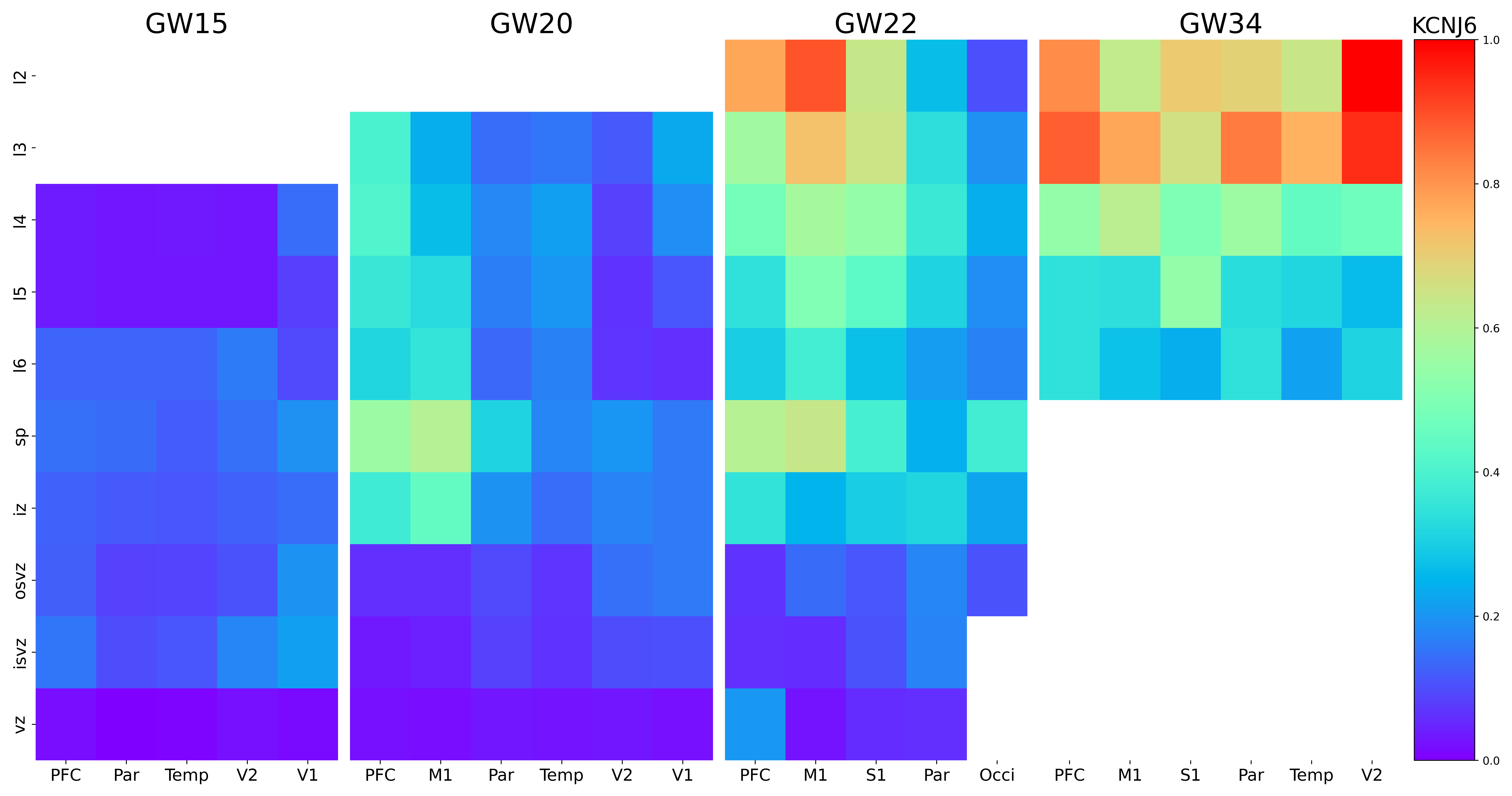

### KCNJ16.png

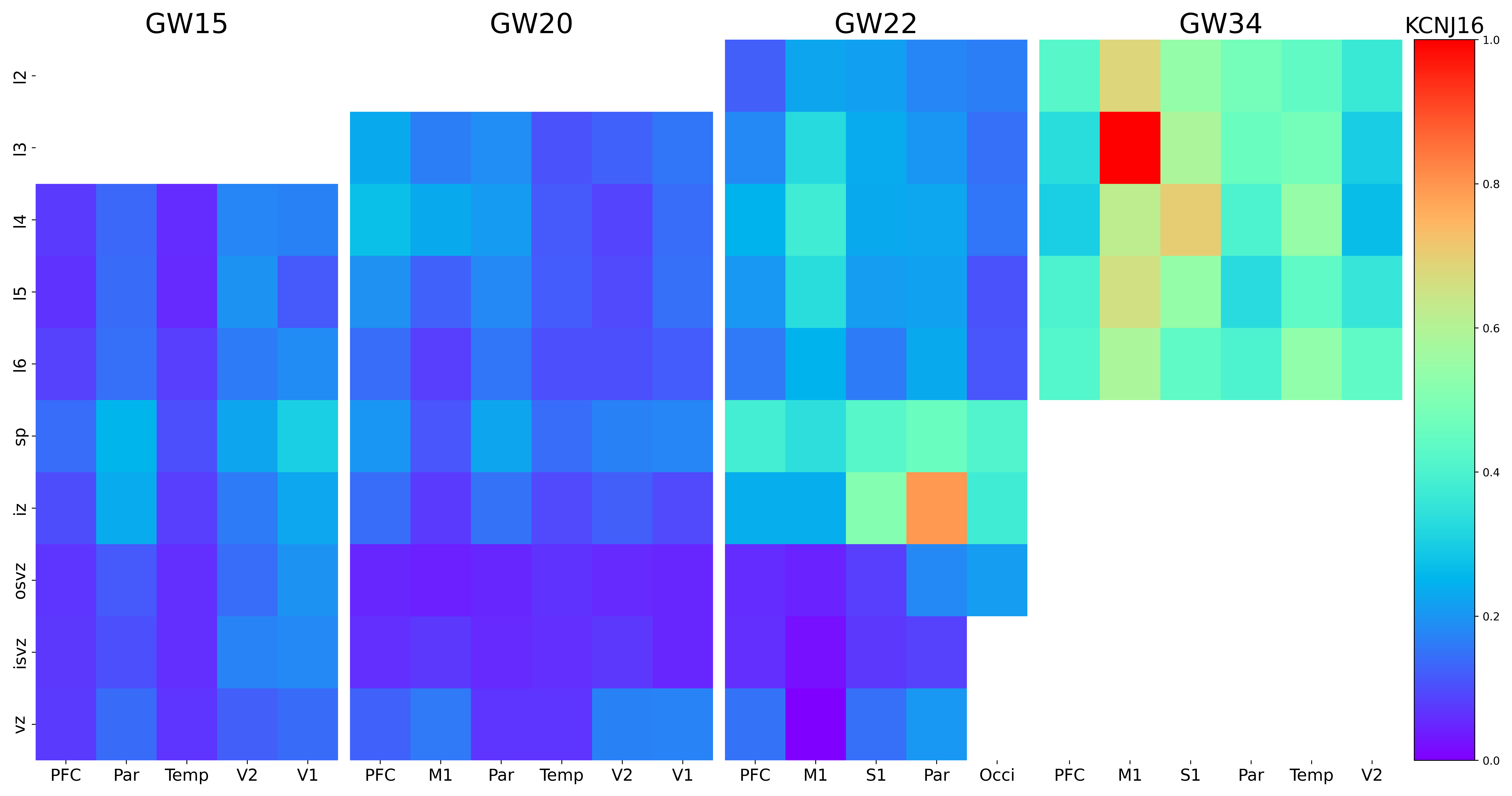

### KCNQ5.png

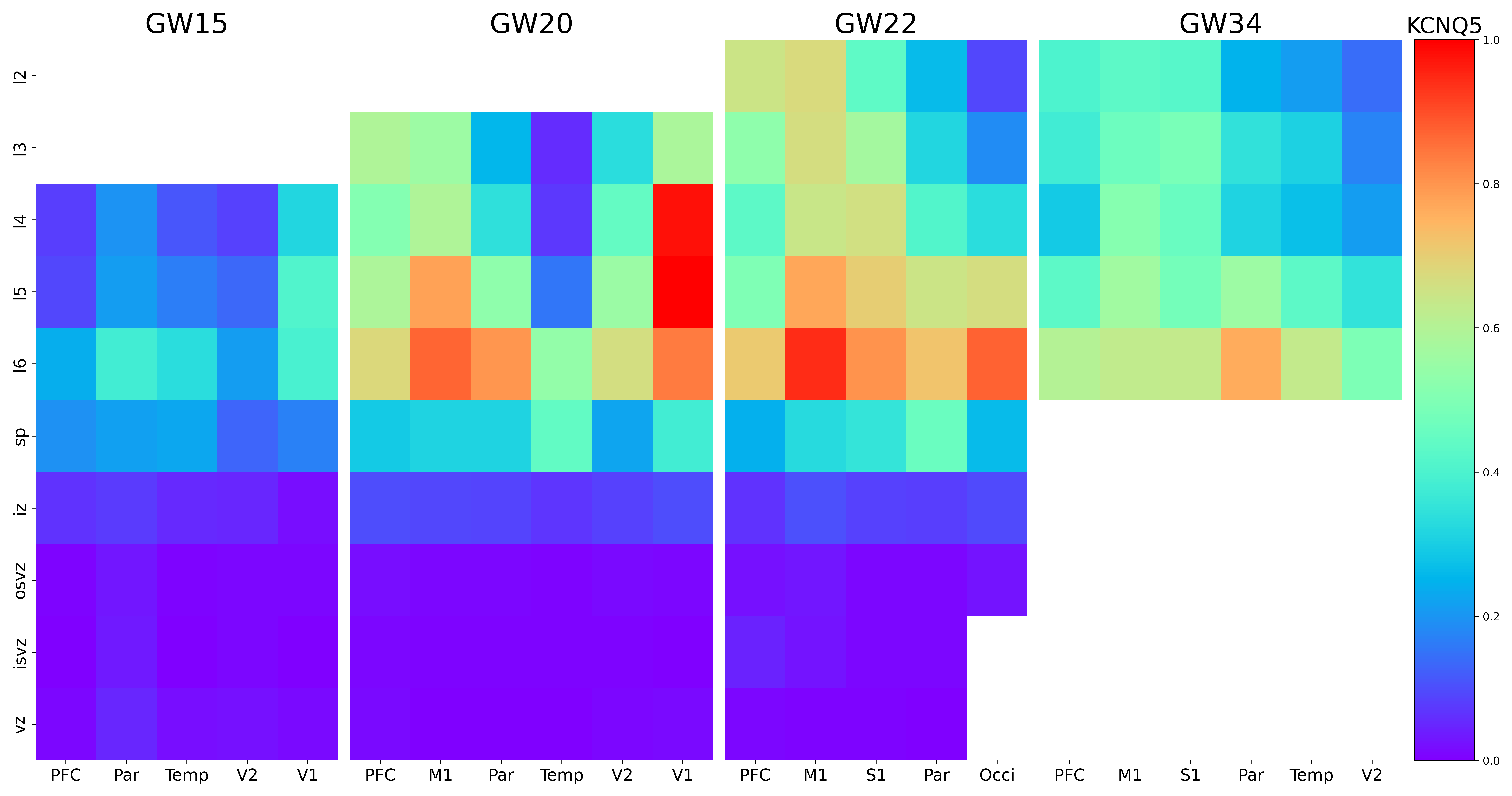

### KCTD12.png

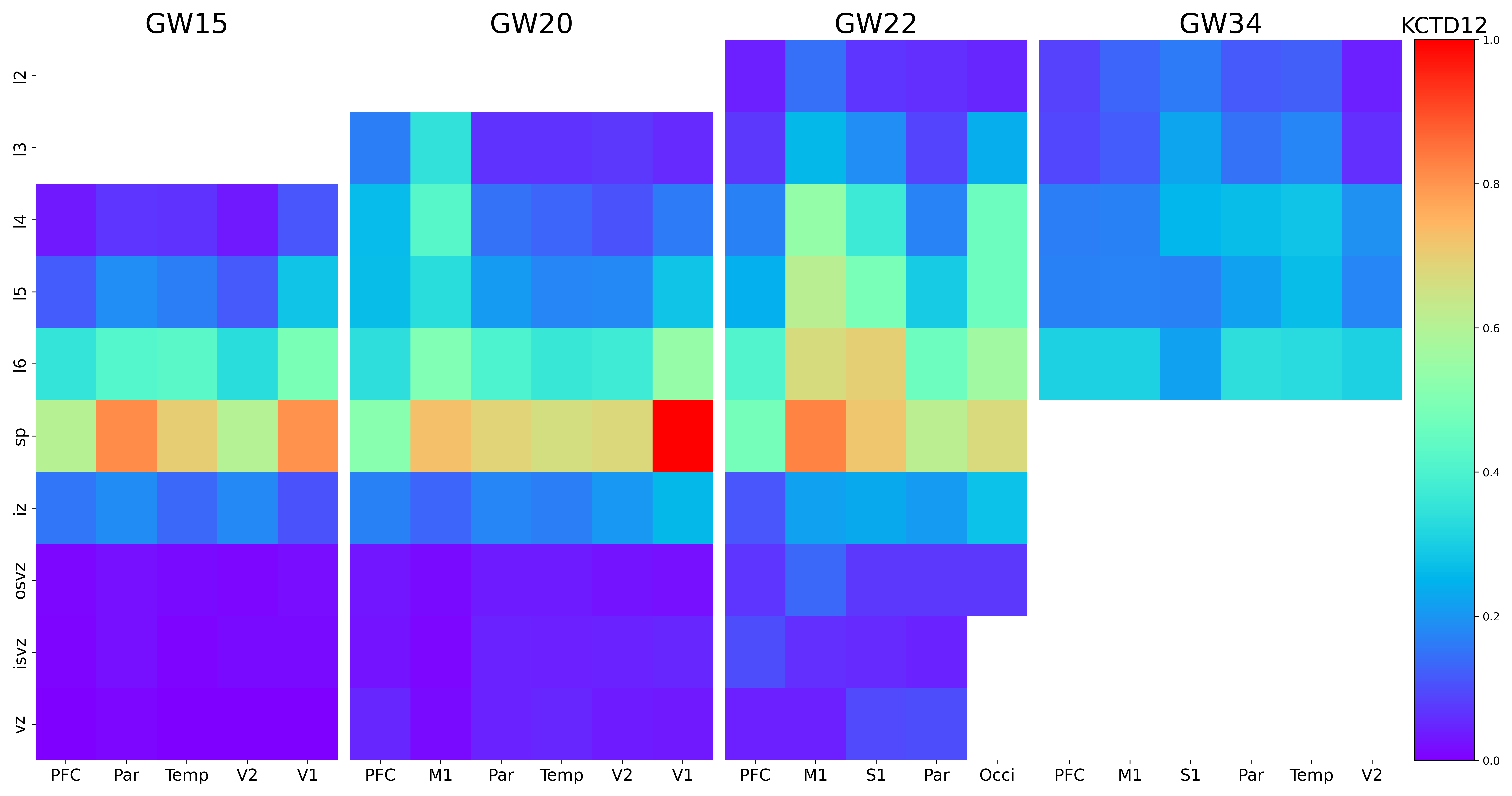

### KDM5B.png

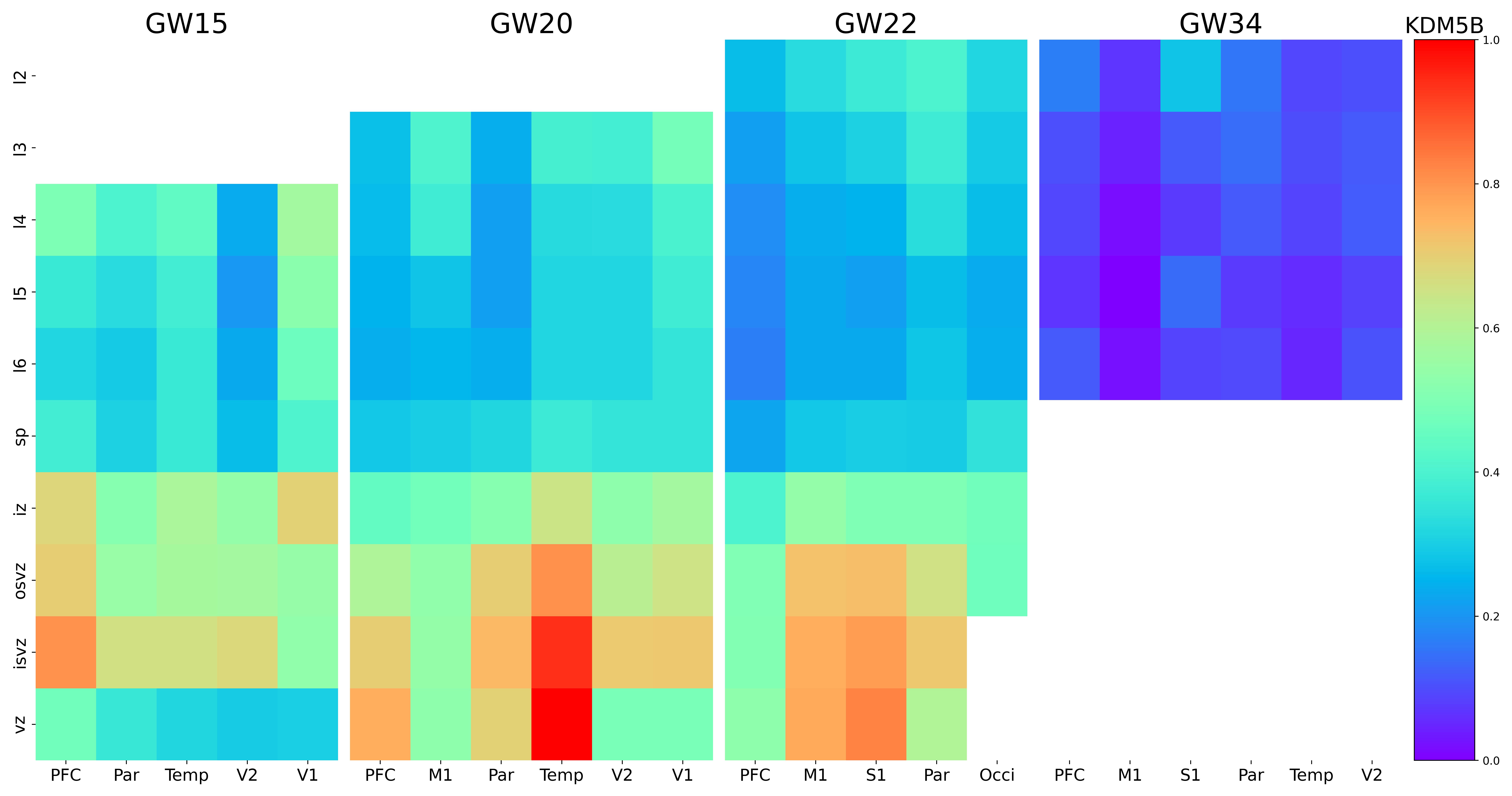

### KIAA0319.png

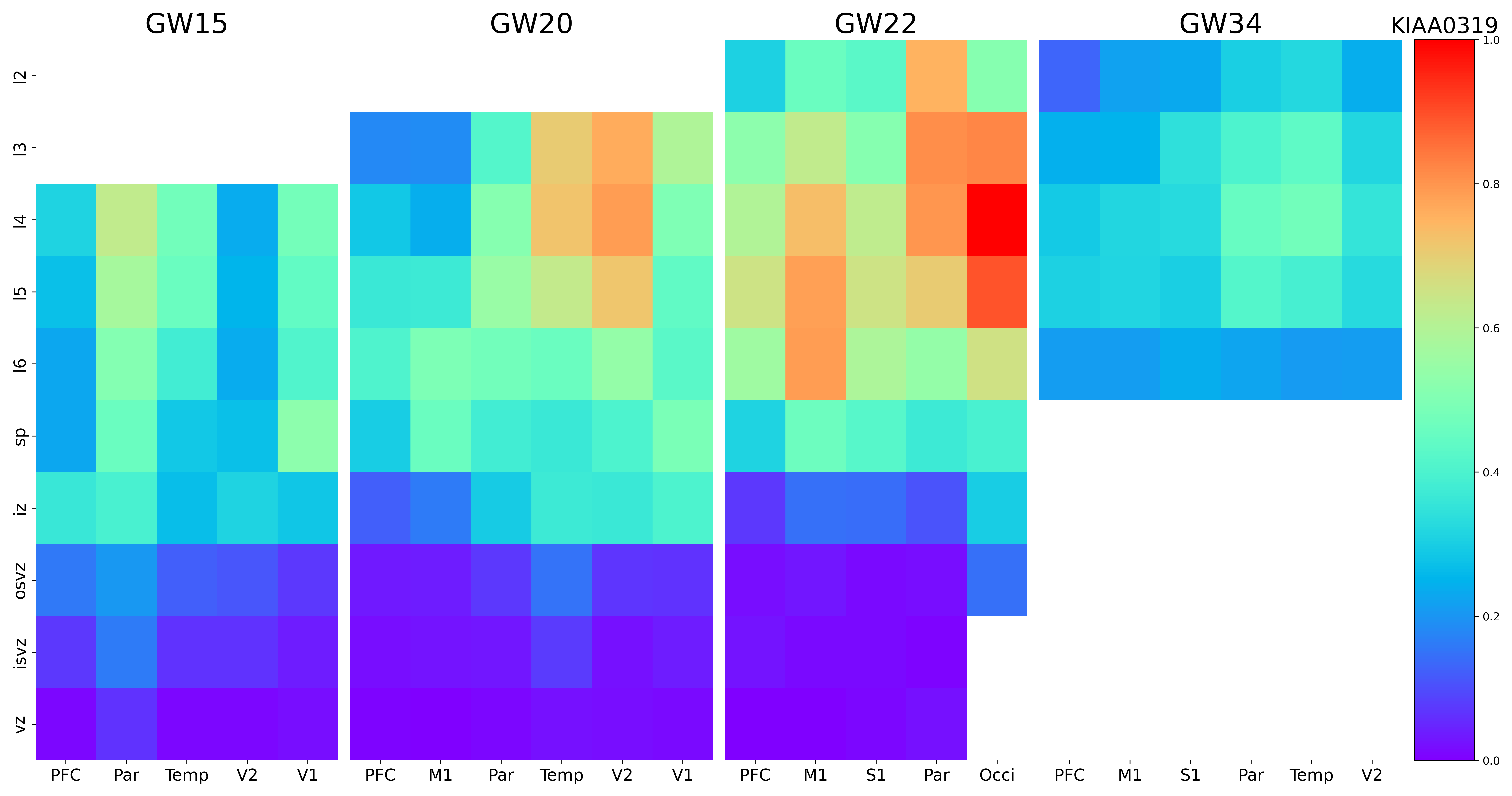

### KIF9-AS1.png

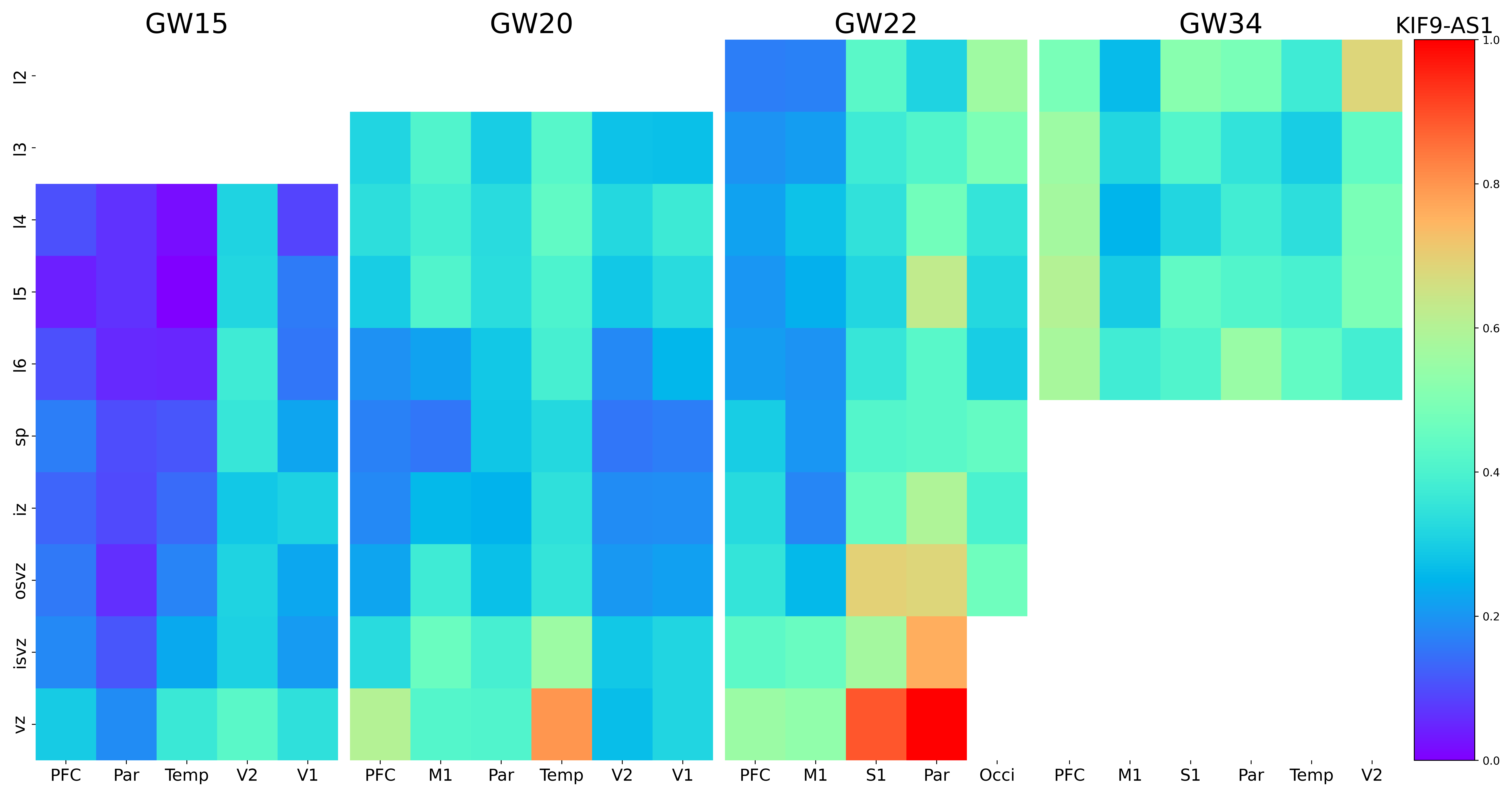

### KIF26B.png

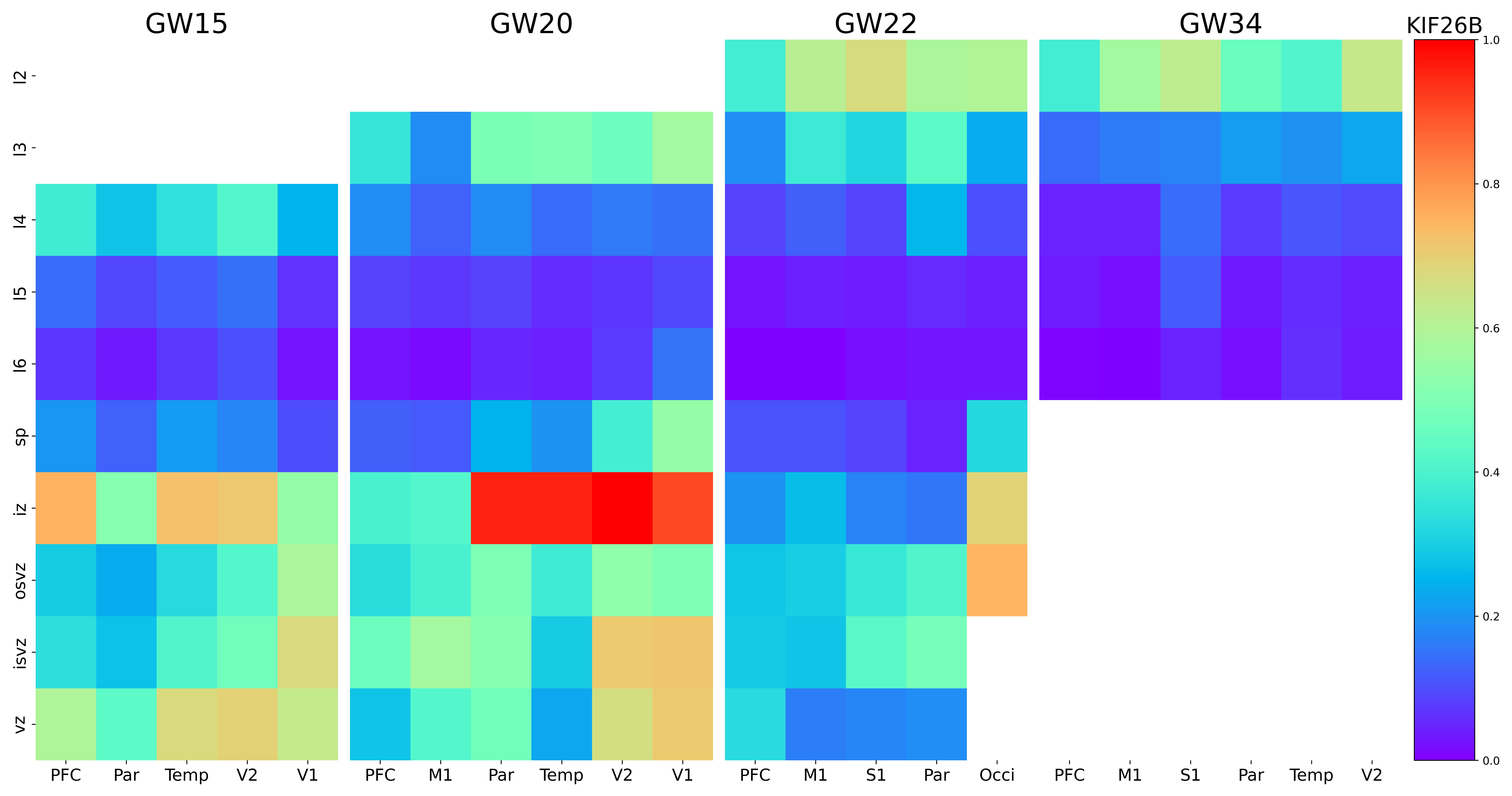

### KITLG.png

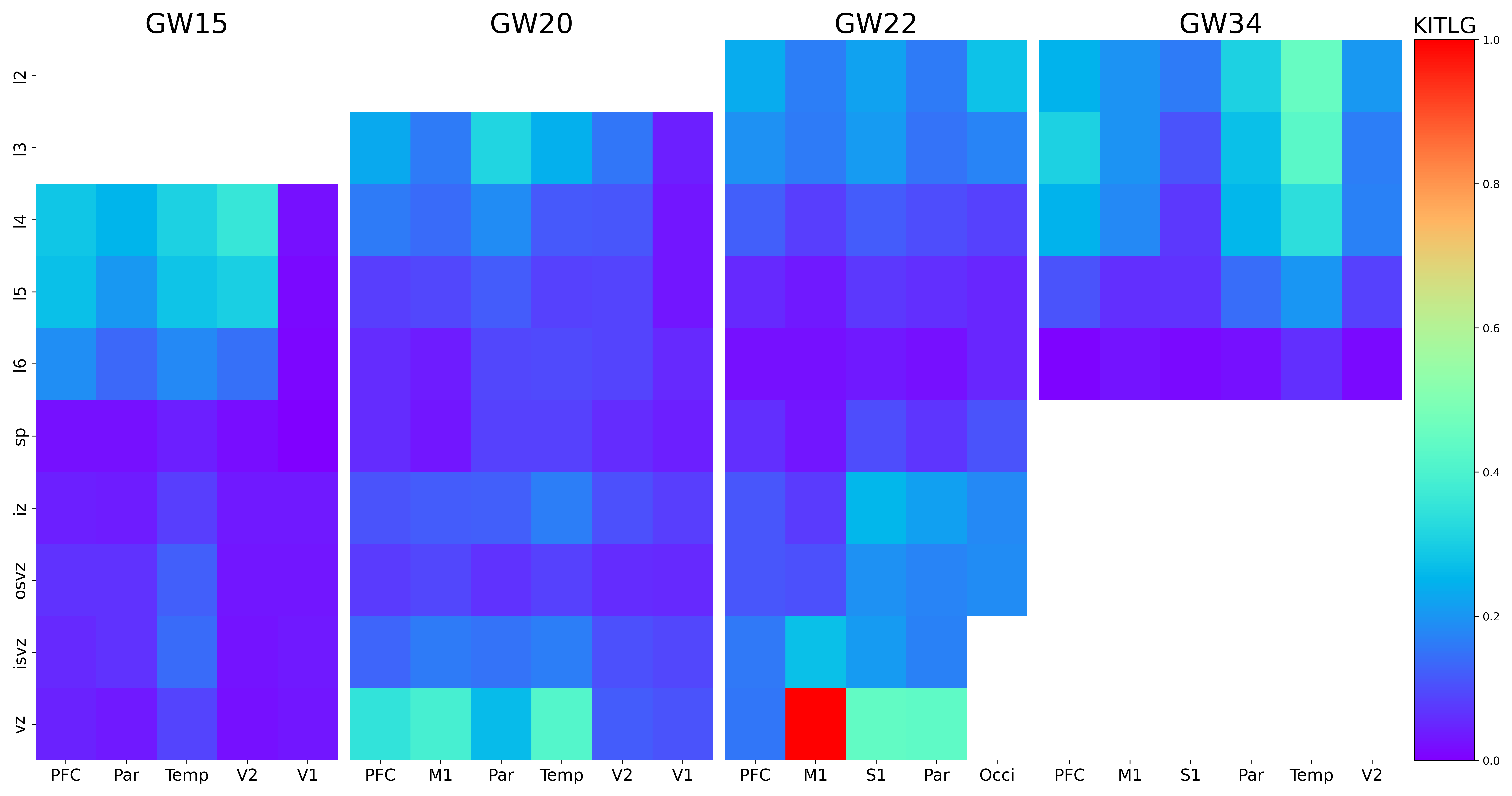

### KLHL1.png

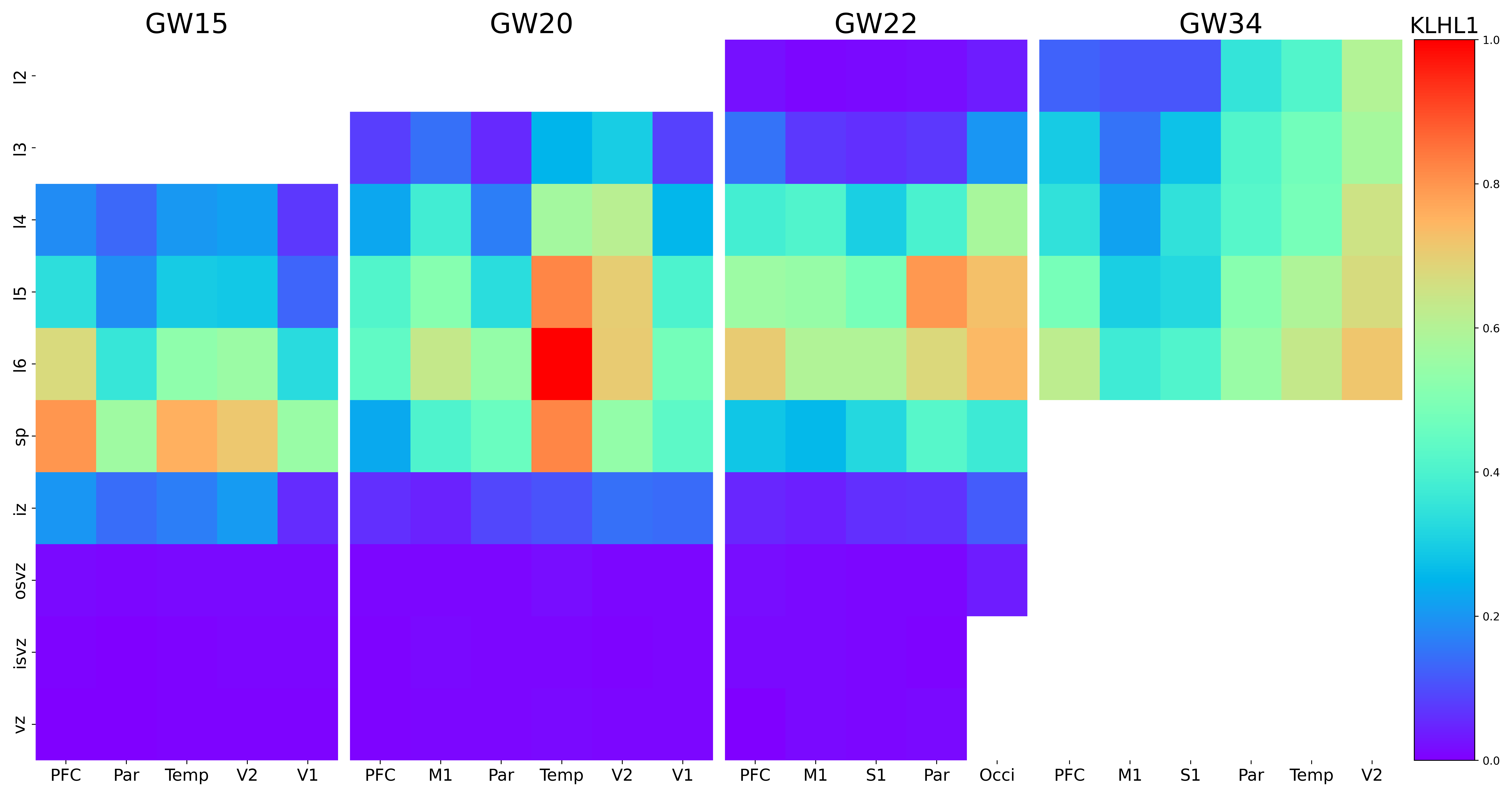

### KMT2E-AS1.png

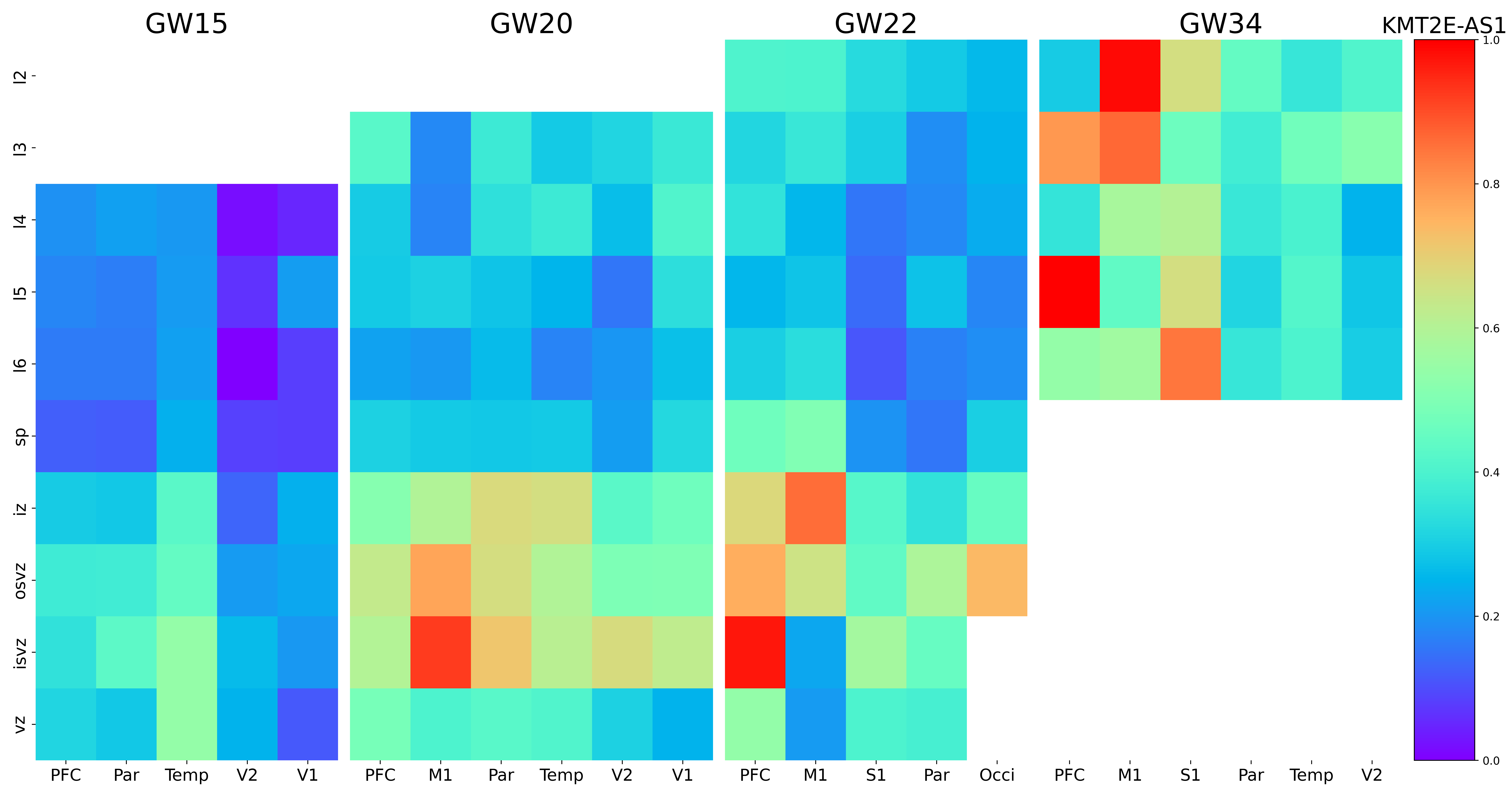

### KMT2E.png

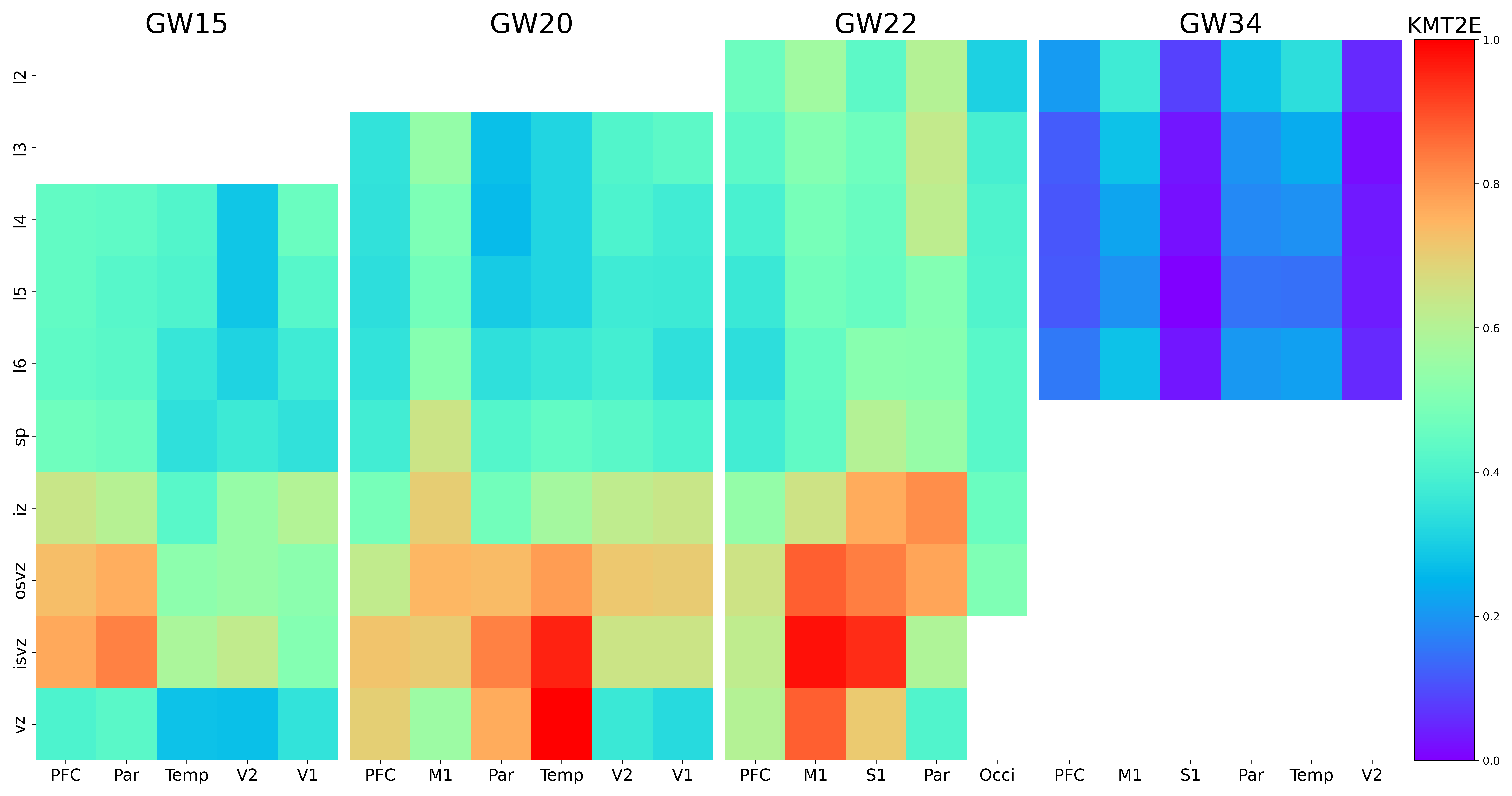

### KPTN.png

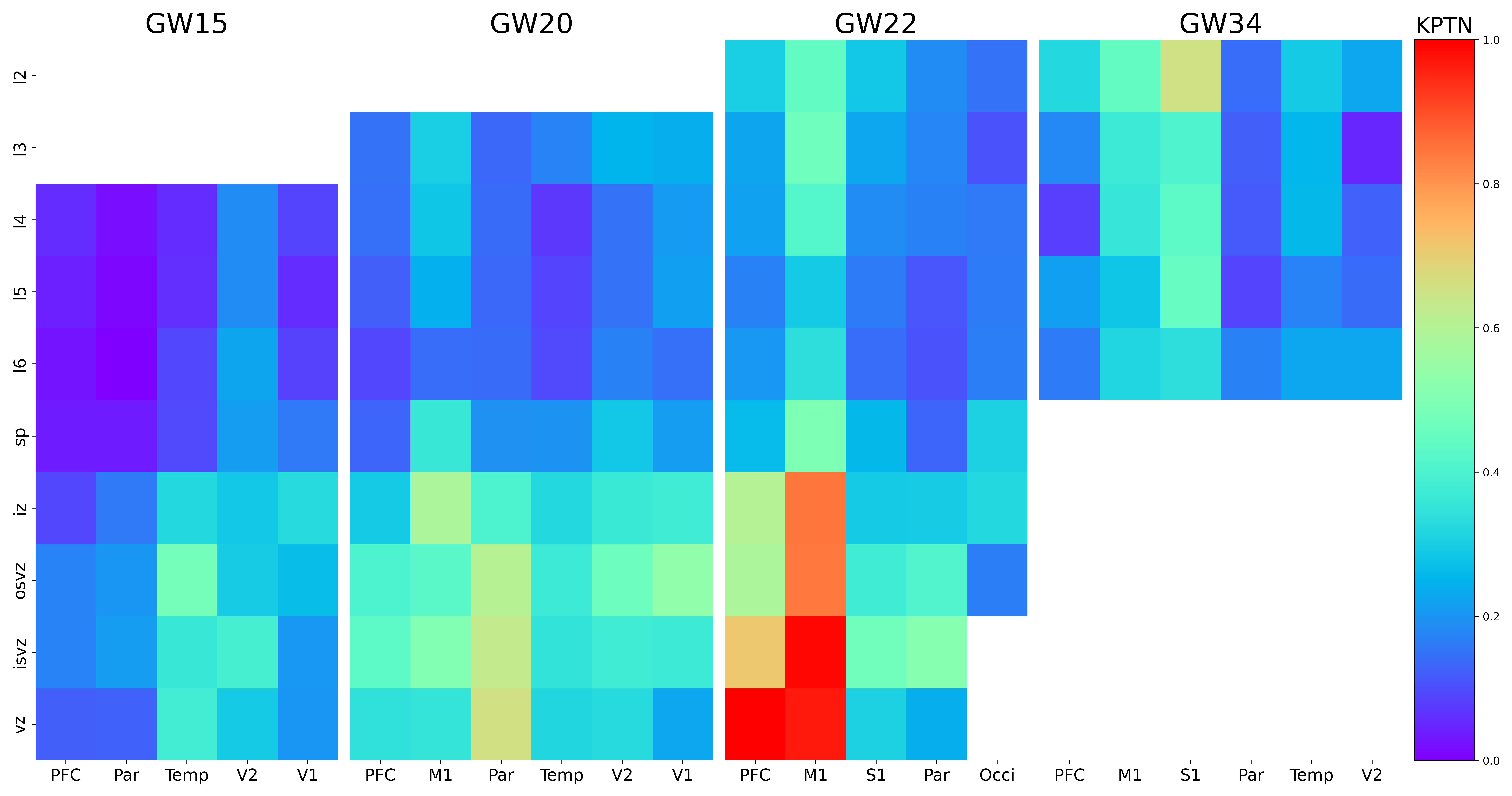

### LAMP5.png

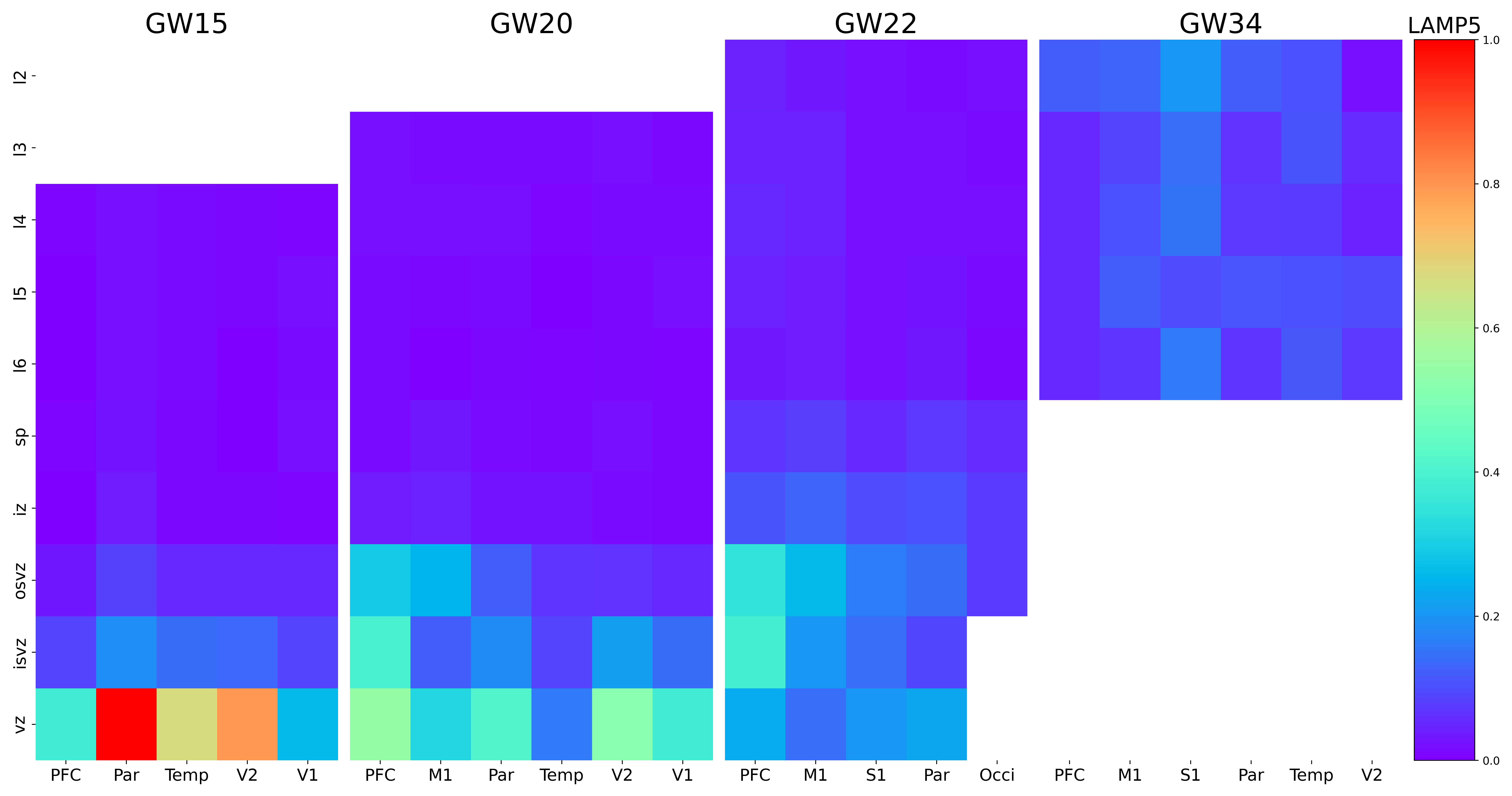

### LGALS1.png

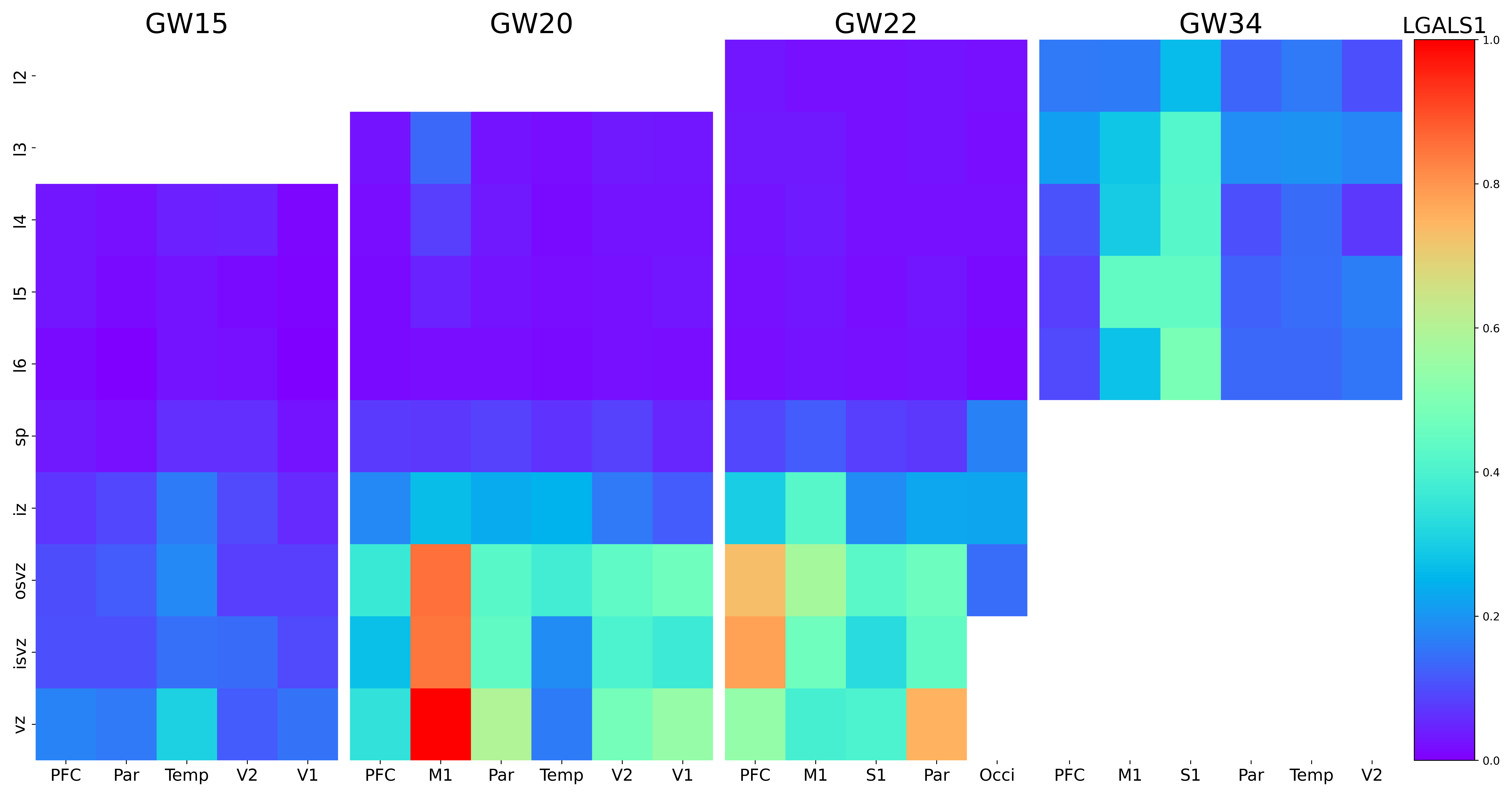

### LGR6.png

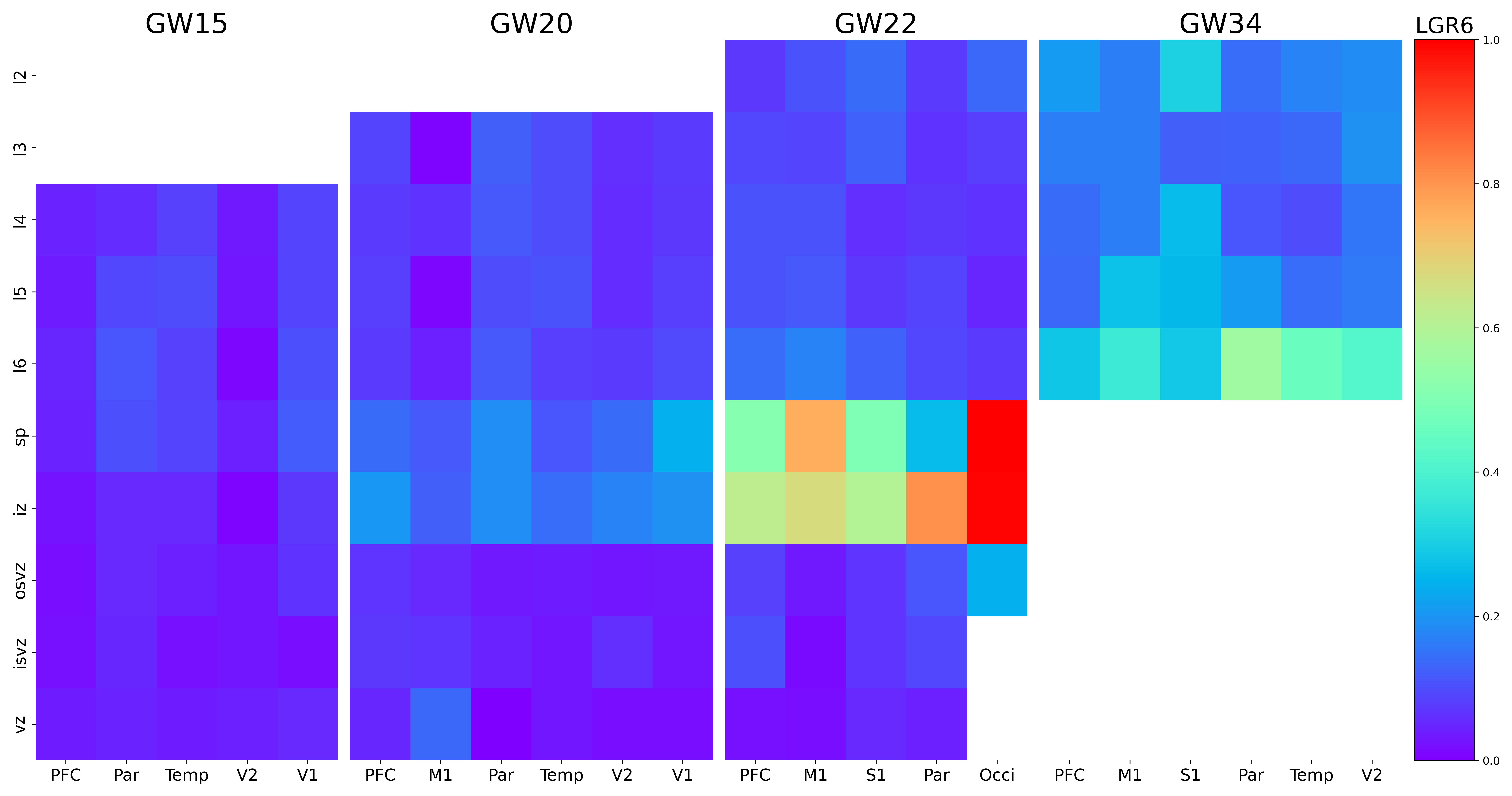

### LHX2.png

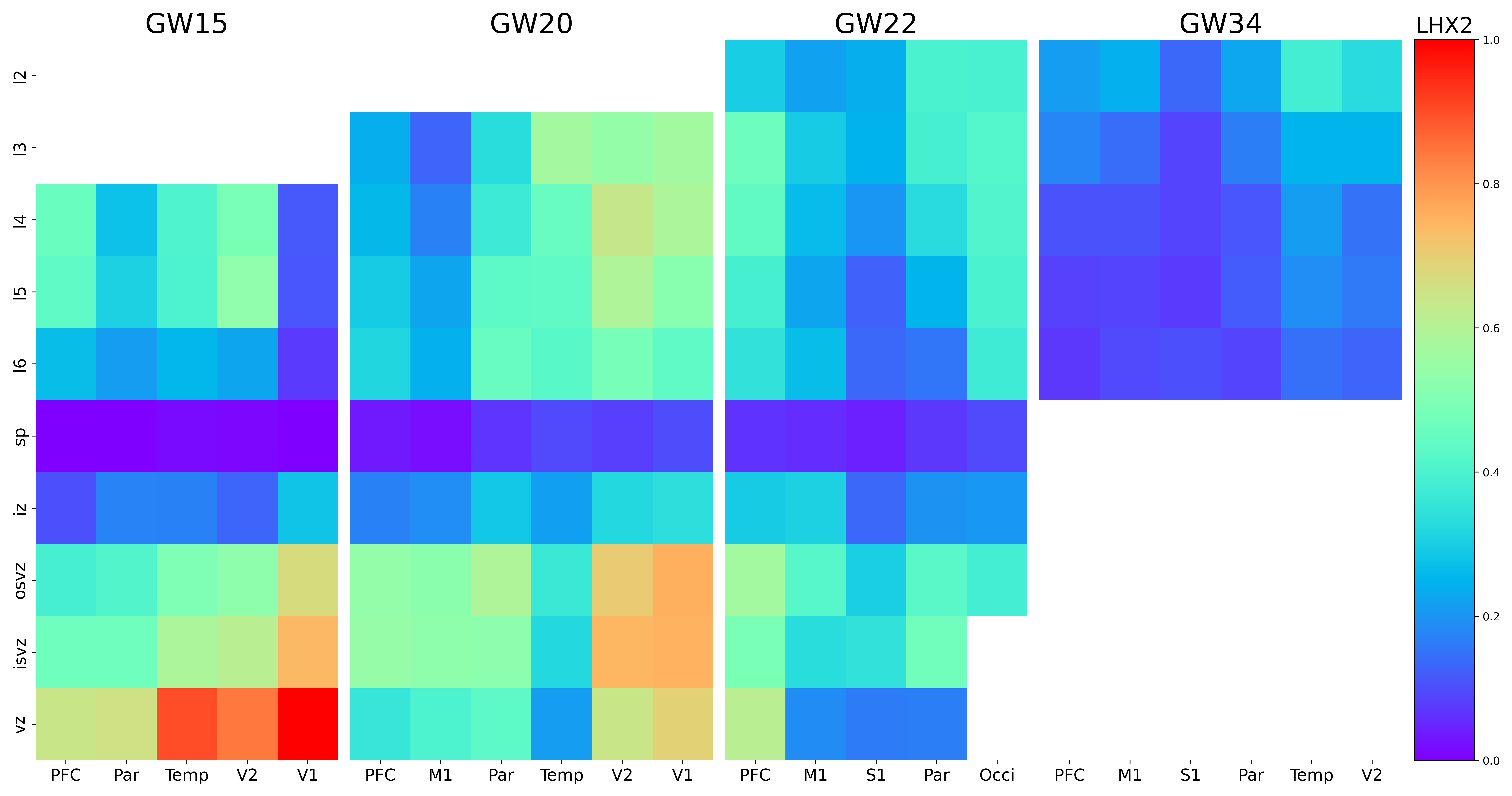

### LHX6.png

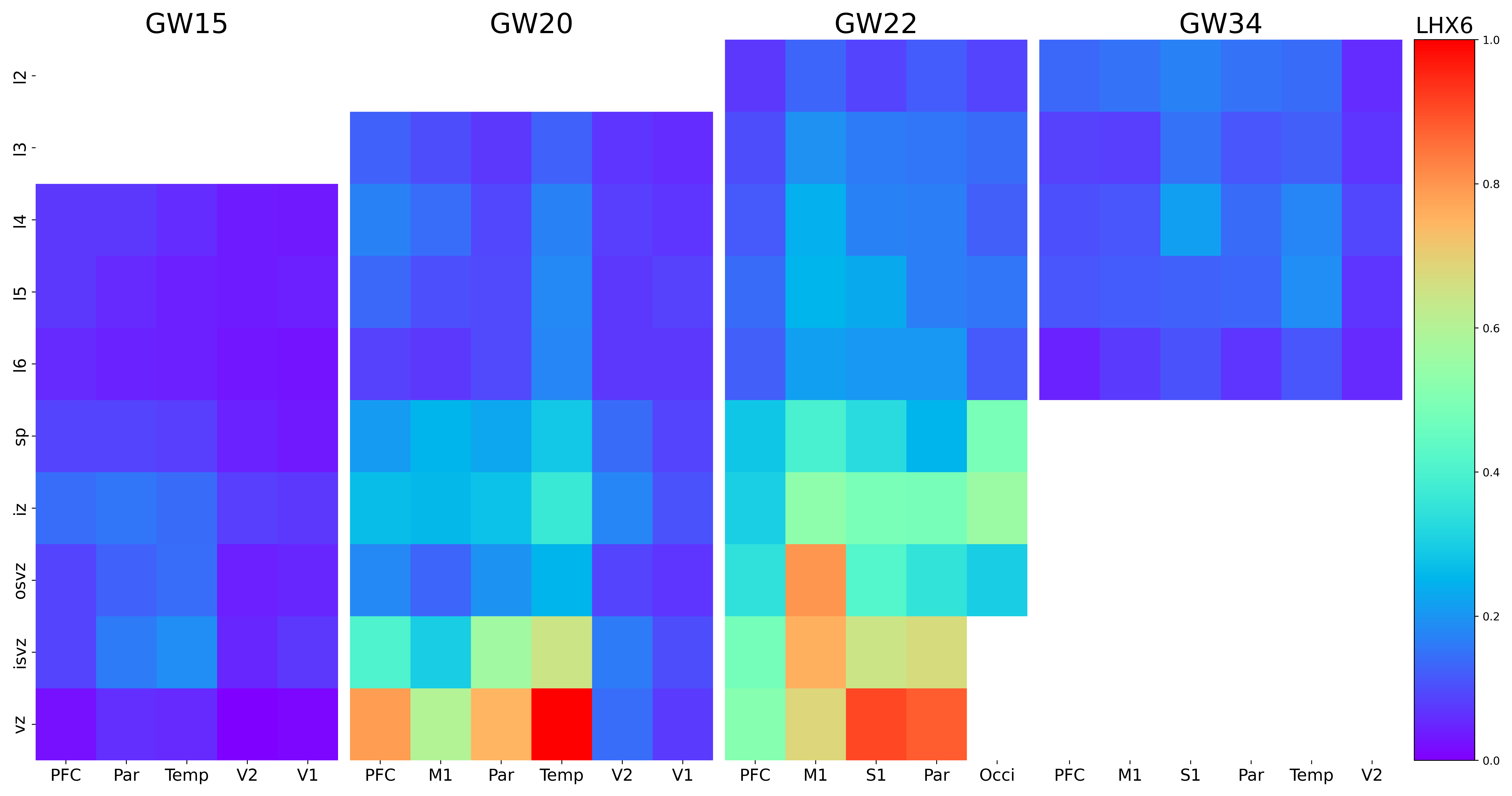

### LINC00271.png

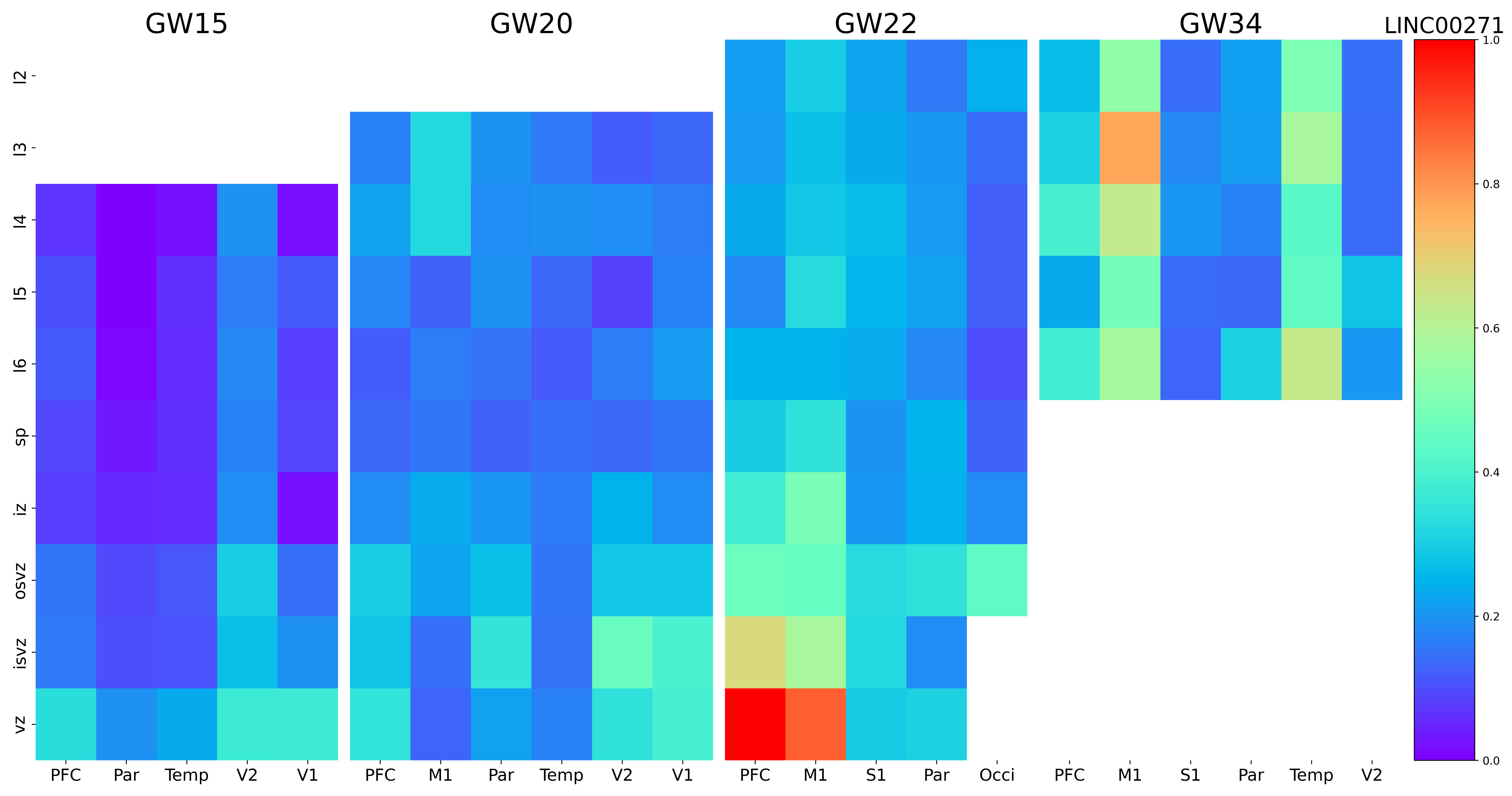

### LINC01089.png

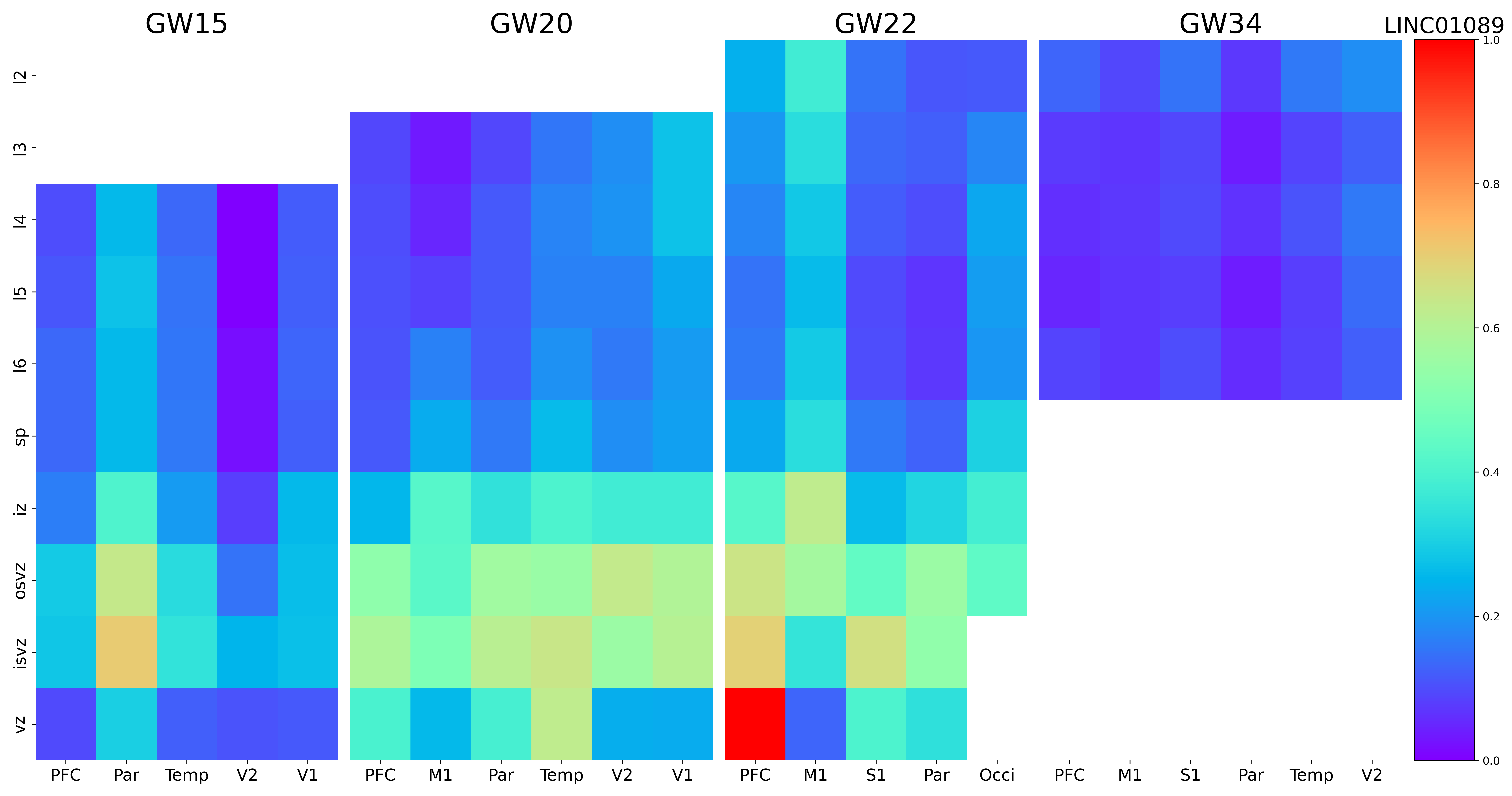

### LINC02593.png

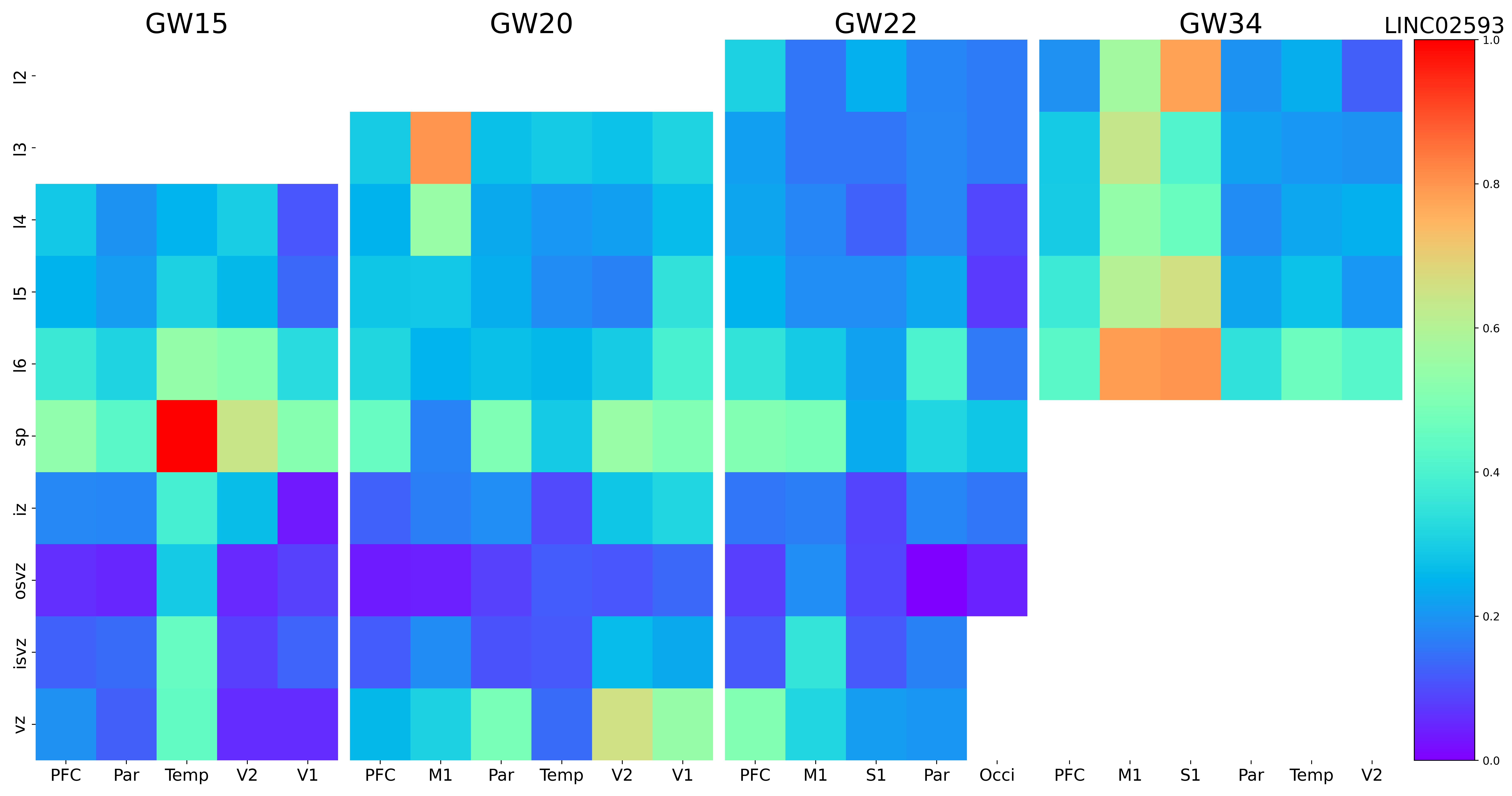

### LINGO1.png

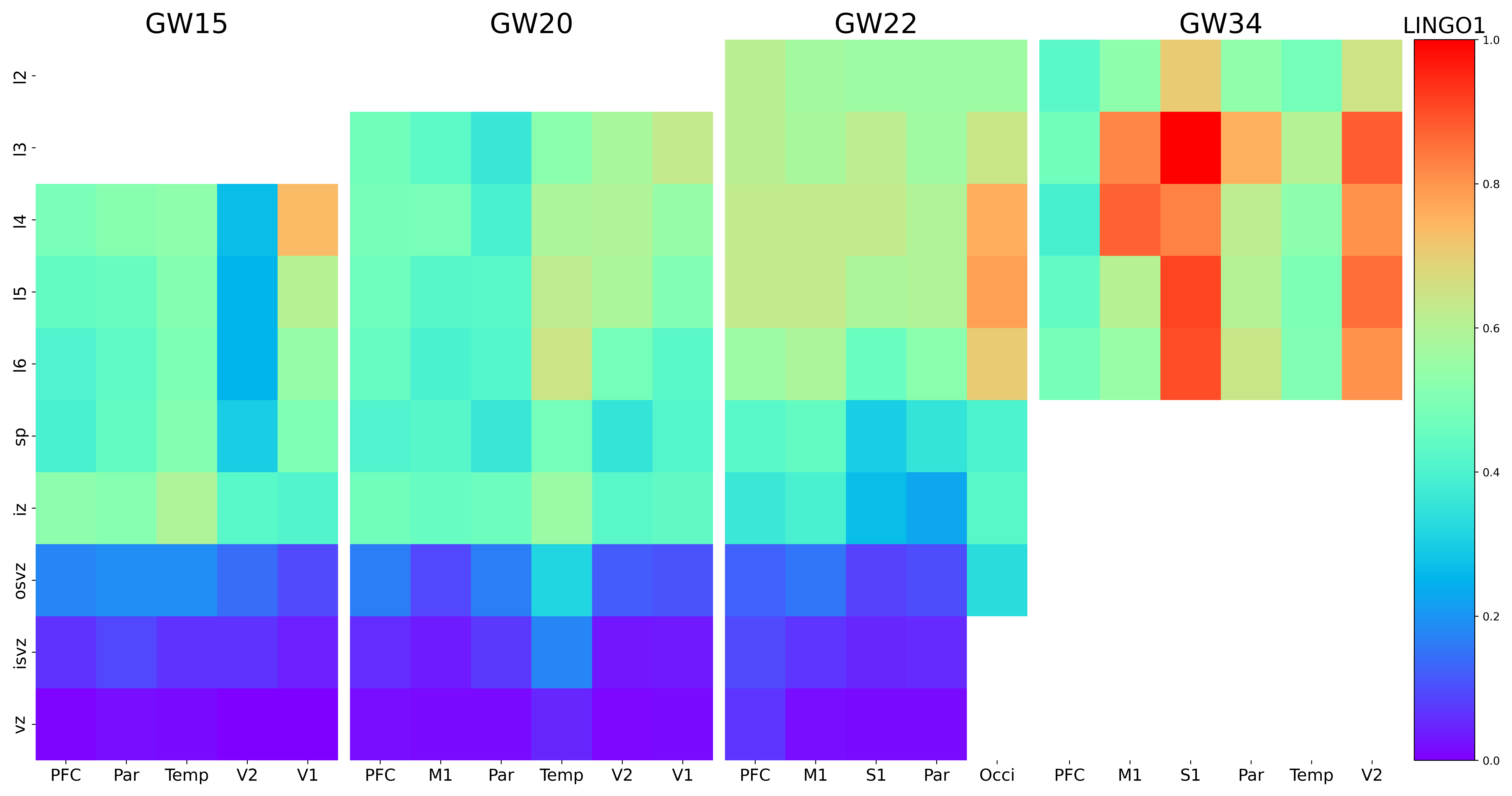

### LPAR1.png

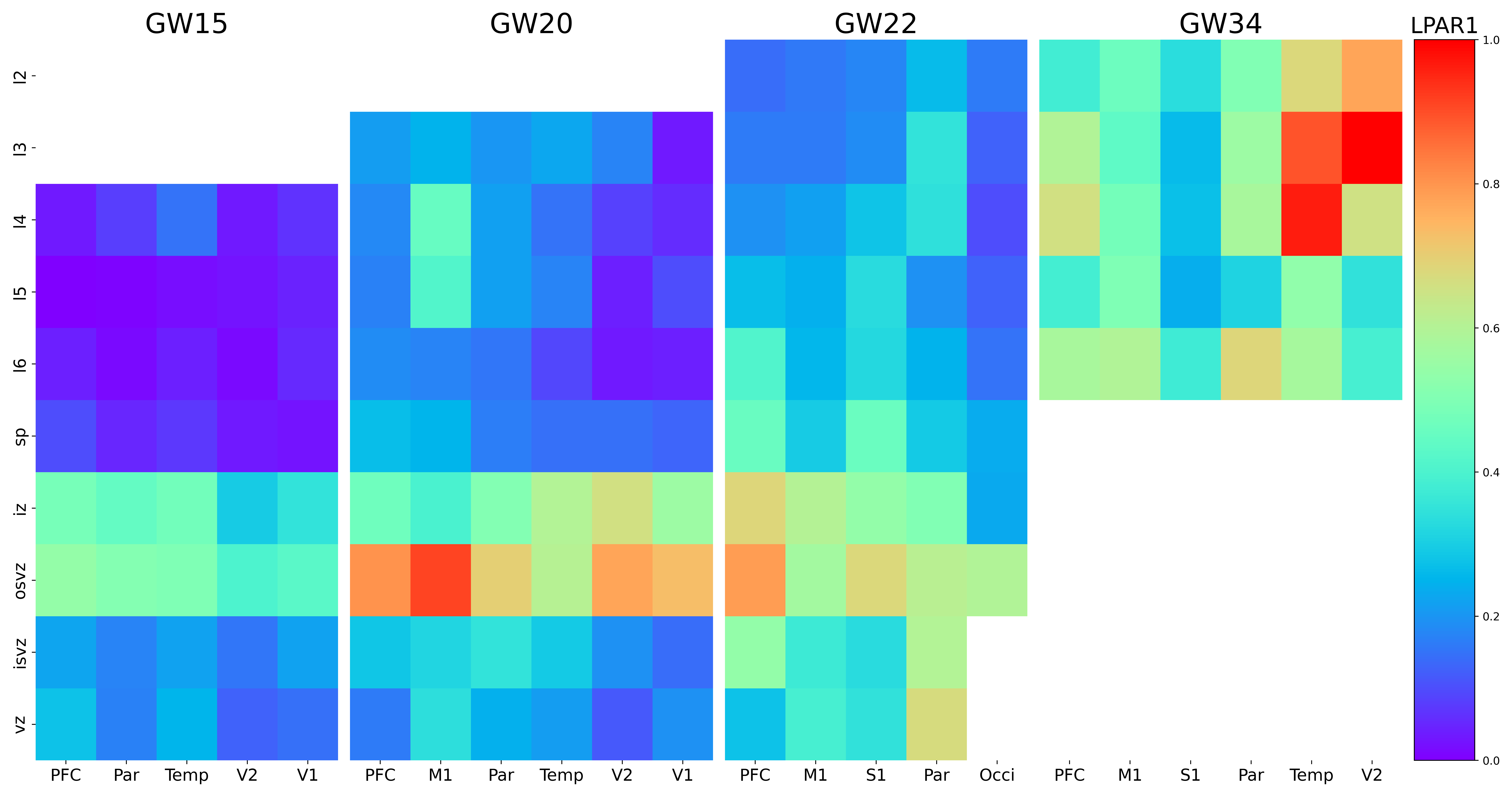

### LPL.png

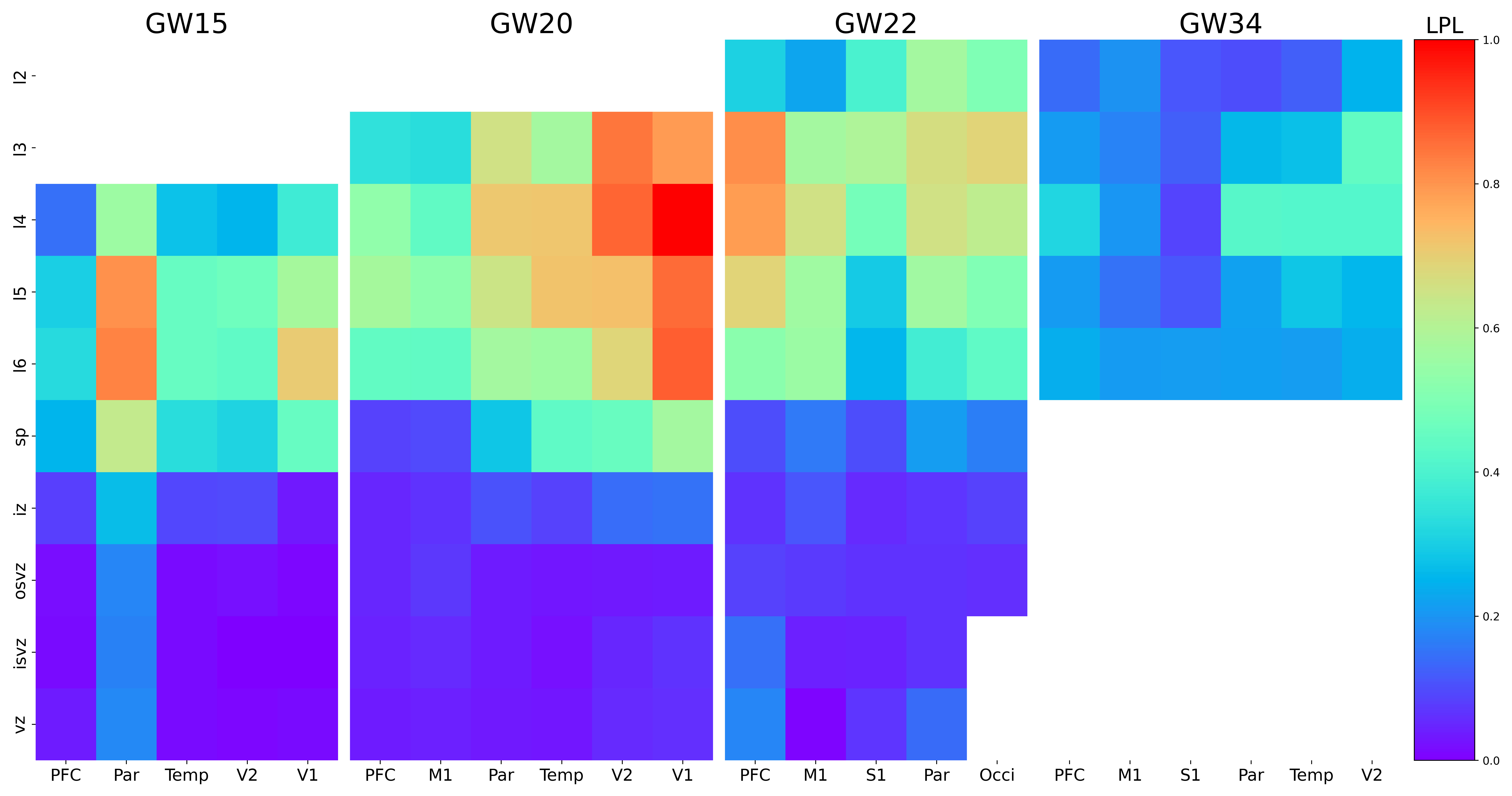

### LRATD2.png

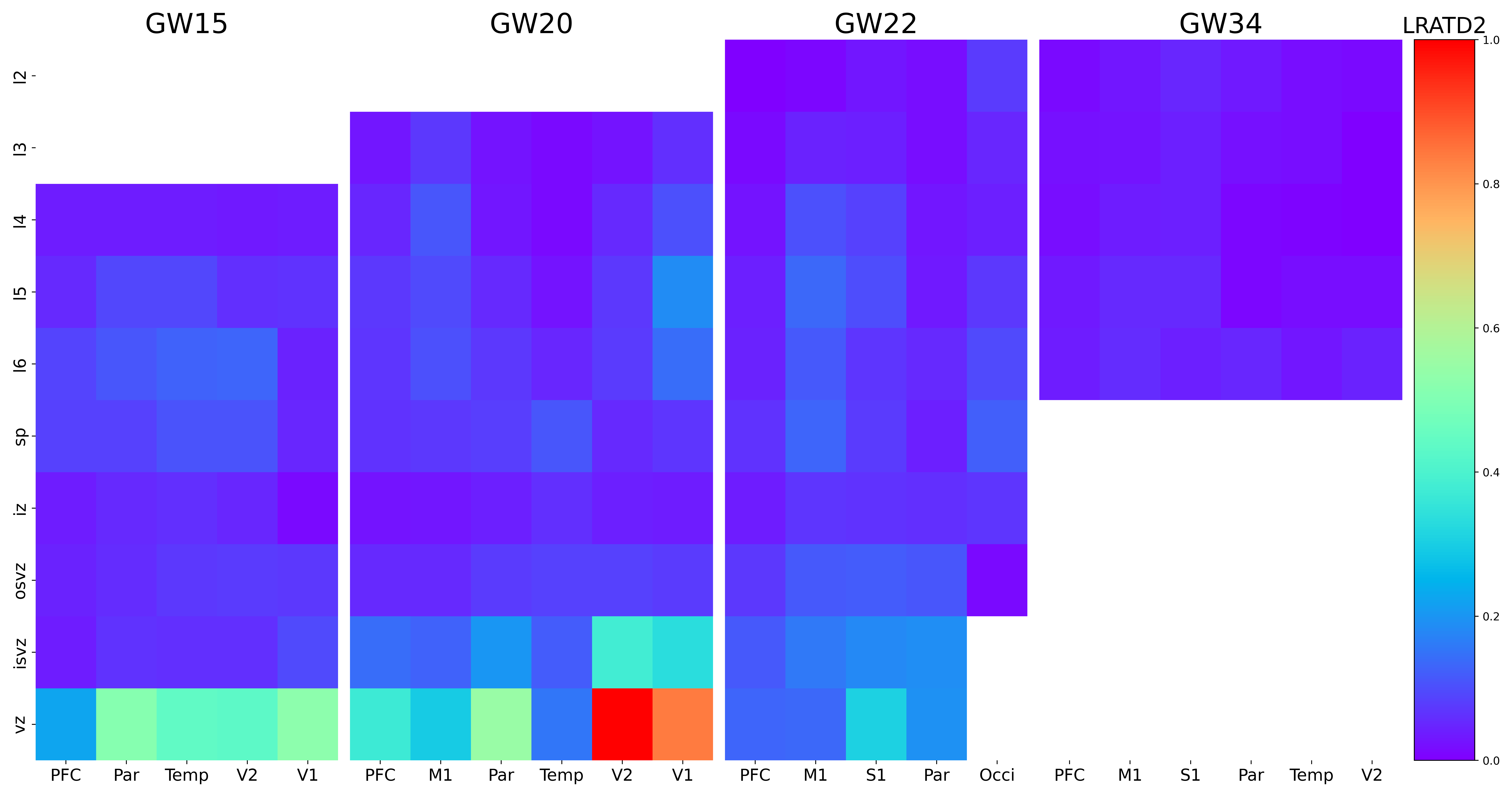

### LRP4.png

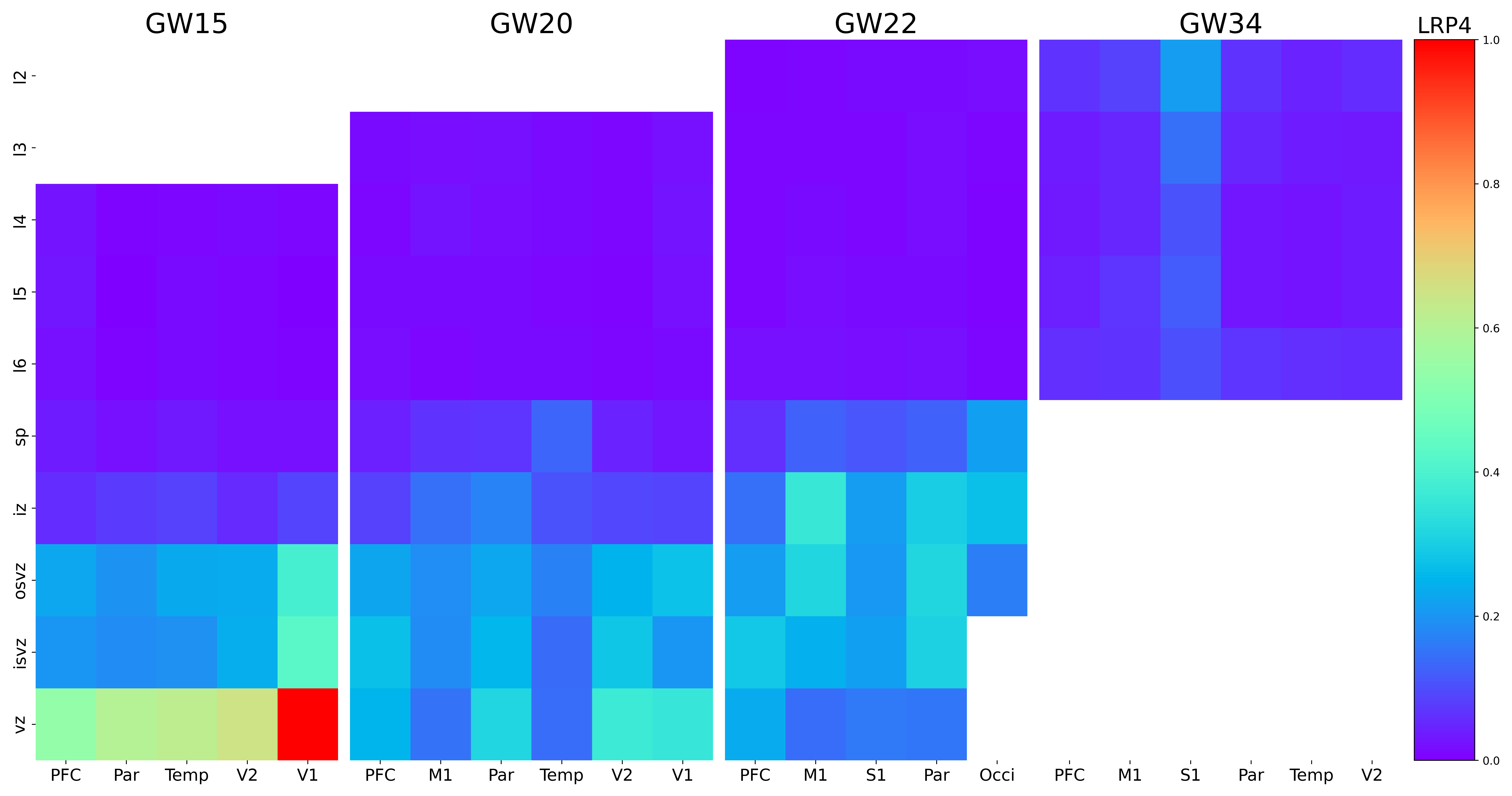

### MCPH1.png

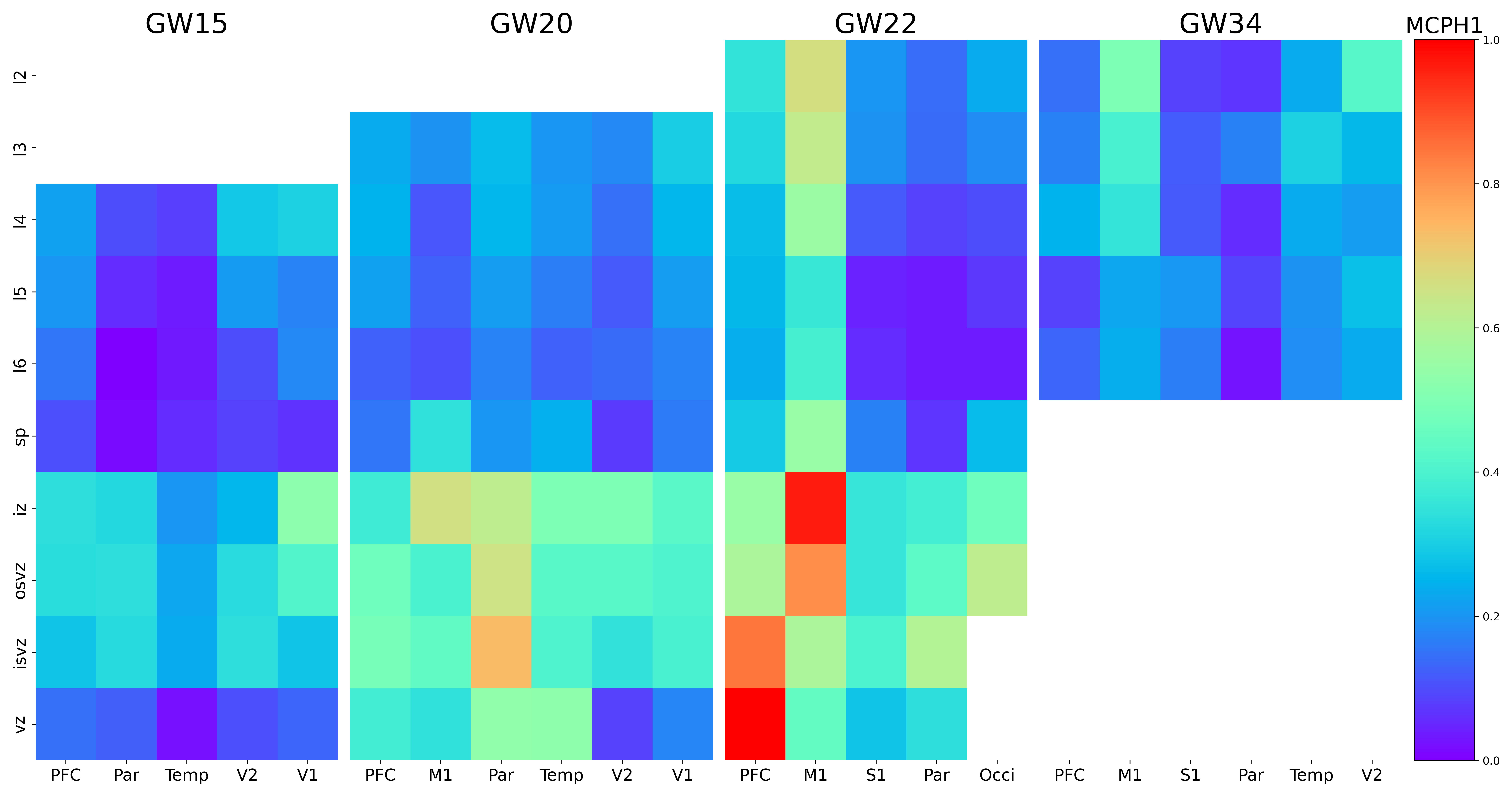

### MEIS1.png

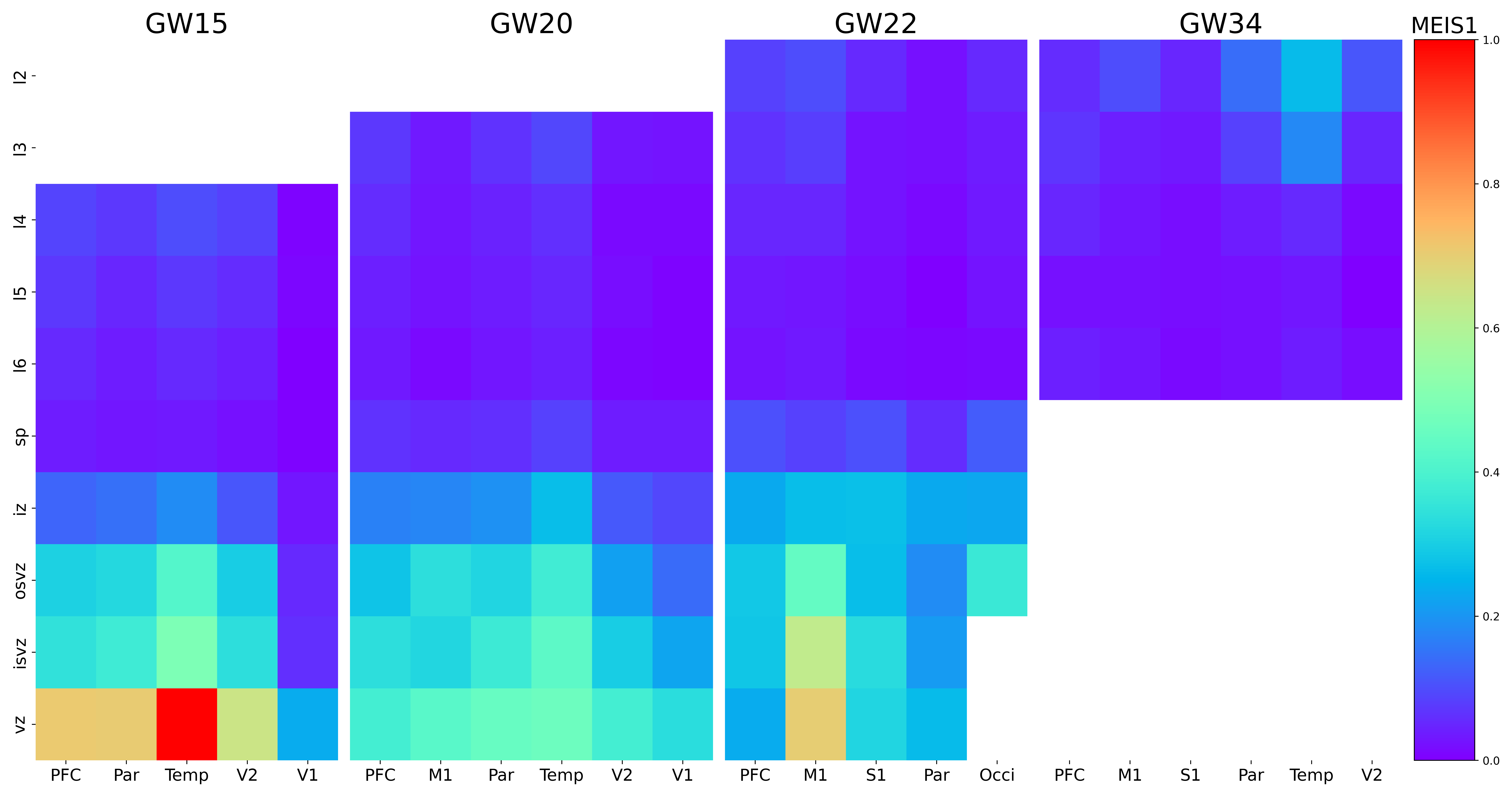

### MET.png

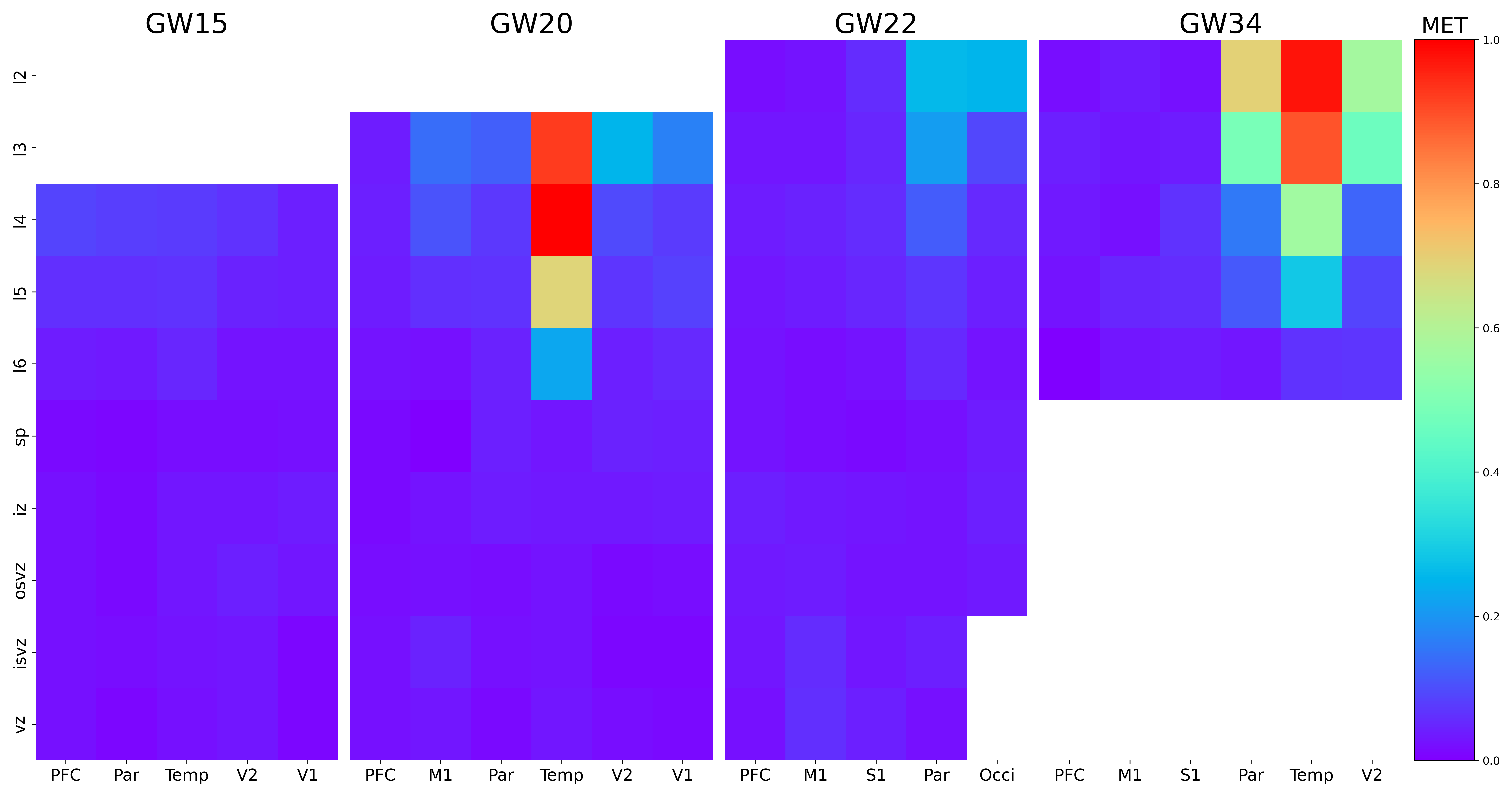

### MLIP.png

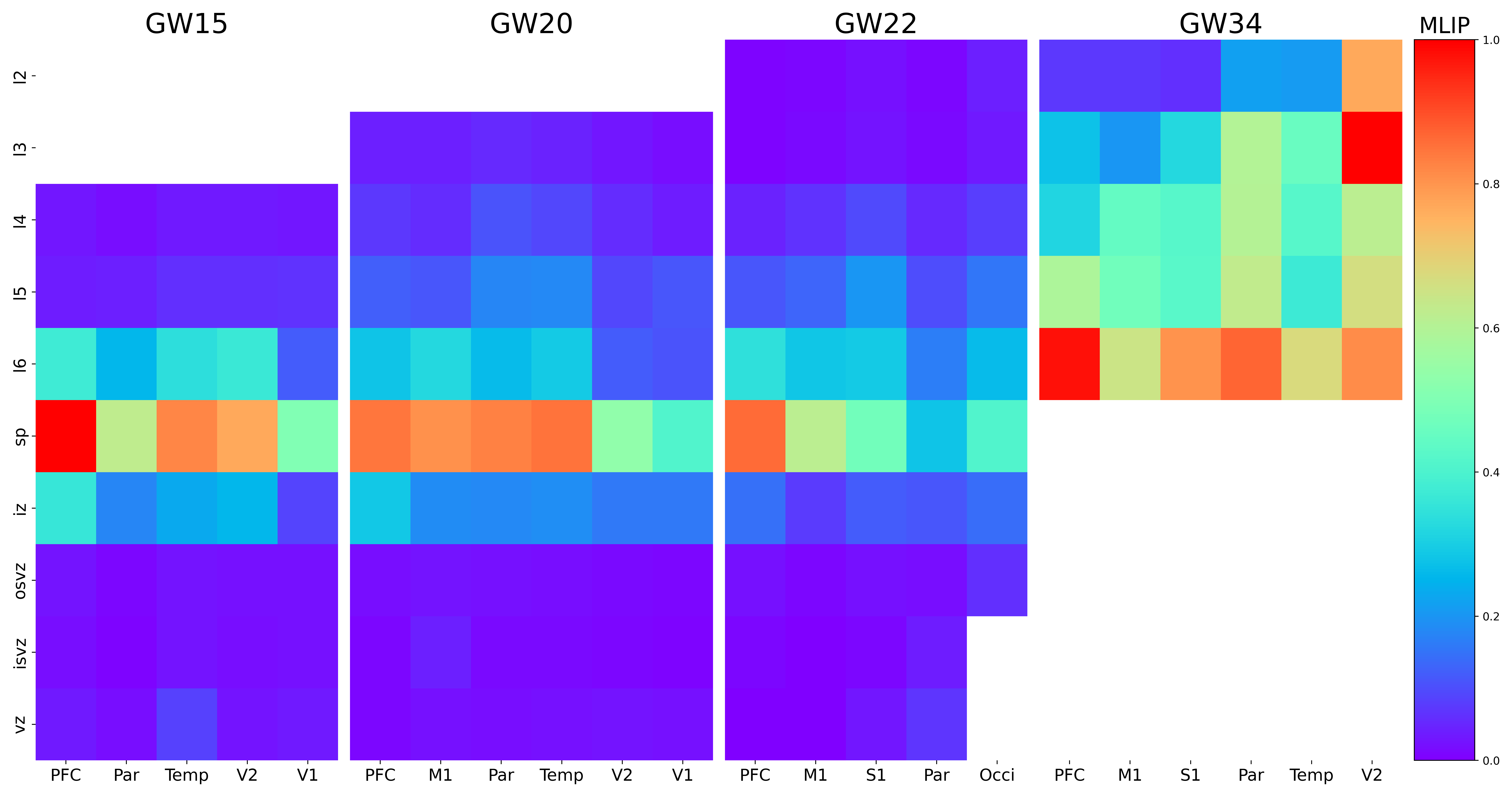

### MOXD1.png

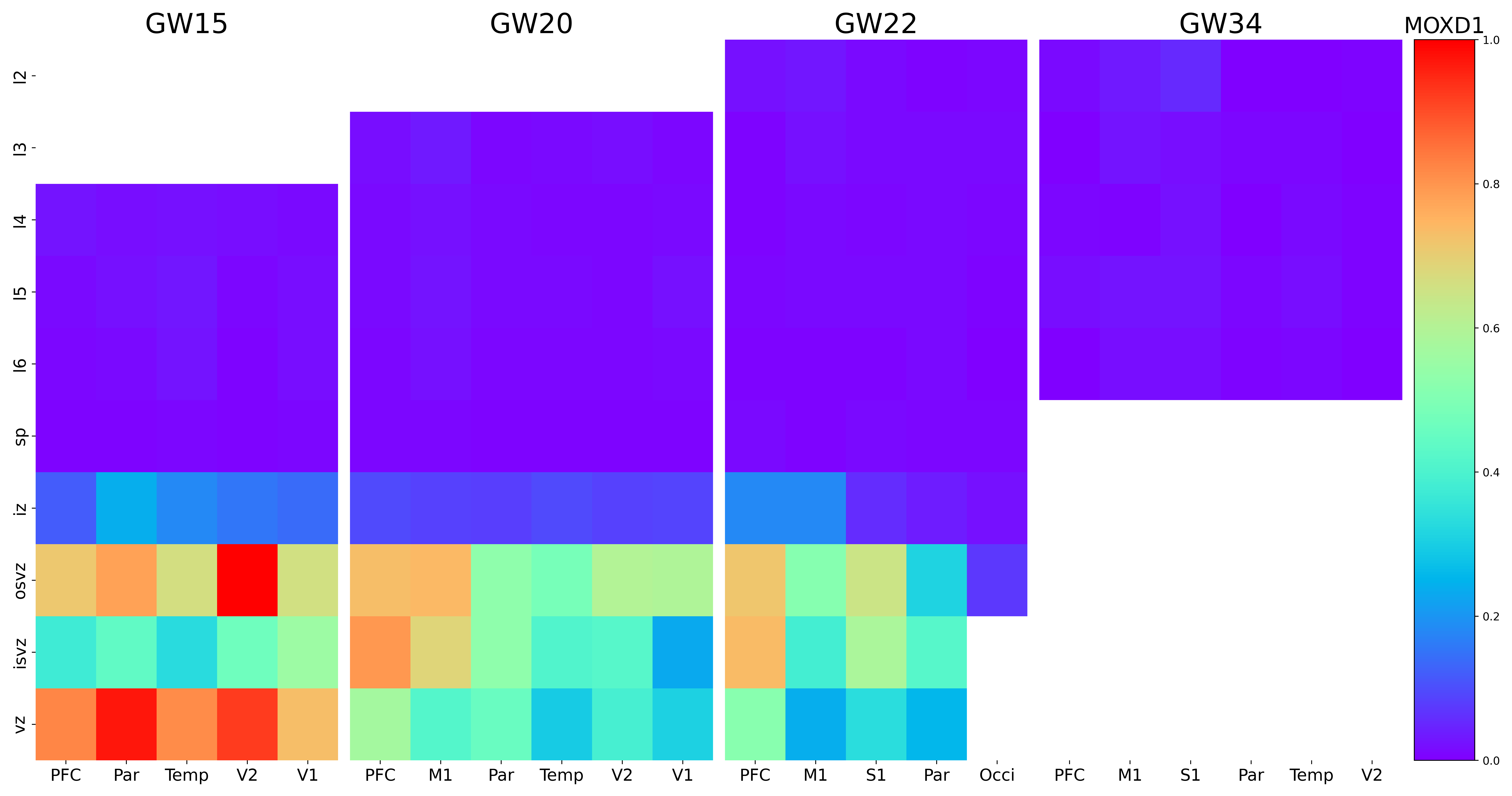

### MSANTD2.png

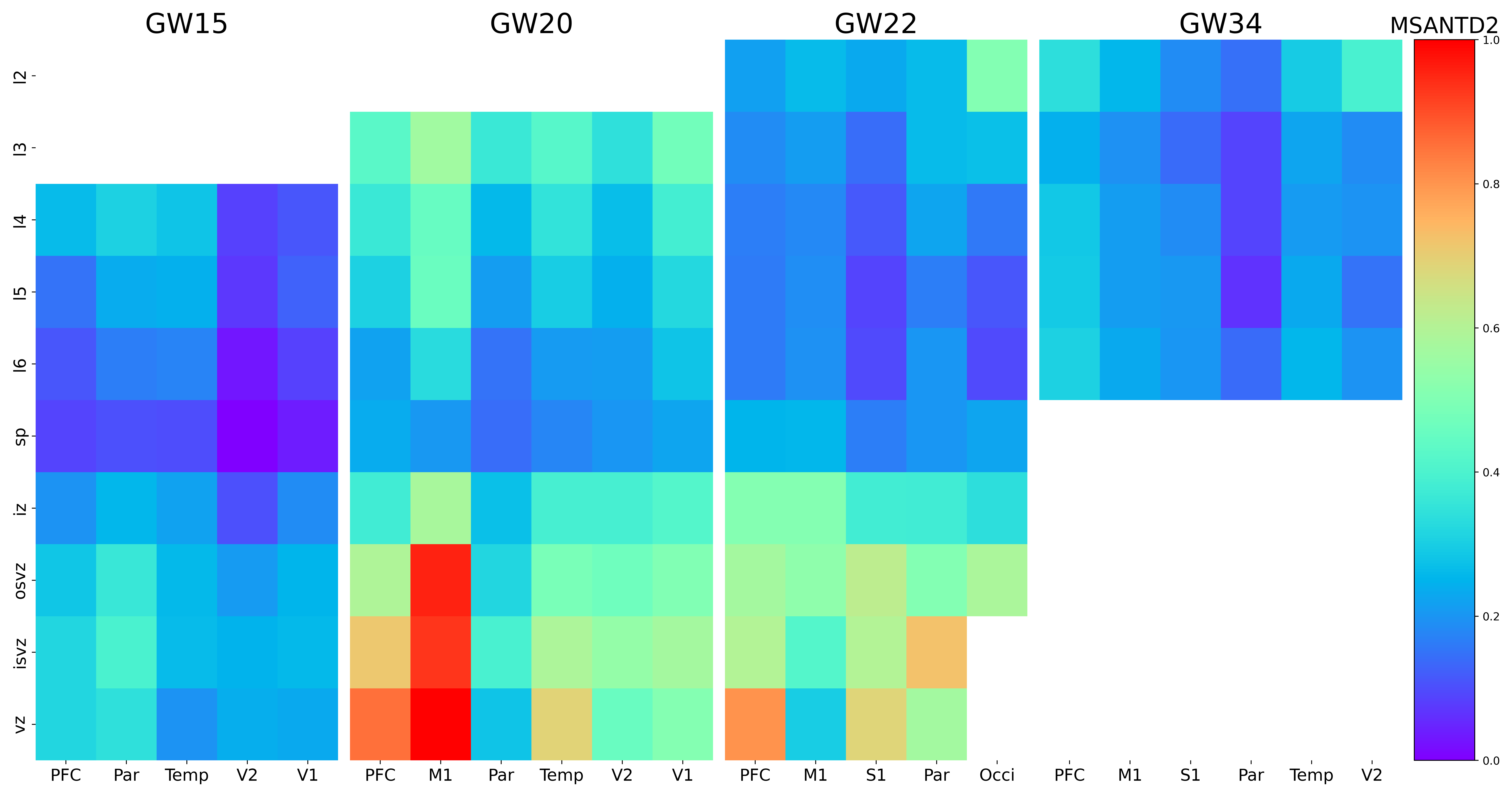

### MSL3.png

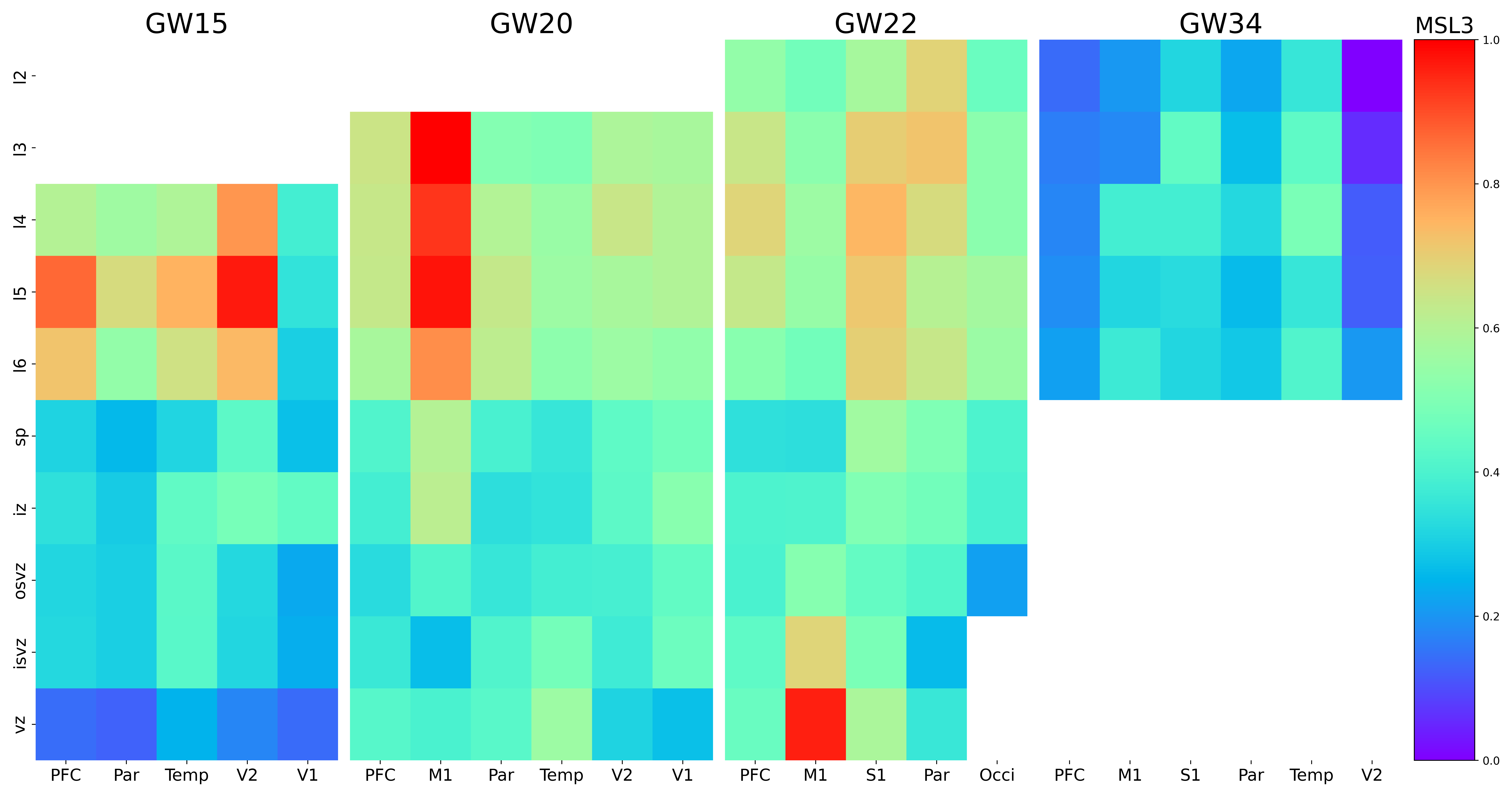

### MT3.png

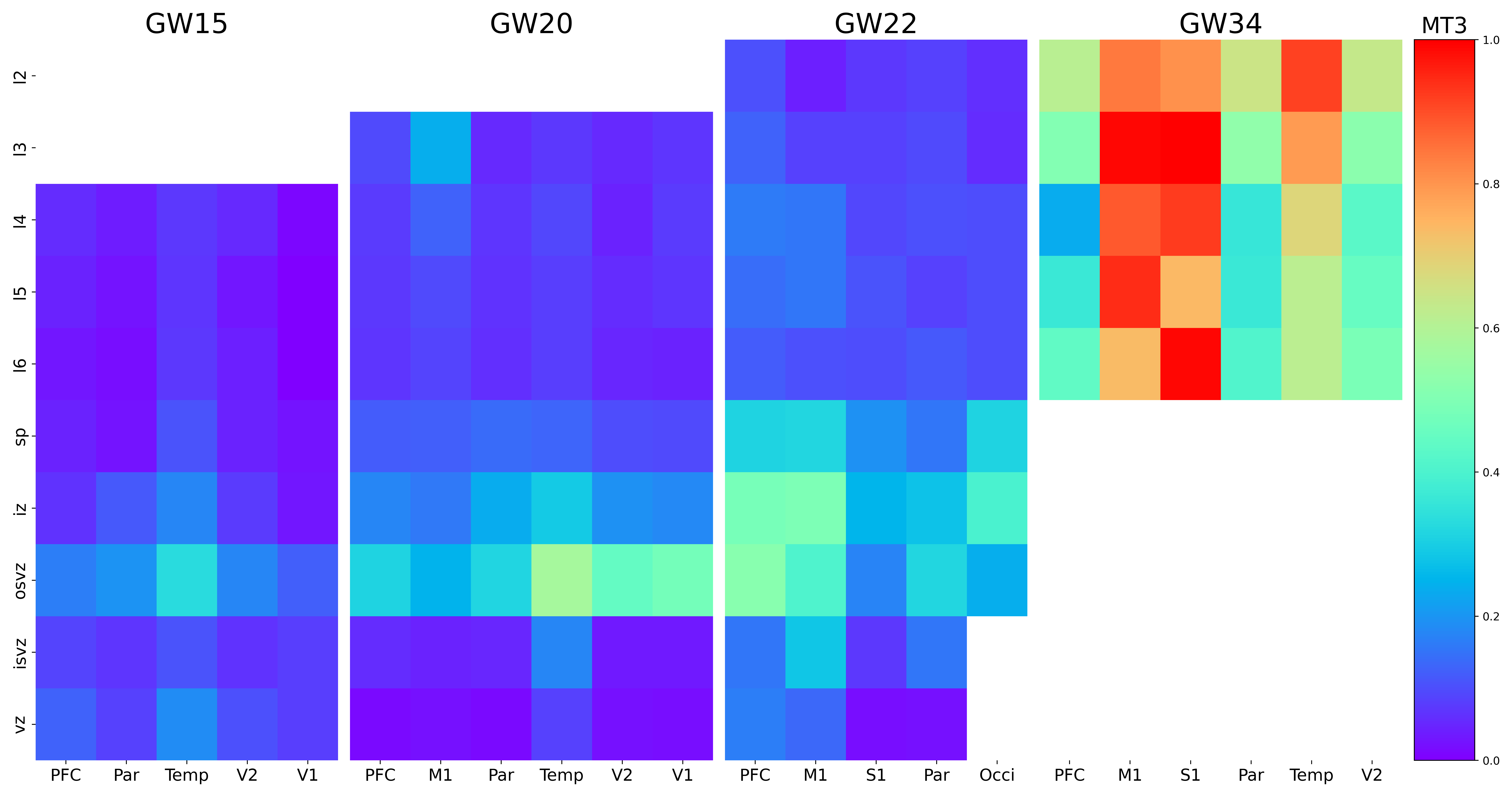

### MYT1L.png

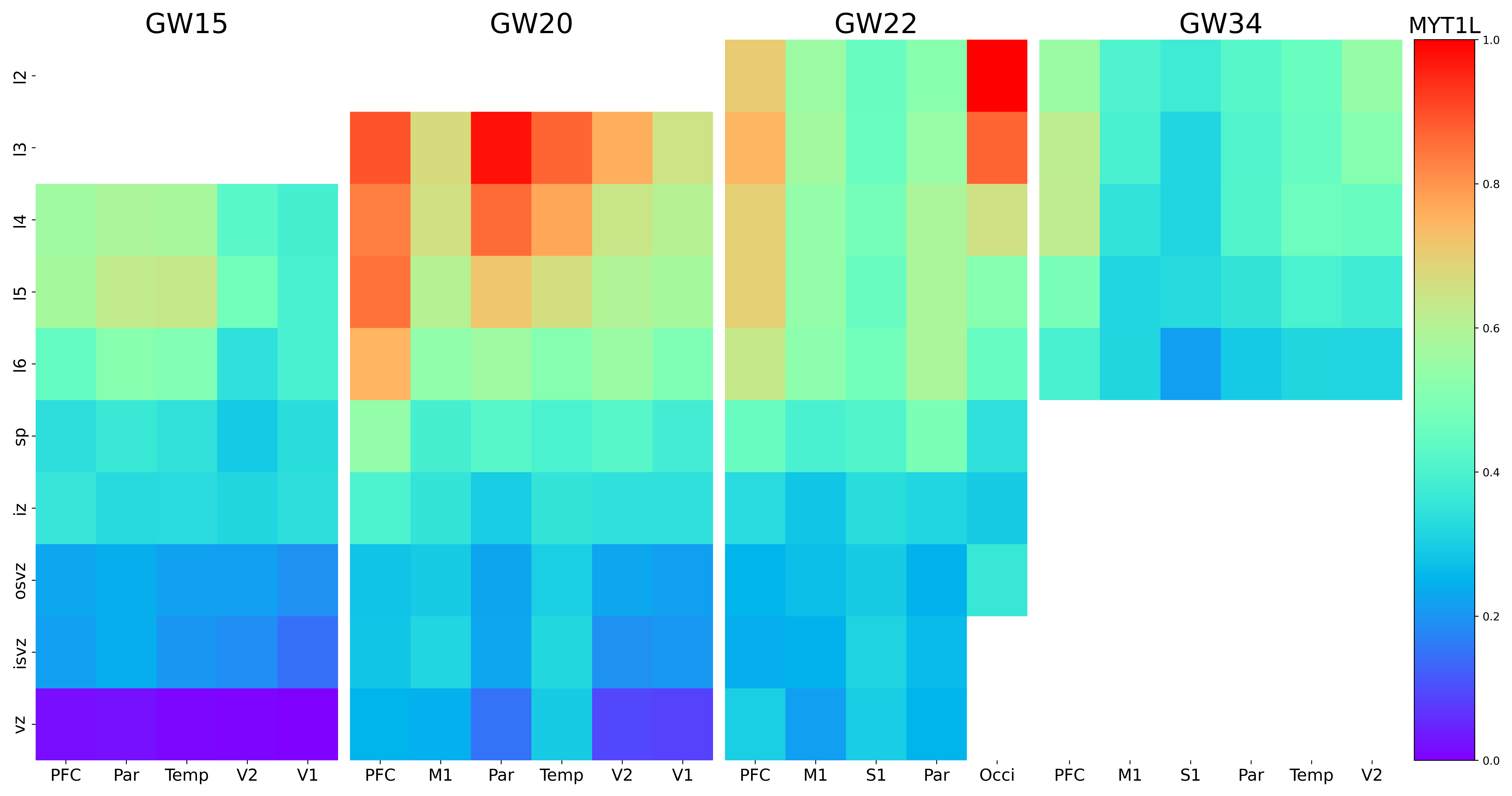

### NAB2.png

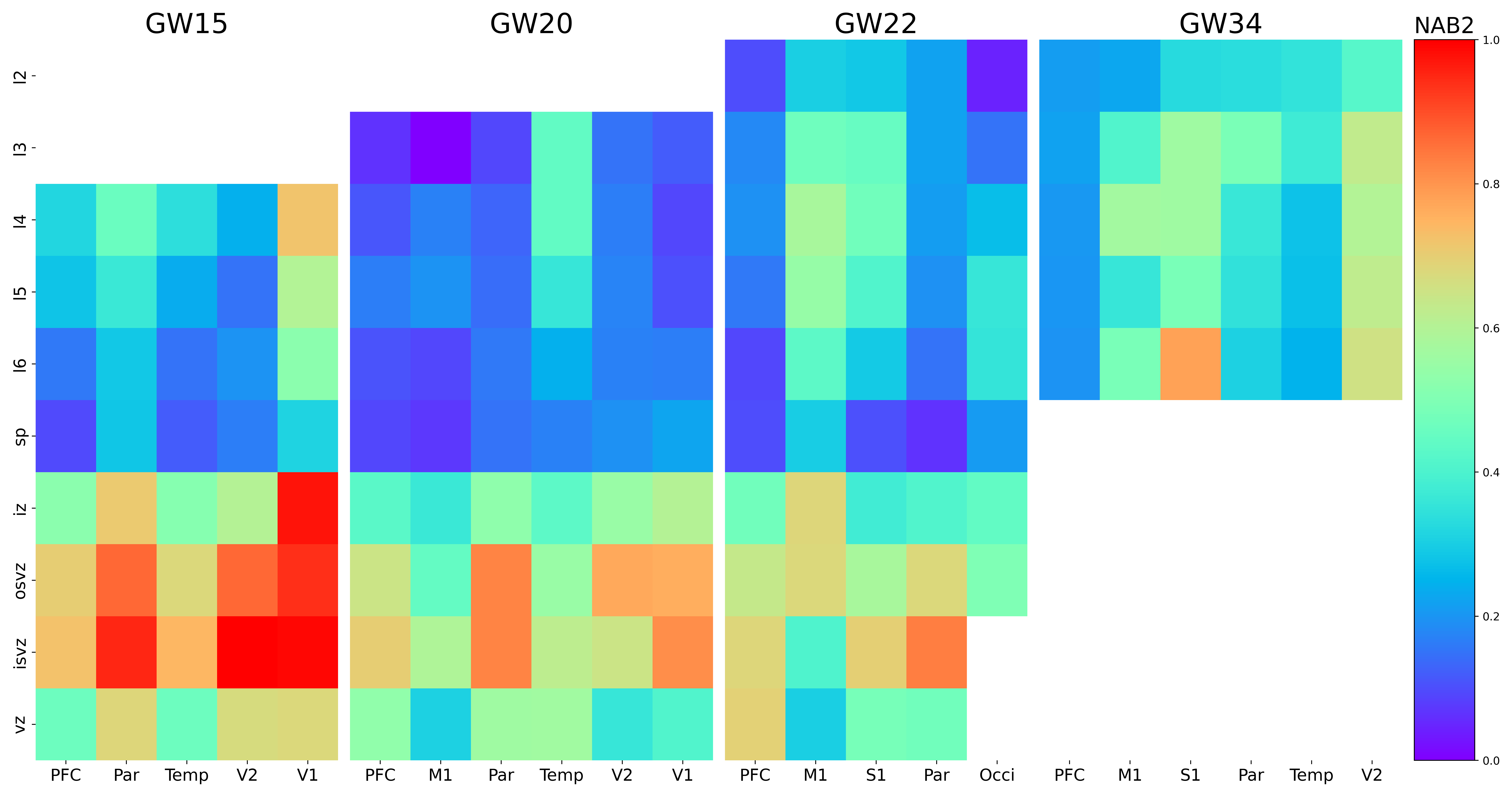

### NAPA-AS1.png

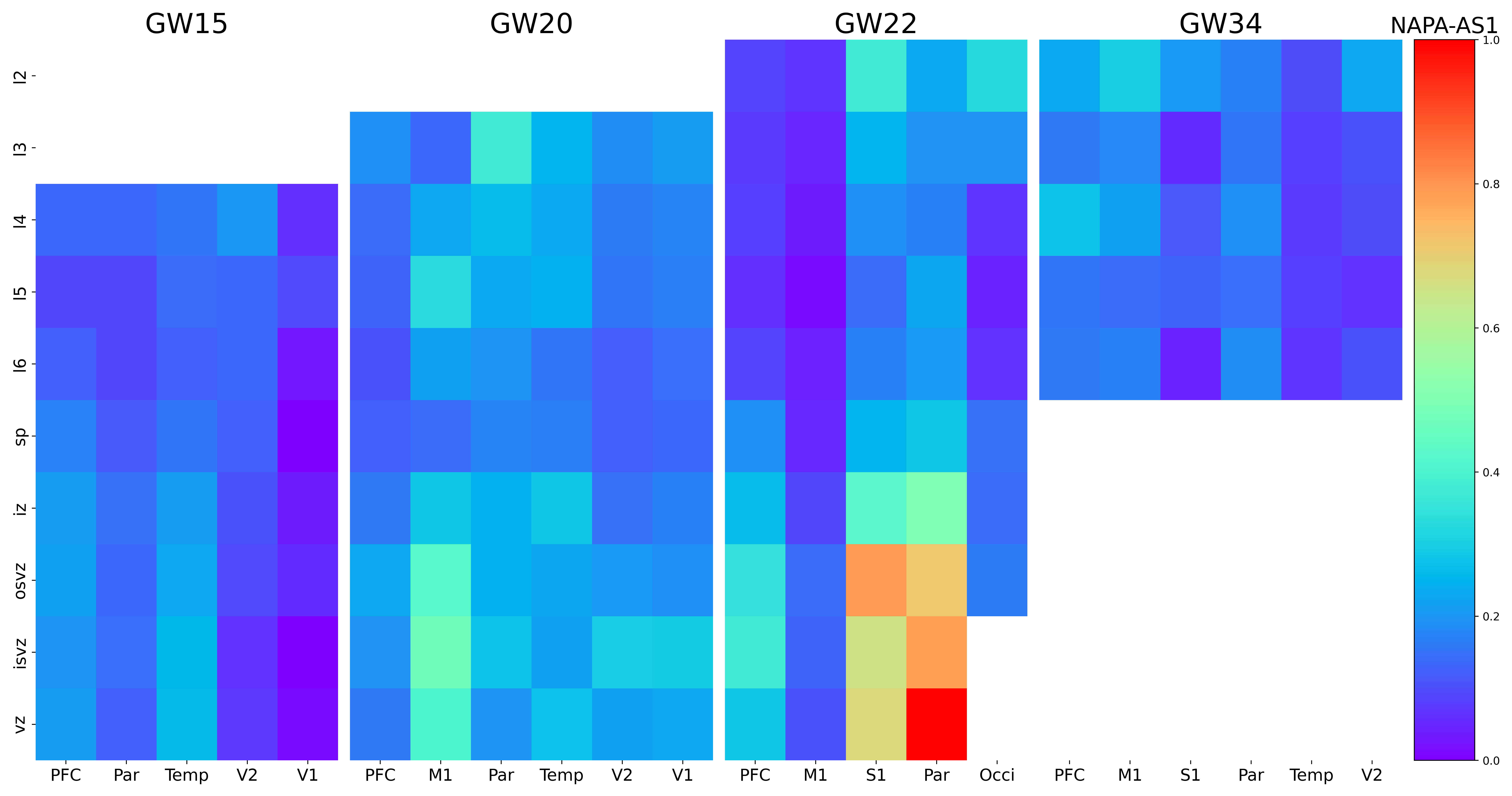

### NDST4.png

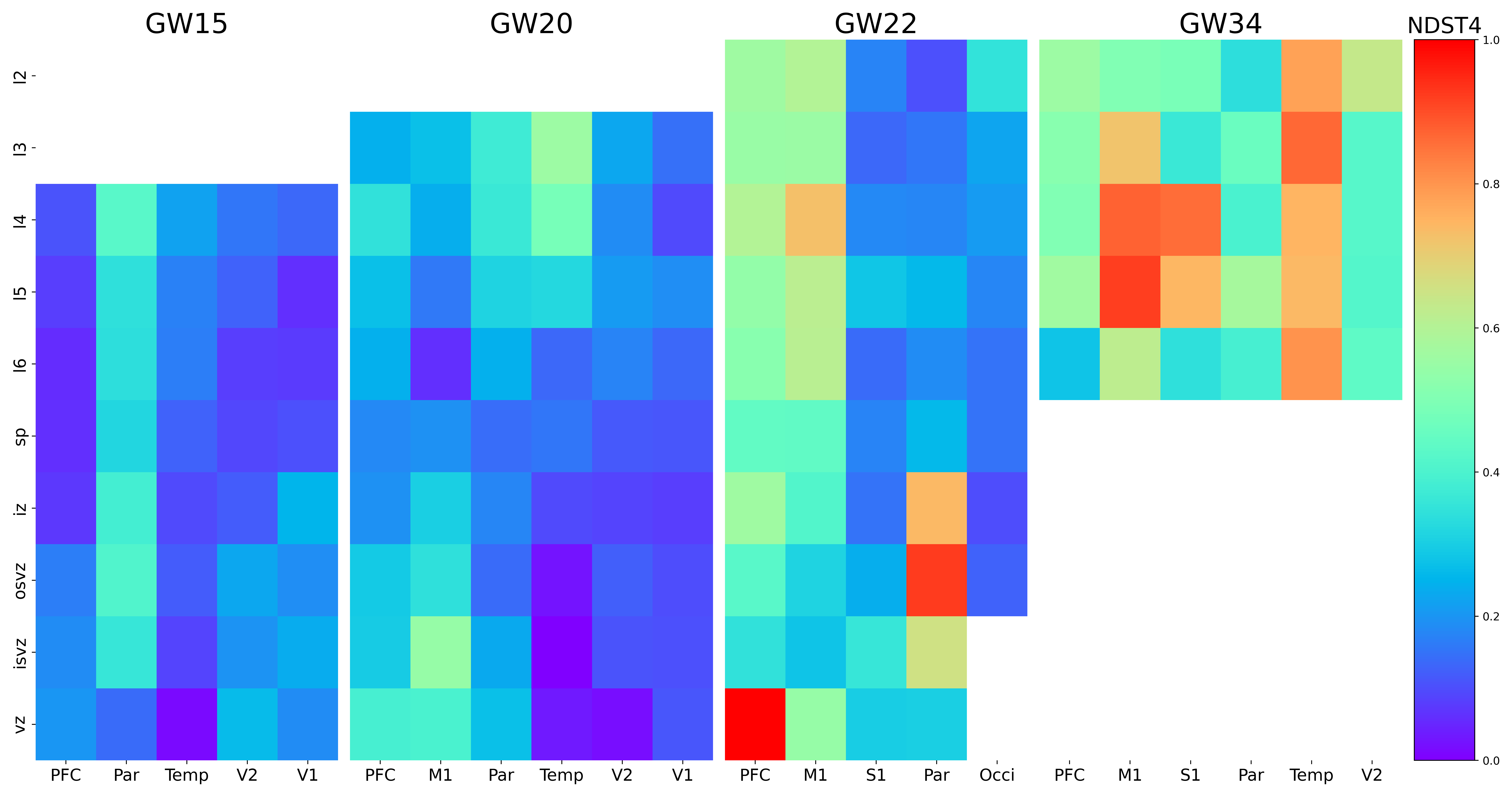

### NECTIN3.png

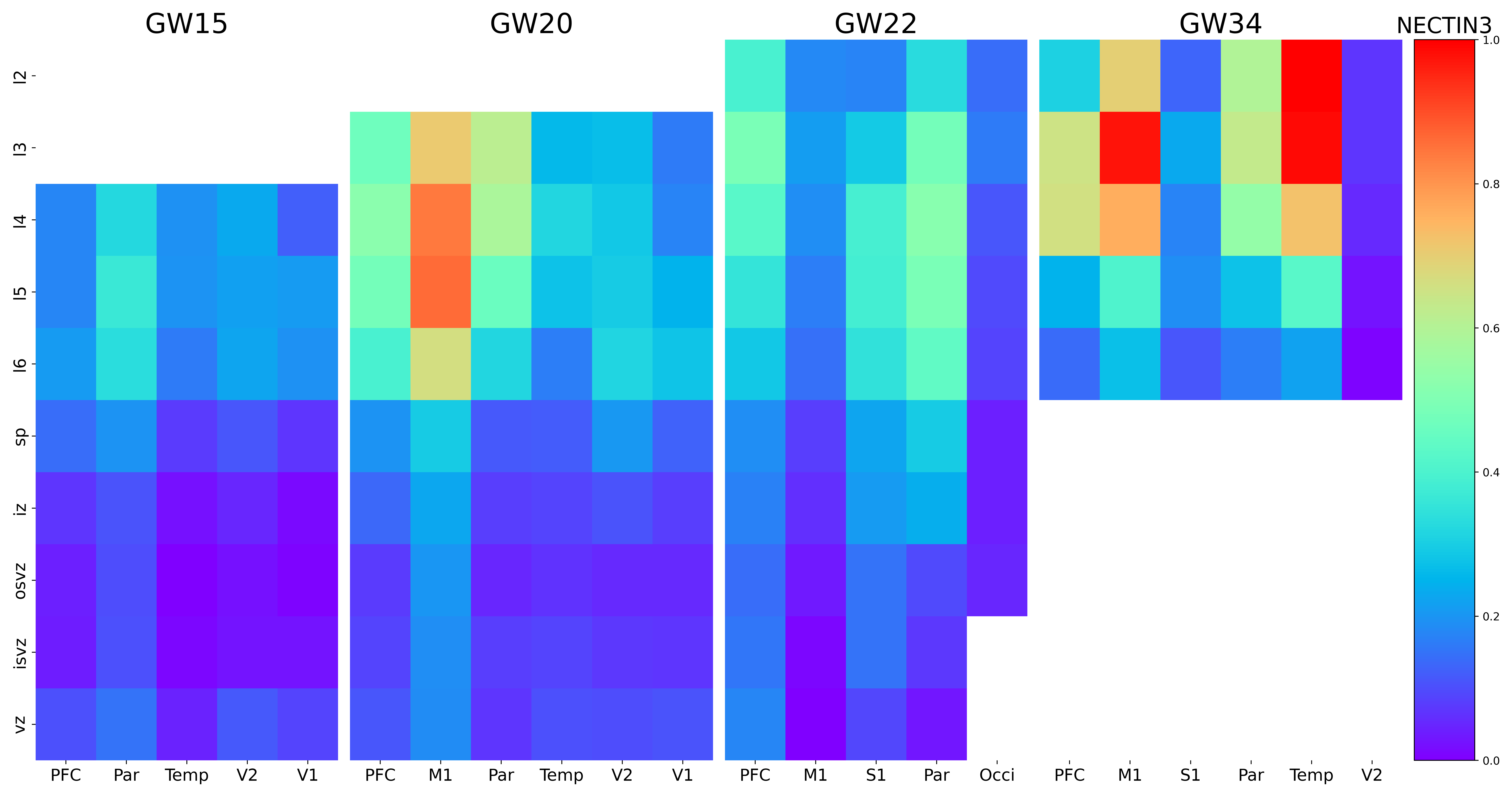

### NEFM.png

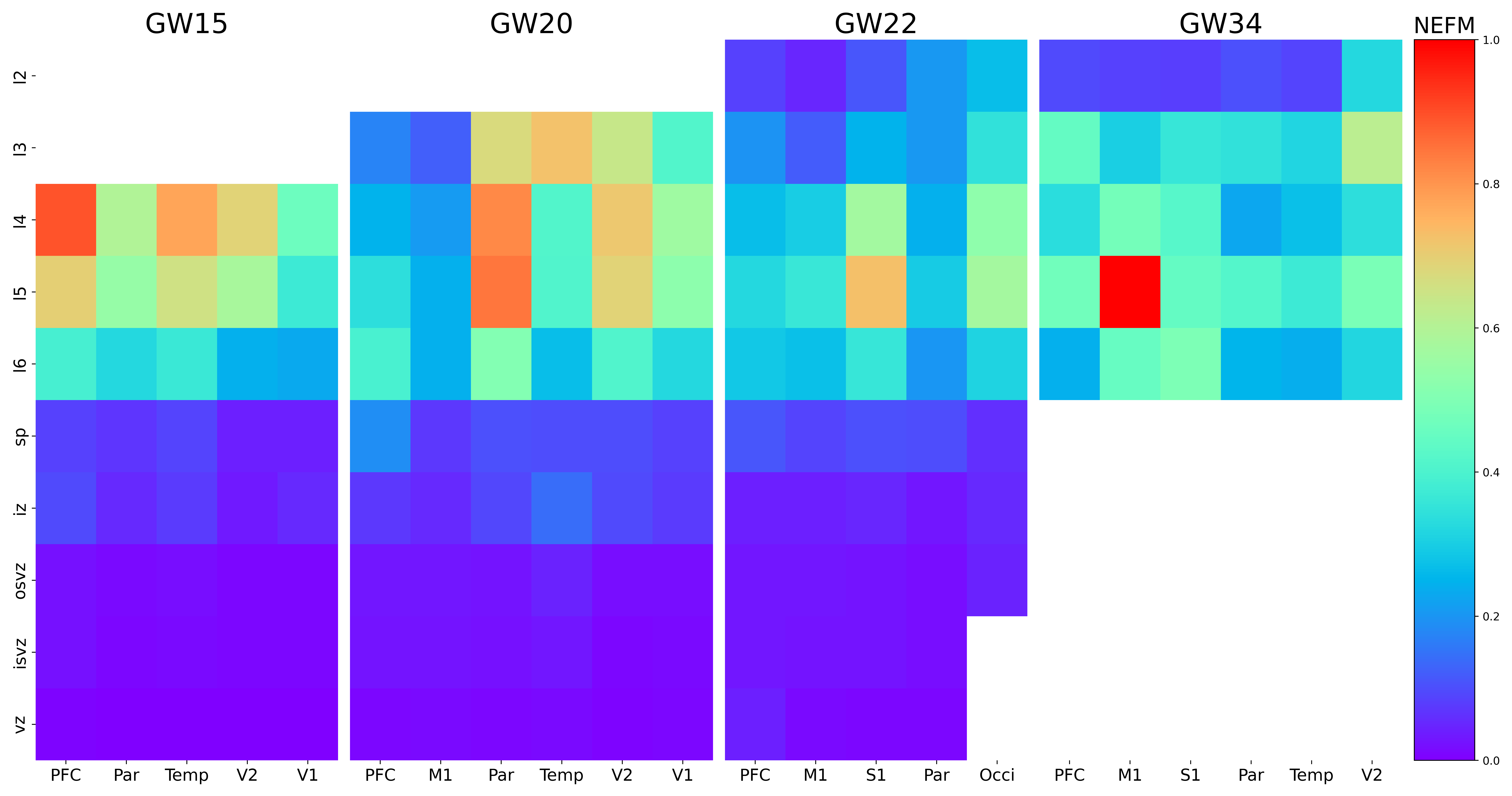

### NEUROD1.png

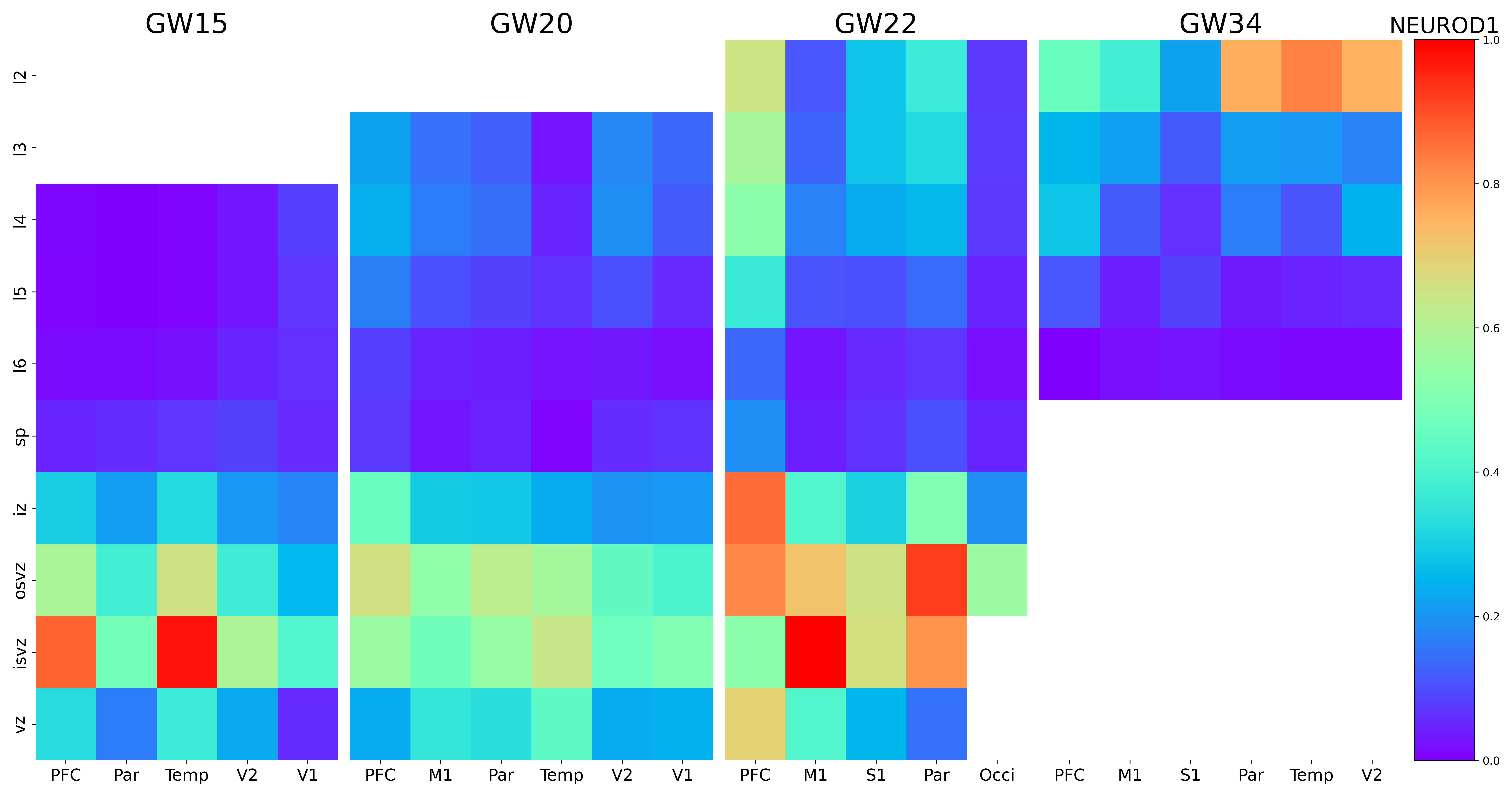

### NEUROD2.png

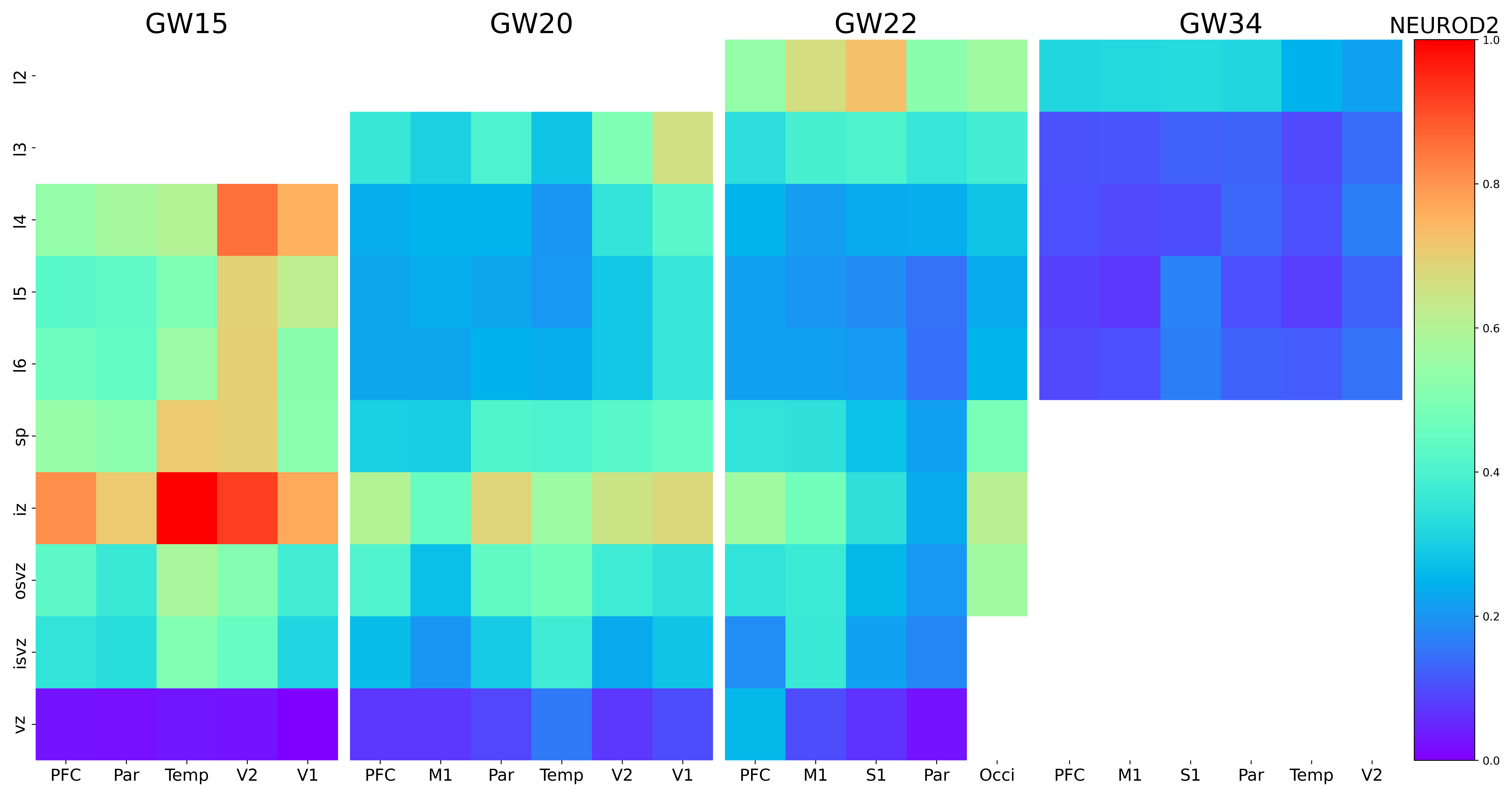

### NEUROD4.png

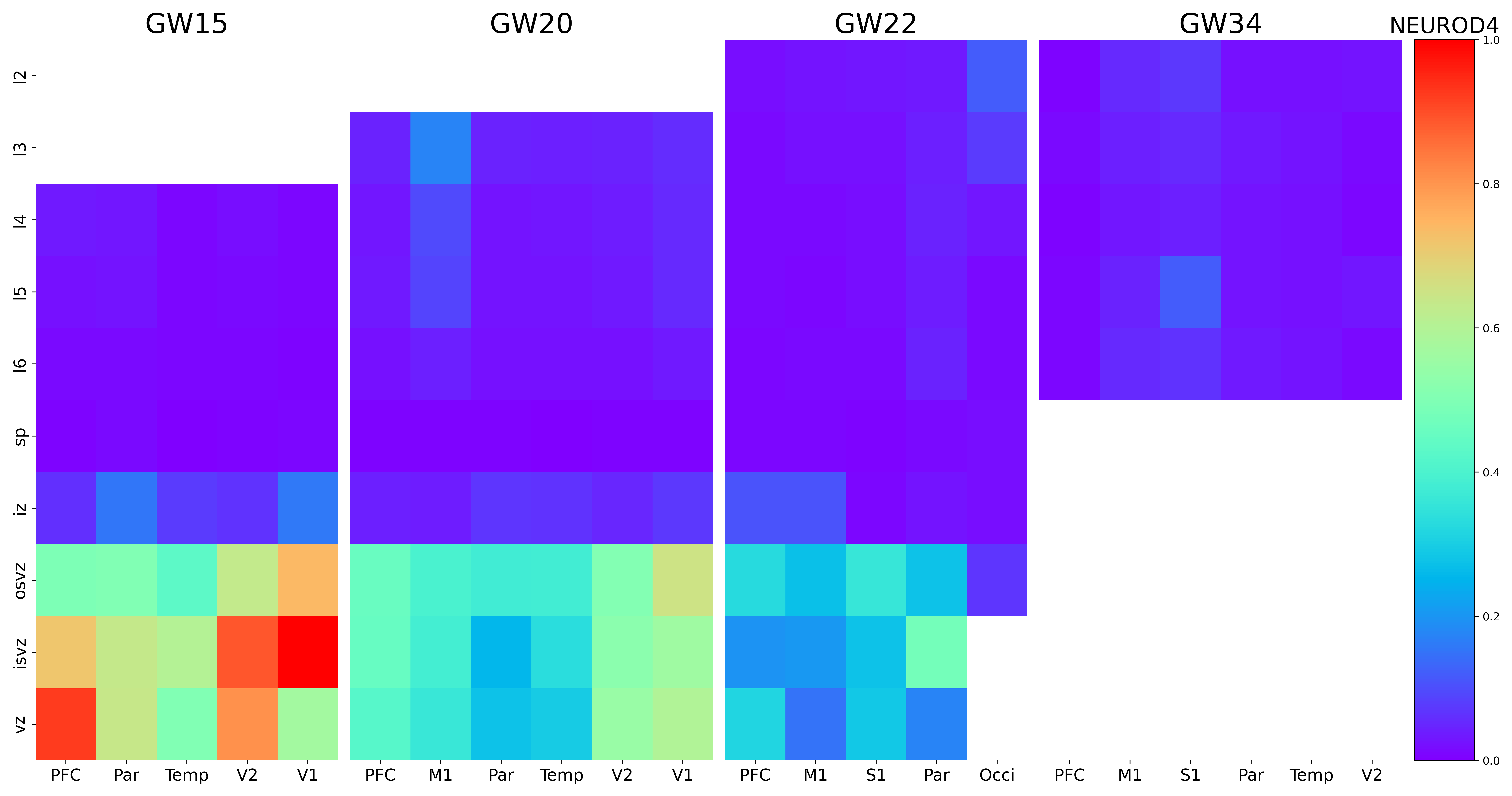

### NEUROD6.png

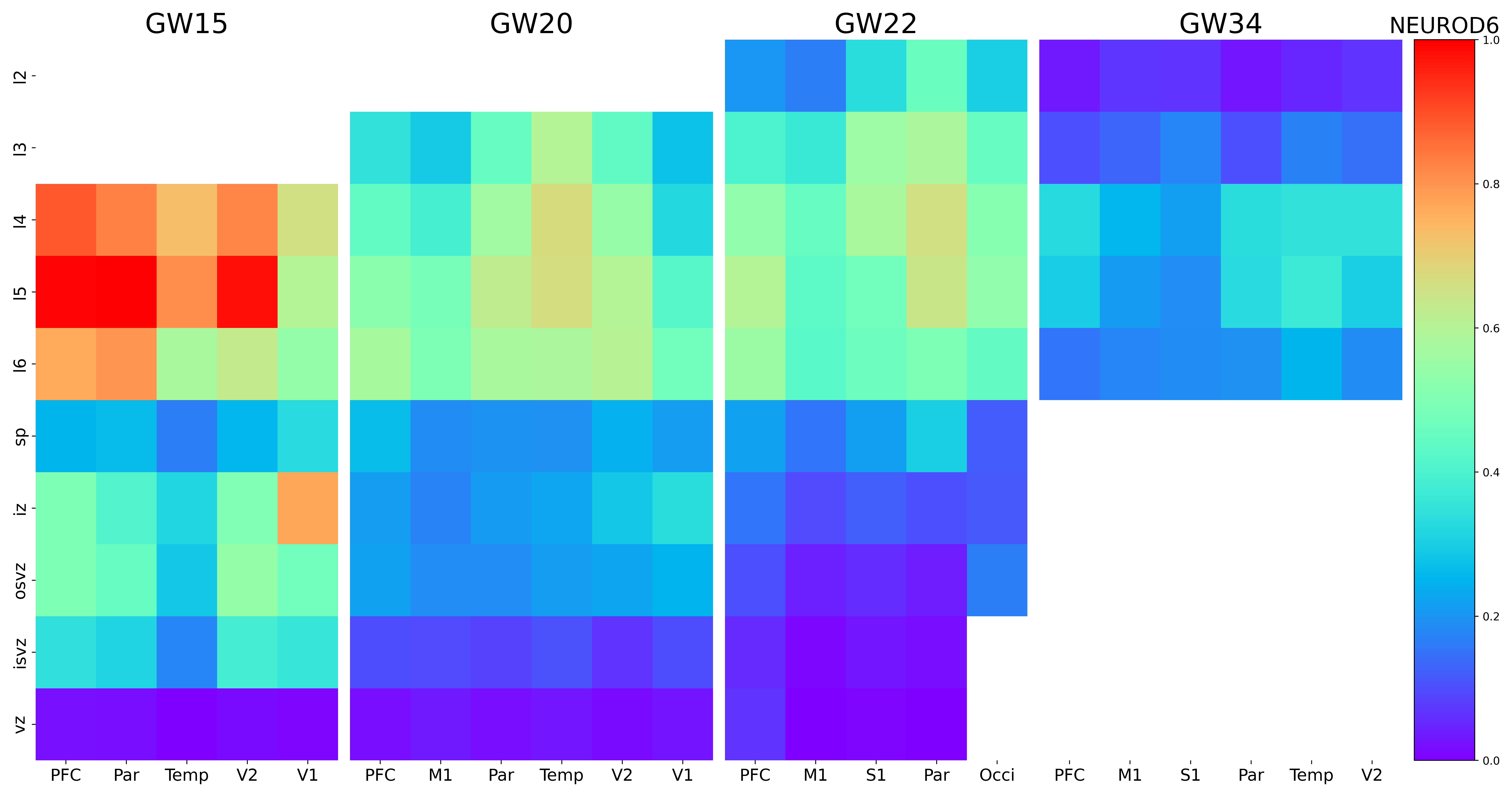

### NEUROG1.png

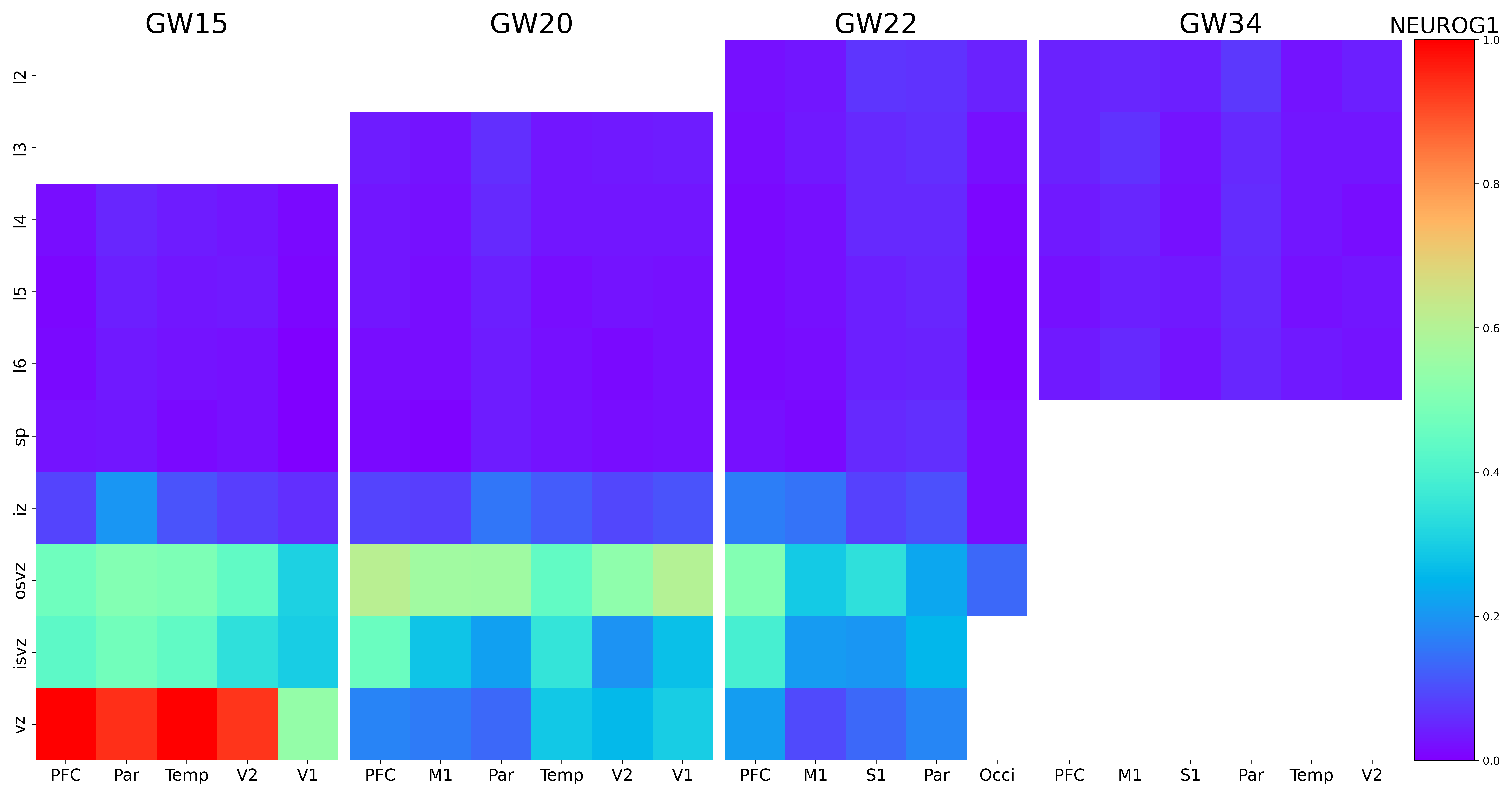

### NEUROG2.png

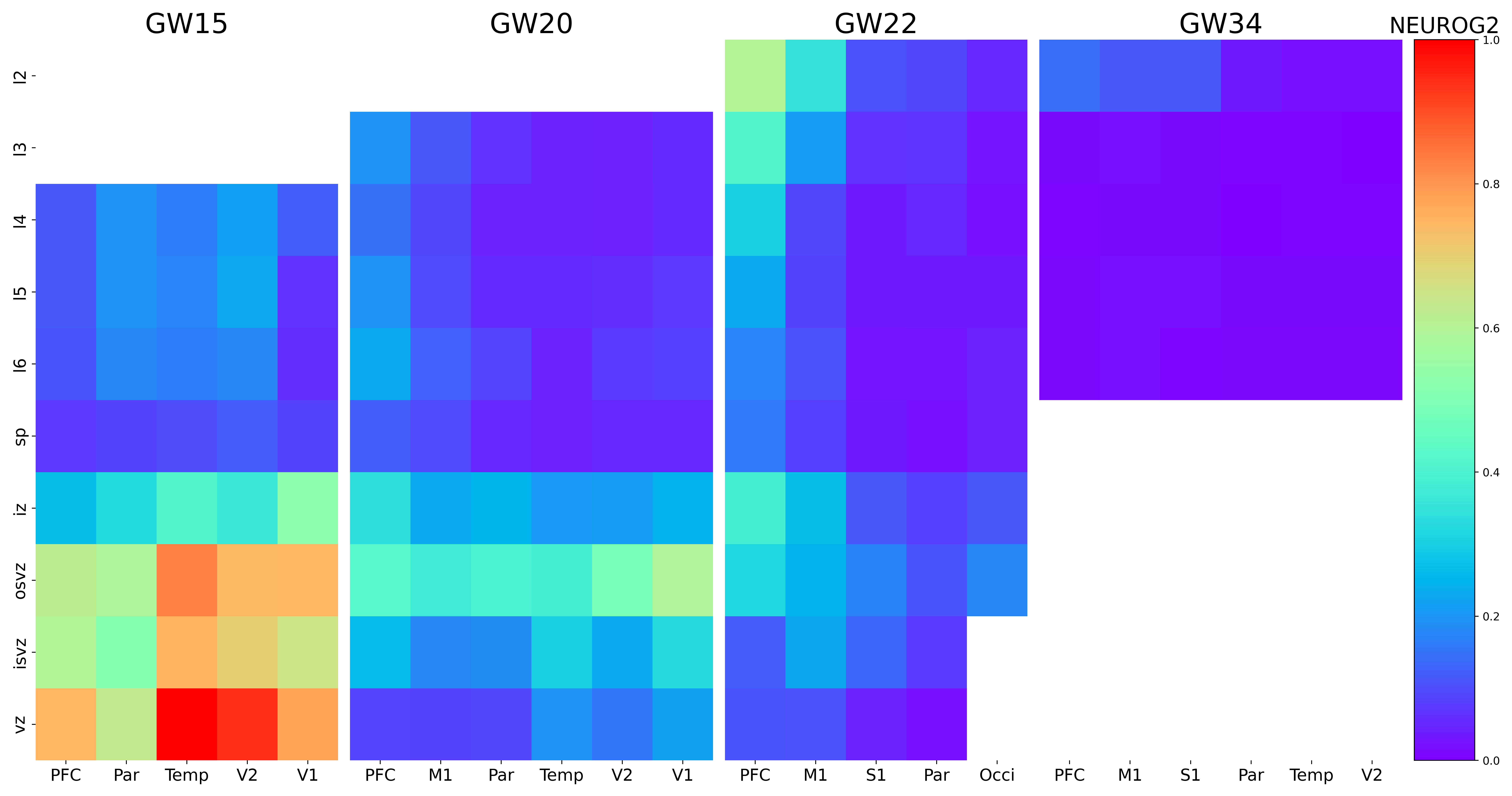

### NFIB.png

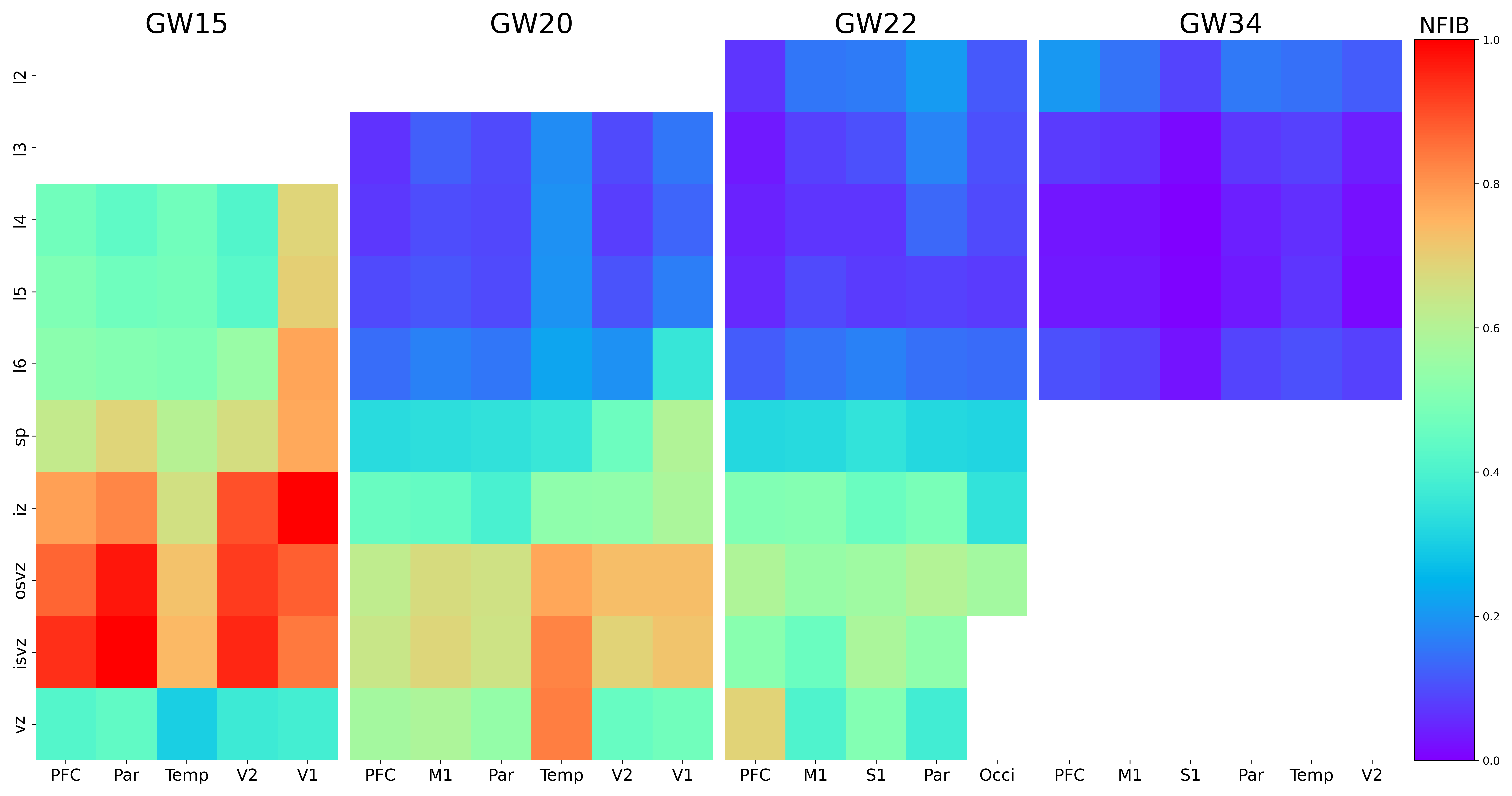

### NHLH1.png

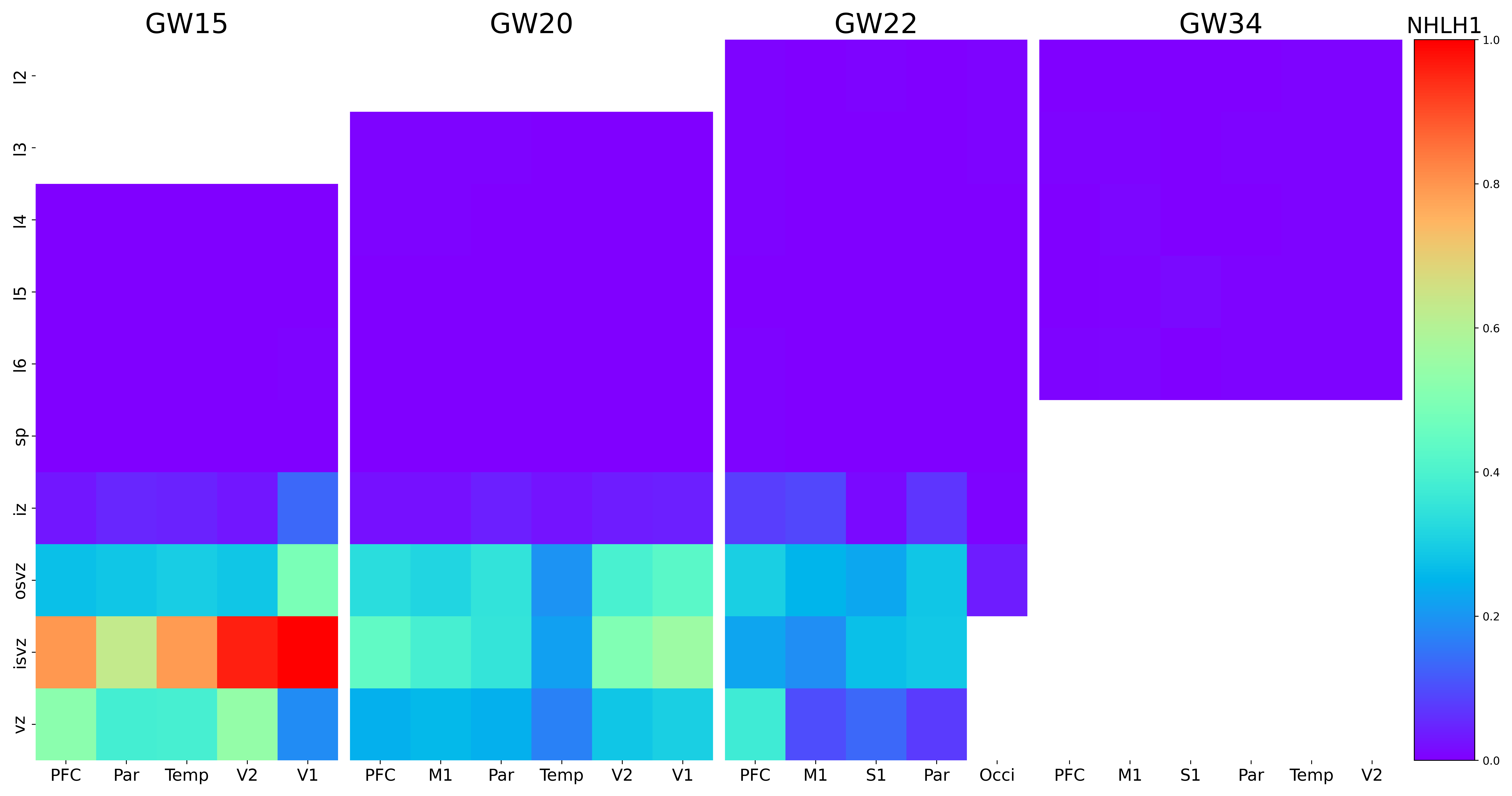

### NIFK-AS1.png

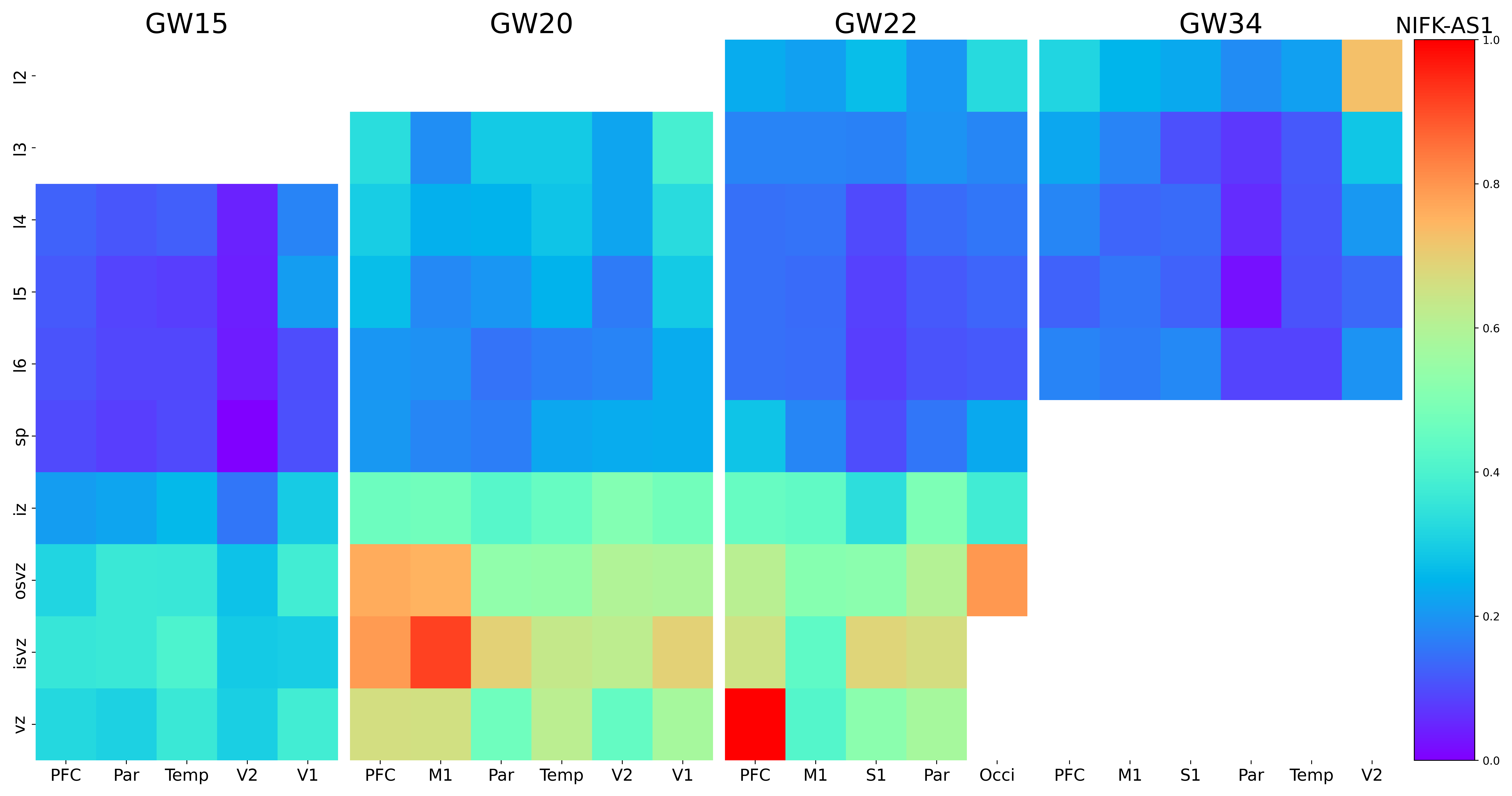

### NIPBL-DT.png

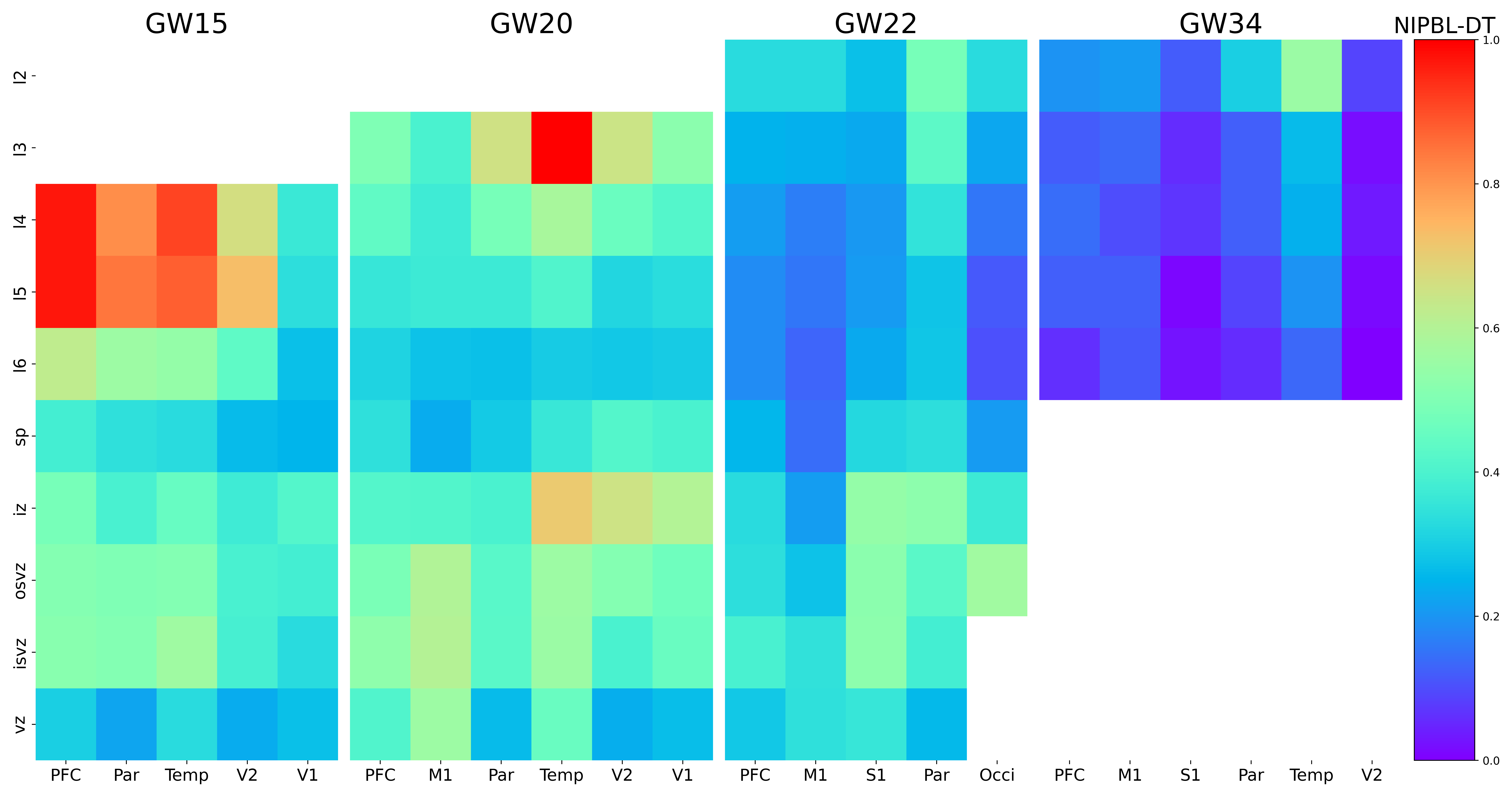

### NIPBL.png

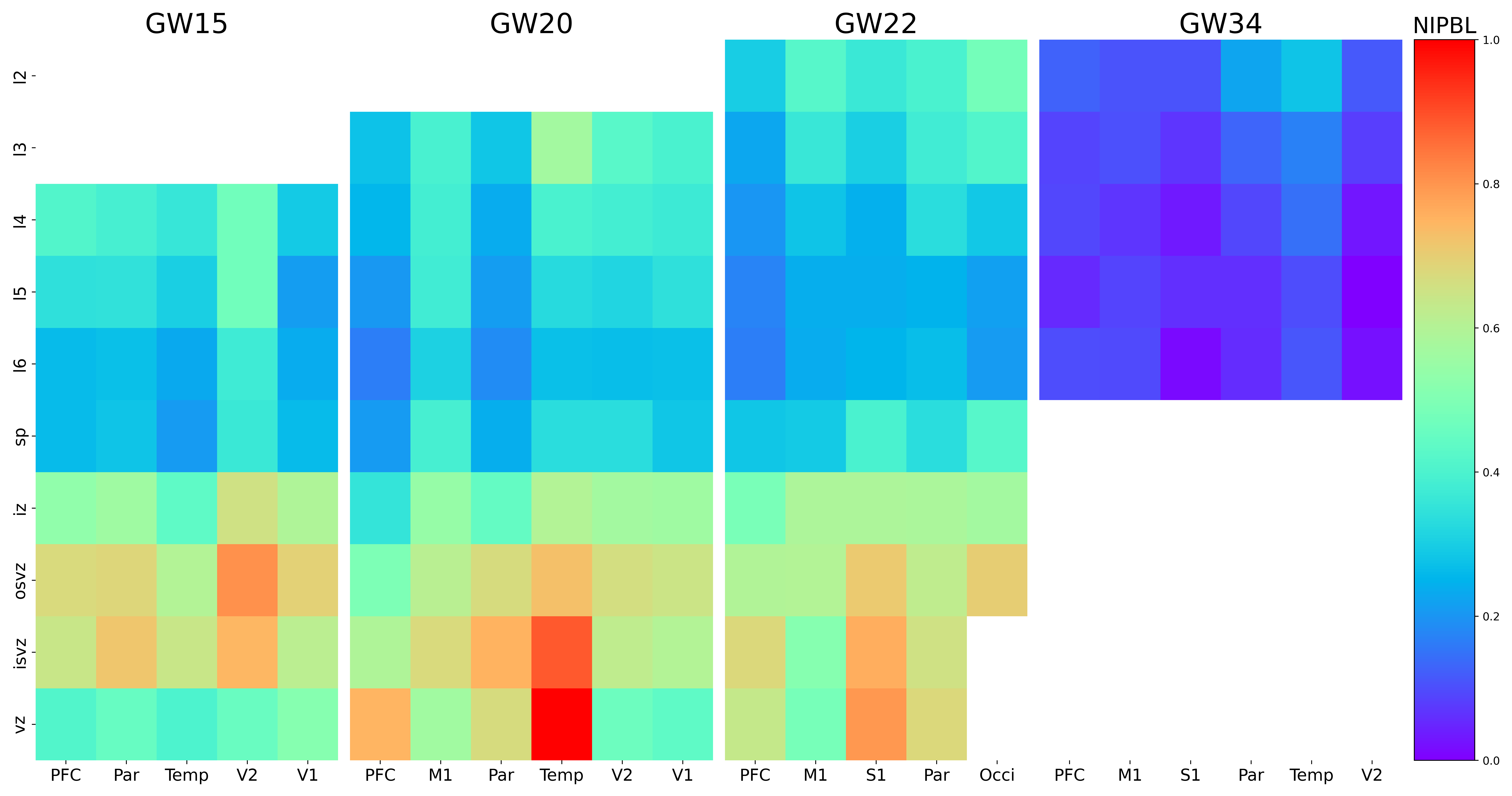

### NKX2-1.png

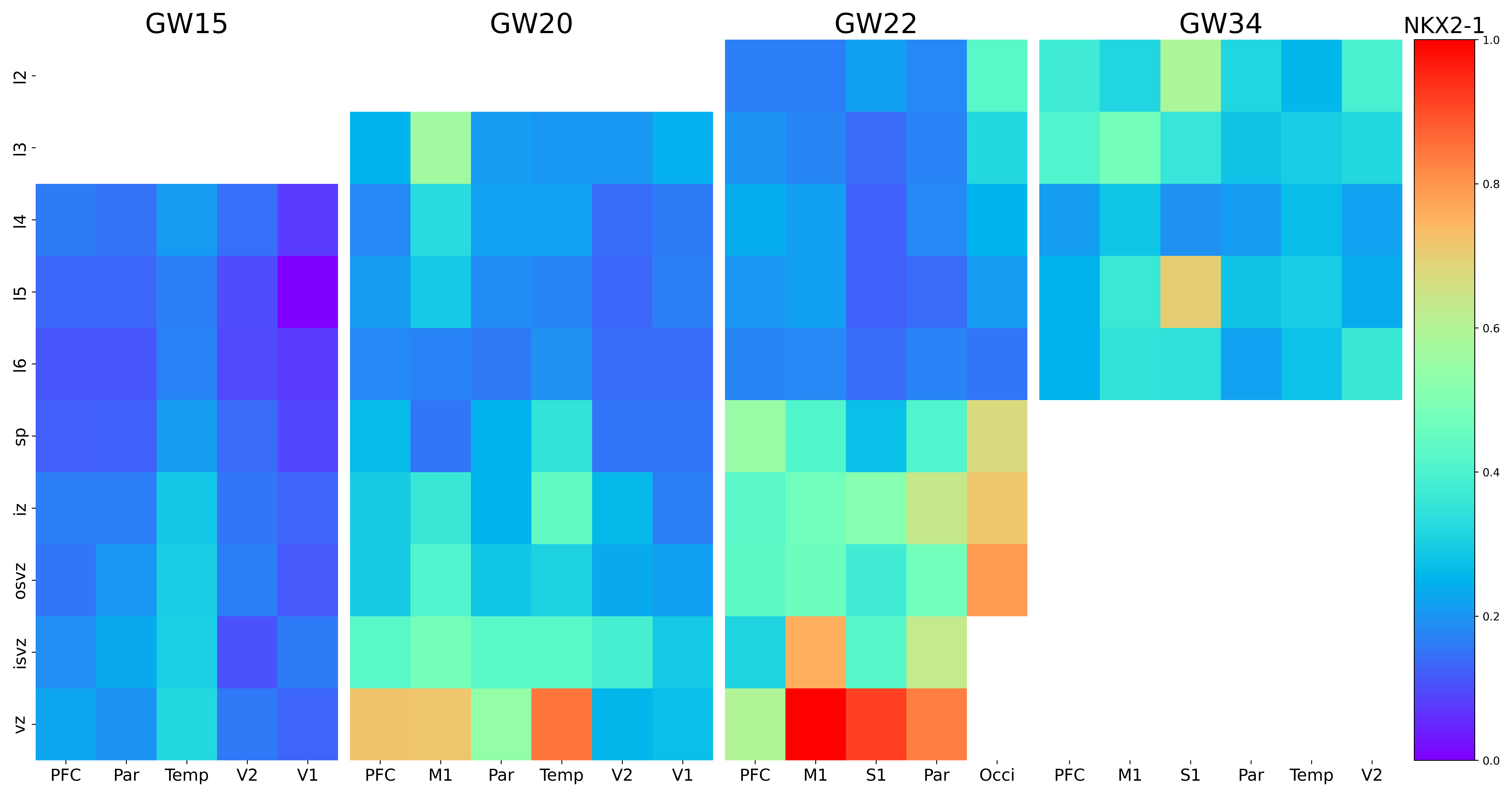

### NPNT.png

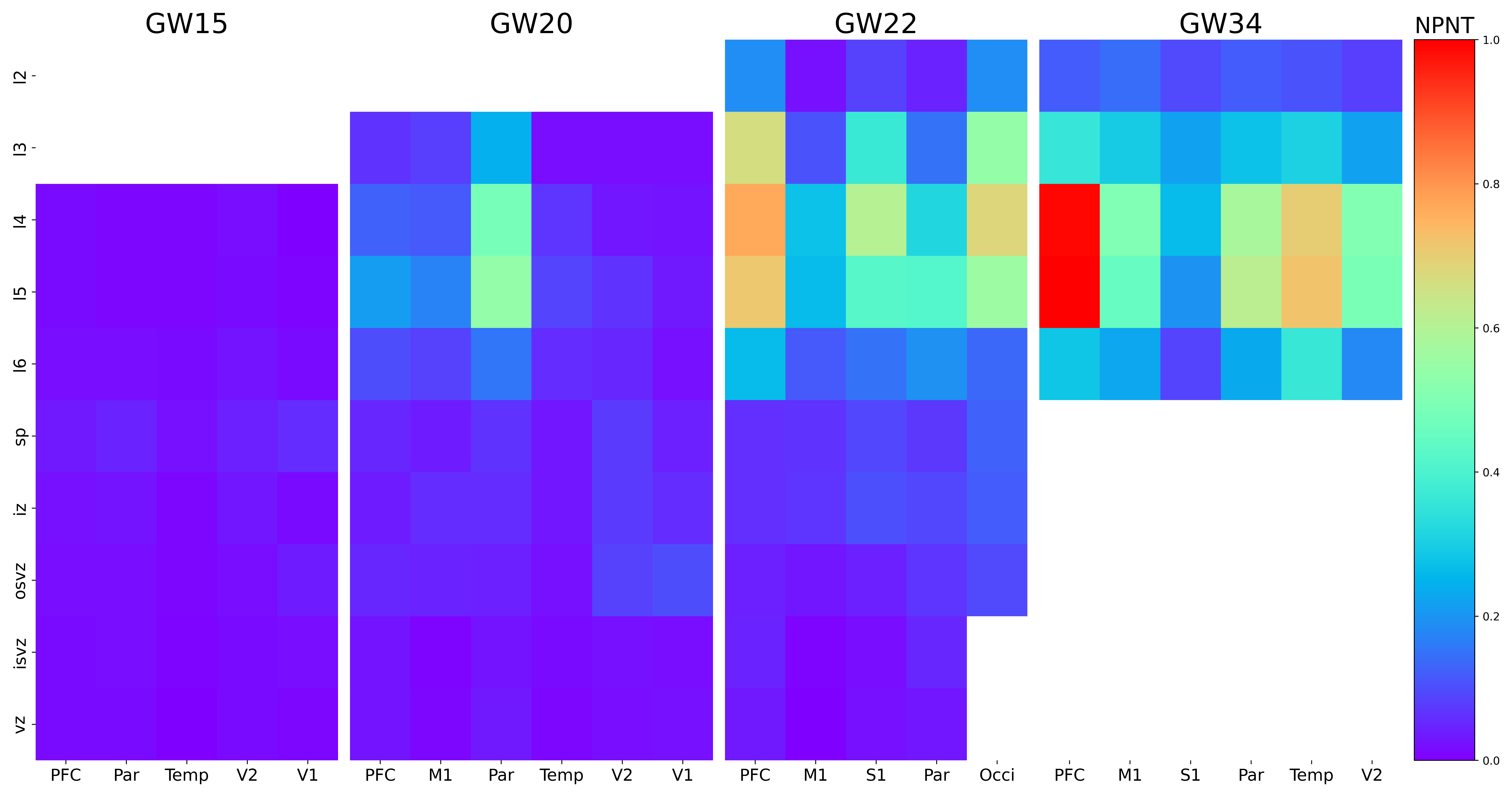

### NPR3.png

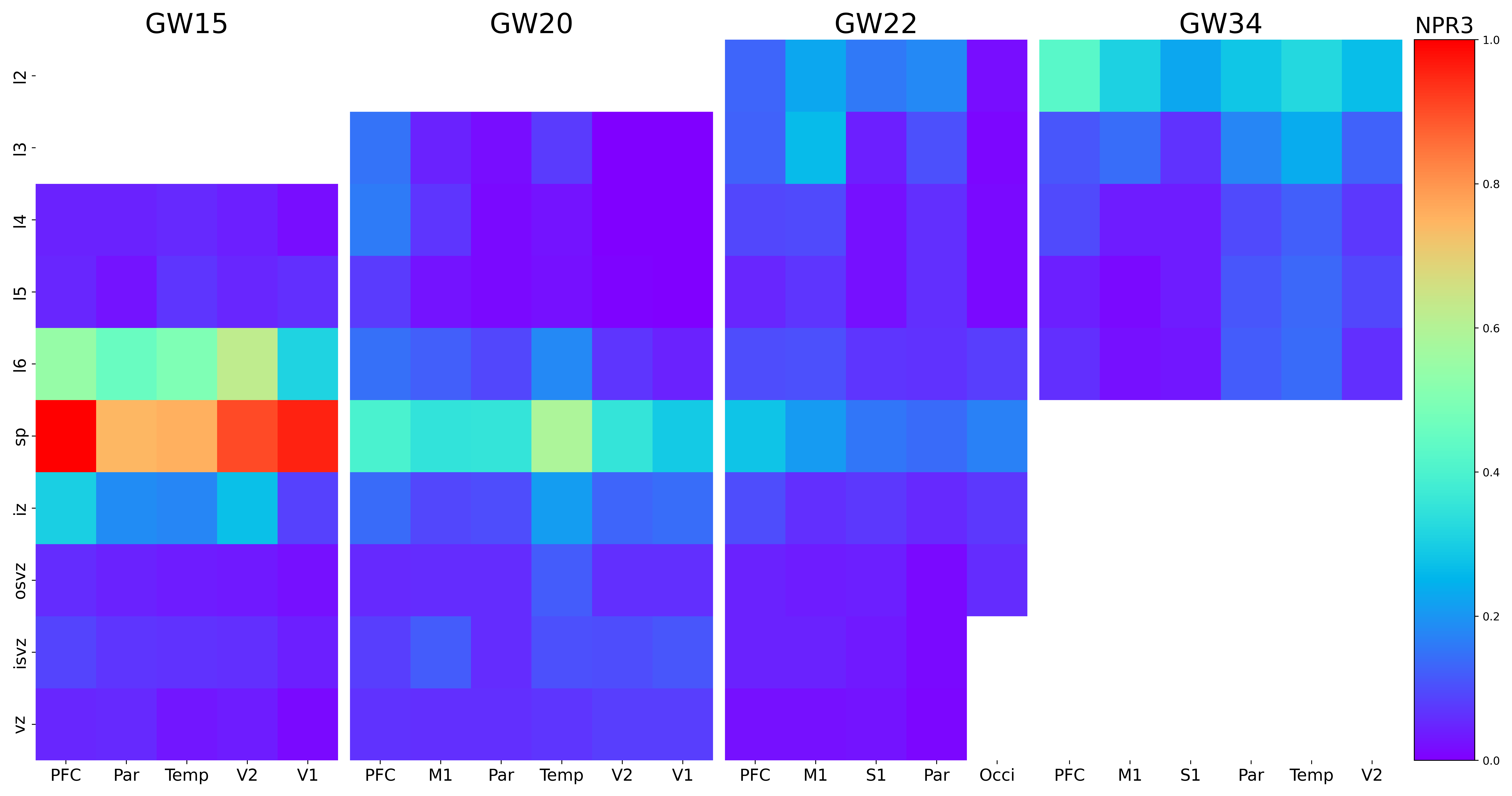

### NPY.png

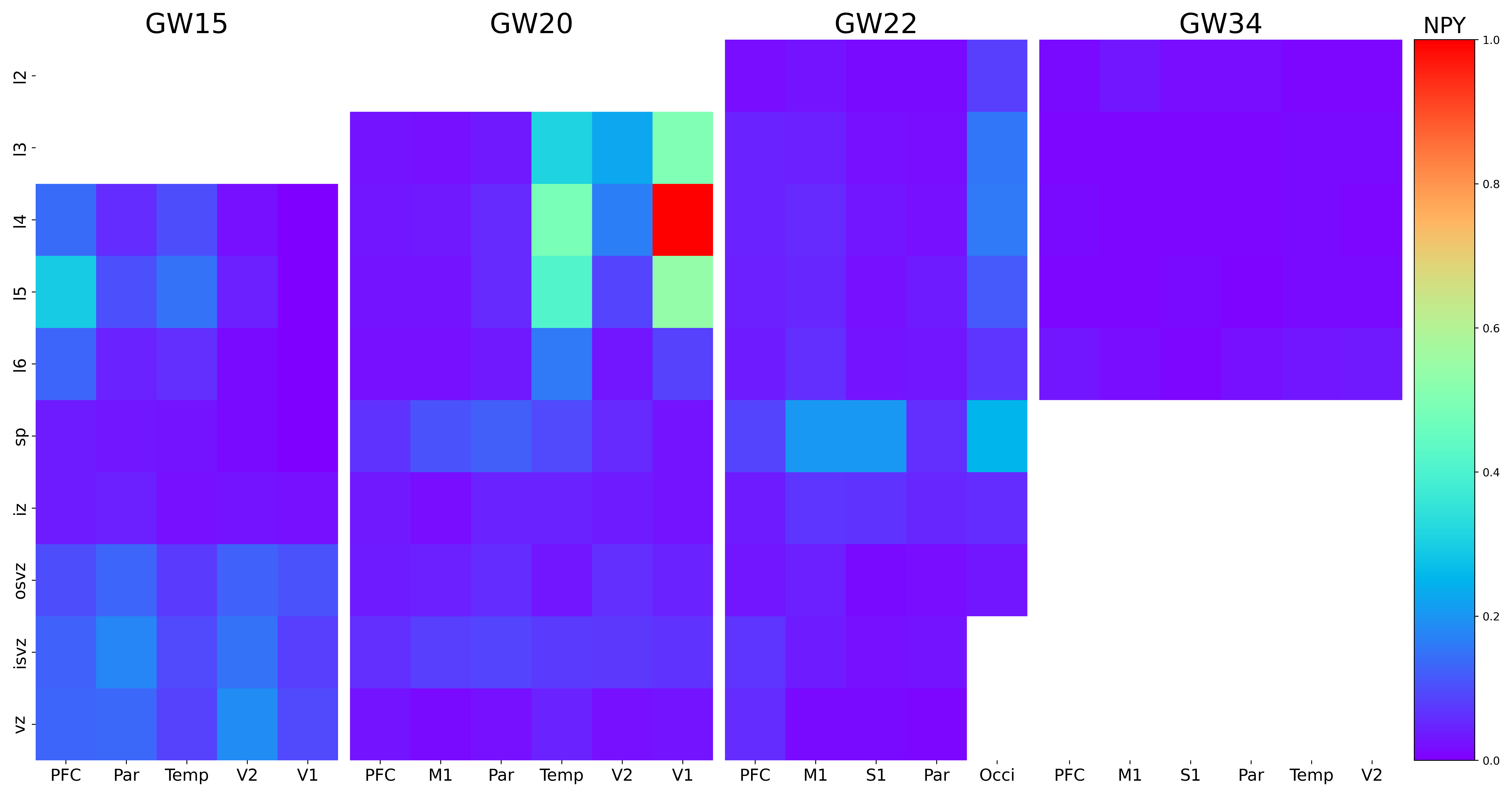

### NR2F1-AS1.png

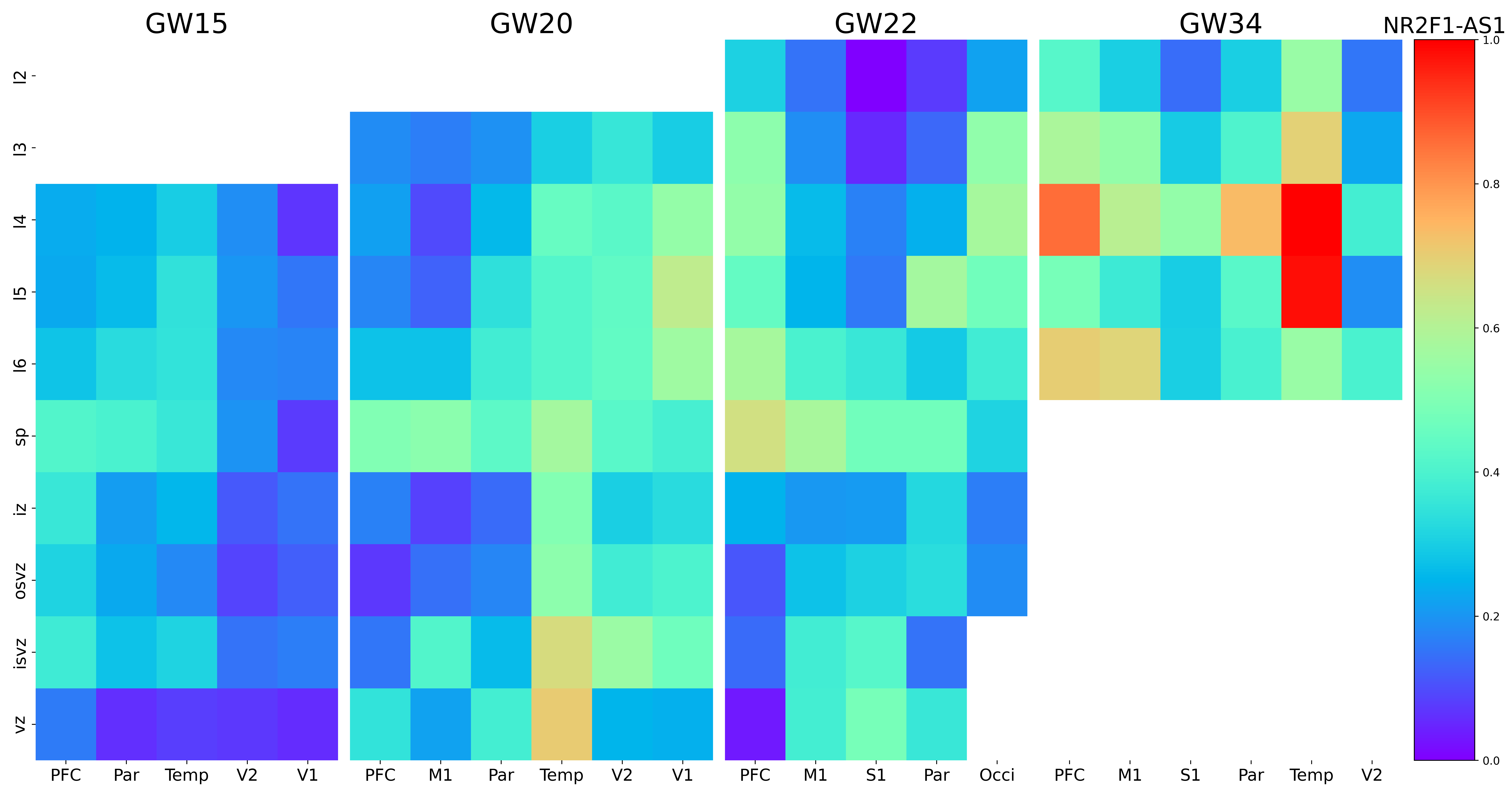

### NR2F1.png

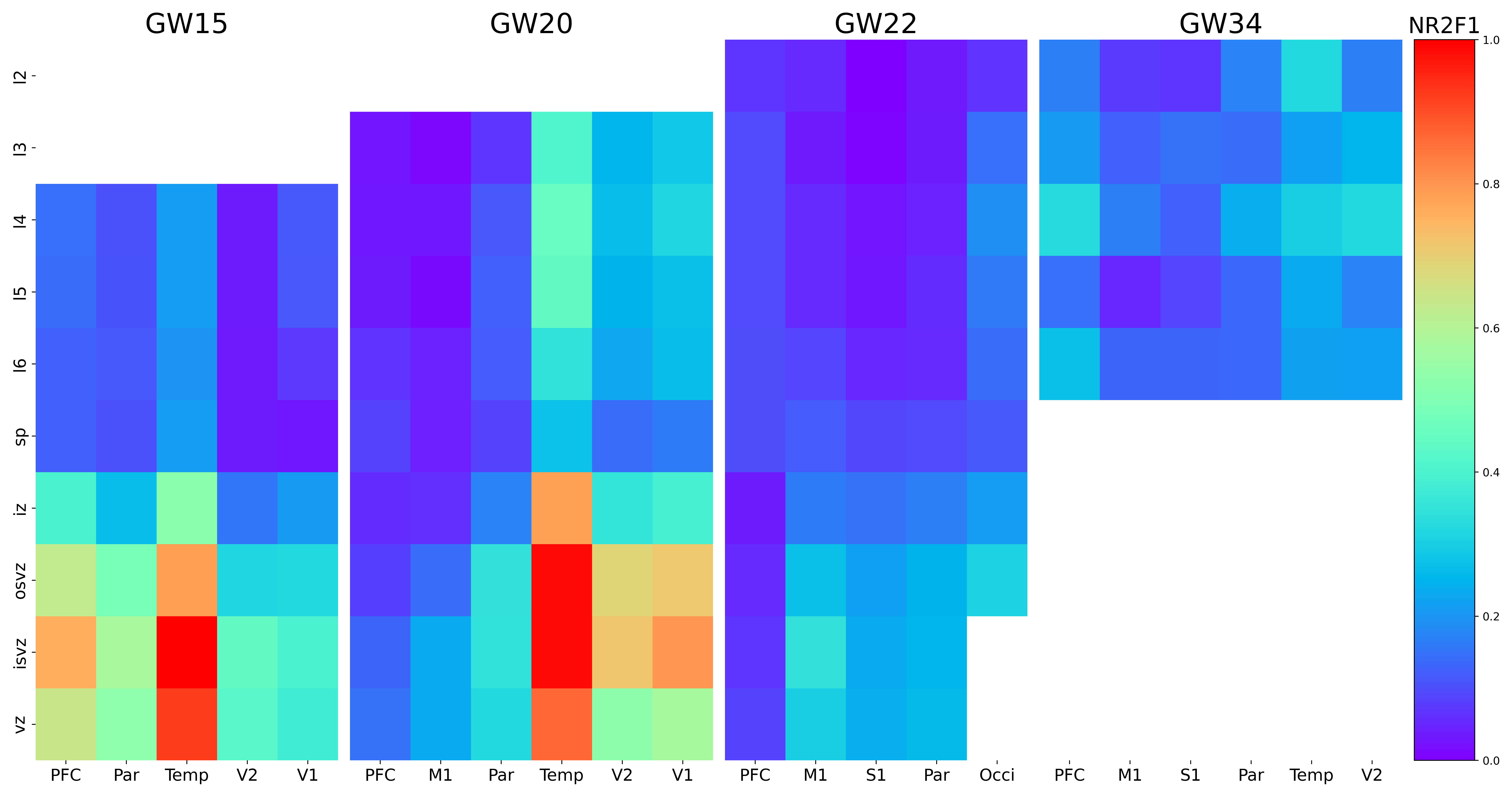

### NR4A2.png

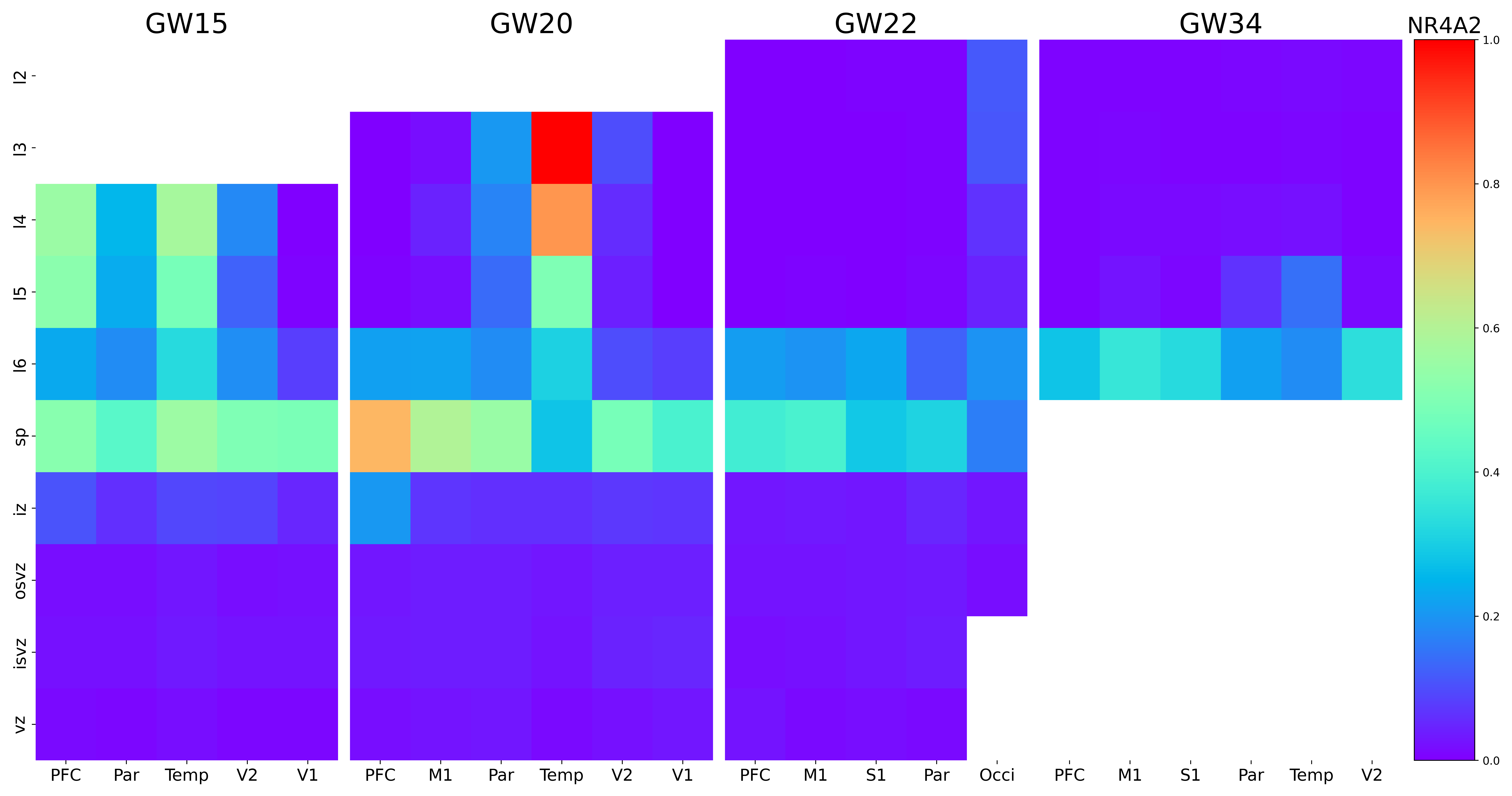

### NRN1.png

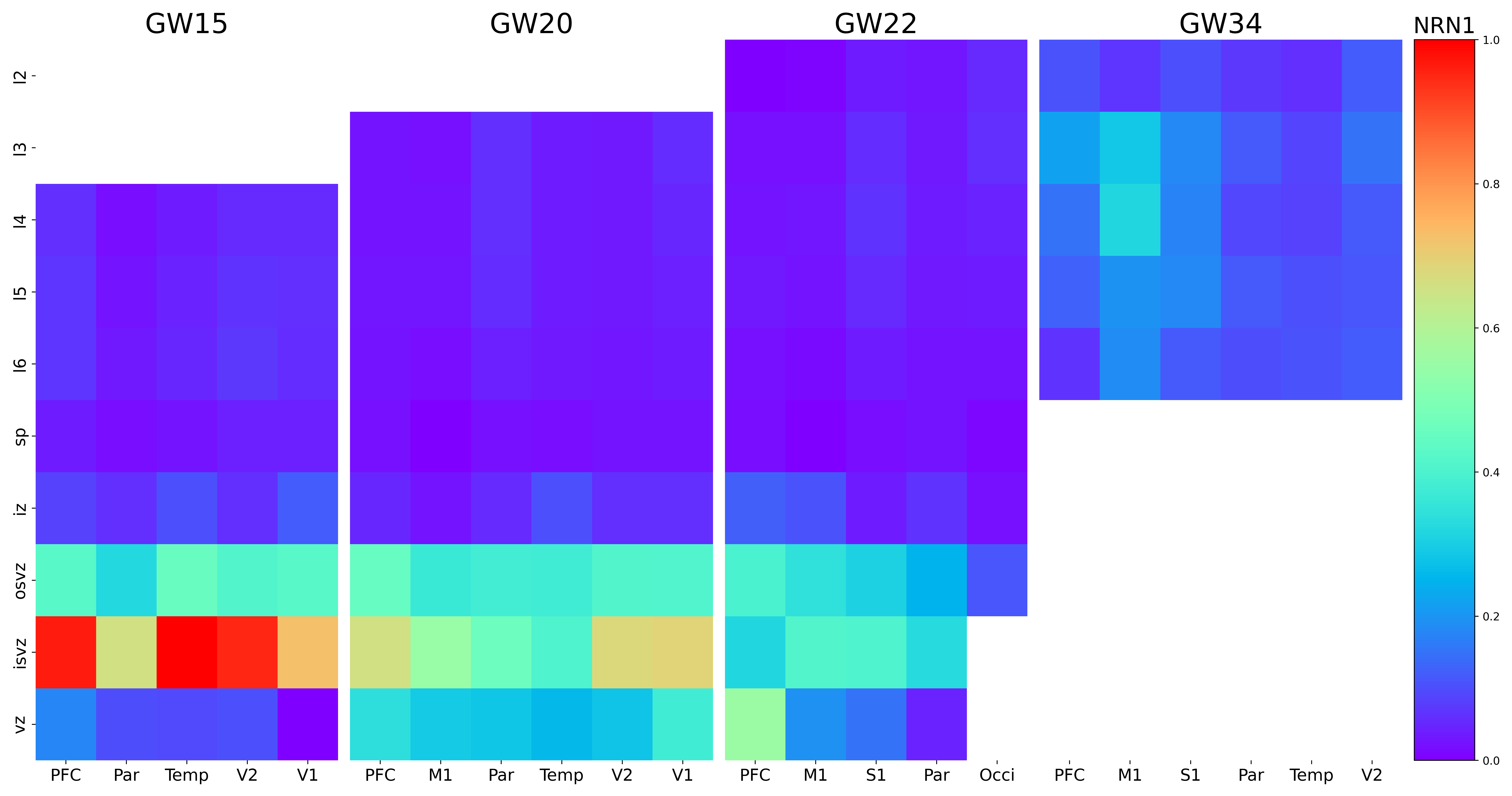
