## Supplementary figures and images for "Spatial Single-cell Analysis Decodes Cortical Layer and Area Specification"

### AC008124.1.png

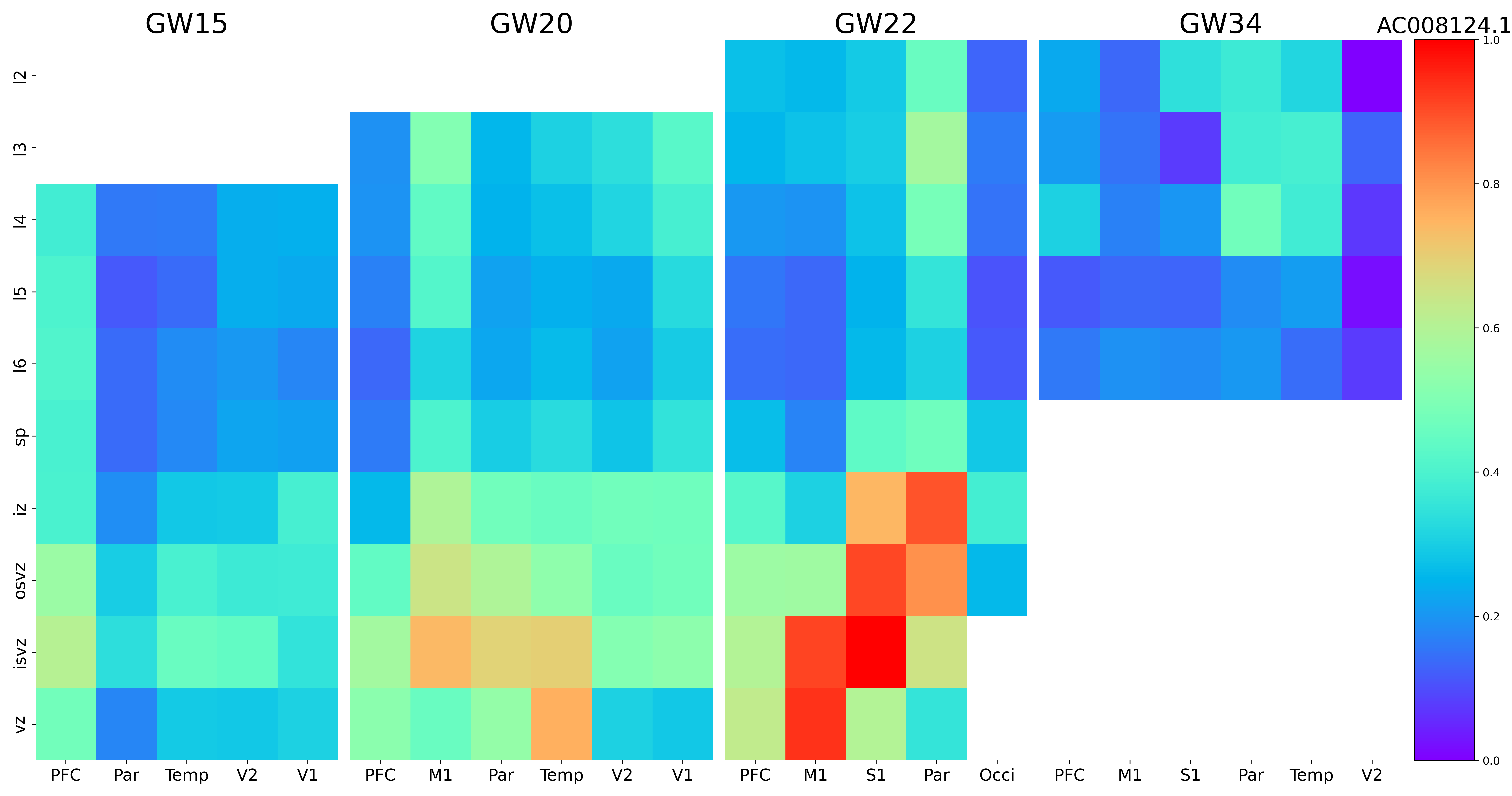

### AC009506.1.png

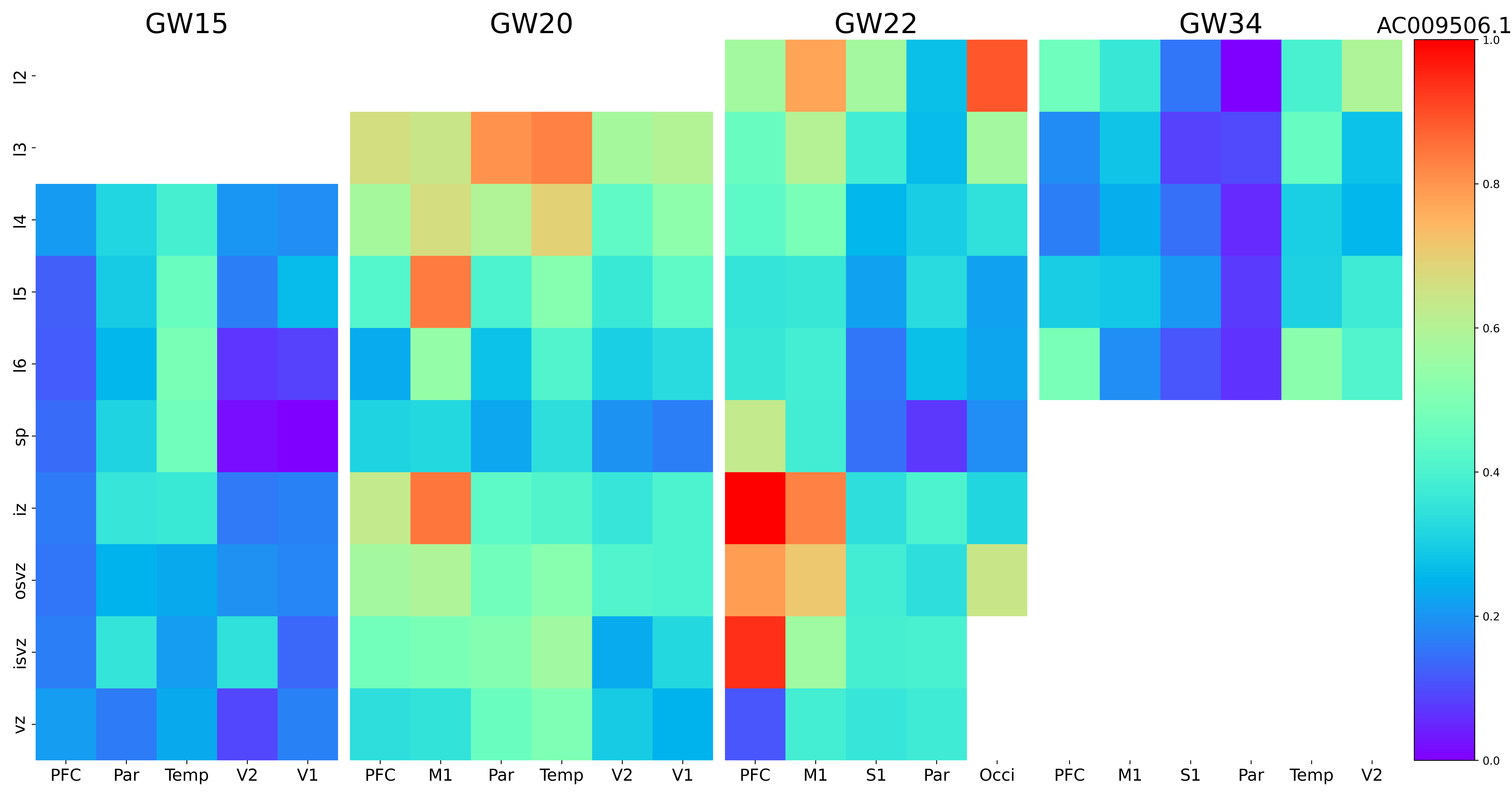

### AC016065.1.png

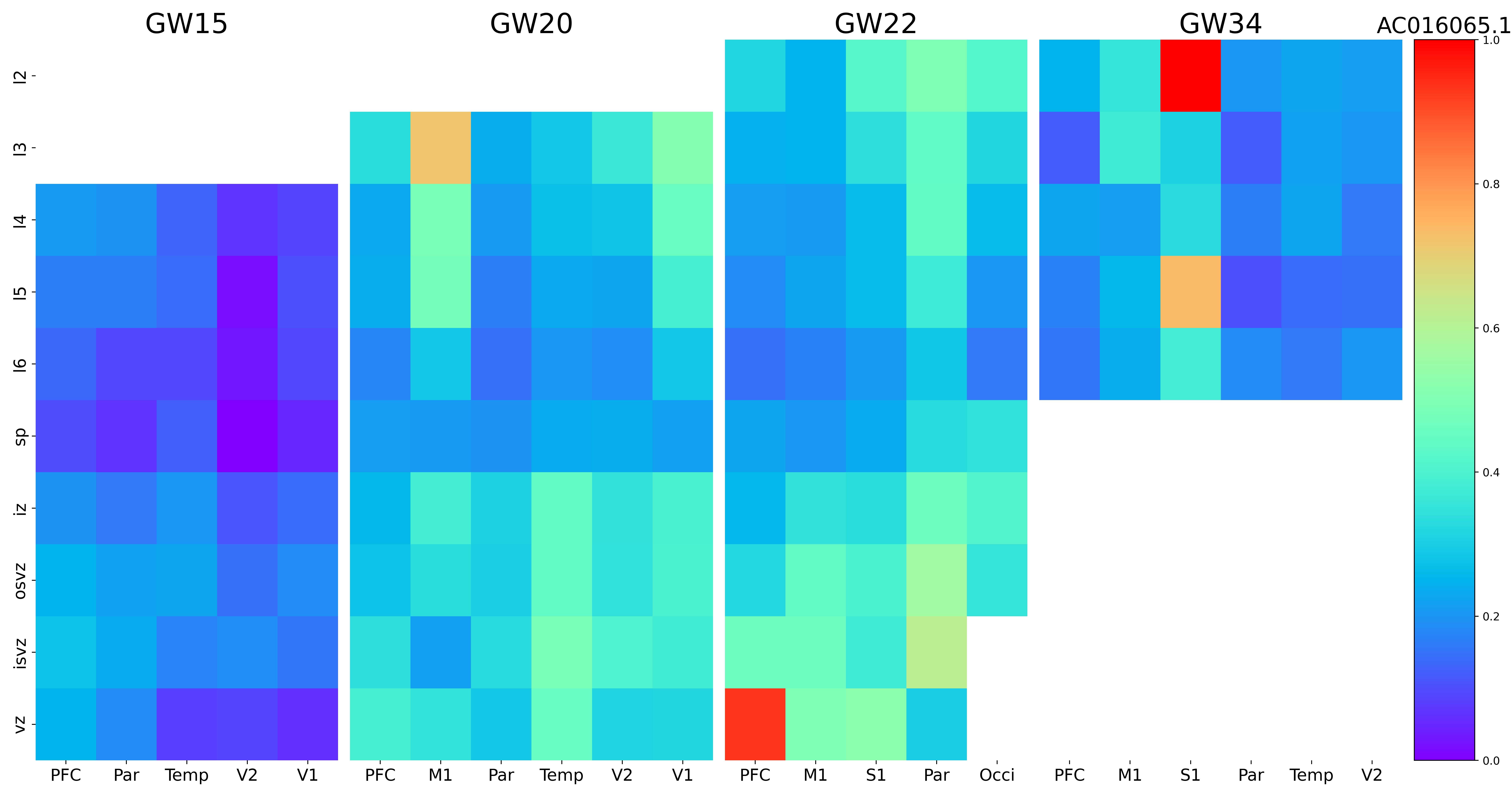

### AC093249.6.png

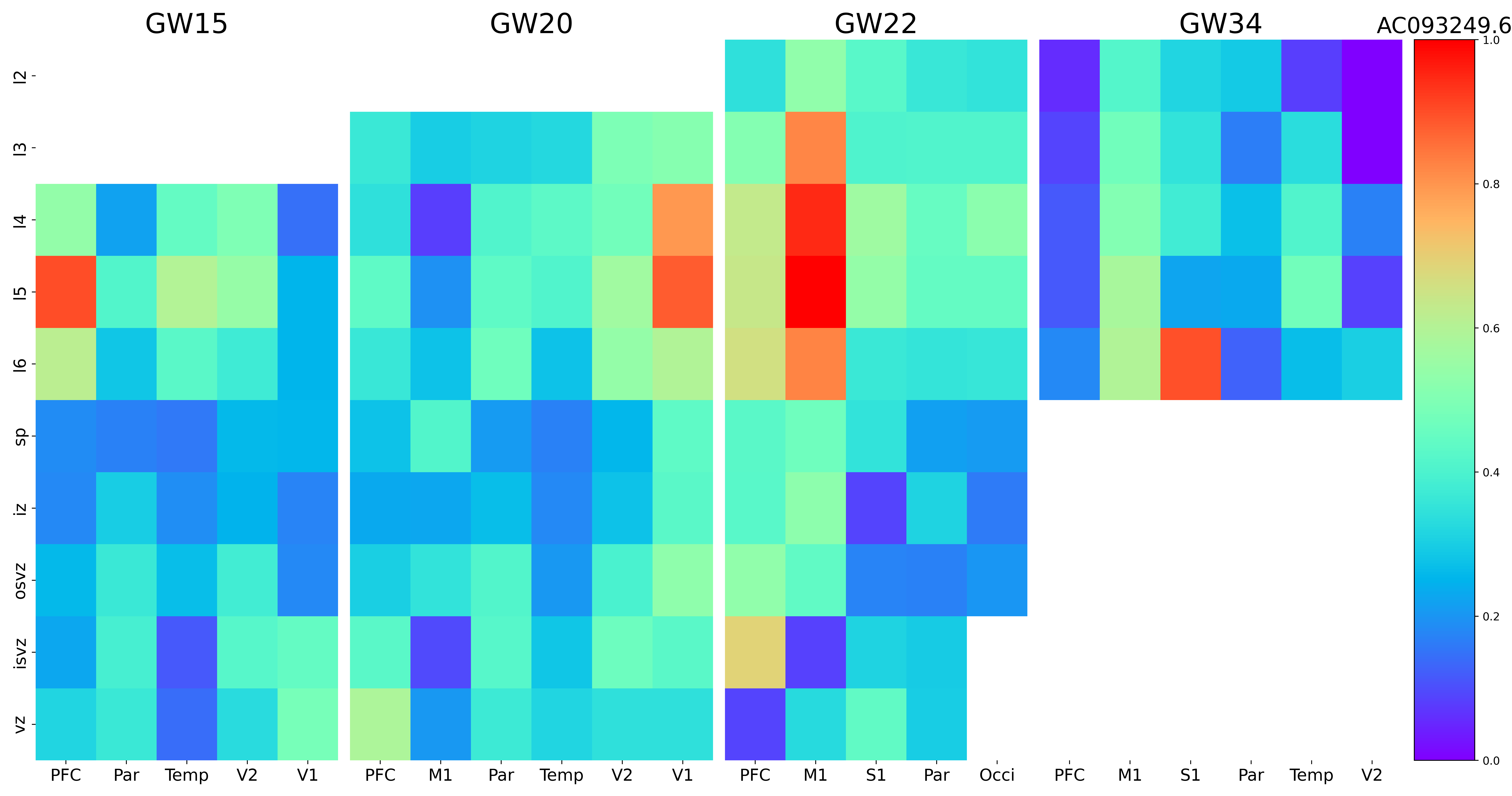

### ACSBG1.png

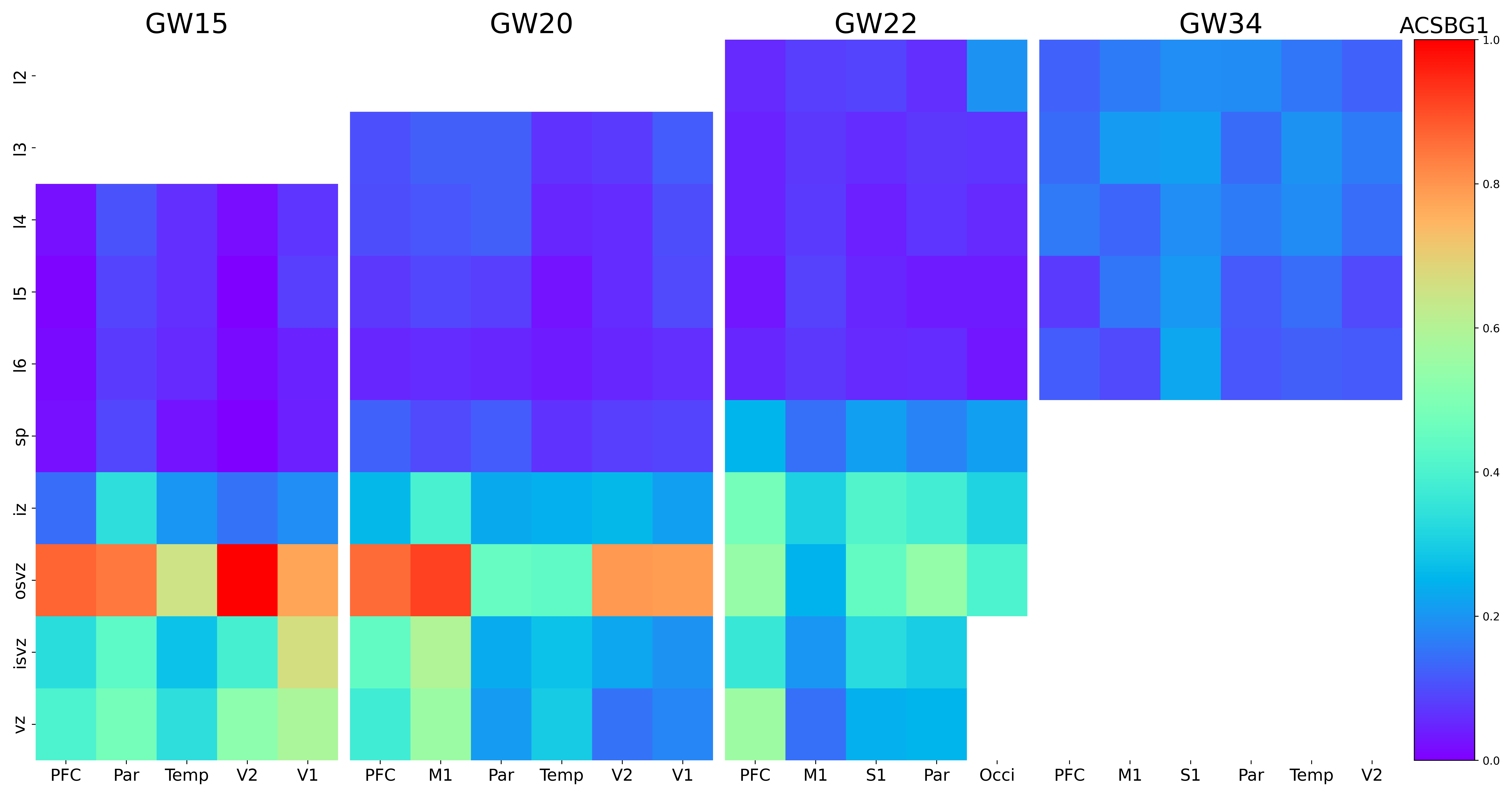

### ACTN2.png

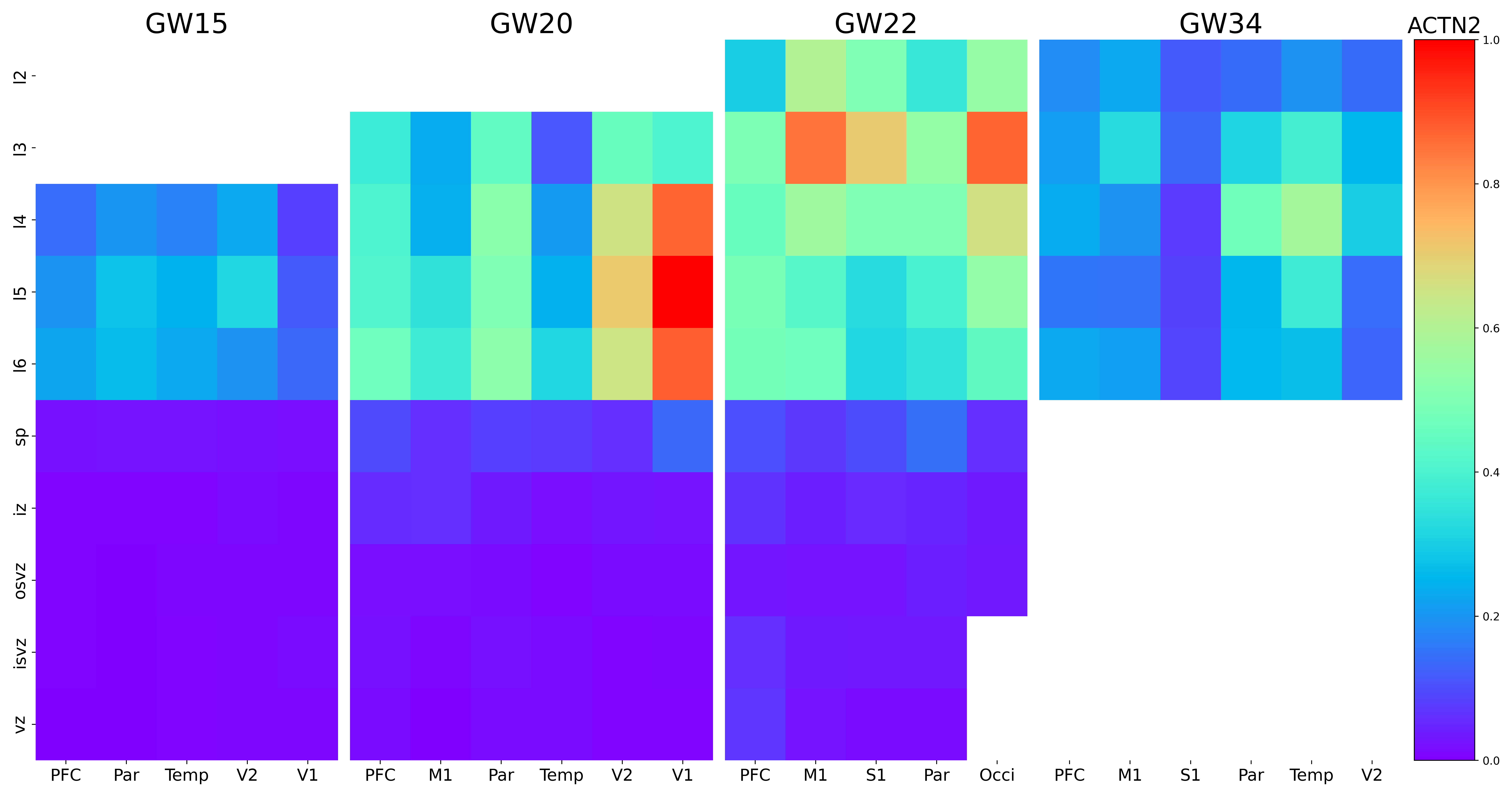

### ADAMTS5.png

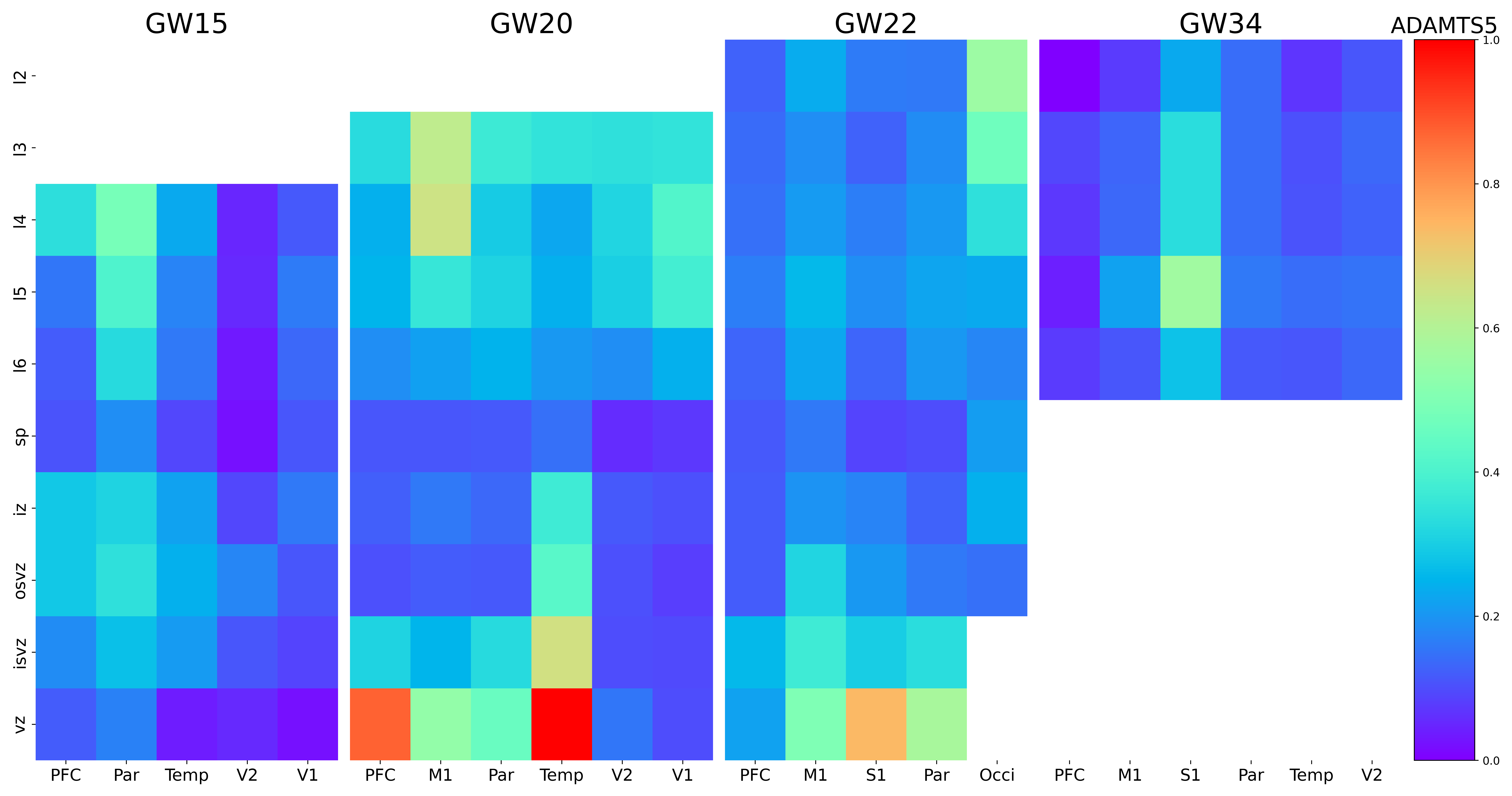

### ADAMTSL3.png

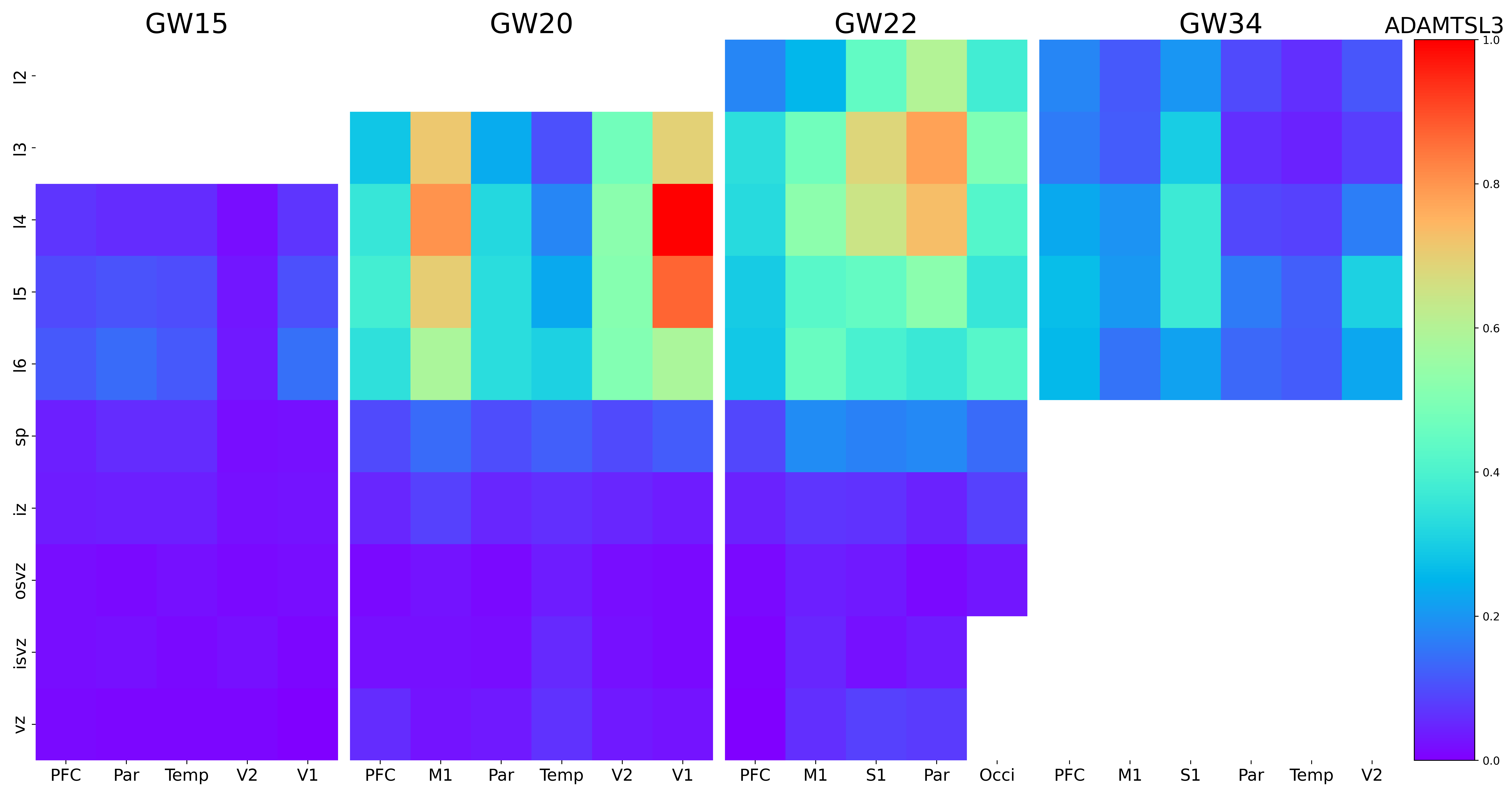

### ADRA2A.png

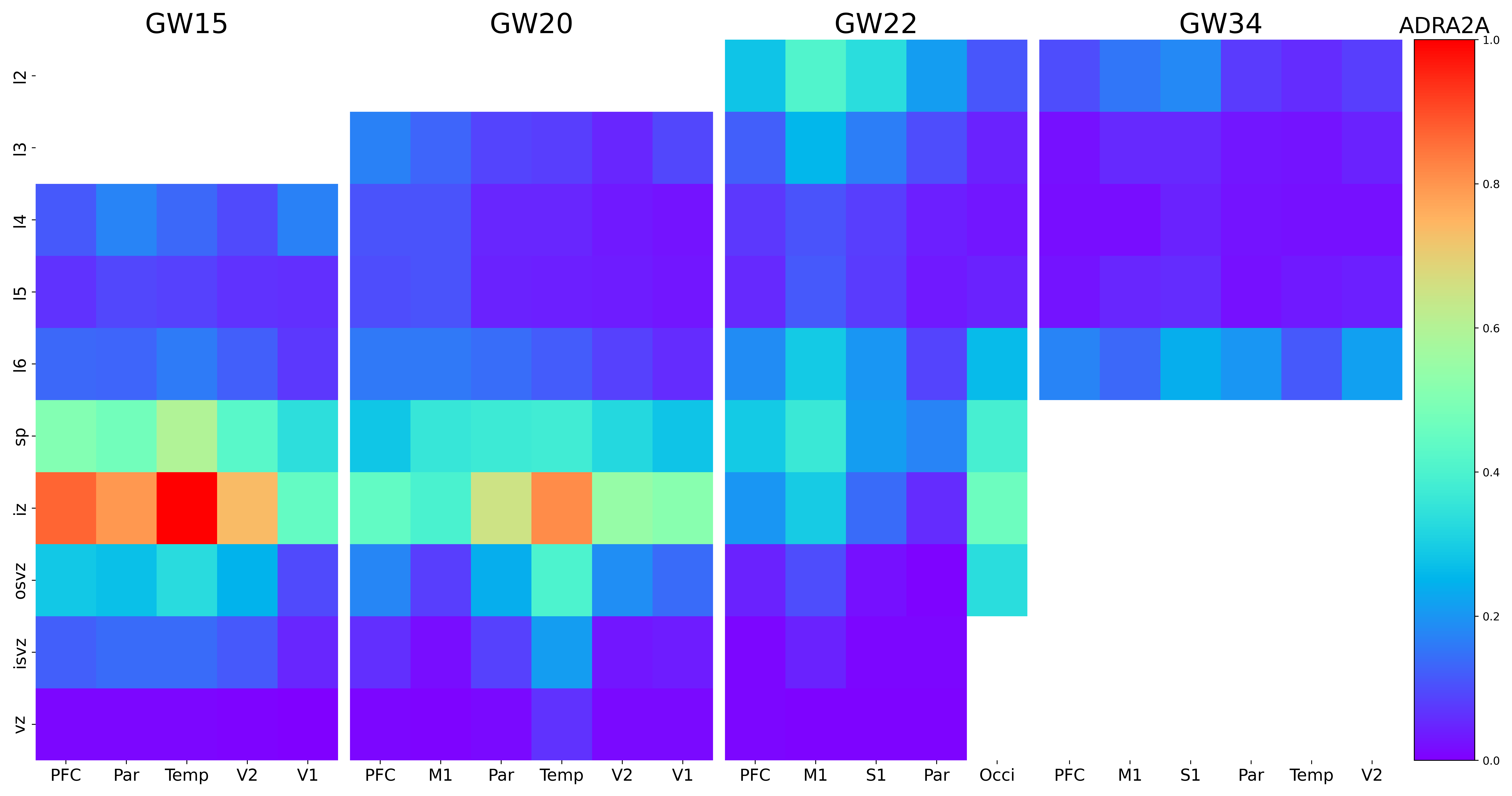

### AGT.png

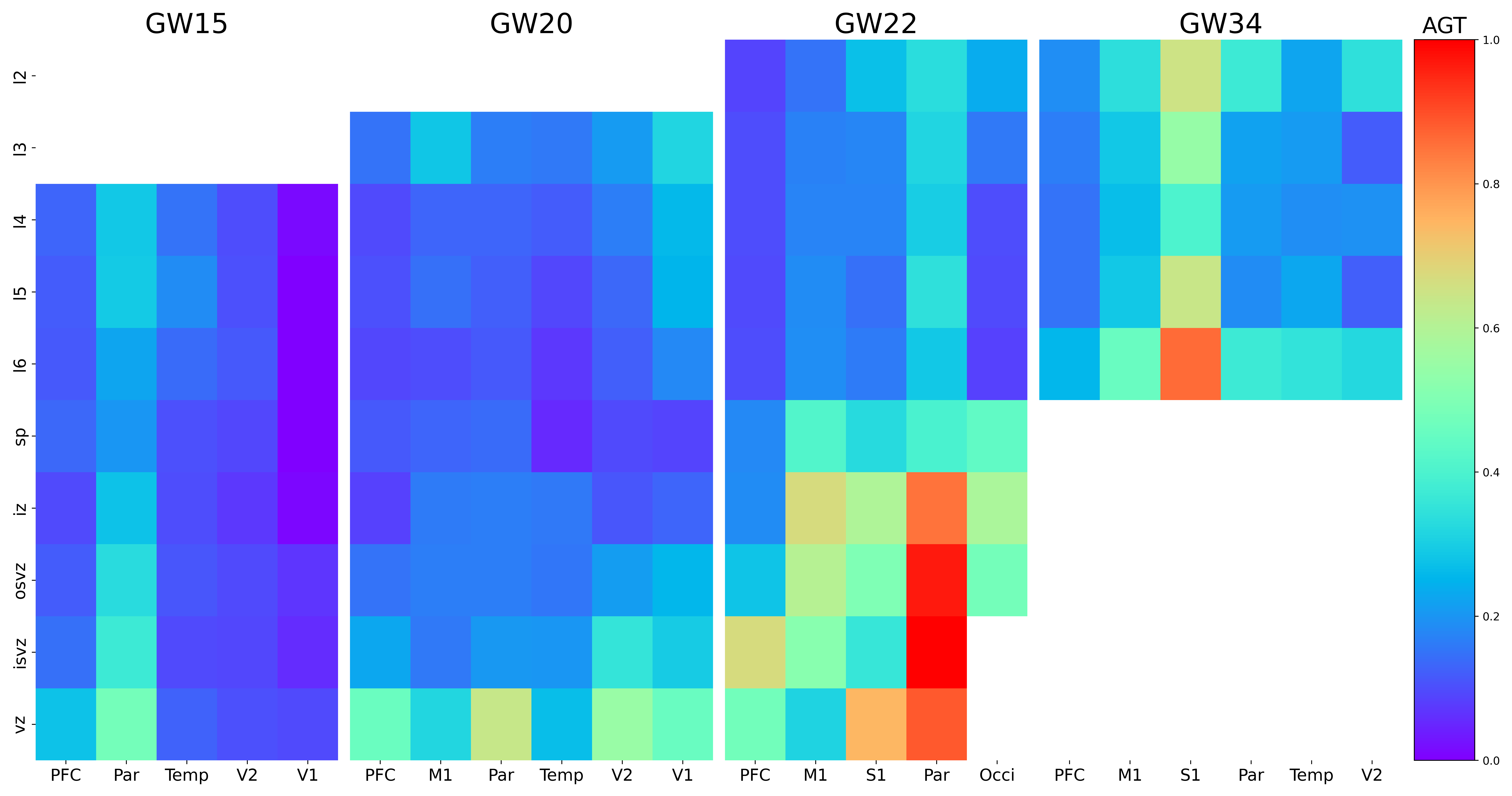

### AHI1.png

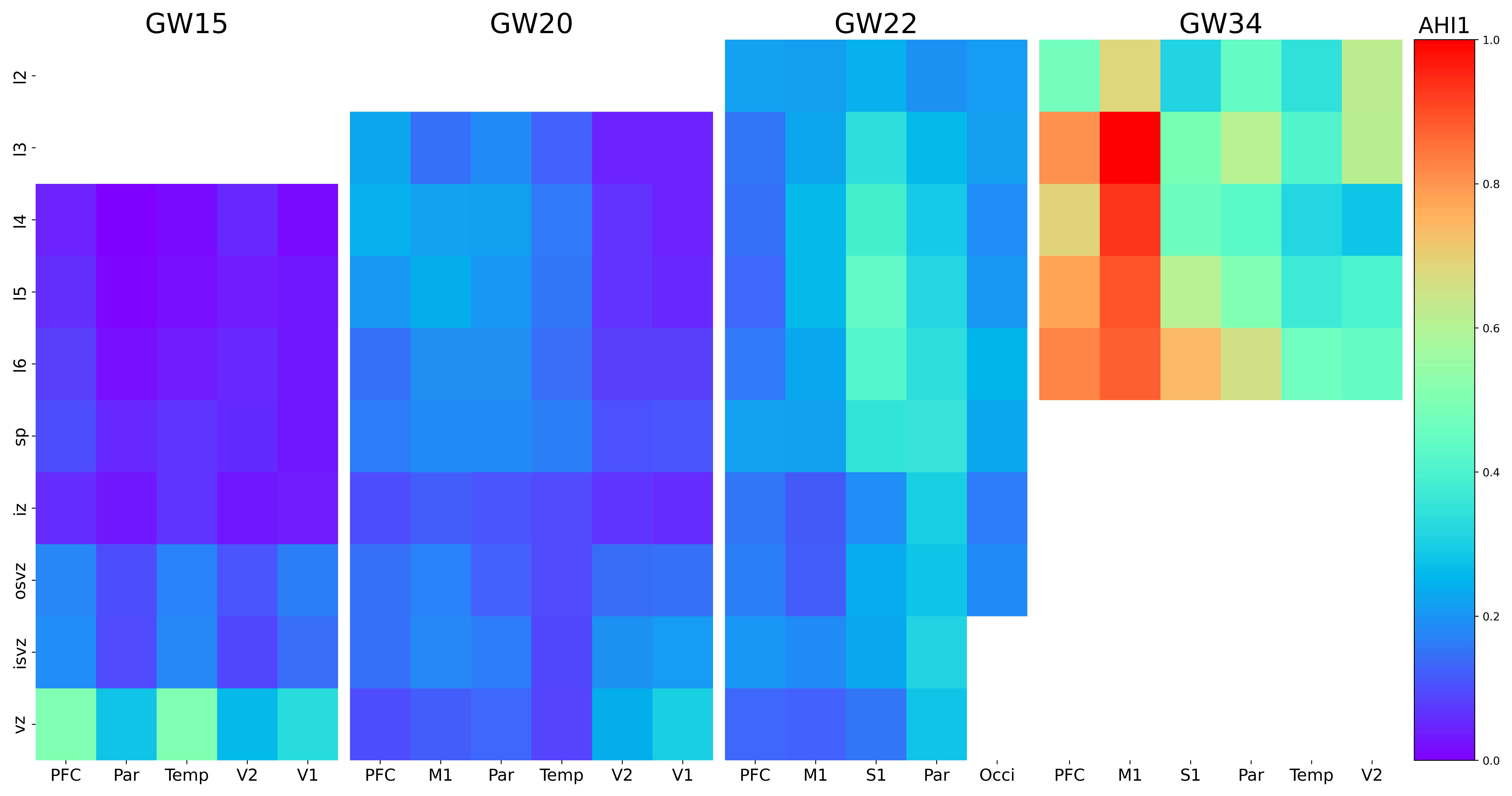

### AKAP7.png

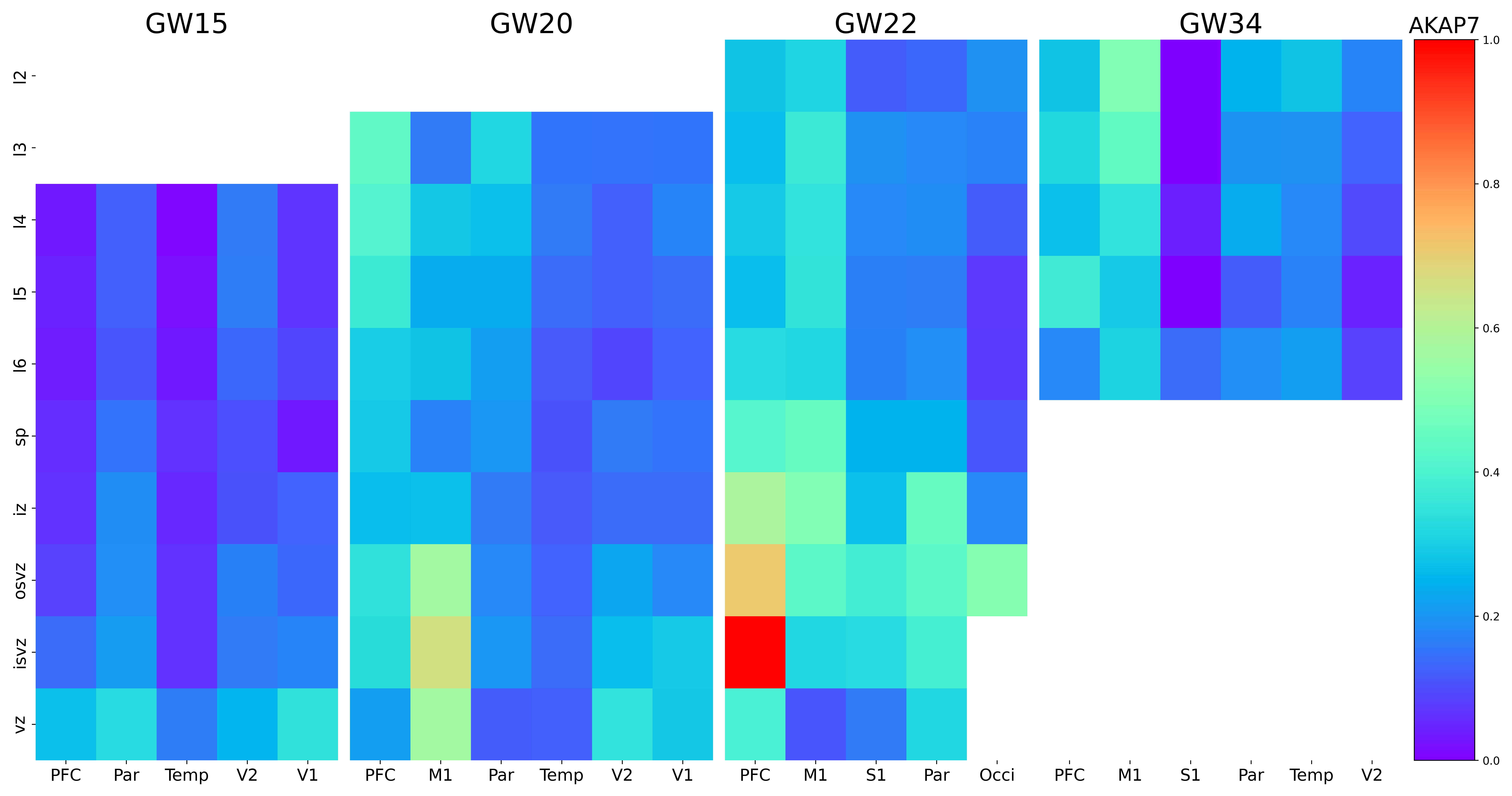

### AKR1C2.png

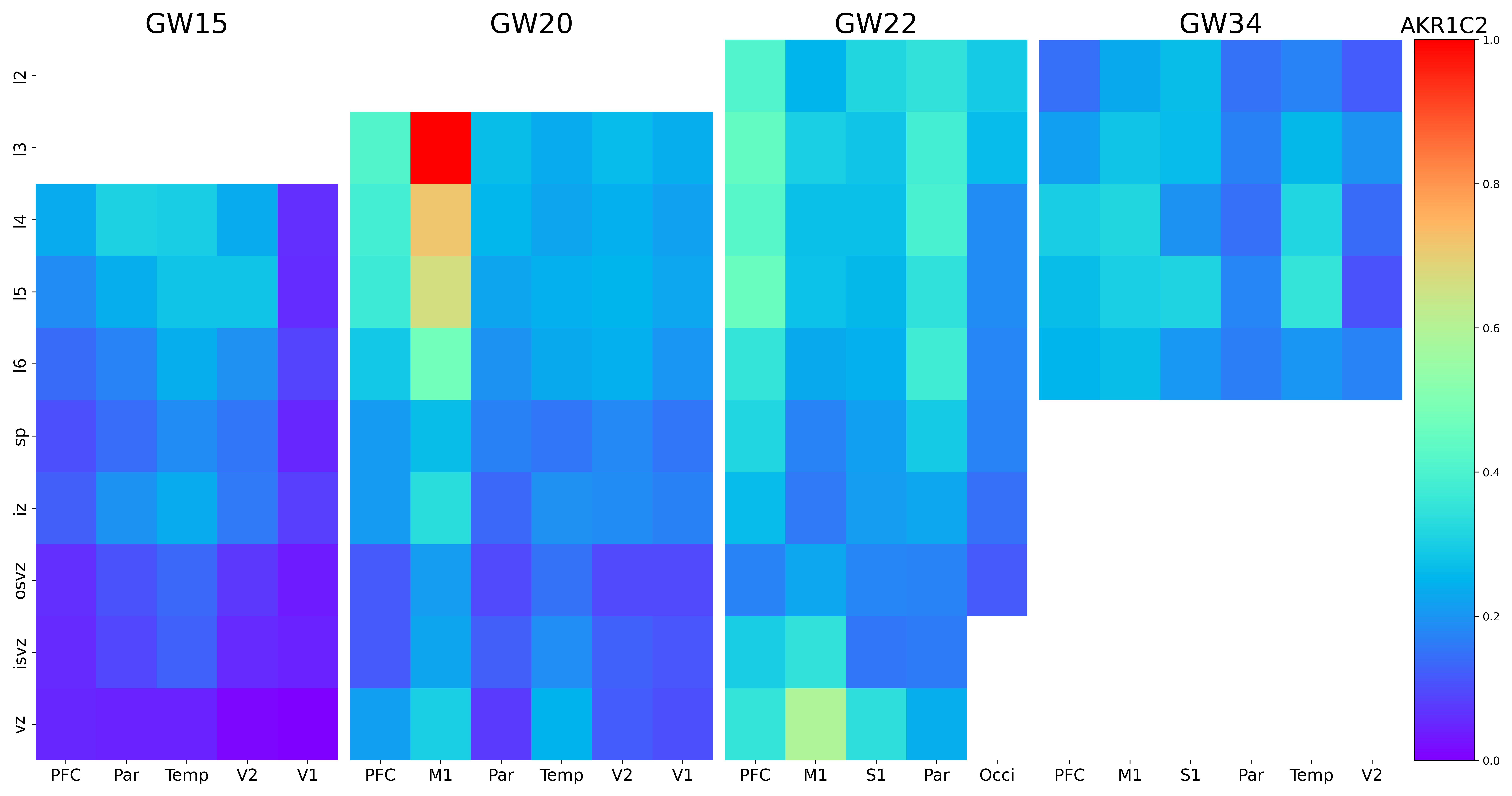

### ALDOC.png

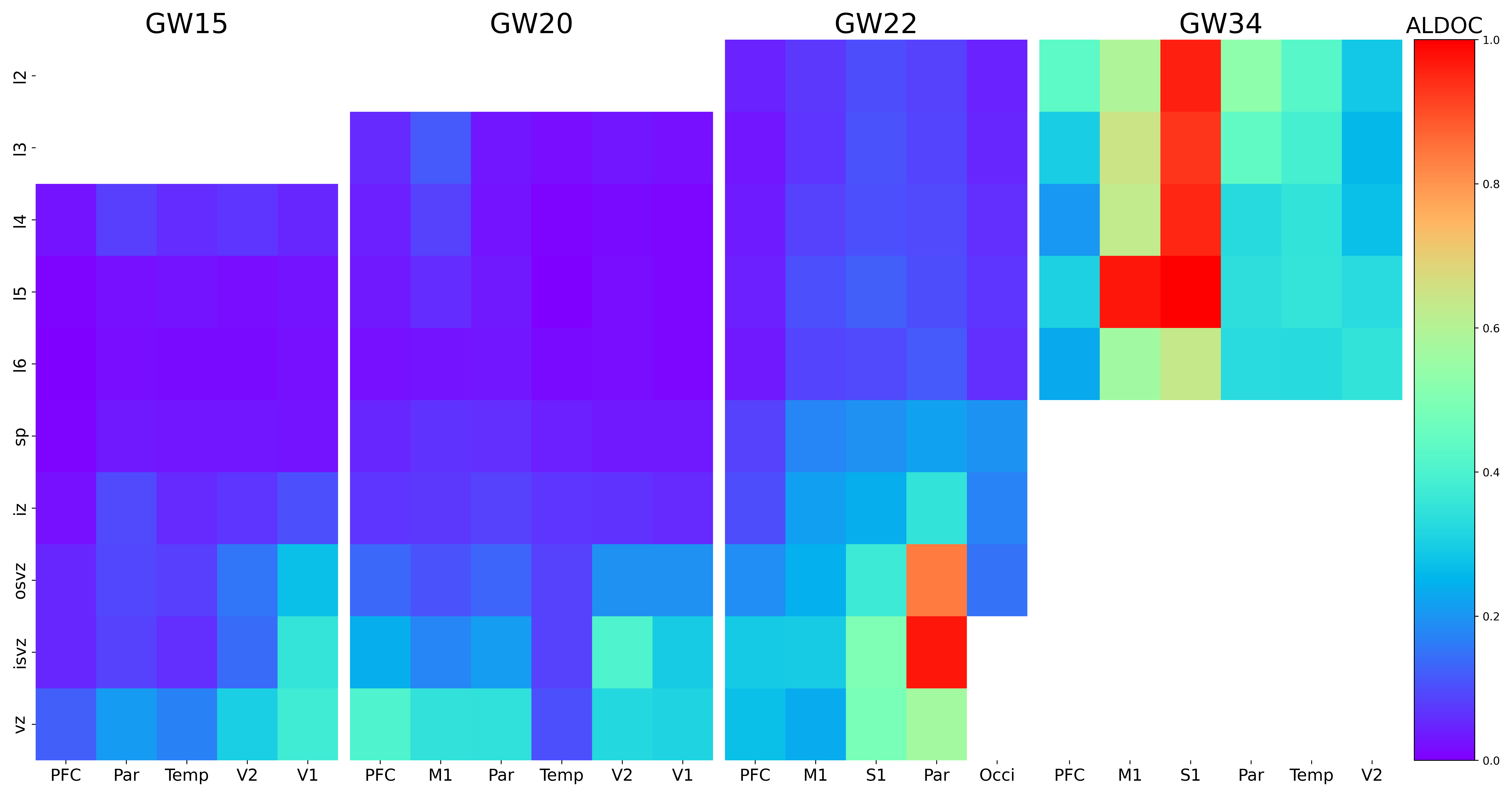

### ALPK1.png

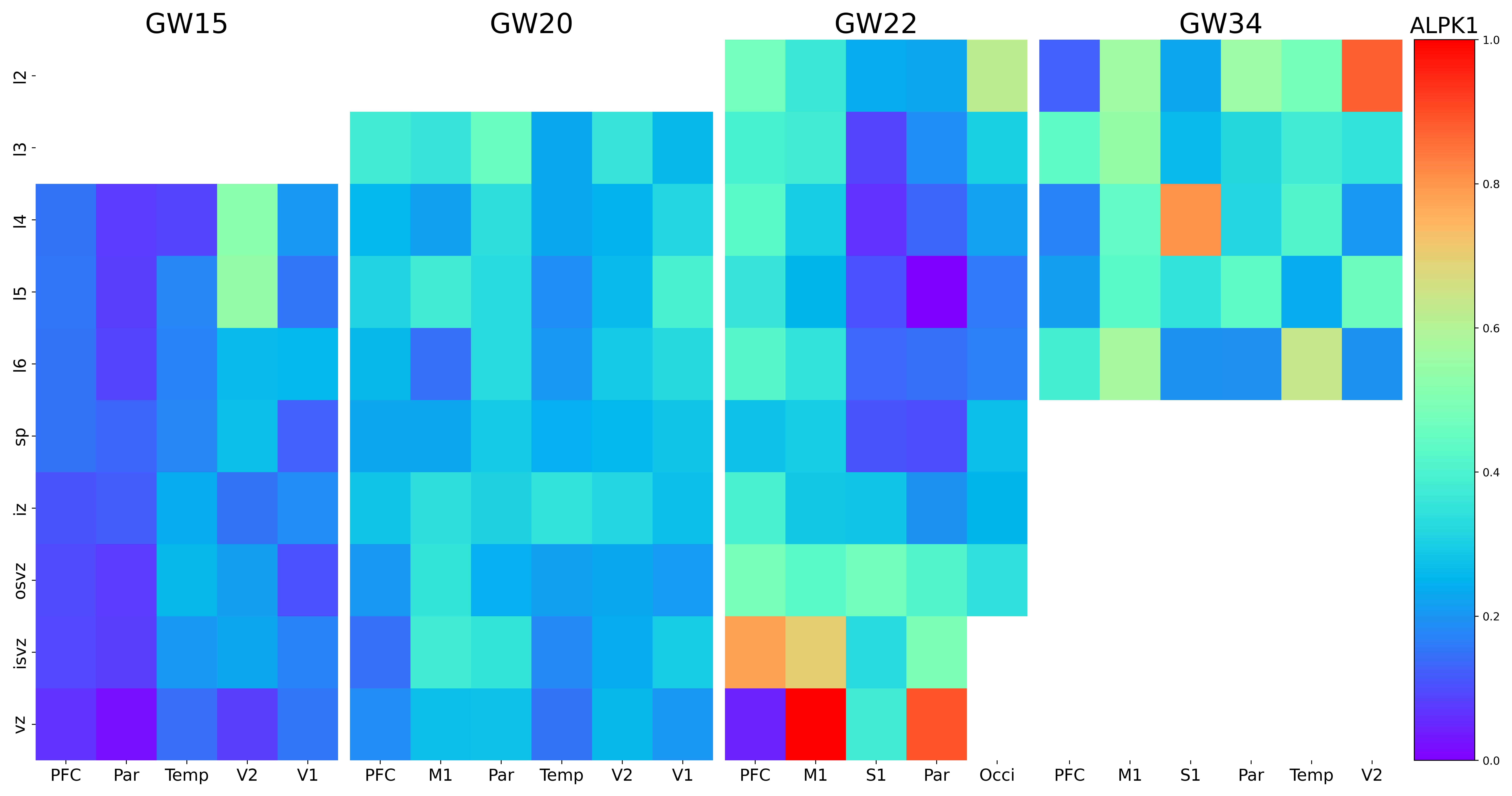

### ANGPT2.png

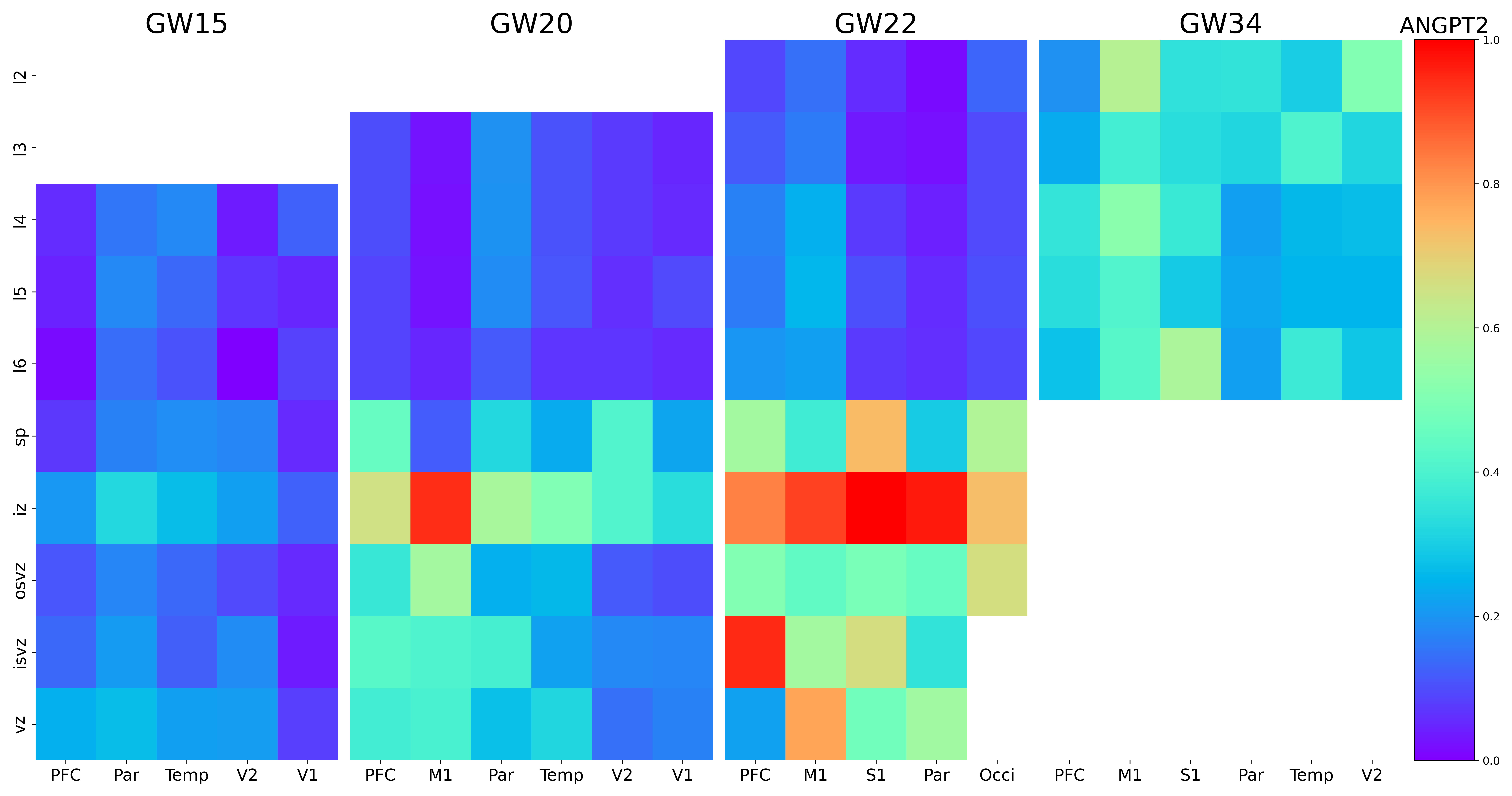

### ANGPTL1.png

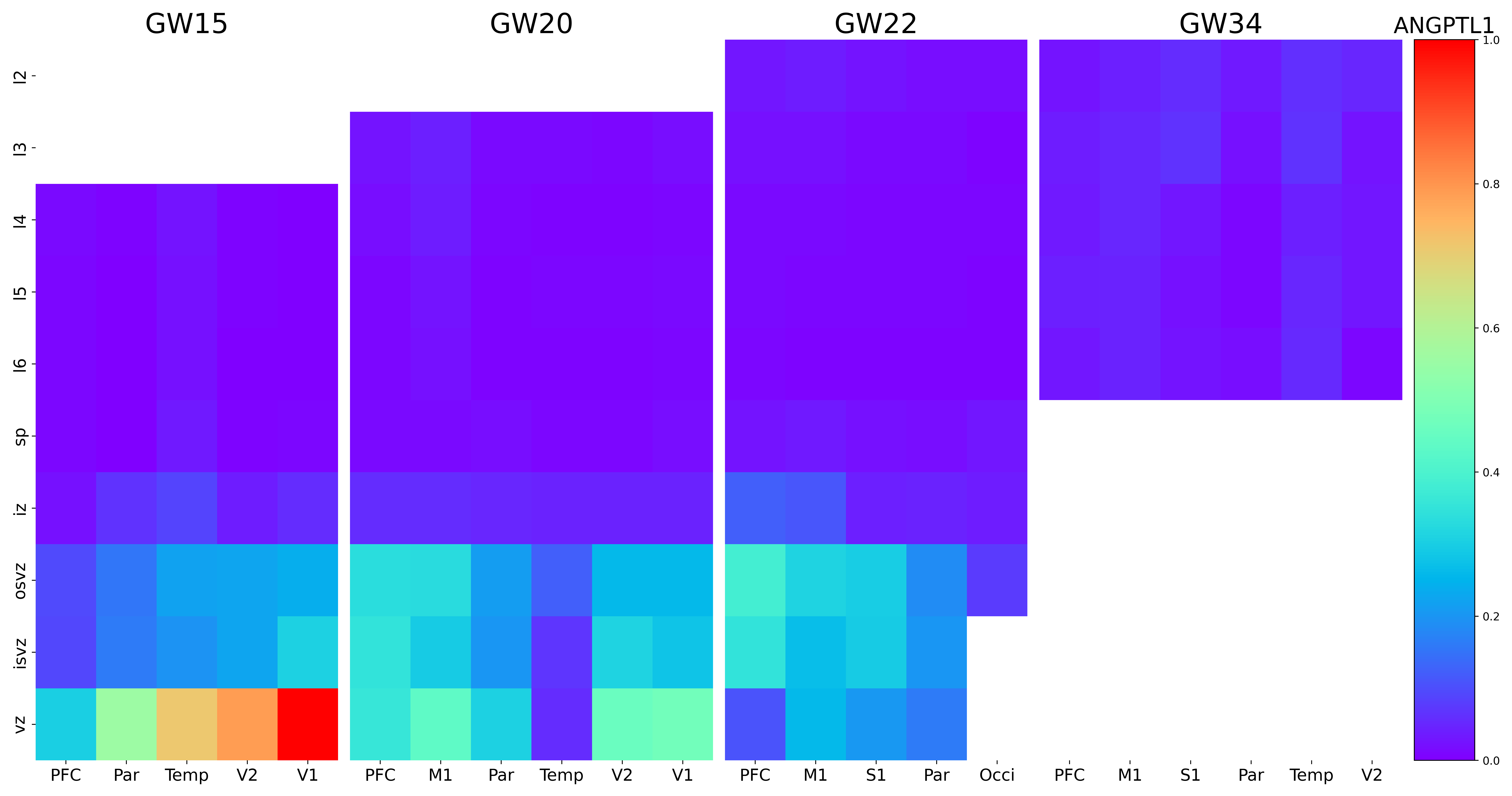

### ANKFN1.png

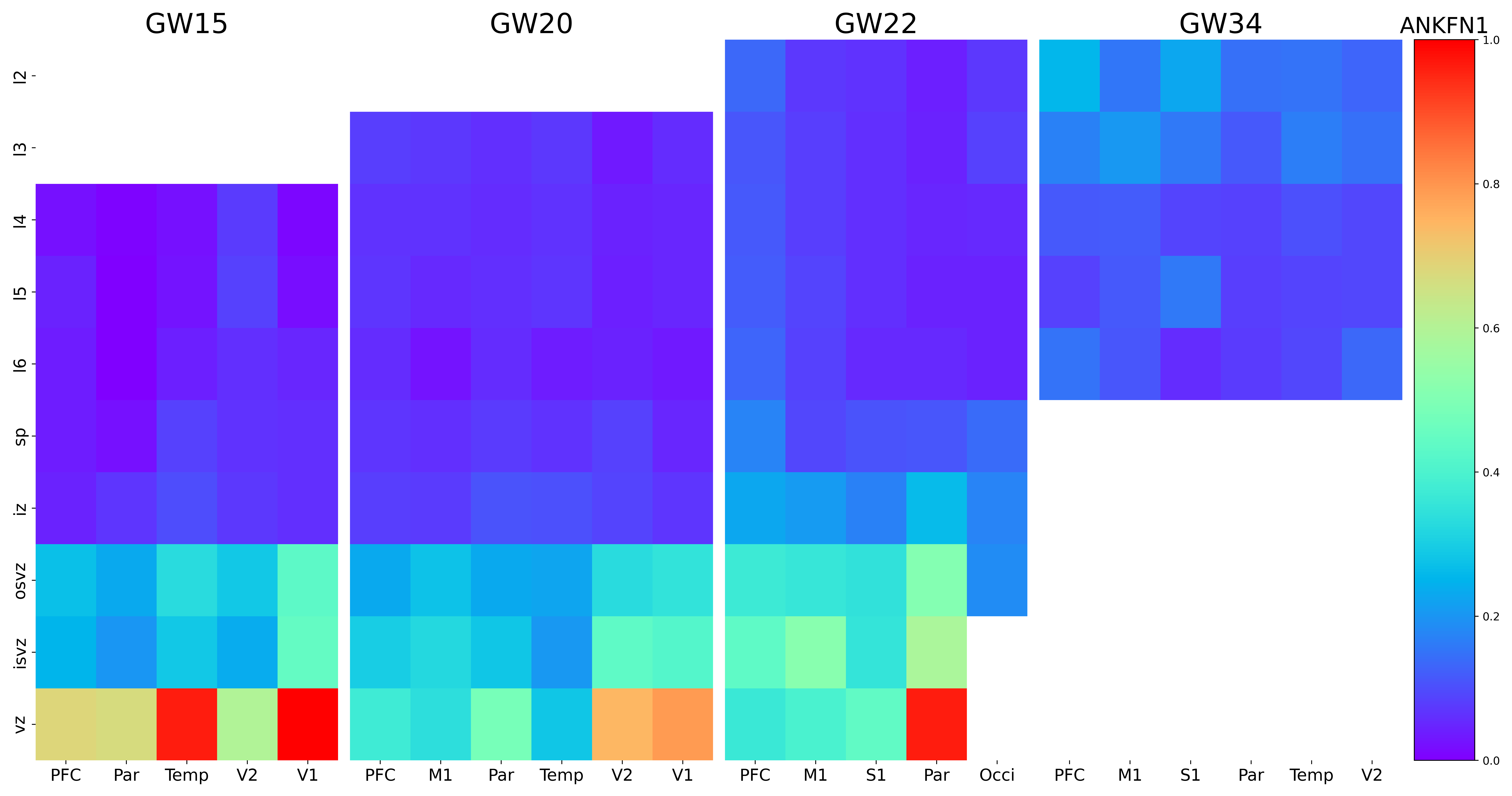

### ANOS1.png

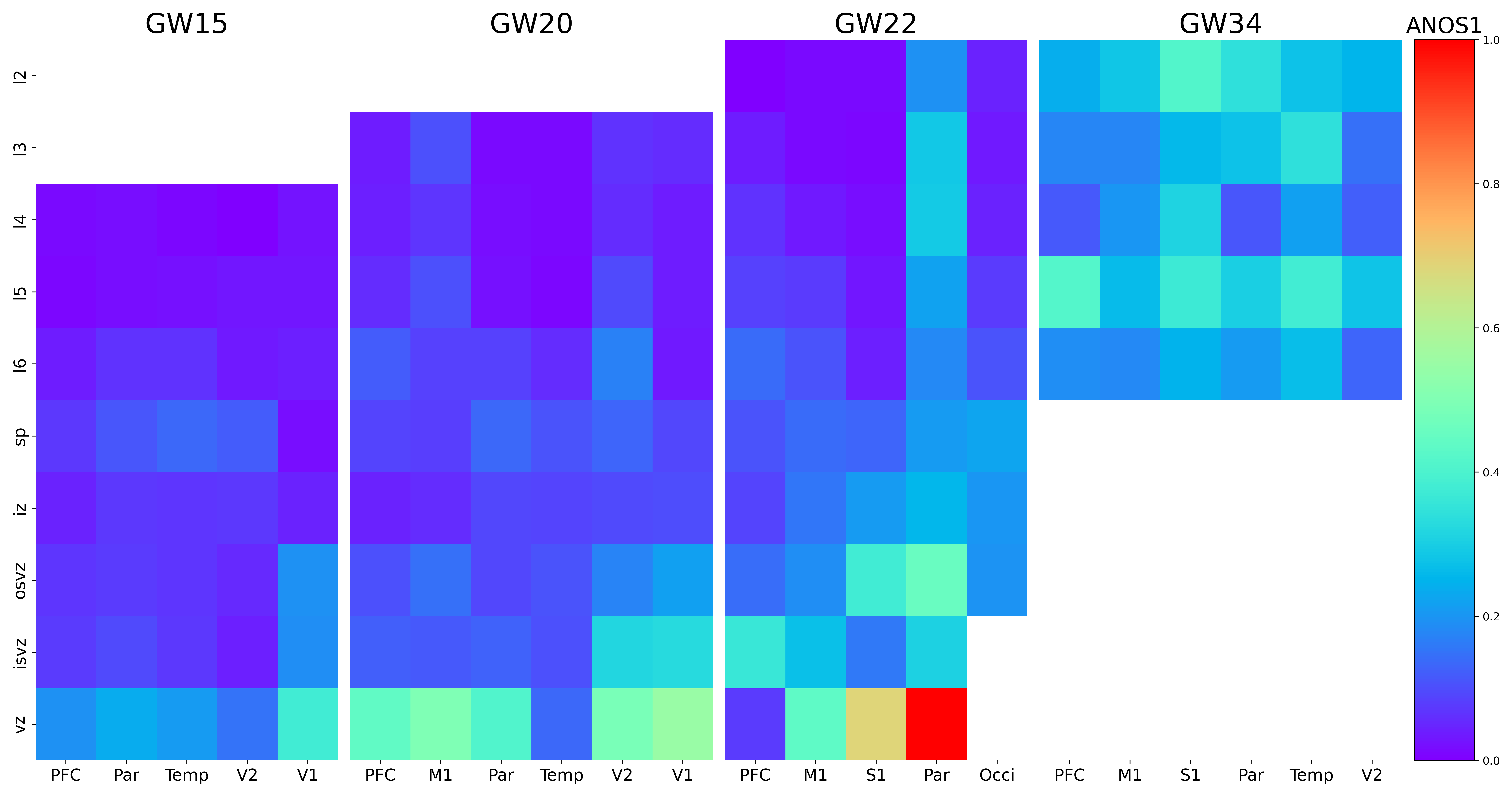

### ANTXR2.png

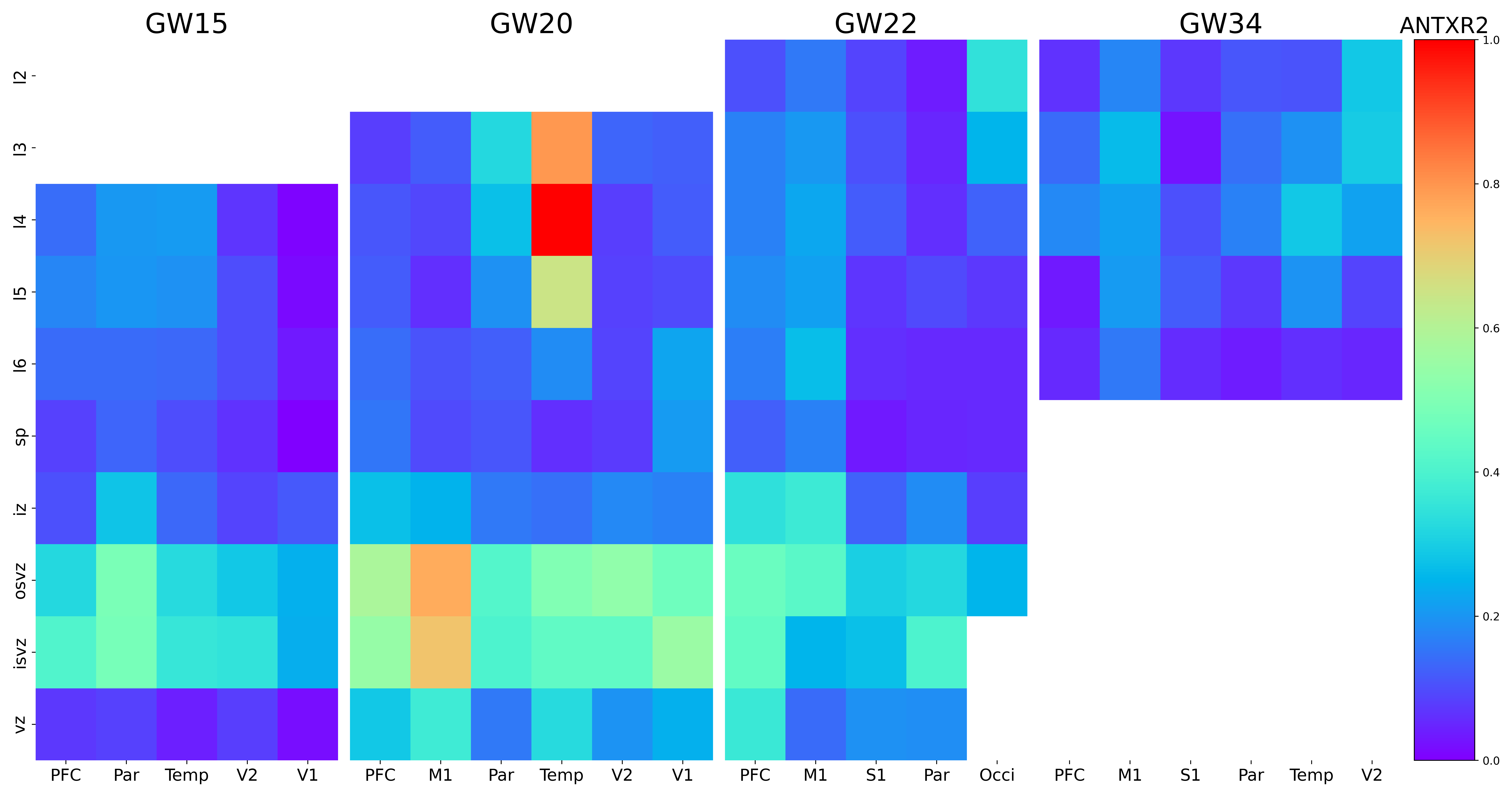

### AP000866.1.png

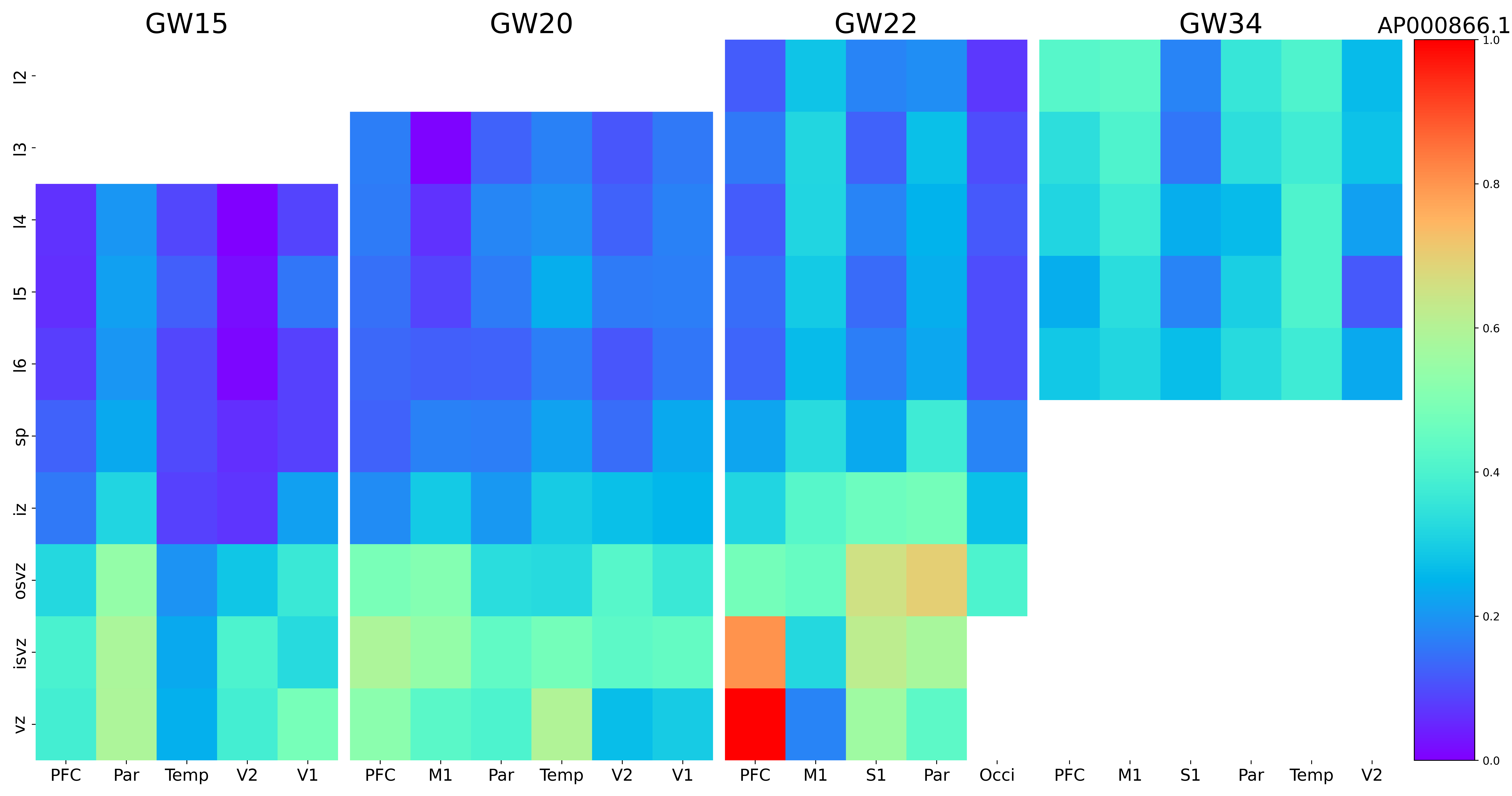

### AP002748.4.png

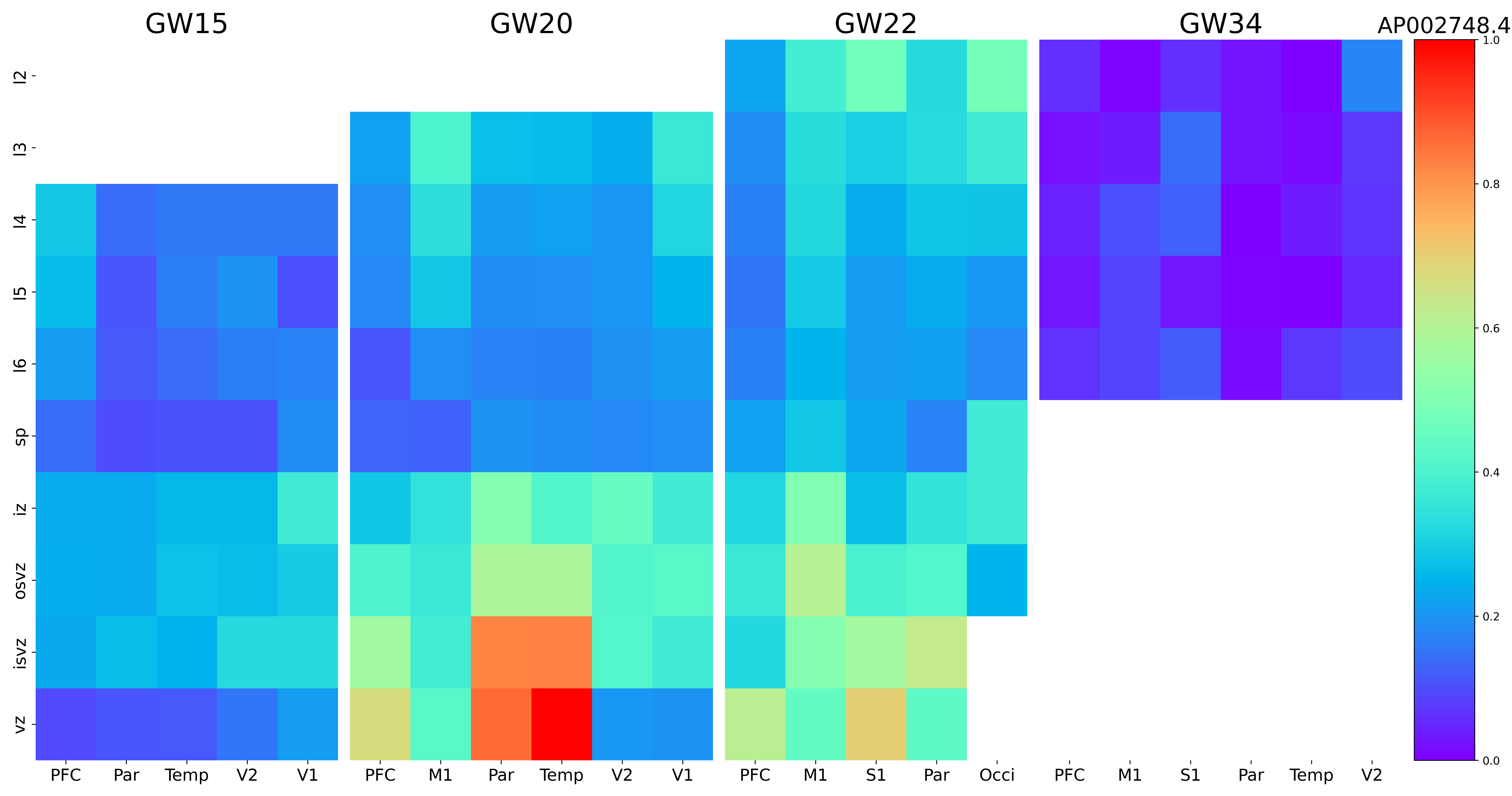

### AQP4.png

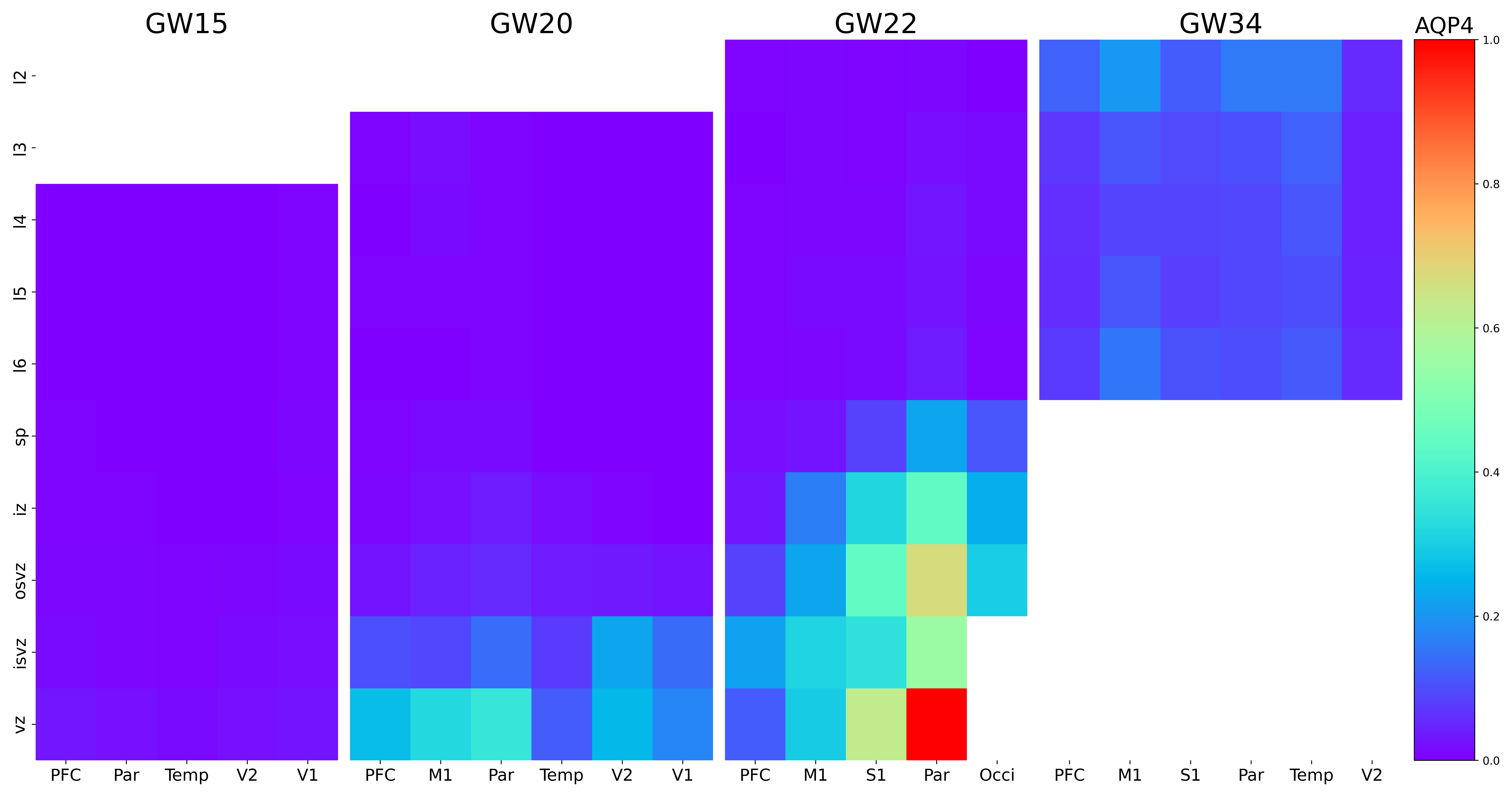

### ARFGEF3.png

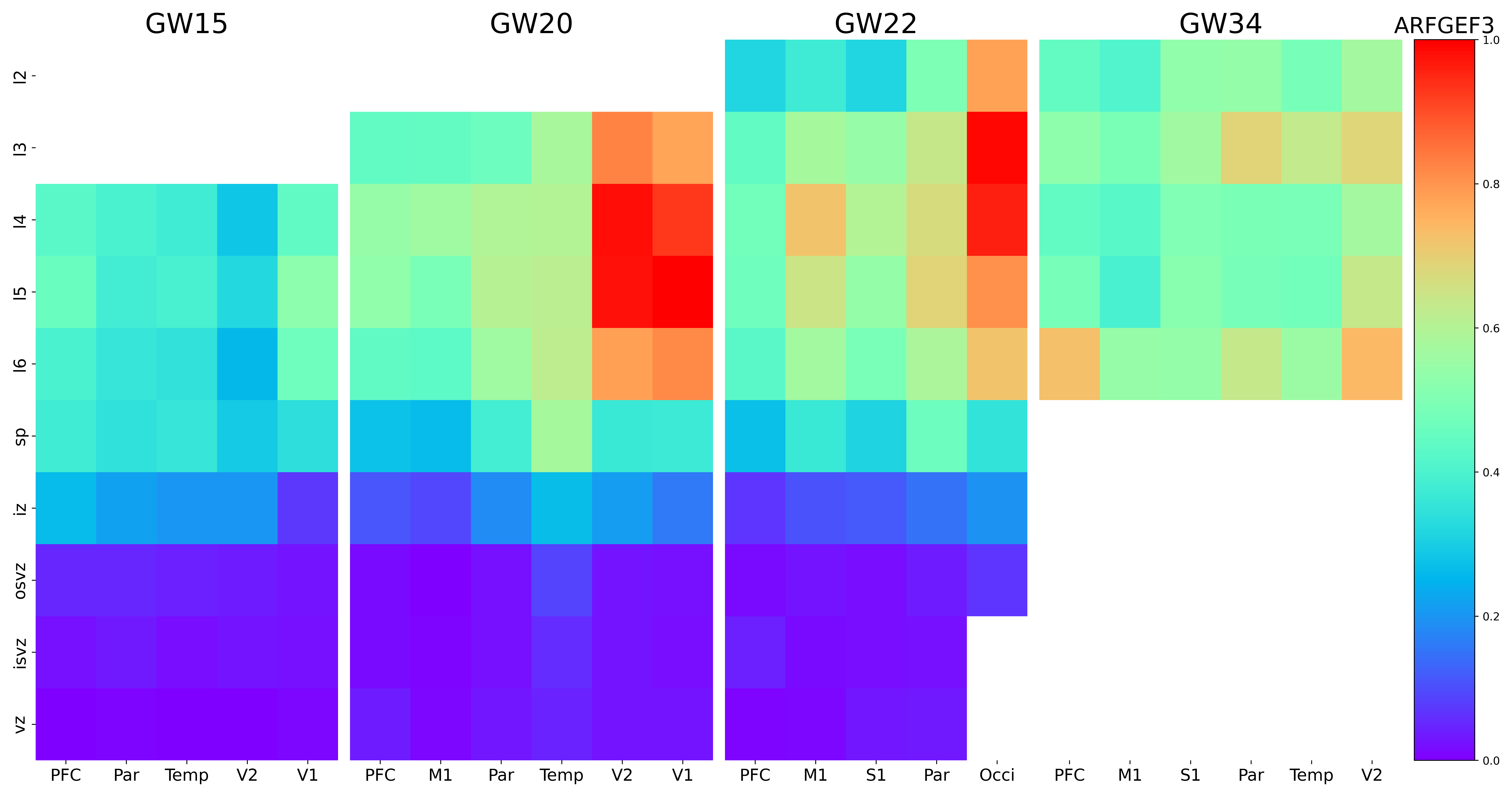

### ARHGAP5-AS1.png

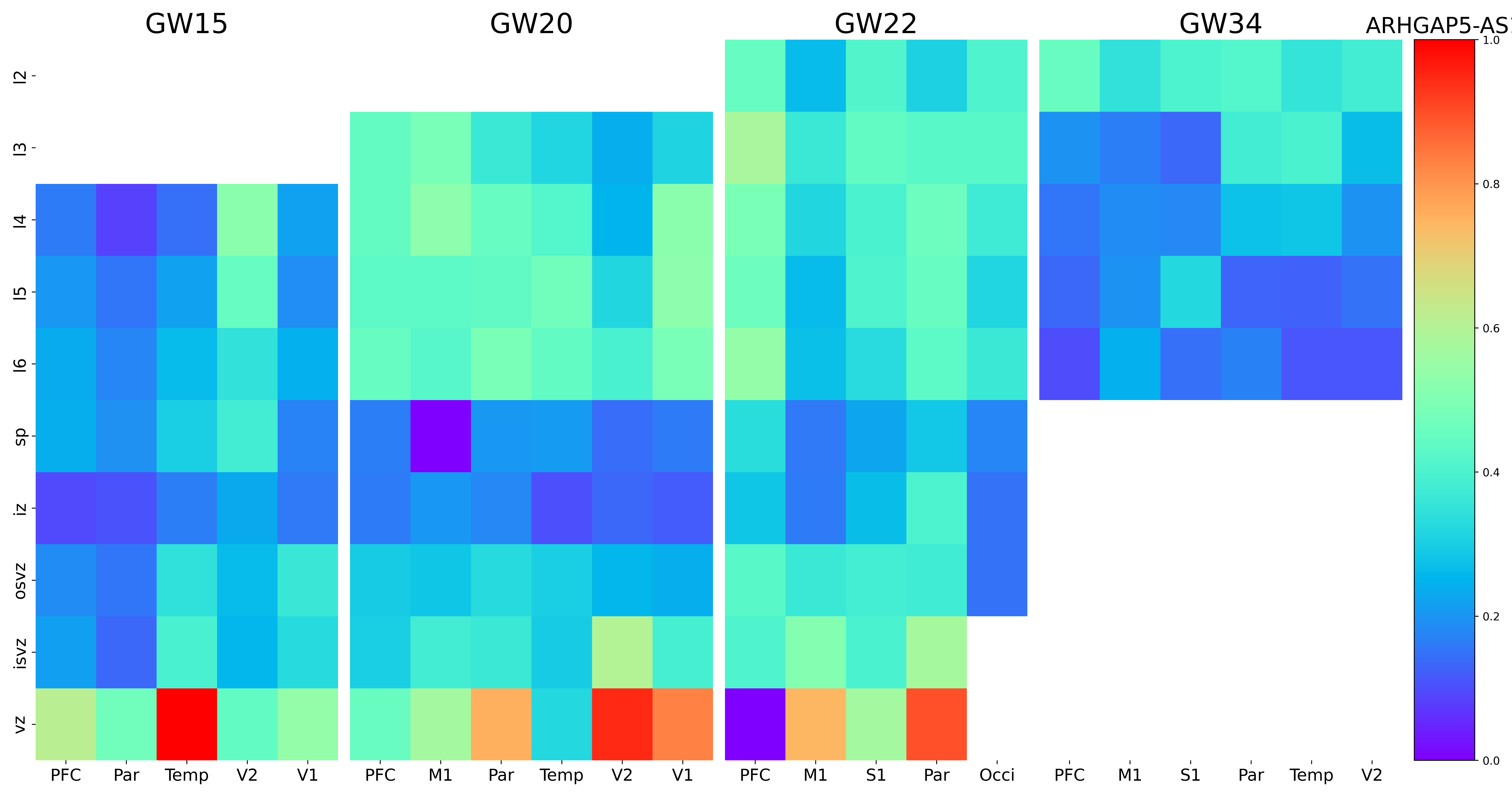

### ARHGAP5.png

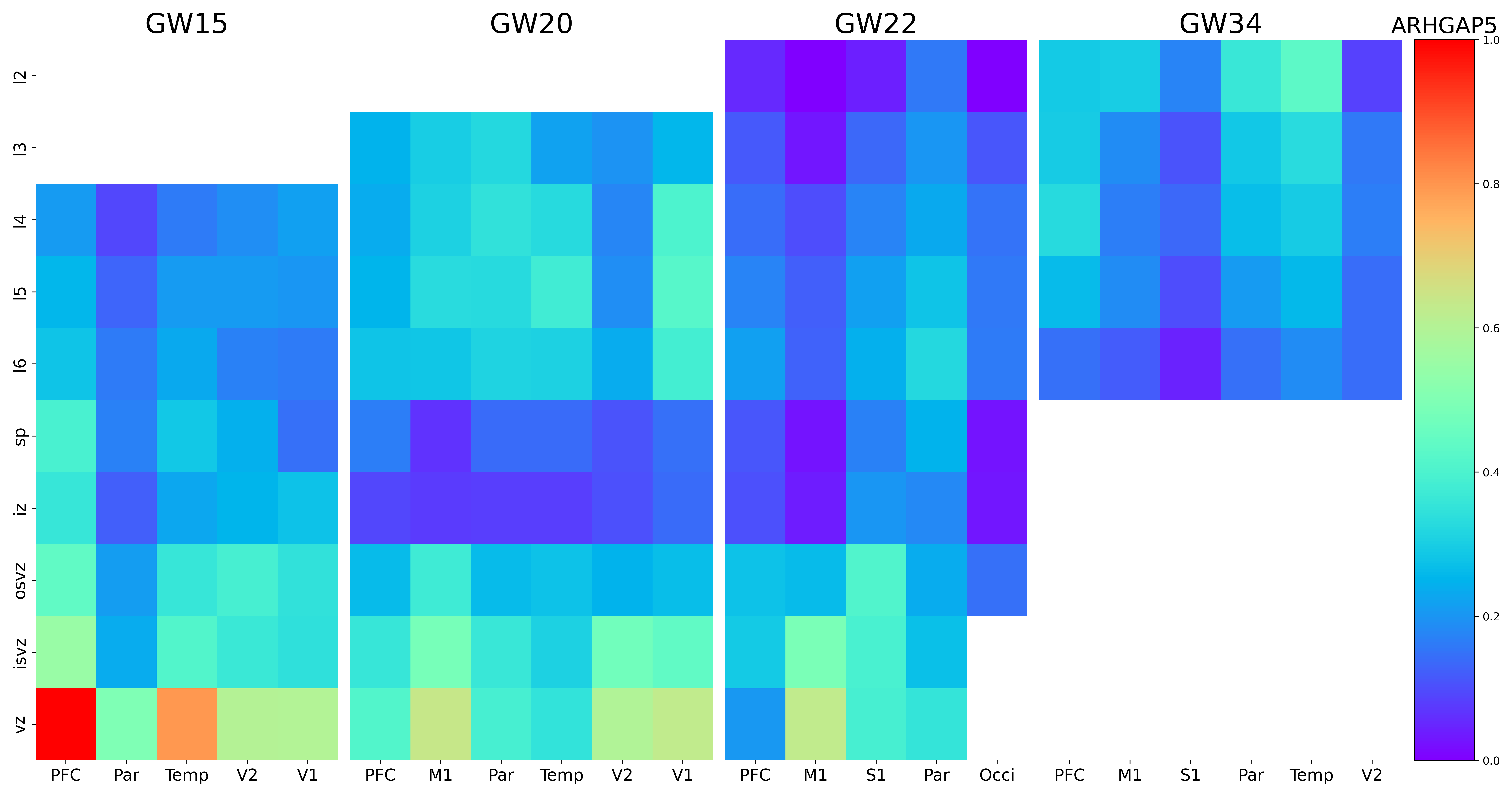

### ARID2.png

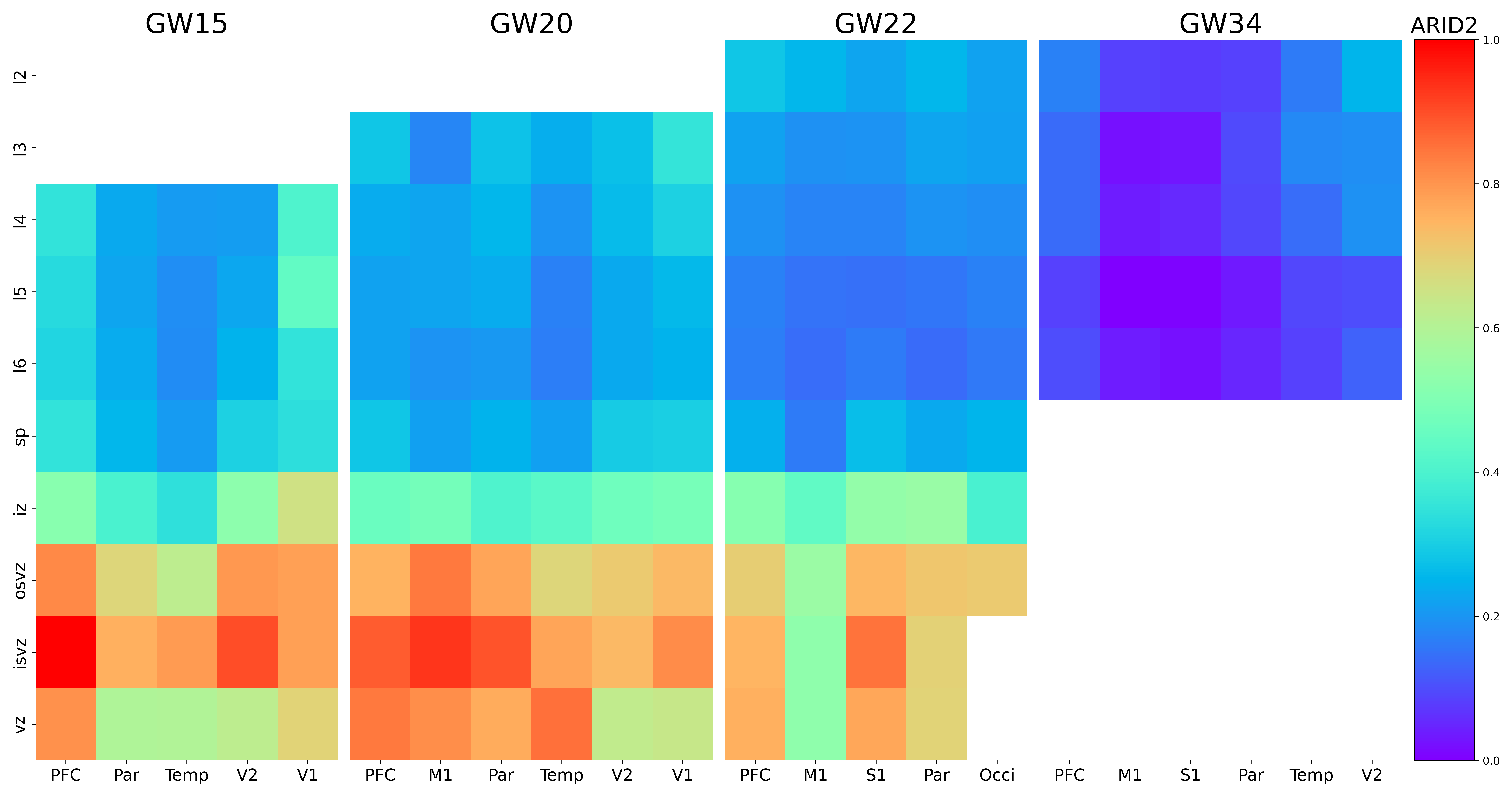

### ARRDC1-AS1.png

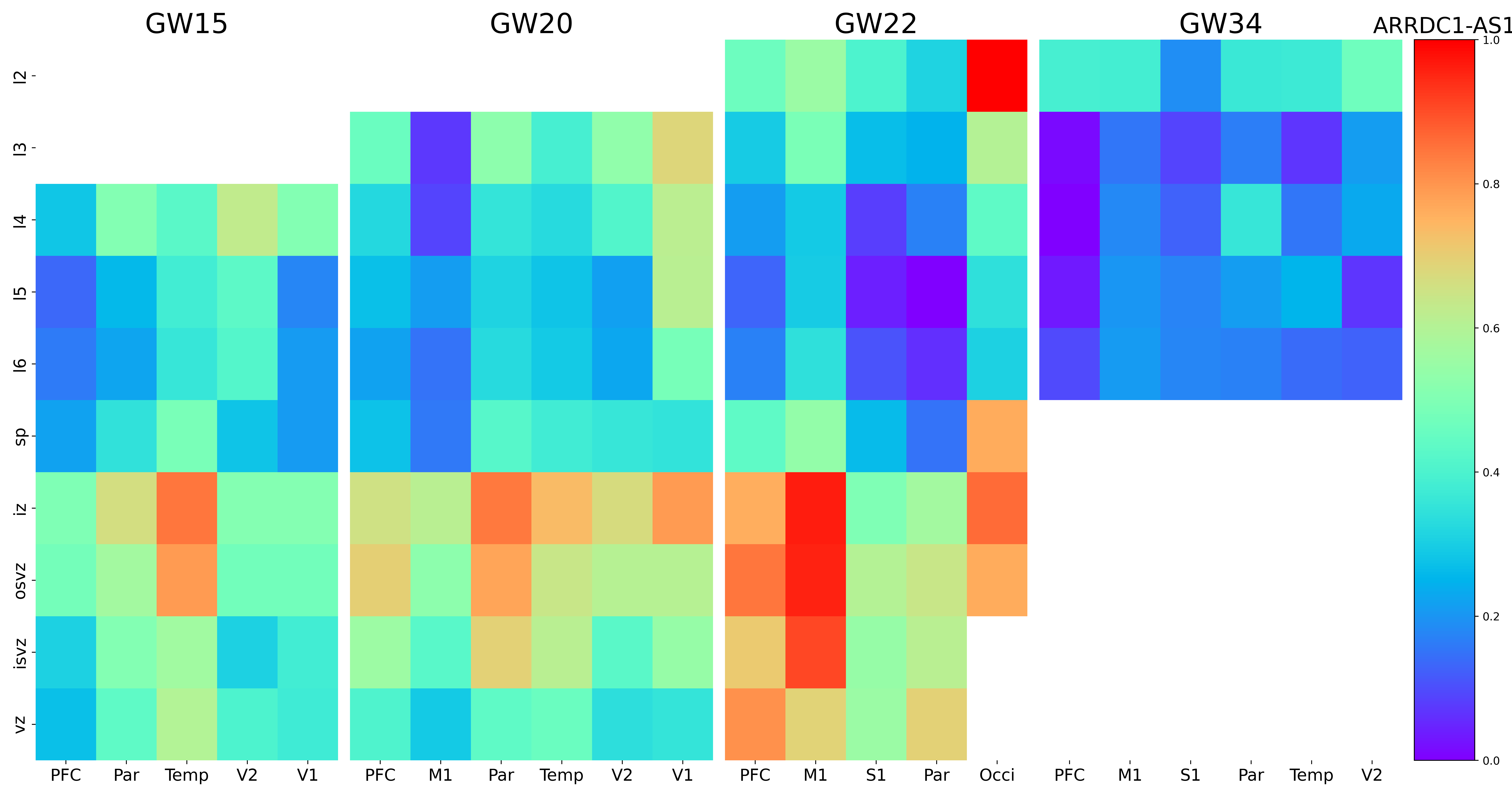

### ASH1L-AS1.png

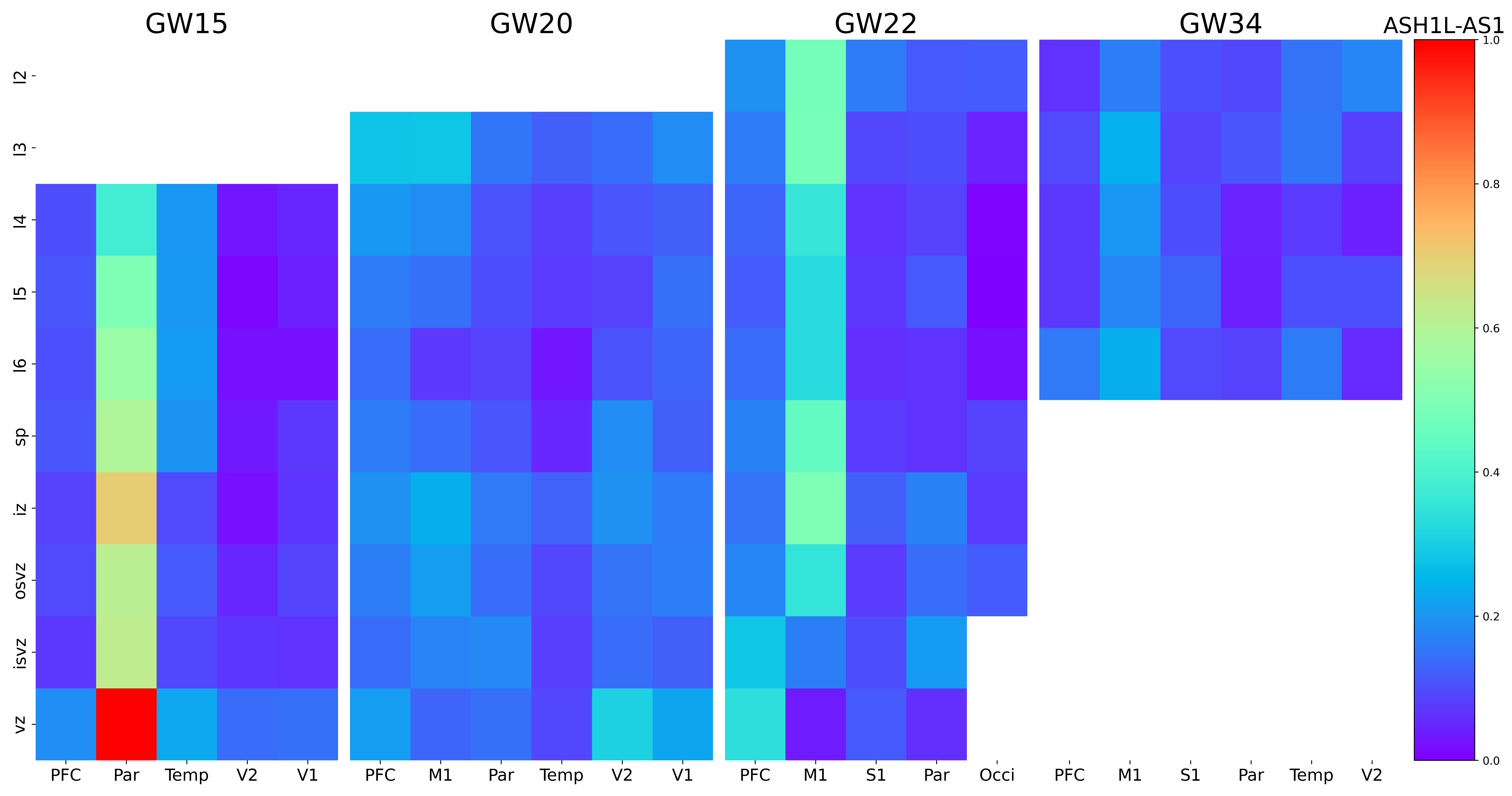

### ASH1L.png

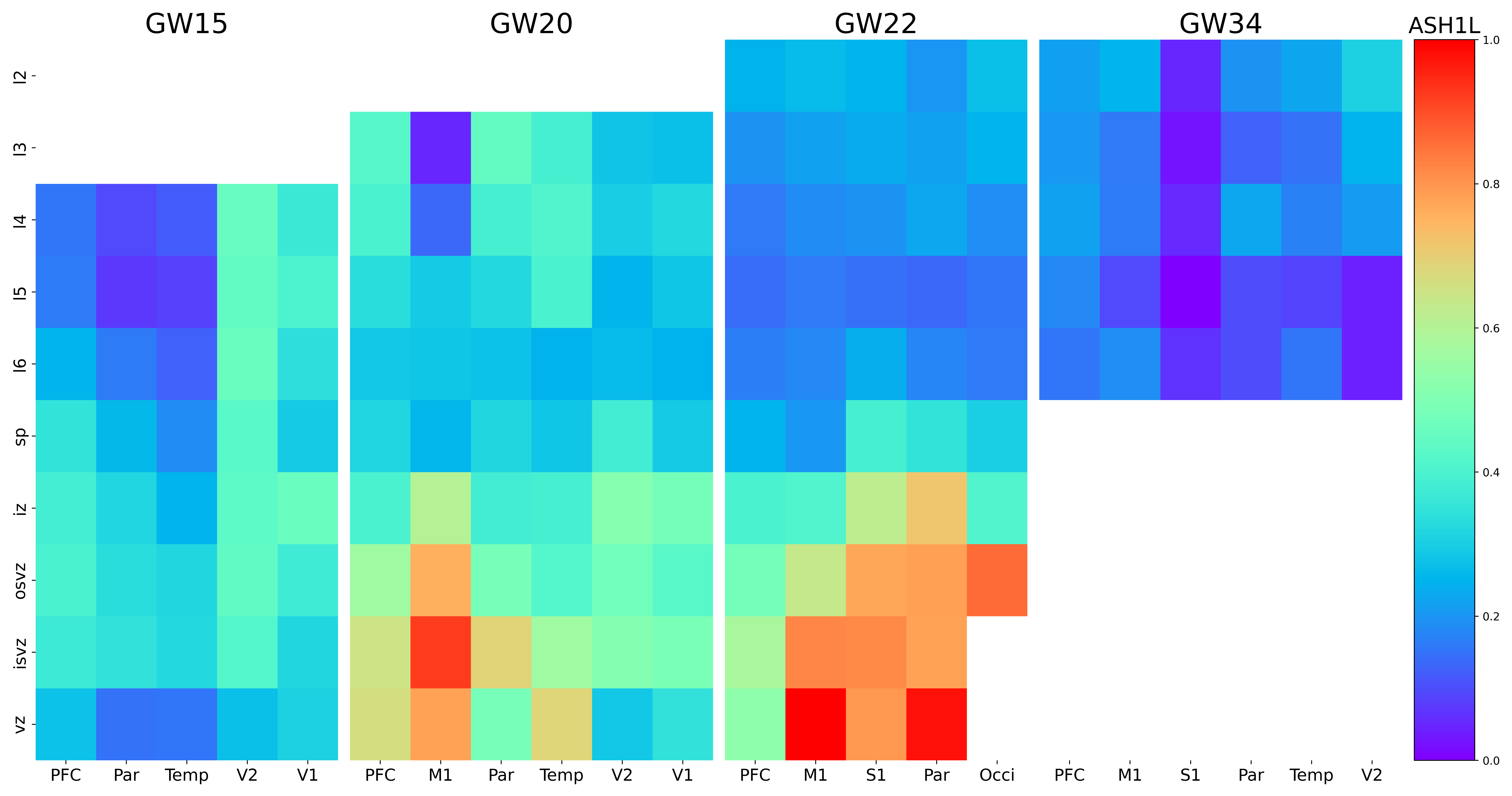

### ATP2B4.png

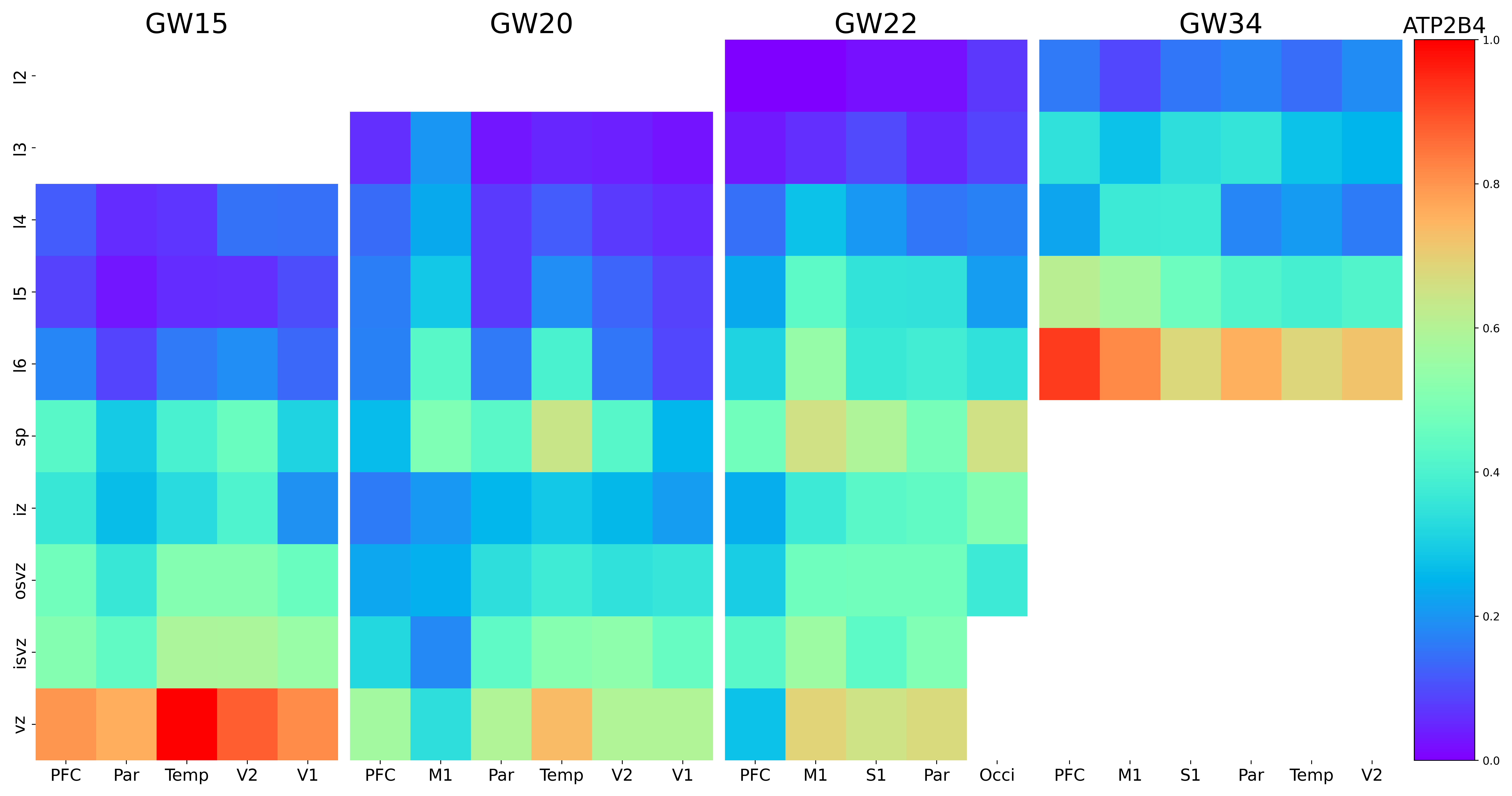

### B3GALT2.png

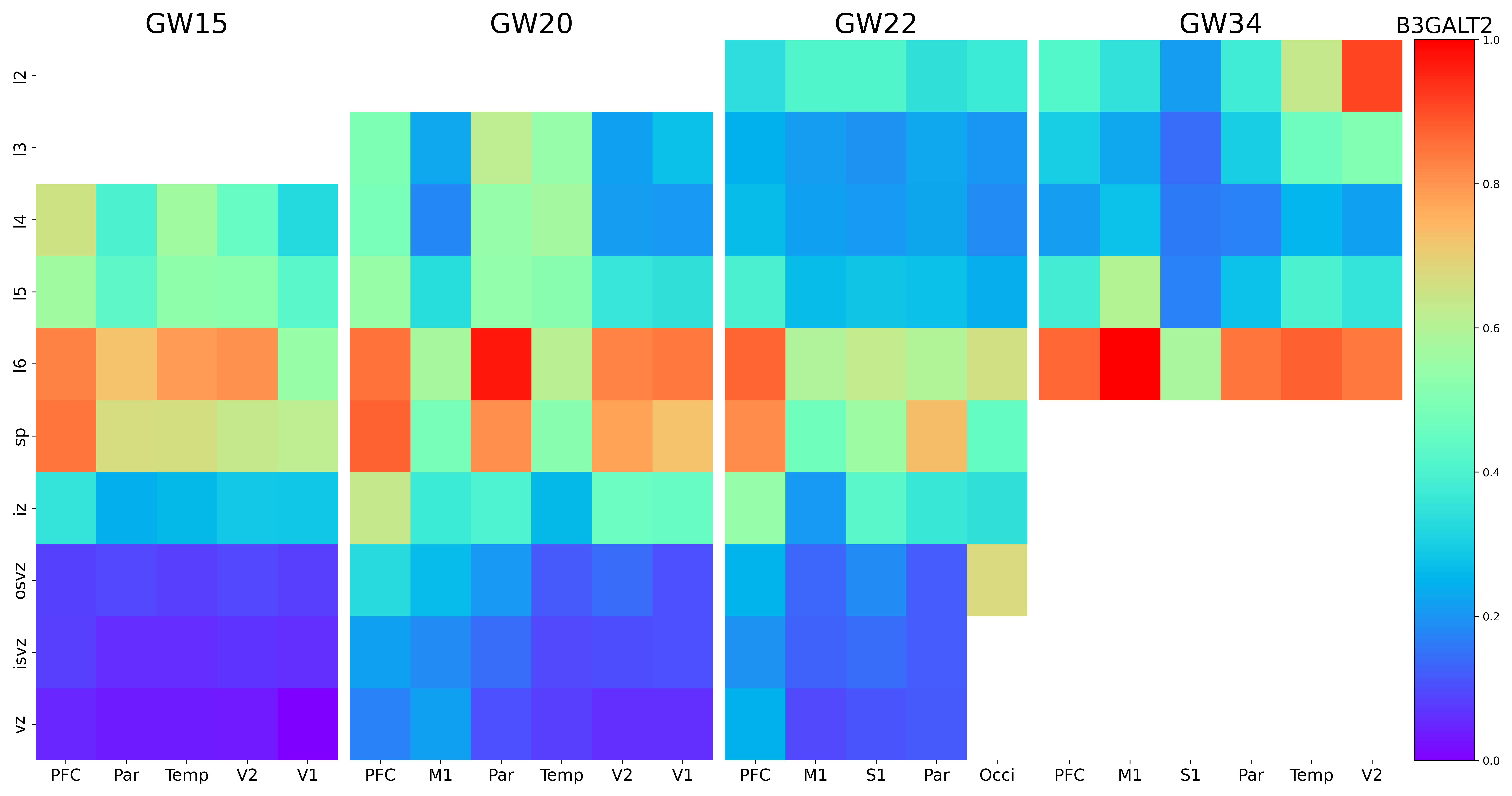

### B3GNT2.png

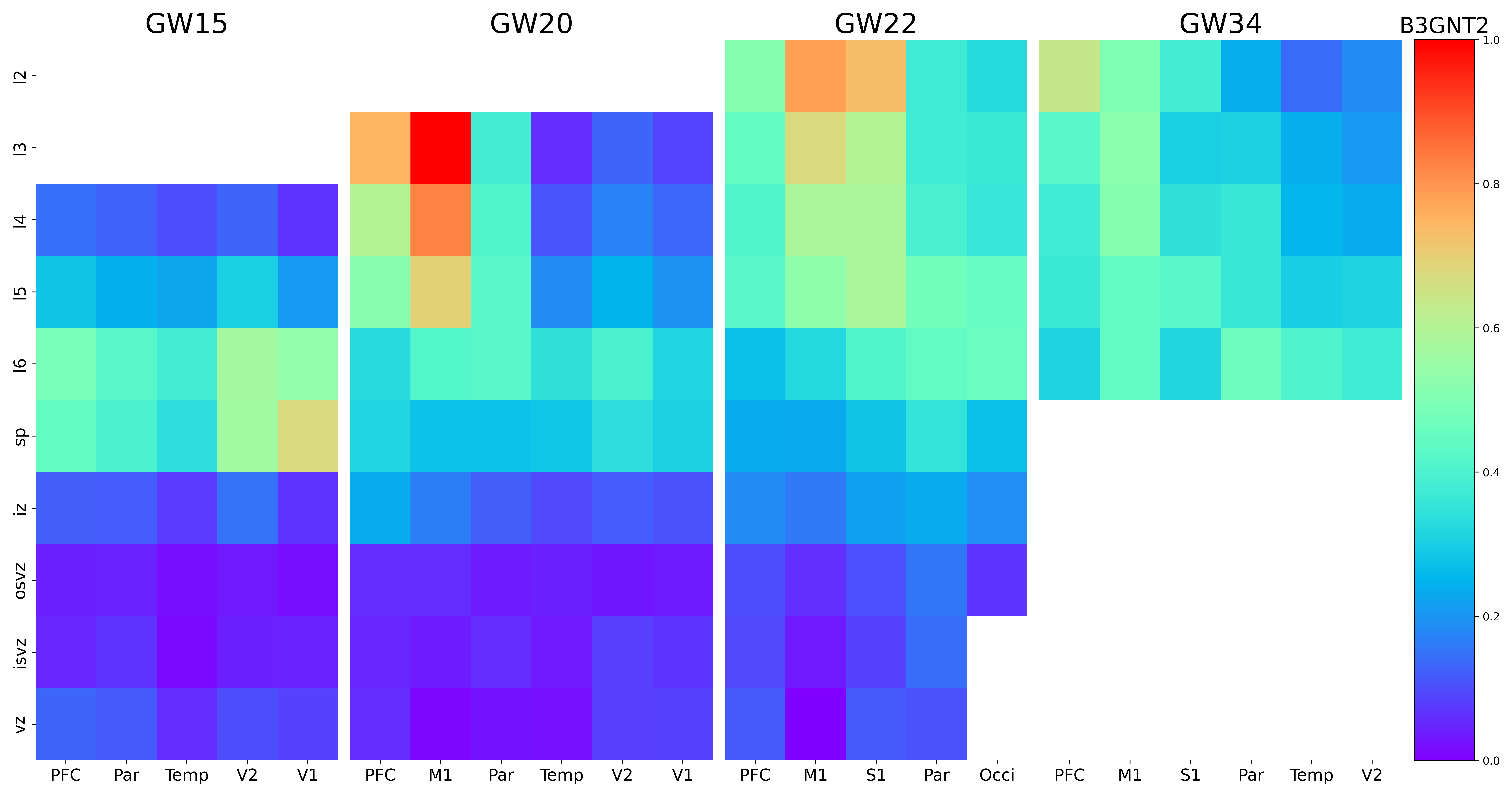

### BACH2.png

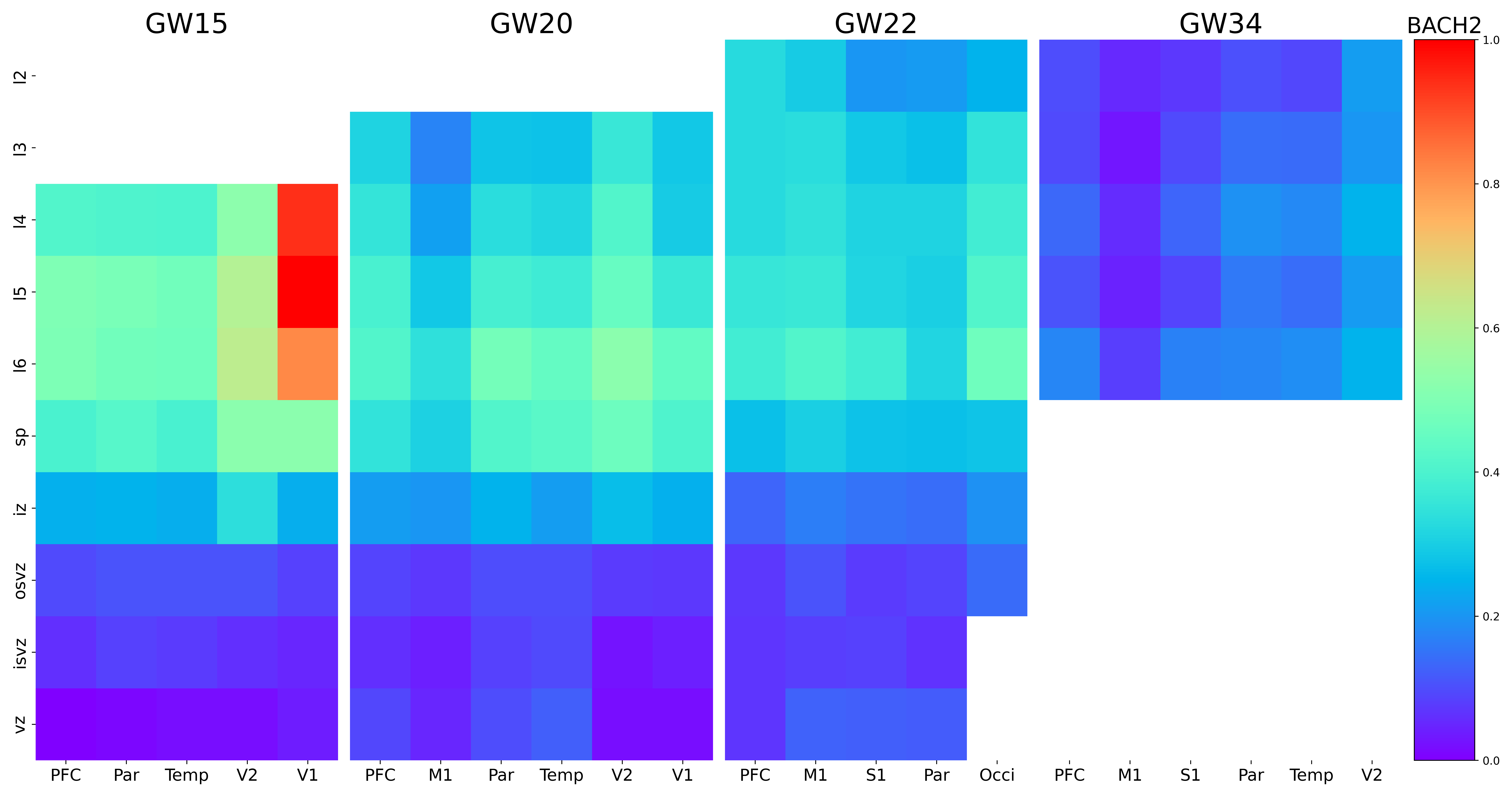

### BAZ2B.png

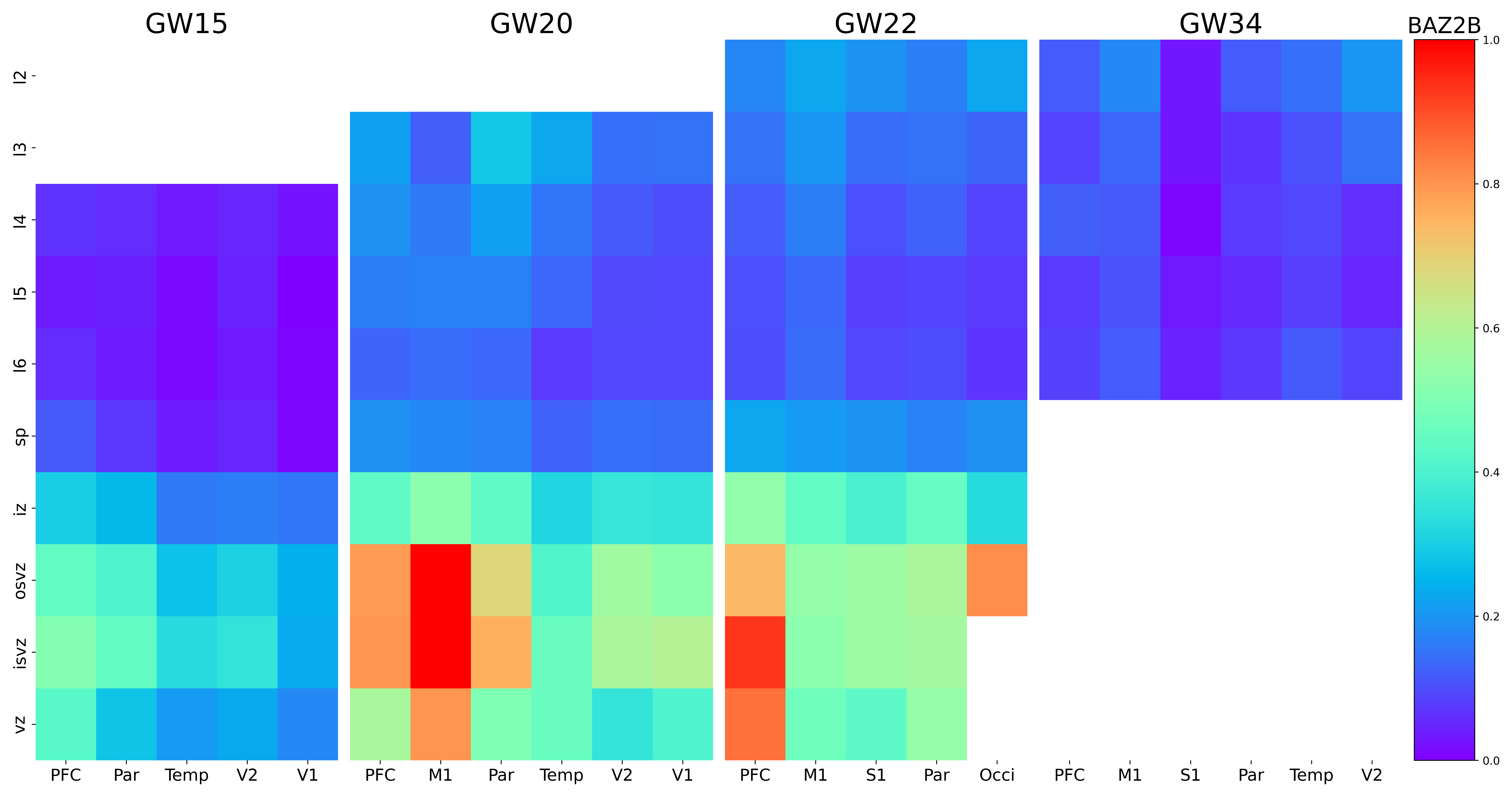

### BCL11B.png

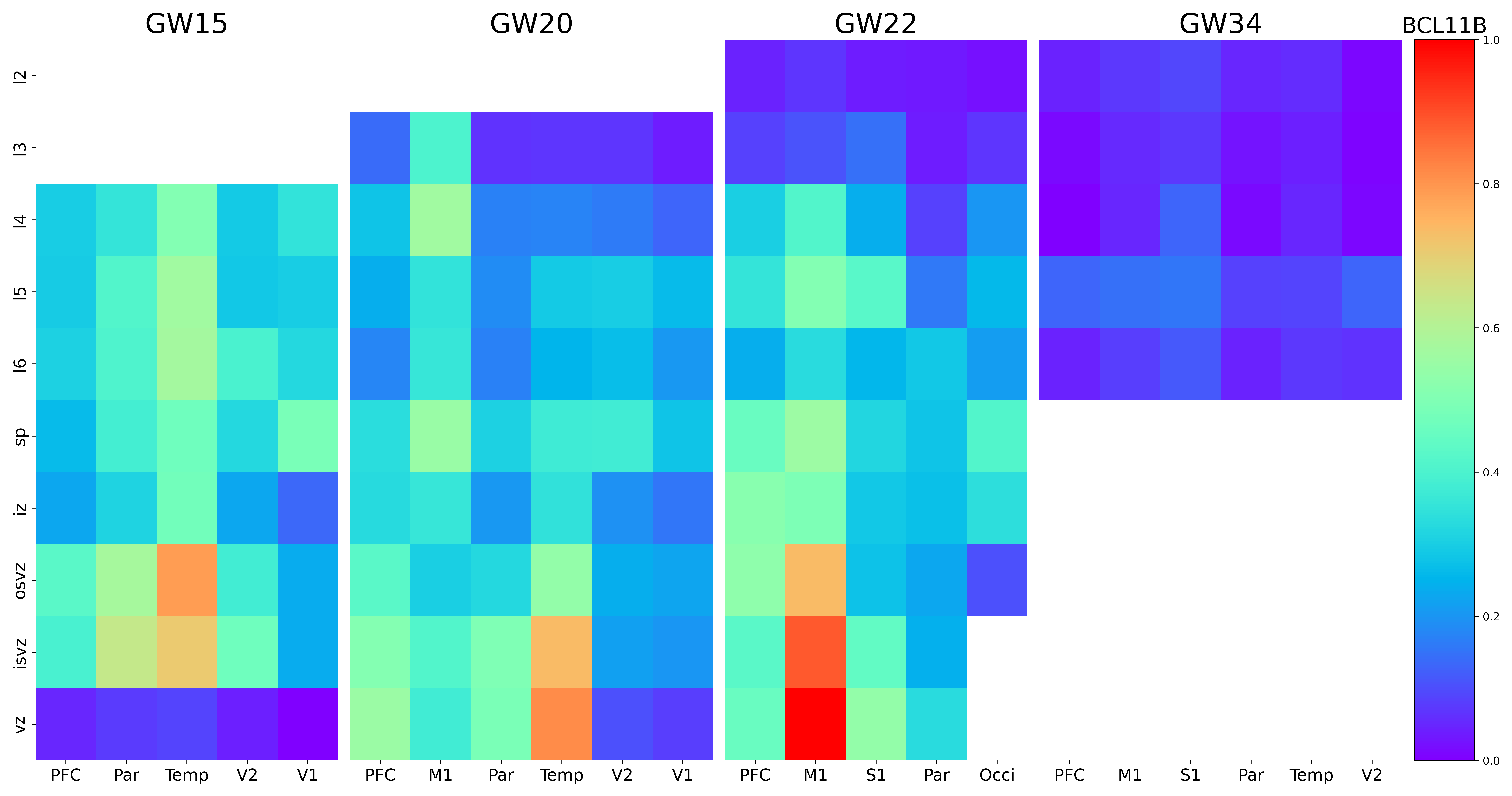

### BEST3.png

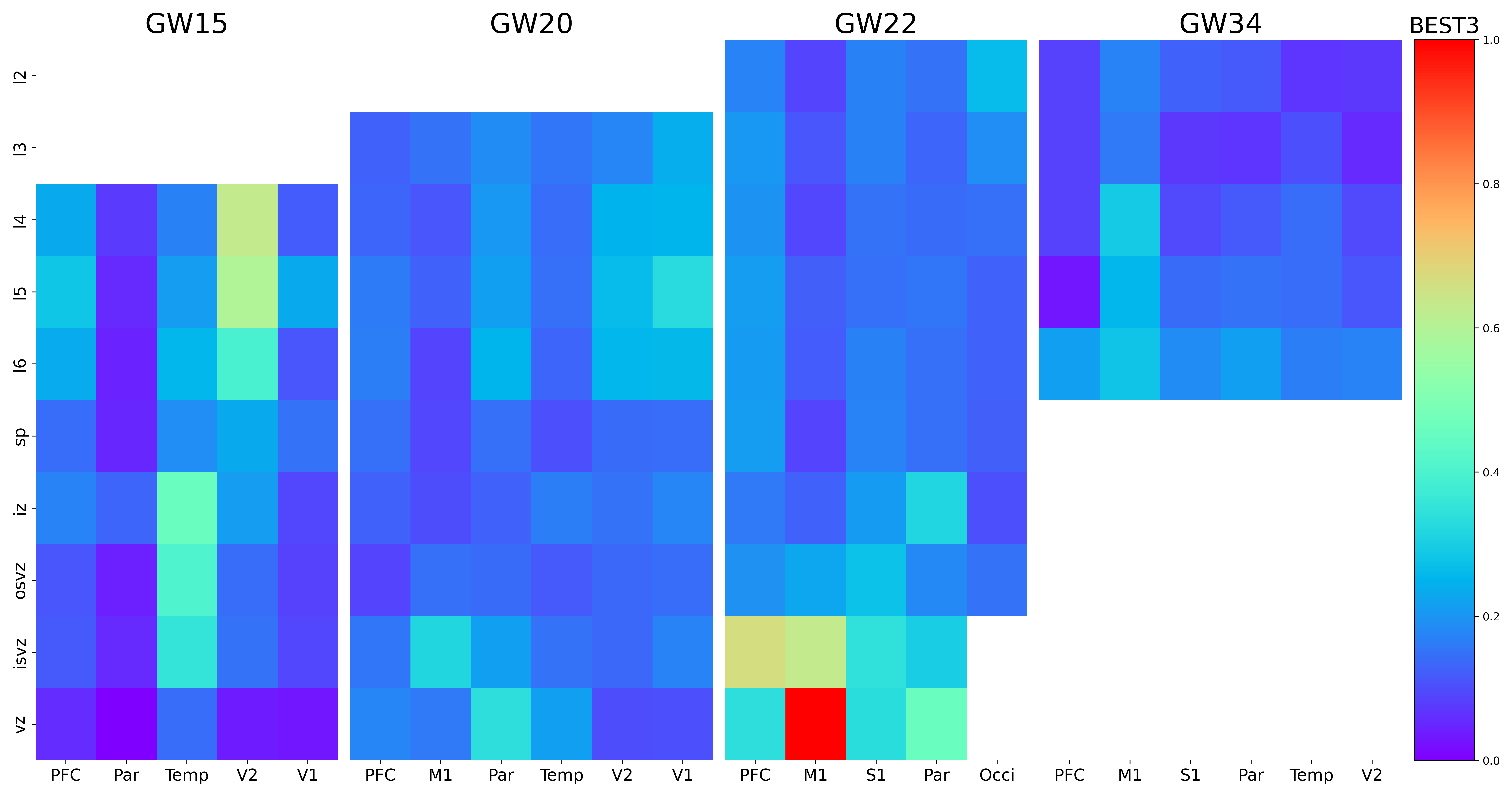

### C8orf34.png

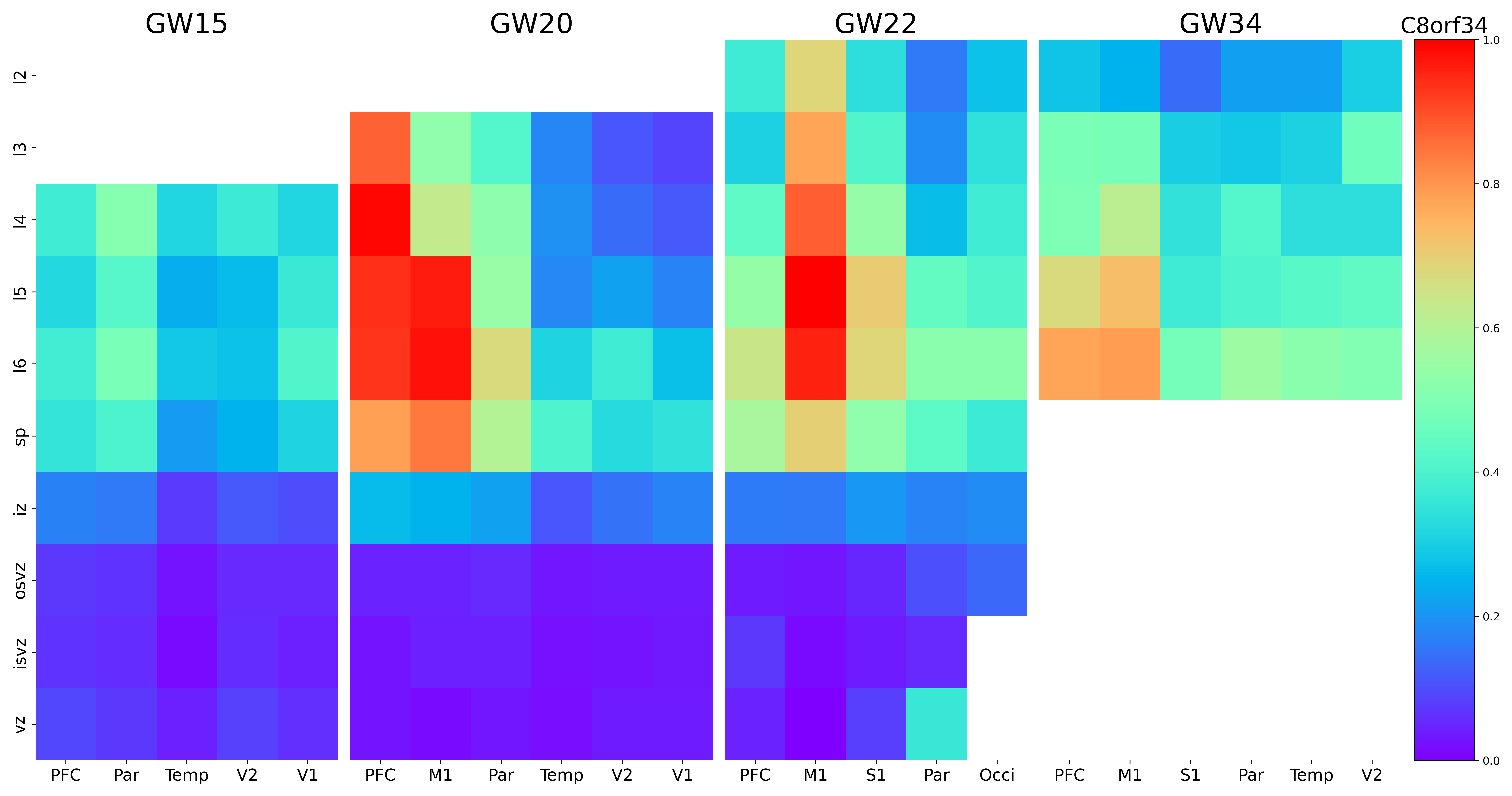

### CA12.png

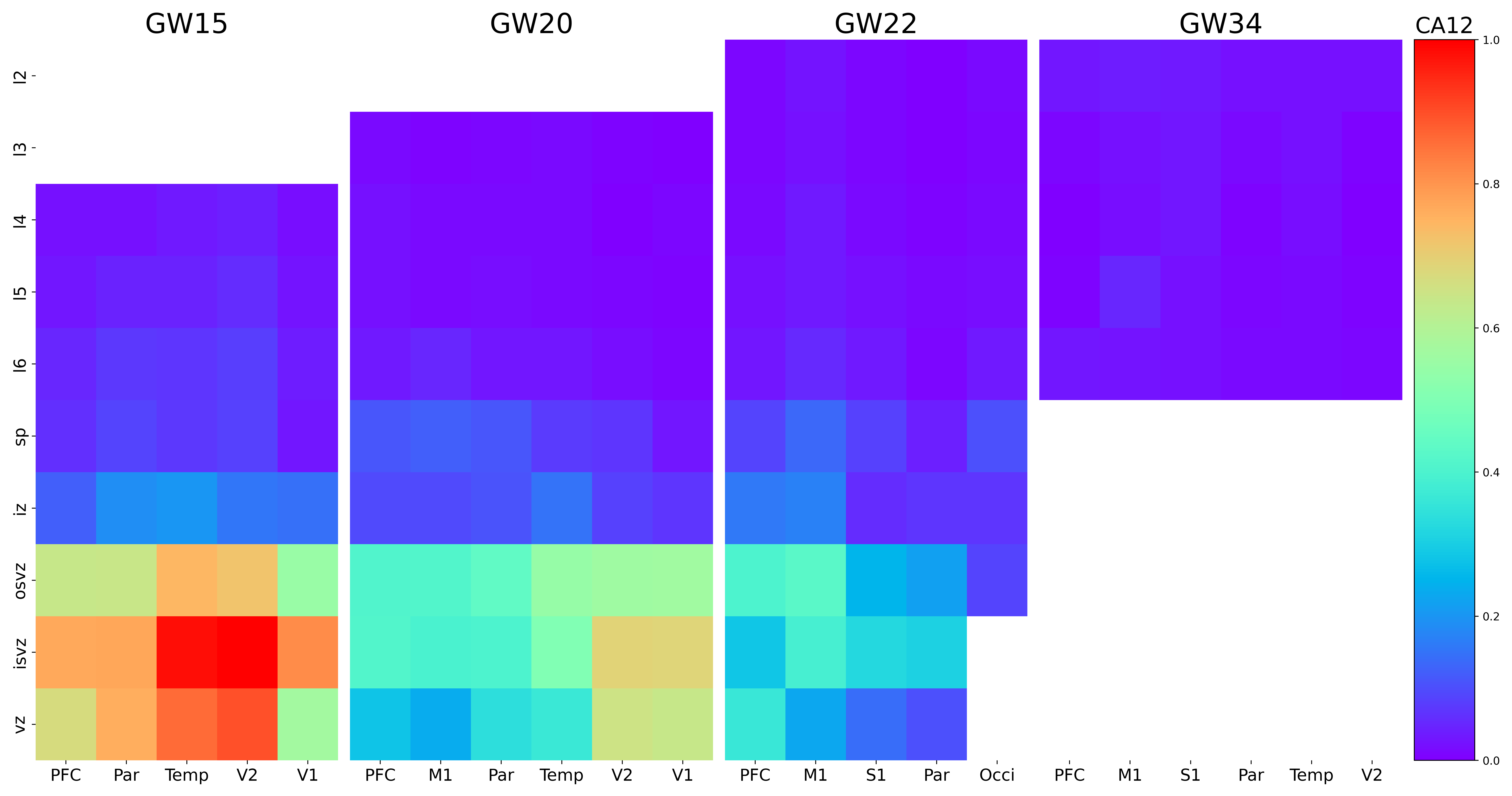

### CACNA1E.png

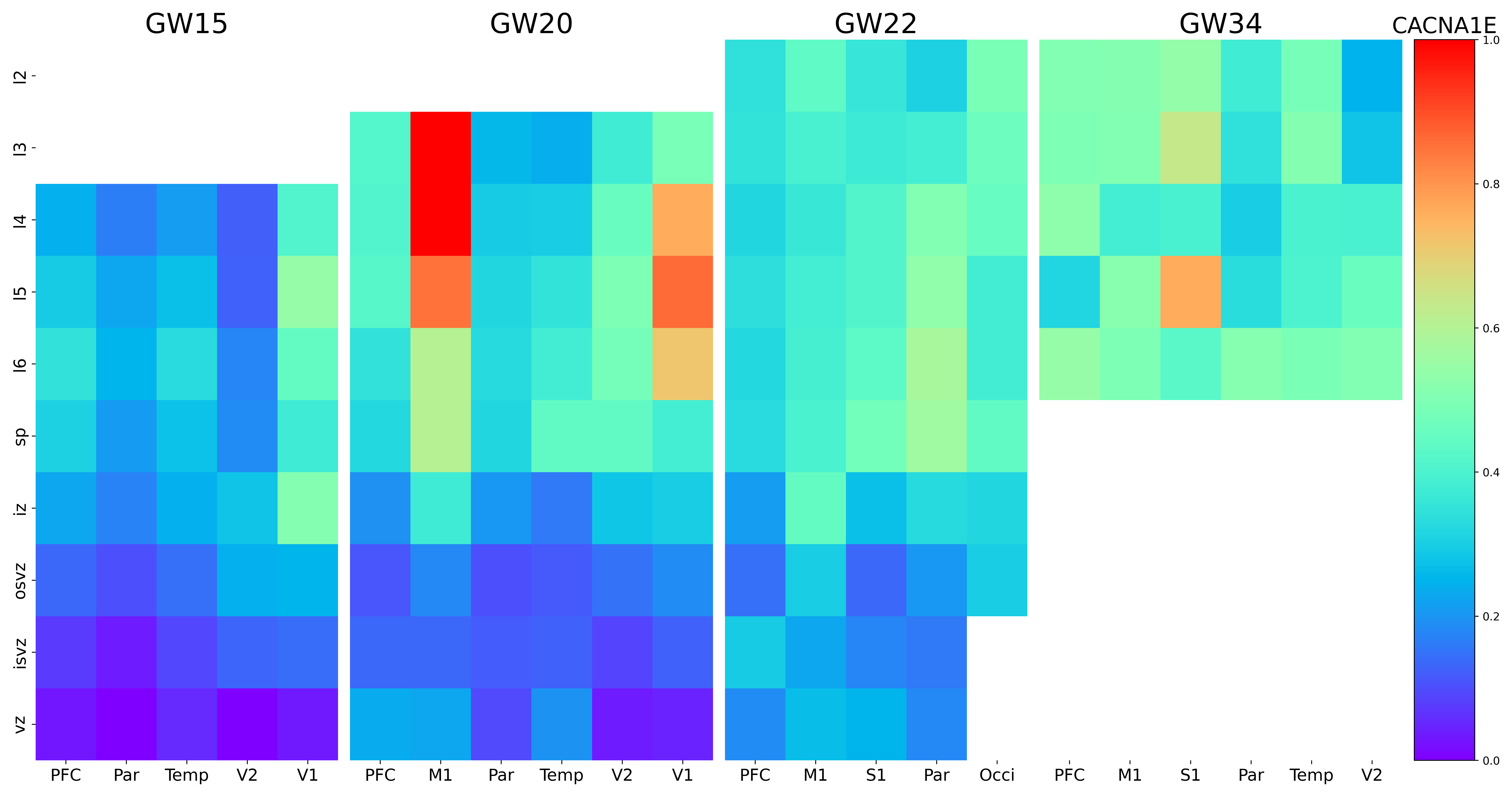

### CALN1.png

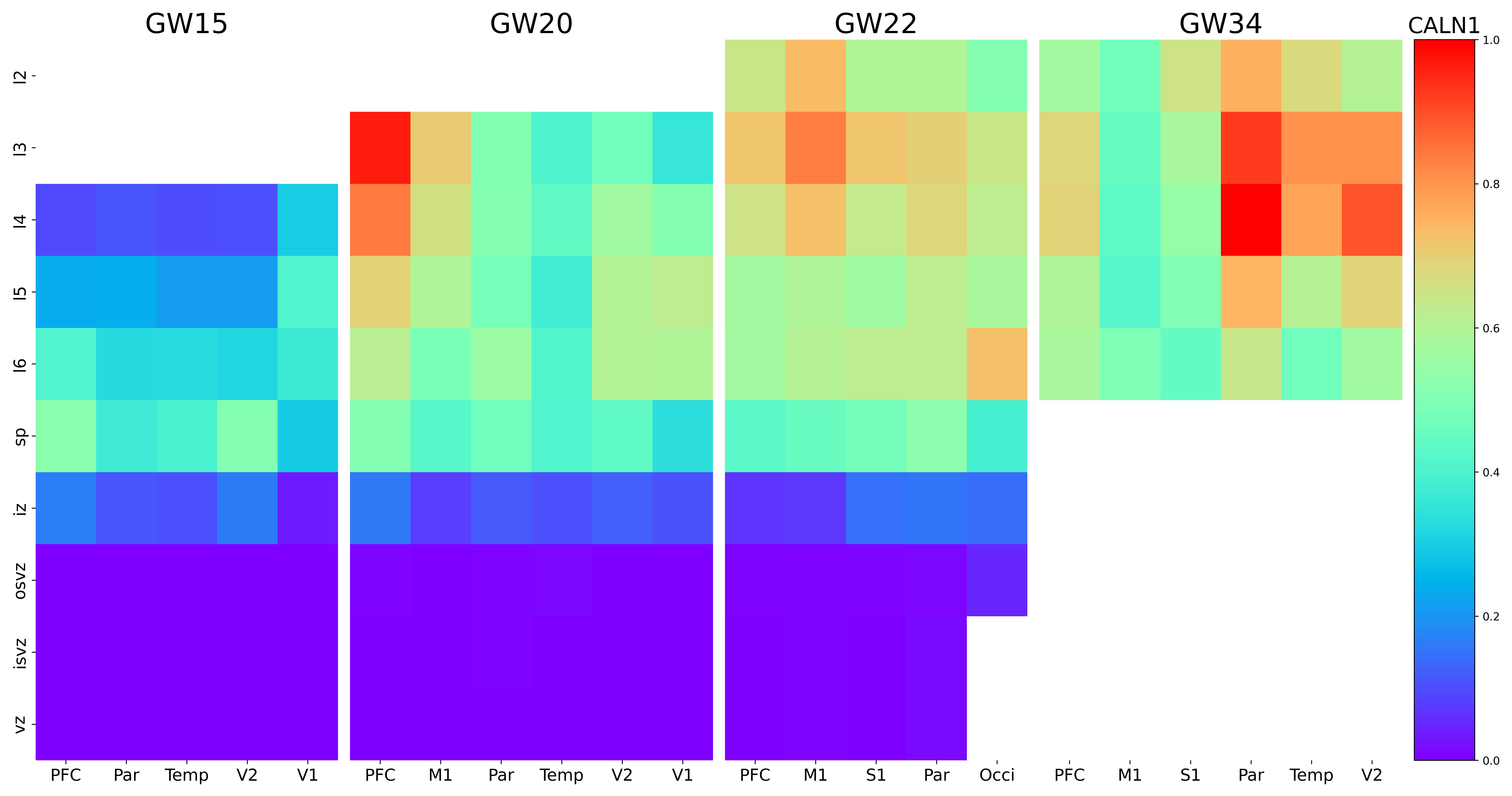

### CAMK2B.png

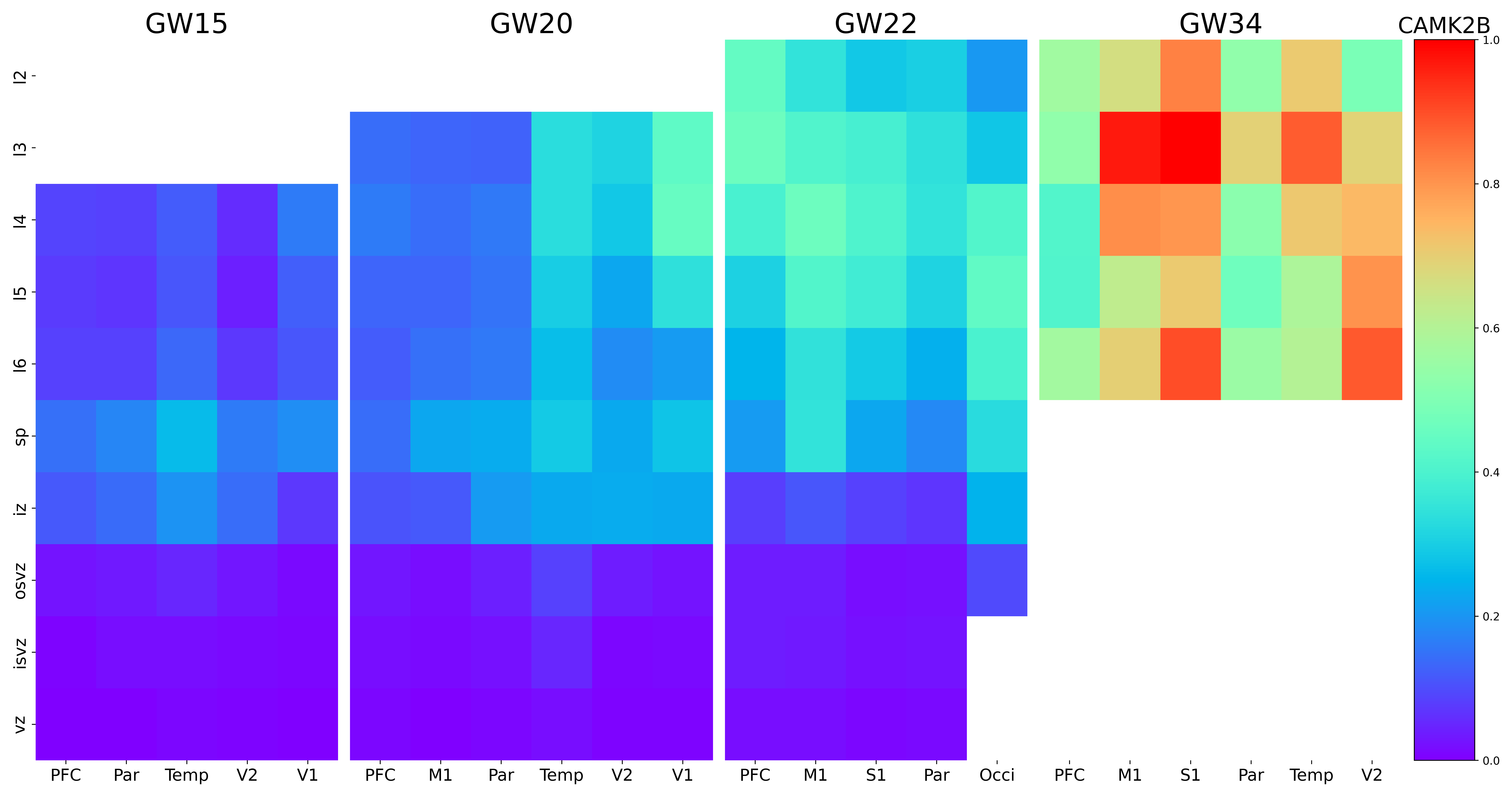

### CBLN2.png

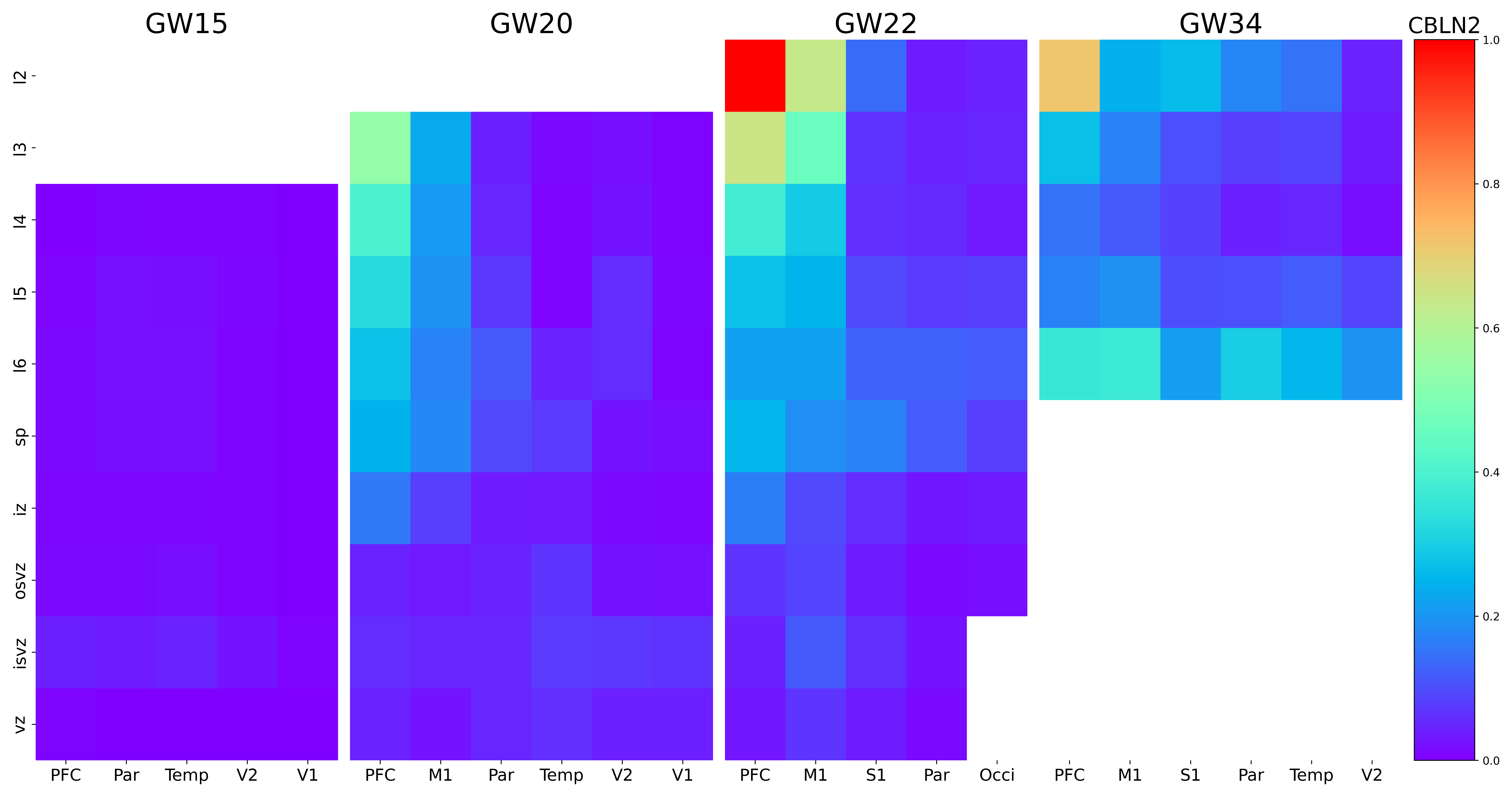

### CBLN4.png

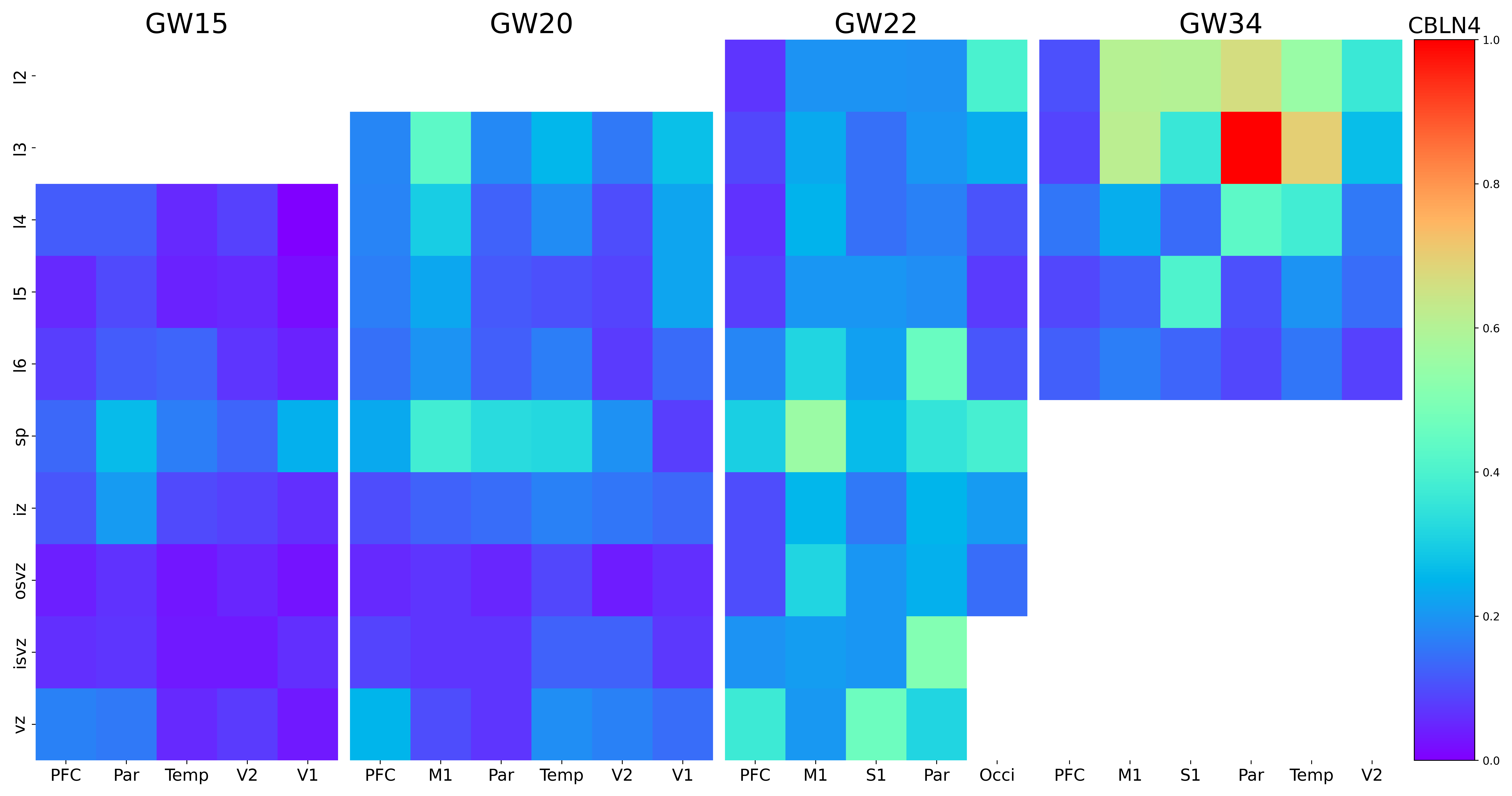

### CCDC144A.png

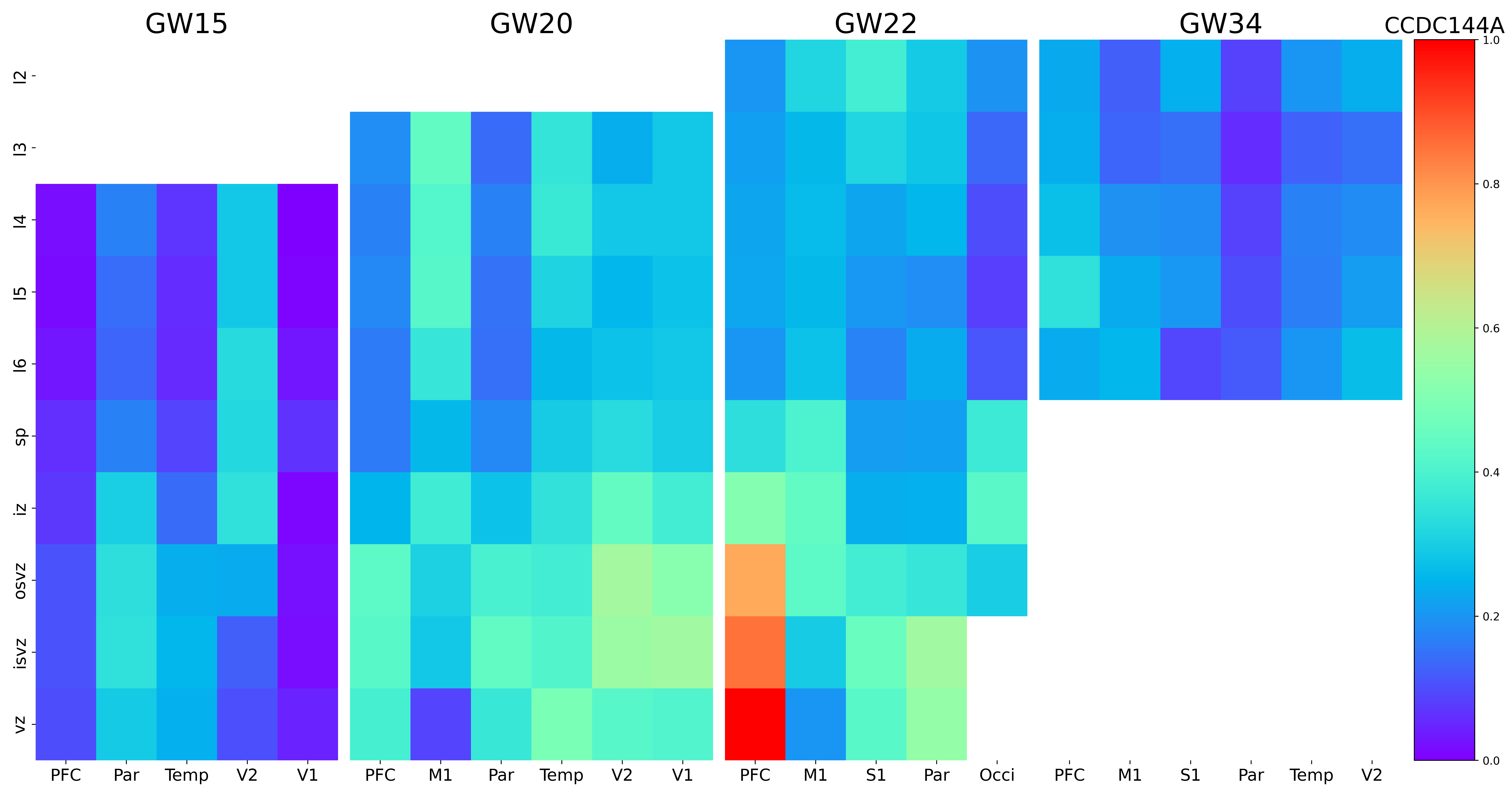

### CCDC175.png

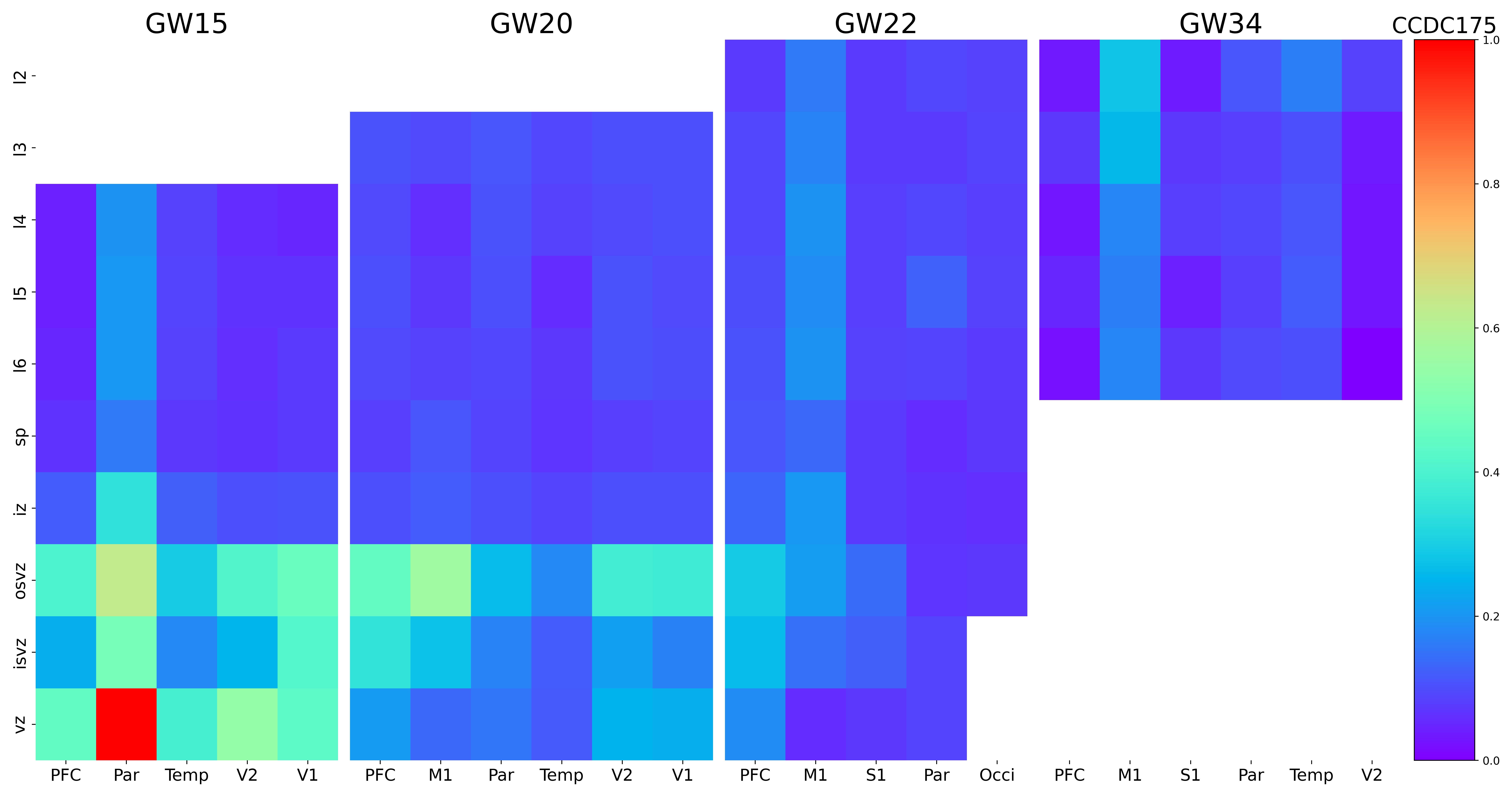

### CCN1.png

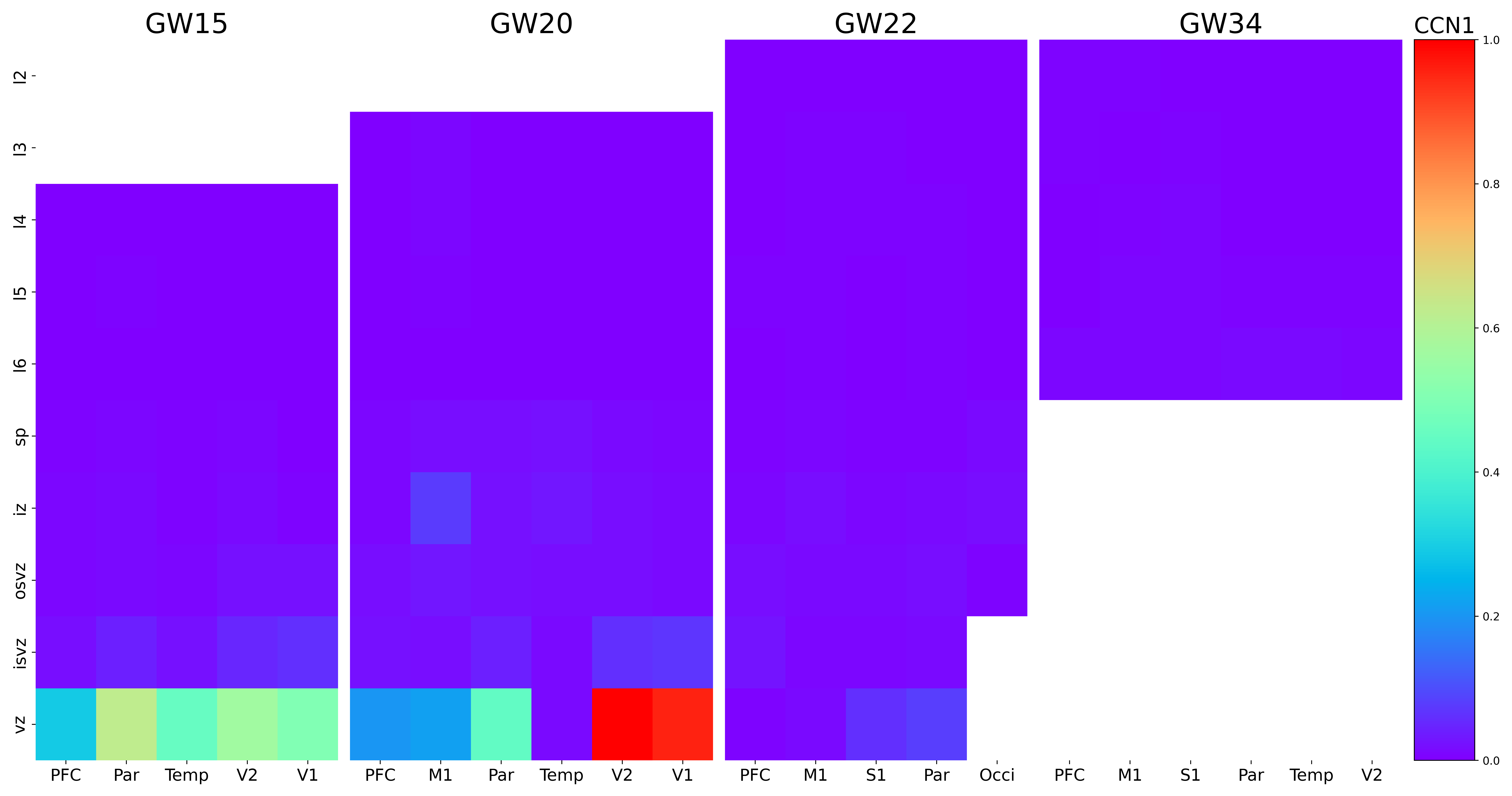

### CCN2.png

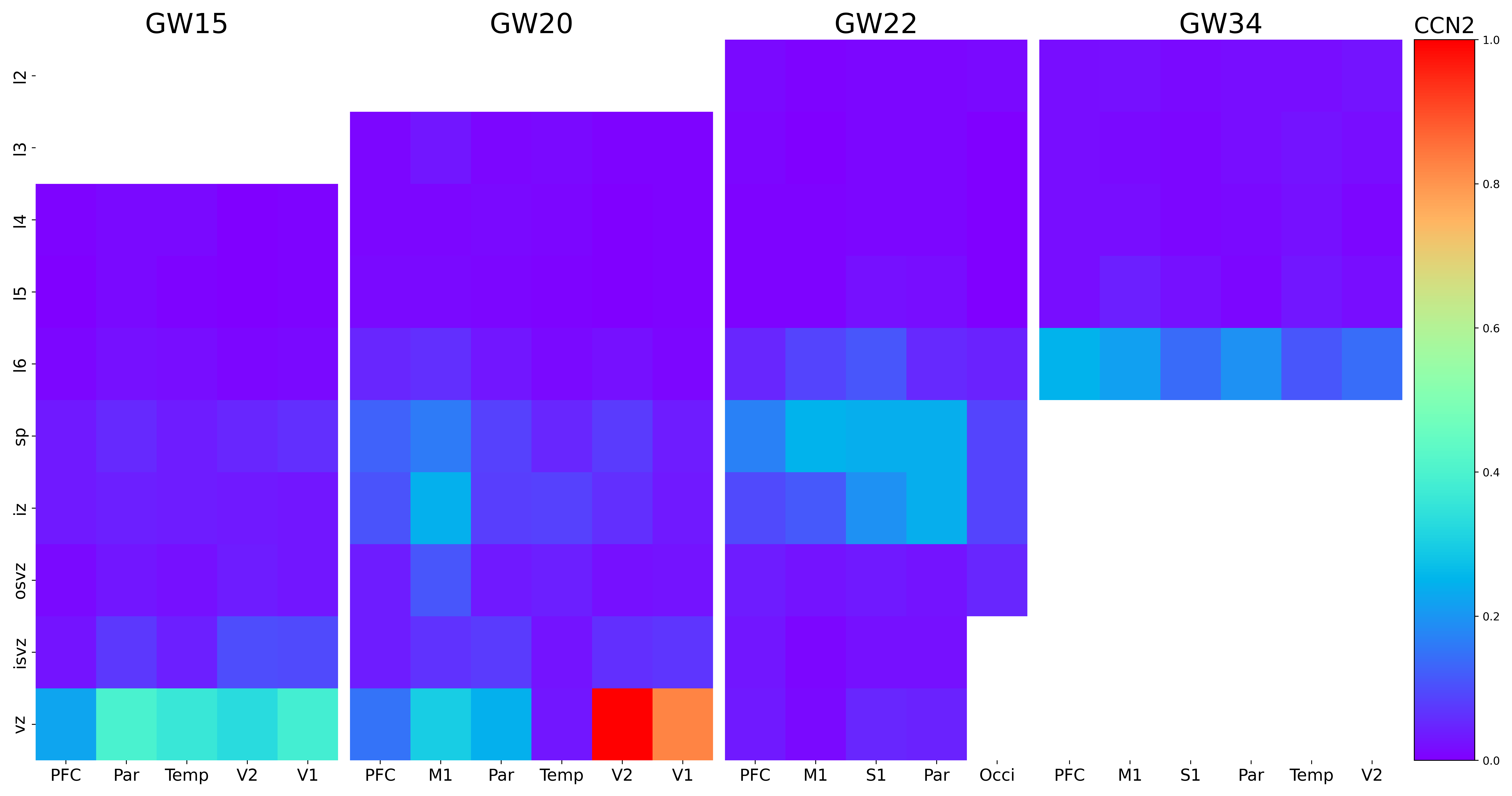

### CD9.png

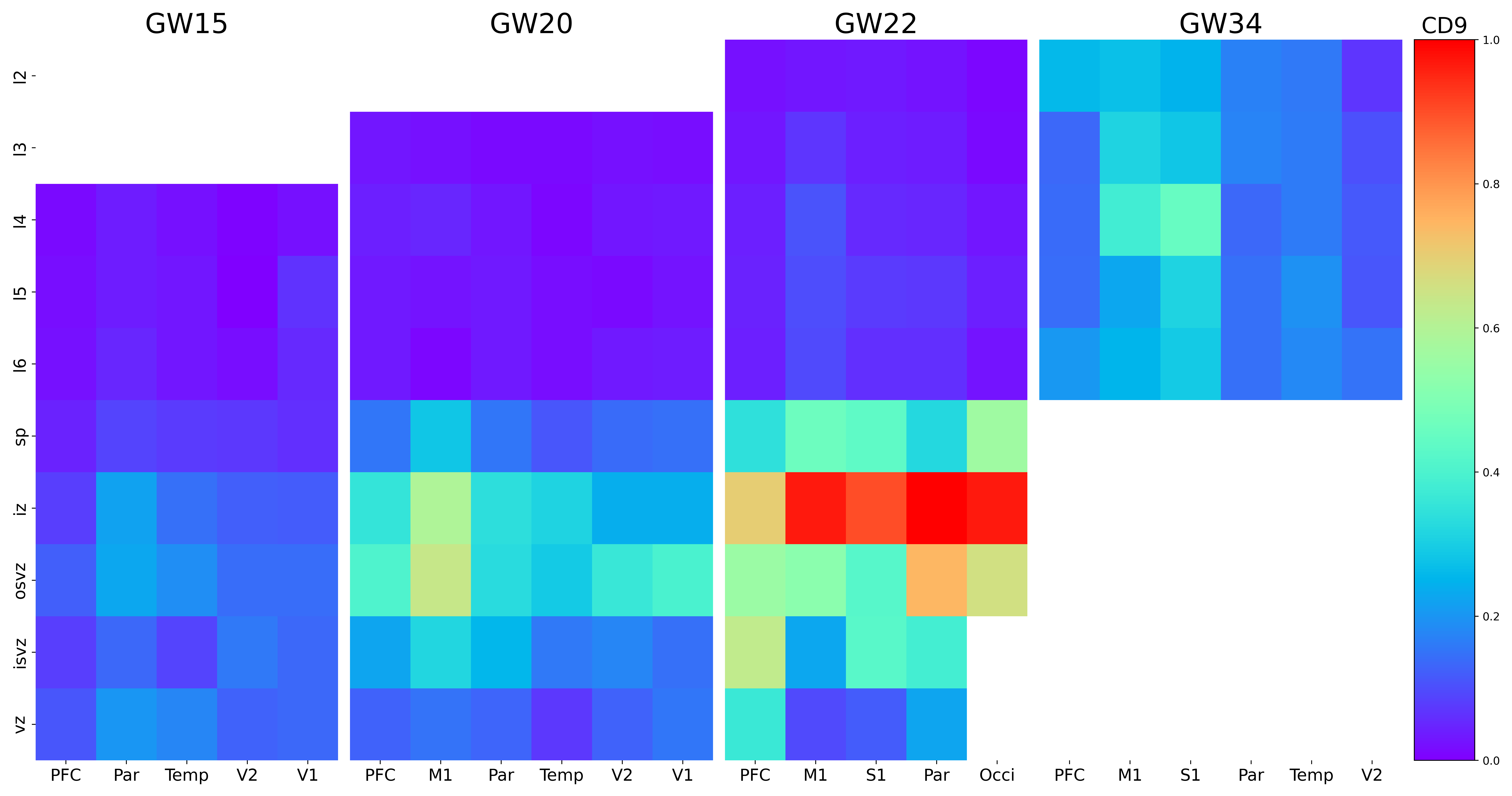

### CD38.png

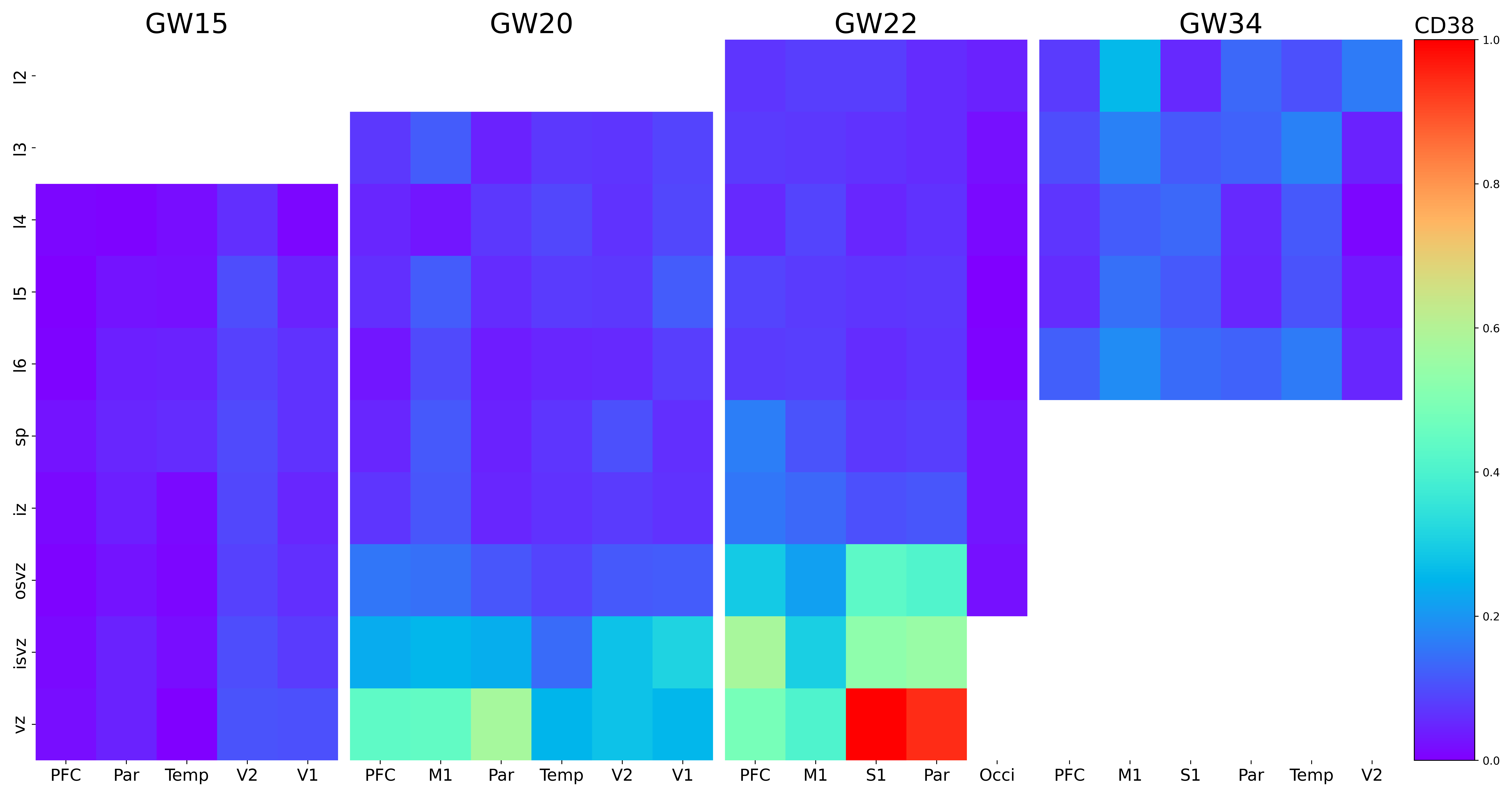

### CD82.png

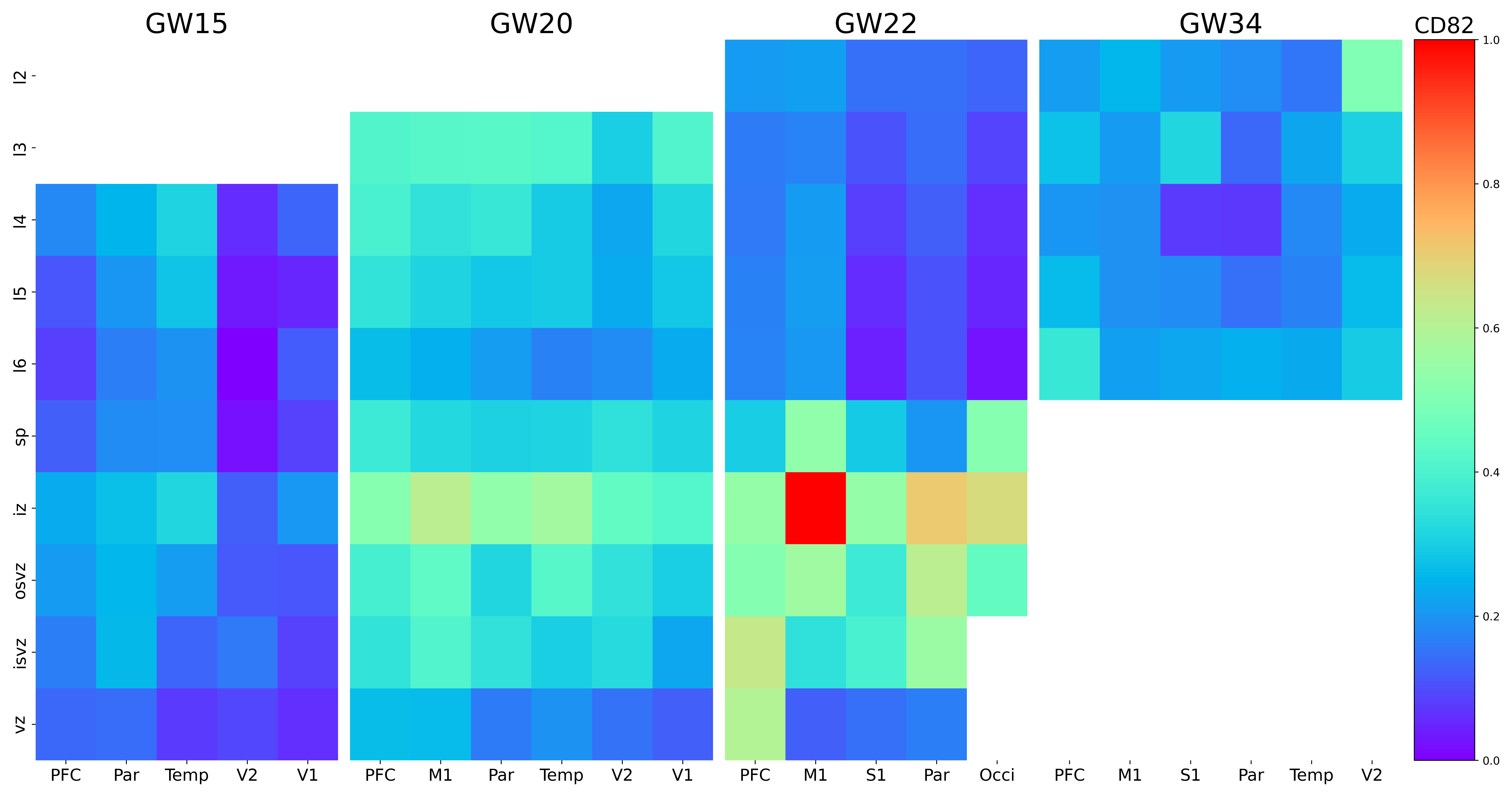

### CDH13.png

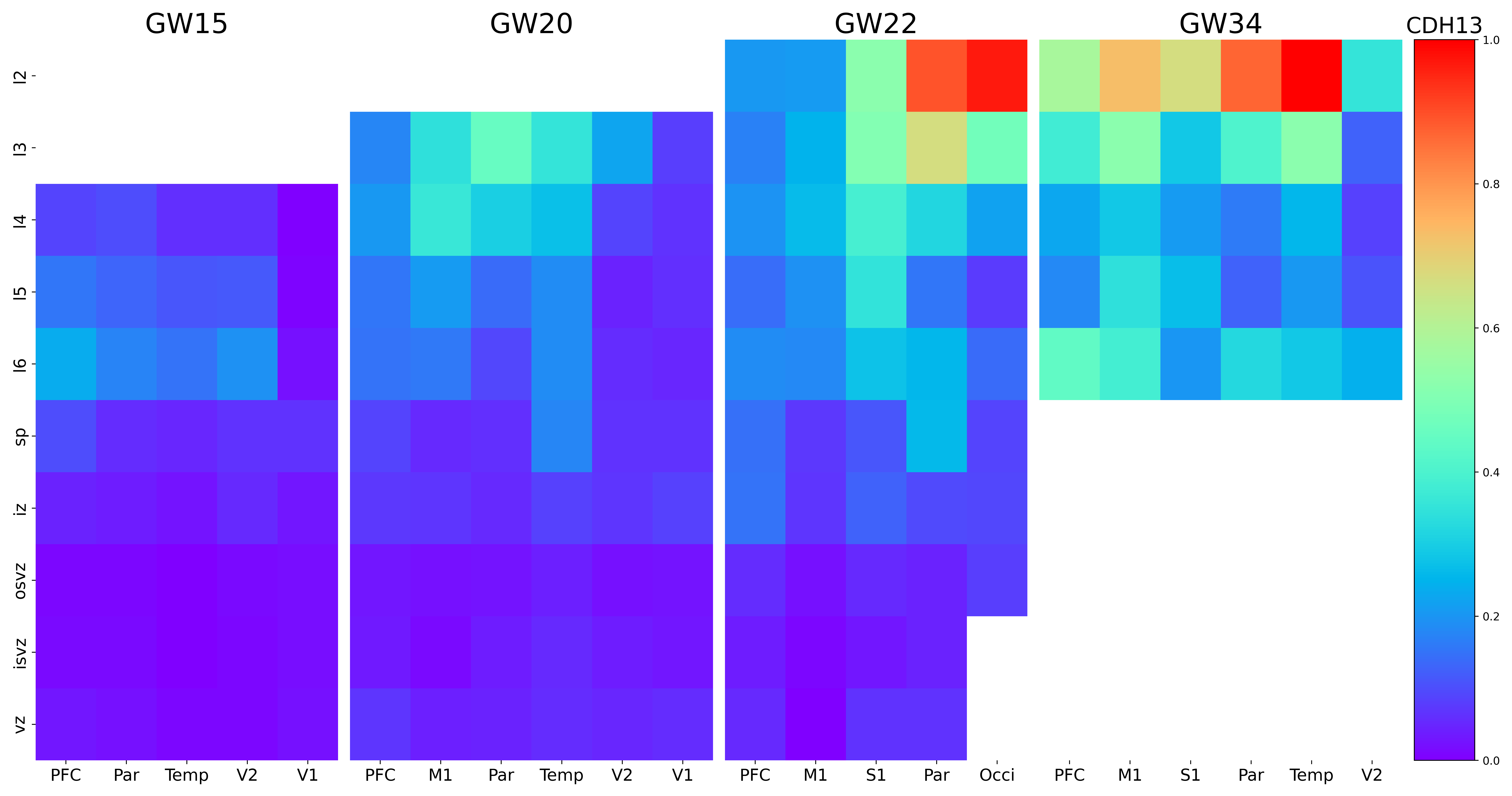

### CFTR.png

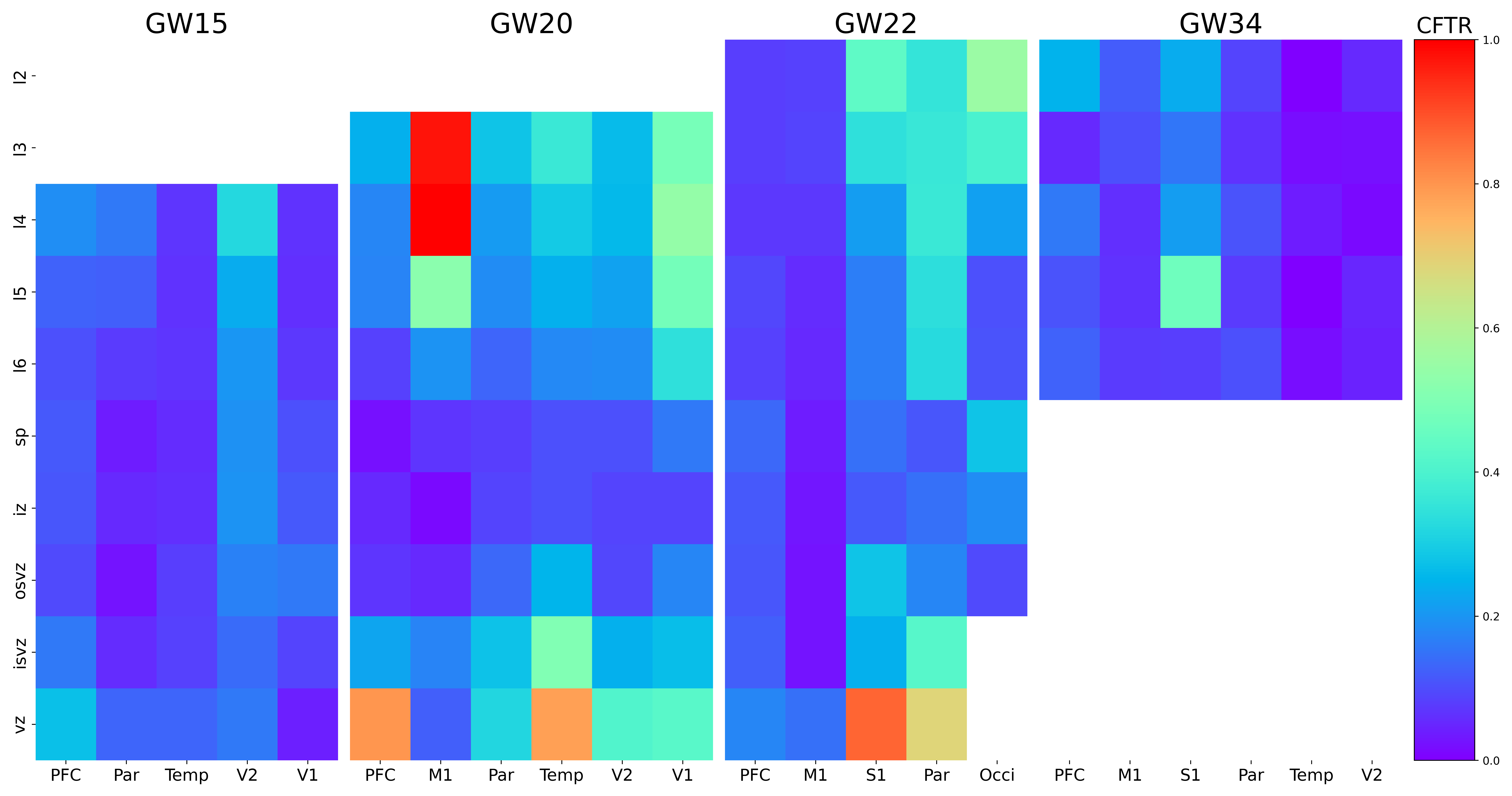

### CHST3.png

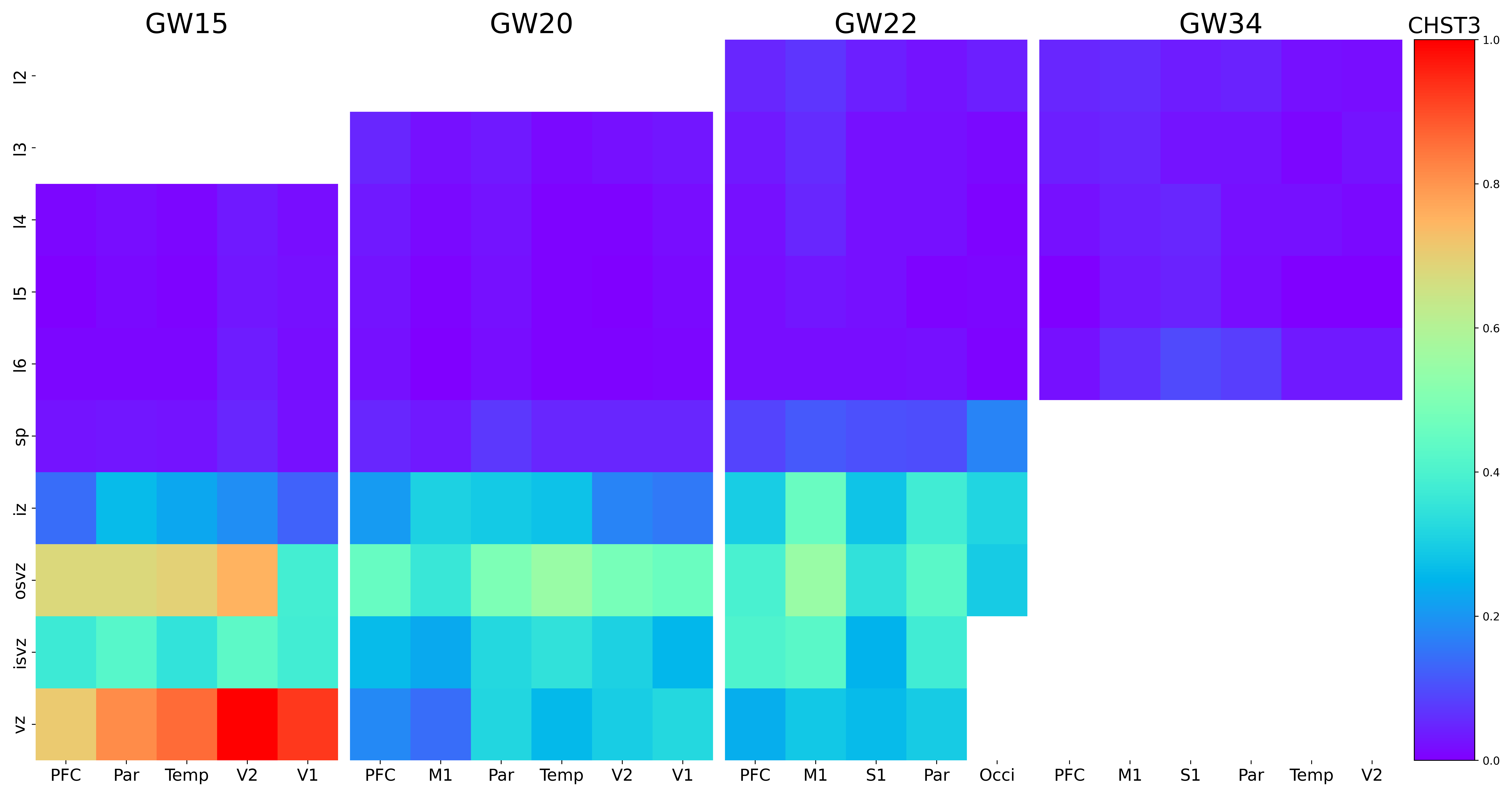

### CLASP1.png

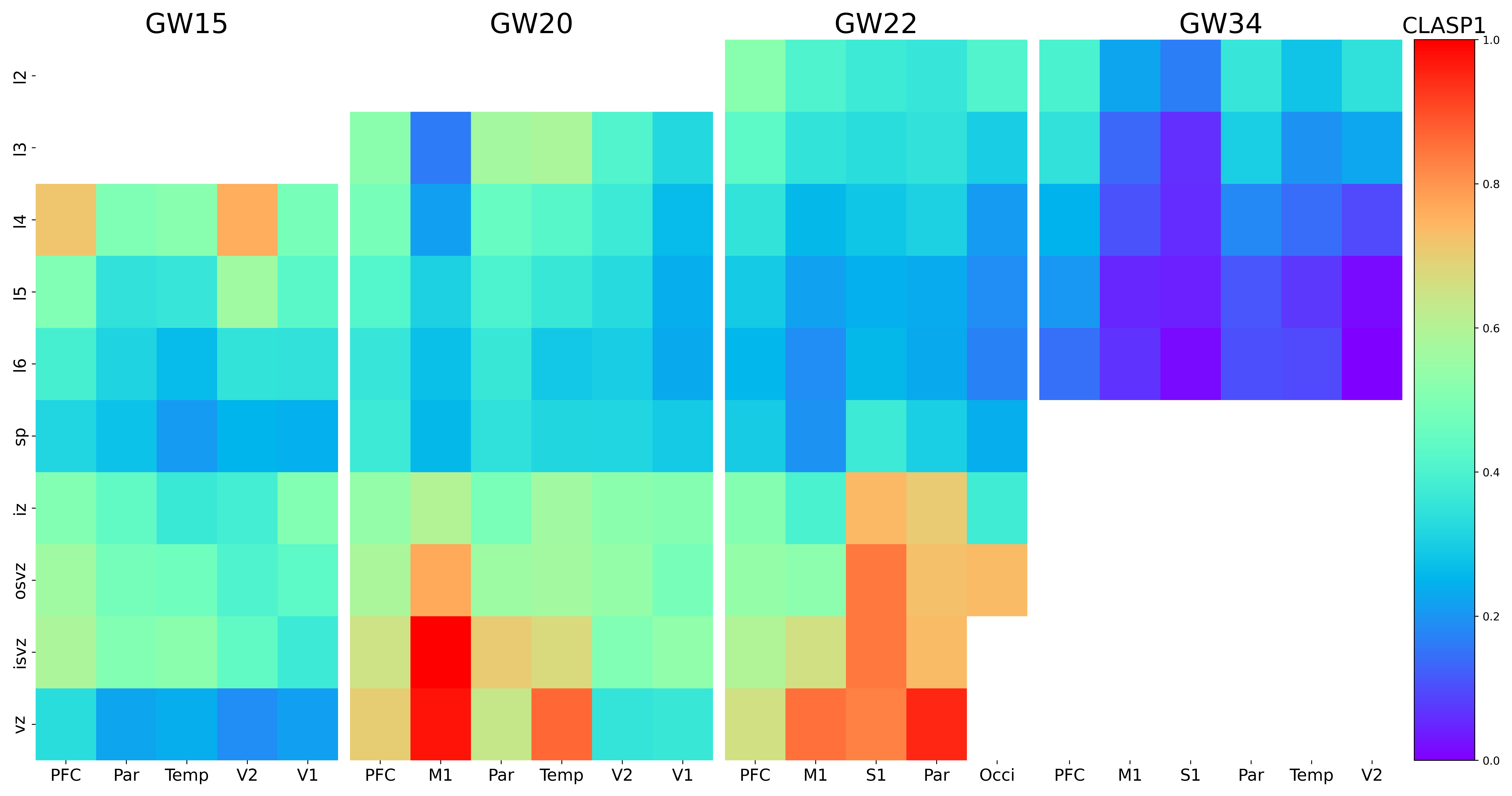

### CLSTN2.png

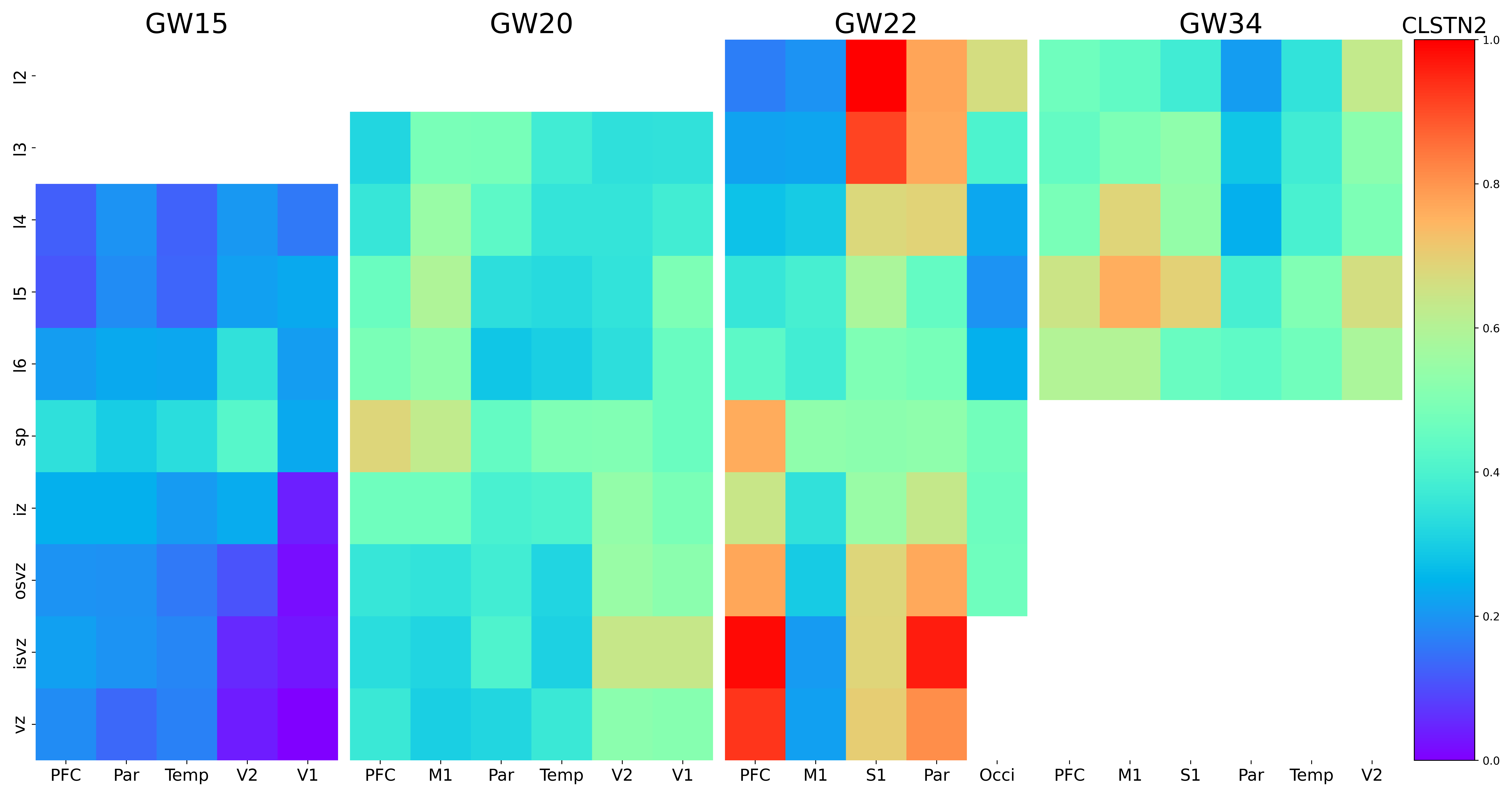

### CNTN2.png

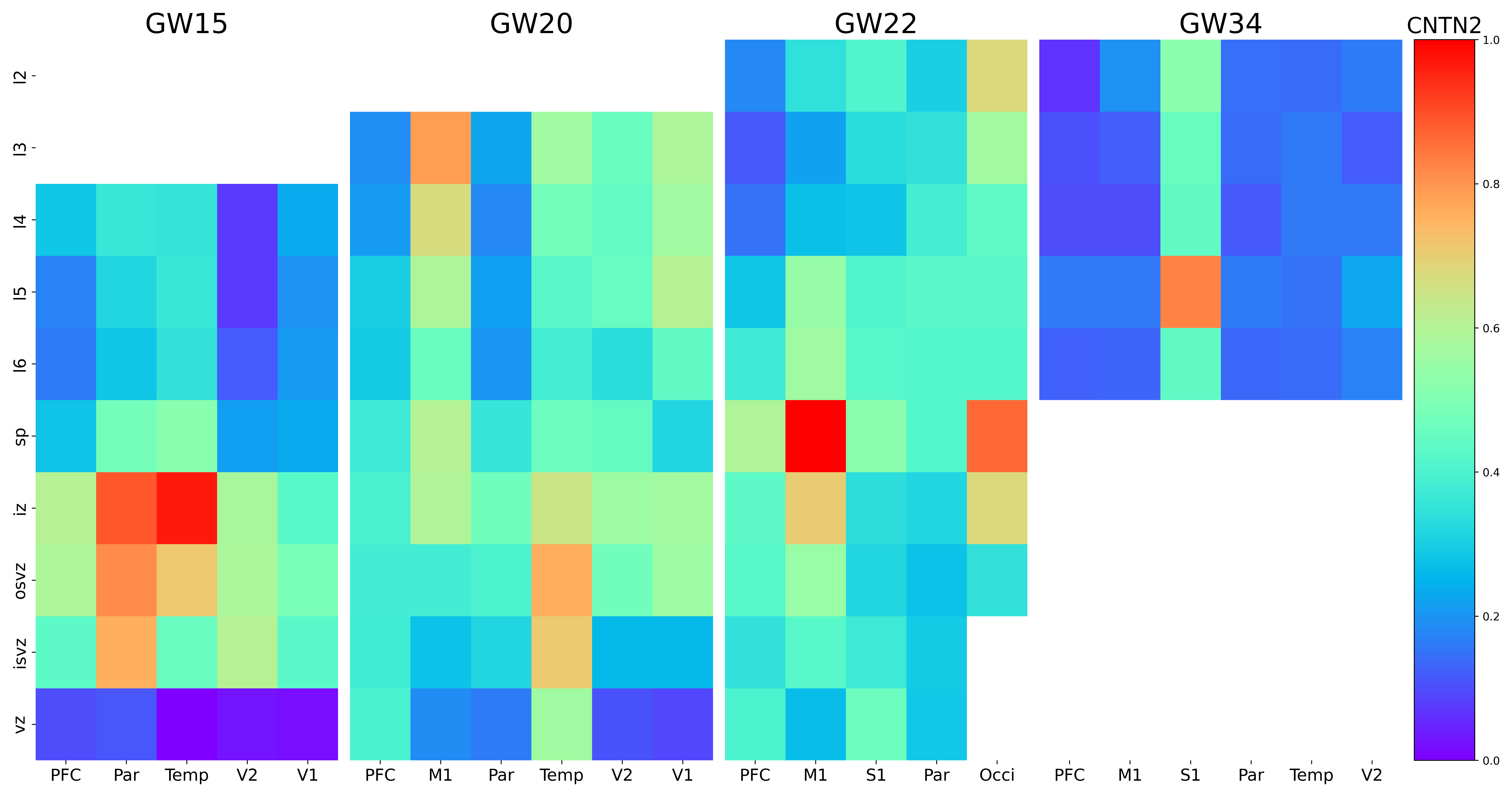

### COL4A5.png

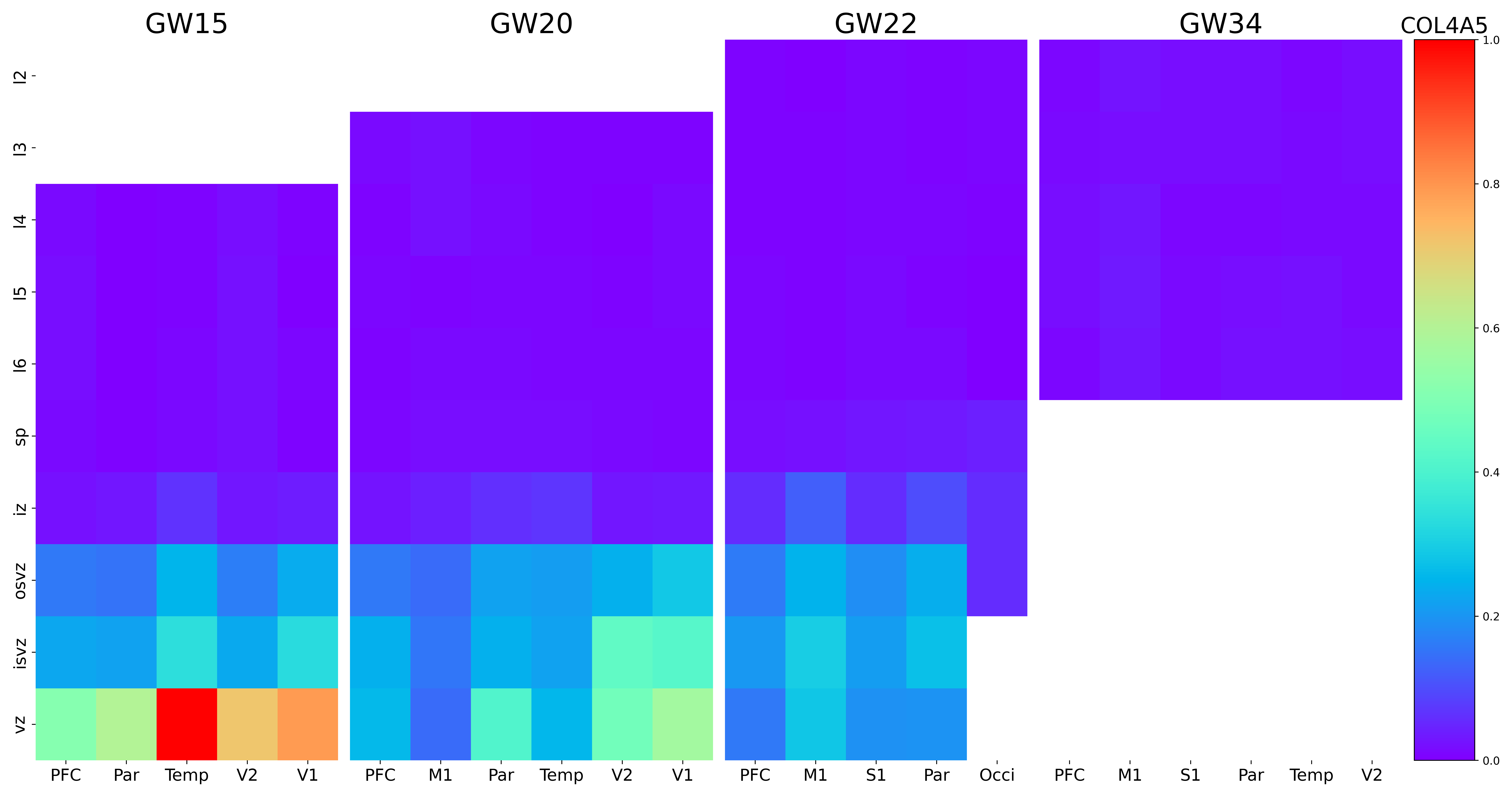

### COL4A6.png

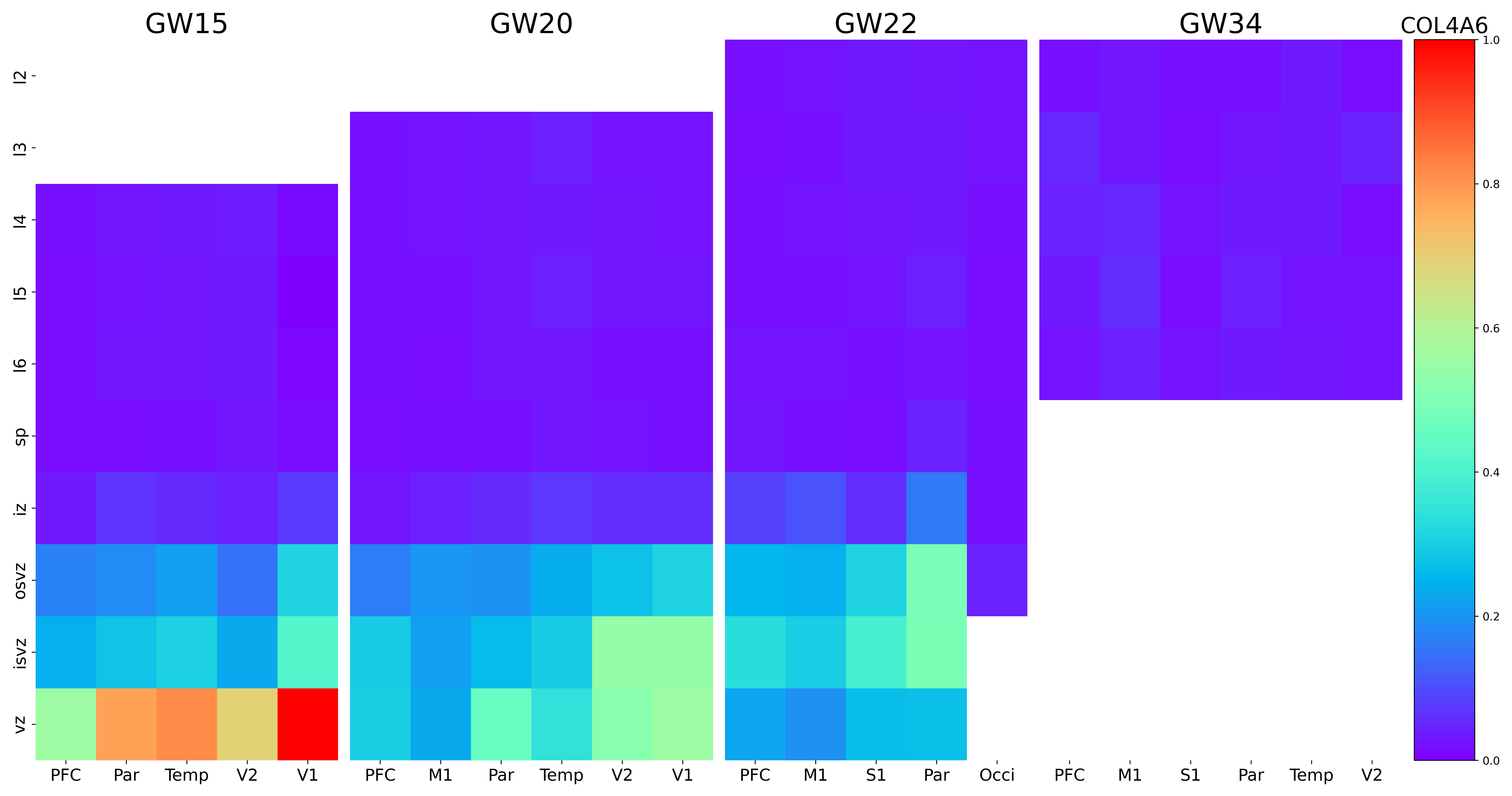

### COL6A1.png

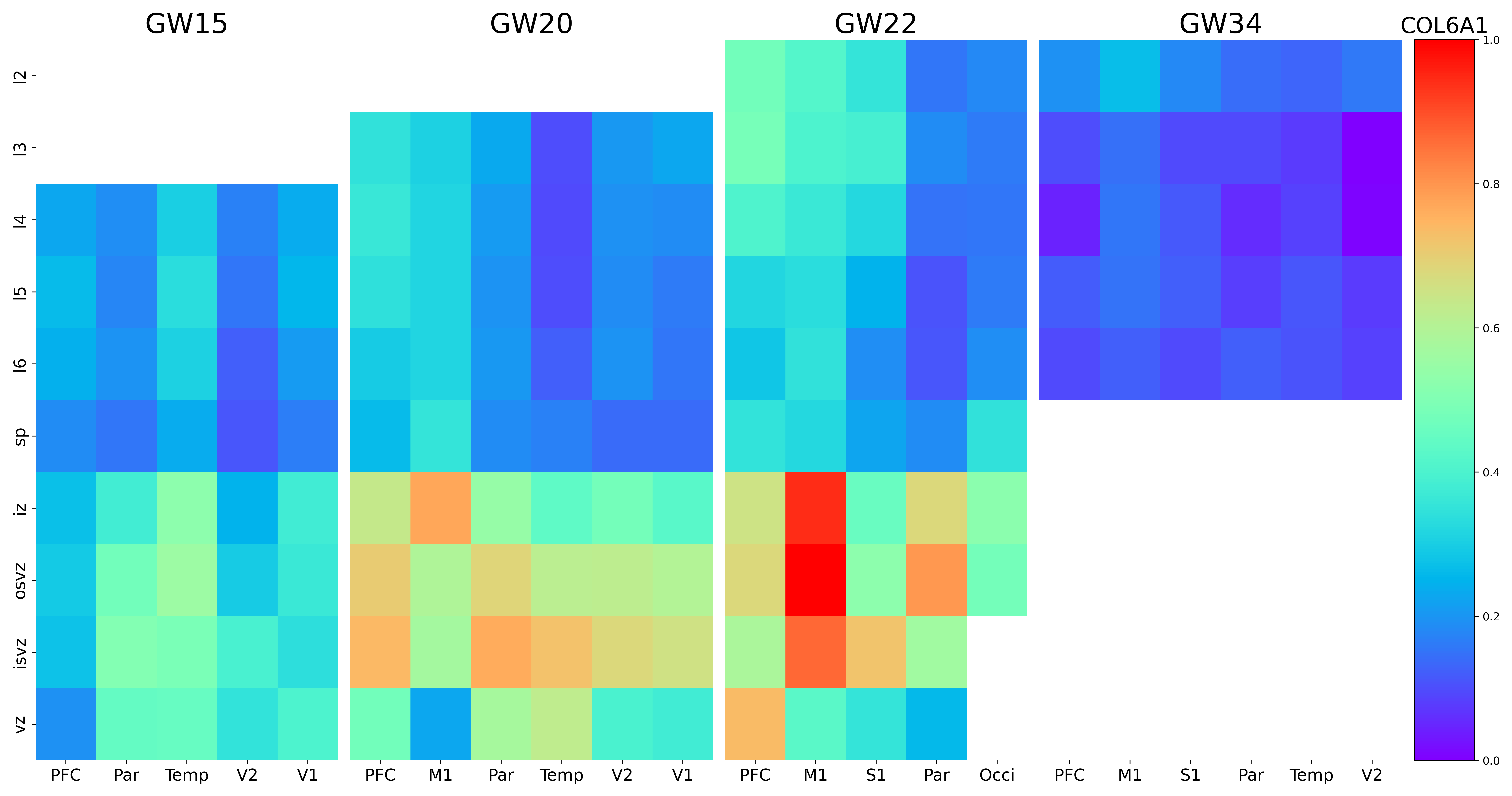

### COL19A1.png

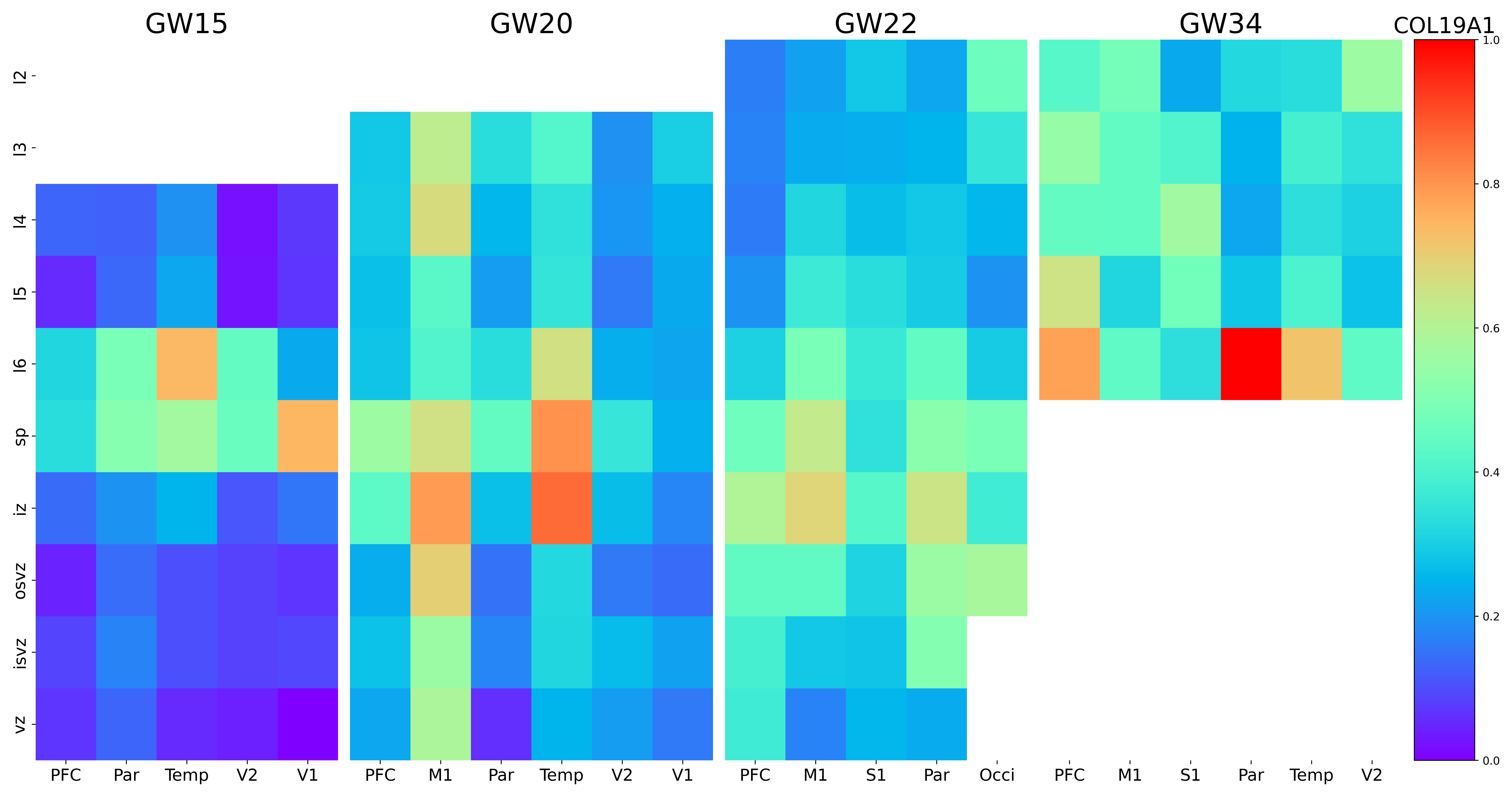

### COL24A1.png

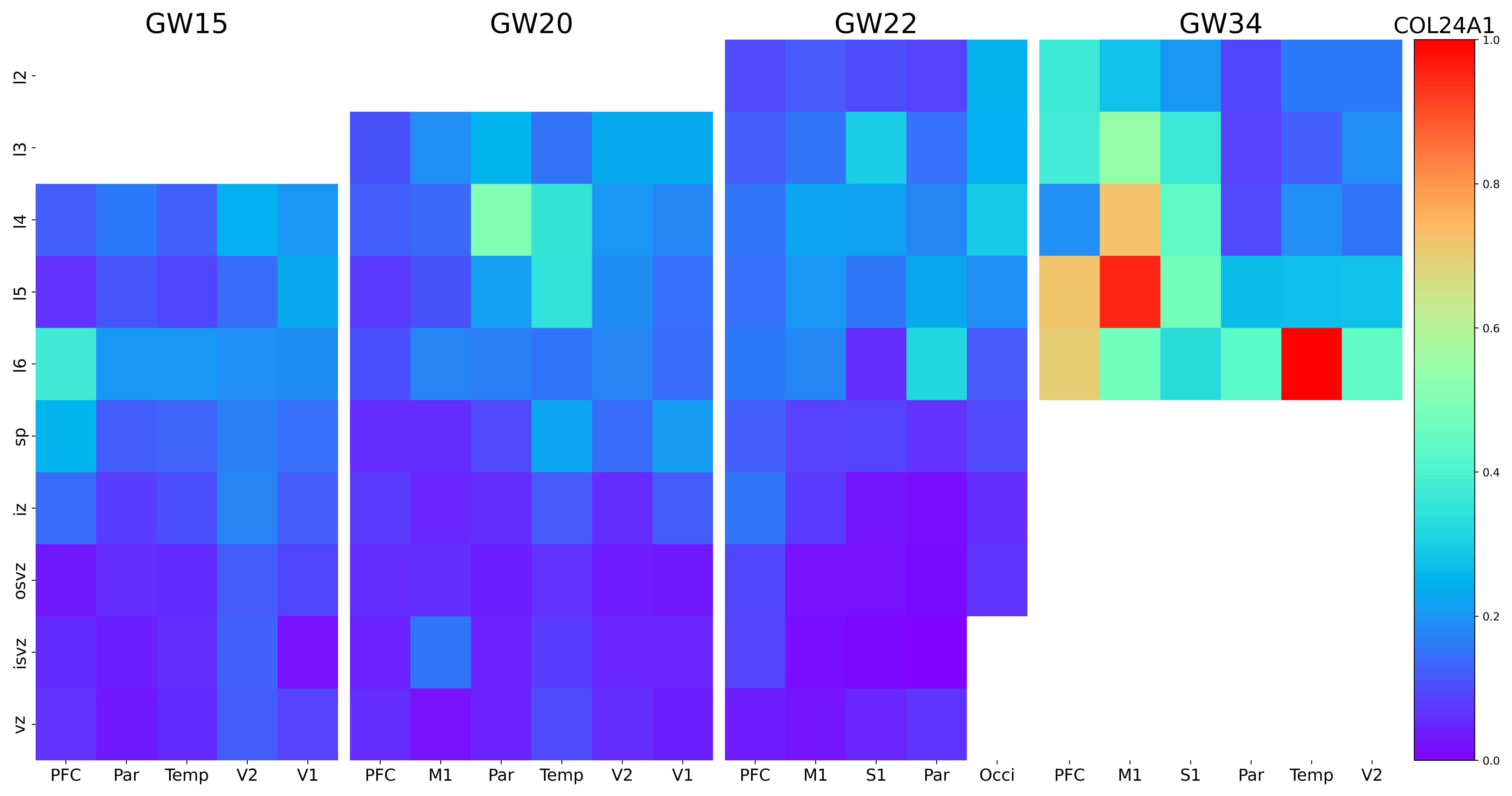

### COL25A1.png

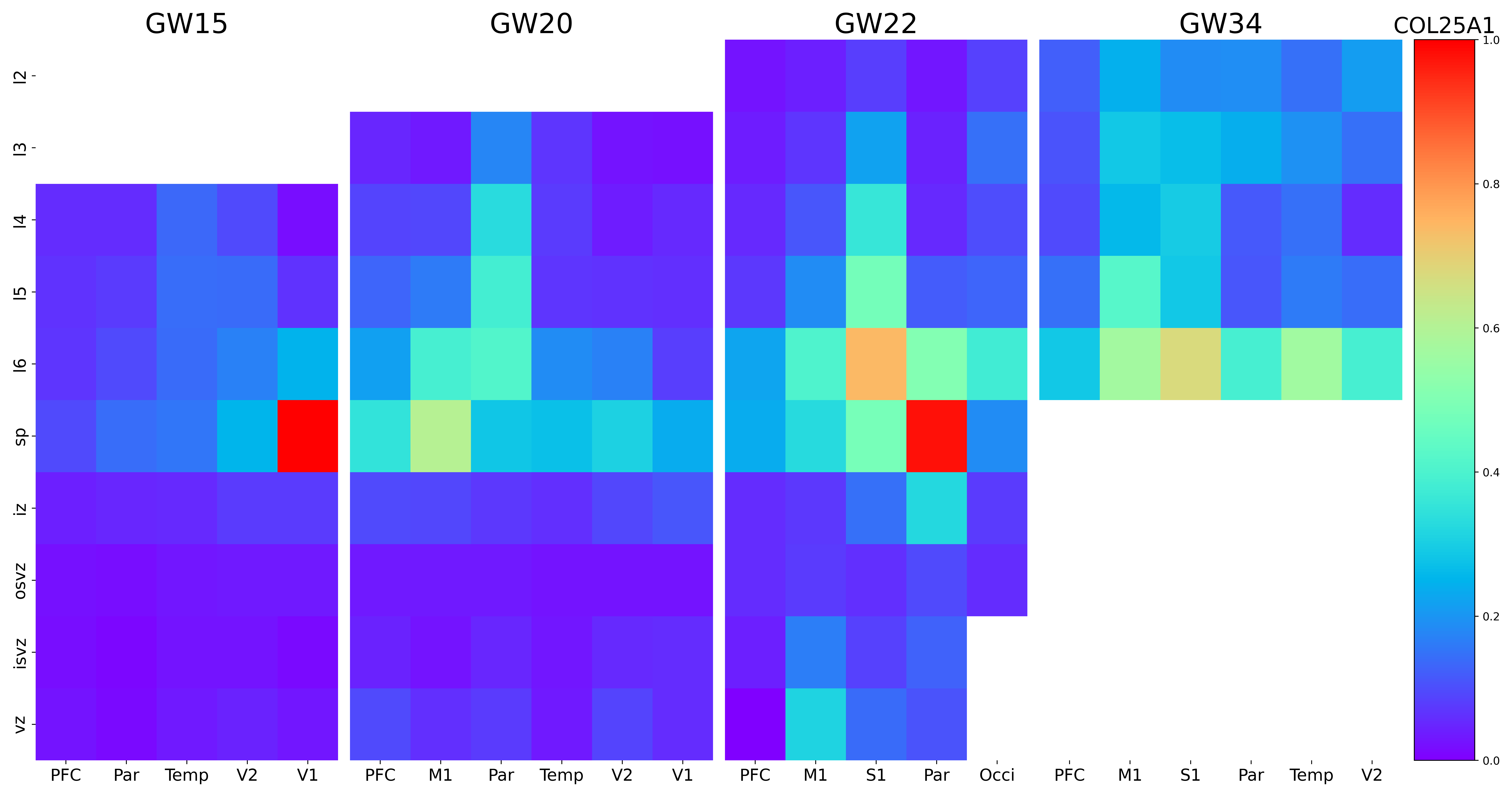

### CPEB2.png

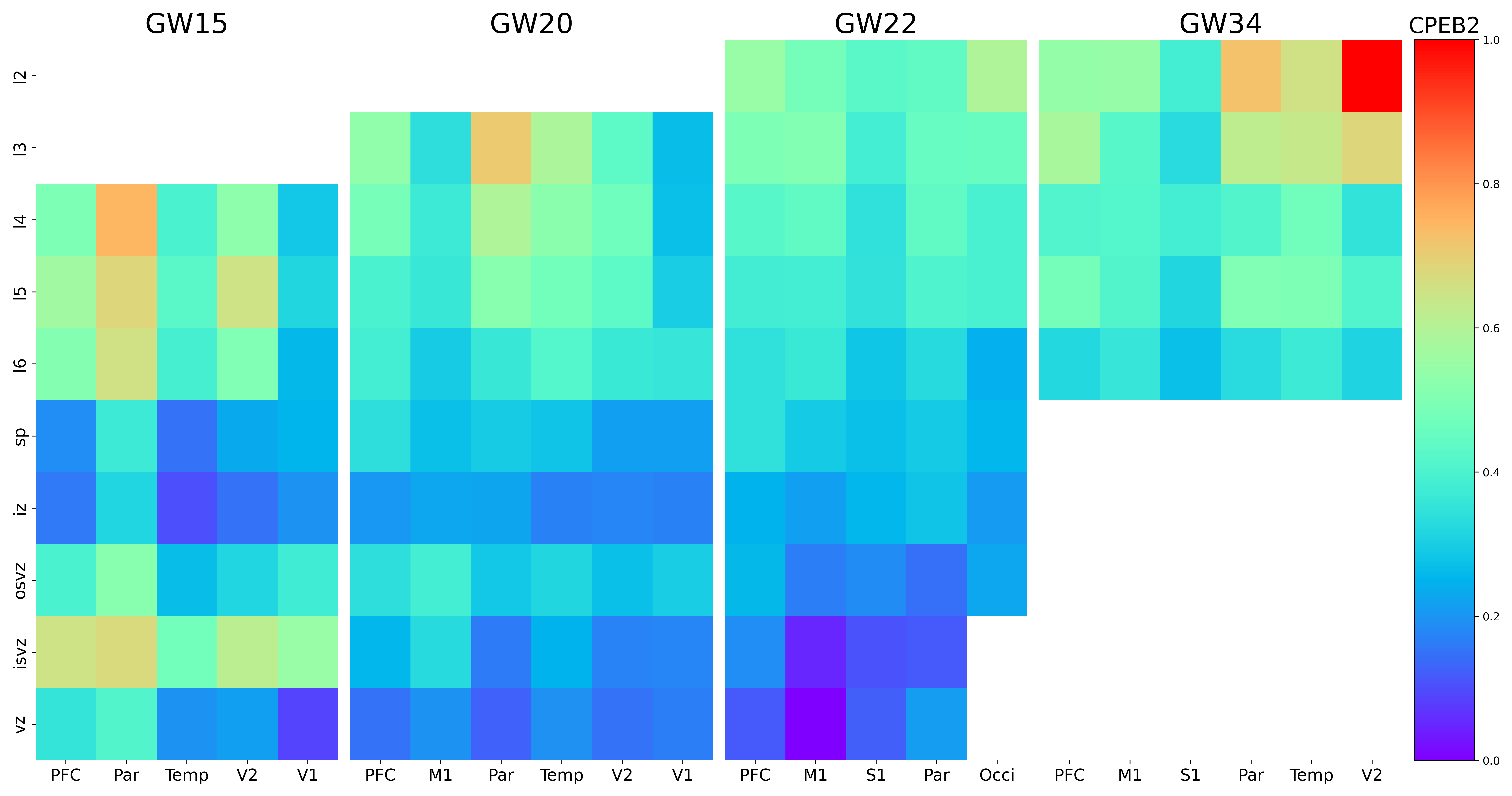

### CPNE8.png

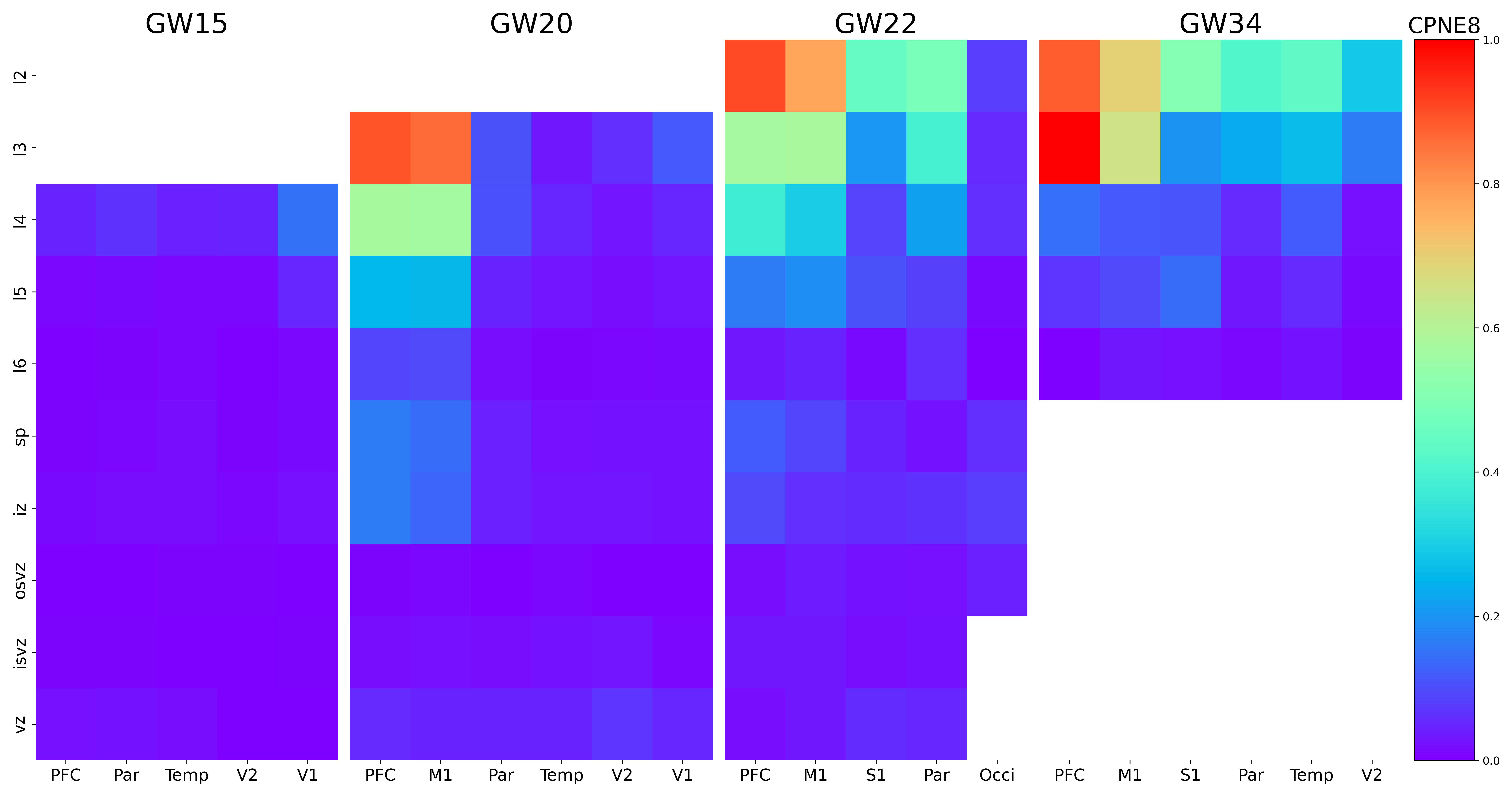

### CRHR2.png

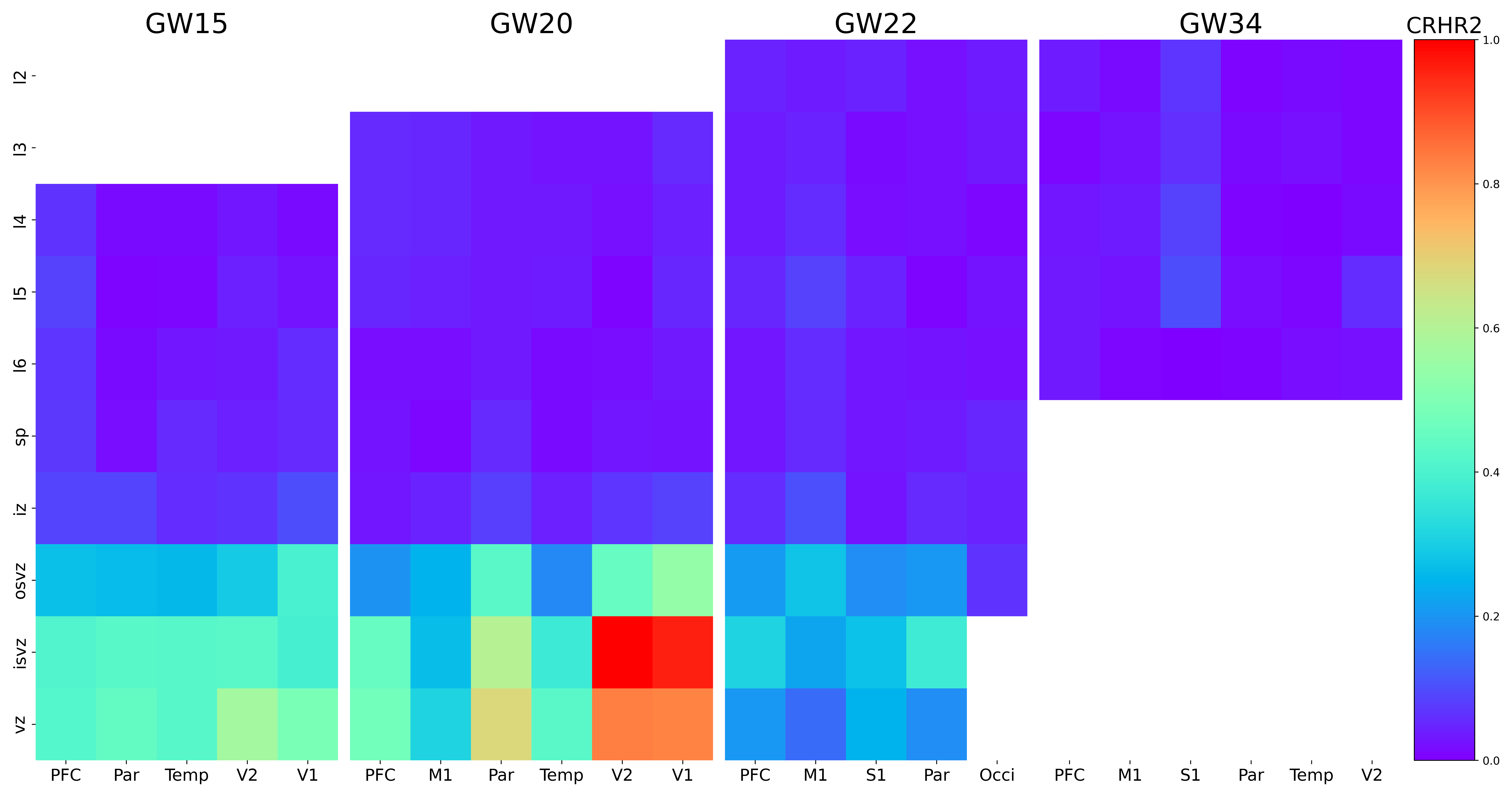

### CRYAB.png

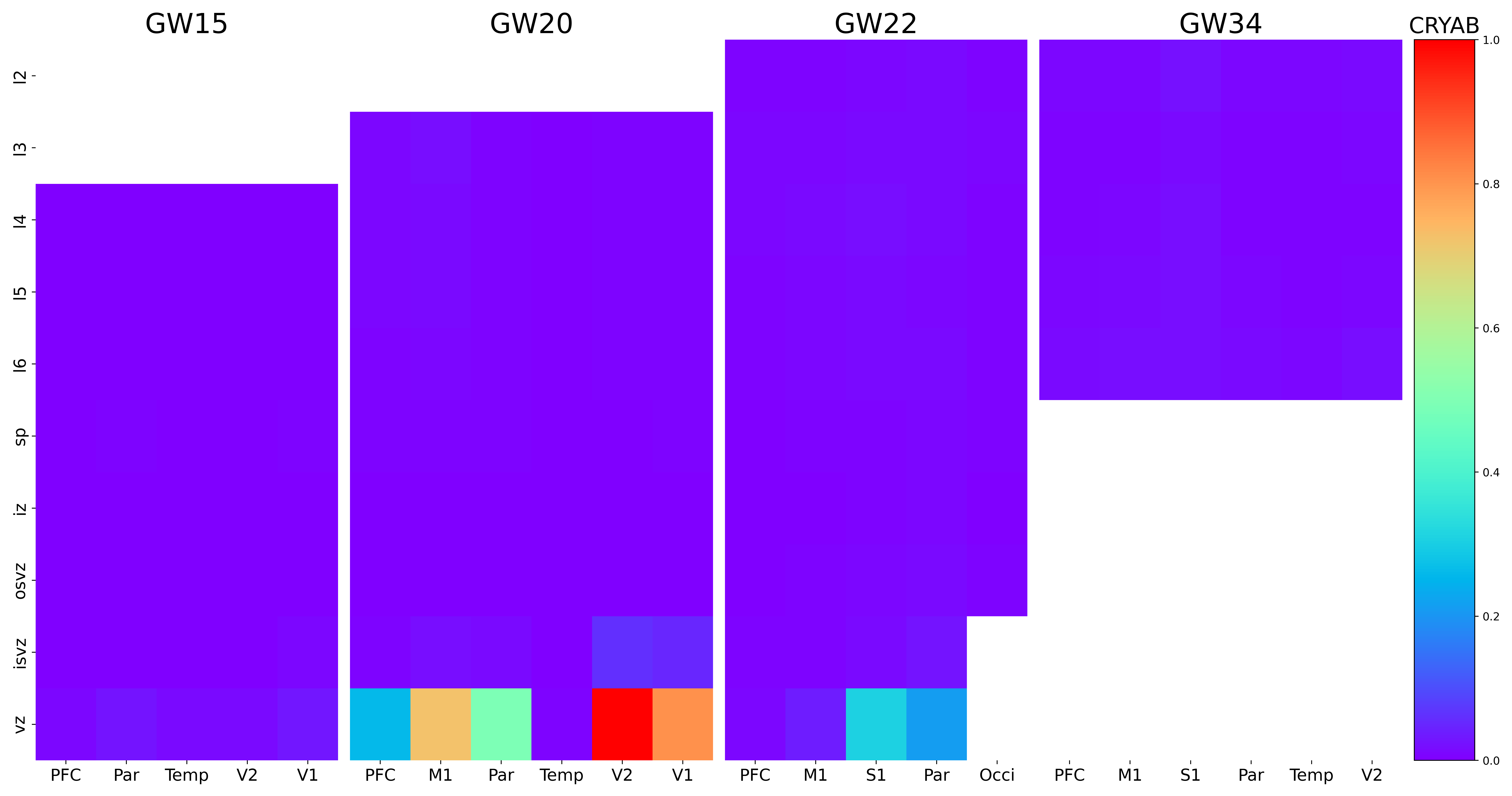

### CRYM.png

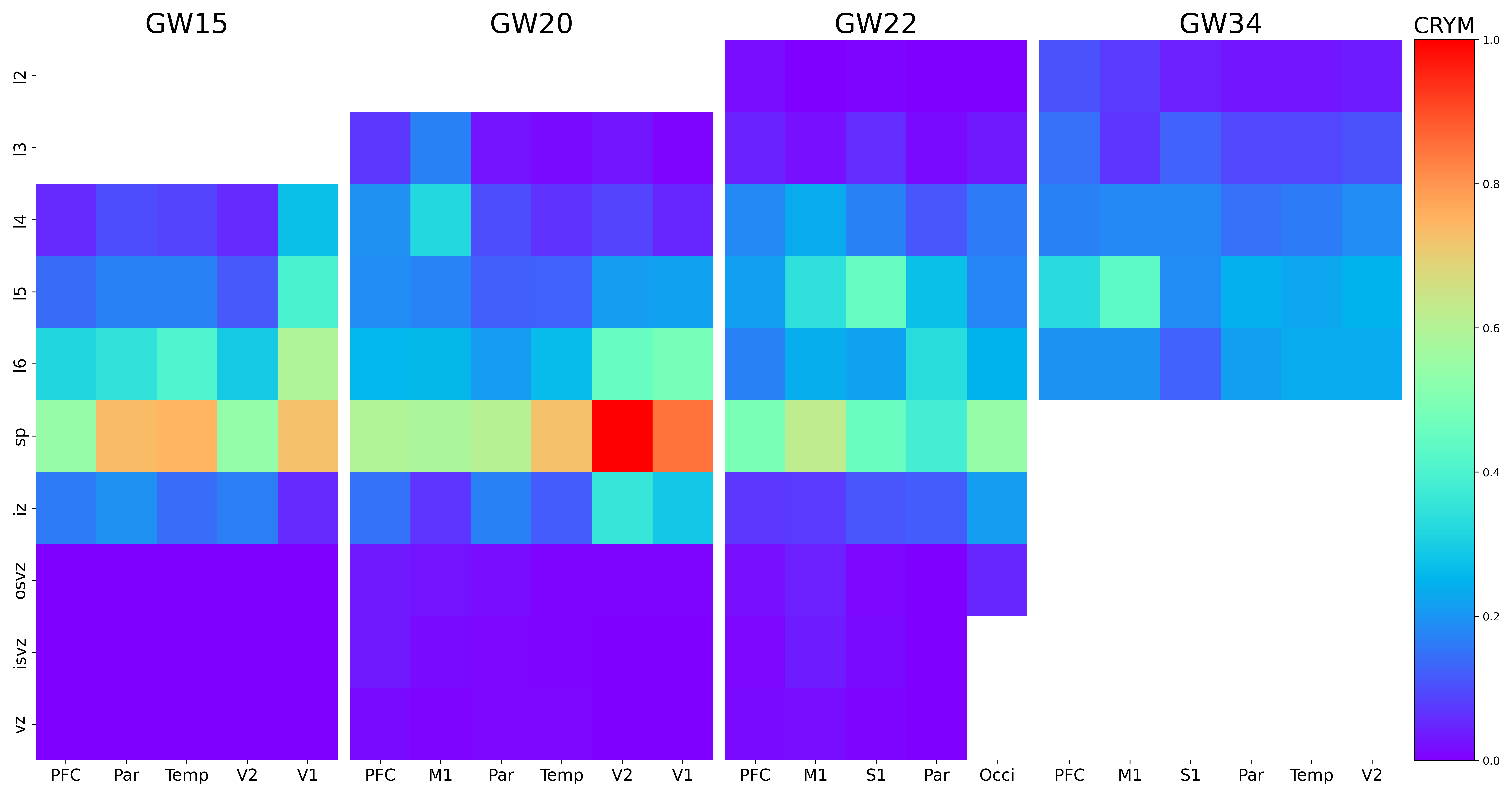

### CSPG5.png

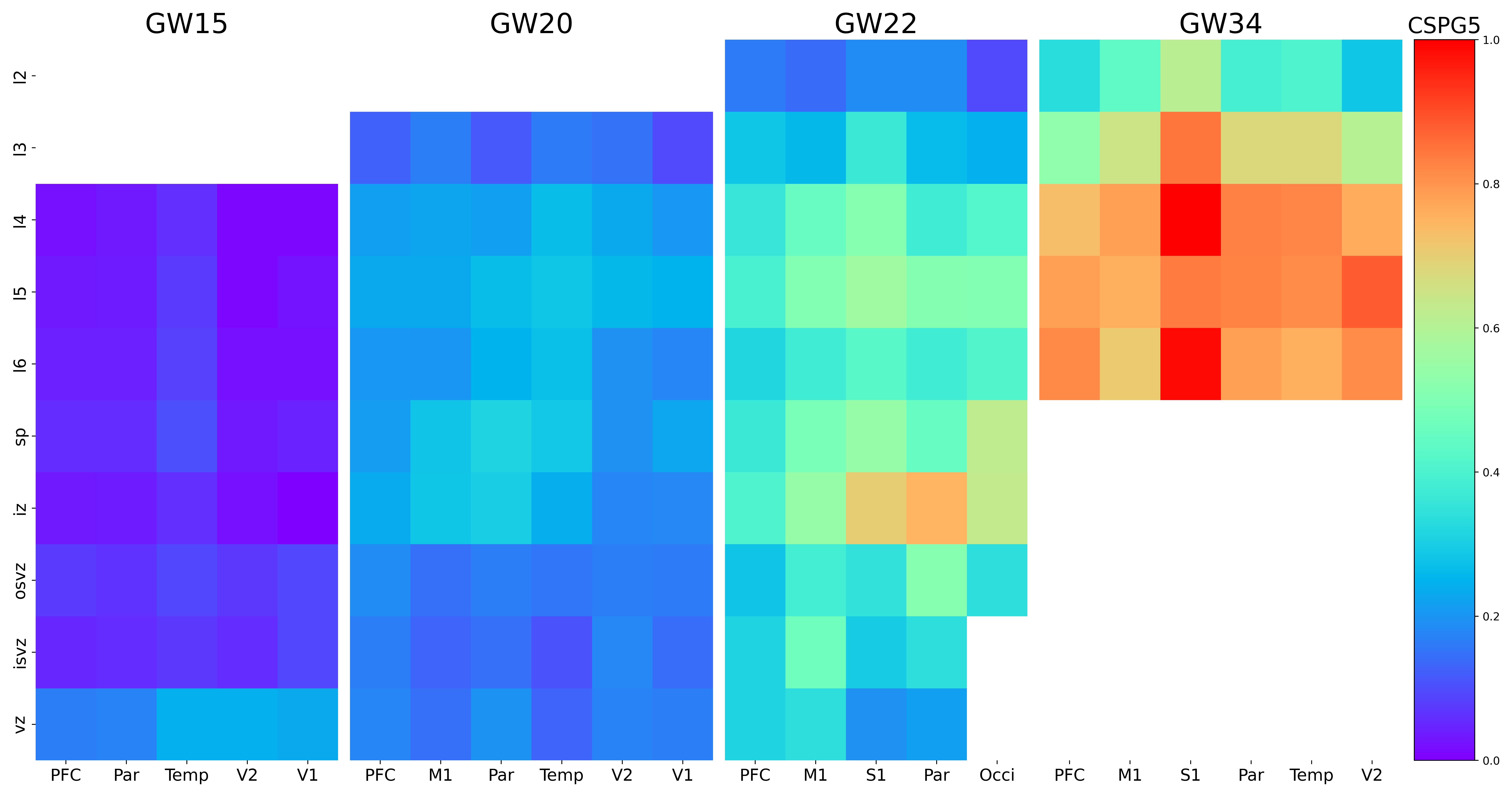

### CST3.png

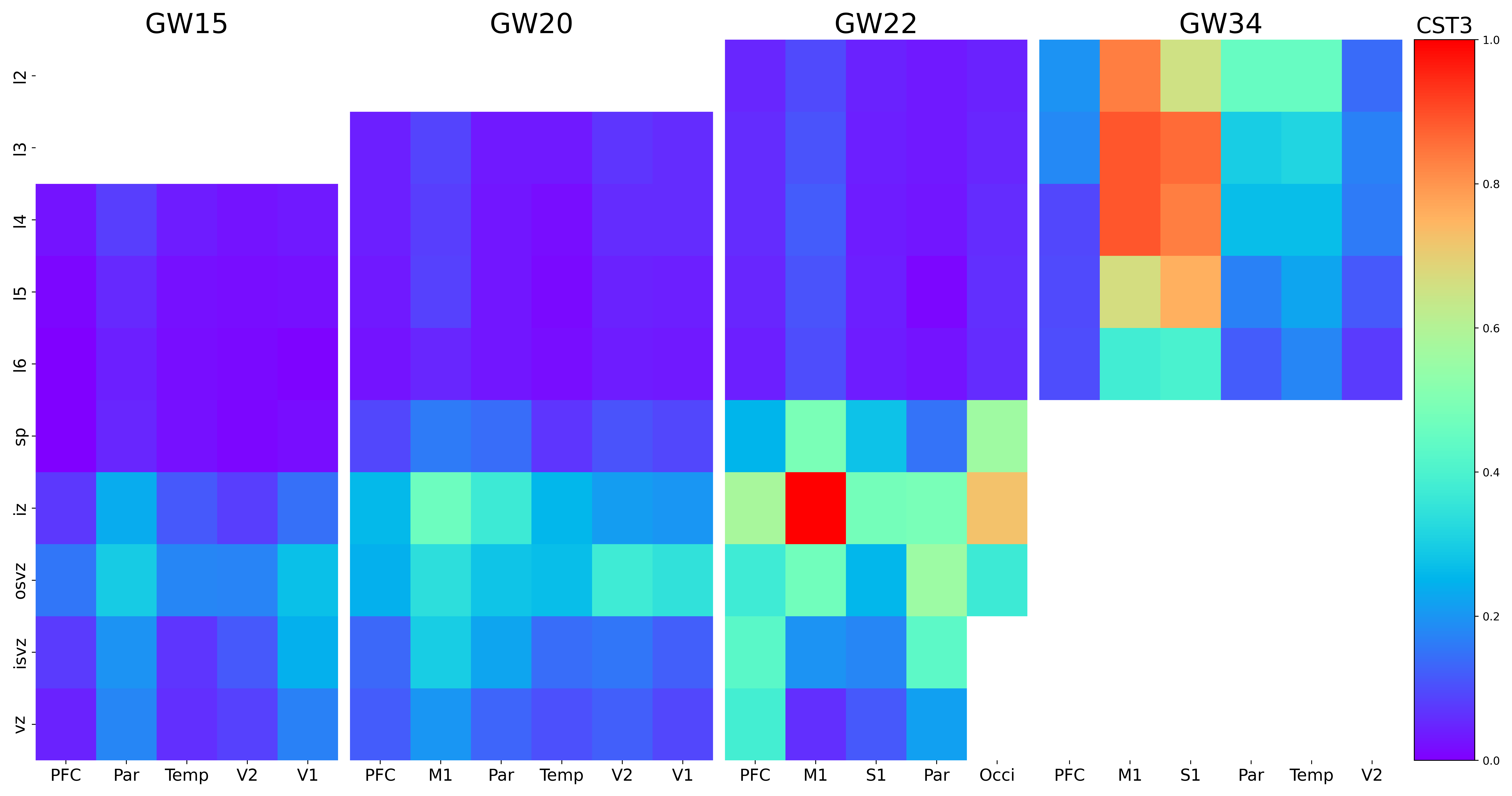

### CUX1.png

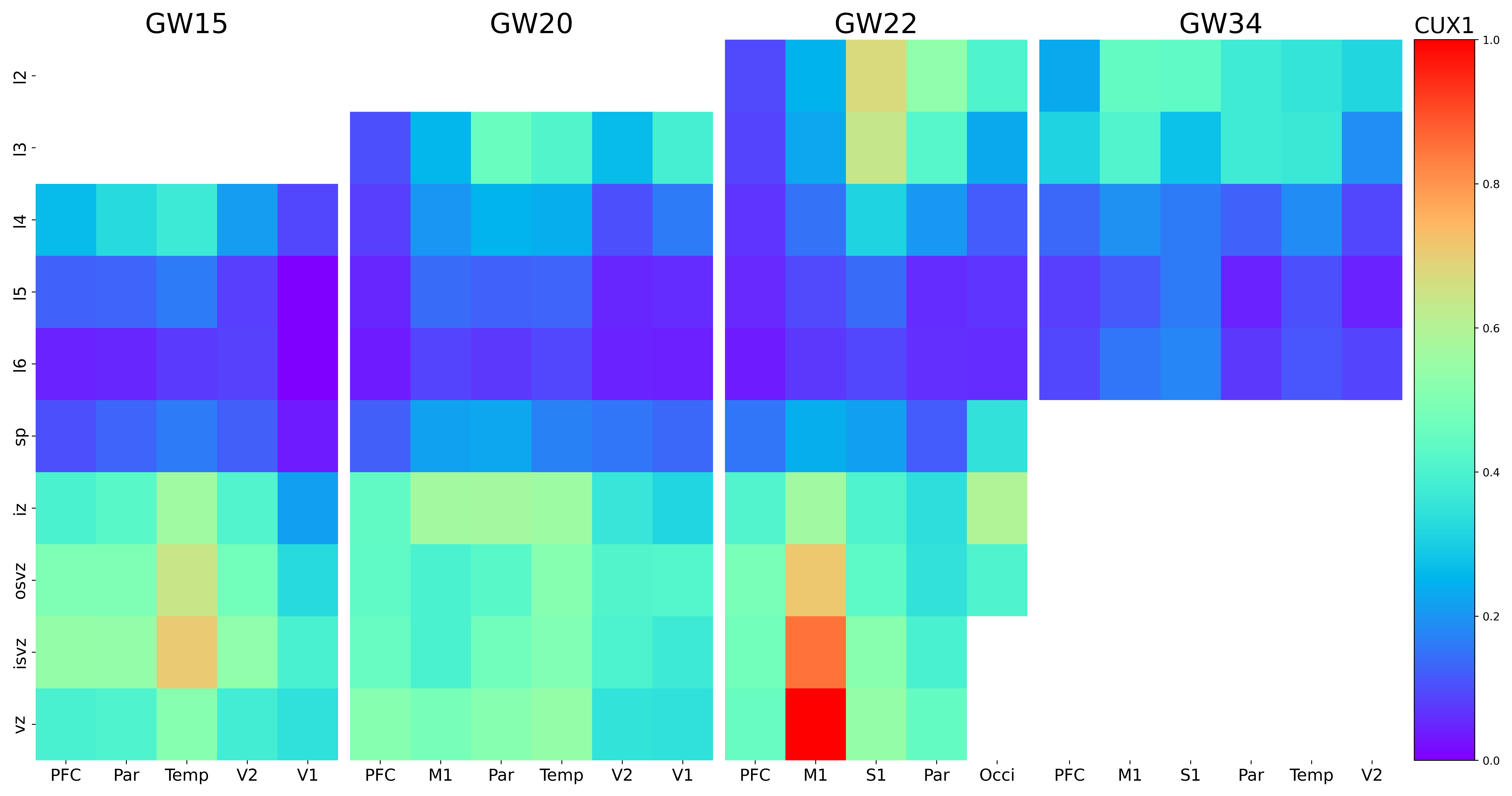

### CUX2.png

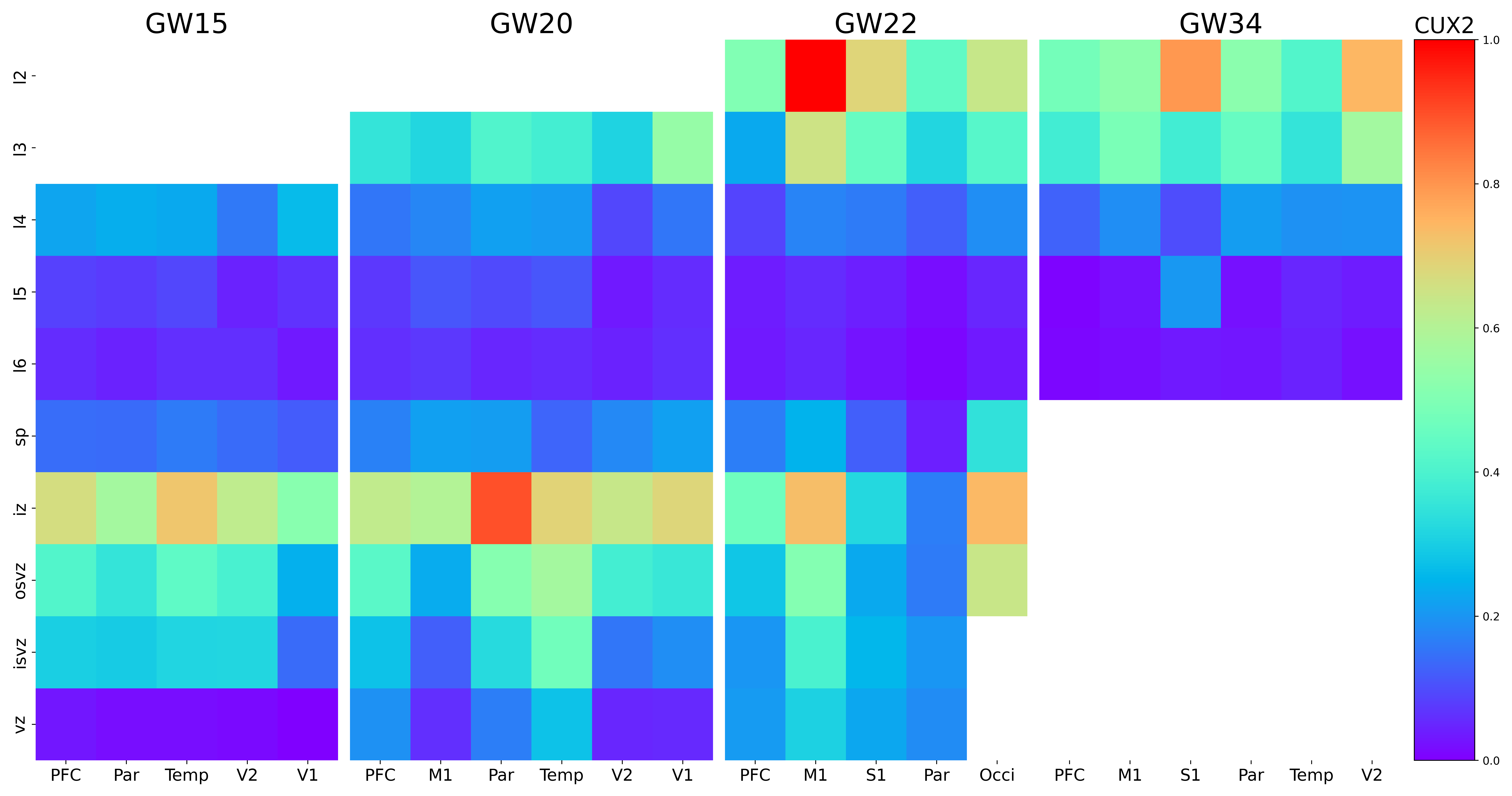

### CXCL14.png

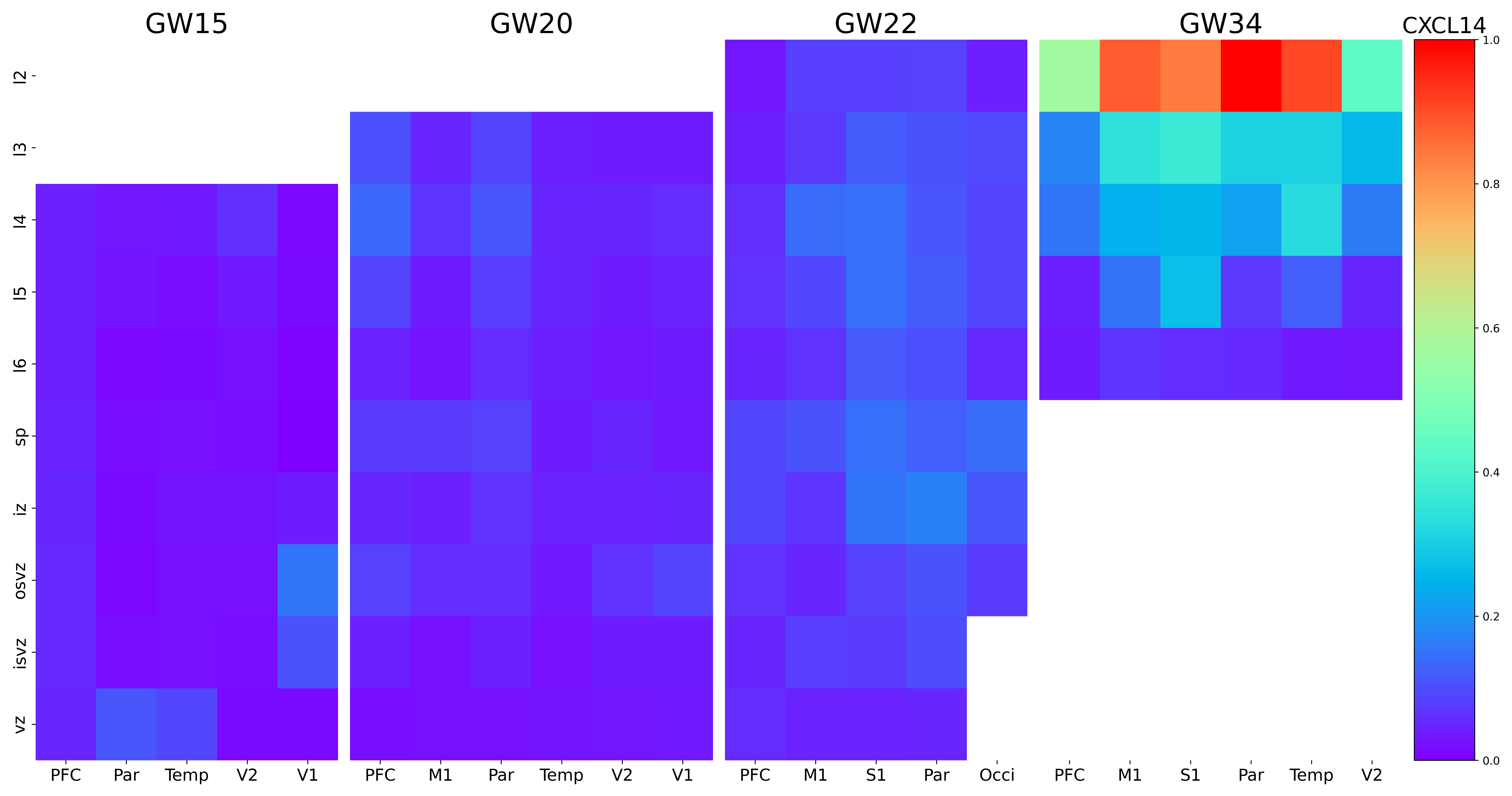

### CYP26A1.png

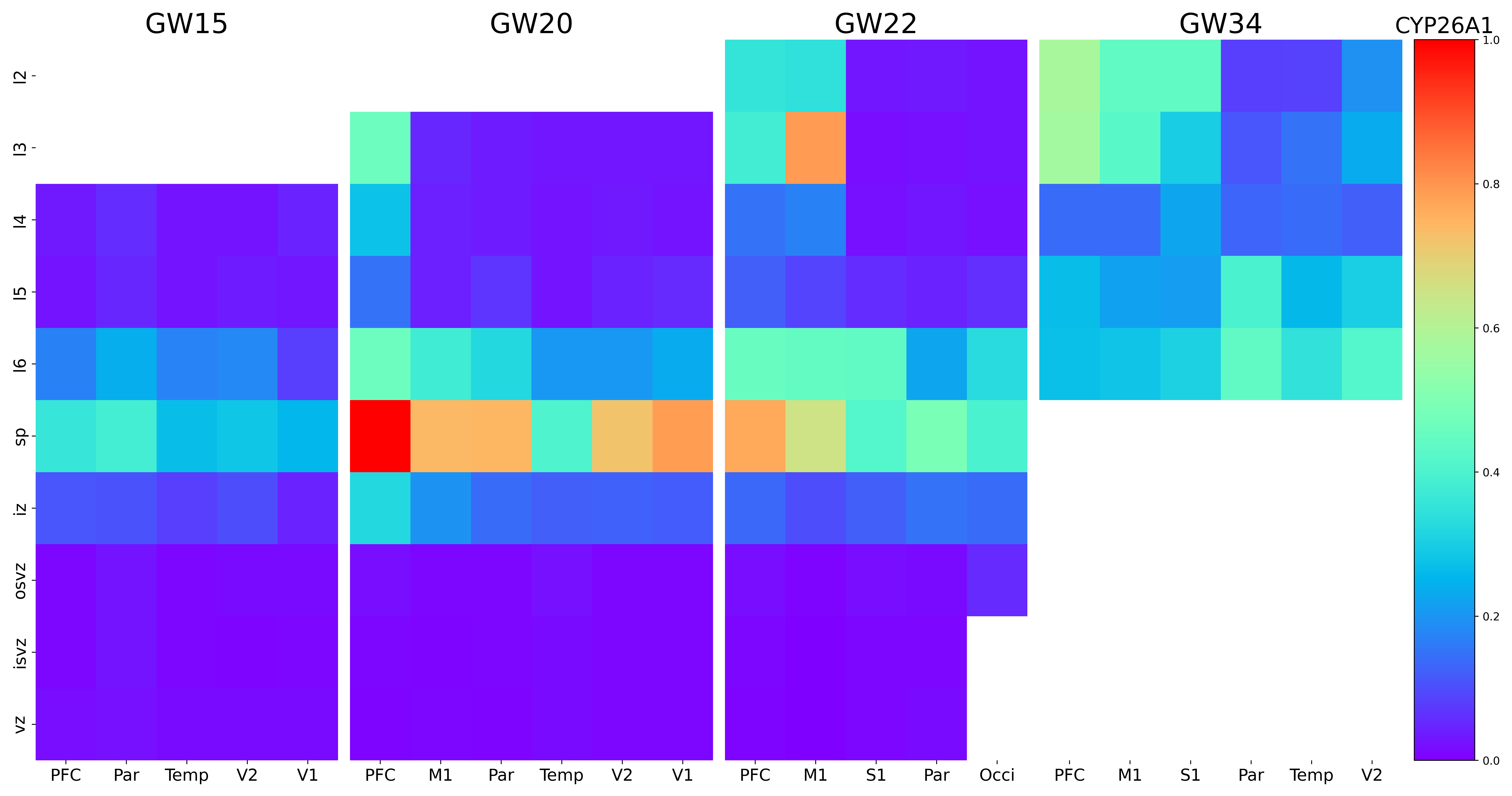

### DACH2.png

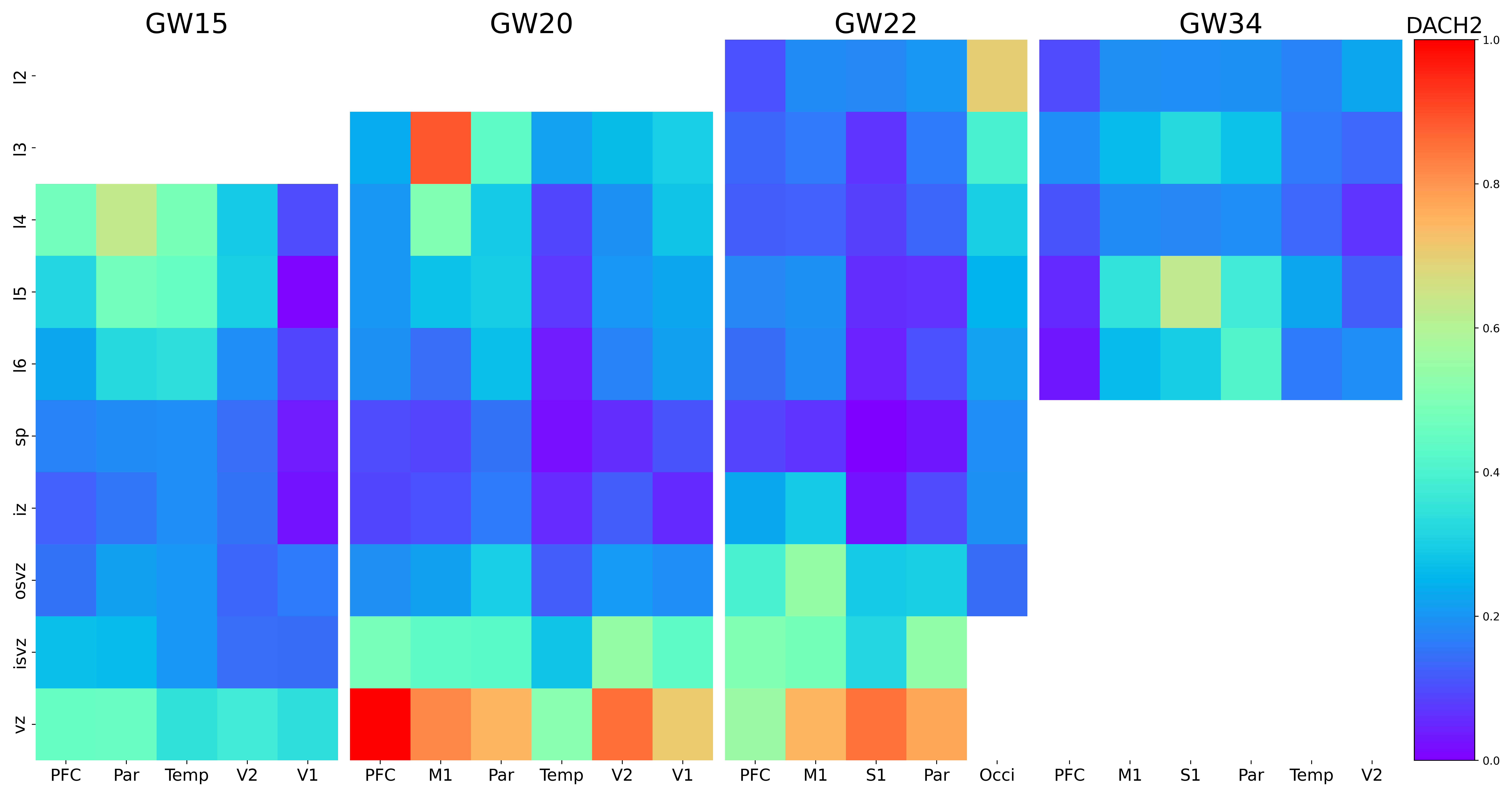

### DENND2B.png

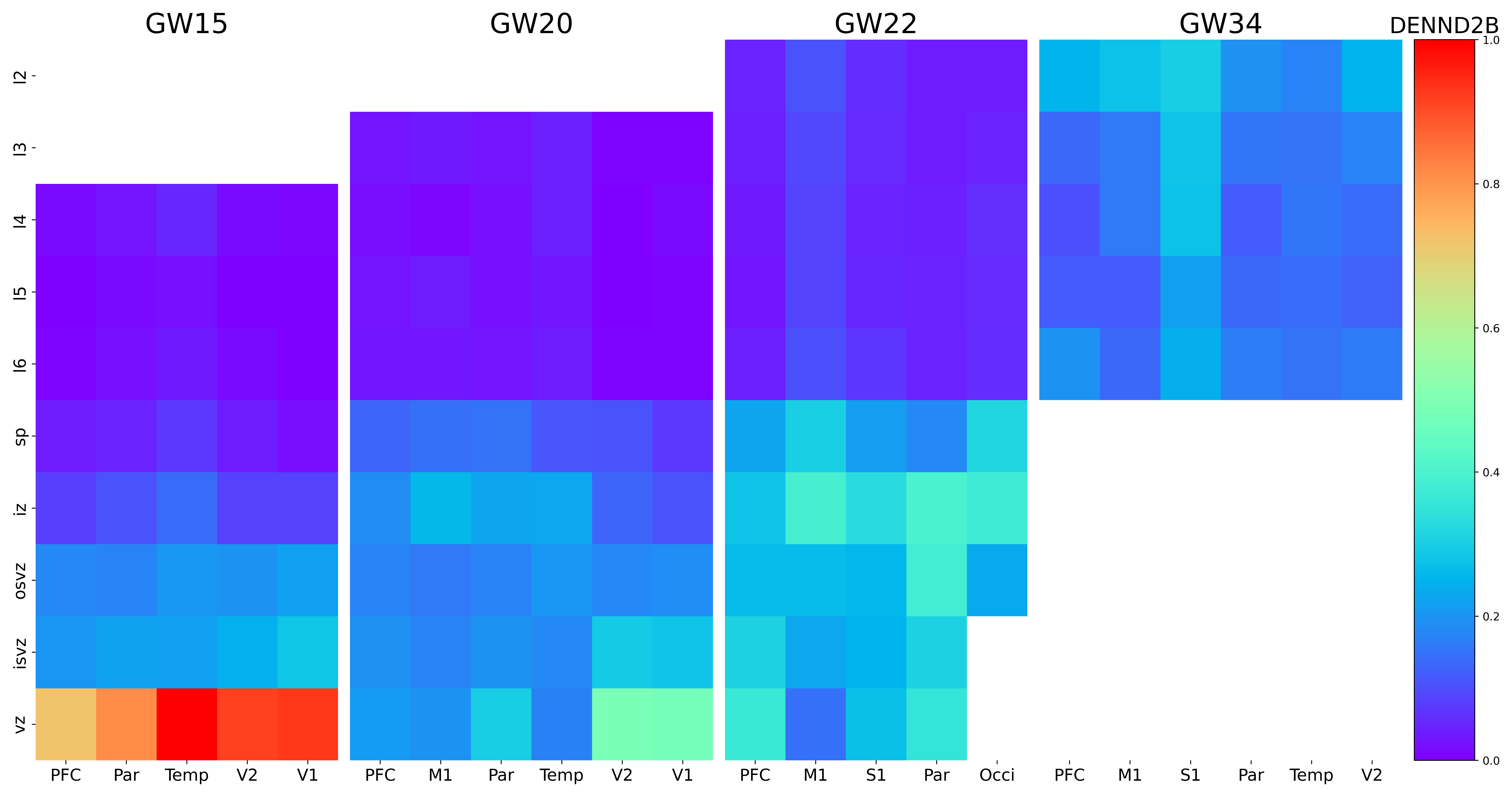

### DIO2.png

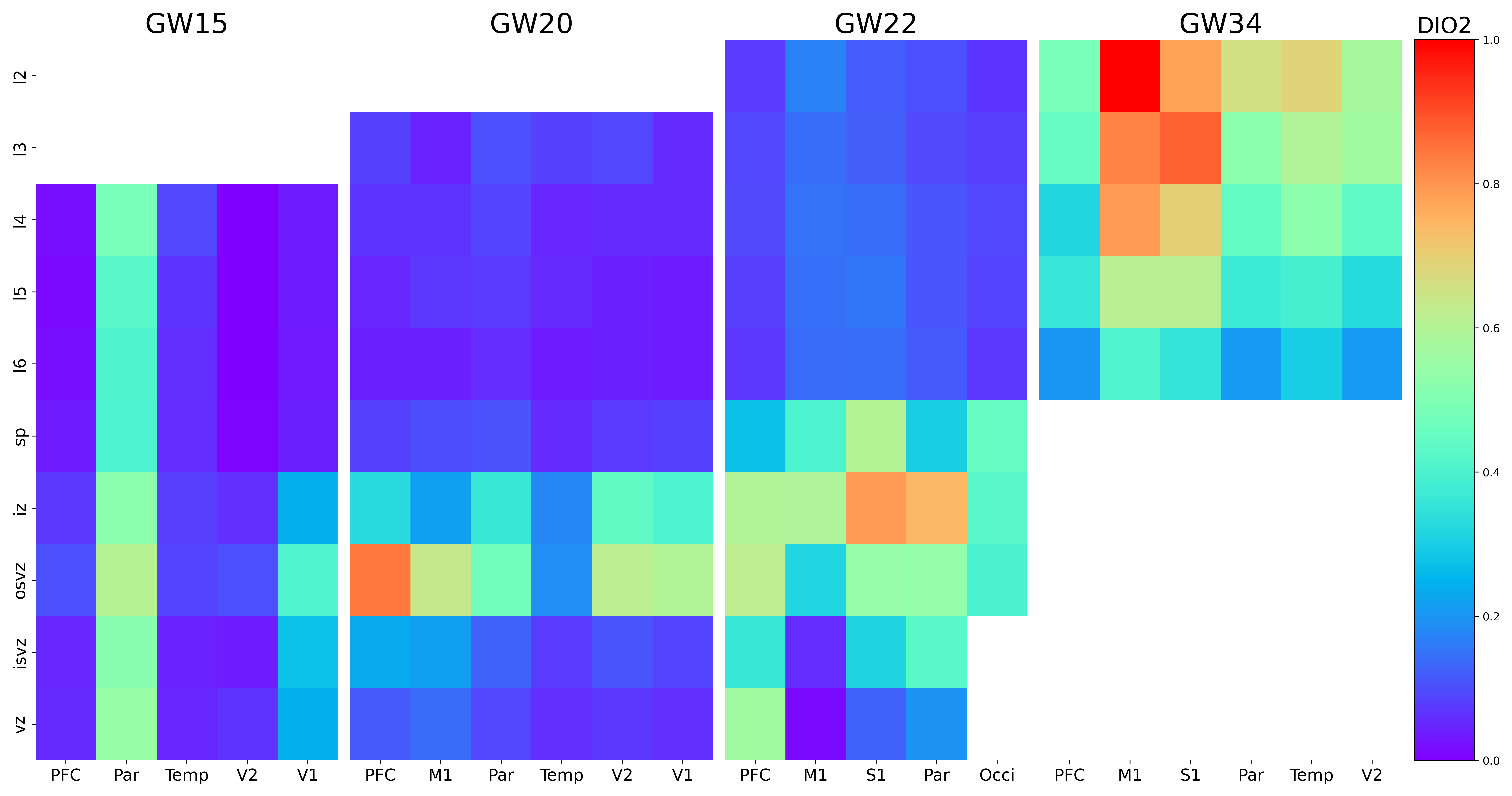

### DISP3.png

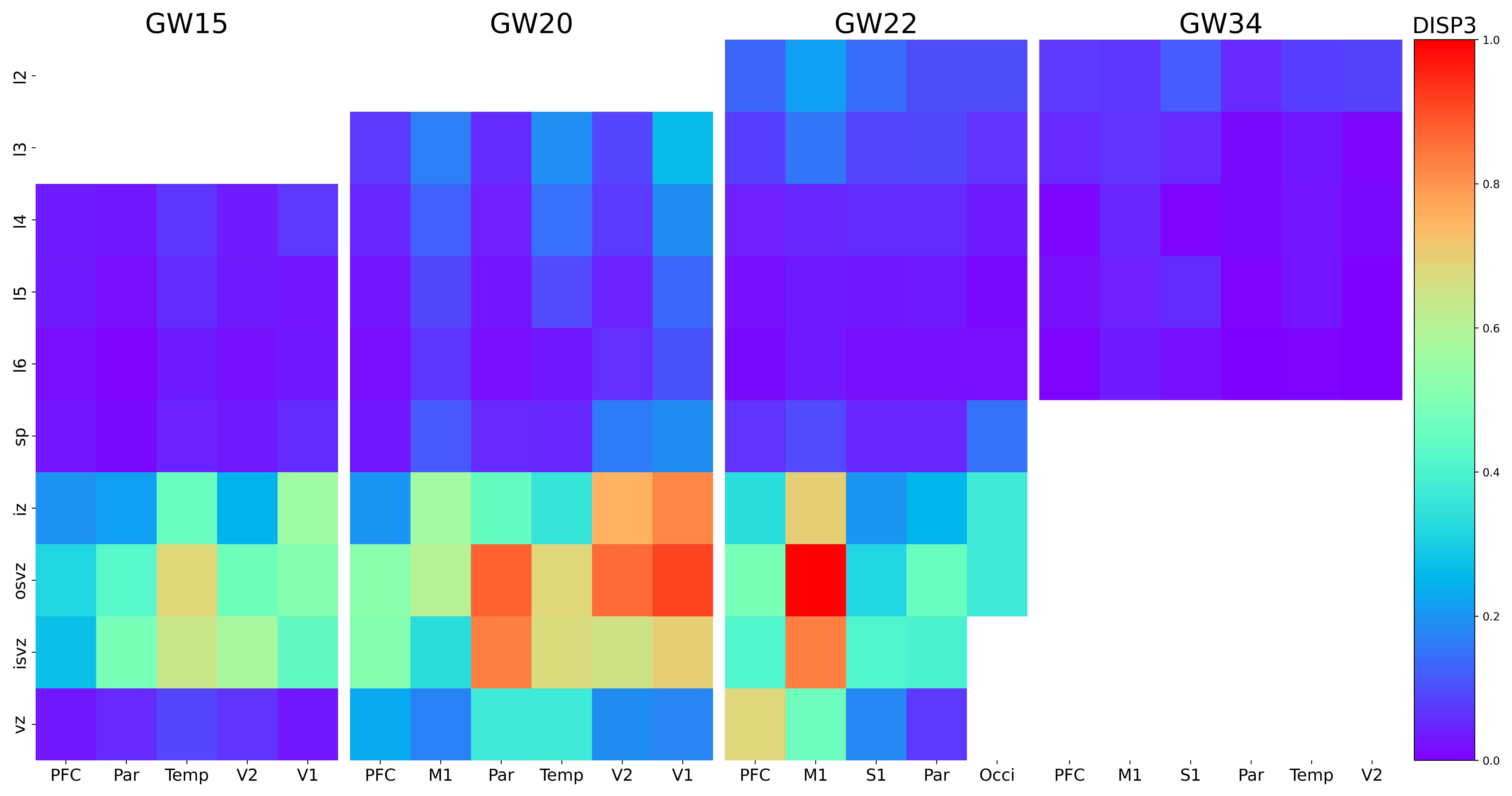

### DNM2.png

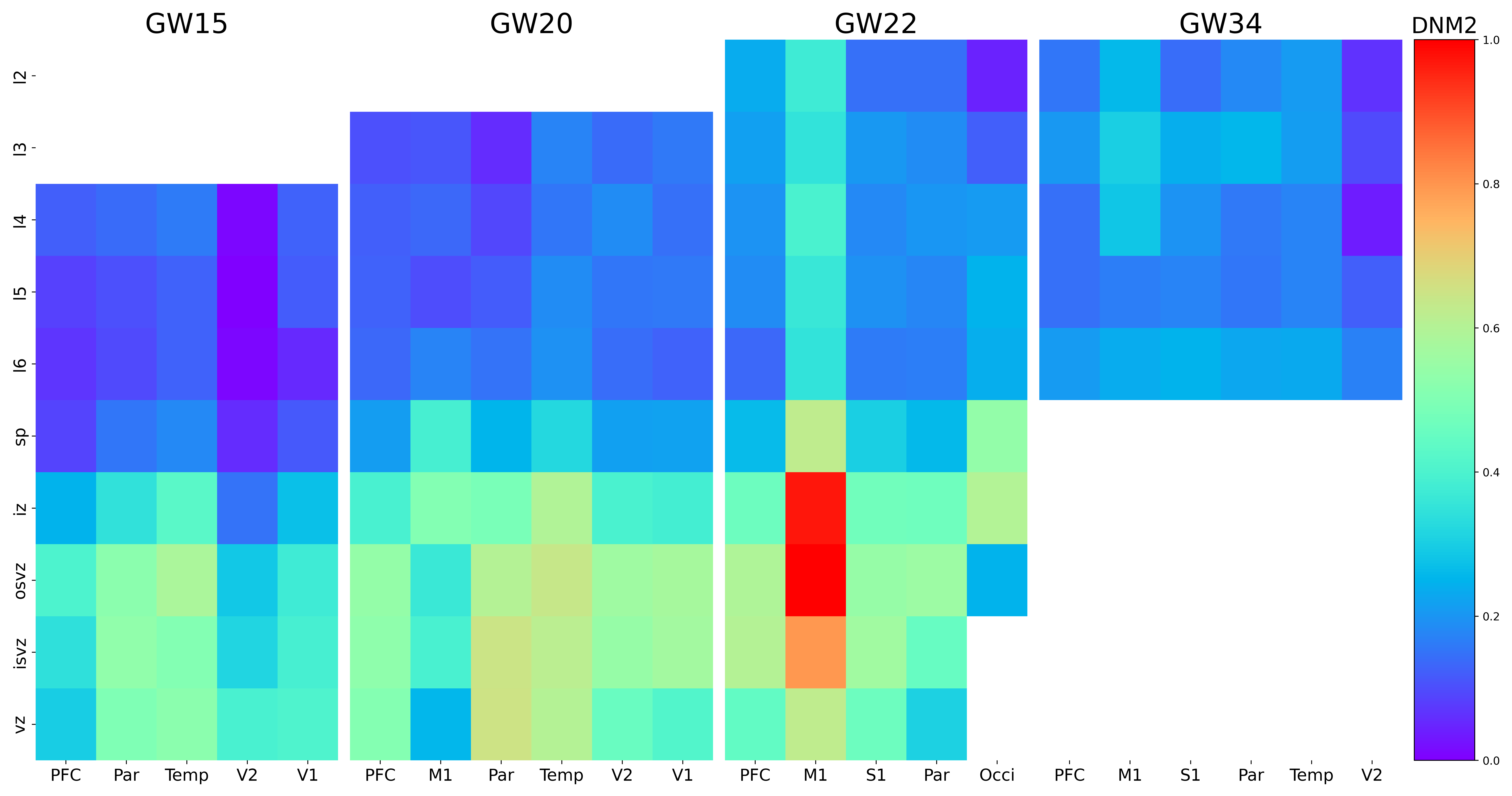

### DPP3.png

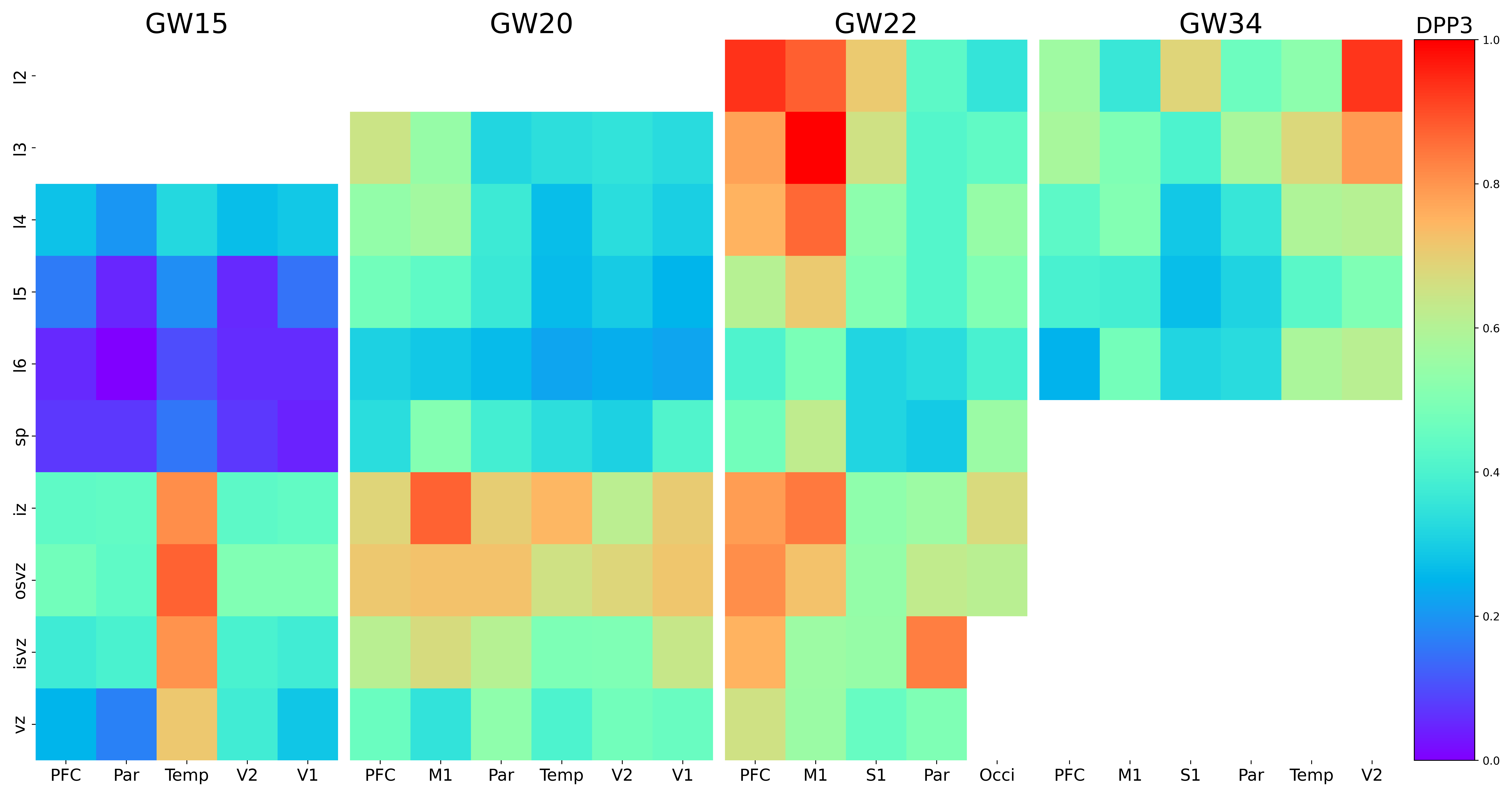

### E2F2.png

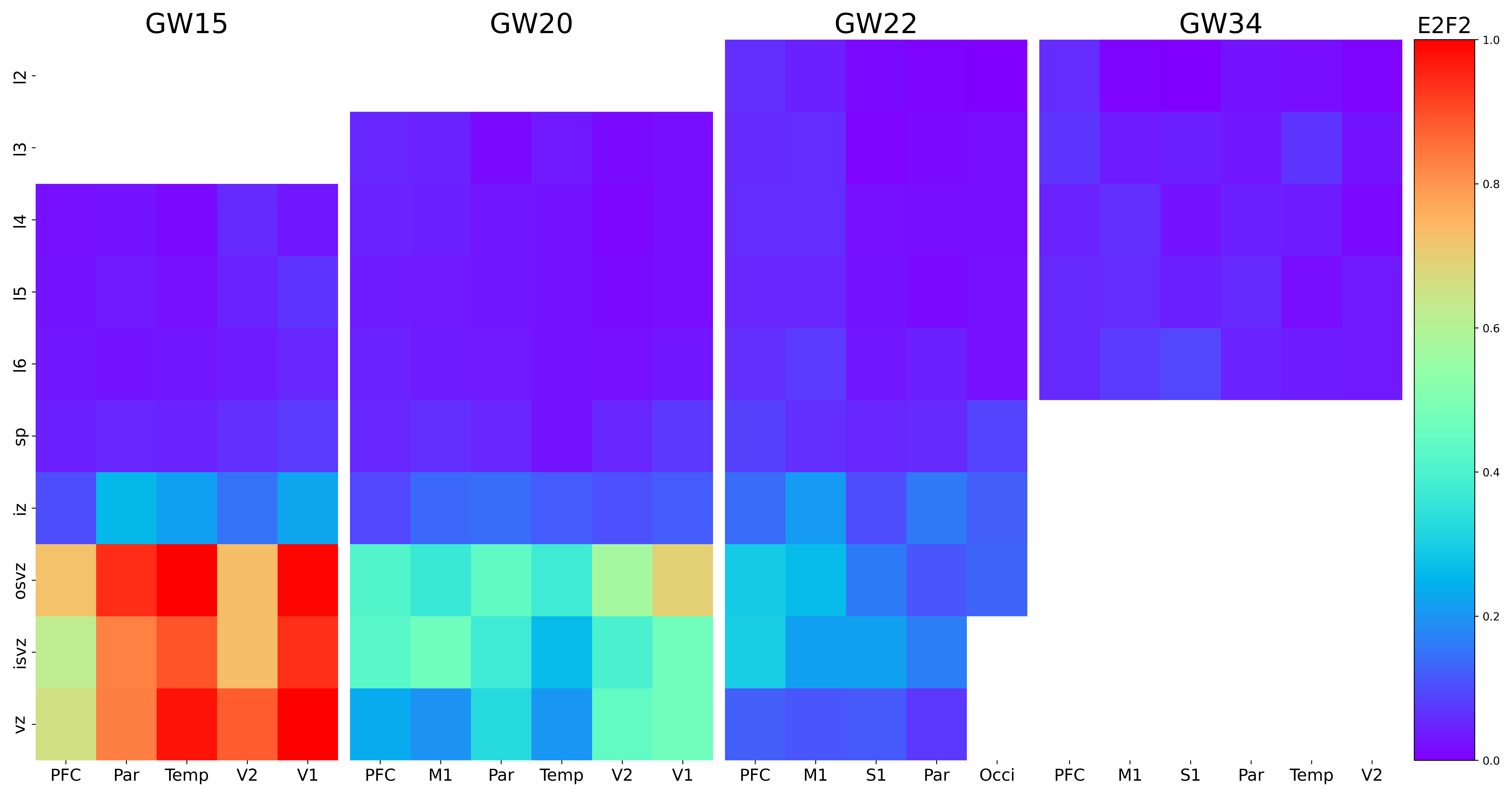

### EEPD1.png

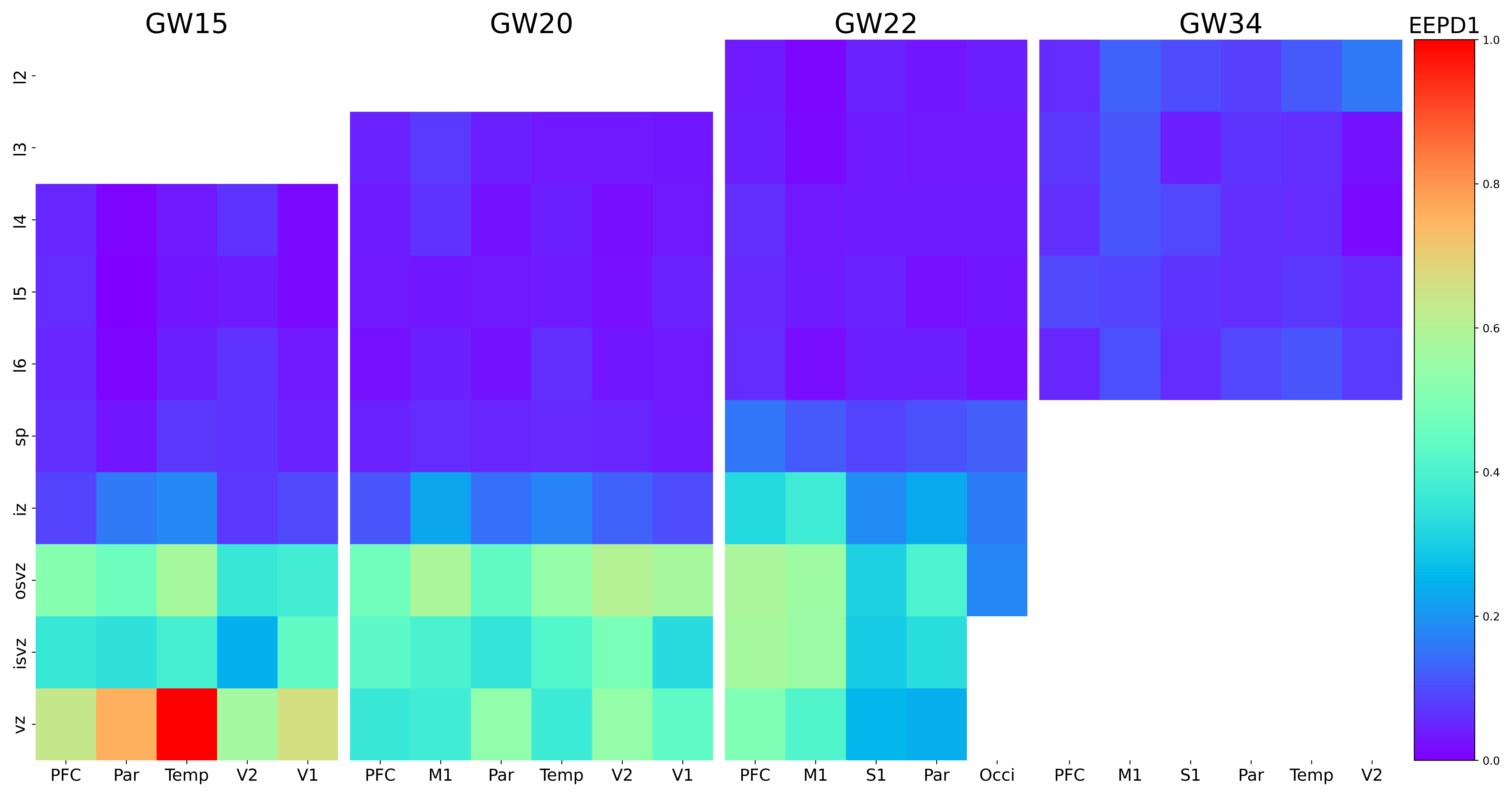

### EGR2.png

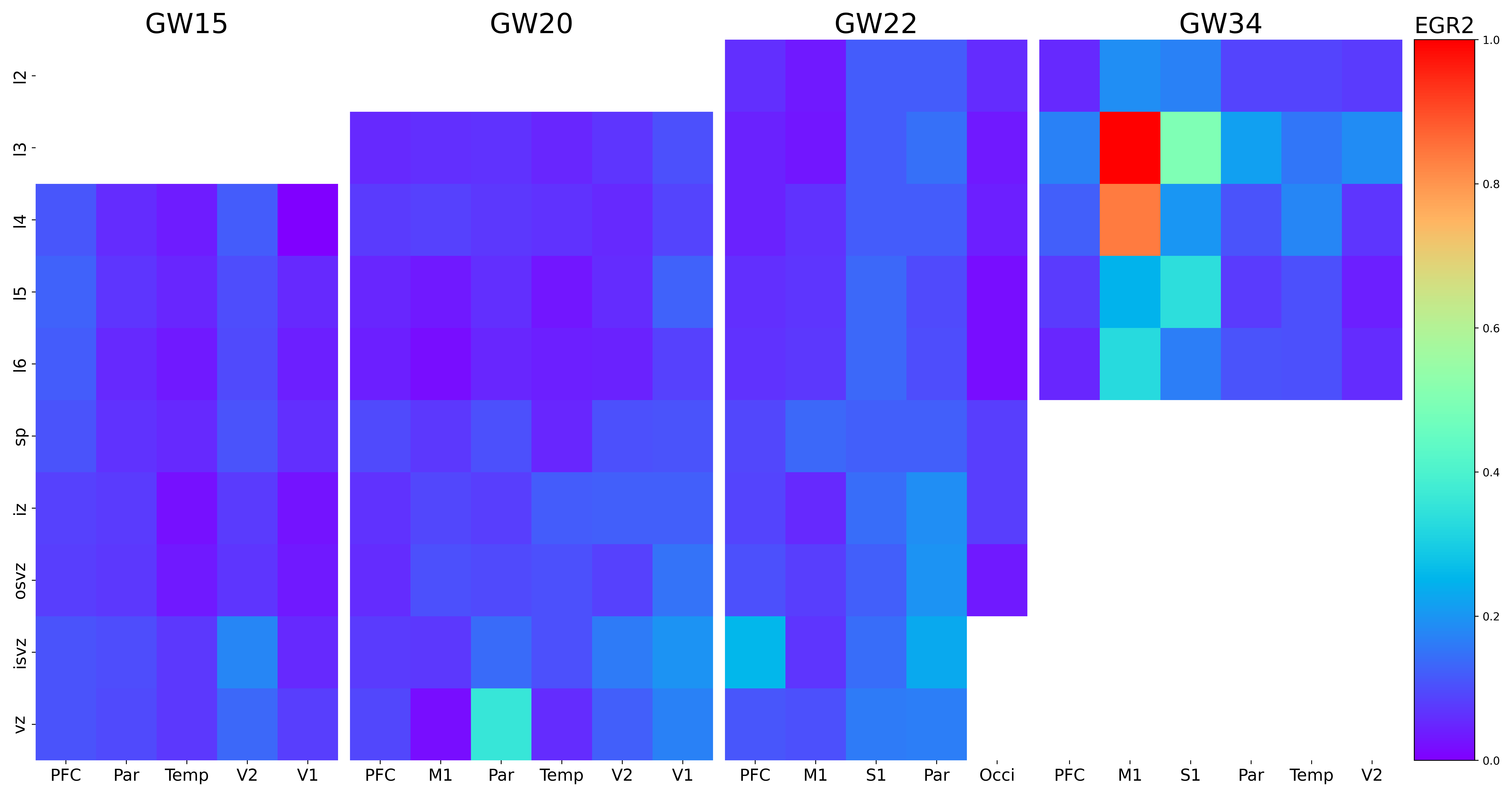

### EHMT1.png

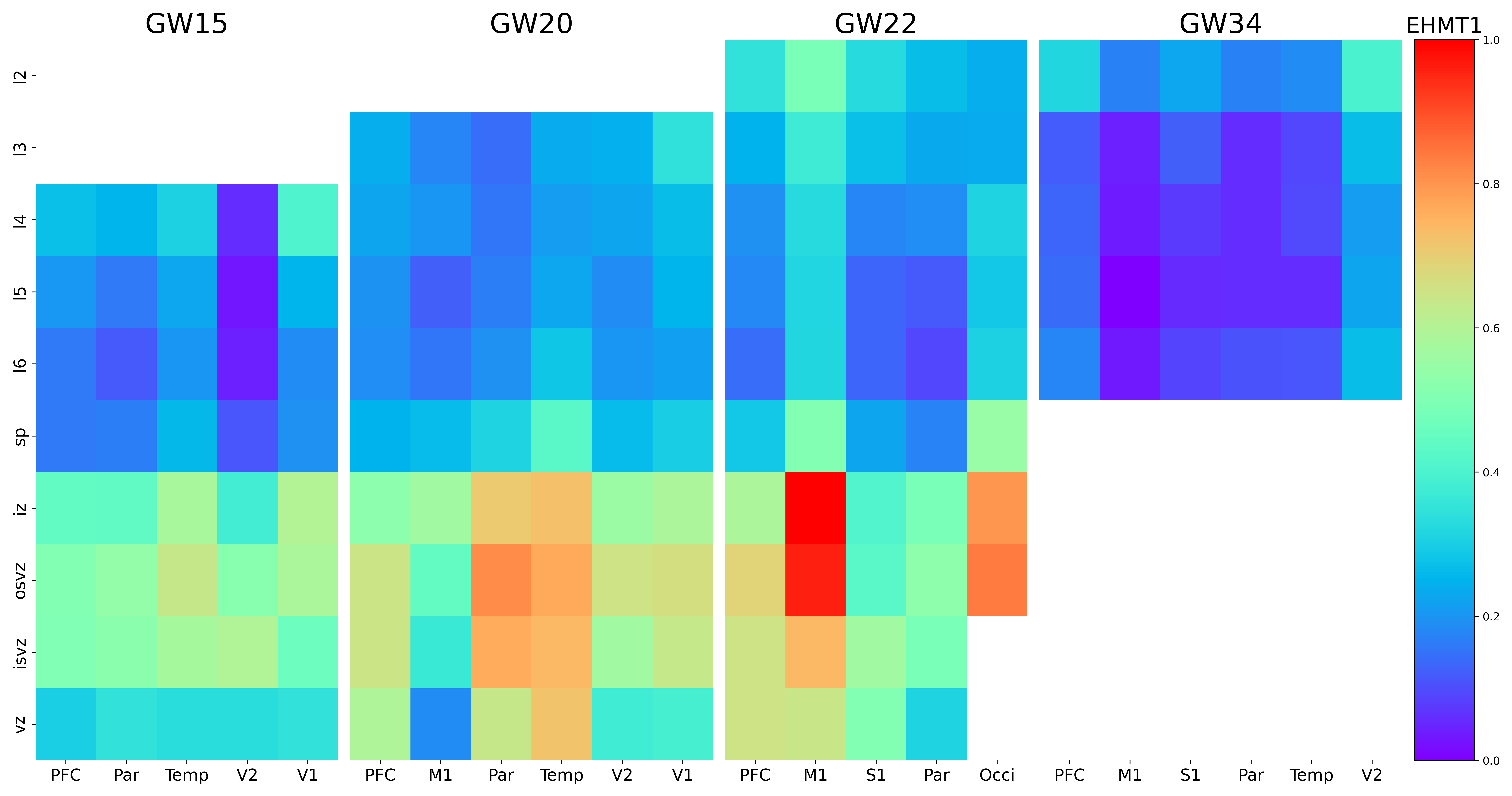

### ELAPOR1.png

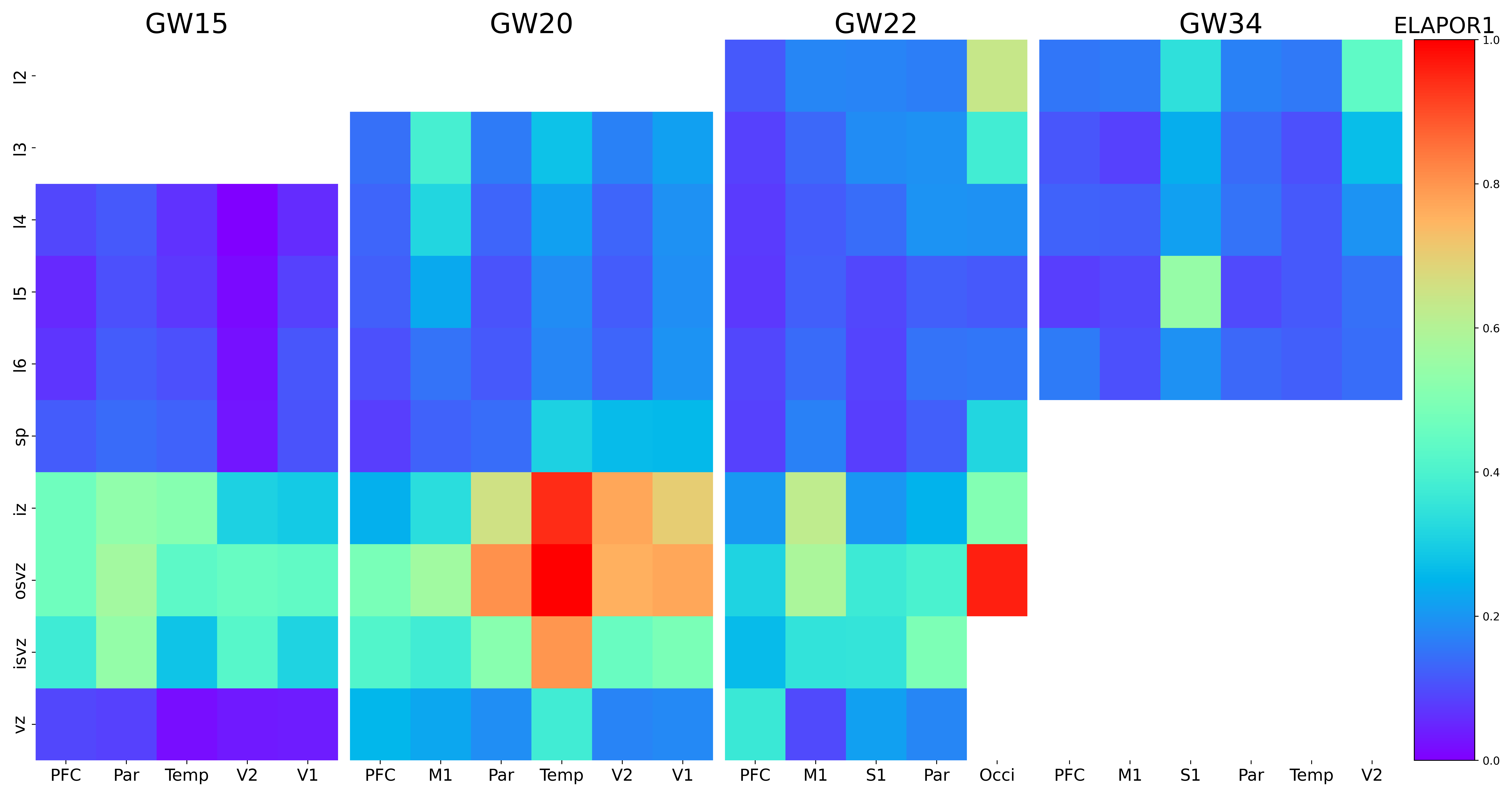

### ELMOD1.png

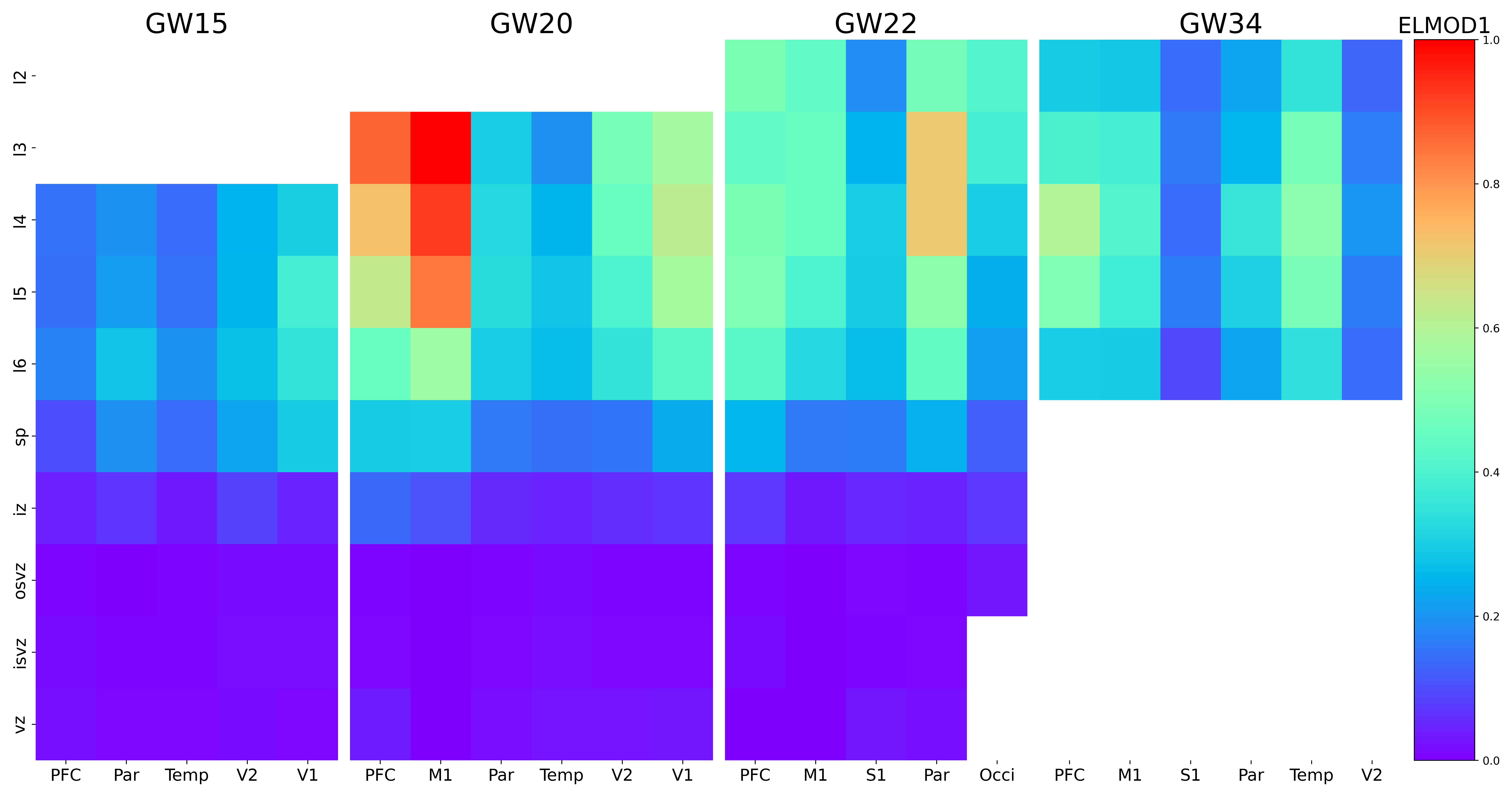

### ELOVL2-AS1.png

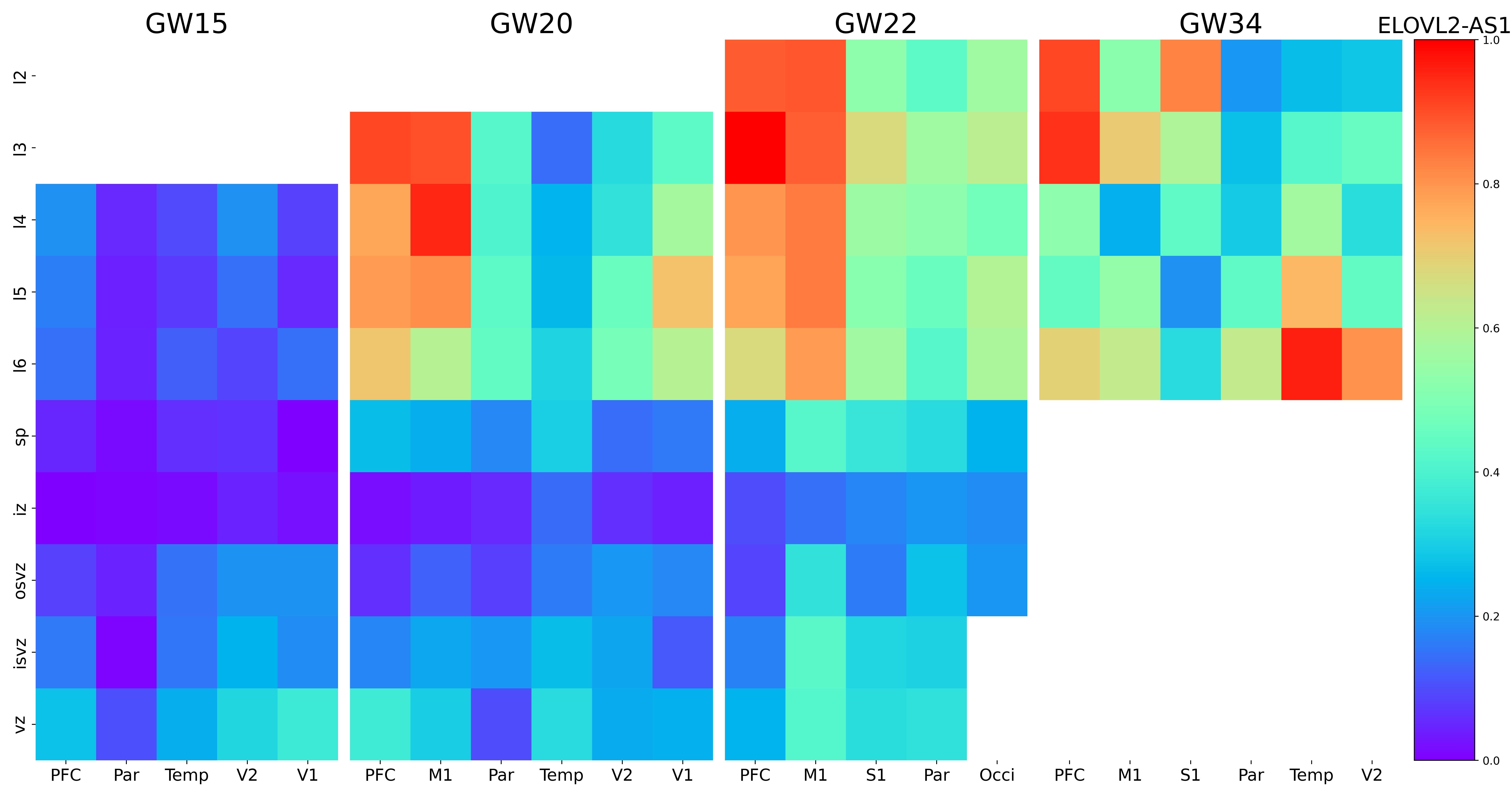

### ELOVL2.png

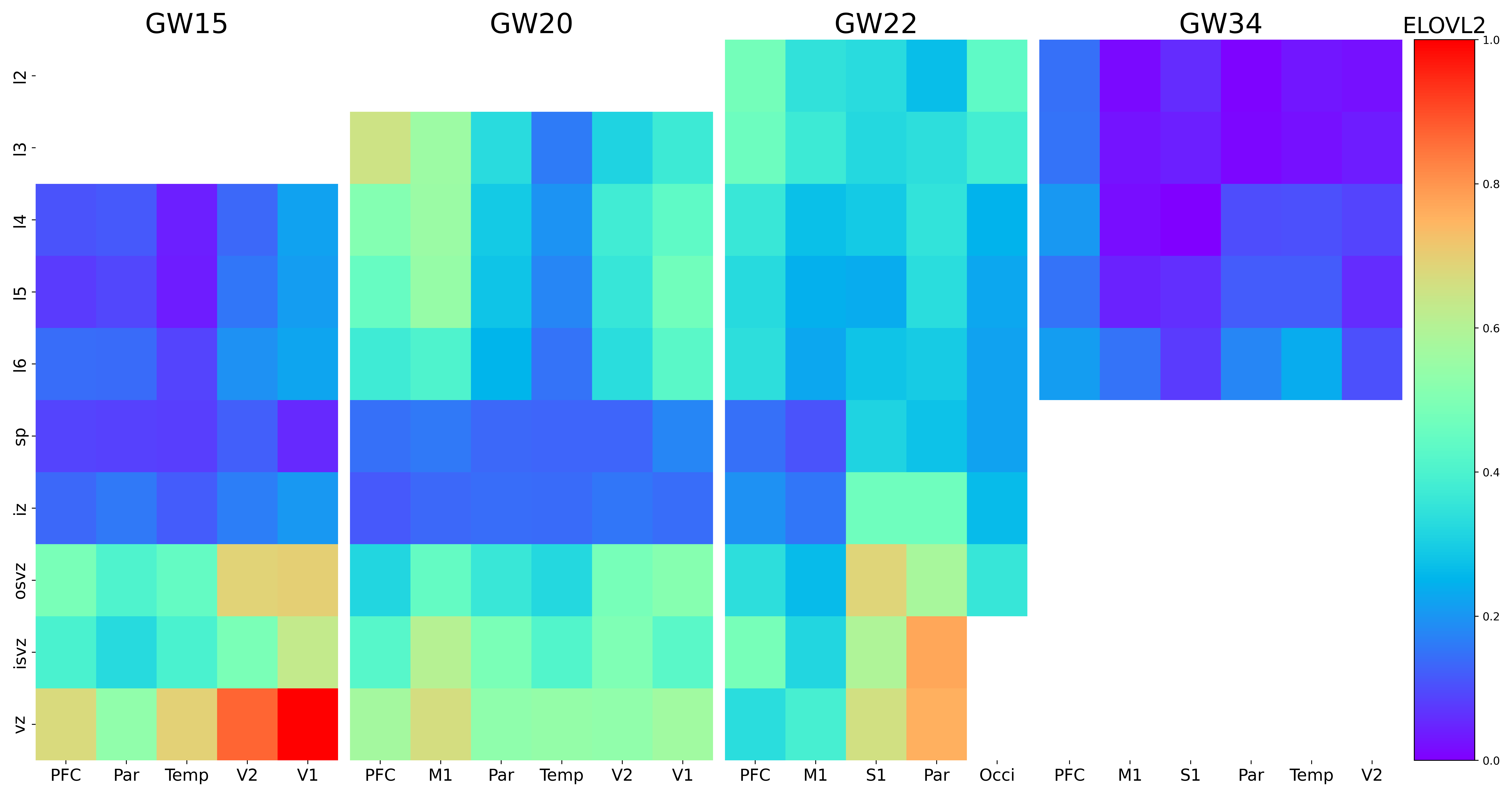

### EML6.png

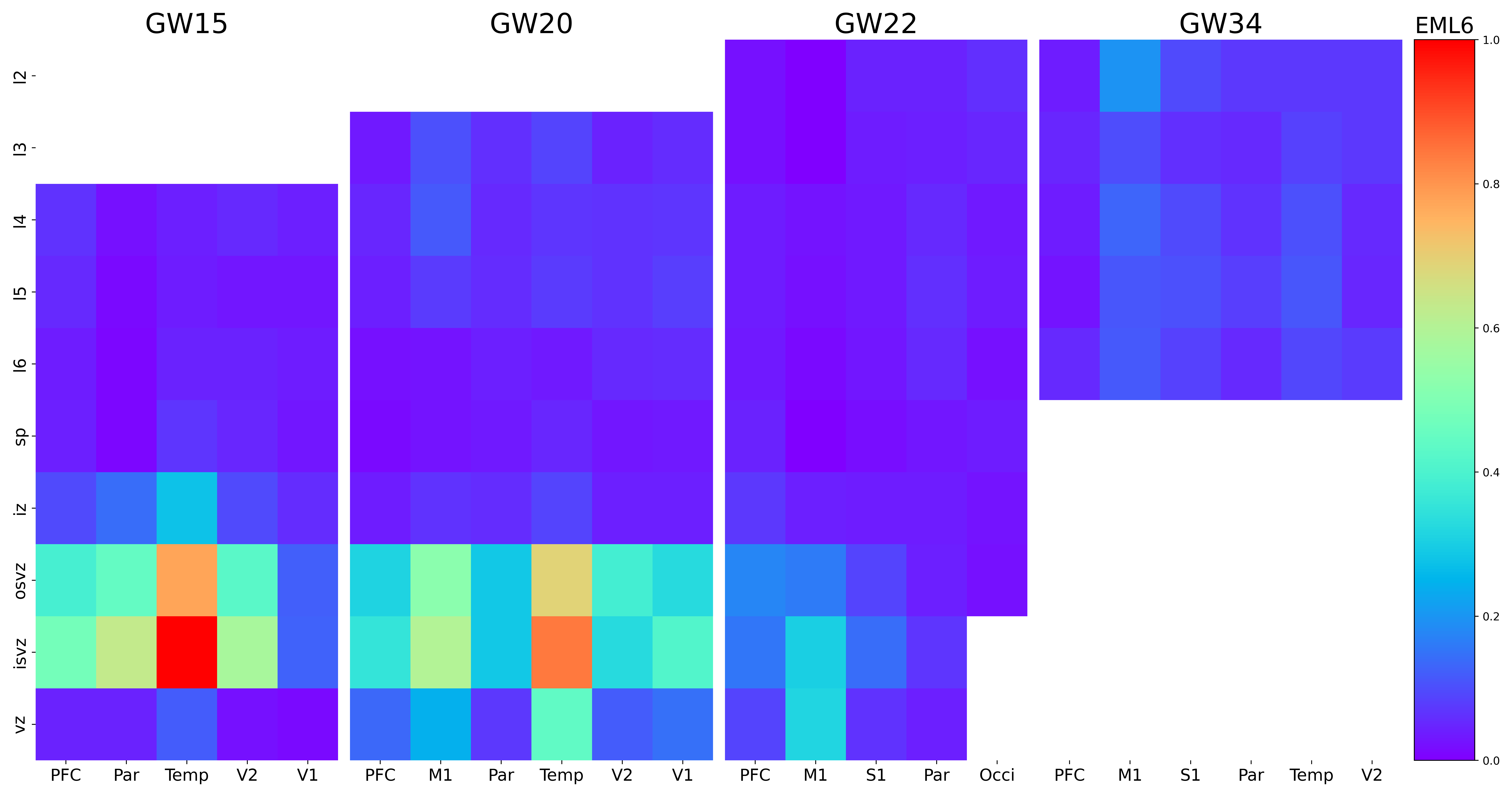

### EMX1.png

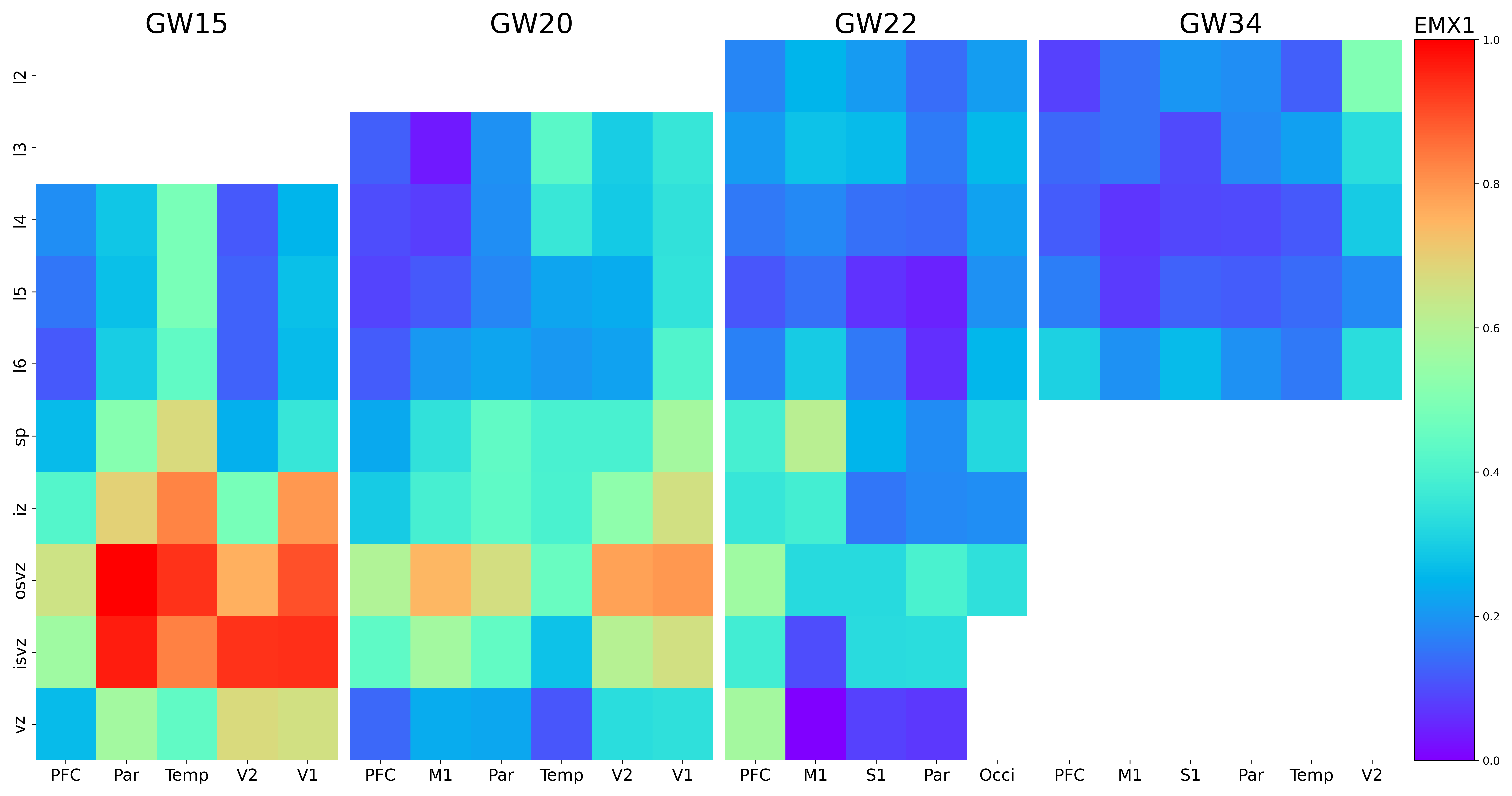

### EOMES.png

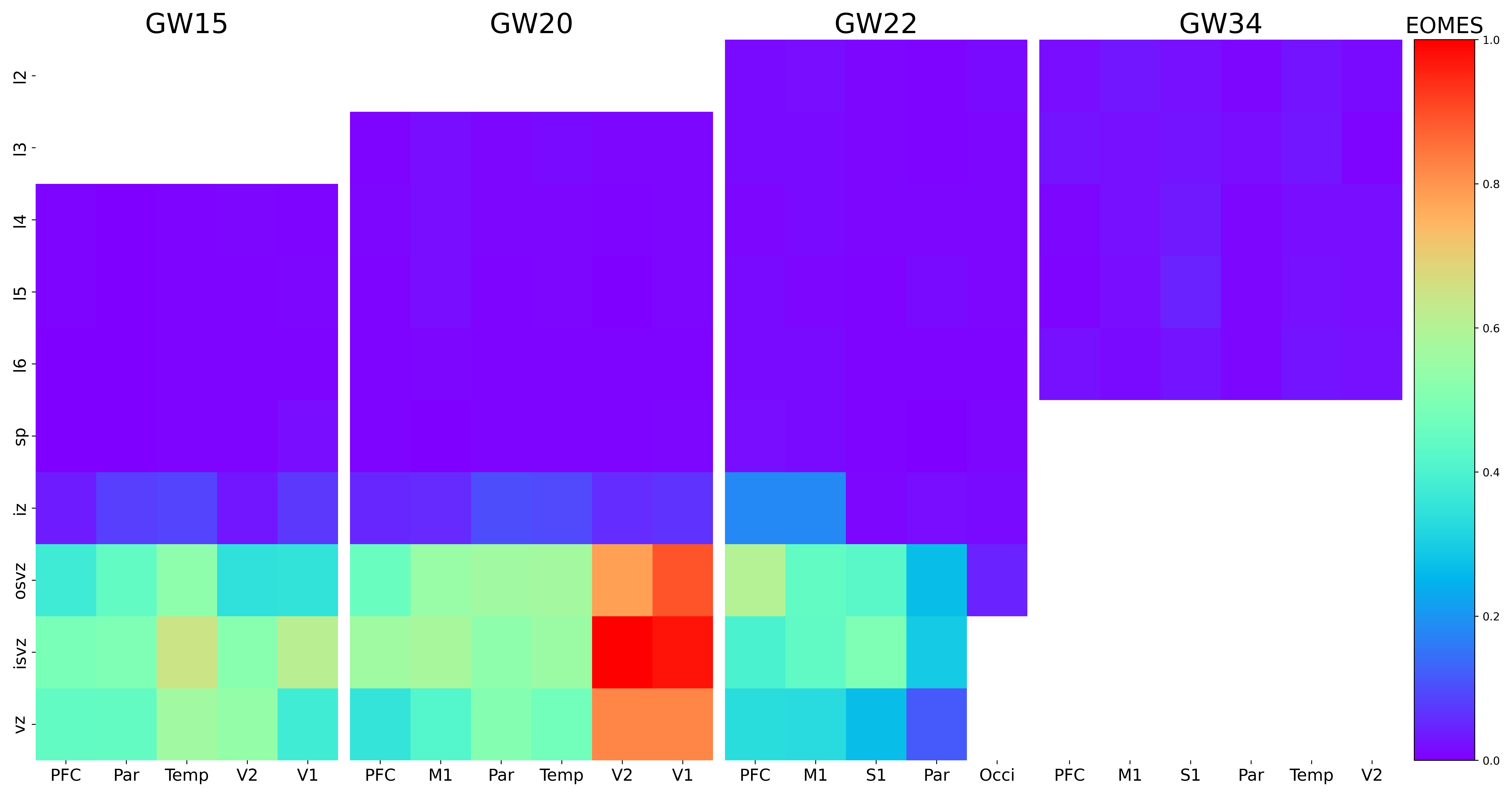

### ETV1.png

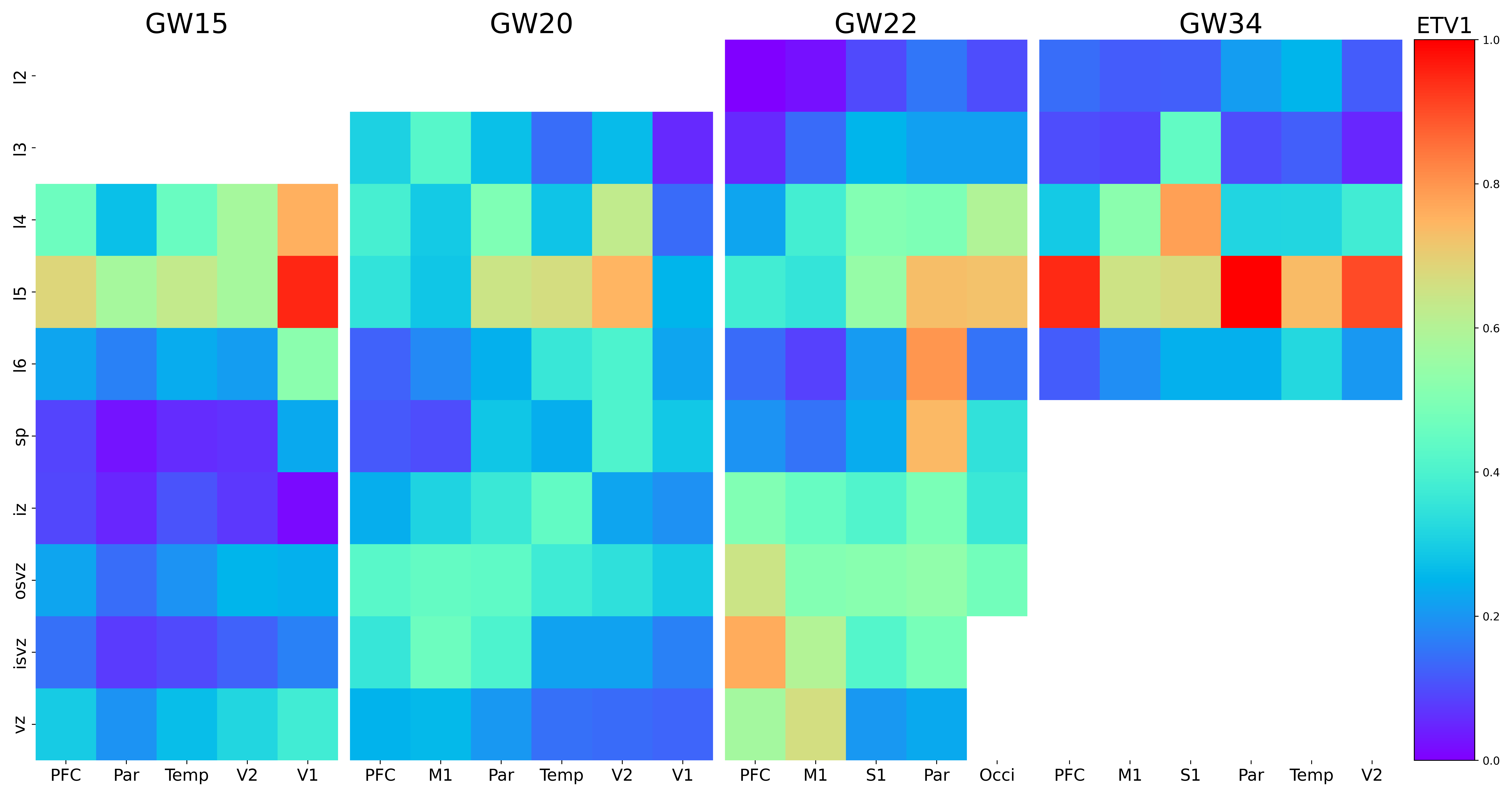

### FAM3C.png

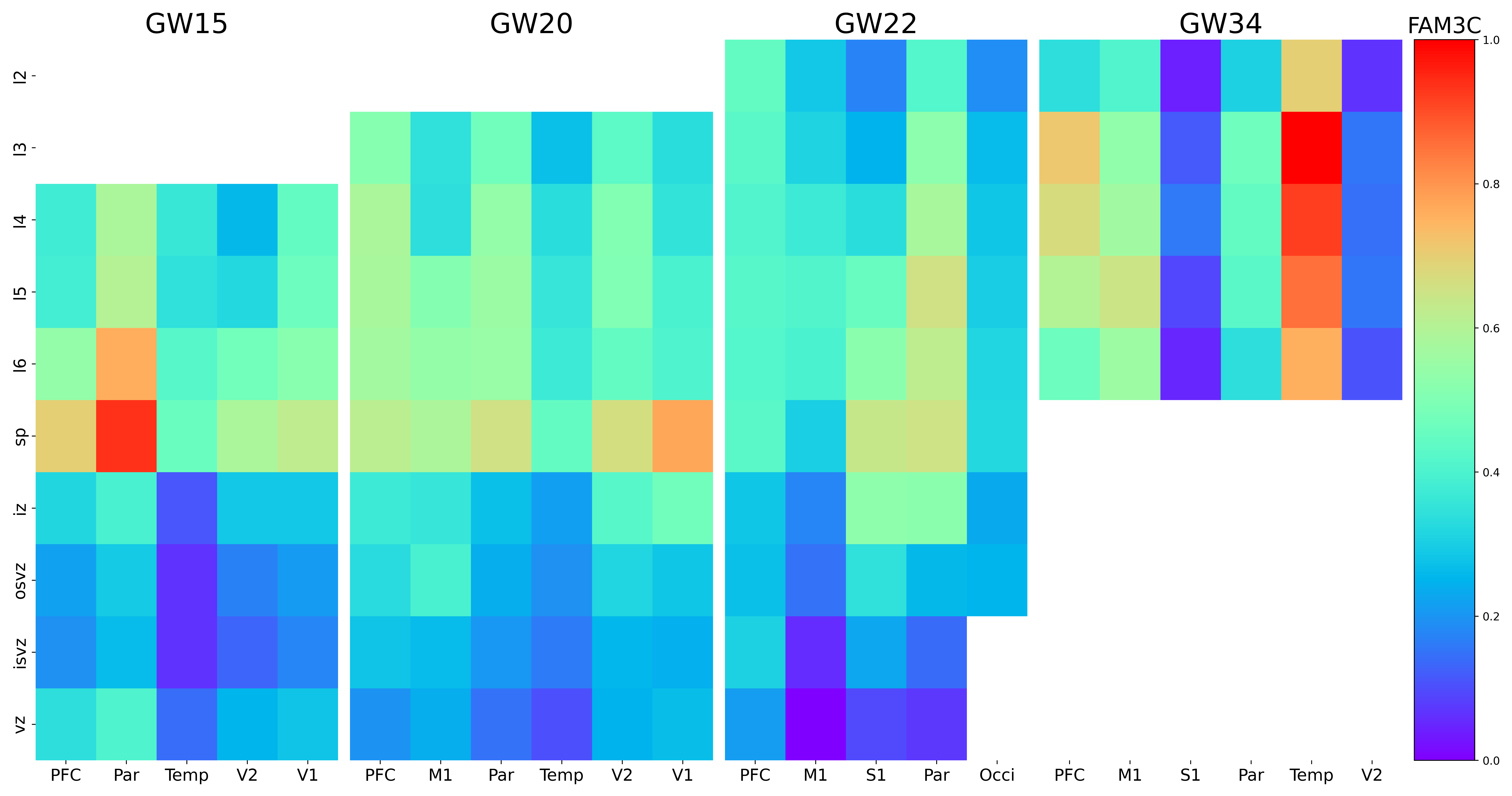

### FAM107A.png

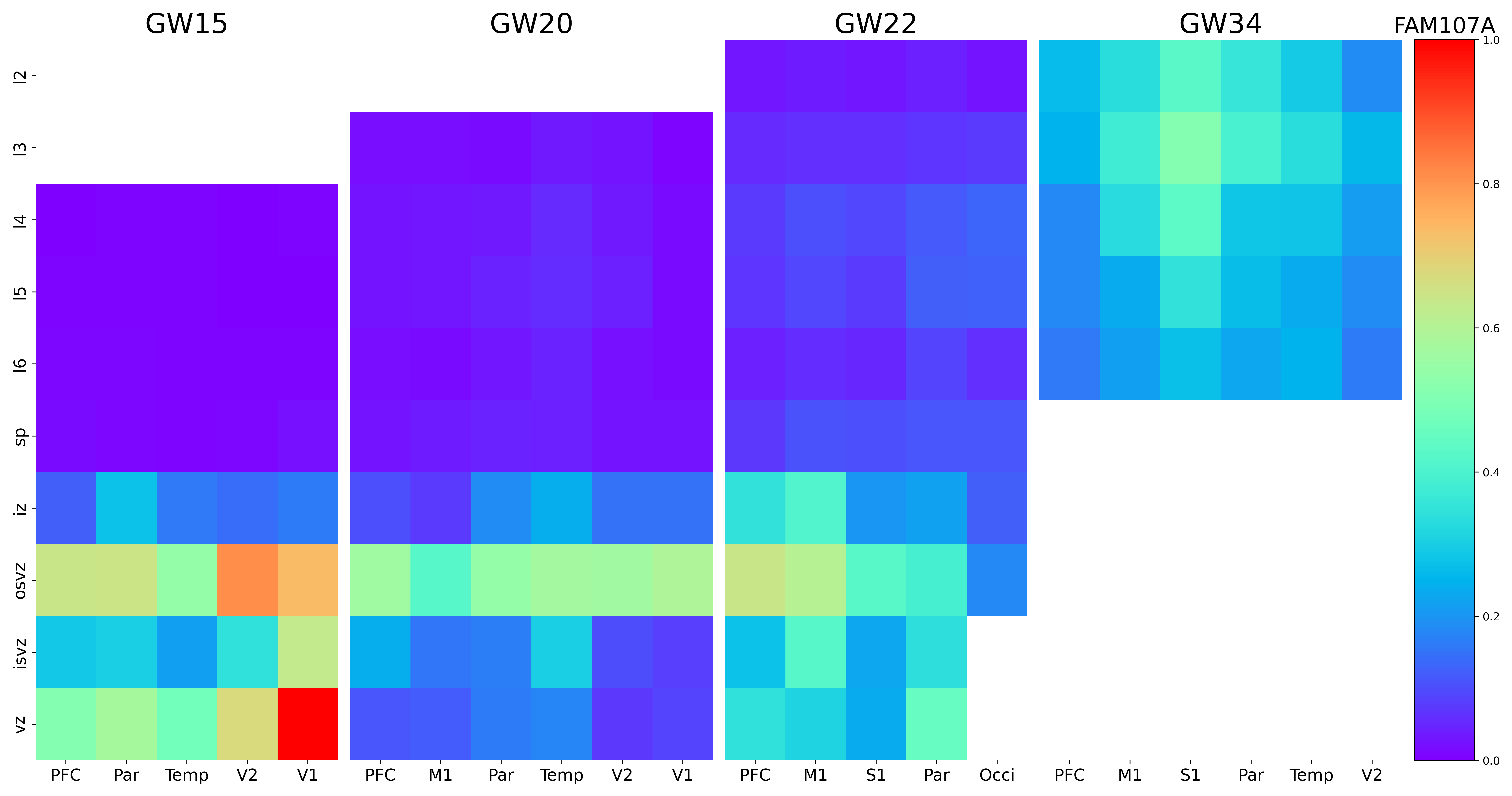

### FAM182B.png

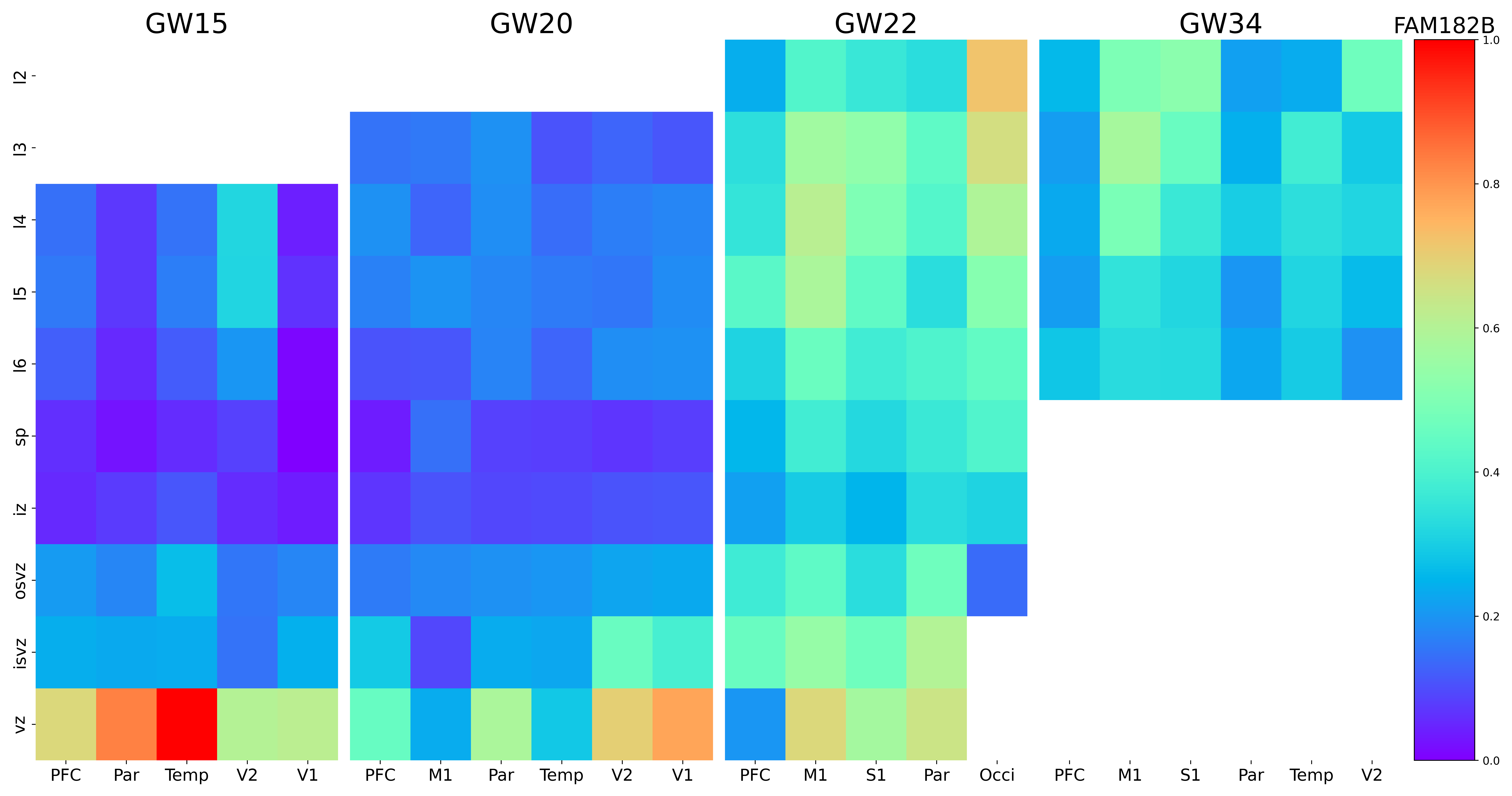

### FBXO32.png

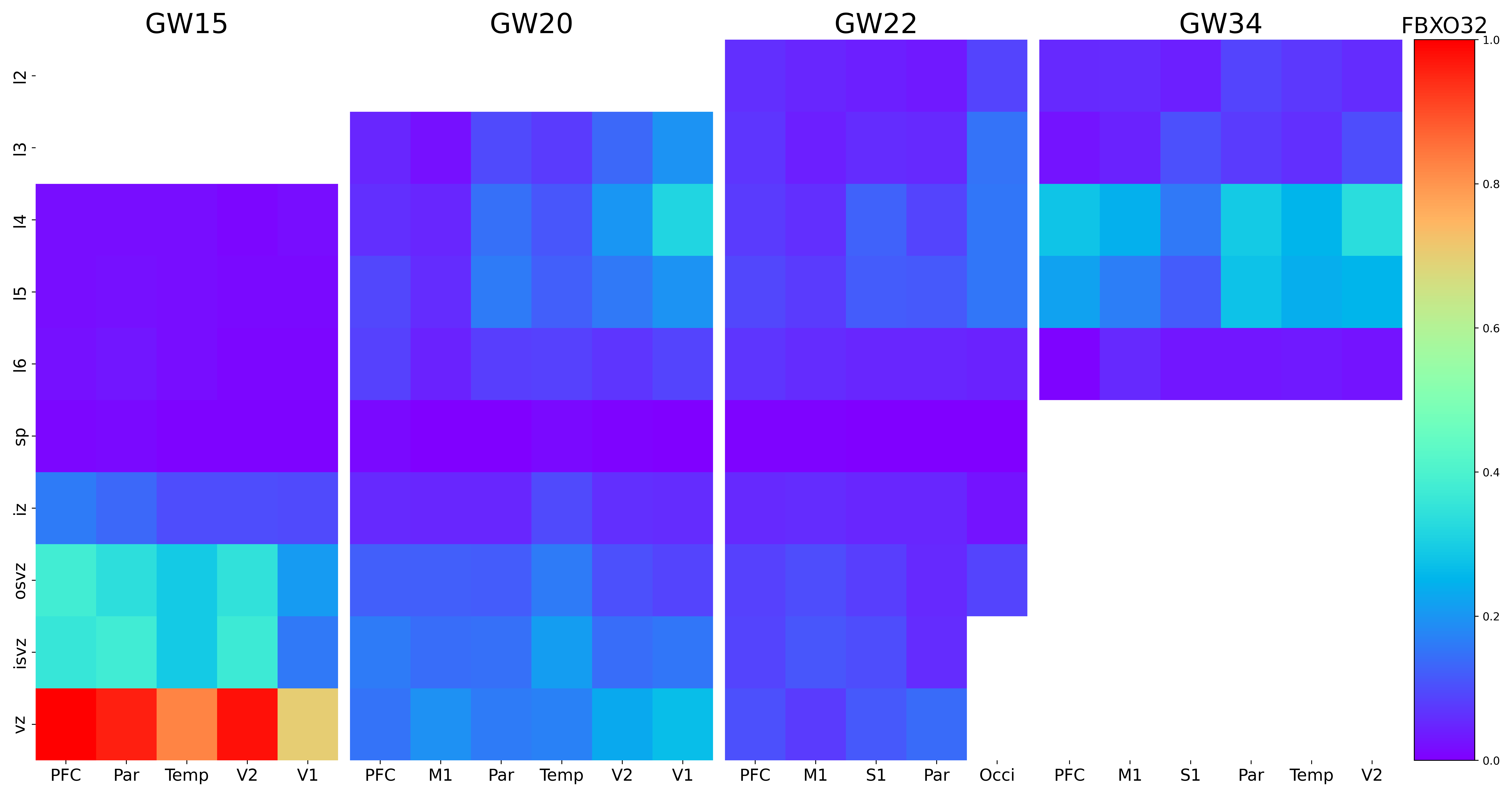

### FEZF2.png

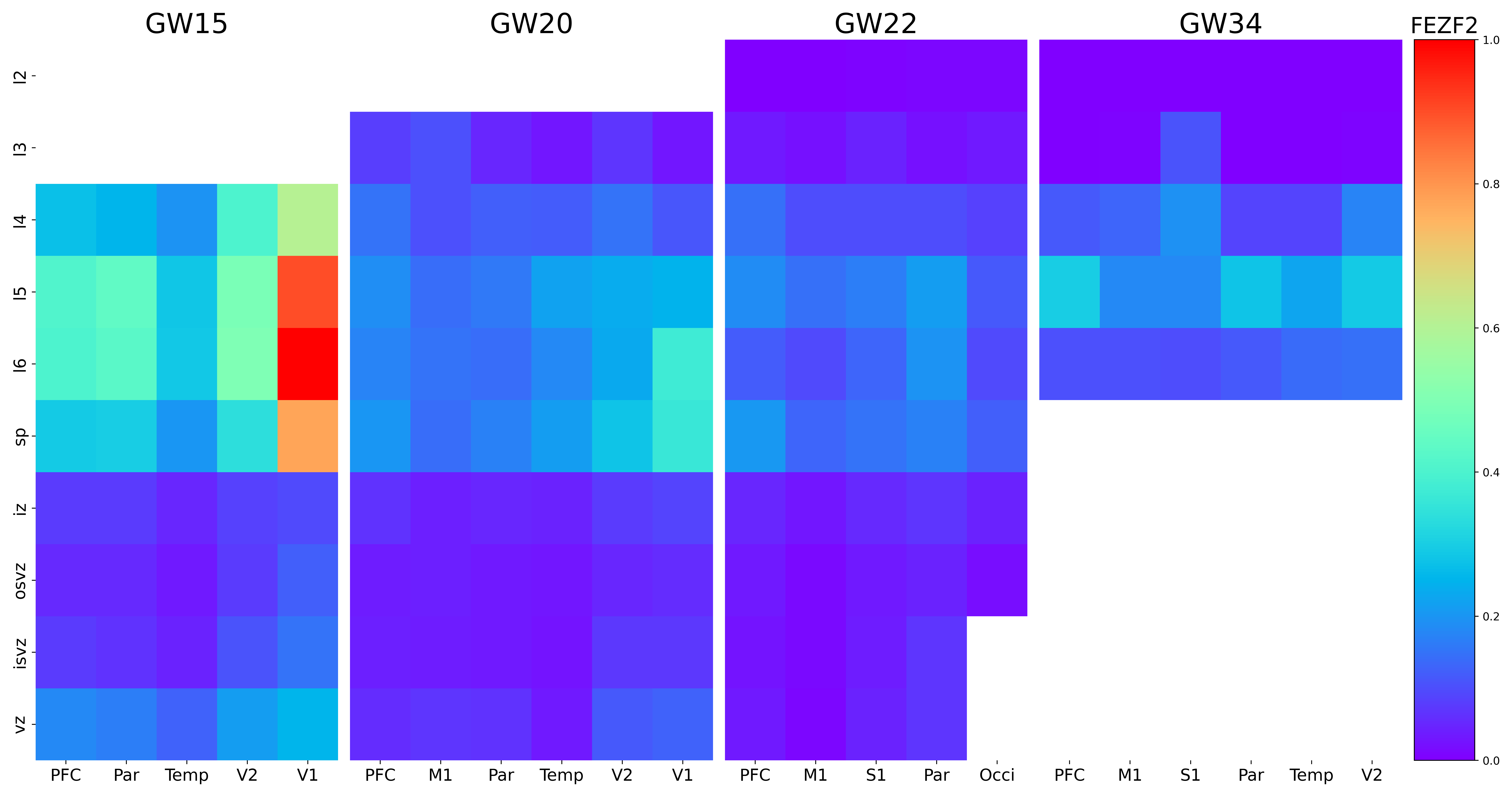

### FGFBP3.png

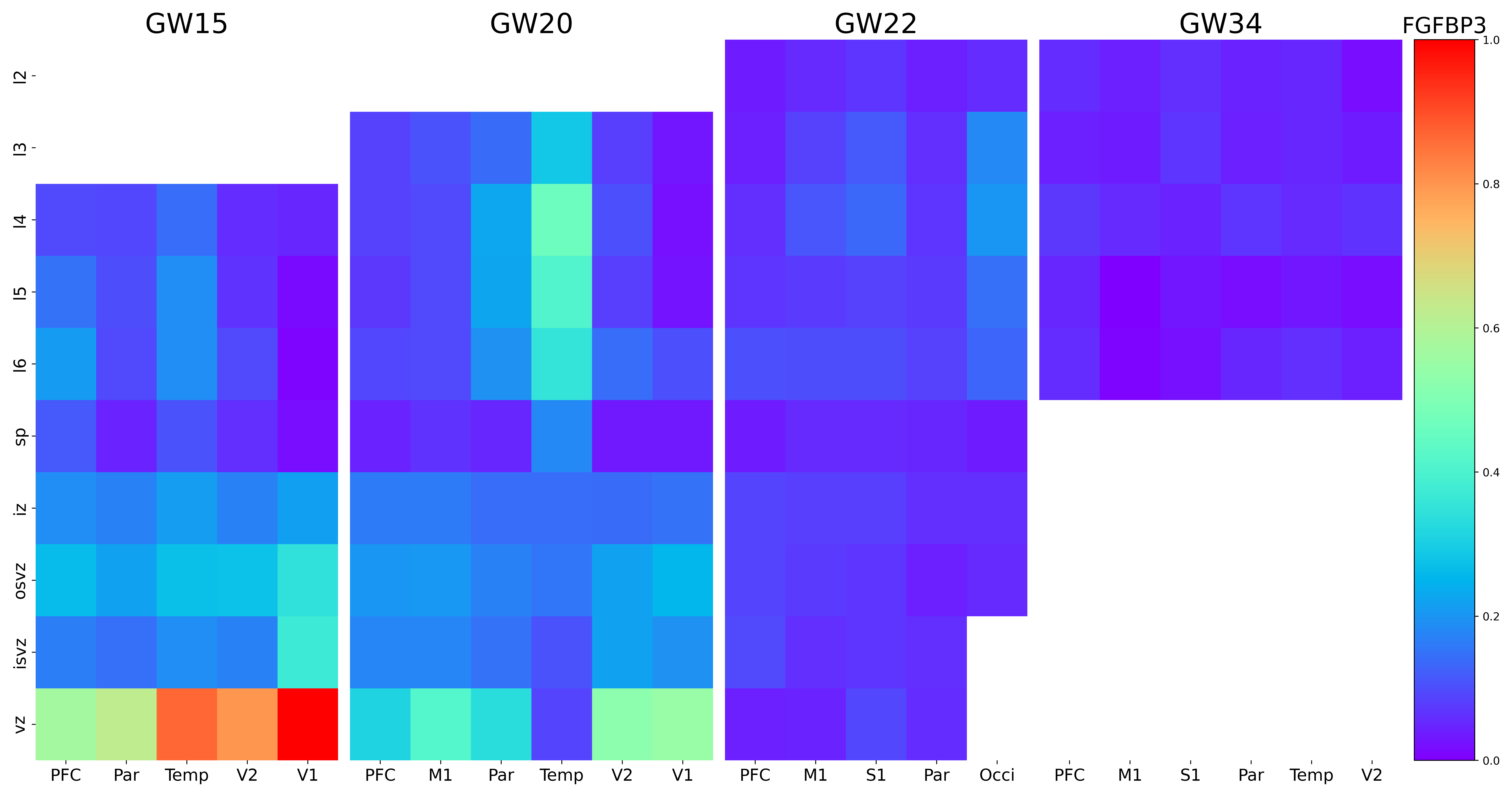

### FLJ20021.png

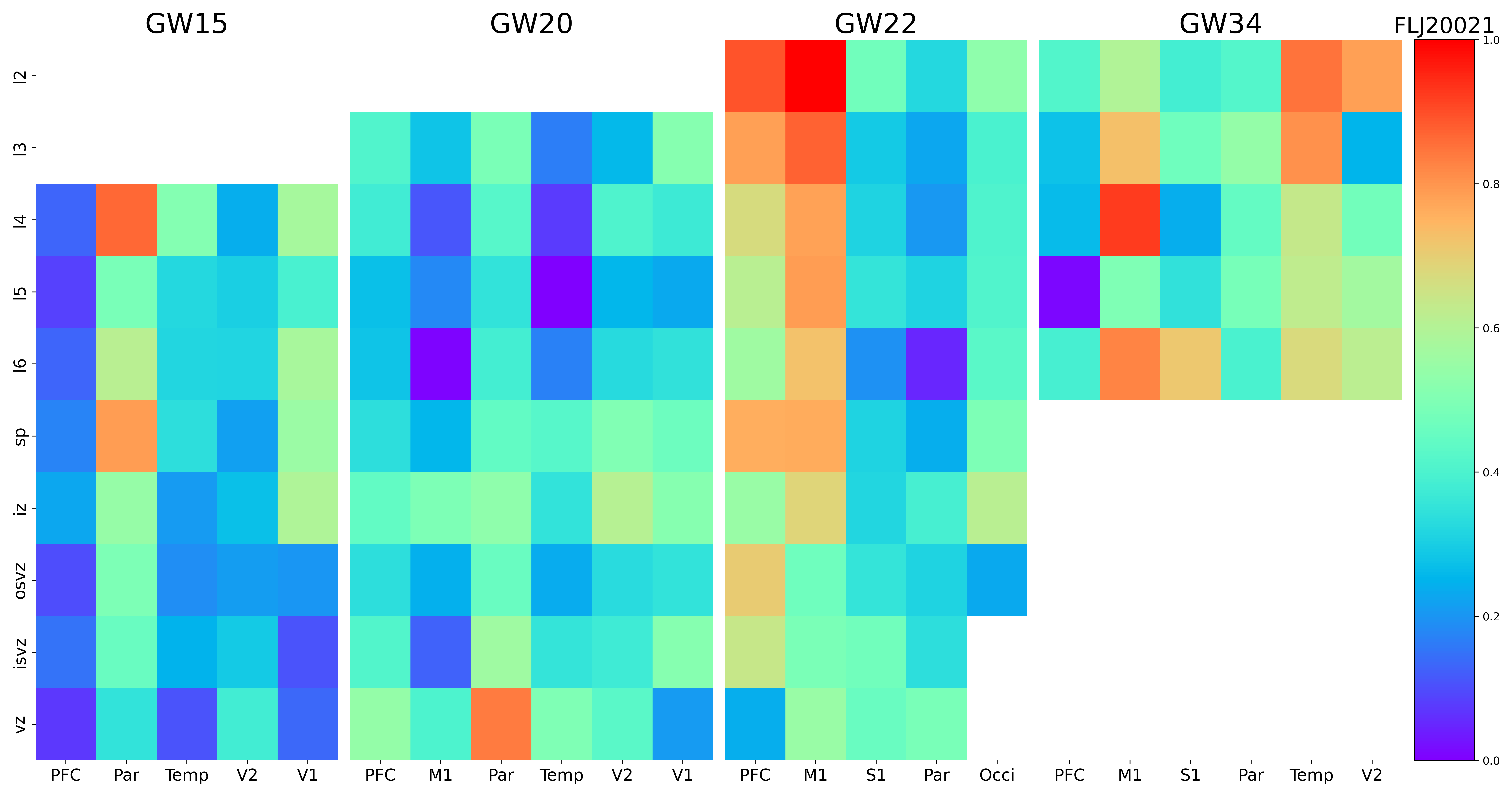

### FOXN4.png

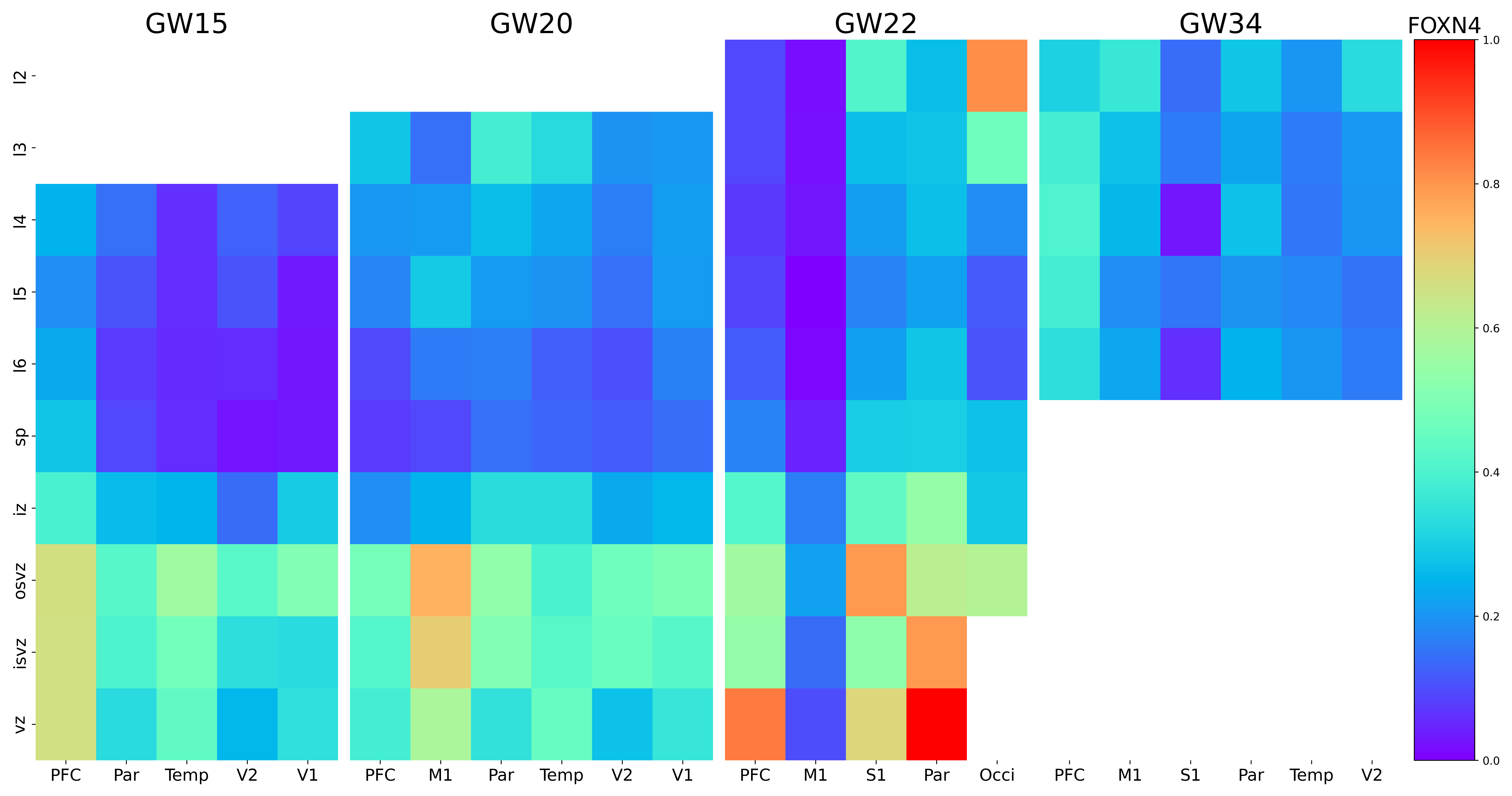
